# AMR - An R Package for Working with Antimicrobial Resistance Data

**DOI:** 10.1101/810622

**Authors:** Matthijs S. Berends, Christian F. Luz, Alexander W. Friedrich, Bhanu N.M. Sinha, Casper J. Albers, Corinna Glasner

## Abstract

Antimicrobial resistance is an increasing threat to global health. Evidence for this trend is generated in microbiological laboratories through testing microorganisms for resistance against antimicrobial agents. International standards and guidelines are in place for this process as well as for reporting data on (inter-)national levels. However, there is a gap in the availability of standardized and reproducible tools for working with laboratory data to produce the required reports. It is known that extensive efforts in data cleaning and validation are required when working with data from laboratory information systems. Furthermore, the global spread and relevance of antimicrobial resistance demands to incorporate international reference data in the analysis process.

In this paper, we introduce the **AMR** package for R that aims at closing this gap by providing tools to simplify antimicrobial resistance data cleaning and analysis, while incorporating international guidelines and scientifically reliable reference data. The **AMR** package enables standardized and reproducible antimicrobial resistance analyses, including the application of evidence-based rules, determination of first isolates, translation of various codes for microorganisms and antimicrobial agents, determination of (multi-drug) resistant microorganisms, and calculation of antimicrobial resistance, prevalence and future trends. The **AMR** package works independently of any laboratory information system and provides several functions to integrate into international workflows (e.g., WHONET software provided by the World Health Organization).

## 1. Introduction

Antimicrobial resistance is a global health problem and of great concern for human medicine, veterinary medicine, and the environment alike. It is associated with significant burdens to both patients and health care systems. Current estimates show the immense dimensions we are already facing, such as claiming at least 50,000 lives due to antimicrobial resistance each year across Europe and the US alone (27). Although estimates for the burden through antimicrobial resistance and their predictions are disputed (11) the rising trend is undeniable (5), thus calling for worldwide efforts on tackling this problem.

Surveillance programs and reliable data are key for controlling and streamlining these efforts. Surveillance data of antimicrobial resistance at higher levels (national or international) usually comprise aggregated numbers. The basis of this information is generated and stored at local microbiological laboratories where isolated microorganisms are tested for their susceptibility to a whole range of antimicrobial agents. The efficacy of these agents against microorganisms is nowadays interpreted as follows (12):

- R (“resistant”) -there is a high likelihood of therapeutic *failure*;
- S (“susceptible, standard dosing regimen”) -there is a high likelihood of therapeutic *success* using a standard dosing regimen of an antimicrobial agent;
- I (“susceptible, increased exposure”) -there is a high likelihood of therapeutic *success*, but only when exposure to an antimicrobial agent is increased by adjusting the dosing regimen or its concentration at the site of infection.

Generally, antimicrobial resistance is defined as the proportion of resistant microorganisms (R) among all tested microorganisms of the same species (R + S + I). Today, the two major guideline institutes to define the international standards on antimicrobial resistance are the European Committee on Antimicrobial Susceptibility Testing (EUCAST) (18) and the Clinical and Laboratory Standards Institute (CLSI) (6). The guidelines from these two institutes are adopted by 94% of all countries reporting antimicrobial resistance to the WHO (45).

Although these standardized guidelines are in place on the laboratory level for the data generation process, stored data in laboratory information systems are often not yet suitable for data analysis. Laboratory information systems are often designed to fit billing purposes rather than epidemiological data analysis. Furthermore, (inter-)national surveillance is hindered by inadequate standardization of epidemiological definitions, different types of samples and data collection, settings included, microbiological testing methods (including susceptibility testing), and data sharing policies (33). The necessity of accurate data analysis in the field of antimicrobial resistance has just recently been further underlined (20). Antimicrobial resistance analyses require a thorough understanding of microbiological tests and their results, the biological taxonomy of microorganisms, the clinical and epidemiological relevance of the results, their pharmaceutical implications, and (inter-)national standards and guidelines for working with and reporting antimicrobial resistance.

Here, we describe the **AMR** package for R (29), which has been developed to standardize clean and reproducible antimicrobial resistance data analyses using international standardized recommendations (18; 6) while incorporating scientifically reliable reference data about valid laboratory outcome, antimicrobial agents, and the complete biological taxonomy of microorganisms. The **AMR** package provides solutions and support for these aspects while being independent of underlying laboratory information systems, thereby democratizing the analysis process. Developed in R and available on the Comprehensive R Archive Network (CRAN) since February 22^nd^2018 (3), the **AMR** package enables reproducible workflows as described in other fields, such as environmental science (22). The **AMR** package provides a new technical instrument to aid in curbing the global threat of antimicrobial resistance. Furthermore, local and regional data in the laboratories can now become relevant in any setting for public health.

While no other packages R package with the purpose of dealing with antimicrobial resistance data are available on CRAN or Bioconductor, the **AMR** package may be integrated in workflows of related packages. For example, the R Epidemics Consortium (RECON) provides high-quality packages for data analysis in infectious disease outbreaks or epidemics (for example incidence and epicontacts) (16; 26). In addition, on the laboratory side the antibioticR package provides approaches to work with disc diffusion zone diameter and minimum inhibitory concentration data from environment samples (28). We aim at providing a comprehensive and standardized toolbox for antimicrobial resistance data processing and analysis, with a focus on microbiological, clinical, and epidemiological purposes that was yet missing.

The following sections describe the functionality of the **AMR** package according to its core functionalities for transforming, enhancing, and analyzing antimicrobial resistance data using scientifically reliable reference data.

## 2. Antimicrobial resistance data

Microbiological tests can be performed on different specimens, such as blood or urine samples or nasal swabs. After arrival at the microbiological laboratory, the specimens are traditionally cultured on specific media, such as blood agar. If a microorganism can be isolated from these media, it is tested against several antimicrobial agents. Based on the minimal inhibitory concentration (MIC) of the respective agent and interpretation guidelines, such as guidelines by EUCAST (18) and CLSI (6), test results are reported as “resistant” (R), “susceptible” or “susceptible, increased exposure” (I). A typical data structure is illustrated in Table 1 (18).

**Table 1:**
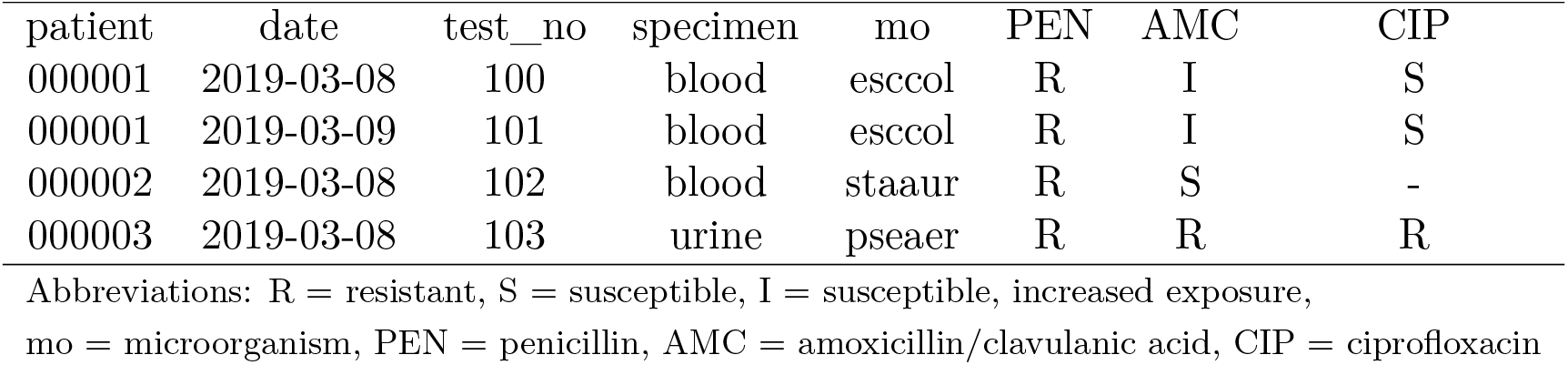
Example of an antimicrobial resistance report.

For the first two rows, the information should be read as: *Escherichia coli* (mo code = esccol) was isolated from blood of patient 000001 and was found to be resistant to penicillin, and susceptible to amoxicillin/clavulanic acid and ciprofloxacin. However, often (especially when merging sources) data is reported in ambiguous formats as exemplified in Table 2. It is crucial that source data can be analyzed in a reliable way, especially when the outcome will be used to evaluate patient treatment options. This requires reproducible and field-specific, specialized data cleaning and transforming.

**Table 2:** Antimicrobial resistance report example - ambiguous formats.

| patient | date | test_no | specimen | mo | PEN | AMC | CIP |
| --- | --- | --- | --- | --- | --- | --- | --- |
| 1 | 2019-03-08 | 100 | blood | esccol | R | I | S |
| 1 | 2019-03-09 | 101 | blood | esccol | R | I | S |
| 2 | 2019-03-08 | 102 | blood | StaAur | >8 (R)* | <0.01 (S)* | . |
| 00003 | 2019-03-08 | 103 | urine | P. aeru. | R | S** | S |

\* Mixed reporting of minimal inhibitory concentration (MIC) and susceptibility interpretation of MIC value
\*\* False reporting; *Pseudomonas aeruginosa* (mo = P. aeru.) is intrinsically resistant to amoxicillin/clavulanic acid (AMC)
Abbreviations: R = resistant, S = susceptible, I = susceptible, increased exposure, mo = microorganism, PEN = penicillin, AMC = amoxicillin/clavulanic acid, CIP = ciprofloxacin

The **AMR** package aims at providing a standardized and automated way of cleaning, transforming, and enhancing these typical data structures (Table 1 and 2), independent of the underlying data source. Processed data would be similar to Table 3 that highlights several package functionalities in the sections below.

**Table 3:**
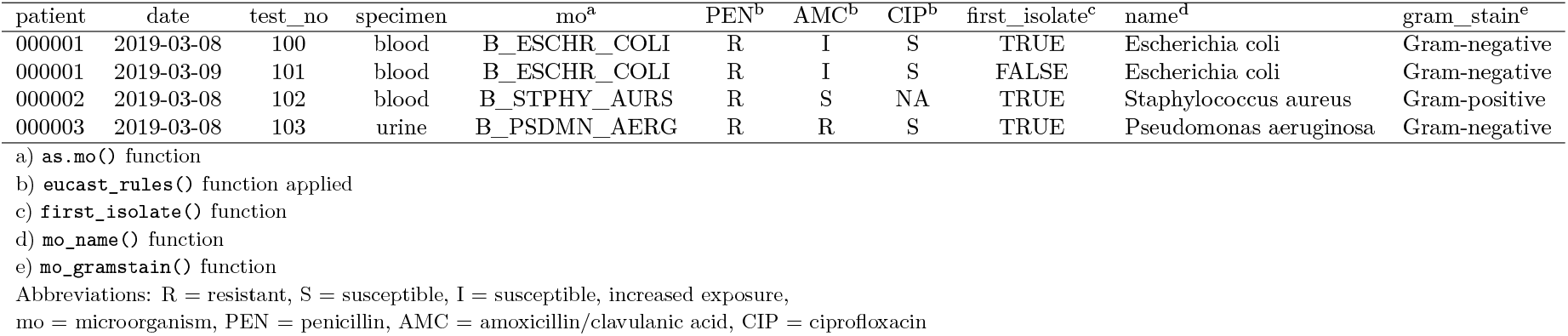
Enhanced antimicrobial resistance report example.

## 3. Antimicrobial resistance data transformation

### 3.1. Working with taxonomically valid microorganism names

Coercing is a computational process of forcing output based on an input. For microorganism names, coercing user input to taxonomically valid microorganism names is crucial to ensure correct interpretation and to enable grouping based on taxonomic properties. To this end, the **AMR** package includes all microbial entries from The Catalogue of Life (http://www.catalogueoflife.org), the most comprehensive and authoritative global index of species currently available (30). It holds essential information on the names, relationships, and distributions of more than 1.9 million species. The integration of it into the **AMR** package is described in the appendix A.

The `as.mo()` function makes use of this underlying data to transform a vector of characters to a new class ‘mo’ of taxonomically valid microorganism name. The resulting values are microbial IDs, which are human-readable for the trained eye and contain information about the taxonomic kingdom, genus, species, and subspecies (Figure 1).

**Figure 1.**
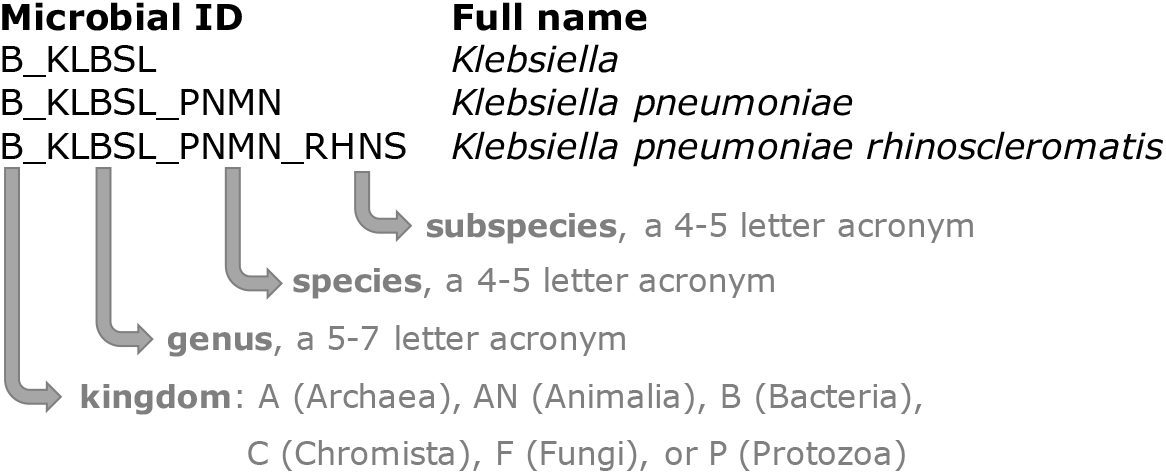
The structure of a typical microbial ID as used in the AMR package. An ID consists of two to four elements, separated by an underscore. The first element is the abbreviation of the taxonomic kingdom. The remaining elements consist of abbreviations of the lowest taxonomic levels of every microorganism: genus, species (if available) and subspecies (if available). Abbreviations used for the microbial IDs of microorganism names were created using the base R function `abbreviate()`.

The `as.mo()` function compares the user input with taxonomically valid microorganism names, rates the matching with a score and returns results based on the highest score. This matching score (*m*), ranging from 0 to 1, is calculated using the following equation:

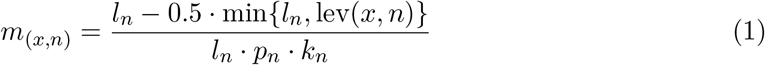

where:

- *x* is the user input;
- *n* is a taxonomic name (genus, species, and subspecies);
- *l*_*n*_ is the length of *n*;
- lev is the Levenshtein distance function (19), which counts any insertion, deletion and substitution as 1 that is needed to change *x* into *n*;
- *p*_*n*_ is the human pathogenic prevalence group of *n*, as described below;
- *k*_*n*_ is the taxonomic kingdom of *n*, set as Bacteria = 1, Fungi = 2, Protozoa = 3, Archaea = 4, others = 5.

The grouping into human pathogenic prevalence (*p*) is based on experience from several microbiological laboratories in the Netherlands in conjunction with international reports on pathogen prevalence (10; 13; 45). <u>Group 1</u> (most prevalent microorganisms) consists of all microorganisms where the taxonomic class is Gammaproteobacteria or where the tax-

onomic genus is *Enterococcus, Staphylococcus* or *Streptococcus*. This group consequently contains all common Gram-negative bacteria, such as *Pseudomonas* and *Legionella* and all species within the order Enterobacterales. <u>Group 2</u> consists of all microorganisms where the taxonomic phylum is Proteobacteria, Firmicutes, Actinobacteria or Sarcomastigophora, or where the taxonomic genus is *Absidia, Acremonium, Actinotignum, Alternaria, Anaerosalibacter, Apophysomyces, Arachnia, Aspergillus, Aureobacterium, Aureobasidium, Bacteroides, Basidiobolus, Beauveria, Blastocystis, Branhamella, Calymmatobacterium, Candida, Capnocytophaga, Catabacter, Chaetomium, Chryseobacterium, Chryseomonas, Chrysonilia, Cladophialophora, Cladosporium, Conidiobolus, Cryptococcus, Curvularia, Exophiala, Exserohilum, Flavobacterium, Fonsecaea, Fusarium, Fusobacterium, Hendersonula, Hypomyces, Koserella, Lelliottia, Leptosphaeria, Leptotrichia, Malassezia, Malbranchea, Mortierella, Mucor, Mycocentrospora, Mycoplasma, Nectria, Ochroconis, Oidiodendron, Phoma, Piedraia, Pithomyces, Pityrosporum, Prevotella,Pseudallescheria, Rhizomucor, Rhizopus, Rhodotorula, Scolecobasidium, Scopulariopsis, Scytalidium,Sporobolomyces, Stachybotrys, Stomatococcus, Treponema, Trichoderma, Trichophyton, Trichosporon, Tritirachium* or *Ureaplasma*. <u>Group 3</u> consists of all other microorganisms.

This will lead to the effect that e.g., `“E. coli”` will return the microbial ID of *Escherichia coli* (*m* = 0.688, a highly prevalent microorganism found in humans) and not *Entamoeba coli* (*m* = 0.079, a less prevalent microorganism in humans), although the latter would alphabetically come first. The matching score function is for users available as `mo_matching_score()`.

If any coercion rules are applied, a warning is printed to the console and scores can be reviewed by calling `mo_uncertainties()`, that prints all other matches with their matching scores. Users can furthermore control the coercion rules by setting the `allow_uncertain` argument in the `as.mo()` function. The following values or levels can be used:

- `0`: no additional rules are applied;
- `1`: allow previously accepted (but now invalid) taxonomic names and minor spelling errors;
- `2`: allow all of 1, strip values between brackets, inverse the words of the input, strip off text elements from the end keeping at least two elements;
- `3`: allow all of level `1` and `2`, strip off text elements from the end, allow any part of a taxonomic name;
- `TRUE` (default): equivalent to 2;
- `FALSE`: equivalent to 0.

To support organization specific microbial IDs, users can specify a custom reference `‘data.frame’`, by using `as.mo(…, reference_df = …)`. This process can also be automated by users with the `set_mo_source()` function.

#### Properties of microorganisms

The package contains functions to return a specific (taxonomic) property of a microorganism from the `microorganisms` data set (see appendix A). Functions that start with `mo_*` can be used to retrieve the most recently defined taxonomic properties of any microorganism quickly and conveniently. These functions rely on the `as.mo()` function internally: mo_name(), mo_fullname(), mo_shortname(), mo_subspecies(), mo_species(), mo_genus(), mo_family(), mo_order(), mo_class(), mo_phylum(), mo_kingdom(), mo_type(), mo_gramstain(), mo_ref(),mo_authors(), mo_year(), mo_rank(), mo_taxonomy(), mo_synonyms(), mo_info() and `mo_url()`. Determination of the Gram stain, by using `mo_gramstain()`, is based on the taxonomic subkingdom and phylum. According to (4), who defined the subkingdoms Negibacteria and Posibacteria, only the following phyla are Posibacteria: Actinobacteria, Chloroflexi, Firmicutes and Tenericutes. Bacteria from these phyla are considered Gram-positive -all other bacteria are considered Gram-negative. Gram stains are only relevant for species within the kingdom of Bacteria. For species outside this kingdom, `mo_gramstain()` will return `NA`.

### 3.2. Working with antimicrobial names or codes

The **AMR** package includes the antibiotics data set, which comprises common laboratory information system codes, official names, ATC (Anatomical Therapeutic Chemical) codes, defined daily doses (DDD) and more than 5,000 trade names of 456 antimicrobial agents (see appendix A). The ATC code system and the reference list for DDDs have been developed and made available by the World Health Organization Collaborating Centre for Drug Statistics Methodology (WHOCC) to standardize pharmaceutical classifications (37). All agents in the `antibiotics` data set have a unique antimicrobial ID, which is based on abbreviations used by the European Antimicrobial Resistance Surveillance Network (EARS-Net), the largest publicly funded system for antimicrobial resistance surveillance in Europe (14). Furthermore, the **AMR** package includes the `antivirals` data seta containing antiviral agents, which is also described in the appendix A.

#### Properties of antimicrobial agents

It is a common task in microbiological data analyses (and other clinical or epidemiological fields) to work with different antimicrobial agents. The **AMR** package provides several functions to translate inputs such as ATC codes, abbreviations, or names in any direction. Using `as.ab()`, any input will be transformed to an antimicrobial ID of class `‘ab’`. Helper functions are available to get specific properties of antimicrobial IDs, such as `ab_name()` for getting the official name, `ab_atc()` for the ATC code, or `ab_cid()` for the CID (Compound ID) used by PubChem (17). Trade names can be also used as input. For example, the input values `“Amoxil”, “dispermox”, “amox”` and `“J01CA04”` all return the ID of amoxicillin (`AMX`):

~~~
*R> as*.*ab(“Amoxicillin”)*
Class <ab>
[1] AMX
*R> as*.*ab(c(“Amoxil”, “dispermox”, “amox”, “J01CA04”))*
Class <ab>
[1] AMX AMX AMX AMX
*R> ab_name(“Amoxil”)*
[1] “Amoxicillin”
*R> ab_atc(“amox”)*
[1] “J01CA04”
*R> ab_name(“J01CA04”)*
[1] “Amoxicillin”
~~~

If more than one antimicrobial agent is found in the input string, a warning with the additional findings is printed to the console.

#### Filtering data based on classes of antimicrobial agents

The application of the ATC classification system also enables grouping of antimicrobial agents for data analyses. Data sets with microbial isolates can be filtered on isolates with specific results for tested antimicrobial agents in a specific antimicrobial class. For example, using `filter_carbapenems(result = “R”)` returns data of all isolates with tested resistance to any of the 14 available antimicrobial agents in in the group of carbapenems according to the antibiotics data set.

### 3.3. Working with antimicrobial susceptibility test results

Minimal inhibitory concentrations (MIC) are susceptibility test results measured by microbiological laboratory equipment to determine at which minimum antimicrobial drug concentration 99.9% of a microorganism is inhibited in growth. These concentrations are often capped at a minimum and maximum, for example `≤0.02 µg/ml` and `≥32 µg/ml`, respectively. The `‘mic’` class, an ordered `‘factor’` containing valid MIC values, keeps these operators while still ordering all possible outcomes correctly so that e.g., `“<= 0.02”` will be considered lower than `“0.04”`.

Another susceptibility testing method is the use of drug diffusion disks, which are small tablets containing a specified concentration of an antimicrobial agent. These disks are applied onto a solid growth medium or a specific agar plate. After 24 hours of incubation, the diameter of the growth inhibition around a disk can be measured in millimeters with a ruler. The `‘disk’` class can be used to clean these kinds of measurements, since they should always be valid numeric values between 6 and 50. The supported minima and maxima of valid values for both classes, `‘mic’` and `‘disk’`, are displayed in Table 4.

**Table 4:** Antimicrobial suceptibility test classes.

| Class | Minimum | Maximum | Unit |
| --- | --- | --- | --- |
| ‘`mic`’ | $\leq 0.001$ | $\geq 1024$ | $\mu\text{g/ml}$ |
| ‘`disk`’ | $\leq 6$ | $\geq 50$ | mm |

The higher the MIC or the smaller the growth inhibition diameter, the more active substance of an antimicrobial agent is needed to inhibit cell growth, i.e. the higher the antimicrobial resistance against the tested antimicrobial agent. At high MICs and small diameters, guidelines interpret the microorganism as “resistant” (R) to the tested antimicrobial agent. At low MICs and wide diameters, guidelines interpret the microorganism as “susceptible” (S) to the tested antimicrobial agent. In between, the microorganism is classified as “susceptible, increased exposure” (I). For these three interpretations the `‘rsi’` class has been developed. When using `as.rsi()` on MIC values (of class `‘mic’`) or disk diffusion diameters (of class ‘disk’), the values will be interpreted according to the guidelines from the CLSI or EUCAST (all guidelines between 2011 and 2020 are included in the **AMR** package) (7; 34). Guidelines can be changed by setting the `guidelines` argument.

~~~
*R> # Low MIC value
R> as.rsi(as.mic(2), “E. coli”, “ampicillin”, guideline = “EUCAST 2020”)*
Class <rsi>
[1] S
*R> # High MIC value
R> as.rsi(as.mic(32), “E. coli”, “ampicillin”, guideline = “EUCAST 2020”)*
Class <rsi>
[1] R
~~~

When using the `as.rsi()` function on existing antimicrobial interpretations, it tries to coerce the input to the values “R”, “S” or “I”. These values can in turn be used to calculate the proportion of antimicrobial resistance.

### 3.4. Interpretative rules by EUCAST

Next to supplying guidelines to interpret raw MIC values, the EUCAST has developed a set of expert rules to assist clinical microbiologists in the interpretation and reporting of antimicrobial susceptibility tests (18). The rules comprise assistance on intrinsic resistance, exceptional phenotypes, and interpretive rules. The **AMR** package covers intrinsic resistant and interpretive rules for data transformation and standardization purposes. The first prevents false susceptibility reporting by providing a list of organisms with known intrinsic resistance to specific antimicrobial agents (e.g., cephalosporin resistance of all enterococci). Interpretative rules apply inference from established resistance mechanisms (43; 8; 9; 21). Both groups of rules are based on classic IF THEN statements (e.g., IF *Enterococcus spp*. resistant to ampicillin THEN also report as resistant to imipenem). Some rules provide assistance for further actions when certain resistance has been detected, i.e., performing additional testing of the isolated microorganism. The **AMR** package function `eucast_rules()` can apply all EUCAST rules that do not rely on additional clinical information, such as additional information on patients’ diagnoses. Table 2 and 3 highlight the transformation for the reporting of AMX = S in patient_id = 000003 to the correct report according to EUCAST rules of AMX = R. Of note, however, EUCAST rules overwrite original data to correct for the difference in how antimicrobial agents affect the tested microorganism *in vitro* (in the laboratory) and *in vivo* (in the human body). This requires users to closely collaborate with the data source provider to ensure correct versioning, backward compatibility, reproducibility, and taking into account specific local regulation for resistance reporting. Typical scenarios where changes to the original data points apply include *in vitro* test results indicating susceptibility when resistance *in vivo* is known. The changes are based on scientific consensus to ensure reliable high-quality reporting of antimicrobial susceptibility results. All changes to the data are printed to the console and can also be reviewed in detail by setting the argument `eucast_rules(…, verbose = TRUE)`.

EUCAST rules are subject to regular updates which are implemented into the **AMR** package by the **AMR** maintenance team shortly after publication. The `eucast_rules()` function supports versioning of the rules. The arguments `version_breakpoints` and `version_expertrules` can be set to current or previous versions. By default, the `eucast_rules()` function uses the latest implemented version.

### 3.5. Working with defined daily doses (DDD)

DDDs are essential for standardizing antimicrobial consumption analysis, for inter-institutional or international comparison. The official DDDs are published by the WHOCC (36). Updates to the official publication are monitored by the **AMR** maintenance team and implemented in the `antibiotics` data set included in the **AMR** package. Other metrics exist such as the recommended daily dose (RDD) or the prescribed daily dose (PDD). However, DDDs are the only metric that is independent of a patient’s disease and therapeutic choices and thus suitable for the **AMR** package.

Functions from the `atc_online_*()` family take any text as input that can be coerced with `as.ab()` (i.e., to class `‘ab’`). Next, the functions access the WHOCC online registry (36) (internet connection required) and download the property defined in the arguments (e.g., `administration = “O” or administration = “P”` for oral or parenteral administration and `property = “ddd” or property = “groups”` to get DDD or the group of the selected antimicrobial defined by its ATC code).

~~~
*R> atc_online_ddd(“amoxicillin”, administration = “O”)*
[1] 1.5
*R> atc_online_groups(“amoxicillin”)*
[1] “ANTIINFECTIVES FOR SYSTEMIC USE”
[2] “ANTIBACTERIALS FOR SYSTEMIC USE”
[3] “BETA-LACTAM ANTIBACTERIALS, PENICILLINS”
[4] “Penicillins with extended spectrum”
~~~

## 4. Enhancing antimicrobial resistance data

### 4.1. Determining first isolates

Determining antimicrobial resistance or susceptibility can be done for a single agent (mono-therapy) or multiple agents (combination therapy). The calculation of antimicrobial resistance statistics is dependent on two prerequisites: the data should only comprise the first isolates and a minimum required number of 30 isolates should be met for every stratum in further analysis (6).

An isolate is a microorganism strain cultivated on specified growth media in a laboratory, so its phenotype can be determined. First isolates are isolates of any species found first in a patient per episode, regardless of the body site or the type of specimen (such as blood or urine) (6). The selection on first isolates (using function `first_isolate()`) is important to prevent selection bias, as it would lead to overestimated or underestimated resistance to an antimicrobial agent. For example, if a patient is admitted with a multi-drug resistant microorganism and that microorganism is found in five different blood cultures the following week, it would overestimate resistance if all isolates were to be included in the analysis. The episode in days can be set with the argument `episode_days`, which defaults to 365 as suggested by the (6) guideline.

### 4.2. Determining multi-drug resistant organisms (MDRO)

Definitions of multi-drug resistant organisms (MDRO) are regulated by national and international expert groups and differ between nations. The **AMR** package provides the functionality to quickly identify MDROs in a data set using the mdro() function. Guidelines can be set with the argument `guideline`. At default, it applies the guideline as proposed by (24). Their work describes the definitions for bacteria being ‘MDR’ (multi-drug-resistant), ‘XDR’ (extensively drug-resistant) or ‘PDR’ (pan-drug-resistant). These definitions are widely adopted (1) and known in the field of medical microbiology.

Other guidelines currently supported are the international EUCAST guideline (`guideline = “EUCAST”`, (15)), the international WHO guideline on the management of drug-resistant tuberculosis

(`guideline = “TB”`, (44)), and the national guidelines of The Netherlands (`guideline = “NL”`, (35)), and Germany (`guideline = “DE”`, (25)).

Some guidelines require a minimum availability of tested antimicrobial agents per isolate. This is needed to prevent false-negatives, since no reliable determination can be performed on only a few test results. This required minimum defaults to 50%, but can be set by the user with the `pct_minimum_classes`. Isolates that do not meet this requirement will be skipped for determination and will return NA (not applicable), with an informative warning printed to the console.

The rules are applied per row of the data. The `mdro()` function automatically identifies the variables containing the microorganism codes and antimicrobial agents based on the `guess_ab_col()` function. Following the guideline set by the user, it analyzes the specific antimicrobial resistance of a microorganism and flags that microorganism accordingly. The outcome is demonstrated in Table 5, where the first row is an MDRO according to the Dutch guidelines (35).

**Table 5:** Example of a multi-drug resistant organism (MDRO) in a data set identified by applying Dutch guidelines.

| mo | AMC | GEN | TOB | CIP | MFX | MDRO |
| --- | --- | --- | --- | --- | --- | --- |
| B_ESCHR_COLI | S | R | R | R | R | Positive |
| B_ESCHR_COLI | R | S | R | R | S | Negative |
| B_ESCHR_COLI | S | S | S | R | S | Negative |

Abbreviations: mo = microorganism, AMC = amoxicilline/clavulanic acid
GEN = gentamicin, TOB = tobramycin, CIP = ciprofloxacin, MFX = moxifloxacin,
MDRO = multi-drug resistant organism, B\_ESCHR\_COLI = microorganism code of
*Escherichia coli*

The returned value is an ordered `‘factor’` with the levels `‘Negative’ < ‘Positive, unconfirmed’ < ‘Positive’`. For some guideline rules that require additional testing (e.g., molecular confirmation), the level `‘Positive, unconfirmed’` is returned.

#### Multi-drug resistant tuberculosis

Tuberculosis is a major threat to global health caused by *Mycobacterium tuberculosis* (MTB) and is one of the top ten causes of death worldwide (46). Exceptional antimicrobial resistance in MTB is therefore of special interest. To this end, the international WHO guideline for the classification of drug resistance in MTB (44) is included in the **AMR** package. `The mdr_tb()` function is a convenient wrapper around `mdro(…, guideline = “TB”)`, which returns an other ordered `‘factor’` than other `mdro()` functions. The output will contain the `‘factor’` levels `‘Negative’ < ‘Mono-resistant’ < ‘Poly-resistant’ < ‘Multi-drug-resistant’ < ‘Extensive drugresistant’`, following the WHO guideline.

## 5. Analyzing antimicrobial resistance data

### 5.1. Calculation of antimicrobial resistance

The **AMR** package contains several functions for fast and simple resistance calculations of bacterial or fungal isolates. A minimum number of available isolates is needed for the reliability of the outcome. The CLSI guideline suggests a minimum of 30 available first isolates irrespective of the type of statistical analysis (6). Therefore, this number is used as the default setting for any function in the package that calculates antimicrobial resistance or susceptibility, which can be changed with the `minimum` argument in all applicable functions.

#### Counts

The **AMR** package relies on the concept of tidy data (39), although not strictly following its rules (one row per test rather than one row per observation). Function names to calculate the number of available isolates follow these general resistance interpretation standards with `count_S(), count_I()`, and `count_R()` respectively. Combinations of antimicrobial resistance interpretations can be counted with `count_SI()` and `count_IR()`. All these functions take a vector of interpretations of the class `‘rsi’` (as discussed above) or are internally trans-formed with `as.rsi()`. The returned value is the sum of the respective interpretation in the selected test column. All count_*() functions support quasi-quotation with pipes, grouped variables, and can be used with `dplyr::summarize()` (42).

#### Proportions

Calculation of antimicrobial resistance is carried out by counting the number of first resistant isolates (interpretation of “R”) and dividing it by the number of all first isolates, see Equation 2. This is implemented in the `proportion_R()` function. To calculate antimicrobial *susceptibility*, the number of susceptible first isolates (interpretation of “S” and “I”) has to be counted and divided by the number of all first isolates, which is implemented in the `proportion_SI()` function. For convenience, the `resistance()` function is an alias of the `proportion_R()` function, and the `susceptibility()` function is an alias of the `proportion_SI()` function.

The functions `proportion_R(), proportion_IR(), proportion_I(), proportion_SI()`, and `proportion_S()` follow the same logic as the `count_*()` functions and all return a vector of class `‘double’` with a value between 0 and 1. The argument `minimum` defines the minimal allowed number of available (tested) isolates (default: `minimum` = 30). Any number below the set `minimum` will return `NA` with a warning.

For calculating the proportion (*P*) of antimicrobial resistance or susceptibility to one antimicrobial agent, the following equation is used:

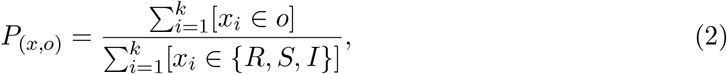

where *P* is the proportion of outcome *o* (that is either “R”, “S”, “I”, or a combination of two of them), where *x* is a character vector of length *k* only consisting of values “R”, “S”, or “I” and [*x*_*i*_ *∈o*] is the indicator function, returning 1 if the indicator function is true and 0 otherwise. The denominator must include the collection *R, S, I* so that ‘wrong’ elements in *x* (i.e., elements not being “R”, “S”, or “I”) will not be counted. Thus, the theoretical antimicrobial susceptibility of the vector *x* = *{S, S, I, R, R}* is:

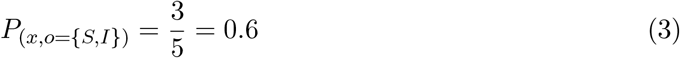

For the proportion of empiric susceptibility (*s*) for more than one antimicrobial agent, the calculation can be carried out in two ways (Table 6). The first method is to count the total number of first isolates where at least one agent was tested as “S” or “I” and divide it by the number of first isolates tested where any of the agents was tested (Equation 4). This method will be used when setting `only_all_tested = FALSE in the susceptibility()` function:

**Table 6:** Example calculation for determining empiric susceptibility (%SI) for more than one antimicrobial agent.

| Antimicrobial agent |  | All isolates<br>(only_all_tested = FALSE) |  | Only isolates tested for both agents<br>(only_all_tested = TRUE) |  |
| --- | --- | --- | --- | --- | --- |
| Agent A | Agent B | Include as<br>numerator | Include as<br>denominator | Include as<br>numerator | Include as<br>denominator |
| S or I | S or I | ✓ | ✓ | ✓ | ✓ |
| R | S or I | ✓ | ✓ | ✓ | ✓ |
| N/A | S or I | ✓ | ✓ |  |  |
| S or I | R | ✓ | ✓ | ✓ | ✓ |
| R | R |  | ✓ |  | ✓ |
| N/A | R |  |  |  |  |
| S or I | N/A | ✓ | ✓ |  |  |
| R | N/A |  |  |  |  |
| N/A | N/A |  |  |  |  |

Abbreviations: R = resistant, S = susceptible, I = susceptible, increased exposure, N/A = not tested or missing

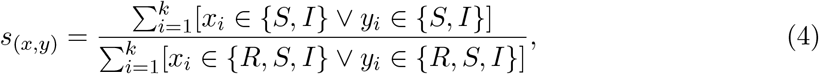

where *x* is a character vector only consisting of values “R”, “S”, or “I” (i.e., ‘agent A’) and *y* is another character vector only consisting of values “R”, “S”, or “I” (i.e., ‘agent B’).

The second method is to count the total number of first isolates where at least one agent was tested as “S” or “I” *and* where all agents were tested divided by the number of first isolates tested where *all* of the agents were tested (Equation 5). This method will be used when setting `only_all_tested = TRUE` in the `susceptibility()` function:

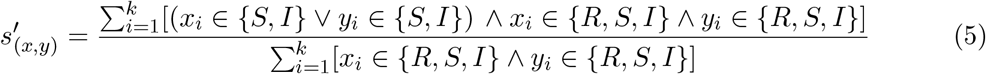

Based on Equation 2, the overall resistance and susceptibility of antimicrobial agents like gentamicin (GEN) and amoxicillin (AMX) can be calculated using the following syntax. The `example_isolates` is an example data set included in the **AMR** package, see appendix A. The n_rsi() function is analogous to the `n()` function of the **dplyr** package. It counts the number of available isolates, but only includes observations with valid antimicrobial results (i.e., “R”, “S”, or “I”).

~~~
*R> library(“dplyr”)
R> example_isolates %>%
+ summarize(r_gen = proportion_R(GEN)*,
~~~

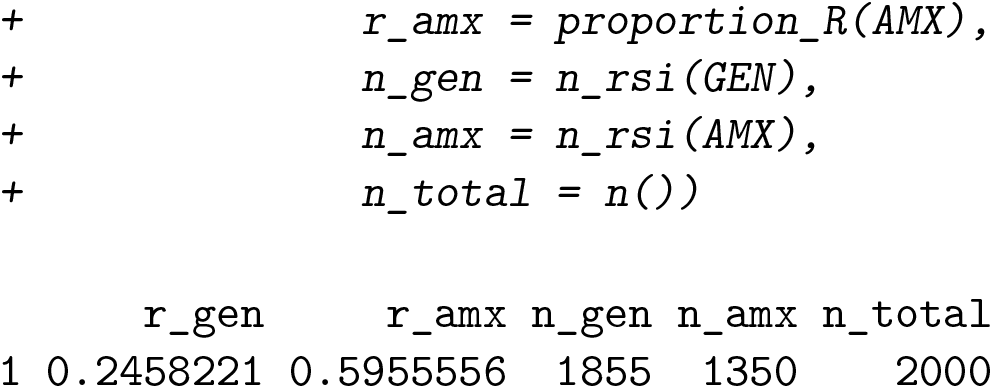

This output reads: the resistance to gentamicin of all isolates in the `example_isolates` data set is *P* (*x* = *GEN, o* = *{R}*) = 24.6%, based on 1855 out of 2000 available isolates. This means that the susceptibility is *P* (*x* = *GEN, o* = *{S, I}*) = 75.4%. The susceptibility to amoxicillin is *P* (*x* = *AMX, o* = *{S, I}*) = 40.4% based on 1350 isolates.

To calculate the effect of combination therapy, i.e., treating patients with multiple agents at the same time, all `proportion_*()` functions can handle multiple variables as arguments as defined in Equation 4 and 5. For example, to calculate the empiric susceptibility of a combination therapy comprising gentamicin (GEN) and amoxicillin (AMX):

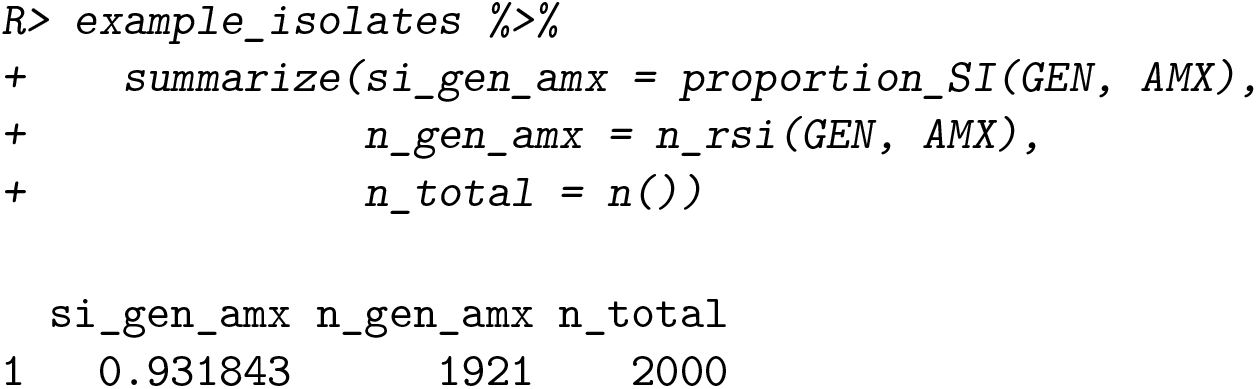

This leads to the conclusion that combining gentamicin with amoxicillin would cover *s*(*x* = *GEN, y* = *AMX*) = 93.2% based on 1921 out of 2000 available isolates, which is 17.8% more than when treating with gentamicin alone (*P* (*x* = *GEN, o* = *{S, I}*) = 75.4%). With these functions, exact calculations can be done to evaluate the empiric success of treating infections with one or more antimicrobial agents.

## 6. Design decisions

The **AMR** package follows the rationale of **tidyverse** packages as authored by (41). Most functions take a `‘data.frame’` or `‘tibble’` as input, support piping (%>%) operations, can work with quasi-quotations, and can be integrated into dplyr workflows, such as `mutate()` to create new variables and `group_by()` to group by variables. Although the **AMR** package integrates well into **tidyverse** workflows, it can also be used with base R only. To this extent, the **AMR** package was developed to be independent of any other R package to ensure and maintain sustainability.

The **AMR** package supports multiple languages. Currently supported languages are English, Dutch, French, German, Italian, Portuguese, and Spanish. The system language will be used if the language is supported but can be overwritten with `options(AMR_locale = …)`. Multi-language support affects language-dependent output of functions such as `mo_name(), mo_gramstain(), mo_type()`, and `ab_name()`.

The **AMR** package uses S3 classes, object oriented (OO) systems available in R. They allow different types of output based on the user input. The **AMR** package introduces 5 S3 classes (`‘mo’, ‘ab’, ‘rsi’, ‘mic’`, and `‘disk’`) to increase the convenience when working with antimicrobial susceptibility data.

## 7. Reproducible example

We consider the problem of working with antimicrobial resistance data from three different hospitals between 2011-01-01 and 2020-01-01. After loading the **AMR** package and additional tidyverse packages to allow transformation and plotting, we load the `example_isolates_unclean` example data from the **AMR** package into the global environment and assign it a new name.

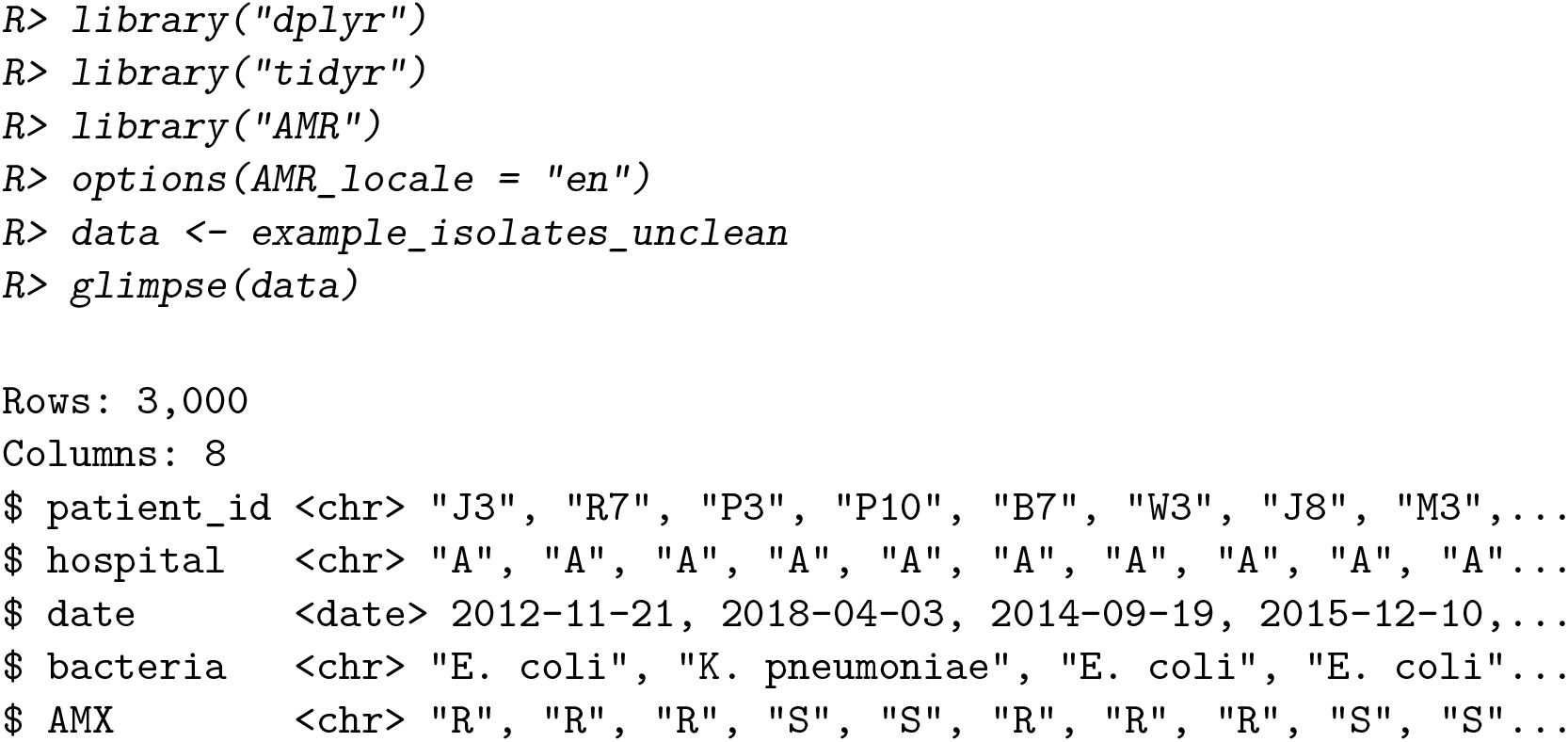
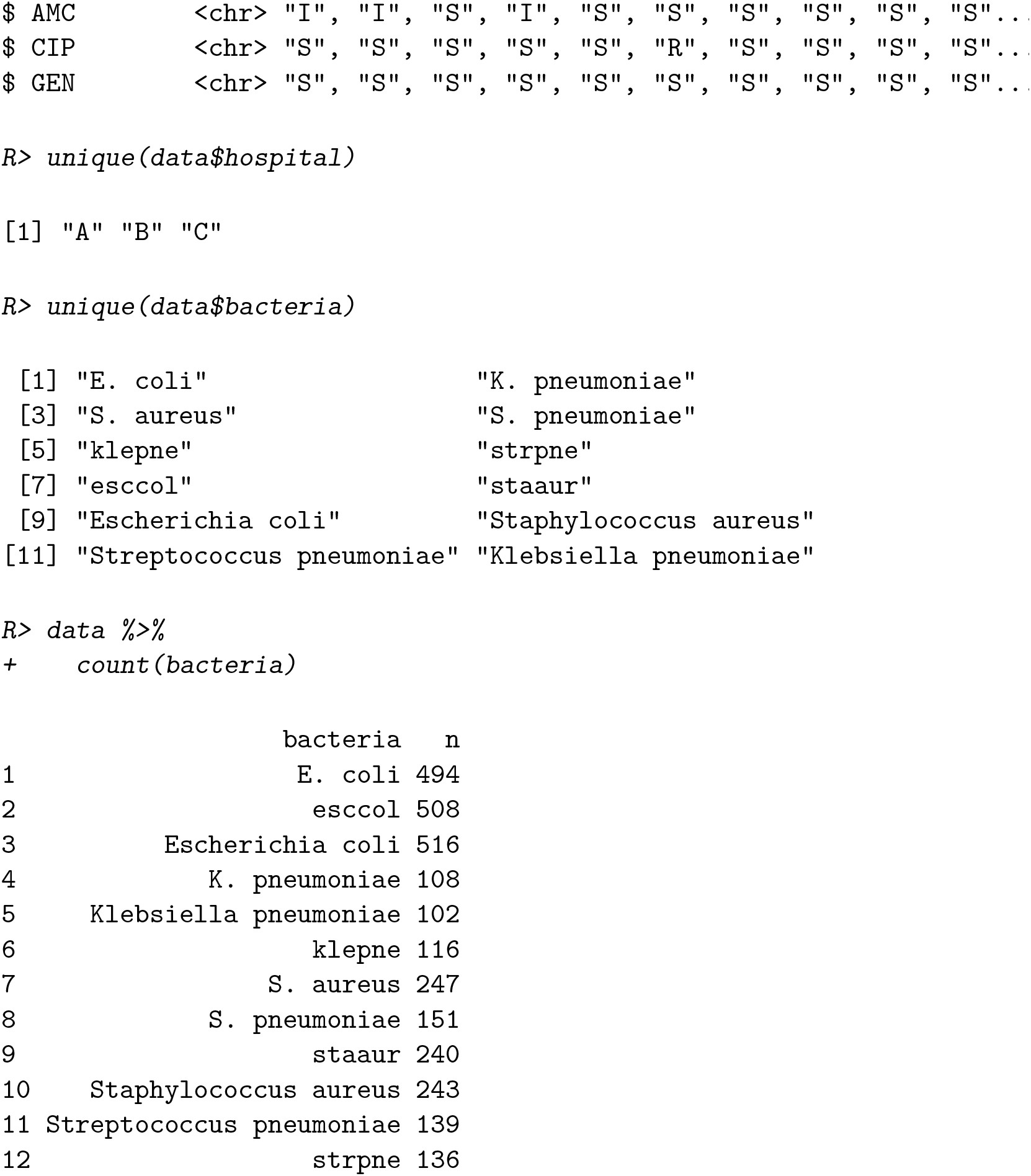

The data contains 3,000 observations of 8 variables from 3 hospitals. The “bacteria” variable comprises 12 unique elements. However, they appear to encode the same information in different formats (‘E. coli’, ‘K. pneumoniae’, ‘S. aureus’, ‘S. pneumoniae’, ‘klepne’, ‘strpne’, ‘esccol’, ‘staaur’, ‘Escherichia coli’, ‘Staphylococcus aureus’, ‘Streptococcus pneumoniae’, ‘Klebsiella pneumoniae’). We can use the `as.mo()` function to standardize the bacterial codes and add a variable with the official scientific name. The correct transformation of the bacterial codes can be reviewed by calling the `mo_uncertainties()` function.

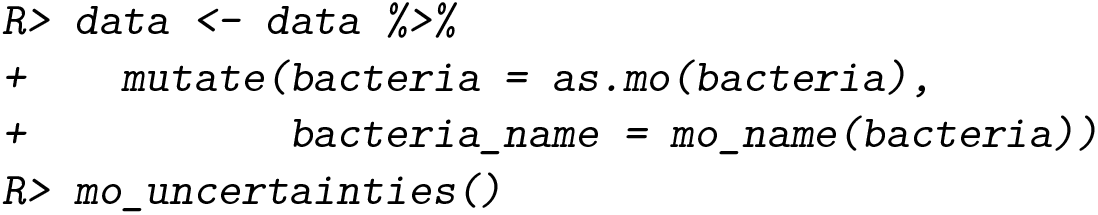
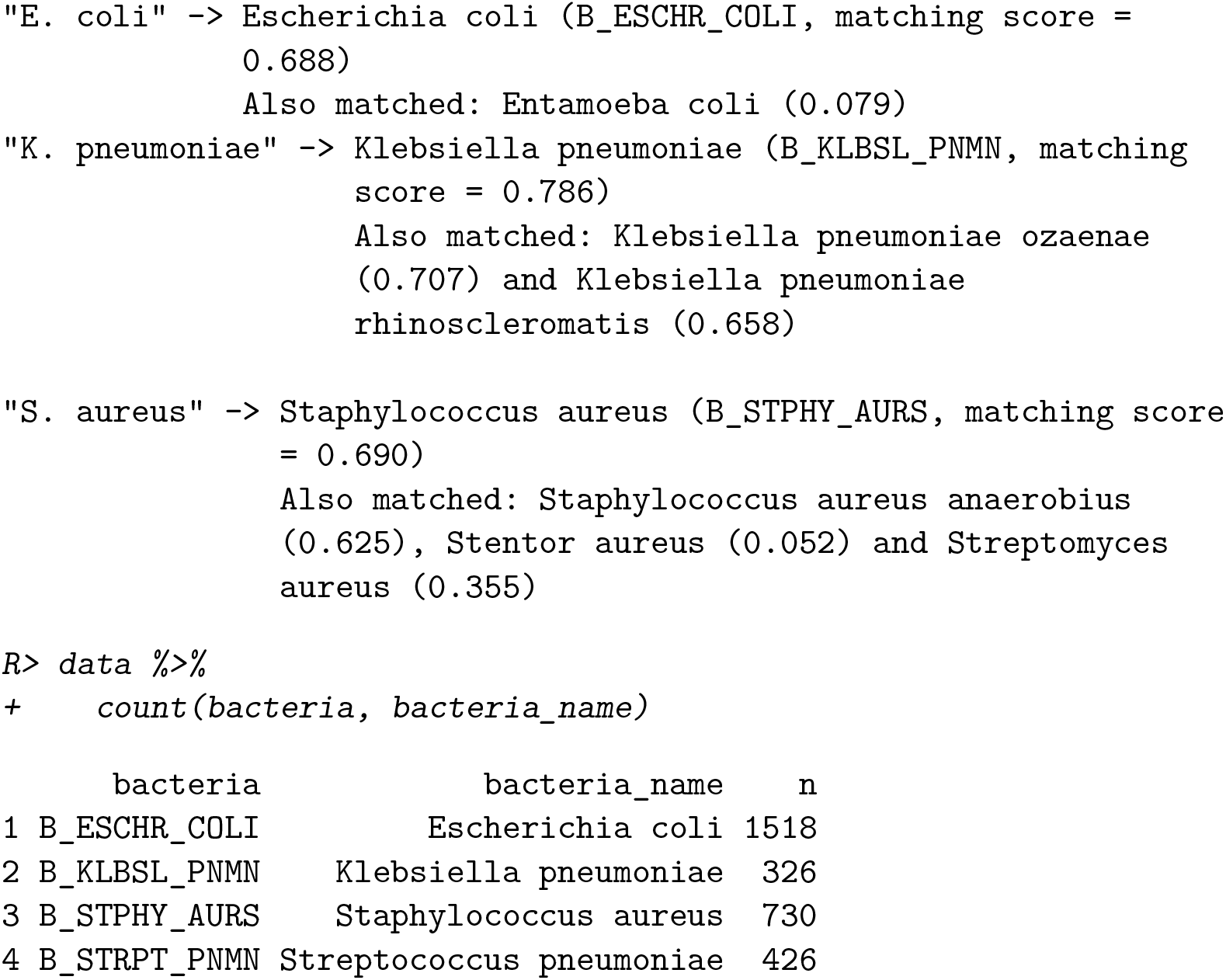

In a next step, we can further enrich the data with additional microbial taxonomic data based on the “bacteria” variable, such as Gram-stain and microorganism family.

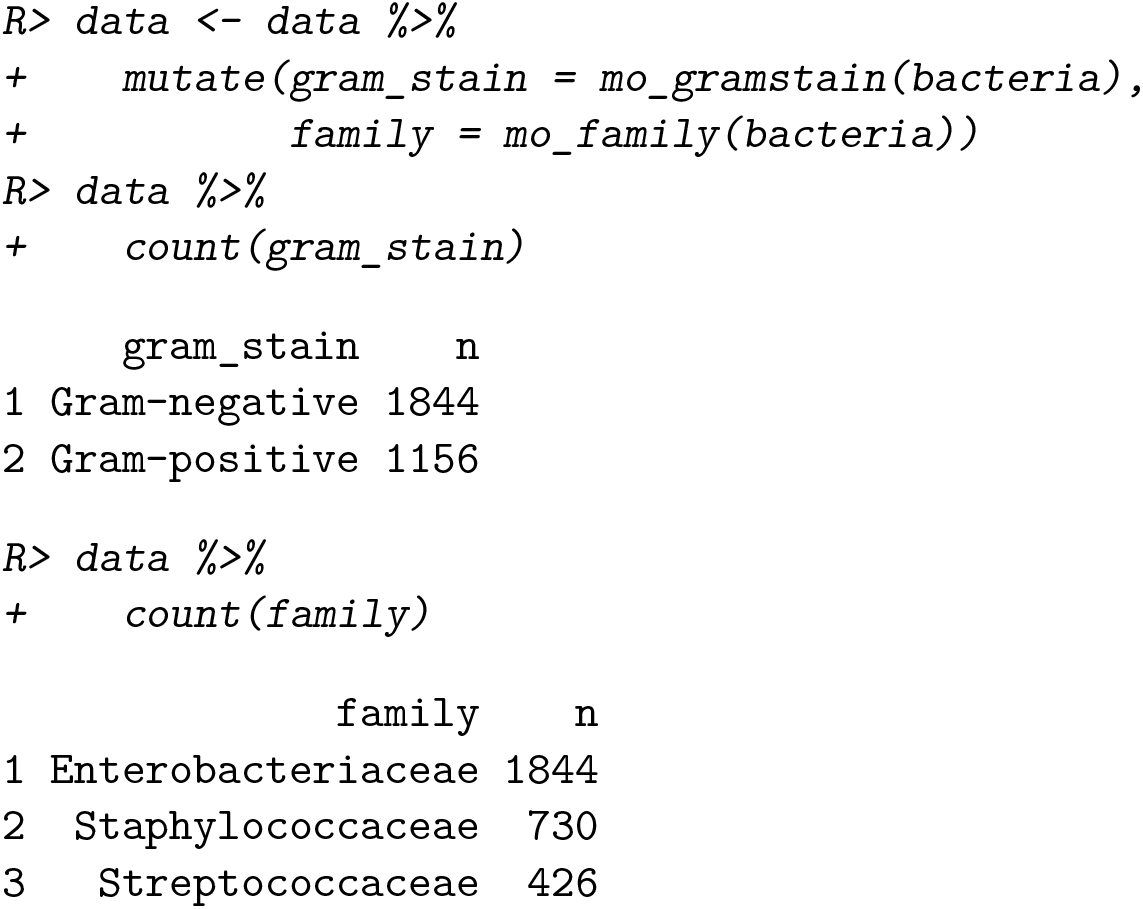

The variables “AMX”, “AMC”, “CIP”, and “GEN” contain antimicrobial susceptibility test results. The abbreviations stand for the tested antimicrobial agent. The official names and additional information about the antimicrobial agents can be checked with the `ab_info()` function from the **AMR** package.

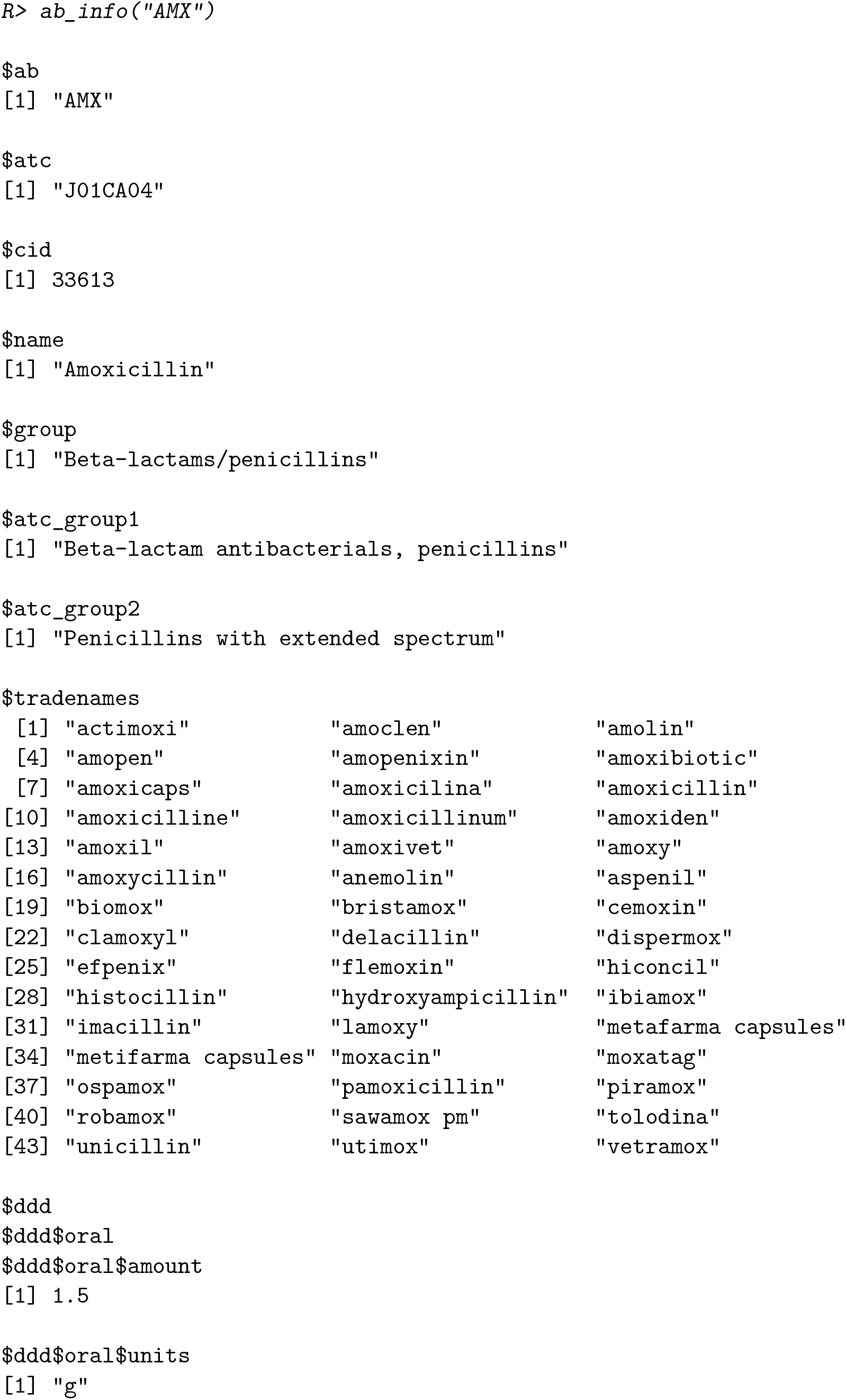
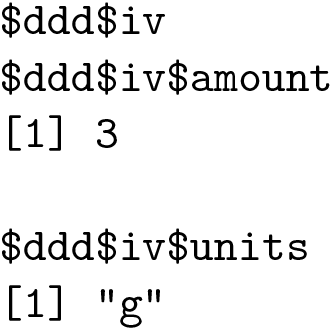

In a data set containing antimicrobial names or codes (e.g., antimicrobial prescription data), the `as.ab()` function can be used to transform all values to valid antimicrobial codes. Extra columns with the official name and the defined daily dose (DDD) for intravenous administration could be added using `ab_name()` and `ab_ddd()`.

|  | agents | agent_names | ddd_iv |
| --- | --- | --- | --- |
| 1 | AMX | Amoxicillin | 3.0 |
| 2 | CRO | Ceftriaxone | 2.0 |
| 3 | CIP | Ciprofloxacin | 0.8 |

Coming back to the cleaning of the data, the columns for the antimicrobial susceptibility test results (“AMX”, “AMC”, “CIP”, “GEN”) need to be checked to contain only standard values (“R”, “S”, “I”).

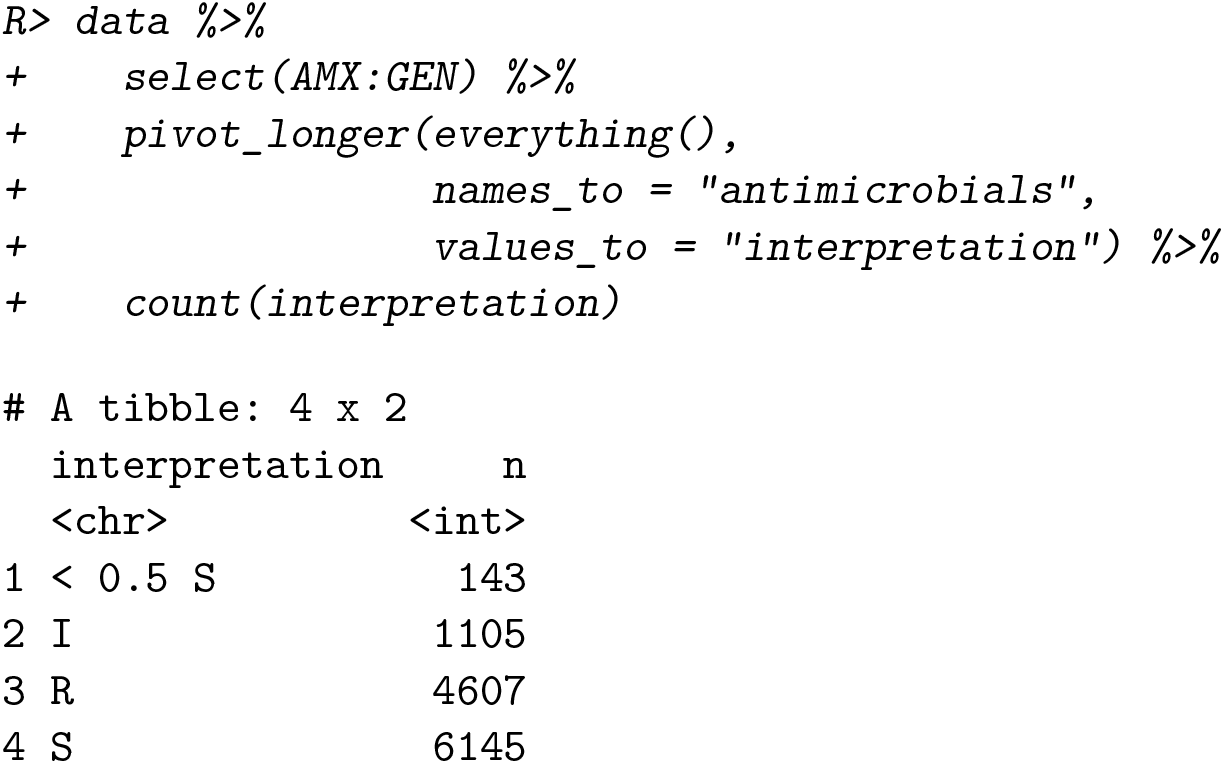

The values contain some mixed values. The `as.rsi()` function can be used to clean these values and to assign a new class (`‘rsi’`) for further use of **AMR** functions.

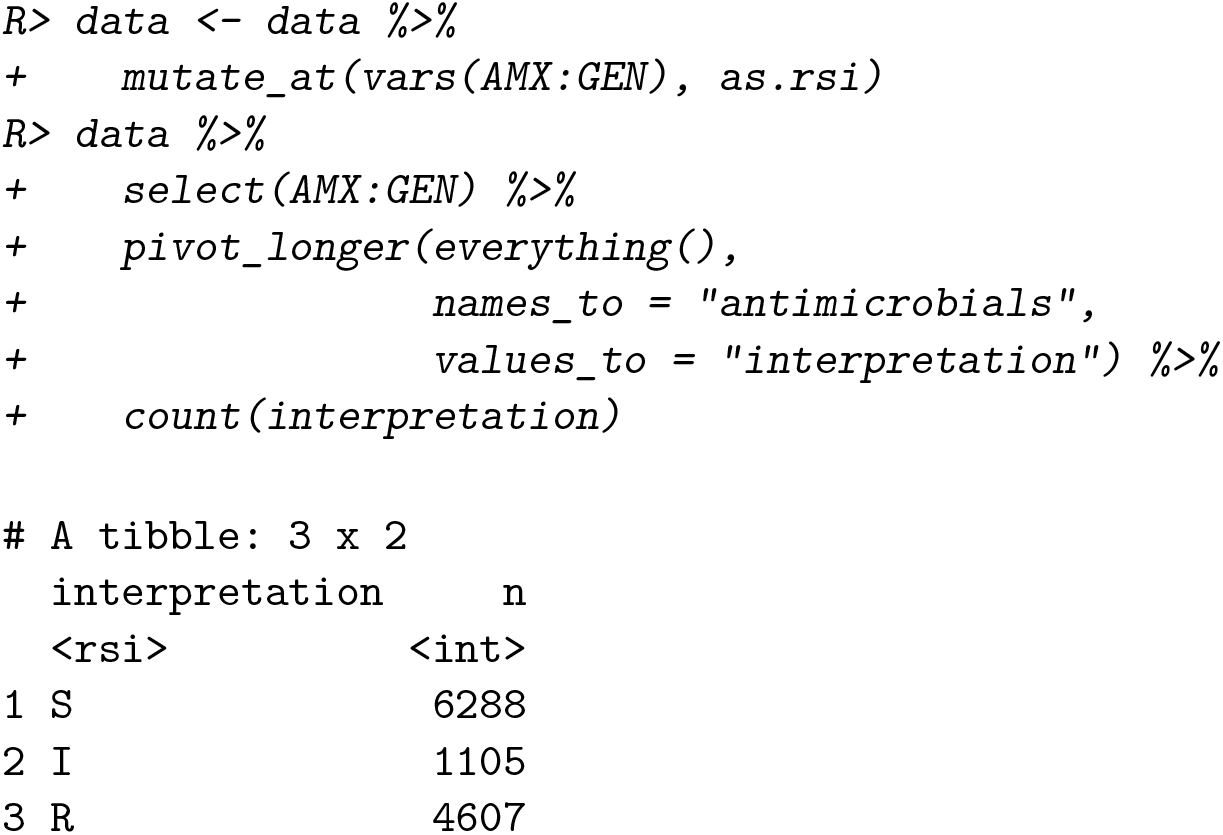

After this transformation, the `eucast_rules()` function can be applied to apply the latest resistance reporting guidelines.

~~~
*R> data <-data %>%
+ eucast_rules()*
~~~

The output to the console lists the changes made to data:

~~~
The rules affected 508 out of 3,000 rows, making a total of 657 edits
=> added 0 test results
=> changed 657 test results
  -11 test results changed from “S” to “I”
  -473 test results changed from “S” to “R”
  -85 test results changed from “I” to “R”
  -19 test results changed from “I” to “S”
  -33 test results changed from “R” to “I”
  -36 test results changed from “R” to “S”
~~~

The data is now clean and ready for further analysis, for example, the identification of multi-drug resistant microorganisms. In this example, we use the Dutch guideline to determine multi-drug resistance (35)).

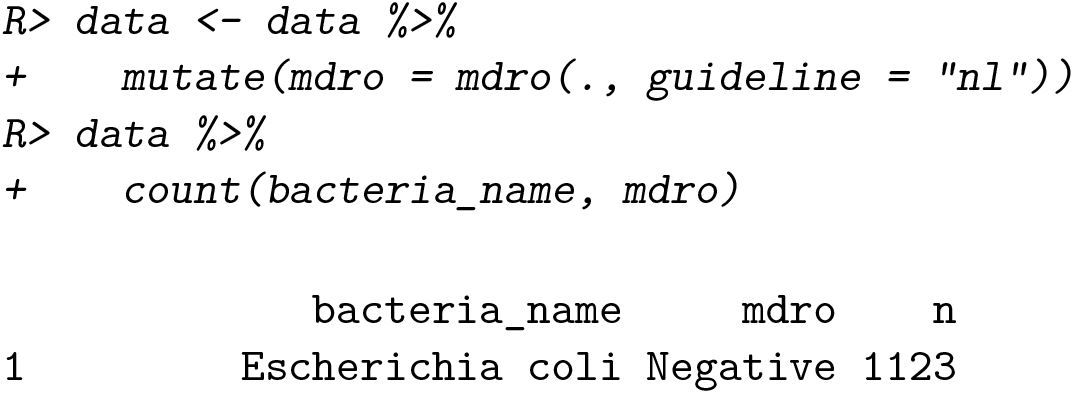
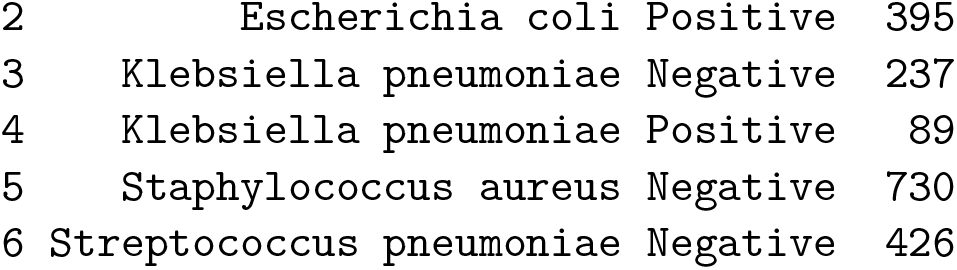

According to the Dutch guideline, 484 multi-drug resistant microorganisms were found in 3000 tested isolates. No multi-drug resistance was found in *Staphylococcus aureus* and *Streptococcus pneumoniae*.

As described in Section 4.1, the identification of first isolates is essential for the reporting of resistance patterns. Using the `filter_first_isolate()` function and `proportion_df()` in combination with `group_by()`, we get a complete resistance analysis per hospital, bacteria, first isolate, and tested antimicrobial agent in one call:

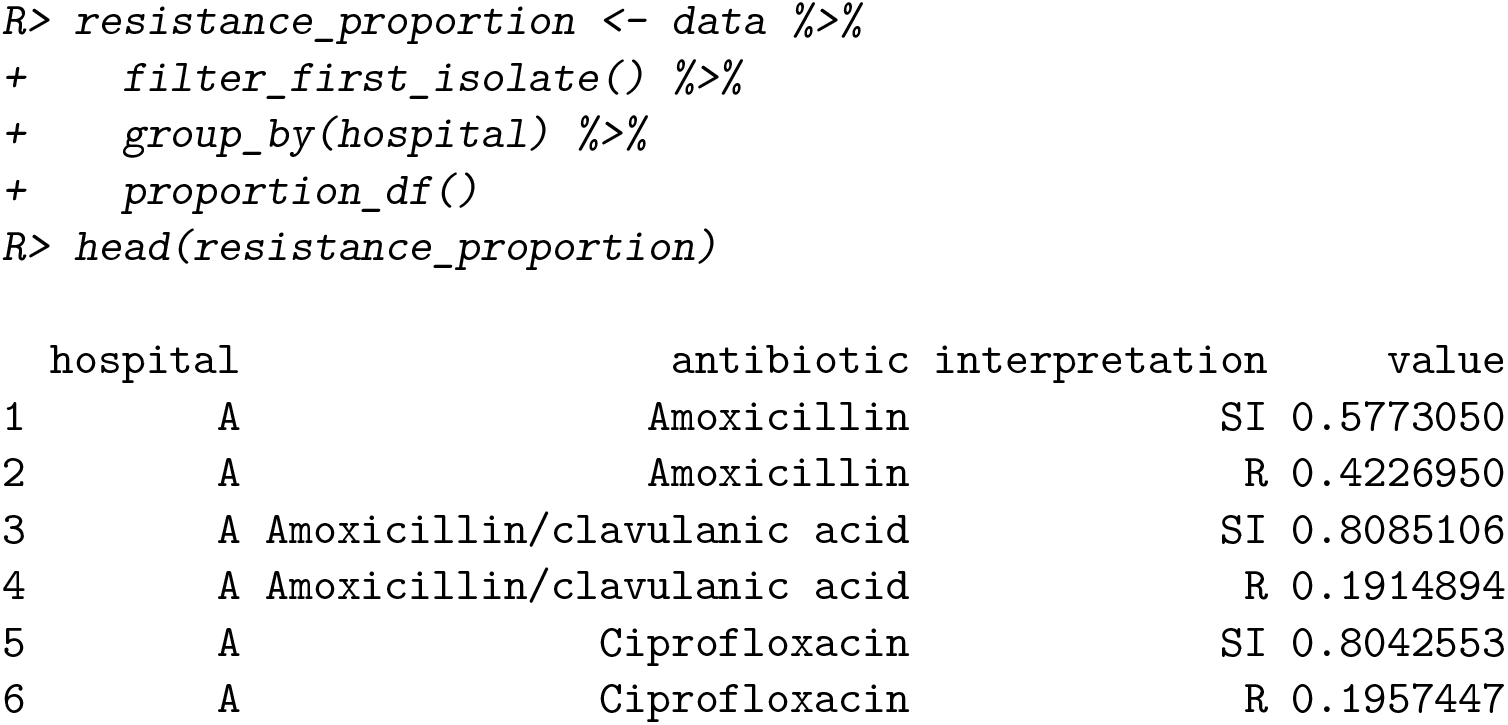

From the console we get the information how many first isolates were identified and used in the filter.

From here on, the data is ready for further analysis with functions for plotting (e.g., the ggplot2 package, (40)), **AMR** extension functions for base R (e.g., `summary(), plot()`), or **AMR** helper functions for plotting and basic modelling (e.g., `ggplot_rsi(), geom_rsi(), resistance_predict()`).

## 8. Discussion

For the first time, a free and open source software solution is available to cover all aspects of working with antimicrobial resistance data. The **AMR** package provides functionalities that enable standardized and reproducible workflows from raw laboratory data to publishable results, for research and clinical workflows alike. In the field of clinical microbiology and infectious diseases, research and clinical workflows are closely linked. For example, a performed research study on the prevalence of antimicrobial-resistant bacteria can have direct implications on the choice of antimicrobial agents for the treatment of patients. The **AMR** package was developed to be used in any research or clinical setting where the data analysis on microorganisms, antimicrobial resistance, antimicrobial agents is required.

Both, researchers and clinicians rely on the data from electronic laboratory information systems (LIS) where laboratory test results are processed, stored, and archived. Although some commercial solutions exist to conduct medical microbiological data analysis, these solutions are not comprehensive enough to apply antimicrobial resistance analysis for any clinical or research setting. Costs of these tools are a further constraint in resource-limited settings. Moreover, researchers and clinicians that require data from multiple LIS sources to be used in multi-center studies experience major barriers which cannot be solved by available commercial solutions.

Firstly, simple codes for microorganisms show substantial differences between different LIS and presumably correct taxonomic names are often misspelled or outdated. We analyzed the taxonomic names of bacteria used in reports from seven different public health institutions that perform microbiological diagnostics in the Netherlands and compared them with an official scientific up-to-date source for microbial taxonomy, the Catalogue of Life (30). These institutions cover microbiological diagnostics for hospitals and primary care for 15% of the total Dutch population. All institutions reported outdated taxonomic names with a maximum lag ranging between 34 and 41 years. Given that antimicrobial resistance guidelines are strongly based on the microbial taxonomy (some rules only apply to a specific genus, other rules apply to a specific family), it is crucial that this information is correct and timely updated. All institutions admitted that there was no standard operating procedure to maintain their taxonomic reference data. Implementing and maintaining the taxonomic data for these and other institutions has been challenging, since no common machine-readable, reliable and up-to-date resource for the microbial taxonomy was publicly available. For reliable reference data about antimicrobial agents, this also holds true. The **AMR** package provides machinereadable reference data files for the complete microbial taxonomy and for more than 500 antimicrobial agents. Using functions starting with `mo_*` and `ab_*`, names of microorganisms and antimicrobial agents can be translated between different LIS codes or other forms of text codes for microorganisms and consequently allows to merge data sets from different sites with little effort.

Secondly, antimicrobial resistance interpretation guidelines (18; 6) and taxonomic definitions of microorganisms are under constant change and are continually published in dedicated peer-reviewed journals. This is further complicated by differences between local, regional, and national guidelines. Yet, comparability and reproducibility across setting and time are key in research and clinics. The **AMR** package functions `eucast_rules()` (to apply guidelines to data), `mdro()` (to check for multi-drug resistance according to guidelines), or `first_isolate()` (to determine first isolates according to guidelines) address the needs to standardize comparability, and empower data analysts beyond the capabilities of their local LIS. The **AMR** package can be used as an extra layer of data validation when retrieving raw data from a LIS. Overall, the functionality of the **AMR** package has the potential to improve data validity in clinical settings, to ease multi-center study workflows, and to foster research reporting practices. The inherent global nature of antimicrobial resistances requires researchers, clinicians, and policy makers to reach beyond the borders of their local laboratory. The **AMR** package can build the bridge to link these sources and further encourages open science principles through its open source approach.

The **AMR** package also has limitations. It does not introduce novel statistical tests or models, nor does it add additional analytical approaches for **AMR** research. The calculation of the proportion of susceptibility for more than one antimicrobial agent simultaneously (see Section 5.1) seems simple but is subject to unclear reporting in clinical practice (32; 23). The lack of clearly defined algorithms can lead to the effect that co-resistance rates for more than one antimicrobial agent are dropped altogether (2). The inclusion of isolates that are tested for some agents (`only_all_tested = FALSE`) or only isolates tested for all agents (`only_all_tested = TRUE`) can have an imminent clinical impact on patient care, if one combination of antimicrobial agents is preferred over another. Therefore, the **AMR** package provides different algorithms to standardize this crucial calculation. Unfortunately, unambiguous methodology for determining the right algorithm is lacking in scientific literature. An analysis on the algorithms used in the **AMR** package and their clinical impact is in preparation.

Reliable information about antimicrobial resistance is vital for clinical decision-making in infectious diseases, since the outcome of local antimicrobial resistance analyses support medical professionals/clinicians in the treatment choices for their patients. Moreover, when this information can be reliably stratified by, for example, year, hospital, and type of patients, new information can lead to new insights for choosing the best antimicrobial therapy for patients suffering from infections. The **AMR** package enables this by providing all required analysis tools and can therefore empower decision-making in infectious management. The **AMR** package is already being applied to this end in six hospitals in the Netherlands. The choice of empirical antimicrobial treatment (meaning; choosing the initial therapy at a time of not knowing the infection-causing pathogen) for septic non-post-surgical patients has been altered in at least one Dutch hospital, by analyzing antimicrobial resistance data with the **AMR** package. The clinical effect of this adjustment is being studied at the moment. To improve the quality of such analyses, planned future developments comprise the implementations of an imputation algorithm specifically for antimicrobial agents, and method guidance for applying prediction modelling in a health care setting based on patient-specific properties.

Since the first package release, users from different public and private settings have been suggesting additional functionalities, in particular, the incorporation of country-or time-specific guidelines (e.g., (24)). This community-centered development will be continued and maintained by researchers at the University Medical Center Groningen and data scientists at Certe Medical Diagnostics and Advice, both non-profit public health organizations located in Groningen, the Netherlands. Moreover, a group of contributors from five different Dutch health care institutions has been formed at the Dutch Association for Medical Microbiology (Nederlandse Vereniging voor Medische Microbiologie -NVMM) that also peer-review major changes to the package, including the implementation of guideline updates. This way, updates required for scientific developments as well as maintaining consistent reproducibility are ensured. Updates to databases and guidelines included in the **AMR** package are incorporated on a regular and automated basis, while preserving version control. Any function making use of guidelines (e.g., `eucast_rules()`) refers to the latest implemented version of the guideline by default.

The aim of the **AMR** package is to provide a comprehensive toolbox of solutions for antimicrobial resistance data processing and analysis on an institution-and country-independent scale for clinical practice and research that are required according to international standards, but were not available to date.

## 9. Summary

This paper demonstrates the **AMR** package and its use for working with antimicrobial resistance data. It can be used to clean, enhance, and analyze such data according to (inter)national recommendations and guidelines while incorporating scientifically reliable reference data on microbiological laboratory test results, antimicrobial agents, and the biological taxonomy of microorganisms. Consequently, it allows for reproducible analyses, regardless of the many possible ways in which raw and uncleaned data are stored in laboratory information systems.

While the burden of antimicrobial resistance is increasing worldwide, reliable data and data analyses are needed to better understand current and future developments. Open source approaches, such as the **AMR** package for R, have the potential to help democratizing the required tools in the field for researchers, clinicians, and policy makers alike. In organizations or countries with very limited resources, this free and open-source package could also overcome a financial limitation that would otherwise hinder antimicrobial resistance analysis in these settings. Across settings, we believe the **AMR** package can be used to support clinical decision-making in infection management by providing improved insight into current local and regional resistance levels. Furthermore, data analysis approaches based on individual patient or microbiological data, which the **AMR** package enables, fosters empowerment of laboratory staff, infection control practitioners, and public health services.

## Computational details

The results in this paper were obtained using 4.0.2 in RStudio 1.3.1093 (31) with the **AMR** package 1.5.0, running under macOS Catalina 10.15.

R itself and all packages used are available from the Comprehensive R Archive Network (CRAN) at https://CRAN.R-project.org/. All development versions of the **AMR** package are available at https://github.com/msberends/AMR/.

## Supporting information

Source package

Complete reproducible example

## Acknowledgments

The authors Matthijs S. Berends and Christian F. Luz contributed equally to this publication. For their contributions to the development of the **AMR** package, we would like to thank (in alphabetical order) Judith M. Fonville, Erwin E.A. Hassing, Eric H.L.C.M. Hazenberg, Gwen Knight, Annick Lenglet, Bart C. Meijer, Sofia Ny, Rogier P. Schade, Dennis Souverein, and Anthony Underwood.

The development of the **AMR** package was partly supported by the INTERREG V A (202085) funded project EurHealth-1Health (http://www.eurhealth1health.eu), part of a Dutch-German cross-border network supported by the European Commission, the Dutch Ministry of Health, Welfare and Sport, the Ministry of Economy, Innovation, Digitalization and Energy of the German Federal State of North Rhine-Westphalia and the Ministry for National and European Affairs and Regional Development of Lower Saxony.

Furthermore, the **AMR** package was developed as part of a project funded by the European Commission Horizon 2020 Framework Marie Skłodowska-Curie Actions (grant agreement number: 713660-PRONKJEWAIL-H2020-MSCA-COFUND-2015).

The funders had no role in study design, data collection and analysis, decision to publish, or preparation of the manuscript.

## A. Included data sets

- `microorganisms` A `‘data.frame’` containing 67,151 (sub)species with 16 columns comprising their complete microbial taxonomy according to the Catalogue of Life (30). Included microorganisms and their complete taxonomic tree of all included (sub)species from kingdom to subspecies with year of scientific publication and responsible author(s):
  – All 55,415 (sub)species from the kingdoms of Archaea, Bacteria, Chromista and Protozoa;
  – All 9,^582^ (sub)species from these orders of the kingdom of Fungi: Eurotiales, Onygenales, Pneumocystales, Saccharomycetales, Schizosaccharomycetales and Tremellales;
  – All 2,153 (sub)species from 47 other relevant genera from the kingdom of Animalia (like *Strongyloides* and *Taenia*);
  – All 12,708 previously accepted names of included (sub)species that have been taxonomically renamed.
  The kingdom of Fungi is a very large taxon with almost 300,000 different (sub)species, of which most are not microbial (but rather macroscopic, such as mushrooms). Therefore, not all fungi fit the scope of the **AMR** package. By only including the aforementioned taxonomic orders, the most relevant fungi are covered (such as all species of *Aspergillus, Candida, Cryptococcus, Histoplasma, Pneumocystis, Saccharomyces* and *Trichophyton*).
- `antibiotics` A `‘data.frame’` containing 456 antibiotic agents with 14 columns. All entries in this data set have three different identifiers: a human readable EARS-Net code (as used by ECDC (13) and WHONET (38) and primarily used by this package), an ATC code (as used by the WHO (37)), and a CID code (Compound ID, as used by PubChem (17)). The data set contains more than 5,000 official brand names from many different countries, as found in PubChem. Other properties in this data set are derived from one or more of these codes, such as official names of pharmacological and chemical subgroups, and defined daily doses (DDD).
- `antivirals` A `‘data.frame’` containing 102 antiviral agents with 9 columns. Like the `antibiotics` data set, it contains ATC codes (as used by the WHO (37)), and a CID code (Compound ID, as used by PubChem (17)), as well as the official name and defined daily dose (DDD) for each antiviral agent.
- `example_isolates` A `‘data.frame’` containing test results of 2,000 microbial isolates. The data set reflects real patient data and can be used to practice **AMR** analysis. It is structured in the typical format of laboratory information systems with one row per isolate and one column per tested antimicrobial agent (i.e., an antibiogram).
- `example_isolates_unclean` A `‘data.frame’` containing test results of 3,000 microbial isolates that require cleaning up before they can be used for analysis. This data set can be used to practice **AMR** analysis and is featured in section 7.
- `WHONET` A `‘data.frame’` containing 500 observations and 53 columns, with the exact same structure as an export file from WHONET 2019 software (38). Such files can be used with the **AMR** package, as this example data set demonstrates. The antibiotic test results are from the `example_isolates` data set. All patient names are created using online surname generators and are only in place for practice purposes.

## References

[1] Abat C, Fournier PE, Jimeno MT, Rolain JM, Raoult D (2018). “Extremely And Pandrug-resistant Bacteria Extra-deaths: Myth Or Reality?” European Journal of Clinical Microbiology & Infectious Diseases, 37(9), 1687–1697. ISSN 0934-9723. doi:10.1007/s10096-018-3300-0. URL www.ncbi.nlm.nih.gov/pubmed/29956024http://link.springer.com/10.1007/s10096-018-3300-0.

[2] Baur D, Gladstone BP, Burkert F, Carrara E, Foschi F, Döbele S, Tacconelli E (2017). “Effect Of Antibiotic Stewardship On The Incidence Of Infection And Colonisation With Antibiotic-resistant Bacteria And Clostridium Difficile Infection: A Systematic Review And Meta-analysis.” The Lancet Infectious Diseases, 17(9), 990–1001. ISSN 14744457. doi:10.1016/S1473-3099(17)30325-0.

[3] Berends MS, Luz CF, Friedrich AW, Sinha BNM, Albers CJ, Glasner C (2020). AMR: Antimicrobial Resistance Analysis. R package version 1.5.0, URL https://CRAN.R-project.org/package=AMR.

[4] Cavalier-Smith T (2002). “The Neomuran Origin Of Archaebacteria, The Negibacterial Root Of The Universal Tree And Bacterial Megaclassification.” International journal of systematic and evolutionary microbiology, 52(Pt 1), 7–76. ISSN 1466-5026. DOI: 10.1099/00207713-52-1-7. URL http://www.ncbi.nlm.nih.gov/pubmed/11837318.

[5] CDC (2019). “AR Threats Report: Antibiotic Resistance Threats In The United States, 2019.” https://www.cdc.gov/drugresistance/pdf/threats-report/ 2019-ar-threats-report-508.pdf.

[6] Clinical and Laboratory Standards Institute (2014). “M39-A4, Analysis And Presentation Of Cumulative Antimicrobial Susceptibility Test Data, 4th Edition.”

[7] Clinical and Laboratory Standards Institute (2019). “Susceptibility Testing Of Infectious Agents And Evaluation Of Performance Of Antimicrobial Susceptibility Test Devices – Part 1, 2nd Edition.”

[8] Courvalin P (1992). “Interpretive Reading Of Antimicrobial Susceptibility Tests. Molecular Analysis And Therapeutic Interpretation Of In Vitro Tests To Improve Antibiotic Therapy.” ASM American Society for Microbiology News, 58(7), 368–375.

[9] Courvalin P (1996). “Interpretive Reading Of In Vitro Antibiotic Susceptibility Tests (the Antibiogramme).” Clin. Microbiol. Infect., 2, S26–S34. doi:10.1111/j.1469-0691.1996.tb00872.x.

[10] de Greeff SC, Mouton JW, Schoffelen AF, Verduin CM (2019). “NethMap 2019: Consumption Of Antimicrobial Agents And Antimicrobial Resistance Among Medically Important Bacteria In The Netherlands / maran 2019: Monitoring Of Antimicrobial Resistance And Antibiotic Usage In Animals In The Netherlands In 2018.”

[11] de Kraker MEA, Stewardson AJ, Harbarth S (2016). “Will 10 Million People Die A Year Due To Antimicrobial Resistance By 2050?” PLoS Med., 13(11), e1002184. DOI: 10.1371/journal.pmed.1002184.

[12] EUCAST (2019). “EUCAST New Definitions Of S, I And R From 2019.” http://www.eucast.org/newsiandr/. Accessed: 2019-12-02.

[13] European Centre for Disease Prevention and Control (2010). “European Antimicrobial Resistance Surveillance Network (ears-net).” https://ecdc.europa.eu/en/about-us/ partnerships-and-networks/disease-and-laboratory-networks/ears-net. Accessed: 2019-4-9.

[14] European Centre for Disease Prevention and Control (2018). “Antimicrobial Resistance (amr) Reporting Protocol 2018. European Antimicrobial Resistance Surveillance Network (ears-net) Surveillance Data For 2017.”

[15] European Committee on Antimicrobial Susceptibility Testing (EUCAST) (2016). “EUCAST Expert Rules, Intrinsic Resistance And Exceptional Phenotypes Tables. Version 3.1, 2016.” http://www.eucast.org/fileadmin/src/media/PDFs/EUCAST_files/Expert_Rules/Expert_rules_intrinsic_exceptional_V3.1.pdf.

[16] Jombart T, Kamvar ZN, FitzJohn R, Cai J, Bhatia S, Schumacher J, Pulliam JR (2020). incidence: Compute, Handle, Plot and Model Incidence of Dated Events. R package version 1.7.3, URL https://doi.org/10.5281/zenodo.2584018.

[17] Kim S, Chen J, Cheng T, Gindulyte A, He J, He S, Li Q, Shoemaker BA, Thiessen PA, Yu B, Zaslavsky L, Zhang J, Bolton EE (2019). “PubChem 2019 Update: Improved Access To Chemical Data.” Nucleic Acids Res., 47(D1), D1102–D1109. doi:10.1093/nar/gky1033.

[18] Leclercq R, Cantón R, Brown DFJ, Giske CG, Heisig P, MacGowan AP, Mouton JW, Nordmann P, Rodloff AC, Rossolini GM, Soussy CJ, Steinbakk M, Winstanley TG, Kahlmeter G (2013). “EUCAST Expert Rules In Antimicrobial Susceptibility Testing.” Clin. Microbiol. Infect., 19(2), 141–160. doi:0.1111/j.1469-0691.2011.03703.x.

[19] Levenshtein VI (1966). “Binary codes capable of correcting deletions, insertions, and reversals.” Soviet physics doklady, 10(8), 707–710.

[20] Limmathurotsakul D, Dunachie S, Fukuda K, Feasey NA, Okeke IN, Holmes AH, Moore CE, Dolecek C, van Doorn HR, Shetty N, Lopez AD, Peacock SJ, Surveillance and Epidemiology of Drug Resistant Infections Consortium (SEDRIC) (2019). “Improving The Estimation Of The Global Burden Of Antimicrobial Resistant Infections.” Lancet Infect. Dis. doi:10.1016/S1473-3099(19)30276-2.

[21] Livermore DM, Winstanley TG, Shannon KP (2001). “Interpretative Reading: Recognizing The Unusual And Inferring Resistance Mechanisms From Resistance Phenotypes.” J. Antimicrob. Chemother., 48 Suppl 1, 87–102. doi:10.1093/jac/48.suppl_1.87.

[22] Lowndes JSS, Best BD, Scarborough C, Afflerbach JC, Frazier MR, O’Hara CC, Jiang N, Halpern BS (2017). “Our Path To Better Science In Less Time Using Open Data Science Tools.” Nat Ecol Evol, 1(6), 160. doi:10.1038/s41559-017-0160.

[23] Ma J, Li N, Liu Y, Wang C, Liu X, Chen S, Xie X, Gan S, Wang M, Cao W, Wang F, Liu Y, Wan D, Sun L, Sun H (2017). “Antimicrobial Resistance Patterns, Clinical Features, And Risk Factors For Septic Shock And Death Of Nosocomial E Coli Bacteremia In Adult Patients With Hematological Disease.” Medicine (United States), 96(21). ISSN 15365964. doi:10.1097/MD.0000000000006959.

[24] Magiorakos AP, Srinivasan A, Carey RB, Carmeli Y, Falagas ME, Giske CG, Harbarth S, Hindler JF, Kahlmeter G, Olsson-Liljequist B, Paterson DL, Rice LB, Stelling J, Struelens MJ, Vatopoulos A, Weber JT, Monnet DL (2012). “Multidrug-resistant, Extensively Drug-resistant And Pandrug-resistant Bacteria: An International Expert Proposal For Interim Standard Definitions For Acquired Resistance.” Clinical microbiology and infection : the official publication of the European Society of Clinical Microbiology and Infectious Diseases, 18(3), 268–81. ISSN 1469-0691. doi:10.1111/j.1469-0691.2011.03570.x. URL dx.doi.org/10.1111/j.1469-0691.2011.03570.xhttp://www.ncbi.nlm.nih.gov/pubmed/21793988.

[25] Müller J, Voss A, Köck R, Sinha B, Rossen JW, Kaase M, Mielke M, Daniels-Haardt I, Jurke A, Hendrix R, Kluytmans JA, Kluytmans-van den Bergh MF, Pulz M, Herrmann J, Kern WV, Wendt C, Friedrich AW (2015). “Cross-border Comparison Of The Dutch And German Guidelines On Multidrug-resistant Gram-negative Microorganisms.” Antimicrobial resistance and infection control, 4, 7. ISSN 2047-2994. doi:10.1186/s13756-015-0047-6. URL www.ncbi.nlm.nih.gov/pubmed/25763183http://www.pubmedcentral.nih.gov/articlerender.fcgi?artid=PMC4355569.

[26] Nagraj V, Jombart T, Randhawa N, Sudre B, Campbell F, Crellen T (2017). epicontacts: Handling, Visualisation and Analysis of Epidemiological Contacts. R package version 1.1.0, URL https://CRAN.R-project.org/package=epicontacts.

[27] O’Neill J (2014). “Antimicrobial Resistance: Tackling A Crisis For The Health And Wealth Of Nations.”

[28] Petzoldt T (2019). antbioticR: Analysis of Antibiotic Resistance Data. R package version 0.3.2, URL https://github.com/tpetzoldt/antibioticR.

[29] R Core Team (2020). R: A Language and Environment for Statistical Computing. R Foundation for Statistical Computing, Vienna, Austria. URL https://www.R-project. org/.

[30] Roskov Y, Ower G, Orrell T, Nicolson D, Bailly N, Kirk PM, Bourgoin T, DeWalt RE, Decock W, Nieukerken Ev, Zarucchi J, Penev L (2019). “Species 2000 & itis Catalogue Of Life, 25th March 2019.” Digital resource at www.catalogueoflife.org/col. Species 2000: Naturalis.

[31] RStudio Team (2020). Rstudio: Integrated Development Environment For R. RStudio, PBC, Boston, MA. URL http://www.rstudio.com/.

[32] Schechner V, Temkin E, Harbarth S, Carmeli Y, Schwaber MJ (2013). “Epidemiological Interpretation Of Studies Examining The Effect Of Antibiotic Usage On Resistance.” Clinical Microbiology Reviews, 26(2), 289–307. ISSN 08938512. doi:10.1128/CMR.00001-13.

[33] Tacconelli E, Sifakis F, Harbarth S, Schrijver R, van Mourik M, Voss A, Sharland M, Rajendran NB, Rodríguez-Baño J, EPI-Net COMBACTE-MAGNET Group (2018). “Surveillance For Control Of Antimicrobial Resistance.” Lancet Infect. Dis., 18(3), e99– e106. doi:10.1016/S1473-3099(17)30485-1.

[34] The European Committee on Antimicrobial Susceptibility Testing (2020). “Breakpoint Tables For Interpretation Of mics And Zone Diameters, Version 10.0, 2020.” http://www.eucast.org/clinical_breakpoints/.

[35] Werkgroep Infectiepreventie (WIP) (2011). “Bijzonder Resistente Micro-organismen (brmo).”

[36] WHO Collaborating Center for Drug Statistics Methodology (2019). “ATC/DDD Index.” https://www.whocc.no/atc_ddd_index/. Accessed: 2019-4-9.

[37] WHO Collaborating Centre for Drug Statistics Methodology (2018). “Guidelines For atc Classification And ddd Assignment.”

[38] WHO Collaborating Centre for Surveillance of Antimicrobial Resistance (2019). “WHONET.” http://www.whonet.org/. Accessed: 2019-07-16.

[39] Wickham H (2014). “Tidy Data.” Journal of Statistical Software, 59(10). ISSN 1548-7660. doi:10.18637/jss.v059.i10. URL http://www.jstatsoft.org/v59/i10/.

[40] Wickham H (2016). ggplot2: Elegant Graphics For Data Analysis. Springer-Verlag New York. ISBN 978-3-319-24277-4. URL https://ggplot2.tidyverse.org.

[41] Wickham H, Averick M, Bryan J, Chang W, McGowan LD, François R, Grolemund G, Hayes A, Henry L, Hester J, Kuhn M, Pedersen TL, Miller E, Bache SM, Müller K, Ooms J, Robinson D, Seidel DP, Spinu V, Takahashi K, Vaughan D, Wilke C, Woo K, Yutani H (2019). “Welcome To The tidyverse.” Journal of Open Source Software, 4(43), 1686. doi:10.21105/joss.01686.

[42] Wickham H, François R, Henry L, Müller K (2020). dplyr: A Grammar of Data Manipulation. R package version 1.0.2, URL https://CRAN.R-project.org/package=dplyr.

[43] Winstanley T, Courvalin P (2011). “Expert Systems In Clinical Microbiology.” Clin. Microbiol. Rev., 24(3), 515–556. doi:10.1128/CMR.00061-10.

[44] World Health Organization (2014). “Companion Handbook To The who Guidelines For The Programmatic Management Of Drug-resistant Tuberculosis.”

[45] World Health Organization (2018). “Global Antimicrobial Resistance Surveillance System (GLASS) Report: Early Implementation 2017-2018.”

[46] World Health Organization (2018). “Global Tuberculosis Report 2018.”

