## Supplementary material for "AMR - An R Package for Working with Antimicrobial Resistance Data": Source package: AMR.pdf

January 5, 2021

**Version** 1.5.0

**Date** 2021-01-05

**Title** Antimicrobial Resistance Analysis

**Description** Functions to simplify the analysis and prediction of Antimicrobial Resistance (AMR) and to work with microbial and antimicrobial properties by using evidence-based methods, like those defined by Leclercq et al. (2013) <doi:10.1111/j.1469-0691.2011.03703.x> and the Clinical and Laboratory Standards Institute (2014) <isbn: 1-56238-899-1>.

**Depends** R (>= 3.0.0)

**Suggests** cleaner,
curl,
dplyr,
ggplot2,
knitr,
microbenchmark,
pillar,
readxl,
rmarkdown,
rstudioapi,
rvest,
skimr,
testthat,
tidyr,
xml2

**VignetteBuilder** knitr,rmarkdown

**URL** https://msberends.github.io/AMR/, https://github.com/msberends/AMR

**BugReports** https://github.com/msberends/AMR/issues

**License** GPL-2 | file LICENSE

**Encoding** UTF-8

**LazyData** true

**RoxygenNote** 7.1.1

**Roxygen** list(markdown = TRUE)

### R topics documented:

**Index** **121**

#### ab_from_text — *Retrieve antimicrobial drug names and doses from clinical text*

##### Description

Use this function on e.g. clinical texts from health care records. It returns a list with all antimicrobial drugs, doses and forms of administration found in the texts.

##### Usage

```
ab_from_text(
  text,
  type = c("drug", "dose", "administration"),
  collapse = NULL,
  translate_ab = FALSE,
  thorough_search = NULL,
  ...
)
```

##### Arguments

| | |
|---|---|
| `text` | text to analyse |
| `type` | type of property to search for, either `"drug"`, `"dose"` or `"administration"`, see *Examples* |
| `collapse` | character to pass on to `paste(...,collapse = ...)` to only return one character per element of `text`, see *Examples* |
| `translate_ab` | if `type = "drug"`: a column name of the antibiotics data set to translate the antibiotic abbreviations to, using `ab_property()`. Defaults to `FALSE`. Using `TRUE` is equal to using "name". |
| `thorough_search` | logical to indicate whether the input must be extensively searched for misspelling and other faulty input values. Setting this to `TRUE` will take considerably more time than when using `FALSE`. At default, it will turn `TRUE` when all input elements contain a maximum of three words. |
| `...` | arguments passed on to `as.ab()` |

##### Details

This function is also internally used by `as.ab()`, although it then only searches for the first drug name and will throw a note if more drug names could have been returned. Note: the `as.ab()` function may use very long regular expression to match brand names of antimicrobial agents. This may fail on some systems.

**Argument `type`:**

At default, the function will search for antimicrobial drug names. All text elements will be searched for official names, ATC codes and brand names. As it uses `as.ab()` internally, it will correct for misspelling.

With `type = "dose"` (or similar, like "dosing", "doses"), all text elements will be searched for numeric values that are higher than 100 and do not resemble years. The output will be numeric. It supports any unit (g, mg, IE, etc.) and multiple values in one clinical text, see *Examples*.

With `type = "administration"` (or abbreviations, like "admin", "adm"), all text elements will be searched for a form of drug administration. It supports the following forms (including common abbreviations): buccal, implant, inhalation, instillation, intravenous, nasal, oral, parenteral, rectal, sublingual, transdermal and vaginal. Abbreviations for oral (such as 'po', 'per os') will become "oral", all values for intravenous (such as 'iv', 'intraven') will become "iv". It supports multiple values in one clinical text, see *Examples*.

**Argument** `collapse`**:**

Without using `collapse`, this function will return a list. This can be convenient to use e.g. inside a `mutate()`):

`df %>% mutate(abx = ab_from_text(clinical_text))`

The returned AB codes can be transformed to official names, groups, etc. with all `ab_*` functions such as `ab_name()` and `ab_group()`, or by using the `translate_ab` argument.

With using `collapse`, this function will return a character:

`df %>% mutate(abx = ab_from_text(clinical_text, collapse = "|"))`

#### Value

A list, or a character if `collapse` is not `NULL`

#### Maturing lifecycle

The lifecycle of this function is **maturing**. The unlying code of a maturing function has been roughed out, but finer details might still change. Since this function needs wider usage and more extensive testing, you are very welcome to suggest changes at our repository or write us an email (see section 'Contact Us').

#### Read more on our website!

On our website `https://msberends.github.io/AMR/` you can find a comprehensive tutorial about how to conduct AMR analysis, the complete documentation of all functions and an example analysis using WHONET data. As we would like to better understand the backgrounds and needs of our users, please participate in our survey!

#### Examples

```
# mind the bad spelling of amoxicillin in this line,
# straight from a true health care record:
ab_from_text("28/03/2020 regular amoxicilliin 500mg po tds")

ab_from_text("500 mg amoxi po and 400mg cipro iv")
ab_from_text("500 mg amoxi po and 400mg cipro iv", type = "dose")
ab_from_text("500 mg amoxi po and 400mg cipro iv", type = "admin")

ab_from_text("500 mg amoxi po and 400mg cipro iv", collapse = ", ")

# if you want to know which antibiotic groups were administered, do e.g.:
abx <- ab_from_text("500 mg amoxi po and 400mg cipro iv")
ab_group(abx[[1]])

if (require("dplyr")) {
```

```
  tibble(clinical_text = c("given 400mg cipro and 500 mg amox",
                           "started on doxy iv today")) %>%
    mutate(abx_codes = ab_from_text(clinical_text),
           abx_doses = ab_from_text(clinical_text, type = "doses"),
           abx_admin = ab_from_text(clinical_text, type = "admin"),
           abx_coll = ab_from_text(clinical_text, collapse = "|"),
           abx_coll_names = ab_from_text(clinical_text,
                                         collapse = "|",
                                         translate_ab = "name"),
           abx_coll_doses = ab_from_text(clinical_text,
                                         type = "doses",
                                         collapse = "|"),
           abx_coll_admin = ab_from_text(clinical_text,
                                         type = "admin",
                                         collapse = "|"))

}
```

---

`ab_property` *Get properties of an antibiotic*

---

#### Description

Use these functions to return a specific property of an antibiotic from the antibiotics data set. All input values will be evaluated internally with `as.ab()`.

#### Usage

```
ab_name(x, language = get_locale(), tolower = FALSE, ...)

ab_atc(x, ...)

ab_cid(x, ...)

ab_synonyms(x, ...)

ab_tradenames(x, ...)

ab_group(x, language = get_locale(), ...)

ab_atc_group1(x, language = get_locale(), ...)

ab_atc_group2(x, language = get_locale(), ...)

ab_loinc(x, ...)

ab_ddd(x, administration = "oral", units = FALSE, ...)

ab_info(x, language = get_locale(), ...)

ab_url(x, open = FALSE, ...)

ab_property(x, property = "name", language = get_locale(), ...)
```

#### Arguments

| | |
|---|---|
| x | any (vector of) text that can be coerced to a valid antibiotic code with `as.ab()` |
| language | language of the returned text, defaults to system language (see `get_locale()`) and can also be set with `getOption("AMR_locale")`. Use `language = NULL` or `language = ""` to prevent translation. |
| tolower | logical to indicate whether the first character of every output should be transformed to a lower case character. This will lead to e.g. "polymyxin B" and not "polymyxin b". |
| ... | other arguments passed on to `as.ab()` |
| administration | way of administration, either `"oral"` or `"iv"` |
| units | a logical to indicate whether the units instead of the DDDs itself must be returned, see Examples |
| open | browse the URL using `utils::browseURL()` |
| property | one of the column names of one of the antibiotics data set |

#### Details

All output will be translated where possible.

The function `ab_url()` will return the direct URL to the official WHO website. A warning will be returned if the required ATC code is not available.

#### Value

- An integer in case of `ab_cid()`
- A named list in case of `ab_info()` and multiple `ab_synonyms()`/`ab_tradenames()`
- A double in case of `ab_ddd()`
- A character in all other cases

#### Stable lifecycle

The lifecycle of this function is **stable**. In a stable function, major changes are unlikely. This means that the unlying code will generally evolve by adding new arguments; removing arguments or changing the meaning of existing arguments will be avoided.

If the unlying code needs breaking changes, they will occur gradually. For example, a argument will be deprecated and first continue to work, but will emit an message informing you of the change. Next, typically after at least one newly released version on CRAN, the message will be transformed to an error.

#### Source

World Health Organization (WHO) Collaborating Centre for Drug Statistics Methodology: https://www.whocc.no/atc_ddd_index/

WHONET 2019 software: http://www.whonet.org/software.html

European Commission Public Health PHARMACEUTICALS - COMMUNITY REGISTER: http://ec.europa.eu/health/documents/community-register/html/atc.htm

##### Reference data publicly available

All reference data sets (about microorganisms, antibiotics, R/SI interpretation, EUCAST rules, etc.) in this AMR package are publicly and freely available. We continually export our data sets to formats for use in R, SPSS, SAS, Stata and Excel. We also supply flat files that are machine-readable and suitable for input in any software program, such as laboratory information systems. Please find all download links on our website, which is automatically updated with every code change.

##### Read more on our website!

On our website https://msberends.github.io/AMR/ you can find a comprehensive tutorial about how to conduct AMR analysis, the complete documentation of all functions and an example analysis using WHONET data. As we would like to better understand the backgrounds and needs of our users, please participate in our survey!

##### See Also

antibiotics

##### Examples

```
# all properties:
ab_name("AMX")       # "Amoxicillin"
ab_atc("AMX")        # J01CA04 (ATC code from the WHO)
ab_cid("AMX")        # 33613 (Compound ID from PubChem)
ab_synonyms("AMX")   # a list with brand names of amoxicillin
ab_tradenames("AMX") # same
ab_group("AMX")      # "Beta-lactams/penicillins"
ab_atc_group1("AMX") # "Beta-lactam antibacterials, penicillins"
ab_atc_group2("AMX") # "Penicillins with extended spectrum"
ab_url("AMX")        # link to the official WHO page

# smart lowercase tranformation
ab_name(x = c("AMC", "PLB"))  # "Amoxicillin/clavulanic acid" "Polymyxin B"
ab_name(x = c("AMC", "PLB"),
        tolower = TRUE)       # "amoxicillin/clavulanic acid" "polymyxin B"

# defined daily doses (DDD)
ab_ddd("AMX", "oral")               #  1
ab_ddd("AMX", "oral", units = TRUE) # "g"
ab_ddd("AMX", "iv")                 #  1
ab_ddd("AMX", "iv", units = TRUE)   # "g"

ab_info("AMX")       # all properties as a list

# all ab_* functions use as.ab() internally, so you can go from 'any' to 'any':
ab_atc("AMP")           # ATC code of AMP (ampicillin)
ab_group("J01CA01")     # Drug group of ampicillins ATC code
ab_loinc("ampicillin")  # LOINC codes of ampicillin
ab_name("21066-6")      # "Ampicillin" (using LOINC)
ab_name(6249)           # "Ampicillin" (using CID)
ab_name("J01CA01")      # "Ampicillin" (using ATC)

# spelling from different languages and dyslexia are no problem
ab_atc("ceftriaxon")
ab_atc("cephtriaxone")
ab_atc("cephthriaxone")
```

```
ab_atc("seephthriaaksone")
```

---

`age` *Age in years of individuals*

---

#### Description

Calculates age in years based on a reference date, which is the sytem date at default.

#### Usage

```
age(x, reference = Sys.Date(), exact = FALSE, na.rm = FALSE, ...)
```

#### Arguments

| | |
|---|---|
| `x` | date(s), will be coerced with `as.POSIXlt()` |
| `reference` | reference date(s) (defaults to today), will be coerced with `as.POSIXlt()` |
| `exact` | a logical to indicate whether age calculation should be exact, i.e. with decimals. It divides the number of days of year-to-date (YTD) of `x` by the number of days in the year of `reference` (either 365 or 366). |
| `na.rm` | a logical to indicate whether missing values should be removed |
| `...` | arguments passed on to `as.POSIXlt()`, such as `origin` |

#### Details

Ages below 0 will be returned as `NA` with a warning. Ages above 120 will only give a warning.

#### Value

An integer (no decimals) if `exact = FALSE`, a double (with decimals) otherwise

#### Stable lifecycle

The lifecycle of this function is **stable**. In a stable function, major changes are unlikely. This means that the unlying code will generally evolve by adding new arguments; removing arguments or changing the meaning of existing arguments will be avoided.

If the unlying code needs breaking changes, they will occur gradually. For example, a argument will be deprecated and first continue to work, but will emit an message informing you of the change. Next, typically after at least one newly released version on CRAN, the message will be transformed to an error.

#### Read more on our website!

On our website `https://msberends.github.io/AMR/` you can find a comprehensive tutorial about how to conduct AMR analysis, the complete documentation of all functions and an example analysis using WHONET data. As we would like to better understand the backgrounds and needs of our users, please participate in our survey!

#### See Also

To split ages into groups, use the `age_groups()` function.

##### Examples

```
# 10 random birth dates
df <- data.frame(birth_date = Sys.Date() - runif(10) * 25000)
# add ages
df$age <- age(df$birth_date)
# add exact ages
df$age_exact <- age(df$birth_date, exact = TRUE)

df
```

---

#### age_groups — *Split ages into age groups*

---

##### Description

Split ages into age groups defined by the `split` argument. This allows for easier demographic (antimicrobial resistance) analysis.

##### Usage

```
age_groups(x, split_at = c(12, 25, 55, 75), na.rm = FALSE)
```

##### Arguments

| | |
|---|---|
| `x` | age, e.g. calculated with `age()` |
| `split_at` | values to split `x` at, defaults to age groups 0-11, 12-24, 25-54, 55-74 and 75+. See Details. |
| `na.rm` | a logical to indicate whether missing values should be removed |

##### Details

To split ages, the input for the `split_at` argument can be:

- A numeric vector. A value of e.g. `c(10,20)` will split `x` on 0-9, 10-19 and 20+. A value of only `50` will split `x` on 0-49 and 50+. The default is to split on young children (0-11), youth (12-24), young adults (25-54), middle-aged adults (55-74) and elderly (75+).
- A character:
  - `"children"` or `"kids"`, equivalent of: `c(0,1,2,4,6,13,18)`. This will split on 0, 1, 2-3, 4-5, 6-12, 13-17 and 18+.
  - `"elderly"` or `"seniors"`, equivalent of: `c(65,75,85)`. This will split on 0-64, 65-74, 75-84, 85+.
  - `"fives"`, equivalent of: `1:20 * 5`. This will split on 0-4, 5-9, ..., 95-99, 100+.
  - `"tens"`, equivalent of: `1:10 * 10`. This will split on 0-9, 10-19, ..., 90-99, 100+.

##### Value

Ordered factor

##### Stable lifecycle

The lifecycle of this function is **stable**. In a stable function, major changes are unlikely. This means that the unlying code will generally evolve by adding new arguments; removing arguments or changing the meaning of existing arguments will be avoided.

If the unlying code needs breaking changes, they will occur gradually. For example, a argument will be deprecated and first continue to work, but will emit an message informing you of the change. Next, typically after at least one newly released version on CRAN, the message will be transformed to an error.

##### Read more on our website!

On our website `https://msberends.github.io/AMR/` you can find a comprehensive tutorial about how to conduct AMR analysis, the complete documentation of all functions and an example analysis using WHONET data. As we would like to better understand the backgrounds and needs of our users, please participate in our survey!

##### See Also

To determine ages, based on one or more reference dates, use the `age()` function.

##### Examples

```
ages <- c(3, 8, 16, 54, 31, 76, 101, 43, 21)

# split into 0-49 and 50+
age_groups(ages, 50)

# split into 0-19, 20-49 and 50+
age_groups(ages, c(20, 50))

# split into groups of ten years
age_groups(ages, 1:10 * 10)
age_groups(ages, split_at = "tens")

# split into groups of five years
age_groups(ages, 1:20 * 5)
age_groups(ages, split_at = "fives")

# split specifically for children
age_groups(ages, c(1, 2, 4, 6, 13, 17))
age_groups(ages, "children")

# resistance of ciprofloxacin per age group
if (require("dplyr")) {
  example_isolates %>%
    filter_first_isolate() %>%
    filter(mo == as.mo("E. coli")) %>%
    group_by(age_group = age_groups(age)) %>%
    select(age_group, CIP) %>%
    ggplot_rsi(x = "age_group", minimum = 0)
}
```

AMR *The* AMR *Package*

#### Description

Welcome to the AMR package.

#### Details

AMR is a free, open-source and independent R package to simplify the analysis and prediction of Antimicrobial Resistance (AMR) and to work with microbial and antimicrobial data and properties, by using evidence-based methods. Our aim is to provide a standard for clean and reproducible antimicrobial resistance data analysis, that can therefore empower epidemiological analyses to continuously enable surveillance and treatment evaluation in any setting.

After installing this package, R knows ~70,000 distinct microbial species and all ~550 antibiotic, antimycotic and antiviral drugs by name and code (including ATC, EARS-NET, LOINC and SNOMED CT), and knows all about valid R/SI and MIC values. It supports any data format, including WHONET/EARS-Net data.

This package is fully independent of any other R package and works on Windows, macOS and Linux with all versions of R since R-3.0.0 (April 2013). It was designed to work in any setting, including those with very limited resources. It was created for both routine data analysis and academic research at the Faculty of Medical Sciences of the University of Groningen, in collaboration with non-profit organisations Certe Medical Diagnostics and Advice and University Medical Center Groningen. This R package is actively maintained and free software; you can freely use and distribute it for both personal and commercial (but not patent) purposes under the terms of the GNU General Public License version 2.0 (GPL-2), as published by the Free Software Foundation.

This package can be used for:

- Reference for the taxonomy of microorganisms, since the package contains all microbial (sub)species from the Catalogue of Life and List of Prokaryotic names with Standing in Nomenclature
- Interpreting raw MIC and disk diffusion values, based on the latest CLSI or EUCAST guidelines
- Retrieving antimicrobial drug names, doses and forms of administration from clinical health care records
- Determining first isolates to be used for AMR analysis
- Calculating antimicrobial resistance
- Determining multi-drug resistance (MDR) / multi-drug resistant organisms (MDRO)
- Calculating (empirical) susceptibility of both mono therapy and combination therapies
- Predicting future antimicrobial resistance using regression models
- Getting properties for any microorganism (such as Gram stain, species, genus or family)
- Getting properties for any antibiotic (such as name, code of EARS-Net/ATC/LOINC/PubChem, defined daily dose or trade name)
- Plotting antimicrobial resistance
- Applying EUCAST expert rules
- Getting SNOMED codes of a microorganism, or getting properties of a microorganism based on a SNOMED code

- Getting LOINC codes of an antibiotic, or getting properties of an antibiotic based on a LOINC code
- Machine reading the EUCAST and CLSI guidelines from 2011-2020 to translate MIC values and disk diffusion diameters to R/SI
- Principal component analysis for AMR

##### Reference data publicly available

All reference data sets (about microorganisms, antibiotics, R/SI interpretation, EUCAST rules, etc.) in this `AMR` package are publicly and freely available. We continually export our data sets to formats for use in R, SPSS, SAS, Stata and Excel. We also supply flat files that are machine-readable and suitable for input in any software program, such as laboratory information systems. Please find all download links on our website, which is automatically updated with every code change.

##### Read more on our website!

On our website `https://msberends.github.io/AMR/` you can find a comprehensive tutorial about how to conduct AMR analysis, the complete documentation of all functions and an example analysis using WHONET data. As we would like to better understand the backgrounds and needs of our users, please participate in our survey!

##### Contact Us

For suggestions, comments or questions, please contact us at:

Matthijs S. Berends
m.s.berends [at] umcg [dot] nl
University of Groningen Department of Medical Microbiology University Medical Center Groningen
Post Office Box 30001
9700 RB Groningen
The Netherlands `https://msberends.github.io/AMR/`

If you have found a bug, please file a new issue at:
`https://github.com/msberends/AMR/issues`

---

`antibiotics` *Data sets with 557 antimicrobials*

---

##### Description

Two data sets containing all antibiotics/antimycotics and antivirals. Use `as.ab()` or one of the `ab_*` functions to retrieve values from the antibiotics data set. Three identifiers are included in this data set: an antibiotic ID (`ab`, primarily used in this package) as defined by WHONET/EARS-Net, an ATC code (`atc`) as defined by the WHO, and a Compound ID (`cid`) as found in PubChem. Other properties in this data set are derived from one or more of these codes.

##### Usage

```
antibiotics

antivirals
```

#### Format

##### For the antibiotics data set: a data.frame with 455 observations and 14 variables::

- ab
  Antibiotic ID as used in this package (such as AMC), using the official EARS-Net (European Antimicrobial Resistance Surveillance Network) codes where available
- atc
  ATC code (Anatomical Therapeutic Chemical) as defined by the WHOCC, like J01CR02
- cid
  Compound ID as found in PubChem
- name
  Official name as used by WHONET/EARS-Net or the WHO
- group
  A short and concise group name, based on WHONET and WHOCC definitions
- atc_group1
  Official pharmacological subgroup (3rd level ATC code) as defined by the WHOCC, like "Macrolides, lincosamides and streptogramins"
- atc_group2
  Official chemical subgroup (4th level ATC code) as defined by the WHOCC, like "Macrolides"
- abbr
  List of abbreviations as used in many countries, also for antibiotic susceptibility testing (AST)
- synonyms
  Synonyms (often trade names) of a drug, as found in PubChem based on their compound ID
- oral_ddd
  Defined Daily Dose (DDD), oral treatment
- oral_units
  Units of oral_ddd
- iv_ddd
  Defined Daily Dose (DDD), parenteral treatment
- iv_units
  Units of iv_ddd
- loinc
  All LOINC codes (Logical Observation Identifiers Names and Codes) associated with the name of the antimicrobial agent. Use ab_loinc() to retrieve them quickly, see ab_property().

##### For the antivirals data set: a data.frame with 102 observations and 9 variables::

- atc
  ATC code (Anatomical Therapeutic Chemical) as defined by the WHOCC
- cid
  Compound ID as found in PubChem
- name
  Official name as used by WHONET/EARS-Net or the WHO
- atc_group
  Official pharmacological subgroup (3rd level ATC code) as defined by the WHOCC
- synonyms
  Synonyms (often trade names) of a drug, as found in PubChem based on their compound ID
- oral_ddd
  Defined Daily Dose (DDD), oral treatment
- oral_units
  Units of oral_ddd

- `iv_ddd`
  Defined Daily Dose (DDD), parenteral treatment
- `iv_units`
  Units of `iv_ddd`

An object of class `data.frame` with 102 rows and 9 columns.

#### Details

Properties that are based on an ATC code are only available when an ATC is available. These properties are: `atc_group1`, `atc_group2`, `oral_ddd`, `oral_units`, `iv_ddd` and `iv_units`.

Synonyms (i.e. trade names) are derived from the Compound ID (`cid`) and consequently only available where a CID is available.

**Direct download:**

These data sets are available as 'flat files' for use even without R - you can find the files here:

- https://github.com/msberends/AMR/raw/master/data-raw/antibiotics.txt
- https://github.com/msberends/AMR/raw/master/data-raw/antivirals.txt

Files in R format (with preserved data structure) can be found here:

- https://github.com/msberends/AMR/raw/master/data/antibiotics.rda
- https://github.com/msberends/AMR/raw/master/data/antivirals.rda

#### Reference data publicly available

All reference data sets (about microorganisms, antibiotics, R/SI interpretation, EUCAST rules, etc.) in this `AMR` package are publicly and freely available. We continually export our data sets to formats for use in R, SPSS, SAS, Stata and Excel. We also supply flat files that are machine-readable and suitable for input in any software program, such as laboratory information systems. Please find all download links on our website, which is automatically updated with every code change.

#### WHOCC

This package contains **all ~550 antibiotic, antimycotic and antiviral drugs** and their Anatomical Therapeutic Chemical (ATC) codes, ATC groups and Defined Daily Dose (DDD) from the World Health Organization Collaborating Centre for Drug Statistics Methodology (WHOCC, https://www.whocc.no) and the Pharmaceuticals Community Register of the European Commission (http://ec.europa.eu/health/documents/community-register/html/atc.htm).

These have become the gold standard for international drug utilisation monitoring and research.

The WHOCC is located in Oslo at the Norwegian Institute of Public Health and funded by the Norwegian government. The European Commission is the executive of the European Union and promotes its general interest.

**NOTE: The WHOCC copyright does not allow use for commercial purposes, unlike any other info from this package.** See https://www.whocc.no/copyright_disclaimer/.

#### Read more on our website!

On our website https://msberends.github.io/AMR/ you can find a comprehensive tutorial about how to conduct AMR analysis, the complete documentation of all functions and an example analysis using WHONET data. As we would like to better understand the backgrounds and needs of our users, please participate in our survey!

#### Source

World Health Organization (WHO) Collaborating Centre for Drug Statistics Methodology (WHOCC): https://www.whocc.no/atc_ddd_index/

WHONET 2019 software: http://www.whonet.org/software.html

European Commission Public Health PHARMACEUTICALS - COMMUNITY REGISTER: http://ec.europa.eu/health/documents/community-register/html/atc.htm

#### See Also

microorganisms, intrinsic_resistant

---

### antibiotic_class_selectors *Antibiotic class selectors*

---

#### Description

These functions help to select the columns of antibiotics that are of a specific antibiotic class, without the need to define the columns or antibiotic abbreviations.

#### Usage

```
ab_class(ab_class)

aminoglycosides()

carbapenems()

cephalosporins()

cephalosporins_1st()

cephalosporins_2nd()

cephalosporins_3rd()

cephalosporins_4th()

cephalosporins_5th()

fluoroquinolones()

glycopeptides()

macrolides()

penicillins()

tetracyclines()
```

#### Arguments

ab_class — an antimicrobial class, like "carbapenems". The columns group, atc_group1 and atc_group2 of the antibiotics data set will be searched (case-insensitive) for this value.

#### Details

All columns will be searched for known antibiotic names, abbreviations, brand names and codes (ATC, EARS-Net, WHO, etc.) in the antibiotics data set. This means that a selector like e.g. aminoglycosides() will pick up column names like 'gen', 'genta', 'J01GB03', 'tobra', 'Tobracin', etc.

#### Reference data publicly available

All reference data sets (about microorganisms, antibiotics, R/SI interpretation, EUCAST rules, etc.) in this AMR package are publicly and freely available. We continually export our data sets to formats for use in R, SPSS, SAS, Stata and Excel. We also supply flat files that are machine-readable and suitable for input in any software program, such as laboratory information systems. Please find all download links on our website, which is automatically updated with every code change.

#### Read more on our website!

On our website https://msberends.github.io/AMR/ you can find a comprehensive tutorial about how to conduct AMR analysis, the complete documentation of all functions and an example analysis using WHONET data. As we would like to better understand the backgrounds and needs of our users, please participate in our survey!

#### See Also

filter_ab_class() for the filter() equivalent.

#### Examples

```
# `example_isolates` is a dataset available in the AMR package.
# See ?example_isolates.

# this will select columns 'IPM' (imipenem) and 'MEM' (meropenem):
example_isolates[, c(carbapenems())]
# this will select columns 'mo', 'AMK', 'GEN', 'KAN' and 'TOB':
example_isolates[, c("mo", aminoglycosides())]

if (require("dplyr")) {

  # this will select columns 'IPM' (imipenem) and 'MEM' (meropenem):
  example_isolates %>%
    select(carbapenems())

  # this will select columns 'mo', 'AMK', 'GEN', 'KAN' and 'TOB':
  example_isolates %>%
    select(mo, aminoglycosides())

  # this will select columns 'mo' and all antimycobacterial drugs ('RIF'):
  example_isolates %>%
    select(mo, ab_class("mycobact"))
```

```
  # get bug/drug combinations for only macrolides in Gram-positives:
  example_isolates %>%
    filter(mo_is_gram_positive()) %>%
    select(mo, macrolides()) %>%
    bug_drug_combinations() %>%
    format()

  data.frame(some_column = "some_value",
             J01CA01 = "S") %>%   # ATC code of ampicillin
    select(penicillins())         # only the 'J01CA01' column will be selected

  # with dplyr 1.0.0 and higher (that adds 'across()'), this is equal:
  # (though the row names on the first are more correct)
  example_isolates %>% filter_carbapenems("R", "all")
  example_isolates %>% filter(across(carbapenems(), ~. == "R"))
}
```

---

`as.ab` *Transform input to an antibiotic ID*

---

#### Description

Use this function to determine the antibiotic code of one or more antibiotics. The data set antibiotics will be searched for abbreviations, official names and synonyms (brand names).

#### Usage

```
as.ab(x, flag_multiple_results = TRUE, info = TRUE, ...)

is.ab(x)
```

#### Arguments

| | |
|---|---|
| `x` | character vector to determine to antibiotic ID |
| `flag_multiple_results` | logical to indicate whether a note should be printed to the console that probably more than one antibiotic code or name can be retrieved from a single input value. |
| `info` | logical to indicate whether a progress bar should be printed |
| `...` | arguments passed on to internal functions |

#### Details

All entries in the antibiotics data set have three different identifiers: a human readable EARS-Net code (column `ab`, used by ECDC and WHONET), an ATC code (column `atc`, used by WHO), and a CID code (column `cid`, Compound ID, used by PubChem). The data set contains more than 5,000 official brand names from many different countries, as found in PubChem.

All these properties will be searched for the user input. The `as.ab()` can correct for different forms of misspelling:

- Wrong spelling of drug names (such as "tobramicin" or "gentamycin"), which corrects for most audible similarities such as f/ph, x/ks, c/z/s, t/th, etc.
- Too few or too many vowels or consonants
- Switching two characters (such as "mreopenem", often the case in clinical data, when doctors typed too fast)
- Digitalised paper records, leaving artefacts like 0/o/O (zero and O's), B/8, n/r, etc.

Use the `ab_*` functions to get properties based on the returned antibiotic ID, see Examples.

Note: the `as.ab()` and `ab_*` functions may use very long regular expression to match brand names of antimicrobial agents. This may fail on some systems.

#### Value

A character vector with additional class `ab`

#### Source

World Health Organization (WHO) Collaborating Centre for Drug Statistics Methodology: `https://www.whocc.no/atc_ddd_index/`

WHONET 2019 software: `http://www.whonet.org/software.html`

European Commission Public Health PHARMACEUTICALS - COMMUNITY REGISTER: `http://ec.europa.eu/health/documents/community-register/html/atc.htm`

#### Stable lifecycle

The lifecycle of this function is **stable**. In a stable function, major changes are unlikely. This means that the unlying code will generally evolve by adding new arguments; removing arguments or changing the meaning of existing arguments will be avoided.

If the unlying code needs breaking changes, they will occur gradually. For example, a argument will be deprecated and first continue to work, but will emit an message informing you of the change. Next, typically after at least one newly released version on CRAN, the message will be transformed to an error.

#### WHOCC

This package contains **all ~550 antibiotic, antimycotic and antiviral drugs** and their Anatomical Therapeutic Chemical (ATC) codes, ATC groups and Defined Daily Dose (DDD) from the World Health Organization Collaborating Centre for Drug Statistics Methodology (WHOCC, `https://www.whocc.no`) and the Pharmaceuticals Community Register of the European Commission (`http://ec.europa.eu/health/documents/community-register/html/atc.htm`).

These have become the gold standard for international drug utilisation monitoring and research.

The WHOCC is located in Oslo at the Norwegian Institute of Public Health and funded by the Norwegian government. The European Commission is the executive of the European Union and promotes its general interest.

**NOTE: The WHOCC copyright does not allow use for commercial purposes, unlike any other info from this package.** See `https://www.whocc.no/copyright_disclaimer/`.

##### Reference data publicly available

All reference data sets (about microorganisms, antibiotics, R/SI interpretation, EUCAST rules, etc.) in this AMR package are publicly and freely available. We continually export our data sets to formats for use in R, SPSS, SAS, Stata and Excel. We also supply flat files that are machine-readable and suitable for input in any software program, such as laboratory information systems. Please find all download links on our website, which is automatically updated with every code change.

##### Read more on our website!

On our website https://msberends.github.io/AMR/ you can find a comprehensive tutorial about how to conduct AMR analysis, the complete documentation of all functions and an example analysis using WHONET data. As we would like to better understand the backgrounds and needs of our users, please participate in our survey!

##### See Also

- antibiotics for the data.frame that is being used to determine ATCs
- `ab_from_text()` for a function to retrieve antimicrobial drugs from clinical text (from health care records)

##### Examples

```
# these examples all return "ERY", the ID of erythromycin:
as.ab("J01FA01")
as.ab("J 01 FA 01")
as.ab("Erythromycin")
as.ab("eryt")
as.ab("   eryt 123")
as.ab("ERYT")
as.ab("ERY")
as.ab("eritromicine") # spelled wrong, yet works
as.ab("Erythrocin")   # trade name
as.ab("Romycin")      # trade name

# spelling from different languages and dyslexia are no problem
ab_atc("ceftriaxon")
ab_atc("cephtriaxone")     # small spelling error
ab_atc("cephthriaxone")    # or a bit more severe
ab_atc("seephthriaaksone") # and even this works

# use ab_* functions to get a specific properties (see ?ab_property);
# they use as.ab() internally:
ab_name("J01FA01")    # "Erythromycin"
ab_name("eryt")       # "Erythromycin"
```

---

#### as.disk — *Transform input to disk diffusion diameters*

---

##### Description

This transforms a vector to a new class `disk`, which is a disk diffusion growth zone size (around an antibiotic disk) in millimetres between 6 and 50.

##### Usage

```
as.disk(x, na.rm = FALSE)

is.disk(x)
```

##### Arguments

| | |
|---|---|
| x | vector |
| na.rm | a logical indicating whether missing values should be removed |

##### Details

Interpret disk values as RSI values with `as.rsi()`. It supports guidelines from EUCAST and CLSI.

##### Value

An integer with additional class `disk`

##### Stable lifecycle

The lifecycle of this function is **stable**. In a stable function, major changes are unlikely. This means that the unlying code will generally evolve by adding new arguments; removing arguments or changing the meaning of existing arguments will be avoided.

If the unlying code needs breaking changes, they will occur gradually. For example, a argument will be deprecated and first continue to work, but will emit an message informing you of the change. Next, typically after at least one newly released version on CRAN, the message will be transformed to an error.

##### Read more on our website!

On our website `https://msberends.github.io/AMR/` you can find a comprehensive tutorial about how to conduct AMR analysis, the complete documentation of all functions and an example analysis using WHONET data. As we would like to better understand the backgrounds and needs of our users, please participate in our survey!

##### See Also

`as.rsi()`

##### Examples

```
# transform existing disk zones to the `disk` class
df <- data.frame(microorganism = "E. coli",
                 AMP = 20,
                 CIP = 14,
                 GEN = 18,
                 TOB = 16)
df[, 2:5] <- lapply(df[, 2:5], as.disk)
# same with dplyr:
# df %>% mutate(across(AMP:TOB, as.disk))

# interpret disk values, see ?as.rsi
as.rsi(x = as.disk(18),
```

```
        mo = "Strep pneu",  # `mo` will be coerced with as.mo()
        ab = "ampicillin",  # and `ab` with as.ab()
        guideline = "EUCAST")

as.rsi(df)
```

`as.mic` *Transform input to minimum inhibitory concentrations (MIC)*

#### Description

This transforms a vector to a new class `mic`, which is an ordered factor with valid minimum inhibitory concentrations (MIC) as levels. Invalid MIC values will be translated as `NA` with a warning.

#### Usage

```
as.mic(x, na.rm = FALSE)

is.mic(x)
```

#### Arguments

| | |
|---|---|
| `x` | vector |
| `na.rm` | a logical indicating whether missing values should be removed |

#### Details

To interpret MIC values as RSI values, use `as.rsi()` on MIC values. It supports guidelines from EUCAST and CLSI.

#### Value

Ordered factor with additional class `mic`

#### Stable lifecycle

The lifecycle of this function is **stable**. In a stable function, major changes are unlikely. This means that the unlying code will generally evolve by adding new arguments; removing arguments or changing the meaning of existing arguments will be avoided.

If the unlying code needs breaking changes, they will occur gradually. For example, a argument will be deprecated and first continue to work, but will emit an message informing you of the change. Next, typically after at least one newly released version on CRAN, the message will be transformed to an error.

#### Read more on our website!

On our website `https://msberends.github.io/AMR/` you can find a comprehensive tutorial about how to conduct AMR analysis, the complete documentation of all functions and an example analysis using WHONET data. As we would like to better understand the backgrounds and needs of our users, please participate in our survey!

#### See Also

as.rsi()

#### Examples

```
mic_data <- as.mic(c(">=32", "1.0", "1", "1.00", 8, "<=0.128", "8", "16", "16"))
is.mic(mic_data)

# this can also coerce combined MIC/RSI values:
as.mic("<=0.002; S") # will return <=0.002

# interpret MIC values
as.rsi(x = as.mic(2),
       mo = as.mo("S. pneumoniae"),
       ab = "AMX",
       guideline = "EUCAST")
as.rsi(x = as.mic(4),
       mo = as.mo("S. pneumoniae"),
       ab = "AMX",
       guideline = "EUCAST")

plot(mic_data)
barplot(mic_data)
```

---

as.mo *Transform input to a microorganism ID*

---

#### Description

Use this function to determine a valid microorganism ID (mo). Determination is done using intelligent rules and the complete taxonomic kingdoms Bacteria, Chromista, Protozoa, Archaea and most microbial species from the kingdom Fungi (see Source). The input can be almost anything: a full name (like "Staphylococcus aureus"), an abbreviated name (such as "S. aureus"), an abbreviation known in the field (such as "MRSA"), or just a genus. Please see *Examples*.

#### Usage

```
as.mo(
  x,
  Becker = FALSE,
  Lancefield = FALSE,
  allow_uncertain = TRUE,
  reference_df = get_mo_source(),
  ignore_pattern = getOption("AMR_ignore_pattern"),
  language = get_locale(),
  ...
)

is.mo(x)

mo_failures()
```

```
mo_uncertainties()

mo_renamed()
```

#### Arguments

`x` a character vector or a data.frame with one or two columns

`Becker` a logical to indicate whether staphylococci should be categorised into coagulase-negative staphylococci ("CoNS") and coagulase-positive staphylococci ("CoPS") instead of their own species, according to Karsten Becker *et al.* (1,2,3).

This excludes *Staphylococcus aureus* at default, use `Becker = "all"` to also categorise *S. aureus* as "CoPS".

`Lancefield` a logical to indicate whether beta-haemolytic *Streptococci* should be categorised into Lancefield groups instead of their own species, according to Rebecca C. Lancefield (4). These *Streptococci* will be categorised in their first group, e.g. *Streptococcus dysgalactiae* will be group C, although officially it was also categorised into groups G and L.

This excludes *Enterococci* at default (who are in group D), use `Lancefield = "all"` to also categorise all *Enterococci* as group D.

`allow_uncertain` a number between `0` (or `"none"`) and `3` (or `"all"`), or `TRUE` (= `2`) or `FALSE` (= `0`) to indicate whether the input should be checked for less probable results, please see *Details*

`reference_df` a data.frame to be used for extra reference when translating `x` to a valid `mo`. See `set_mo_source()` and `get_mo_source()` to automate the usage of your own codes (e.g. used in your analysis or organisation).

`ignore_pattern` a regular expression (case-insensitive) of which all matches in `x` must return `NA`. This can be convenient to exclude known non-relevant input and can also be set with the option `AMR_ignore_pattern`, e.g. `options(AMR_ignore_pattern = "(not reported|contaminated flora)")`.

`language` language to translate text like "no growth", which defaults to the system language (see `get_locale()`)

`...` other arguments passed on to functions

#### Details

**General info:**

A microorganism ID from this package (class: `mo`) is human readable and typically looks like these examples:

```
Code               Full name
---------------    --------------------------------------
B_KLBSL            Klebsiella
B_KLBSL_PNMN       Klebsiella pneumoniae
B_KLBSL_PNMN_RHNS  Klebsiella pneumoniae rhinoscleromatis
|   |    |    |
|   |    |    |
|   |    |    \---> subspecies, a 4-5 letter acronym
|   |    \----> species, a 4-5 letter acronym
|   \----> genus, a 5-7 letter acronym
\----> taxonomic kingdom: A (Archaea), AN (Animalia), B (Bacteria),
                          C (Chromista), F (Fungi), P (Protozoa)
```

Values that cannot be coerced will be considered 'unknown' and will get the MO code `UNKNOWN`.

Use the `mo_*` functions to get properties based on the returned code, see Examples.

The algorithm uses data from the Catalogue of Life (see below) and from one other source (see microorganisms).

The `as.mo()` function uses several coercion rules for fast and logical results. It assesses the input matching criteria in the following order:

1. Human pathogenic prevalence: the function starts with more prevalent microorganisms, followed by less prevalent ones;
2. Taxonomic kingdom: the function starts with determining Bacteria, then Fungi, then Protozoa, then others;
3. Breakdown of input values to identify possible matches.

This will lead to the effect that e.g. `"E. coli"` (a microorganism highly prevalent in humans) will return the microbial ID of *Escherichia coli* and not *Entamoeba coli* (a microorganism less prevalent in humans), although the latter would alphabetically come first.

**Coping with uncertain results:**

In addition, the `as.mo()` function can differentiate four levels of uncertainty to guess valid results:

- Uncertainty level 0: no additional rules are applied;
- Uncertainty level 1: allow previously accepted (but now invalid) taxonomic names and minor spelling errors;
- Uncertainty level 2: allow all of level 1, strip values between brackets, inverse the words of the input, strip off text elements from the end keeping at least two elements;
- Uncertainty level 3: allow all of level 1 and 2, strip off text elements from the end, allow any part of a taxonomic name.

The level of uncertainty can be set using the argument `allow_uncertain`. The default is `allow_uncertain = TRUE`, which is equal to uncertainty level 2. Using `allow_uncertain = FALSE` is equal to uncertainty level 0 and will skip all rules. You can also use e.g. `as.mo(..., allow_uncertain = 1)` to only allow up to level 1 uncertainty.

With the default setting (`allow_uncertain = TRUE`, level 2), below examples will lead to valid results:

- `"Streptococcus group B (known as S. agalactiae)"`. The text between brackets will be removed and a warning will be thrown that the result *Streptococcus group B* (`B_STRPT_GRPB`) needs review.
- `"S. aureus - please mind: MRSA"`. The last word will be stripped, after which the function will try to find a match. If it does not, the second last word will be stripped, etc. Again, a warning will be thrown that the result *Staphylococcus aureus* (`B_STPHY_AURS`) needs review.
- `"Fluoroquinolone-resistant Neisseria gonorrhoeae"`. The first word will be stripped, after which the function will try to find a match. A warning will be thrown that the result *Neisseria gonorrhoeae* (`B_NESSR_GNRR`) needs review.

There are three helper functions that can be run after using the `as.mo()` function:

- Use `mo_uncertainties()` to get a data.frame that prints in a pretty format with all taxonomic names that were guessed. The output contains the matching score for all matches (see *Background on matching score*).
- Use `mo_failures()` to get a character vector with all values that could not be coerced to a valid value.
- Use `mo_renamed()` to get a data.frame with all values that could be coerced based on old, previously accepted taxonomic names.

**Microbial prevalence of pathogens in humans:**
The intelligent rules consider the prevalence of microorganisms in humans grouped into three groups, which is available as the `prevalence` columns in the microorganisms and microorganisms.old data sets. The grouping into human pathogenic prevalence is explained in the section *Matching score for microorganisms* below.

#### Value

A character vector with additional class `mo`

#### Source

1. Becker K *et al.* **Coagulase-Negative Staphylococci**. 2014. Clin Microbiol Rev. 27(4): 870–926; doi: 10.1128/CMR.0010913
2. Becker K *et al.* **Implications of identifying the recently defined members of the *S. aureus* complex, *S. argenteus* and *S. schweitzeri*: A position paper of members of the ESCMID Study Group for staphylococci and Staphylococcal Diseases (ESGS).** 2019. Clin Microbiol Infect; doi: 10.1016/j.cmi.2019.02.028
3. Becker K *et al.* **Emergence of coagulase-negative staphylococci** 2020. Expert Rev Anti Infect Ther. 18(4):349-366; doi: 10.1080/14787210.2020.1730813
4. Lancefield RC **A serological differentiation of human and other groups of hemolytic streptococci**. 1933. J Exp Med. 57(4): 571–95; doi: 10.1084/jem.57.4.571
5. Catalogue of Life: Annual Checklist (public online taxonomic database), `http://www.catalogueoflife.org` (check included annual version with `catalogue_of_life_version()`).

#### Stable lifecycle

The lifecycle of this function is **stable**. In a stable function, major changes are unlikely. This means that the unlying code will generally evolve by adding new arguments; removing arguments or changing the meaning of existing arguments will be avoided.

If the unlying code needs breaking changes, they will occur gradually. For example, a argument will be deprecated and first continue to work, but will emit an message informing you of the change. Next, typically after at least one newly released version on CRAN, the message will be transformed to an error.

#### Matching score for microorganisms

With ambiguous user input in `as.mo()` and all the `mo_*` functions, the returned results are chosen based on their matching score using `mo_matching_score()`. This matching score $m$, is calculated as:

$$m_{(x,n)} = \frac{l_n - 0.5 \cdot \min \{ \, l_n \, \text{lev}(x, n)}{l_n \cdot p_n \cdot k_n}$$

where:

- $x$ is the user input;
- $n$ is a taxonomic name (genus, species, and subspecies);
- $l_n$ is the length of $n$;
- lev is the Levenshtein distance function, which counts any insertion, deletion and substitution as 1 that is needed to change $x$ into $n$;

- $p_n$ is the human pathogenic prevalence group of $n$, as described below;
- $k_n$ is the taxonomic kingdom of $n$, set as Bacteria = 1, Fungi = 2, Protozoa = 3, Archaea = 4, others = 5.

The grouping into human pathogenic prevalence ($p$) is based on experience from several microbiological laboratories in the Netherlands in conjunction with international reports on pathogen prevalence. **Group 1** (most prevalent microorganisms) consists of all microorganisms where the taxonomic class is Gammaproteobacteria or where the taxonomic genus is *Enterococcus*, *Staphylococcus* or *Streptococcus*. This group consequently contains all common Gram-negative bacteria, such as *Pseudomonas* and *Legionella* and all species within the order Enterobacterales. **Group 2** consists of all microorganisms where the taxonomic phylum is Proteobacteria, Firmicutes, Actinobacteria or Sarcomastigophora, or where the taxonomic genus is *Absidia*, *Acremonium*, *Actinotignum*, *Alternaria*, *Anaerosalibacter*, *Apophysomyces*, *Arachnia*, *Aspergillus*, *Aureobacterium*, *Aureobasidium*, *Bacteroides*, *Basidiobolus*, *Beauveria*, *Blastocystis*, *Branhamella*, *Calymmatobacterium*, *Candida*, *Capnocytophaga*, *Catabacter*, *Chaetomium*, *Chryseobacterium*, *Chryseomonas*, *Chrysonilia*, *Cladophialophora*, *Cladosporium*, *Conidiobolus*, *Cryptococcus*, *Curvularia*, *Exophiala*, *Exserohilum*, *Flavobacterium*, *Fonsecaea*, *Fusarium*, *Fusobacterium*, *Hendersonula*, *Hypomyces*, *Koserella*, *Lelliottia*, *Leptosphaeria*, *Leptotrichia*, *Malassezia*, *Malbranchea*, *Mortierella*, *Mucor*, *Mycocentrospora*, *Mycoplasma*, *Nectria*, *Ochroconis*, *Oidiodendron*, *Phoma*, *Piedraia*, *Pithomyces*, *Pityrosporum*, *Prevotella,\Pseudallescheria*, *Rhizomucor*, *Rhizopus*, *Rhodotorula*, *Scolecobasidium*, *Scopulariopsis*, *Scytalidium,Sporobolomyces*, *Stachybotrys*, *Stomatococcus*, *Treponema*, *Trichoderma*, *Trichophyton*, *Trichosporon*, *Tritirachium* or *Ureaplasma*. **Group 3** consists of all other microorganisms.

All matches are sorted descending on their matching score and for all user input values, the top match will be returned. This will lead to the effect that e.g., `"E. coli"` will return the microbial ID of *Escherichia coli* ($m = 0.688$, a highly prevalent microorganism found in humans) and not *Entamoeba coli* ($m = 0.079$, a less prevalent microorganism in humans), although the latter would alphabetically come first.

##### Catalogue of Life

This package contains the complete taxonomic tree of almost all microorganisms (~70,000 species) from the authoritative and comprehensive Catalogue of Life (CoL, `http://www.catalogueoflife.org`). The CoL is the most comprehensive and authoritative global index of species currently available. Nonetheless, we supplemented the CoL data with data from the List of Prokaryotic names with Standing in Nomenclature (LPSN, lpsn.dsmz.de). This supplementation is needed until the CoL+ project is finished, which we await.

Click here for more information about the included taxa. Check which versions of the CoL and LSPN were included in this package with `catalogue_of_life_version()`.

##### Reference data publicly available

All reference data sets (about microorganisms, antibiotics, R/SI interpretation, EUCAST rules, etc.) in this `AMR` package are publicly and freely available. We continually export our data sets to formats for use in R, SPSS, SAS, Stata and Excel. We also supply flat files that are machine-readable and suitable for input in any software program, such as laboratory information systems. Please find all download links on our website, which is automatically updated with every code change.

##### Read more on our website!

On our website `https://msberends.github.io/AMR/` you can find a comprehensive tutorial about how to conduct AMR analysis, the complete documentation of all functions and an example analysis using WHONET data. As we would like to better understand the backgrounds and needs of our users, please participate in our survey!

#### See Also

microorganisms for the data.frame that is being used to determine ID's.

The mo_* functions (such as mo_genus(), mo_gramstain()) to get properties based on the returned code.

#### Examples

```
# These examples all return "B_STPHY_AURS", the ID of S. aureus:
as.mo("sau") # WHONET code
as.mo("stau")
as.mo("STAU")
as.mo("staaur")
as.mo("S. aureus")
as.mo("S aureus")
as.mo("Staphylococcus aureus")
as.mo("Staphylococcus aureus (MRSA)")
as.mo("Zthafilokkoockus oureuz") # handles incorrect spelling
as.mo("MRSA")    # Methicillin Resistant S. aureus
as.mo("VISA")    # Vancomycin Intermediate S. aureus
as.mo("VRSA")    # Vancomycin Resistant S. aureus
as.mo(115329001) # SNOMED CT code

# Dyslexia is no problem - these all work:
as.mo("Ureaplasma urealyticum")
as.mo("Ureaplasma urealyticus")
as.mo("Ureaplasmium urealytica")
as.mo("Ureaplazma urealitycium")

as.mo("Streptococcus group A")
as.mo("GAS") # Group A Streptococci
as.mo("GBS") # Group B Streptococci

as.mo("S. epidermidis")                 # will remain species: B_STPHY_EPDR
as.mo("S. epidermidis", Becker = TRUE)  # will not remain species: B_STPHY_CONS

as.mo("S. pyogenes")                    # will remain species: B_STRPT_PYGN
as.mo("S. pyogenes", Lancefield = TRUE) # will not remain species: B_STRPT_GRPA

# All mo_* functions use as.mo() internally too (see ?mo_property):
mo_genus("E. coli")                           # returns "Escherichia"
mo_gramstain("E. coli")                       # returns "Gram negative"
mo_is_intrinsic_resistant("E. coli", "vanco") # returns TRUE
```

---

as.rsi *Interpret MIC and disk values, or clean raw R/SI data*

---

##### Description

Interpret minimum inhibitory concentration (MIC) values and disk diffusion diameters according to EUCAST or CLSI, or clean up existing R/SI values. This transforms the input to a new class `rsi`, which is an ordered factor with levels S < I < R. Values that cannot be interpreted will be returned as `NA` with a warning.

##### Usage

```
as.rsi(x, ...)

is.rsi(x)

is.rsi.eligible(x, threshold = 0.05)

## S3 method for class 'mic'
as.rsi(
  x,
  mo = NULL,
  ab = deparse(substitute(x)),
  guideline = "EUCAST",
  uti = FALSE,
  conserve_capped_values = FALSE,
  add_intrinsic_resistance = FALSE,
  reference_data = AMR::rsi_translation,
  ...
)

## S3 method for class 'disk'
as.rsi(
  x,
  mo = NULL,
  ab = deparse(substitute(x)),
  guideline = "EUCAST",
  uti = FALSE,
  add_intrinsic_resistance = FALSE,
  reference_data = AMR::rsi_translation,
  ...
)

## S3 method for class 'data.frame'
as.rsi(
  x,
  ...,
  col_mo = NULL,
  guideline = "EUCAST",
  uti = NULL,
  conserve_capped_values = FALSE,
  add_intrinsic_resistance = FALSE,
  reference_data = rsi_translation
)
```

#### Arguments

| | |
|---|---|
| x | vector of values (for class mic: an MIC value in mg/L, for class disk: a disk diffusion radius in millimetres) |
| ... | for using on a data.frame: names of columns to apply as.rsi() on (supports tidy selection like AMX:VAN). Otherwise: arguments passed on to methods. |
| threshold | maximum fraction of invalid antimicrobial interpretations of x, please see *Examples* |
| mo | any (vector of) text that can be coerced to a valid microorganism code with as.mo(), will be determined automatically if the dplyr package is installed |
| ab | any (vector of) text that can be coerced to a valid antimicrobial code with as.ab() |
| guideline | defaults to the latest included EUCAST guideline, see Details for all options |
| uti | (Urinary Tract Infection) A vector with logicals (TRUE or FALSE) to specify whether a UTI specific interpretation from the guideline should be chosen. For using as.rsi() on a data.frame, this can also be a column containing logicals or when left blank, the data set will be searched for a 'specimen' and rows containing 'urin' (such as 'urine', 'urina') in that column will be regarded isolates from a UTI. See *Examples*. |
| conserve_capped_values | a logical to indicate that MIC values starting with ">" (but not ">=") must always return "R" , and that MIC values starting with "<" (but not "<=") must always return "S" |
| add_intrinsic_resistance | *(only useful when using a EUCAST guideline)* a logical to indicate whether intrinsic antibiotic resistance must also be considered for applicable bug-drug combinations, meaning that e.g. ampicillin will always return "R" in *Klebsiella* species. Determination is based on the intrinsic_resistant data set, that itself is based on 'EUCAST Expert Rules' and 'EUCAST Intrinsic Resistance and Unusual Phenotypes' v3.2 from 2020. |
| reference_data | a data.frame to be used for interpretation, which defaults to the rsi_translation data set. Changing this argument allows for using own interpretation guidelines. This argument must contain a data set that is equal in structure to the rsi_translation data set (same column names and column types). Please note that the guideline argument will be ignored when reference_data is manually set. |
| col_mo | column name of the IDs of the microorganisms (see as.mo()), defaults to the first column of class mo. Values will be coerced using as.mo(). |

#### Details

##### How it works:

The as.rsi() function works in four ways:

1. For **cleaning raw / untransformed data**. The data will be cleaned to only contain values S, I and R and will try its best to determine this with some intelligence. For example, mixed values with R/SI interpretations and MIC values such as "<0.25; S" will be coerced to "S". Combined interpretations for multiple test methods (as seen in laboratory records) such as "S; S" will be coerced to "S", but a value like "S; I" will return NA with a warning that the input is unclear.

2. For **interpreting minimum inhibitory concentration (MIC) values** according to EUCAST or CLSI. You must clean your MIC values first using `as.mic()`, that also gives your columns the new data class `mic`. Also, be sure to have a column with microorganism names or codes. It will be found automatically, but can be set manually using the `mo` argument.
   - Using `dplyr`, R/SI interpretation can be done very easily with either:

     ```
     your_data %>% mutate_if(is.mic, as.rsi)             # until dplyr 1.0.0
     your_data %>% mutate(across(where(is.mic), as.rsi)) # since dplyr 1.0.0
     ```

   - Operators like "<=" will be stripped before interpretation. When using `conserve_capped_values = TRUE`, an MIC value of e.g. ">2" will always return "R", even if the breakpoint according to the chosen guideline is ">=4". This is to prevent that capped values from raw laboratory data would not be treated conservatively. The default behaviour (`conserve_capped_values = FALSE`) considers ">2" to be lower than ">=4" and might in this case return "S" or "I".
3. For **interpreting disk diffusion diameters** according to EUCAST or CLSI. You must clean your disk zones first using `as.disk()`, that also gives your columns the new data class `disk`. Also, be sure to have a column with microorganism names or codes. It will be found automatically, but can be set manually using the `mo` argument.
   - Using `dplyr`, R/SI interpretation can be done very easily with either:

     ```
     your_data %>% mutate_if(is.disk, as.rsi)             # until dplyr 1.0.0
     your_data %>% mutate(across(where(is.disk), as.rsi)) # since dplyr 1.0.0
     ```

4. For **interpreting a complete data set**, with automatic determination of MIC values, disk diffusion diameters, microorganism names or codes, and antimicrobial test results. This is done very simply by running `as.rsi(data)`.

**Supported guidelines:**

For interpreting MIC values as well as disk diffusion diameters, supported guidelines to be used as input for the `guideline` argument are: "CLSI 2010", "CLSI 2011", "CLSI 2012", "CLSI 2013", "CLSI 2014", "CLSI 2015", "CLSI 2016", "CLSI 2017", "CLSI 2018", "CLSI 2019", "EUCAST 2011", "EUCAST 2012", "EUCAST 2013", "EUCAST 2014", "EUCAST 2015", "EUCAST 2016", "EUCAST 2017", "EUCAST 2018", "EUCAST 2019", "EUCAST 2020".

Simply using `"CLSI"` or `"EUCAST"` as input will automatically select the latest version of that guideline. You can set your own data set using the `reference_data` argument. The `guideline` argument will then be ignored.

**After interpretation:**

After using `as.rsi()`, you can use the `eucast_rules()` defined by EUCAST to (1) apply inferred susceptibility and resistance based on results of other antimicrobials and (2) apply intrinsic resistance based on taxonomic properties of a microorganism.

**Machine readable interpretation guidelines:**

The repository of this package contains a machine readable version of all guidelines. This is a CSV file consisting of 18,650 rows and 10 columns. This file is machine readable, since it contains one row for every unique combination of the test method (MIC or disk diffusion), the antimicrobial agent and the microorganism. **This allows for easy implementation of these rules in laboratory information systems (LIS)**. Note that it only contains interpretation guidelines for humans - interpretation guidelines from CLSI for animals were removed.

**Other:**

The function `is.rsi.eligible()` returns `TRUE` when a columns contains at most 5% invalid antimicrobial interpretations (not S and/or I and/or R), and `FALSE` otherwise. The threshold of 5% can be set with the `threshold` argument.

##### Value

Ordered factor with new class rsi

##### Interpretation of R and S/I

In 2019, the European Committee on Antimicrobial Susceptibility Testing (EUCAST) has decided to change the definitions of susceptibility testing categories R and S/I as shown below (https://www.eucast.org/newsiandr/).

- **R = Resistant**
  A microorganism is categorised as *Resistant* when there is a high likelihood of therapeutic failure even when there is increased exposure. Exposure is a function of how the mode of administration, dose, dosing interval, infusion time, as well as distribution and excretion of the antimicrobial agent will influence the infecting organism at the site of infection.
- **S = Susceptible**
  A microorganism is categorised as *Susceptible, standard dosing regimen*, when there is a high likelihood of therapeutic success using a standard dosing regimen of the agent.
- **I = Increased exposure, but still susceptible**
  A microorganism is categorised as *Susceptible, Increased exposure* when there is a high likelihood of therapeutic success because exposure to the agent is increased by adjusting the dosing regimen or by its concentration at the site of infection.

This AMR package honours this new insight. Use susceptibility() (equal to proportion_SI()) to determine antimicrobial susceptibility and count_susceptible() (equal to count_SI()) to count susceptible isolates.

##### Stable lifecycle

The lifecycle of this function is **stable**. In a stable function, major changes are unlikely. This means that the unlying code will generally evolve by adding new arguments; removing arguments or changing the meaning of existing arguments will be avoided.

If the unlying code needs breaking changes, they will occur gradually. For example, a argument will be deprecated and first continue to work, but will emit an message informing you of the change. Next, typically after at least one newly released version on CRAN, the message will be transformed to an error.

##### Reference data publicly available

All reference data sets (about microorganisms, antibiotics, R/SI interpretation, EUCAST rules, etc.) in this AMR package are publicly and freely available. We continually export our data sets to formats for use in R, SPSS, SAS, Stata and Excel. We also supply flat files that are machine-readable and suitable for input in any software program, such as laboratory information systems. Please find all download links on our website, which is automatically updated with every code change.

##### Read more on our website!

On our website https://msberends.github.io/AMR/ you can find a comprehensive tutorial about how to conduct AMR analysis, the complete documentation of all functions and an example analysis using WHONET data. As we would like to better understand the backgrounds and needs of our users, please participate in our survey!

#### See Also

as.mic(), as.disk(), as.mo()

#### Examples

```
summary(example_isolates) # see all R/SI results at a glance

if (require("skimr")) {
  # class <rsi> supported in skim() too:
  skim(example_isolates)
}

# For INTERPRETING disk diffusion and MIC values -----------------------

# a whole data set, even with combined MIC values and disk zones
df <- data.frame(microorganism = "Escherichia coli",
                 AMP = as.mic(8),
                 CIP = as.mic(0.256),
                 GEN = as.disk(18),
                 TOB = as.disk(16),
                 NIT = as.mic(32),
                 ERY = "R")
as.rsi(df)

# for single values
as.rsi(x = as.mic(2),
       mo = as.mo("S. pneumoniae"),
       ab = "AMP",
       guideline = "EUCAST")

as.rsi(x = as.disk(18),
       mo = "Strep pneu",  # `mo` will be coerced with as.mo()
       ab = "ampicillin",  # and `ab` with as.ab()
       guideline = "EUCAST")

# the dplyr way
if (require("dplyr")) {
  df %>% mutate_if(is.mic, as.rsi)
  df %>% mutate_if(function(x) is.mic(x) | is.disk(x), as.rsi)
  df %>% mutate(across(where(is.mic), as.rsi))
  df %>% mutate_at(vars(AMP:TOB), as.rsi)
  df %>% mutate(across(AMP:TOB, as.rsi))

  df %>%
    mutate_at(vars(AMP:TOB), as.rsi, mo = .$microorganism)

  # to include information about urinary tract infections (UTI)
  data.frame(mo = "E. coli",
             NIT = c("<= 2", 32),
             from_the_bladder = c(TRUE, FALSE)) %>%
    as.rsi(uti = "from_the_bladder")

  data.frame(mo = "E. coli",
             NIT = c("<= 2", 32),
             specimen = c("urine", "blood")) %>%
```

```
    as.rsi() # automatically determines urine isolates

  df %>%
    mutate_at(vars(AMP:NIT), as.rsi, mo = "E. coli", uti = TRUE)
}

# For CLEANING existing R/SI values ------------------------------------

as.rsi(c("S", "I", "R", "A", "B", "C"))
as.rsi("<= 0.002; S") # will return "S"
rsi_data <- as.rsi(c(rep("S", 474), rep("I", 36), rep("R", 370)))
is.rsi(rsi_data)
plot(rsi_data)    # for percentages
barplot(rsi_data) # for frequencies

# the dplyr way
if (require("dplyr")) {
  example_isolates %>%
    mutate_at(vars(PEN:RIF), as.rsi)
  # same:
  example_isolates %>%
    as.rsi(PEN:RIF)

  # fastest way to transform all columns with already valid AMR results to class `rsi`:
  example_isolates %>%
    mutate_if(is.rsi.eligible, as.rsi)

  # note: from dplyr 1.0.0 on, this will be:
  # example_isolates %>%
  #   mutate(across(where(is.rsi.eligible), as.rsi))
}
```

---

`atc_online_property` *Get ATC properties from WHOCC website*

---

#### Description

Gets data from the WHO to determine properties of an ATC (e.g. an antibiotic), such as the name, defined daily dose (DDD) or standard unit.

#### Usage

```
atc_online_property(
  atc_code,
  property,
  administration = "O",
  url = "https://www.whocc.no/atc_ddd_index/?code=%s&showdescription=no",
 url_vet = "https://www.whocc.no/atcvet/atcvet_index/?code=%s&showdescription=no"
)

atc_online_groups(atc_code, ...)

atc_online_ddd(atc_code, ...)
```

#### Arguments

| | |
|---|---|
| `atc_code` | a character or character vector with ATC code(s) of antibiotic(s) |
| `property` | property of an ATC code. Valid values are `"ATC"`, `"Name"`, `"DDD"`, `"U"` (`"unit"`), `"Adm.R"`, `"Note"` and `groups`. For this last option, all hierarchical groups of an ATC code will be returned, see Examples. |
| `administration` | type of administration when using `property = "Adm.R"`, see Details |
| `url` | url of website of the WHOCC. The sign %s can be used as a placeholder for ATC codes. |
| `url_vet` | url of website of the WHOCC for veterinary medicine. The sign %s can be used as a placeholder for ATC_vet codes (that all start with "Q"). |
| `...` | arguments to pass on to `atc_property` |

#### Details

Options for argument `administration`:

- `"Implant"` = Implant
- `"Inhal"` = Inhalation
- `"Instill"` = Instillation
- `"N"` = nasal
- `"O"` = oral
- `"P"` = parenteral
- `"R"` = rectal
- `"SL"` = sublingual/buccal
- `"TD"` = transdermal
- `"V"` = vaginal

Abbreviations of return values when using `property = "U"` (unit):

- `"g"` = gram
- `"mg"` = milligram
- ‘"mcg"‘ = microgram
- `"U"` = unit
- `"TU"` = thousand units
- `"MU"` = million units
- `"mmol"` = millimole
- `"ml"` = milliliter (e.g. eyedrops)

**N.B. This function requires an internet connection and only works if the following packages are installed: `curl`, `rvest`, `xml2`.**

#### Stable lifecycle

The lifecycle of this function is **stable**. In a stable function, major changes are unlikely. This means that the unlying code will generally evolve by adding new arguments; removing arguments or changing the meaning of existing arguments will be avoided.

If the unlying code needs breaking changes, they will occur gradually. For example, a argument will be deprecated and first continue to work, but will emit an message informing you of the change. Next, typically after at least one newly released version on CRAN, the message will be transformed to an error.

##### Read more on our website!

On our website `https://msberends.github.io/AMR/` you can find a comprehensive tutorial about how to conduct AMR analysis, the complete documentation of all functions and an example analysis using WHONET data. As we would like to better understand the backgrounds and needs of our users, please participate in our survey!

##### Source

`https://www.whocc.no/atc_ddd_alterations__cumulative/ddd_alterations/abbrevations/`

##### Examples

```
# oral DDD (Defined Daily Dose) of amoxicillin
atc_online_property("J01CA04", "DDD", "O")

# parenteral DDD (Defined Daily Dose) of amoxicillin
atc_online_property("J01CA04", "DDD", "P")

atc_online_property("J01CA04", property = "groups") # search hierarchical groups of amoxicillin
```

---

#### `availability` *Check availability of columns*

---

##### Description

Easy check for data availability of all columns in a data set. This makes it easy to get an idea of which antimicrobial combinations can be used for calculation with e.g. `susceptibility()` and `resistance()`.

##### Usage

```
availability(tbl, width = NULL)
```

##### Arguments

| | |
|---|---|
| `tbl` | a data.frame or list |
| `width` | number of characters to present the visual availability, defaults to filling the width of the console |

##### Details

The function returns a data.frame with columns `"resistant"` and `"visual_resistance"`. The values in that columns are calculated with `resistance()`.

##### Value

data.frame with column names of `tbl` as row names

##### Stable lifecycle

The lifecycle of this function is **stable**. In a stable function, major changes are unlikely. This means that the unlying code will generally evolve by adding new arguments; removing arguments or changing the meaning of existing arguments will be avoided.

If the unlying code needs breaking changes, they will occur gradually. For example, a argument will be deprecated and first continue to work, but will emit an message informing you of the change. Next, typically after at least one newly released version on CRAN, the message will be transformed to an error.

##### Read more on our website!

On our website `https://msberends.github.io/AMR/` you can find a comprehensive tutorial about how to conduct AMR analysis, the complete documentation of all functions and an example analysis using WHONET data. As we would like to better understand the backgrounds and needs of our users, please participate in our survey!

##### Examples

```
availability(example_isolates)

if (require("dplyr")) {
  example_isolates %>%
    filter(mo == as.mo("E. coli")) %>%
    select_if(is.rsi) %>%
    availability()
}
```

---

#### bug_drug_combinations *Determine bug-drug combinations*

---

##### Description

Determine antimicrobial resistance (AMR) of all bug-drug combinations in your data set where at least 30 (default) isolates are available per species. Use `format()` on the result to prettify it to a publicable/printable format, see Examples.

##### Usage

```
bug_drug_combinations(x, col_mo = NULL, FUN = mo_shortname, ...)

## S3 method for class 'bug_drug_combinations'
format(
  x,
  translate_ab = "name (ab, atc)",
  language = get_locale(),
  minimum = 30,
  combine_SI = TRUE,
  combine_IR = FALSE,
  add_ab_group = TRUE,
  remove_intrinsic_resistant = FALSE,
  decimal.mark = getOption("OutDec"),
```

```
  big.mark = ifelse(decimal.mark == ",", ".", ","),
  ...
)
```

#### Arguments

- `x` — data with antibiotic columns, such as amox, AMX and AMC
- `col_mo` — column name of the IDs of the microorganisms (see `as.mo()`), defaults to the first column of class `mo`. Values will be coerced using `as.mo()`.
- `FUN` — function to call on the mo column to transform the microorganism IDs, defaults to `mo_shortname()`
- `...` — arguments passed on to FUN
- `translate_ab` — character of length 1 containing column names of the antibiotics data set
- `language` — language of the returned text, defaults to system language (see `get_locale()`) and can also be set with `getOption("AMR_locale")`. Use `language = NULL` or `language = ""` to prevent translation.
- `minimum` — the minimum allowed number of available (tested) isolates. Any isolate count lower than `minimum` will return `NA` with a warning. The default number of `30` isolates is advised by the Clinical and Laboratory Standards Institute (CLSI) as best practice, see Source.
- `combine_SI` — a logical to indicate whether all values of S and I must be merged into one, so the output only consists of S+I vs. R (susceptible vs. resistant). This used to be the argument `combine_IR`, but this now follows the redefinition by EUCAST about the interpretation of I (increased exposure) in 2019, see section 'Interpretation of S, I and R' below. Default is `TRUE`.
- `combine_IR` — logical to indicate whether values R and I should be summed
- `add_ab_group` — logical to indicate where the group of the antimicrobials must be included as a first column
- `remove_intrinsic_resistant` — logical to indicate that rows and columns with 100% resistance for all tested antimicrobials must be removed from the table
- `decimal.mark` — the character to be used to indicate the numeric decimal point.
- `big.mark` — character; if not empty used as mark between every `big.interval` decimals *before* (hence `big`) the decimal point.

#### Details

The function `format()` calculates the resistance per bug-drug combination. Use `combine_IR = FALSE` (default) to test R vs. S+I and `combine_IR = TRUE` to test R+I vs. S.

#### Value

The function `bug_drug_combinations()` returns a data.frame with columns "mo", "ab", "S", "I", "R" and "total".

#### Stable lifecycle

The lifecycle of this function is **stable**. In a stable function, major changes are unlikely. This means that the unlying code will generally evolve by adding new arguments; removing arguments or changing the meaning of existing arguments will be avoided.

If the unlying code needs breaking changes, they will occur gradually. For example, a argument will be deprecated and first continue to work, but will emit an message informing you of the change. Next, typically after at least one newly released version on CRAN, the message will be transformed to an error.

#### Read more on our website!

On our website `https://msberends.github.io/AMR/` you can find a comprehensive tutorial about how to conduct AMR analysis, the complete documentation of all functions and an example analysis using WHONET data. As we would like to better understand the backgrounds and needs of our users, please participate in our survey!

#### Source

**M39 Analysis and Presentation of Cumulative Antimicrobial Susceptibility Test Data, 4th Edition**, 2014, *Clinical and Laboratory Standards Institute (CLSI)*. `https://clsi.org/standards/products/microbiology/documents/m39/`.

#### Examples

```
x <- bug_drug_combinations(example_isolates)
x
format(x, translate_ab = "name (atc)")

# Use FUN to change to transformation of microorganism codes
bug_drug_combinations(example_isolates,
                      FUN = mo_gramstain)

bug_drug_combinations(example_isolates,
                      FUN = function(x) ifelse(x == as.mo("E. coli"),
                                               "E. coli",
                                               "Others"))
```

---

### `catalogue_of_life` *The Catalogue of Life*

---

#### Description

This package contains the complete taxonomic tree of almost all microorganisms from the authoritative and comprehensive Catalogue of Life.

#### Catalogue of Life

This package contains the complete taxonomic tree of almost all microorganisms (~70,000 species) from the authoritative and comprehensive Catalogue of Life (CoL, `http://www.catalogueoflife.org`). The CoL is the most comprehensive and authoritative global index of species currently available. Nonetheless, we supplemented the CoL data with data from the List of Prokaryotic names with Standing in Nomenclature (LPSN, lpsn.dsmz.de). This supplementation is needed until the CoL+ project is finished, which we await.

Click here for more information about the included taxa. Check which versions of the CoL and LSPN were included in this package with `catalogue_of_life_version()`.

#### Included taxa

Included are:

- All ~55,000 (sub)species from the kingdoms of Archaea, Bacteria, Chromista and Protozoa
- All ~5,000 (sub)species from these orders of the kingdom of Fungi: Eurotiales, Microascales, Mucorales, Onygenales, Pneumocystales, Saccharomycetales, Schizosaccharomycetales and Tremellales, as well as ~4,600 other fungal (sub)species. The kingdom of Fungi is a very large taxon with almost 300,000 different (sub)species, of which most are not microbial (but rather macroscopic, like mushrooms). Because of this, not all fungi fit the scope of this package and including everything would tremendously slow down our algorithms too. By only including the aforementioned taxonomic orders, the most relevant fungi are covered (such as all species of *Aspergillus*, *Candida*, *Cryptococcus*, *Histplasma*, *Pneumocystis*, *Saccharomyces* and *Trichophyton*).
- All ~2,200 (sub)species from ~50 other relevant genera from the kingdom of Animalia (such as *Strongyloides* and *Taenia*)
- All ~13,000 previously accepted names of all included (sub)species (these were taxonomically renamed)
- The complete taxonomic tree of all included (sub)species: from kingdom to subspecies
- The responsible author(s) and year of scientific publication

The Catalogue of Life (`http://www.catalogueoflife.org`) is the most comprehensive and authoritative global index of species currently available. It holds essential information on the names, relationships and distributions of over 1.9 million species. The Catalogue of Life is used to support the major biodiversity and conservation information services such as the Global Biodiversity Information Facility (GBIF), Encyclopedia of Life (EoL) and the International Union for Conservation of Nature Red List. It is recognised by the Convention on Biological Diversity as a significant component of the Global Taxonomy Initiative and a contribution to Target 1 of the Global Strategy for Plant Conservation.

The syntax used to transform the original data to a cleansed R format, can be found here: `https://github.com/msberends/AMR/blob/master/data-raw/reproduction_of_microorganisms.R`.

#### Read more on our website!

On our website `https://msberends.github.io/AMR/` you can find a comprehensive tutorial about how to conduct AMR analysis, the complete documentation of all functions and an example analysis using WHONET data. As we would like to better understand the backgrounds and needs of our users, please participate in our survey!

#### See Also

Data set microorganisms for the actual data.
Function `as.mo()` to use the data for intelligent determination of microorganisms.

#### Examples

```
# Get version info of included data set
catalogue_of_life_version()

# Get a note when a species was renamed
mo_shortname("Chlamydophila psittaci")
# Note: 'Chlamydophila psittaci' (Everett et al., 1999) was renamed back to
```

```
#       'Chlamydia psittaci' (Page, 1968)
#> [1] "C. psittaci"

# Get any property from the entire taxonomic tree for all included species
mo_class("E. coli")
#> [1] "Gammaproteobacteria"

mo_family("E. coli")
#> [1] "Enterobacteriaceae"

mo_gramstain("E. coli") # based on kingdom and phylum, see ?mo_gramstain
#> [1] "Gram-negative"

mo_ref("E. coli")
#> [1] "Castellani et al., 1919"

# Do not get mistaken - this package is about microorganisms
mo_kingdom("C. elegans")
#> [1] "Fungi"                # Fungi?!
mo_name("C. elegans")
#> [1] "Cladosporium elegans"  # Because a microorganism was found
```

---

catalogue_of_life_version
*Version info of included Catalogue of Life*

---

#### Description

This function returns information about the included data from the Catalogue of Life.

#### Usage

```
catalogue_of_life_version()
```

#### Details

For DSMZ, see microorganisms.

#### Value

a list, which prints in pretty format

#### Catalogue of Life

This package contains the complete taxonomic tree of almost all microorganisms (~70,000 species) from the authoritative and comprehensive Catalogue of Life (CoL, `http://www.catalogueoflife.org`). The CoL is the most comprehensive and authoritative global index of species currently available. Nonetheless, we supplemented the CoL data with data from the List of Prokaryotic names with Standing in Nomenclature (LPSN, lpsn.dsmz.de). This supplementation is needed until the CoL+ project is finished, which we await.

Click here for more information about the included taxa. Check which versions of the CoL and LSPN were included in this package with `catalogue_of_life_version()`.

##### Read more on our website!

On our website `https://msberends.github.io/AMR/` you can find a comprehensive tutorial about how to conduct AMR analysis, the complete documentation of all functions and an example analysis using WHONET data. As we would like to better understand the backgrounds and needs of our users, please participate in our survey!

##### See Also

microorganisms

---

#### count *Count available isolates*

---

##### Description

These functions can be used to count resistant/susceptible microbial isolates. All functions support quasiquotation with pipes, can be used in `summarise()` from the `dplyr` package and also support grouped variables, please see *Examples*.

`count_resistant()` should be used to count resistant isolates, `count_susceptible()` should be used to count susceptible isolates.

##### Usage

```
count_resistant(..., only_all_tested = FALSE)

count_susceptible(..., only_all_tested = FALSE)

count_R(..., only_all_tested = FALSE)

count_IR(..., only_all_tested = FALSE)

count_I(..., only_all_tested = FALSE)

count_SI(..., only_all_tested = FALSE)

count_S(..., only_all_tested = FALSE)

count_all(..., only_all_tested = FALSE)

n_rsi(..., only_all_tested = FALSE)

count_df(
  data,
  translate_ab = "name",
  language = get_locale(),
  combine_SI = TRUE,
  combine_IR = FALSE
)
```

#### Arguments

`...` — one or more vectors (or columns) with antibiotic interpretations. They will be transformed internally with `as.rsi()` if needed.

`only_all_tested` — (for combination therapies, i.e. using more than one variable for `...`): a logical to indicate that isolates must be tested for all antibiotics, see section *Combination therapy* below

`data` — a data.frame containing columns with class `rsi` (see `as.rsi()`)

`translate_ab` — a column name of the antibiotics data set to translate the antibiotic abbreviations to, using `ab_property()`

`language` — language of the returned text, defaults to system language (see `get_locale()`) and can also be set with `getOption("AMR_locale")`. Use `language = NULL` or `language = ""` to prevent translation.

`combine_SI` — a logical to indicate whether all values of S and I must be merged into one, so the output only consists of S+I vs. R (susceptible vs. resistant). This used to be the argument `combine_IR`, but this now follows the redefinition by EUCAST about the interpretation of I (increased exposure) in 2019, see section 'Interpretation of S, I and R' below. Default is `TRUE`.

`combine_IR` — a logical to indicate whether all values of I and R must be merged into one, so the output only consists of S vs. I+R (susceptible vs. non-susceptible). This is outdated, see argument `combine_SI`.

#### Details

These functions are meant to count isolates. Use the `resistance()`/`susceptibility()` functions to calculate microbial resistance/susceptibility.

The function `count_resistant()` is equal to the function `count_R()`. The function `count_susceptible()` is equal to the function `count_SI()`.

The function `n_rsi()` is an alias of `count_all()`. They can be used to count all available isolates, i.e. where all input antibiotics have an available result (S, I or R). Their use is equal to `n_distinct()`. Their function is equal to `count_susceptible(...) + count_resistant(...)`.

The function `count_df()` takes any variable from `data` that has an `rsi` class (created with `as.rsi()`) and counts the number of S's, I's and R's. It also supports grouped variables. The function `rsi_df()` works exactly like `count_df()`, but adds the percentage of S, I and R.

#### Value

An integer

#### Stable lifecycle

The lifecycle of this function is **stable**. In a stable function, major changes are unlikely. This means that the unlying code will generally evolve by adding new arguments; removing arguments or changing the meaning of existing arguments will be avoided.

If the unlying code needs breaking changes, they will occur gradually. For example, a argument will be deprecated and first continue to work, but will emit an message informing you of the change. Next, typically after at least one newly released version on CRAN, the message will be transformed to an error.

##### Interpretation of R and S/I

In 2019, the European Committee on Antimicrobial Susceptibility Testing (EUCAST) has decided to change the definitions of susceptibility testing categories R and S/I as shown below (https://www.eucast.org/newsiandr/).

- **R = Resistant**
  A microorganism is categorised as *Resistant* when there is a high likelihood of therapeutic failure even when there is increased exposure. Exposure is a function of how the mode of administration, dose, dosing interval, infusion time, as well as distribution and excretion of the antimicrobial agent will influence the infecting organism at the site of infection.
- **S = Susceptible**
  A microorganism is categorised as *Susceptible, standard dosing regimen*, when there is a high likelihood of therapeutic success using a standard dosing regimen of the agent.
- **I = Increased exposure, but still susceptible**
  A microorganism is categorised as *Susceptible, Increased exposure* when there is a high likelihood of therapeutic success because exposure to the agent is increased by adjusting the dosing regimen or by its concentration at the site of infection.

This AMR package honours this new insight. Use `susceptibility()` (equal to `proportion_SI()`) to determine antimicrobial susceptibility and `count_susceptible()` (equal to `count_SI()`) to count susceptible isolates.

##### Combination therapy

When using more than one variable for ... (= combination therapy), use `only_all_tested` to only count isolates that are tested for all antibiotics/variables that you test them for. See this example for two antibiotics, Drug A and Drug B, about how `susceptibility()` works to calculate the %SI:

```
--------------------------------------------------------------------
                    only_all_tested = FALSE  only_all_tested = TRUE
                    -----------------------  -----------------------
 Drug A    Drug B   include as  include as   include as  include as
                    numerator   denominator  numerator   denominator
--------  --------  ----------  -----------  ----------  -----------
 S or I    S or I       X            X           X            X
   R       S or I       X            X           X            X
  <NA>     S or I       X            X           -            -
 S or I      R          X            X           X            X
   R         R          -            X           -            X
  <NA>       R          -            -           -            -
 S or I     <NA>        X            X           -            -
   R        <NA>        -            -           -            -
  <NA>      <NA>        -            -           -            -
--------------------------------------------------------------------
```

Please note that, in combination therapies, for `only_all_tested = TRUE` applies that:

```
    count_S()    +   count_I()    +   count_R()    = count_all()
  proportion_S() + proportion_I() + proportion_R() = 1
```

and that, in combination therapies, for `only_all_tested = FALSE` applies that:

```
  count_S()    +   count_I()    +   count_R()    >= count_all()
proportion_S() + proportion_I() + proportion_R() >= 1
```

Using only_all_tested has no impact when only using one antibiotic as input.

##### Read more on our website!

On our website https://msberends.github.io/AMR/ you can find a comprehensive tutorial about how to conduct AMR analysis, the complete documentation of all functions and an example analysis using WHONET data. As we would like to better understand the backgrounds and needs of our users, please participate in our survey!

##### See Also

proportion_* to calculate microbial resistance and susceptibility.

##### Examples

```
# example_isolates is a data set available in the AMR package.
?example_isolates

count_resistant(example_isolates$AMX)   # counts "R"
count_susceptible(example_isolates$AMX) # counts "S" and "I"
count_all(example_isolates$AMX)         # counts "S", "I" and "R"

# be more specific
count_S(example_isolates$AMX)
count_SI(example_isolates$AMX)
count_I(example_isolates$AMX)
count_IR(example_isolates$AMX)
count_R(example_isolates$AMX)

# Count all available isolates
count_all(example_isolates$AMX)
n_rsi(example_isolates$AMX)

# n_rsi() is an alias of count_all().
# Since it counts all available isolates, you can
# calculate back to count e.g. susceptible isolates.
# These results are the same:
count_susceptible(example_isolates$AMX)
susceptibility(example_isolates$AMX) * n_rsi(example_isolates$AMX)

if (require("dplyr")) {
  example_isolates %>%
    group_by(hospital_id) %>%
    summarise(R  = count_R(CIP),
              I  = count_I(CIP),
              S  = count_S(CIP),
              n1 = count_all(CIP),  # the actual total; sum of all three
              n2 = n_rsi(CIP),      # same - analogous to n_distinct
              total = n())          # NOT the number of tested isolates!

  # Count co-resistance between amoxicillin/clav acid and gentamicin,
  # so we can see that combination therapy does a lot more than mono therapy.
```

```
  # Please mind that `susceptibility()` calculates percentages right away instead.
  example_isolates %>% count_susceptible(AMC) # 1433
  example_isolates %>% count_all(AMC)         # 1879

  example_isolates %>% count_susceptible(GEN) # 1399
  example_isolates %>% count_all(GEN)         # 1855

  example_isolates %>% count_susceptible(AMC, GEN) # 1764
  example_isolates %>% count_all(AMC, GEN)         # 1936

  # Get number of S+I vs. R immediately of selected columns
  example_isolates %>%
    select(AMX, CIP) %>%
    count_df(translate = FALSE)

  # It also supports grouping variables
  example_isolates %>%
    select(hospital_id, AMX, CIP) %>%
    group_by(hospital_id) %>%
    count_df(translate = FALSE)
}
```

#### eucast_rules *Apply EUCAST rules*

##### Description

Apply rules for clinical breakpoints and intrinsic resistance as defined by the European Committee on Antimicrobial Susceptibility Testing (EUCAST, https://eucast.org), see *Source*.

To improve the interpretation of the antibiogram before EUCAST rules are applied, some non-EUCAST rules can applied at default, see Details.

##### Usage

```
eucast_rules(
  x,
  col_mo = NULL,
  info = interactive(),
  rules = getOption("AMR_eucastrules", default = c("breakpoints", "expert")),
  verbose = FALSE,
  version_breakpoints = 10,
  version_expertrules = 3.2,
  ampc_cephalosporin_resistance = NA,
  ...
)
```

##### Arguments

| | |
|---|---|
| x | data with antibiotic columns, such as amox, AMX and AMC |
| col_mo | column name of the IDs of the microorganisms (see as.mo()), defaults to the first column of class mo. Values will be coerced using as.mo(). |

| | |
|---|---|
| `info` | a logical to indicate whether progress should be printed to the console, defaults to only print while in interactive sessions |
| `rules` | a character vector that specifies which rules should be applied. Must be one or more of `"breakpoints"`, `"expert"`, `"other"`, `"all"`, and defaults to `c("breakpoints","expert")`. The default value can be set to another value, e.g. using `options(AMR_eucastrules = "all")`. |
| `verbose` | a logical to turn Verbose mode on and off (default is off). In Verbose mode, the function does not apply rules to the data, but instead returns a data set in logbook form with extensive info about which rows and columns would be effected and in which way. Using Verbose mode takes a lot more time. |
| `version_breakpoints` | the version number to use for the EUCAST Clinical Breakpoints guideline. Currently supported: 10.0. |
| `version_expertrules` | the version number to use for the EUCAST Expert Rules and Intrinsic Resistance guideline. Currently supported: 3.1, 3.2. |
| `ampc_cephalosporin_resistance` | a character value that should be applied for AmpC de-repressed cephalosporin-resistant mutants, defaults to `NA`. Currently only works when `version_expertrules` is `3.2`; '*EUCAST Expert Rules v3.2 on Enterobacterales*' states that susceptible (S) results of cefotaxime, ceftriaxone and ceftazidime should be reported with a note, or results should be suppressed (emptied) for these agents. A value of `NA` for this argument will remove results for these agents, while e.g. a value of `"R"` will make the results for these agents resistant. Use `NULL` to not alter the results for AmpC de-repressed cephalosporin-resistant mutants.<br>For *EUCAST Expert Rules* v3.2, this rule applies to: *Enterobacter, Klebsiella aerogenes, Citrobacter braakii, freundii, gillenii, murliniae, rodenticum, sedlakii, werkmanii, youngae, Hafnia alvei, Serratia, Morganella morganii, Providencia.* |
| `...` | column name of an antibiotic, please see section *Antibiotics* below |

#### Details

**Note:** This function does not translate MIC values to RSI values. Use `as.rsi()` for that.
**Note:** When ampicillin (AMP, J01CA01) is not available but amoxicillin (AMX, J01CA04) is, the latter will be used for all rules where there is a dependency on ampicillin. These drugs are interchangeable when it comes to expression of antimicrobial resistance.

The file containing all EUCAST rules is located here: https://github.com/msberends/AMR/blob/master/data-raw/eucast_rules.tsv.

##### 'Other' rules:

Before further processing, two non-EUCAST rules about drug combinations can be applied to improve the efficacy of the EUCAST rules, and the reliability of your data (analysis). These rules are:

1. A drug **with** enzyme inhibitor will be set to S if the same drug **without** enzyme inhibitor is S
2. A drug **without** enzyme inhibitor will be set to R if the same drug **with** enzyme inhibitor is R

Important examples include amoxicillin and amoxicillin/clavulanic acid, and trimethoprim and trimethoprim/sulfamethoxazole. Needless to say, for these rules to work, both drugs must be available in the data set.

Since these rules are not officially approved by EUCAST, they are not applied at default. To use these rules, include "other" to the rules argument, or use eucast_rules(...,rules = "all"). You can also set the option AMR_eucastrules, i.e. run options(AMR_eucastrules = "all").

#### Value

The input of x, possibly with edited values of antibiotics. Or, if verbose = TRUE, a data.frame with all original and new values of the affected bug-drug combinations.

#### Antibiotics

To define antibiotics column names, leave as it is to determine it automatically with guess_ab_col() or input a text (case-insensitive), or use NULL to skip a column (e.g. TIC = NULL to skip ticarcillin). Manually defined but non-existing columns will be skipped with a warning.

The following antibiotics are used for the functions eucast_rules() and mdro(). These are shown below in the format 'name (antimicrobial ID, ATC code)', sorted alphabetically:

Amikacin (AMK, J01GB06), amoxicillin (AMX, J01CA04), amoxicillin/clavulanic acid (AMC, J01CR02), ampicillin (AMP, J01CA01), ampicillin/sulbactam (SAM, J01CR01), avoparcin (AVO, no ATC code), azithromycin (AZM, J01FA10), azlocillin (AZL, J01CA09), aztreonam (ATM, J01DF01), bacampicillin (BAM, J01CA06), benzylpenicillin (PEN, J01CE01), cadazolid (CDZ, J01DD09), carbenicillin (CRB, J01CA03), carindacillin (CRN, J01CA05), cefacetrile (CAC, J01DB10), cefaclor (CEC, J01DC04), cefadroxil (CFR, J01DB05), cefaloridine (RID, J01DB02), cefamandole (MAN, J01DC03), cefatrizine (CTZ, J01DB07), cefazedone (CZD, J01DB06), cefazolin (CZO, J01DB04), cefcapene (CCP, no ATC code), cefcapene pivoxil (CCX, no ATC code), cefdinir (CDR, J01DD15), cefditoren (DIT, J01DD16), cefditoren pivoxil (DIX, no ATC code), cefepime (FEP, J01DE01), cefetamet (CAT, J01DD10), cefetamet pivoxil (CPI, no ATC code), cefixime (CFM, J01DD08), cefmenoxime (CMX, J01DD05), cefmetazole (CMZ, J01DC09), cefodizime (DIZ, J01DD09), cefonicid (CID, J01DC06), cefoperazone (CFP, J01DD12), cefoperazone/sulbactam (CSL, J01DD62), ceforanide (CND, J01DC11), cefotaxime (CTX, J01DD01), cefotaxime/clavulanic acid (CTC, no ATC code), cefotaxime/sulbactam (CTS, no ATC code), cefotetan (CTT, J01DC05), cefotiam (CTF, J01DC07), cefotiam hexetil (CHE, no ATC code), cefovecin (FOV, no ATC code), cefoxitin (FOX, J01DC01), cefoxitin screening (FOX1, no ATC code), cefpimizole (CFZ, no ATC code), cefpiramide (CPM, J01DD11), cefpirome (CPO, J01DE02), cefpodoxime (CPD, J01DD13), cefpodoxime proxetil (CPX, no ATC code), cefpodoxime/clavulanic acid (CDC, no ATC code), cefprozil (CPR, J01DC10), cefroxadine (CRD, J01DB11), cefsulodin (CFS, J01DD03), ceftaroline (CPT, J01DI02), ceftazidime (CAZ, J01DD02), ceftazidime/avibactam (CZA, no ATC code), ceftazidime/clavulanic acid (CCV, J01DD52), cefteram (CEM, no ATC code), cefteram pivoxil (CPL, no ATC code), ceftezole (CTL, J01DB12), ceftibuten (CTB, J01DD14), ceftiofur (TIO, no ATC code), ceftizoxime (CZX, J01DD07), ceftizoxime alapivoxil (CZP, no ATC code), ceftobiprole (BPR, J01DI01), ceftobiprole medocaril (CFM1, J01DI01), ceftolozane/enzyme inhibitor (CEI, J01DI54), ceftriaxone (CRO, J01DD04), cefuroxime (CXM, J01DC02), cephalexin (LEX, J01DB01), cephalothin (CEP, J01DB03), cephapirin (HAP, J01DB08), cephradine (CED, J01DB09), chloramphenicol (CHL, J01BA01), ciprofloxacin (CIP, J01MA02), clarithromycin (CLR, J01FA09), clindamycin (CLI, J01FF01), colistin (COL, J01XB01), cycloserine (CYC, J04AB01), dalbavancin (DAL, J01XA04), daptomycin (DAP, J01XX09), dibekacin (DKB, J01GB09), dirithromycin (DIR, J01FA13), doripenem (DOR, J01DH04), doxycycline (DOX, J01AA02), enoxacin (ENX, J01MA04), epicillin (EPC, J01CA07), ertapenem (ETP, J01DH03), erythromycin (ERY, J01FA01), fleroxacin (FLE, J01MA08), flucloxacillin (FLC, J01CF05), flurithromycin (FLR1, J01FA14), fosfomycin (FOS, J01XX01), fusidic acid (FUS, J01XC01), gatifloxacin (GAT, J01MA16), gemifloxacin (GEM, J01MA15), gentamicin (GEN, J01GB03), grepafloxacin (GRX, J01MA11), hetacillin (HET, J01CA18), imipenem (IPM, J01DH51), isepamicin (ISE, J01GB11), josamycin (JOS, J01FA07), kanamycin (KAN, J01GB04), latamoxef (LTM, J01DD06), levofloxacin (LVX, J01MA12), lincomycin (LIN, J01FF02), linezolid (LNZ, J01XX08), lomefloxacin (LOM, J01MA07), loracarbef (LOR, J01DC08), mecillinam (Amdinocillin)

(MEC, J01CA11), meropenem (MEM, J01DH02), meropenem/vaborbactam (MEV, J01DH52), metampicillin (MTM, J01CA14), mezlocillin (MEZ, J01CA10), midecamycin (MID, J01FA03), minocycline (MNO, J01AA08), miocamycin (MCM, J01FA11), moxifloxacin (MFX, J01MA14), nalidixic acid (NAL, J01MB02), neomycin (NEO, J01GB05), netilmicin (NET, J01GB07), nitrofurantoin (NIT, J01XE01), norfloxacin (NOR, J01MA06), norvancomycin (NVA, no ATC code), novobiocin (NOV, QJ01XX95), ofloxacin (OFX, J01MA01), oleandomycin (OLE, J01FA05), oritavancin (ORI, J01XA05), oxacillin (OXA, J01CF04), pazufloxacin (PAZ, J01MA18), pefloxacin (PEF, J01MA03), phenoxymethylpenicillin (PHN, J01CE02), piperacillin (PIP, J01CA12), piperacillin/tazobactam (TZP, J01CR05), pirlimycin (PRL, no ATC code), pivampicillin (PVM, J01CA02), pivmecillinam (PME, J01CA08), polymyxin B (PLB, J01XB02), pristinamycin (PRI, J01FG01), prulifloxacin (PRU, J01MA17), quinupristin/dalfopristin (QDA, J01FG02), ramoplanin (RAM, no ATC code), ribostamycin (RST, J01GB10), rifampicin (RIF, J04AB02), rokitamycin (ROK, J01FA12), roxithromycin (RXT, J01FA06), rufloxacin (RFL, J01MA10), sisomicin (SIS, J01GB08), sparfloxacin (SPX, J01MA09), spiramycin (SPI, J01FA02), streptoduocin (STR, J01GA02), streptomycin (STR1, J01GA01), sulbenicillin (SBC, J01CA16), sulfadiazine (SDI, J01EC02), sulfadiazine/trimethoprim (SLT1, J01EE02), sulfadimethoxine (SUD, J01ED01), sulfadimidine (SDM, J01EB03), sulfadimidine/trimethoprim (SLT2, J01EE05), sulfafurazole (SLF, J01EB05), sulfaisodimidine (SLF1, J01EB01), sulfalene (SLF2, J01ED02), sulfamazone (SZO, J01ED09), sulfamerazine (SLF3, J01ED07), sulfamerazine/trimethoprim (SLT3, J01EE07), sulfamethizole (SLF4, J01EB02), sulfamethoxazole (SMX, J01EC01), sulfamethoxypyridazine (SLF5, J01ED05), sulfametomidine (SLF6, J01ED03), sulfametoxydiazine (SLF7, J01ED04), sulfametrole/trimethoprim (SLT4, J01EE03), sulfamoxole (SLF8, J01EC03), sulfamoxole/trimethoprim (SLT5, J01EE04), sulfanilamide (SLF9, J01EB06), sulfaperin (SLF10, J01ED06), sulfaphenazole (SLF11, J01ED08), sulfapyridine (SLF12, J01EB04), sulfathiazole (SUT, J01EB07), sulfathiourea (SLF13, J01EB08), talampicillin (TAL, J01CA15), tedizolid (TZD, J01XX11), teicoplanin (TEC, J01XA02), teicoplanin-macromethod (TCM, no ATC code), telavancin (TLV, J01XA03), telithromycin (TLT, J01FA15), temafloxacin (TMX, J01MA05), temocillin (TEM, J01CA17), tetracycline (TCY, J01AA07), thiacetazone (THA, no ATC code), ticarcillin (TIC, J01CA13), ticarcillin/clavulanic acid (TCC, J01CR03), tigecycline (TGC, J01AA12), tobramycin (TOB, J01GB01), trimethoprim (TMP, J01EA01), trimethoprim/sulfamethoxazole (SXT, J01EE01), troleandomycin (TRL, J01FA08), trovafloxacin (TVA, J01MA13), vancomycin (VAN, J01XA01)

##### Stable lifecycle

The lifecycle of this function is **stable**. In a stable function, major changes are unlikely. This means that the unlying code will generally evolve by adding new arguments; removing arguments or changing the meaning of existing arguments will be avoided.

If the unlying code needs breaking changes, they will occur gradually. For example, a argument will be deprecated and first continue to work, but will emit an message informing you of the change. Next, typically after at least one newly released version on CRAN, the message will be transformed to an error.

##### Reference data publicly available

All reference data sets (about microorganisms, antibiotics, R/SI interpretation, EUCAST rules, etc.) in this AMR package are publicly and freely available. We continually export our data sets to formats for use in R, SPSS, SAS, Stata and Excel. We also supply flat files that are machine-readable and suitable for input in any software program, such as laboratory information systems. Please find all download links on our website, which is automatically updated with every code change.

##### Read more on our website!

On our website `https://msberends.github.io/AMR/` you can find a comprehensive tutorial about how to conduct AMR analysis, the complete documentation of all functions and an example analysis using WHONET data. As we would like to better understand the backgrounds and needs of our users, please participate in our survey!

##### Source

- EUCAST Expert Rules. Version 2.0, 2012.
  Leclercq et al. **EUCAST expert rules in antimicrobial susceptibility testing.** *Clin Microbiol Infect.* 2013;19(2):141-60; doi: 10.1111/j.14690691.2011.03703.x
- EUCAST Expert Rules, Intrinsic Resistance and Exceptional Phenotypes Tables. Version 3.1, 2016. (link)
- EUCAST Intrinsic Resistance and Unusual Phenotypes. Version 3.2, 2020. (link)
- EUCAST Breakpoint tables for interpretation of MICs and zone diameters. Version 9.0, 2019. (link)
- EUCAST Breakpoint tables for interpretation of MICs and zone diameters. Version 10.0, 2020. (link)

##### Examples

```
a <- data.frame(mo = c("Staphylococcus aureus",
                       "Enterococcus faecalis",
                       "Escherichia coli",
                       "Klebsiella pneumoniae",
                       "Pseudomonas aeruginosa"),
                VAN = "-",       # Vancomycin
                AMX = "-",       # Amoxicillin
                COL = "-",       # Colistin
                CAZ = "-",       # Ceftazidime
                CXM = "-",       # Cefuroxime
                PEN = "S",       # Benzylpenicillin
                FOX = "S",       # Cefoxitin
                stringsAsFactors = FALSE)

a
#                       mo  VAN  AMX  COL  CAZ  CXM  PEN  FOX
# 1  Staphylococcus aureus    -    -    -    -    -    S    S
# 2  Enterococcus faecalis    -    -    -    -    -    S    S
# 3       Escherichia coli    -    -    -    -    -    S    S
# 4  Klebsiella pneumoniae    -    -    -    -    -    S    S
# 5 Pseudomonas aeruginosa    -    -    -    -    -    S    S

# apply EUCAST rules: some results wil be changed
b <- eucast_rules(a)

b
#                       mo  VAN  AMX  COL  CAZ  CXM  PEN  FOX
# 1  Staphylococcus aureus    -    S    R    R    S    S    S
# 2  Enterococcus faecalis    -    -    R    R    R    S    R
# 3       Escherichia coli    R    -    -    -    -    R    S
# 4  Klebsiella pneumoniae    R    R    -    -    -    R    S
# 5 Pseudomonas aeruginosa    R    R    -    -    R    R    R

# do not apply EUCAST rules, but rather get a data.frame
```

```
# containing all details about the transformations:
c <- eucast_rules(a, verbose = TRUE)
```

---

example_isolates *Data set with 2,000 example isolates*

---

#### Description

A data set containing 2,000 microbial isolates with their full antibiograms. The data set reflects reality and can be used to practice AMR analysis. For examples, please read the tutorial on our website.

#### Usage

```
example_isolates
```

#### Format

A data.frame with 2,000 observations and 49 variables:

- `date`
  date of receipt at the laboratory
- `hospital_id`
  ID of the hospital, from A to D
- `ward_icu`
  logical to determine if ward is an intensive care unit
- `ward_clinical`
  logical to determine if ward is a regular clinical ward
- `ward_outpatient`
  logical to determine if ward is an outpatient clinic
- `age`
  age of the patient
- `gender`
  gender of the patient
- `patient_id`
  ID of the patient
- `mo`
  ID of microorganism created with `as.mo()`, see also microorganisms
- `PEN:RIF`
  40 different antibiotics with class `rsi` (see `as.rsi()`); these column names occur in the antibiotics data set and can be translated with `ab_name()`

#### Reference data publicly available

All reference data sets (about microorganisms, antibiotics, R/SI interpretation, EUCAST rules, etc.) in this `AMR` package are publicly and freely available. We continually export our data sets to formats for use in R, SPSS, SAS, Stata and Excel. We also supply flat files that are machine-readable and suitable for input in any software program, such as laboratory information systems. Please find all download links on our website, which is automatically updated with every code change.

##### Read more on our website!

On our website `https://msberends.github.io/AMR/` you can find a comprehensive tutorial about how to conduct AMR analysis, the complete documentation of all functions and an example analysis using WHONET data. As we would like to better understand the backgrounds and needs of our users, please participate in our survey!

---

`example_isolates_unclean`

*Data set with unclean data*

---

##### Description

A data set containing 3,000 microbial isolates that are not cleaned up and consequently not ready for AMR analysis. This data set can be used for practice.

##### Usage

```
example_isolates_unclean
```

##### Format

A data.frame with 3,000 observations and 8 variables:

- `patient_id`
  ID of the patient
- `date`
  date of receipt at the laboratory
- `hospital`
  ID of the hospital, from A to C
- `bacteria`
  info about microorganism that can be transformed with `as.mo()`, see also microorganisms
- `AMX:GEN`
  4 different antibiotics that have to be transformed with `as.rsi()`

##### Reference data publicly available

All reference data sets (about microorganisms, antibiotics, R/SI interpretation, EUCAST rules, etc.) in this `AMR` package are publicly and freely available. We continually export our data sets to formats for use in R, SPSS, SAS, Stata and Excel. We also supply flat files that are machine-readable and suitable for input in any software program, such as laboratory information systems. Please find all download links on our website, which is automatically updated with every code change.

##### Read more on our website!

On our website `https://msberends.github.io/AMR/` you can find a comprehensive tutorial about how to conduct AMR analysis, the complete documentation of all functions and an example analysis using WHONET data. As we would like to better understand the backgrounds and needs of our users, please participate in our survey!

filter_ab_class *Filter isolates on result in antimicrobial class*

#### Description

Filter isolates on results in specific antimicrobial classes. This makes it easy to filter on isolates that were tested for e.g. any aminoglycoside, or to filter on carbapenem-resistant isolates without the need to specify the drugs.

#### Usage

```
filter_ab_class(x, ab_class, result = NULL, scope = "any", ...)

filter_aminoglycosides(x, result = NULL, scope = "any", ...)

filter_carbapenems(x, result = NULL, scope = "any", ...)

filter_cephalosporins(x, result = NULL, scope = "any", ...)

filter_1st_cephalosporins(x, result = NULL, scope = "any", ...)

filter_2nd_cephalosporins(x, result = NULL, scope = "any", ...)

filter_3rd_cephalosporins(x, result = NULL, scope = "any", ...)

filter_4th_cephalosporins(x, result = NULL, scope = "any", ...)

filter_5th_cephalosporins(x, result = NULL, scope = "any", ...)

filter_fluoroquinolones(x, result = NULL, scope = "any", ...)

filter_glycopeptides(x, result = NULL, scope = "any", ...)

filter_macrolides(x, result = NULL, scope = "any", ...)

filter_penicillins(x, result = NULL, scope = "any", ...)

filter_tetracyclines(x, result = NULL, scope = "any", ...)
```

#### Arguments

| | |
|---|---|
| x | a data set |
| ab_class | an antimicrobial class, like "carbapenems". The columns group, atc_group1 and atc_group2 of the antibiotics data set will be searched (case-insensitive) for this value. |
| result | an antibiotic result: S, I or R (or a combination of more of them) |
| scope | the scope to check which variables to check, can be "any" (default) or "all" |
| ... | previously used when this package still depended on the dplyr package, now ignored |

#### Details

All columns of x will be searched for known antibiotic names, abbreviations, brand names and codes (ATC, EARS-Net, WHO, etc.). This means that a filter function like e.g. `filter_aminoglycosides()` will include column names like 'gen', 'genta', 'J01GB03', 'tobra', 'Tobracin', etc.

#### Stable lifecycle

The lifecycle of this function is **stable**. In a stable function, major changes are unlikely. This means that the unlying code will generally evolve by adding new arguments; removing arguments or changing the meaning of existing arguments will be avoided.

If the unlying code needs breaking changes, they will occur gradually. For example, a argument will be deprecated and first continue to work, but will emit an message informing you of the change. Next, typically after at least one newly released version on CRAN, the message will be transformed to an error.

#### See Also

`antibiotic_class_selectors()` for the `select()` equivalent.

#### Examples

```
filter_aminoglycosides(example_isolates)

if (require("dplyr")) {

  # filter on isolates that have any result for any aminoglycoside
  example_isolates %>% filter_aminoglycosides()
  example_isolates %>% filter_ab_class("aminoglycoside")

  # this is essentially the same as (but without determination of column names):
  example_isolates %>%
    filter_at(.vars = vars(c("GEN", "TOB", "AMK", "KAN")),
              .vars_predicate = any_vars(. %in% c("S", "I", "R")))

  # filter on isolates that show resistance to ANY aminoglycoside
  example_isolates %>% filter_aminoglycosides("R", "any")

  # filter on isolates that show resistance to ALL aminoglycosides
  example_isolates %>% filter_aminoglycosides("R", "all")

  # filter on isolates that show resistance to
  # any aminoglycoside and any fluoroquinolone
  example_isolates %>%
    filter_aminoglycosides("R") %>%
    filter_fluoroquinolones("R")

  # filter on isolates that show resistance to
  # all aminoglycosides and all fluoroquinolones
  example_isolates %>%
    filter_aminoglycosides("R", "all") %>%
    filter_fluoroquinolones("R", "all")

  # with dplyr 1.0.0 and higher (that adds 'across()'), this is equal:
```

```
  # (though the row names on the first are more correct)
  example_isolates %>% filter_carbapenems("R", "all")
  example_isolates %>% filter(across(carbapenems(), ~. == "R"))
}
```

---

first_isolate *Determine first (weighted) isolates*

---

#### Description

Determine first (weighted) isolates of all microorganisms of every patient per episode and (if needed) per specimen type. To determine patient episodes not necessarily based on microorganisms, use `is_new_episode()` that also supports grouping with the `dplyr` package.

#### Usage

```
first_isolate(
  x,
  col_date = NULL,
  col_patient_id = NULL,
  col_mo = NULL,
  col_testcode = NULL,
  col_specimen = NULL,
  col_icu = NULL,
  col_keyantibiotics = NULL,
  episode_days = 365,
  testcodes_exclude = NULL,
  icu_exclude = FALSE,
  specimen_group = NULL,
  type = "keyantibiotics",
  ignore_I = TRUE,
  points_threshold = 2,
  info = interactive(),
  include_unknown = FALSE,
  ...
)

filter_first_isolate(
  x,
  col_date = NULL,
  col_patient_id = NULL,
  col_mo = NULL,
  ...
)

filter_first_weighted_isolate(
  x,
  col_date = NULL,
  col_patient_id = NULL,
  col_mo = NULL,
```

```
  col_keyantibiotics = NULL,
  ...
)
```

#### Arguments

| | |
|---|---|
| x | a data.frame containing isolates. Can be left blank when used inside dplyr verbs, such as filter(), mutate() and summarise(). |
| col_date | column name of the result date (or date that is was received on the lab), defaults to the first column with a date class |
| col_patient_id | column name of the unique IDs of the patients, defaults to the first column that starts with 'patient' or 'patid' (case insensitive) |
| col_mo | column name of the IDs of the microorganisms (see as.mo()), defaults to the first column of class mo. Values will be coerced using as.mo(). |
| col_testcode | column name of the test codes. Use col_testcode = NULL to **not** exclude certain test codes (such as test codes for screening). In that case testcodes_exclude will be ignored. |
| col_specimen | column name of the specimen type or group |
| col_icu | column name of the logicals (TRUE/FALSE) whether a ward or department is an Intensive Care Unit (ICU) |
| col_keyantibiotics | column name of the key antibiotics to determine first *weighted* isolates, see key_antibiotics(). Defaults to the first column that starts with 'key' followed by 'ab' or 'antibiotics' (case insensitive). Use col_keyantibiotics = FALSE to prevent this. |
| episode_days | episode in days after which a genus/species combination will be determined as 'first isolate' again. The default of 365 days is based on the guideline by CLSI, see Source. |
| testcodes_exclude | character vector with test codes that should be excluded (case-insensitive) |
| icu_exclude | logical whether ICU isolates should be excluded (rows with value TRUE in the column set with col_icu) |
| specimen_group | value in the column set with col_specimen to filter on |
| type | type to determine weighed isolates; can be "keyantibiotics" or "points", see Details |
| ignore_I | logical to determine whether antibiotic interpretations with "I" will be ignored when type = "keyantibiotics", see Details |
| points_threshold | points until the comparison of key antibiotics will lead to inclusion of an isolate when type = "points", see Details |
| info | print progress |
| include_unknown | logical to determine whether 'unknown' microorganisms should be included too, i.e. microbial code "UNKNOWN", which defaults to FALSE. For WHONET users, this means that all records with organism code "con" (*contamination*) will be excluded at default. Isolates with a microbial ID of NA will always be excluded as first isolate. |
| ... | arguments passed on to first_isolate() when using filter_first_isolate(), or arguments passed on to key_antibiotics() when using filter_first_weighted_isolate() |

#### Details

These functions are context-aware when used inside dplyr verbs, such as filter(), mutate() and summarise(). This means that then the x argument can be left blank, please see *Examples*.

The first_isolate() function is a wrapper around the is_new_episode() function, but more efficient for data sets containing microorganism codes or names.

All isolates with a microbial ID of NA will be excluded as first isolate.

##### Why this is so important:

To conduct an analysis of antimicrobial resistance, you should only include the first isolate of every patient per episode (Hindler *et al.* 2007). If you would not do this, you could easily get an overestimate or underestimate of the resistance of an antibiotic. Imagine that a patient was admitted with an MRSA and that it was found in 5 different blood cultures the following week. The resistance percentage of oxacillin of all *S. aureus* isolates would be overestimated, because you included this MRSA more than once. It would be selection bias.

##### filter_*() shortcuts:

The functions filter_first_isolate() and filter_first_weighted_isolate() are helper functions to quickly filter on first isolates.

The function filter_first_isolate() is essentially equal to either:

```
x[first_isolate(x, ...), ]

x %>% filter(first_isolate(...))
```

The function filter_first_weighted_isolate() is essentially equal to:

```
x %>%
  mutate(keyab = key_antibiotics(.)) %>%
  mutate(only_weighted_firsts = first_isolate(x,
                                   col_keyantibiotics = "keyab", ...)) %>%
  filter(only_weighted_firsts == TRUE) %>%
  select(-only_weighted_firsts, -keyab)
```

#### Value

A logical vector

#### Key antibiotics

There are two ways to determine whether isolates can be included as first *weighted* isolates which will give generally the same results:

1. Using type = "keyantibiotics" and argument ignore_I

   Any difference from S to R (or vice versa) will (re)select an isolate as a first weighted isolate. With ignore_I = FALSE, also differences from I to S|R (or vice versa) will lead to this. This is a reliable method and 30-35 times faster than method 2. Read more about this in the key_antibiotics() function.

2. Using type = "points" and argument points_threshold

   A difference from I to S|R (or vice versa) means 0.5 points, a difference from S to R (or vice versa) means 1 point. When the sum of points exceeds points_threshold, which default to 2, an isolate will be (re)selected as a first weighted isolate.

##### Stable lifecycle

The lifecycle of this function is **stable**. In a stable function, major changes are unlikely. This means that the unlying code will generally evolve by adding new arguments; removing arguments or changing the meaning of existing arguments will be avoided.

If the unlying code needs breaking changes, they will occur gradually. For example, a argument will be deprecated and first continue to work, but will emit an message informing you of the change. Next, typically after at least one newly released version on CRAN, the message will be transformed to an error.

##### Read more on our website!

On our website `https://msberends.github.io/AMR/` you can find a comprehensive tutorial about how to conduct AMR analysis, the complete documentation of all functions and an example analysis using WHONET data. As we would like to better understand the backgrounds and needs of our users, please participate in our survey!

##### Source

Methodology of this function is strictly based on:

**M39 Analysis and Presentation of Cumulative Antimicrobial Susceptibility Test Data, 4th Edition**, 2014, *Clinical and Laboratory Standards Institute (CLSI)*. `https://clsi.org/standards/products/microbiology/documents/m39/`.

##### See Also

`key_antibiotics()`

##### Examples

```
# `example_isolates` is a dataset available in the AMR package.
# See ?example_isolates.

# basic filtering on first isolates
example_isolates[first_isolate(example_isolates), ]

# filtering based on isolates ----------------------------------------------

if (require("dplyr")) {
  # filter on first isolates:
  example_isolates %>%
    mutate(first_isolate = first_isolate(.)) %>%
    filter(first_isolate == TRUE)

  # short-hand versions:
  example_isolates %>%
    filter(first_isolate())
  example_isolates %>%
    filter_first_isolate()

  example_isolates %>%
    filter_first_weighted_isolate()

  # now let's see if first isolates matter:
  A <- example_isolates %>%
```

```
    group_by(hospital_id) %>%
    summarise(count = n_rsi(GEN),            # gentamicin availability
              resistance = resistance(GEN))  # gentamicin resistance

  B <- example_isolates %>%
    filter_first_weighted_isolate() %>%      # the 1st isolate filter
    group_by(hospital_id) %>%
    summarise(count = n_rsi(GEN),            # gentamicin availability
              resistance = resistance(GEN))  # gentamicin resistance

  # Have a look at A and B.
  # B is more reliable because every isolate is counted only once.
  # Gentamicin resistance in hospital D appears to be 3.7% higher than
  # when you (erroneously) would have used all isolates for analysis.
}
```

---

g.test *G-test for Count Data*

---

#### Description

g.test() performs chi-squared contingency table tests and goodness-of-fit tests, just like chisq.test() but is more reliable (1). A *G*-test can be used to see whether the number of observations in each category fits a theoretical expectation (called a ***G*-test of goodness-of-fit**), or to see whether the proportions of one variable are different for different values of the other variable (called a ***G*-test of independence**).

#### Usage

```
g.test(x, y = NULL, p = rep(1/length(x), length(x)), rescale.p = FALSE)
```

#### Arguments

| | |
|---|---|
| x | a numeric vector or matrix. x and y can also both be factors. |
| y | a numeric vector; ignored if x is a matrix. If x is a factor, y should be a factor of the same length. |
| p | a vector of probabilities of the same length of x. An error is given if any entry of p is negative. |
| rescale.p | a logical scalar; if TRUE then p is rescaled (if necessary) to sum to 1. If rescale.p is FALSE, and p does not sum to 1, an error is given. |

#### Details

If x is a matrix with one row or column, or if x is a vector and y is not given, then a *goodness-of-fit test* is performed (x is treated as a one-dimensional contingency table). The entries of x must be non-negative integers. In this case, the hypothesis tested is whether the population probabilities equal those in p, or are all equal if p is not given.

If x is a matrix with at least two rows and columns, it is taken as a two-dimensional contingency table: the entries of x must be non-negative integers. Otherwise, x and y must be vectors or factors of the same length; cases with missing values are removed, the objects are coerced to factors, and the contingency table is computed from these. Then Pearson's chi-squared test is performed of the null hypothesis that the joint distribution of the cell counts in a 2-dimensional contingency table is the product of the row and column marginals.

The p-value is computed from the asymptotic chi-squared distribution of the test statistic.

In the contingency table case simulation is done by random sampling from the set of all contingency tables with given marginals, and works only if the marginals are strictly positive. Note that this is not the usual sampling situation assumed for a chi-squared test (such as the *G*-test) but rather that for Fisher's exact test.

In the goodness-of-fit case simulation is done by random sampling from the discrete distribution specified by `p`, each sample being of size `n = sum(x)`. This simulation is done in R and may be slow.

**G-test of goodness-of-fit (likelihood ratio test):**

Use the *G*-test of goodness-of-fit when you have one nominal variable with two or more values (such as male and female, or red, pink and white flowers). You compare the observed counts of numbers of observations in each category with the expected counts, which you calculate using some kind of theoretical expectation (such as a 1:1 sex ratio or a 1:2:1 ratio in a genetic cross).

If the expected number of observations in any category is too small, the *G*-test may give inaccurate results, and you should use an exact test instead (`fisher.test()`).

The *G*-test of goodness-of-fit is an alternative to the chi-square test of goodness-of-fit (`chisq.test()`); each of these tests has some advantages and some disadvantages, and the results of the two tests are usually very similar.

**G-test of independence:**

Use the *G*-test of independence when you have two nominal variables, each with two or more possible values. You want to know whether the proportions for one variable are different among values of the other variable.

It is also possible to do a *G*-test of independence with more than two nominal variables. For example, Jackson et al. (2013) also had data for children under 3, so you could do an analysis of old vs. young, thigh vs. arm, and reaction vs. no reaction, all analyzed together.

Fisher's exact test (`fisher.test()`) is an **exact** test, where the *G*-test is still only an **approximation**. For any 2x2 table, Fisher's Exact test may be slower but will still run in seconds, even if the sum of your observations is multiple millions.

The *G*-test of independence is an alternative to the chi-square test of independence (`chisq.test()`), and they will give approximately the same results.

**How the test works:**

Unlike the exact test of goodness-of-fit (`fisher.test()`), the *G*-test does not directly calculate the probability of obtaining the observed results or something more extreme. Instead, like almost all statistical tests, the *G*-test has an intermediate step; it uses the data to calculate a test statistic that measures how far the observed data are from the null expectation. You then use a mathematical relationship, in this case the chi-square distribution, to estimate the probability of obtaining that value of the test statistic.

The *G*-test uses the log of the ratio of two likelihoods as the test statistic, which is why it is also called a likelihood ratio test or log-likelihood ratio test. The formula to calculate a *G*-statistic is:

$G = 2 * sum(x * log(x/E))$

where `E` are the expected values. Since this is chi-square distributed, the p value can be calculated in R with:

```
p <- stats::pchisq(G, df, lower.tail = FALSE)
```

where df are the degrees of freedom.

If there are more than two categories and you want to find out which ones are significantly different from their null expectation, you can use the same method of testing each category vs. the sum of all categories, with the Bonferroni correction. You use *G*-tests for each category, of course.

#### Value

A list with class "htest" containing the following components:

| | |
|---|---|
| statistic | the value the chi-squared test statistic. |
| parameter | the degrees of freedom of the approximate chi-squared distribution of the test statistic, NA if the p-value is computed by Monte Carlo simulation. |
| p.value | the p-value for the test. |
| method | a character string indicating the type of test performed, and whether Monte Carlo simulation or continuity correction was used. |
| data.name | a character string giving the name(s) of the data. |
| observed | the observed counts. |
| expected | the expected counts under the null hypothesis. |
| residuals | the Pearson residuals, (observed -expected) / sqrt(expected). |
| stdres | standardized residuals, (observed -expected) / sqrt(V), where V is the residual cell variance (Agresti, 2007, section 2.4.5 for the case where x is a matrix, n * p * (1 -p) otherwise). |

#### Questioning lifecycle

The lifecycle of this function is **questioning**. This function might be no longer be optimal approach, or is it questionable whether this function should be in this AMR package at all.

#### Read more on our website!

On our website https://msberends.github.io/AMR/ you can find a comprehensive tutorial about how to conduct AMR analysis, the complete documentation of all functions and an example analysis using WHONET data. As we would like to better understand the backgrounds and needs of our users, please participate in our survey!

#### Source

The code for this function is identical to that of chisq.test(), except that:

- The calculation of the statistic was changed to $2 * sum(x * log(x/E))$
- Yates' continuity correction was removed as it does not apply to a *G*-test
- The possibility to simulate p values with simulate.p.value was removed

#### References

1. McDonald, J.H. 2014. **Handbook of Biological Statistics (3rd ed.)**. Sparky House Publishing, Baltimore, Maryland. http://www.biostathandbook.com/gtestgof.html.

#### See Also

chisq.test()

#### Examples

```
# = EXAMPLE 1 =
# Shivrain et al. (2006) crossed clearfield rice (which are resistant
# to the herbicide imazethapyr) with red rice (which are susceptible to
# imazethapyr). They then crossed the hybrid offspring and examined the
# F2 generation, where they found 772 resistant plants, 1611 moderately
# resistant plants, and 737 susceptible plants. If resistance is controlled
# by a single gene with two co-dominant alleles, you would expect a 1:2:1
# ratio.

x <- c(772, 1611, 737)
G <- g.test(x, p = c(1, 2, 1) / 4)
# G$p.value = 0.12574.

# There is no significant difference from a 1:2:1 ratio.
# Meaning: resistance controlled by a single gene with two co-dominant
# alleles, is plausible.

# = EXAMPLE 2 =
# Red crossbills (Loxia curvirostra) have the tip of the upper bill either
# right or left of the lower bill, which helps them extract seeds from pine
# cones. Some have hypothesized that frequency-dependent selection would
# keep the number of right and left-billed birds at a 1:1 ratio. Groth (1992)
# observed 1752 right-billed and 1895 left-billed crossbills.

x <- c(1752, 1895)
g.test(x)
# p = 0.01787343

# There is a significant difference from a 1:1 ratio.
# Meaning: there are significantly more left-billed birds.
```

---

`get_episode` *Determine (new) episodes for patients*

---

#### Description

These functions determine which items in a vector can be considered (the start of) a new episode, based on the argument `episode_days`. This can be used to determine clinical episodes for any epidemiological analysis. The `get_episode()` function returns the index number of the episode per group, while the `is_new_episode()` function returns values `TRUE`/`FALSE` to indicate whether an item in a vector is the start of a new episode.

#### Usage

```
get_episode(x, episode_days, ...)

is_new_episode(x, episode_days, ...)
```

##### Arguments

| | |
|---|---|
| `x` | vector of dates (class `Date` or `POSIXt`) |
| `episode_days` | length of the required episode in days, please see *Details* |
| `...` | arguments passed on to `as.Date()` |

##### Details

Dates are first sorted from old to new. The oldest date will mark the start of the first episode. After this date, the next date will be marked that is at least `episode_days` days later than the start of the first episode. From that second marked date on, the next date will be marked that is at least `episode_days` days later than the start of the second episode which will be the start of the third episode, and so on. Before the vector is being returned, the original order will be restored.

The `first_isolate()` function is a wrapper around the `is_new_episode()` function, but is more efficient for data sets containing microorganism codes or names.

The `dplyr` package is not required for these functions to work, but these functions support variable grouping and work conveniently inside `dplyr` verbs such as `filter()`, `mutate()` and `summarise()`.

##### Value

- `get_episode()`: a double vector
- `is_new_episode()`: a logical vector

##### Stable lifecycle

The lifecycle of this function is **stable**. In a stable function, major changes are unlikely. This means that the unlying code will generally evolve by adding new arguments; removing arguments or changing the meaning of existing arguments will be avoided.

If the unlying code needs breaking changes, they will occur gradually. For example, a argument will be deprecated and first continue to work, but will emit an message informing you of the change. Next, typically after at least one newly released version on CRAN, the message will be transformed to an error.

##### Read more on our website!

On our website `https://msberends.github.io/AMR/` you can find a comprehensive tutorial about how to conduct AMR analysis, the complete documentation of all functions and an example analysis using WHONET data. As we would like to better understand the backgrounds and needs of our users, please participate in our survey!

##### See Also

`first_isolate()`

##### Examples

```
# `example_isolates` is a dataset available in the AMR package.
# See ?example_isolates.

get_episode(example_isolates$date, episode_days = 60)
is_new_episode(example_isolates$date, episode_days = 60)

# filter on results from the third 60-day episode only, using base R
```

```
example_isolates[which(get_episode(example_isolates$date, 60) == 3), ]

if (require("dplyr")) {
  # is_new_episode() can also be used in dplyr verbs to determine patient
  # episodes based on any (combination of) grouping variables:
  example_isolates %>%
    mutate(condition = sample(x = c("A", "B", "C"),
                              size = 2000,
                              replace = TRUE)) %>%
    group_by(condition) %>%
    mutate(new_episode = is_new_episode(date, 365))

  example_isolates %>%
    group_by(hospital_id, patient_id) %>%
    transmute(date,
              patient_id,
              new_index = get_episode(date, 60),
              new_logical = is_new_episode(date, 60))

  example_isolates %>%
    group_by(hospital_id) %>%
    summarise(patients = n_distinct(patient_id),
              n_episodes_365 = sum(is_new_episode(date, episode_days = 365)),
              n_episodes_60  = sum(is_new_episode(date, episode_days = 60)),
              n_episodes_30  = sum(is_new_episode(date, episode_days = 30)))

  # grouping on patients and microorganisms leads to the same results
  # as first_isolate():
  x <- example_isolates %>%
    filter(first_isolate(., include_unknown = TRUE))

  y <- example_isolates %>%
    group_by(patient_id, mo) %>%
    filter(is_new_episode(date, 365))

  identical(x$patient_id, y$patient_id)

  # but is_new_episode() has a lot more flexibility than first_isolate(),
  # since you can now group on anything that seems relevant:
  example_isolates %>%
    group_by(patient_id, mo, hospital_id, ward_icu) %>%
    mutate(flag_episode = is_new_episode(date, 365))
}
```

---

#### ggplot_pca *PCA biplot with* ggplot2

---

##### Description

Produces a ggplot2 variant of a so-called biplot for PCA (principal component analysis), but is more flexible and more appealing than the base R biplot() function.

##### Usage

```
ggplot_pca(
  x,
  choices = 1:2,
  scale = 1,
  pc.biplot = TRUE,
  labels = NULL,
  labels_textsize = 3,
  labels_text_placement = 1.5,
  groups = NULL,
  ellipse = TRUE,
  ellipse_prob = 0.68,
  ellipse_size = 0.5,
  ellipse_alpha = 0.5,
  points_size = 2,
  points_alpha = 0.25,
  arrows = TRUE,
  arrows_colour = "darkblue",
  arrows_size = 0.5,
  arrows_textsize = 3,
  arrows_textangled = TRUE,
  arrows_alpha = 0.75,
  base_textsize = 10,
  ...
)
```

##### Arguments

`x` an object returned by `pca()`, `prcomp()` or `princomp()`

`choices` length 2 vector specifying the components to plot. Only the default is a biplot in the strict sense.

`scale` The variables are scaled by `lambda ^ scale` and the observations are scaled by `lambda ^ (1-scale)` where `lambda` are the singular values as computed by `princomp`. Normally `0 <= scale <= 1`, and a warning will be issued if the specified `scale` is outside this range.

`pc.biplot` If true, use what Gabriel (1971) refers to as a "principal component biplot", with `lambda = 1` and observations scaled up by sqrt(n) and variables scaled down by sqrt(n). Then inner products between variables approximate covariances and distances between observations approximate Mahalanobis distance.

`labels` an optional vector of labels for the observations. If set, the labels will be placed below their respective points. When using the `pca()` function as input for `x`, this will be determined automatically based on the attribute `non_numeric_cols`, see `pca()`.

`labels_textsize`
the size of the text used for the labels

`labels_text_placement`
adjustment factor the placement of the variable names (>=1 means further away from the arrow head)

`groups` an optional vector of groups for the labels, with the same length as `labels`. If set, the points and labels will be coloured according to these groups. When

| | |
|---|---|
| | using the `pca()` function as input for `x`, this will be determined automatically based on the attribute `non_numeric_cols`, see `pca()`. |
| `ellipse` | a logical to indicate whether a normal data ellipse should be drawn for each group (set with `groups`) |
| `ellipse_prob` | statistical size of the ellipse in normal probability |
| `ellipse_size` | the size of the ellipse line |
| `ellipse_alpha` | the alpha (transparency) of the ellipse line |
| `points_size` | the size of the points |
| `points_alpha` | the alpha (transparency) of the points |
| `arrows` | a logical to indicate whether arrows should be drawn |
| `arrows_colour` | the colour of the arrow and their text |
| `arrows_size` | the size (thickness) of the arrow lines |
| `arrows_textsize` | the size of the text at the end of the arrows |
| `arrows_textangled` | a logical whether the text at the end of the arrows should be angled |
| `arrows_alpha` | the alpha (transparency) of the arrows and their text |
| `base_textsize` | the text size for all plot elements except the labels and arrows |
| `...` | Arguments passed on to functions |

#### Details

The colours for labels and points can be changed by adding another scale layer for colour, like `scale_colour_viridis_d()` or `scale_colour_brewer()`.

#### Maturing lifecycle

The lifecycle of this function is **maturing**. The unlying code of a maturing function has been roughed out, but finer details might still change. Since this function needs wider usage and more extensive testing, you are very welcome to suggest changes at our repository or write us an email (see section 'Contact Us').

#### Source

The `ggplot_pca()` function is based on the `ggbiplot()` function from the `ggbiplot` package by Vince Vu, as found on GitHub: `https://github.com/vqv/ggbiplot` (retrieved: 2 March 2020, their latest commit: `7325e88`; 12 February 2015).

As per their GPL-2 licence that demands documentation of code changes, the changes made based on the source code were:

1. Rewritten code to remove the dependency on packages `plyr`, `scales` and `grid`
2. Parametrised more options, like arrow and ellipse settings
3. Hardened all input possibilities by defining the exact type of user input for every argument
4. Added total amount of explained variance as a caption in the plot
5. Cleaned all syntax based on the `lintr` package, fixed grammatical errors and added integrity checks
6. Updated documentation

#### Examples

```
# `example_isolates` is a dataset available in the AMR package.
# See ?example_isolates.

# See ?pca for more info about Principal Component Analysis (PCA).
if (require("dplyr")) {
  pca_model <- example_isolates %>%
    filter(mo_genus(mo) == "Staphylococcus") %>%
    group_by(species = mo_shortname(mo)) %>%
    summarise_if (is.rsi, resistance) %>%
    pca(FLC, AMC, CXM, GEN, TOB, TMP, SXT, CIP, TEC, TCY, ERY)

  # old (base R)
  biplot(pca_model)

  # new
  ggplot_pca(pca_model)

  if (require("ggplot2")) {
    ggplot_pca(pca_model) +
      scale_colour_viridis_d() +
      labs(title = "Title here")
  }
}
```

---

ggplot_rsi — *AMR plots with* ggplot2

---

#### Description

Use these functions to create bar plots for antimicrobial resistance analysis. All functions rely on ggplot2 functions.

#### Usage

```
ggplot_rsi(
  data,
  position = NULL,
  x = "antibiotic",
  fill = "interpretation",
  facet = NULL,
  breaks = seq(0, 1, 0.1),
  limits = NULL,
  translate_ab = "name",
  combine_SI = TRUE,
  combine_IR = FALSE,
  minimum = 30,
  language = get_locale(),
  nrow = NULL,
  colours = c(S = "#61a8ff", SI = "#61a8ff", I = "#61f7ff", IR = "#ff6961", R =
    "#ff6961"),
  datalabels = TRUE,
```

```
    datalabels.size = 2.5,
    datalabels.colour = "grey15",
    title = NULL,
    subtitle = NULL,
    caption = NULL,
    x.title = "Antimicrobial",
    y.title = "Proportion",
    ...
  )

  geom_rsi(
    position = NULL,
    x = c("antibiotic", "interpretation"),
    fill = "interpretation",
    translate_ab = "name",
    minimum = 30,
    language = get_locale(),
    combine_SI = TRUE,
    combine_IR = FALSE,
    ...
  )

  facet_rsi(facet = c("interpretation", "antibiotic"), nrow = NULL)

  scale_y_percent(breaks = seq(0, 1, 0.1), limits = NULL)

  scale_rsi_colours(
    colours = c(S = "#61a8ff", SI = "#61a8ff", I = "#61f7ff", IR = "#ff6961", R =
      "#ff6961")
  )

  theme_rsi()

  labels_rsi_count(
    position = NULL,
    x = "antibiotic",
    translate_ab = "name",
    minimum = 30,
    language = get_locale(),
    combine_SI = TRUE,
    combine_IR = FALSE,
    datalabels.size = 3,
    datalabels.colour = "grey15"
  )
```

#### Arguments

| | |
|---|---|
| `data` | a data.frame with column(s) of class `rsi` (see `as.rsi()`) |
| `position` | position adjustment of bars, either `"fill"`, `"stack"` or `"dodge"` |
| `x` | variable to show on x axis, either `"antibiotic"` (default) or `"interpretation"` or a grouping variable |

| | |
|---|---|
| fill | variable to categorise using the plots legend, either "antibiotic" (default) or "interpretation" or a grouping variable |
| facet | variable to split plots by, either "interpretation" (default) or "antibiotic" or a grouping variable |
| breaks | numeric vector of positions |
| limits | numeric vector of length two providing limits of the scale, use NA to refer to the existing minimum or maximum |
| translate_ab | a column name of the antibiotics data set to translate the antibiotic abbreviations to, using ab_property() |
| combine_SI | a logical to indicate whether all values of S and I must be merged into one, so the output only consists of S+I vs. R (susceptible vs. resistant). This used to be the argument combine_IR, but this now follows the redefinition by EUCAST about the interpretation of I (increased exposure) in 2019, see section 'Interpretation of S, I and R' below. Default is TRUE. |
| combine_IR | a logical to indicate whether all values of I and R must be merged into one, so the output only consists of S vs. I+R (susceptible vs. non-susceptible). This is outdated, see argument combine_SI. |
| minimum | the minimum allowed number of available (tested) isolates. Any isolate count lower than minimum will return NA with a warning. The default number of 30 isolates is advised by the Clinical and Laboratory Standards Institute (CLSI) as best practice, see Source. |
| language | language of the returned text, defaults to system language (see get_locale()) and can also be set with getOption("AMR_locale"). Use language = NULL or language = "" to prevent translation. |
| nrow | (when using facet) number of rows |
| colours | a named vector with colours for the bars. The names must be one or more of: S, SI, I, IR, R or be FALSE to use default ggplot2 colours. |
| datalabels | show datalabels using labels_rsi_count() |
| datalabels.size | size of the datalabels |
| datalabels.colour | colour of the datalabels |
| title | text to show as title of the plot |
| subtitle | text to show as subtitle of the plot |
| caption | text to show as caption of the plot |
| x.title | text to show as x axis description |
| y.title | text to show as y axis description |
| ... | other arguments passed on to geom_rsi() |

#### Details

At default, the names of antibiotics will be shown on the plots using ab_name(). This can be set with the translate_ab argument. See count_df().

**The functions:**

geom_rsi() will take any variable from the data that has an rsi class (created with as.rsi()) using rsi_df() and will plot bars with the percentage R, I and S. The default behaviour is to have the bars stacked and to have the different antibiotics on the x axis.

facet_rsi() creates 2d plots (at default based on S/I/R) using ggplot2::facet_wrap().

scale_y_percent() transforms the y axis to a 0 to 100% range using ggplot2::scale_y_continuous().

scale_rsi_colours() sets colours to the bars: pastel blue for S, pastel turquoise for I and pastel red for R, using ggplot2::scale_fill_manual().

theme_rsi() is a [ggplot2 theme][ggplot2::theme() with minimal distraction.

labels_rsi_count() print datalabels on the bars with percentage and amount of isolates using ggplot2::geom_text().

ggplot_rsi() is a wrapper around all above functions that uses data as first input. This makes it possible to use this function after a pipe (%>%). See Examples.

##### Maturing lifecycle

The lifecycle of this function is **maturing**. The unlying code of a maturing function has been roughed out, but finer details might still change. Since this function needs wider usage and more extensive testing, you are very welcome to suggest changes at our repository or write us an email (see section 'Contact Us').

##### Read more on our website!

On our website https://msberends.github.io/AMR/ you can find a comprehensive tutorial about how to conduct AMR analysis, the complete documentation of all functions and an example analysis using WHONET data. As we would like to better understand the backgrounds and needs of our users, please participate in our survey!

##### Examples

```
if (require("ggplot2") & require("dplyr")) {

  # get antimicrobial results for drugs against a UTI:
  ggplot(example_isolates %>% select(AMX, NIT, FOS, TMP, CIP)) +
    geom_rsi()

  # prettify the plot using some additional functions:
  df <- example_isolates %>% select(AMX, NIT, FOS, TMP, CIP)
  ggplot(df) +
    geom_rsi() +
    scale_y_percent() +
    scale_rsi_colours() +
    labels_rsi_count() +
    theme_rsi()

  # or better yet, simplify this using the wrapper function - a single command:
  example_isolates %>%
    select(AMX, NIT, FOS, TMP, CIP) %>%
    ggplot_rsi()

  # get only proportions and no counts:
  example_isolates %>%
    select(AMX, NIT, FOS, TMP, CIP) %>%
    ggplot_rsi(datalabels = FALSE)

  # add other ggplot2 arguments as you like:
  example_isolates %>%
    select(AMX, NIT, FOS, TMP, CIP) %>%
    ggplot_rsi(width = 0.5,
```

```
                 colour = "black",
                 size = 1,
                 linetype = 2,
                 alpha = 0.25)

    example_isolates %>%
      select(AMX) %>%
      ggplot_rsi(colours = c(SI = "yellow"))

  }

  # resistance of ciprofloxacine per age group
  example_isolates %>%
    mutate(first_isolate = first_isolate(.)) %>%
    filter(first_isolate == TRUE,
           mo == as.mo("E. coli")) %>%
    # age_groups() is also a function in this AMR package:
    group_by(age_group = age_groups(age)) %>%
    select(age_group,
           CIP) %>%
    ggplot_rsi(x = "age_group")

  # for colourblind mode, use divergent colours from the viridis package:
  example_isolates %>%
    select(AMX, NIT, FOS, TMP, CIP) %>%
    ggplot_rsi() +
    scale_fill_viridis_d()
  # a shorter version which also adjusts data label colours:
  example_isolates %>%
    select(AMX, NIT, FOS, TMP, CIP) %>%
    ggplot_rsi(colours = FALSE)

  # it also supports groups (don't forget to use the group var on `x` or `facet`):
  example_isolates %>%
    select(hospital_id, AMX, NIT, FOS, TMP, CIP) %>%
    group_by(hospital_id) %>%
    ggplot_rsi(x = "hospital_id",
               facet = "antibiotic",
               nrow = 1,
               title = "AMR of Anti-UTI Drugs Per Hospital",
               x.title = "Hospital",
               datalabels = FALSE)
```

---

guess_ab_col *Guess antibiotic column*

---

#### Description

This tries to find a column name in a data set based on information from the antibiotics data set. Also supports WHONET abbreviations.

#### Usage

```
guess_ab_col(x = NULL, search_string = NULL, verbose = FALSE)
```

#### Arguments

| | |
|---|---|
| x | a data.frame |
| search_string | a text to search x for, will be checked with as.ab() if this value is not a column in x |
| verbose | a logical to indicate whether additional info should be printed |

#### Details

You can look for an antibiotic (trade) name or abbreviation and it will search x and the antibiotics data set for any column containing a name or code of that antibiotic. **Longer columns names take precedence over shorter column names.**

#### Value

A column name of x, or NULL when no result is found.

#### Stable lifecycle

The lifecycle of this function is **stable**. In a stable function, major changes are unlikely. This means that the unlying code will generally evolve by adding new arguments; removing arguments or changing the meaning of existing arguments will be avoided.

If the unlying code needs breaking changes, they will occur gradually. For example, a argument will be deprecated and first continue to work, but will emit an message informing you of the change. Next, typically after at least one newly released version on CRAN, the message will be transformed to an error.

#### Read more on our website!

On our website https://msberends.github.io/AMR/ you can find a comprehensive tutorial about how to conduct AMR analysis, the complete documentation of all functions and an example analysis using WHONET data. As we would like to better understand the backgrounds and needs of our users, please participate in our survey!

#### Examples

```
df <- data.frame(amox = "S",
                 tetr = "R")

guess_ab_col(df, "amoxicillin")
# [1] "amox"
guess_ab_col(df, "J01AA07") # ATC code of tetracycline
# [1] "tetr"

guess_ab_col(df, "J01AA07", verbose = TRUE)
# NOTE: Using column 'tetr' as input for J01AA07 (tetracycline).
# [1] "tetr"

# WHONET codes
df <- data.frame(AMP_ND10 = "R",
                 AMC_ED20 = "S")
```

```
guess_ab_col(df, "ampicillin")
# [1] "AMP_ND10"
guess_ab_col(df, "J01CR02")
# [1] "AMC_ED20"
guess_ab_col(df, as.ab("augmentin"))
# [1] "AMC_ED20"

# Longer names take precendence:
df <- data.frame(AMP_ED2 = "S",
                 AMP_ED20 = "S")
guess_ab_col(df, "ampicillin")
# [1] "AMP_ED20"
```

---

#### intrinsic_resistant *Data set with bacterial intrinsic resistance*

---

##### Description

Data set containing defined intrinsic resistance by EUCAST of all bug-drug combinations.

##### Usage

```
intrinsic_resistant
```

##### Format

A data.frame with 93,892 observations and 2 variables:

- `microorganism`
  Name of the microorganism
- `antibiotic`
  Name of the antibiotic drug

##### Details

The repository of this AMR package contains a file comprising this exact data set: `https://github.com/msberends/AMR/blob/master/data-raw/intrinsic_resistant.txt`. This file **allows for machine reading EUCAST guidelines about intrinsic resistance**, which is almost impossible with the Excel and PDF files distributed by EUCAST. The file is updated automatically.

This data set is based on 'EUCAST Expert Rules' and 'EUCAST Intrinsic Resistance and Unusual Phenotypes' v3.2 from 2020.

##### Reference data publicly available

All reference data sets (about microorganisms, antibiotics, R/SI interpretation, EUCAST rules, etc.) in this AMR package are publicly and freely available. We continually export our data sets to formats for use in R, SPSS, SAS, Stata and Excel. We also supply flat files that are machine-readable and suitable for input in any software program, such as laboratory information systems. Please find all download links on our website, which is automatically updated with every code change.

##### Read more on our website!

On our website `https://msberends.github.io/AMR/` you can find a comprehensive tutorial about how to conduct AMR analysis, the complete documentation of all functions and an example analysis using WHONET data. As we would like to better understand the backgrounds and needs of our users, please participate in our survey!

##### Examples

```
if (require("dplyr")) {
  intrinsic_resistant %>%
    filter(antibiotic == "Vancomycin", microorganism %like% "Enterococcus") %>%
    pull(microorganism)
  # [1] "Enterococcus casseliflavus" "Enterococcus gallinarum"
}
```

---

`join` *Join microorganisms to a data set*

---

##### Description

Join the data set microorganisms easily to an existing table or character vector.

##### Usage

```
inner_join_microorganisms(x, by = NULL, suffix = c("2", ""), ...)

left_join_microorganisms(x, by = NULL, suffix = c("2", ""), ...)

right_join_microorganisms(x, by = NULL, suffix = c("2", ""), ...)

full_join_microorganisms(x, by = NULL, suffix = c("2", ""), ...)

semi_join_microorganisms(x, by = NULL, ...)

anti_join_microorganisms(x, by = NULL, ...)
```

##### Arguments

| | |
|---|---|
| `x` | existing table to join, or character vector |
| `by` | a variable to join by - if left empty will search for a column with class `mo` (created with `as.mo()`) or will be `"mo"` if that column name exists in `x`, could otherwise be a column name of `x` with values that exist in `microorganisms$mo` (such as `by = "bacteria_id"`), or another column in microorganisms (but then it should be named, like `by = c("bacteria_id" = "fullname")`) |
| `suffix` | if there are non-joined duplicate variables in `x` and `y`, these suffixes will be added to the output to disambiguate them. Should be a character vector of length 2. |
| `...` | ignored |

#### Details

**Note:** As opposed to the `join()` functions of `dplyr`, character vectors are supported and at default existing columns will get a suffix `"2"` and the newly joined columns will not get a suffix.

If the `dplyr` package is installed, their join functions will be used. Otherwise, the much slower `merge()` function from base R will be used.

#### Stable lifecycle

The lifecycle of this function is **stable**. In a stable function, major changes are unlikely. This means that the unlying code will generally evolve by adding new arguments; removing arguments or changing the meaning of existing arguments will be avoided.

If the unlying code needs breaking changes, they will occur gradually. For example, a argument will be deprecated and first continue to work, but will emit an message informing you of the change. Next, typically after at least one newly released version on CRAN, the message will be transformed to an error.

#### Read more on our website!

On our website `https://msberends.github.io/AMR/` you can find a comprehensive tutorial about how to conduct AMR analysis, the complete documentation of all functions and an example analysis using WHONET data. As we would like to better understand the backgrounds and needs of our users, please participate in our survey!

#### Examples

```
left_join_microorganisms(as.mo("K. pneumoniae"))
left_join_microorganisms("B_KLBSL_PNE")

if (require("dplyr")) {
  example_isolates %>%
    left_join_microorganisms() %>%
    colnames()

  df <- data.frame(date = seq(from = as.Date("2018-01-01"),
                              to = as.Date("2018-01-07"),
                              by = 1),
                   bacteria = as.mo(c("S. aureus", "MRSA", "MSSA", "STAAUR",
                                      "E. coli", "E. coli", "E. coli")),
                   stringsAsFactors = FALSE)
  colnames(df)
  df_joined <- left_join_microorganisms(df, "bacteria")
  colnames(df_joined)
}
```

---

`key_antibiotics` *Key antibiotics for first* weighted *isolates*

---

#### Description

These function can be used to determine first isolates (see `first_isolate()`). Using key antibiotics to determine first isolates is more reliable than without key antibiotics. These selected isolates can then be called first *weighted* isolates.

#### Usage

```
key_antibiotics(
  x,
  col_mo = NULL,
  universal_1 = guess_ab_col(x, "amoxicillin"),
  universal_2 = guess_ab_col(x, "amoxicillin/clavulanic acid"),
  universal_3 = guess_ab_col(x, "cefuroxime"),
  universal_4 = guess_ab_col(x, "piperacillin/tazobactam"),
  universal_5 = guess_ab_col(x, "ciprofloxacin"),
  universal_6 = guess_ab_col(x, "trimethoprim/sulfamethoxazole"),
  GramPos_1 = guess_ab_col(x, "vancomycin"),
  GramPos_2 = guess_ab_col(x, "teicoplanin"),
  GramPos_3 = guess_ab_col(x, "tetracycline"),
  GramPos_4 = guess_ab_col(x, "erythromycin"),
  GramPos_5 = guess_ab_col(x, "oxacillin"),
  GramPos_6 = guess_ab_col(x, "rifampin"),
  GramNeg_1 = guess_ab_col(x, "gentamicin"),
  GramNeg_2 = guess_ab_col(x, "tobramycin"),
  GramNeg_3 = guess_ab_col(x, "colistin"),
  GramNeg_4 = guess_ab_col(x, "cefotaxime"),
  GramNeg_5 = guess_ab_col(x, "ceftazidime"),
  GramNeg_6 = guess_ab_col(x, "meropenem"),
  warnings = TRUE,
  ...
)

key_antibiotics_equal(
  y,
  z,
  type = c("keyantibiotics", "points"),
  ignore_I = TRUE,
  points_threshold = 2,
  info = FALSE
)
```

#### Arguments

`x` a data.frame with antibiotics columns, like `AMX` or `amox`. Can be left blank when used inside `dplyr` verbs, such as `filter()`, `mutate()` and `summarise()`.

`col_mo` column name of the IDs of the microorganisms (see `as.mo()`), defaults to the first column of class `mo`. Values will be coerced using `as.mo()`.

`universal_1, universal_2, universal_3, universal_4, universal_5, universal_6`
column names of **broad-spectrum** antibiotics, case-insensitive. See details for which antibiotics will be used at default (which are guessed with `guess_ab_col()`).

GramPos_1, GramPos_2, GramPos_3, GramPos_4, GramPos_5, GramPos_6
: column names of antibiotics for **Gram-positives**, case-insensitive. See details for which antibiotics will be used at default (which are guessed with guess_ab_col()).

GramNeg_1, GramNeg_2, GramNeg_3, GramNeg_4, GramNeg_5, GramNeg_6
: column names of antibiotics for **Gram-negatives**, case-insensitive. See details for which antibiotics will be used at default (which are guessed with guess_ab_col()).

warnings
: give a warning about missing antibiotic columns (they will be ignored)

...
: other arguments passed on to functions

y, z
: character vectors to compare

type
: type to determine weighed isolates; can be "keyantibiotics" or "points", see Details

ignore_I
: logical to determine whether antibiotic interpretations with "I" will be ignored when type = "keyantibiotics", see Details

points_threshold
: points until the comparison of key antibiotics will lead to inclusion of an isolate when type = "points", see Details

info
: print progress

#### Details

The key_antibiotics() function is context-aware when used inside dplyr verbs, such as filter(), mutate() and summarise(). This means that then the x argument can be left blank, please see *Examples*.

The function key_antibiotics() returns a character vector with 12 antibiotic results for every isolate. These isolates can then be compared using key_antibiotics_equal(), to check if two isolates have generally the same antibiogram. Missing and invalid values are replaced with a dot (".") by key_antibiotics() and ignored by key_antibiotics_equal().

The first_isolate() function only uses this function on the same microbial species from the same patient. Using this, e.g. an MRSA will be included after a susceptible *S. aureus* (MSSA) is found within the same patient episode. Without key antibiotic comparison it would not. See first_isolate() for more info.

At default, the antibiotics that are used for **Gram-positive bacteria** are:

- Amoxicillin
- Amoxicillin/clavulanic acid
- Cefuroxime
- Piperacillin/tazobactam
- Ciprofloxacin
- Trimethoprim/sulfamethoxazole
- Vancomycin
- Teicoplanin
- Tetracycline
- Erythromycin
- Oxacillin
- Rifampin

At default the antibiotics that are used for **Gram-negative bacteria** are:

- Amoxicillin
- Amoxicillin/clavulanic acid
- Cefuroxime
- Piperacillin/tazobactam
- Ciprofloxacin
- Trimethoprim/sulfamethoxazole
- Gentamicin
- Tobramycin
- Colistin
- Cefotaxime
- Ceftazidime
- Meropenem

The function `key_antibiotics_equal()` checks the characters returned by `key_antibiotics()` for equality, and returns a `logical` vector.

#### Stable lifecycle

The lifecycle of this function is **stable**. In a stable function, major changes are unlikely. This means that the unlying code will generally evolve by adding new arguments; removing arguments or changing the meaning of existing arguments will be avoided.

If the unlying code needs breaking changes, they will occur gradually. For example, a argument will be deprecated and first continue to work, but will emit an message informing you of the change. Next, typically after at least one newly released version on CRAN, the message will be transformed to an error.

#### Key antibiotics

There are two ways to determine whether isolates can be included as first *weighted* isolates which will give generally the same results:

1. Using `type = "keyantibiotics"` and argument `ignore_I`
   Any difference from S to R (or vice versa) will (re)select an isolate as a first weighted isolate. With `ignore_I = FALSE`, also differences from I to S|R (or vice versa) will lead to this. This is a reliable method and 30-35 times faster than method 2. Read more about this in the `key_antibiotics()` function.
2. Using `type = "points"` and argument `points_threshold`
   A difference from I to S|R (or vice versa) means 0.5 points, a difference from S to R (or vice versa) means 1 point. When the sum of points exceeds `points_threshold`, which default to 2, an isolate will be (re)selected as a first weighted isolate.

#### Read more on our website!

On our website `https://msberends.github.io/AMR/` you can find a comprehensive tutorial about how to conduct AMR analysis, the complete documentation of all functions and an example analysis using WHONET data. As we would like to better understand the backgrounds and needs of our users, please participate in our survey!

#### See Also

`first_isolate()`

#### Examples

```
# `example_isolates` is a dataset available in the AMR package.
# See ?example_isolates.

# output of the `key_antibiotics()` function could be like this:
strainA <- "SSSRR.S.R..S"
strainB <- "SSSIRSSSRSSS"

# those strings can be compared with:
key_antibiotics_equal(strainA, strainB)
# TRUE, because I is ignored (as well as missing values)

key_antibiotics_equal(strainA, strainB, ignore_I = FALSE)
# FALSE, because I is not ignored and so the 4th character differs

if (require("dplyr")) {
  # set key antibiotics to a new variable
  my_patients <- example_isolates %>%
    mutate(keyab = key_antibiotics()) %>% # no need to define `x`
    mutate(
      # now calculate first isolates
      first_regular = first_isolate(col_keyantibiotics = FALSE),
      # and first WEIGHTED isolates
      first_weighted = first_isolate(col_keyantibiotics = "keyab")
    )

  # Check the difference, in this data set it results in a lot more isolates:
  sum(my_patients$first_regular, na.rm = TRUE)
  sum(my_patients$first_weighted, na.rm = TRUE)
}
```

---

### kurtosis *Kurtosis of the sample*

---

#### Description

Kurtosis is a measure of the "tailedness" of the probability distribution of a real-valued random variable. A normal distribution has a kurtosis of 3 and a excess kurtosis of 0.

#### Usage

```
kurtosis(x, na.rm = FALSE, excess = FALSE)

## Default S3 method:
kurtosis(x, na.rm = FALSE, excess = FALSE)

## S3 method for class 'matrix'
kurtosis(x, na.rm = FALSE, excess = FALSE)

## S3 method for class 'data.frame'
kurtosis(x, na.rm = FALSE, excess = FALSE)
```

#### Arguments

| | |
|---|---|
| `x` | a vector of values, a matrix or a data.frame |
| `na.rm` | a logical to indicate whether `NA` values should be stripped before the computation proceeds |
| `excess` | a logical to indicate whether the *excess kurtosis* should be returned, defined as the kurtosis minus 3. |

#### Stable lifecycle

The lifecycle of this function is **stable**. In a stable function, major changes are unlikely. This means that the unlying code will generally evolve by adding new arguments; removing arguments or changing the meaning of existing arguments will be avoided.

If the unlying code needs breaking changes, they will occur gradually. For example, a argument will be deprecated and first continue to work, but will emit an message informing you of the change. Next, typically after at least one newly released version on CRAN, the message will be transformed to an error.

#### Read more on our website!

On our website `https://msberends.github.io/AMR/` you can find a comprehensive tutorial about how to conduct AMR analysis, the complete documentation of all functions and an example analysis using WHONET data. As we would like to better understand the backgrounds and needs of our users, please participate in our survey!

#### See Also

`skewness()`

---

`lifecycle` *Lifecycles of functions in the* `AMR` *package*

---

#### Description

Functions in this `AMR` package are categorised using the lifecycle circle of the Tidyverse as found on www.tidyverse.org/lifecycle.

This page contains a section for every lifecycle (with text borrowed from the aforementioned Tidyverse website), so they can be used in the manual pages of the functions.

#### Experimental lifecycle

The lifecycle of this function is **experimental**. An experimental function is in early stages of development. The unlying code might be changing frequently. Experimental functions might be removed without deprecation, so you are generally best off waiting until a function is more mature before you use it in production code. Experimental functions are only available in development versions of this `AMR` package and will thus not be included in releases that are submitted to CRAN, since such functions have not yet matured enough.

##### Maturing lifecycle

The lifecycle of this function is **maturing**. The unlying code of a maturing function has been roughed out, but finer details might still change. Since this function needs wider usage and more extensive testing, you are very welcome to suggest changes at our repository or write us an email (see section 'Contact Us').

##### Stable lifecycle

The lifecycle of this function is **stable**. In a stable function, major changes are unlikely. This means that the unlying code will generally evolve by adding new arguments; removing arguments or changing the meaning of existing arguments will be avoided.

If the unlying code needs breaking changes, they will occur gradually. For example, a argument will be deprecated and first continue to work, but will emit an message informing you of the change. Next, typically after at least one newly released version on CRAN, the message will be transformed to an error.

##### Retired lifecycle

The lifecycle of this function is **retired**. A retired function is no longer under active development, and (if appropiate) a better alternative is available. No new arguments will be added, and only the most critical bugs will be fixed. In a future version, this function will be removed.

##### Questioning lifecycle

The lifecycle of this function is **questioning**. This function might be no longer be optimal approach, or is it questionable whether this function should be in this `AMR` package at all.

---

#### `like` *Pattern matching with keyboard shortcut*

---

##### Description

Convenient wrapper around `grep()` to match a pattern: `x %like% pattern`. It always returns a `logical` vector and is always case-insensitive (use `x %like_case% pattern` for case-sensitive matching). Also, `pattern` can be as long as `x` to compare items of each index in both vectors, or they both can have the same length to iterate over all cases.

##### Usage

```
like(x, pattern, ignore.case = TRUE)

x %like% pattern

x %like_case% pattern
```

#### Arguments

| | |
|---|---|
| `x` | a character vector where matches are sought, or an object which can be coerced by `as.character()` to a character vector. |
| `pattern` | a character string containing a regular expression (or character string for `fixed = TRUE`) to be matched in the given character vector. Coerced by `as.character()` to a character string if possible. If a character vector of length 2 or more is supplied, the first element is used with a warning. |
| `ignore.case` | if `FALSE`, the pattern matching is *case sensitive* and if `TRUE`, case is ignored during matching. |

#### Details

The %like% function:

- Is case-insensitive (use %like_case% for case-sensitive matching)
- Supports multiple patterns
- Checks if `pattern` is a regular expression and sets `fixed = TRUE` if not, to greatly improve speed
- Tries again with `perl = TRUE` if regex fails

Using RStudio? The text %like% can also be directly inserted in your code from the Addins menu and can have its own Keyboard Shortcut like `Ctrl+Shift+L` or `Cmd+Shift+L` (see `Tools` > Modify Keyboard Shortcuts...).

#### Value

A `logical` vector

#### Stable lifecycle

The lifecycle of this function is **stable**. In a stable function, major changes are unlikely. This means that the unlying code will generally evolve by adding new arguments; removing arguments or changing the meaning of existing arguments will be avoided.

If the unlying code needs breaking changes, they will occur gradually. For example, a argument will be deprecated and first continue to work, but will emit an message informing you of the change. Next, typically after at least one newly released version on CRAN, the message will be transformed to an error.

#### Read more on our website!

On our website `https://msberends.github.io/AMR/` you can find a comprehensive tutorial about how to conduct AMR analysis, the complete documentation of all functions and an example analysis using WHONET data. As we would like to better understand the backgrounds and needs of our users, please participate in our survey!

#### Source

Idea from the `like` function from the `data.table` package

#### See Also

`grep()`

##### Examples

```
# simple test
a <- "This is a test"
b <- "TEST"
a %like% b
#> TRUE
b %like% a
#> FALSE

# also supports multiple patterns, length must be equal to x
a <- c("Test case", "Something different", "Yet another thing")
b <- c(     "case",           "diff",      "yet")
a %like% b
#> TRUE TRUE TRUE

# get isolates whose name start with 'Ent' or 'ent'

if (require("dplyr")) {
  example_isolates %>%
    filter(mo_name(mo) %like% "^ent")
}
```

---

#### mdro — *Determine multidrug-resistant organisms (MDRO)*

---

##### Description

Determine which isolates are multidrug-resistant organisms (MDRO) according to international and national guidelines.

##### Usage

```
mdro(
  x,
  guideline = "CMI2012",
  col_mo = NULL,
  info = interactive(),
  pct_required_classes = 0.5,
  combine_SI = TRUE,
  verbose = FALSE,
  ...
)

brmo(x, guideline = "BRMO", ...)

mrgn(x, guideline = "MRGN", ...)

mdr_tb(x, guideline = "TB", ...)

mdr_cmi2012(x, guideline = "CMI2012", ...)

eucast_exceptional_phenotypes(x, guideline = "EUCAST", ...)
```

#### Arguments

- `x` — a data.frame with antibiotics columns, like `AMX` or `amox`. Can be left blank when used inside `dplyr` verbs, such as `filter()`, `mutate()` and `summarise()`.
- `guideline` — a specific guideline to follow. When left empty, the publication by Magiorakos *et al.* (2012, Clinical Microbiology and Infection) will be followed, please see *Details*.
- `col_mo` — column name of the IDs of the microorganisms (see `as.mo()`), defaults to the first column of class `mo`. Values will be coerced using `as.mo()`.
- `info` — a logical to indicate whether progress should be printed to the console, defaults to only print while in interactive sessions
- `pct_required_classes` — minimal required percentage of antimicrobial classes that must be available per isolate, rounded down. For example, with the default guideline, 17 antimicrobial classes must be available for *S. aureus*. Setting this `pct_required_classes` argument to `0.5` (default) means that for every *S. aureus* isolate at least 8 different classes must be available. Any lower number of available classes will return `NA` for that isolate.
- `combine_SI` — a logical to indicate whether all values of S and I must be merged into one, so resistance is only considered when isolates are R, not I. As this is the default behaviour of the `mdro()` function, it follows the redefinition by EUCAST about the interpretation of I (increased exposure) in 2019, see section 'Interpretation of S, I and R' below. When using `combine_SI = FALSE`, resistance is considered when isolates are R or I.
- `verbose` — a logical to turn Verbose mode on and off (default is off). In Verbose mode, the function does not return the MDRO results, but instead returns a data set in logbook form with extensive info about which isolates would be MDRO-positive, or why they are not.
- `...` — column name of an antibiotic, please see section *Antibiotics* below

#### Details

These functions are context-aware when used inside `dplyr` verbs, such as `filter()`, `mutate()` and `summarise()`. This means that then the `x` argument can be left blank, please see *Examples*.

For the `pct_required_classes` argument, values above 1 will be divided by 100. This is to support both fractions (`0.75` or `3/4`) and percentages (`75`).

Currently supported guidelines are (case-insensitive):

- `guideline = "CMI2012"` (default)

  Magiorakos AP, Srinivasan A *et al.* "Multidrug-resistant, extensively drug-resistant and pandrug-resistant bacteria: an international expert proposal for interim standard definitions for acquired resistance." Clinical Microbiology and Infection (2012) (link)
- `guideline = "EUCAST3.2"` (or simply `guideline = "EUCAST"`)

  The European international guideline - EUCAST Expert Rules Version 3.2 "Intrinsic Resistance and Unusual Phenotypes" (link)
- `guideline = "EUCAST3.1"`

  The European international guideline - EUCAST Expert Rules Version 3.1 "Intrinsic Resistance and Exceptional Phenotypes Tables" (link)

- `guideline = "TB"`
  The international guideline for multi-drug resistant tuberculosis - World Health Organization "Companion handbook to the WHO guidelines for the programmatic management of drug-resistant tuberculosis" (link)
- `guideline = "MRGN"`
  The German national guideline - Mueller et al. (2015) Antimicrobial Resistance and Infection Control 4:7; doi: 10.1186/s1375601500476
- `guideline = "BRMO"`
  The Dutch national guideline - Rijksinstituut voor Volksgezondheid en Milieu "WIP-richtlijn BRMO (Bijzonder Resistente Micro-Organismen) (ZKH)" (link)

Please suggest your own (country-specific) guidelines by letting us know: `https://github.com/msberends/AMR/issues/new`.

**Note:** Every test that involves the Enterobacteriaceae family, will internally be performed using its newly named *order* Enterobacterales, since the Enterobacteriaceae family has been taxonomically reclassified by Adeolu *et al.* in 2016. Before that, Enterobacteriaceae was the only family under the Enterobacteriales (with an i) order. All species under the old Enterobacteriaceae family are still under the new Enterobacterales (without an i) order, but divided into multiple families. The way tests are performed now by this `mdro()` function makes sure that results from before 2016 and after 2016 are identical.

#### Value

- CMI 2012 paper - function `mdr_cmi2012()` or `mdro()`:
  Ordered factor with levels `Negative` < `Multi-drug-resistant (MDR)` < Extensively drug-resistant (XDR) < `Pandrug-resistant (PDR)`
- TB guideline - function `mdr_tb()` or `mdro(..., guideline = "TB")`:
  Ordered factor with levels `Negative` < `Mono-resistant` < `Poly-resistant` < `Multi-drug-resistant` < Extensively drug-resistant
- German guideline - function `mrgn()` or `mdro(..., guideline = "MRGN")`:
  Ordered factor with levels `Negative` < 3MRGN < 4MRGN
- Everything else:
  Ordered factor with levels `Negative` < Positive, unconfirmed < `Positive`. The value `"Positive, unconfirmed"` means that, according to the guideline, it is not entirely sure if the isolate is multi-drug resistant and this should be confirmed with additional (e.g. molecular) tests

#### Stable lifecycle

The lifecycle of this function is **stable**. In a stable function, major changes are unlikely. This means that the unlying code will generally evolve by adding new arguments; removing arguments or changing the meaning of existing arguments will be avoided.

If the unlying code needs breaking changes, they will occur gradually. For example, a argument will be deprecated and first continue to work, but will emit an message informing you of the change. Next, typically after at least one newly released version on CRAN, the message will be transformed to an error.

#### Antibiotics

To define antibiotics column names, leave as it is to determine it automatically with `guess_ab_col()` or input a text (case-insensitive), or use `NULL` to skip a column (e.g. `TIC = NULL` to skip ticarcillin). Manually defined but non-existing columns will be skipped with a warning.

The following antibiotics are used for the functions `eucast_rules()` and `mdro()`. These are shown below in the format 'name (antimicrobial ID, ATC code)', sorted alphabetically:

Amikacin (AMK, J01GB06), amoxicillin (AMX, J01CA04), amoxicillin/clavulanic acid (AMC, J01CR02), ampicillin (AMP, J01CA01), ampicillin/sulbactam (SAM, J01CR01), avoparcin (AVO, no ATC code), azithromycin (AZM, J01FA10), azlocillin (AZL, J01CA09), aztreonam (ATM, J01DF01), bacampicillin (BAM, J01CA06), benzylpenicillin (PEN, J01CE01), cadazolid (CDZ, J01DD09), carbenicillin (CRB, J01CA03), carindacillin (CRN, J01CA05), cefacetrile (CAC, J01DB10), cefaclor (CEC, J01DC04), cefadroxil (CFR, J01DB05), cefaloridine (RID, J01DB02), cefamandole (MAN, J01DC03), cefatrizine (CTZ, J01DB07), cefazedone (CZD, J01DB06), cefazolin (CZO, J01DB04), cefcapene (CCP, no ATC code), cefcapene pivoxil (CCX, no ATC code), cefdinir (CDR, J01DD15), cefditoren (DIT, J01DD16), cefditoren pivoxil (DIX, no ATC code), cefepime (FEP, J01DE01), cefetamet (CAT, J01DD10), cefetamet pivoxil (CPI, no ATC code), cefixime (CFM, J01DD08), cefmenoxime (CMX, J01DD05), cefmetazole (CMZ, J01DC09), cefodizime (DIZ, J01DD09), cefonicid (CID, J01DC06), cefoperazone (CFP, J01DD12), cefoperazone/sulbactam (CSL, J01DD62), ceforanide (CND, J01DC11), cefotaxime (CTX, J01DD01), cefotaxime/clavulanic acid (CTC, no ATC code), cefotaxime/sulbactam (CTS, no ATC code), cefotetan (CTT, J01DC05), cefotiam (CTF, J01DC07), cefotiam hexetil (CHE, no ATC code), cefovecin (FOV, no ATC code), cefoxitin (FOX, J01DC01), cefoxitin screening (FOX1, no ATC code), cefpimizole (CFZ, no ATC code), cefpiramide (CPM, J01DD11), cefpirome (CPO, J01DE02), cefpodoxime (CPD, J01DD13), cefpodoxime proxetil (CPX, no ATC code), cefpodoxime/clavulanic acid (CDC, no ATC code), cefprozil (CPR, J01DC10), cefroxadine (CRD, J01DB11), cefsulodin (CFS, J01DD03), ceftaroline (CPT, J01DI02), ceftazidime (CAZ, J01DD02), ceftazidime/avibactam (CZA, no ATC code), ceftazidime/clavulanic acid (CCV, J01DD52), cefteram (CEM, no ATC code), cefteram pivoxil (CPL, no ATC code), ceftezole (CTL, J01DB12), ceftibuten (CTB, J01DD14), ceftiofur (TIO, no ATC code), ceftizoxime (CZX, J01DD07), ceftizoxime alapivoxil (CZP, no ATC code), ceftobiprole (BPR, J01DI01), ceftobiprole medocaril (CFM1, J01DI01), ceftolozane/enzyme inhibitor (CEI, J01DI54), ceftriaxone (CRO, J01DD04), cefuroxime (CXM, J01DC02), cephalexin (LEX, J01DB01), cephalothin (CEP, J01DB03), cephapirin (HAP, J01DB08), cephradine (CED, J01DB09), chloramphenicol (CHL, J01BA01), ciprofloxacin (CIP, J01MA02), clarithromycin (CLR, J01FA09), clindamycin (CLI, J01FF01), colistin (COL, J01XB01), cycloserine (CYC, J04AB01), dalbavancin (DAL, J01XA04), daptomycin (DAP, J01XX09), dibekacin (DKB, J01GB09), dirithromycin (DIR, J01FA13), doripenem (DOR, J01DH04), doxycycline (DOX, J01AA02), enoxacin (ENX, J01MA04), epicillin (EPC, J01CA07), ertapenem (ETP, J01DH03), erythromycin (ERY, J01FA01), fleroxacin (FLE, J01MA08), flucloxacillin (FLC, J01CF05), flurithromycin (FLR1, J01FA14), fosfomycin (FOS, J01XX01), fusidic acid (FUS, J01XC01), gatifloxacin (GAT, J01MA16), gemifloxacin (GEM, J01MA15), gentamicin (GEN, J01GB03), grepafloxacin (GRX, J01MA11), hetacillin (HET, J01CA18), imipenem (IPM, J01DH51), isepamicin (ISE, J01GB11), josamycin (JOS, J01FA07), kanamycin (KAN, J01GB04), latamoxef (LTM, J01DD06), levofloxacin (LVX, J01MA12), lincomycin (LIN, J01FF02), linezolid (LNZ, J01XX08), lomefloxacin (LOM, J01MA07), loracarbef (LOR, J01DC08), mecillinam (Amdinocillin) (MEC, J01CA11), meropenem (MEM, J01DH02), meropenem/vaborbactam (MEV, J01DH52), metampicillin (MTM, J01CA14), mezlocillin (MEZ, J01CA10), midecamycin (MID, J01FA03), minocycline (MNO, J01AA08), miocamycin (MCM, J01FA11), moxifloxacin (MFX, J01MA14), nalidixic acid (NAL, J01MB02), neomycin (NEO, J01GB05), netilmicin (NET, J01GB07), nitrofurantoin (NIT, J01XE01), norfloxacin (NOR, J01MA06), norvancomycin (NVA, no ATC code), novobiocin (NOV, QJ01XX95), ofloxacin (OFX, J01MA01), oleandomycin (OLE, J01FA05), oritavancin (ORI, J01XA05), oxacillin (OXA, J01CF04), pazufloxacin (PAZ, J01MA18), pefloxacin (PEF, J01MA03), phenoxymethylpenicillin (PHN, J01CE02), piperacillin (PIP, J01CA12), piperacillin/tazobactam (TZP, J01CR05), pirlimycin (PRL, no ATC code), pivampicillin (PVM, J01CA02), pivmecillinam (PME, J01CA08), polymyxin B (PLB, J01XB02), pristinamycin (PRI, J01FG01), prulifloxacin (PRU, J01MA17), quinupristin/dalfopristin (QDA, J01FG02), ramoplanin (RAM, no ATC code), ribostamycin (RST, J01GB10), rifampicin (RIF, J04AB02), rokitamycin (ROK, J01FA12), roxithromycin (RXT, J01FA06), rufloxacin (RFL, J01MA10), sisomicin (SIS, J01GB08), sparfloxacin (SPX, J01MA09), spiramycin (SPI, J01FA02), streptoduocin (STR, J01GA02), streptomycin (STR1, J01GA01), sulbenicillin (SBC, J01CA16), sulfadiazine

(SDI, J01EC02), sulfadiazine/trimethoprim (SLT1, J01EE02), sulfadimethoxine (SUD, J01ED01), sulfadimidine (SDM, J01EB03), sulfadimidine/trimethoprim (SLT2, J01EE05), sulfafurazole (SLF, J01EB05), sulfaisodimidine (SLF1, J01EB01), sulfalene (SLF2, J01ED02), sulfamazone (SZO, J01ED09), sulfamerazine (SLF3, J01ED07), sulfamerazine/trimethoprim (SLT3, J01EE07), sulfamethizole (SLF4, J01EB02), sulfamethoxazole (SMX, J01EC01), sulfamethoxypyridazine (SLF5, J01ED05), sulfametomidine (SLF6, J01ED03), sulfametoxydiazine (SLF7, J01ED04), sulfametrole/trimethoprim (SLT4, J01EE03), sulfamoxole (SLF8, J01EC03), sulfamoxole/trimethoprim (SLT5, J01EE04), sulfanilamide (SLF9, J01EB06), sulfaperin (SLF10, J01ED06), sulfaphenazole (SLF11, J01ED08), sulfapyridine (SLF12, J01EB04), sulfathiazole (SUT, J01EB07), sulfathiourea (SLF13, J01EB08), talampicillin (TAL, J01CA15), tedizolid (TZD, J01XX11), teicoplanin (TEC, J01XA02), teicoplanin-macromethod (TCM, no ATC code), telavancin (TLV, J01XA03), telithromycin (TLT, J01FA15), temafloxacin (TMX, J01MA05), temocillin (TEM, J01CA17), tetracycline (TCY, J01AA07), thiacetazone (THA, no ATC code), ticarcillin (TIC, J01CA13), ticarcillin/clavulanic acid (TCC, J01CR03), tigecycline (TGC, J01AA12), tobramycin (TOB, J01GB01), trimethoprim (TMP, J01EA01), trimethoprim/sulfamethoxazole (SXT, J01EE01), troleandomycin (TRL, J01FA08), trovafloxacin (TVA, J01MA13), vancomycin (VAN, J01XA01)

##### Interpretation of R and S/I

In 2019, the European Committee on Antimicrobial Susceptibility Testing (EUCAST) has decided to change the definitions of susceptibility testing categories R and S/I as shown below (https://www.eucast.org/newsiandr/).

- **R = Resistant**
  A microorganism is categorised as *Resistant* when there is a high likelihood of therapeutic failure even when there is increased exposure. Exposure is a function of how the mode of administration, dose, dosing interval, infusion time, as well as distribution and excretion of the antimicrobial agent will influence the infecting organism at the site of infection.
- **S = Susceptible**
  A microorganism is categorised as *Susceptible, standard dosing regimen*, when there is a high likelihood of therapeutic success using a standard dosing regimen of the agent.
- **I = Increased exposure, but still susceptible**
  A microorganism is categorised as *Susceptible, Increased exposure* when there is a high likelihood of therapeutic success because exposure to the agent is increased by adjusting the dosing regimen or by its concentration at the site of infection.

This AMR package honours this new insight. Use `susceptibility()` (equal to `proportion_SI()`) to determine antimicrobial susceptibility and `count_susceptible()` (equal to `count_SI()`) to count susceptible isolates.

##### Read more on our website!

On our website https://msberends.github.io/AMR/ you can find a comprehensive tutorial about how to conduct AMR analysis, the complete documentation of all functions and an example analysis using WHONET data. As we would like to better understand the backgrounds and needs of our users, please participate in our survey!

##### Source

Please see *Details* for the list of publications used for this function.

#### Examples

```
mdro(example_isolates, guideline = "EUCAST")

if (require("dplyr")) {
  example_isolates %>%
    mdro() %>%
    table()

  # no need to define `x` when used inside dplyr verbs:
  example_isolates %>%
    mutate(MDRO = mdro(),
           EUCAST = eucast_exceptional_phenotypes(),
           BRMO = brmo(),
           MRGN = mrgn())
}
```

---

`microorganisms` *Data set with 67,151 microorganisms*

---

#### Description

A data set containing the microbial taxonomy of six kingdoms from the Catalogue of Life. MO codes can be looked up using `as.mo()`.

#### Usage

```
microorganisms
```

#### Format

A data.frame with 67,151 observations and 16 variables:

- `mo`
  ID of microorganism as used by this package
- `fullname`
  Full name, like `"Escherichia coli"`
- `kingdom`, `phylum`, `class`, `order`, `family`, `genus`, `species`, `subspecies`
  Taxonomic rank of the microorganism
- `rank`
  Text of the taxonomic rank of the microorganism, like `"species"` or `"genus"`
- `ref`
  Author(s) and year of concerning scientific publication
- `species_id`
  ID of the species as used by the Catalogue of Life
- `source`
  Either "CoL", "DSMZ" (see Source) or "manually added"
- `prevalence`
  Prevalence of the microorganism, see `as.mo()`
- `snomed`
  SNOMED code of the microorganism. Use `mo_snomed()` to retrieve it quickly, see `mo_property()`.

#### Details

Manually added were:

- 11 entries of *Streptococcus* (beta-haemolytic: groups A, B, C, D, F, G, H, K and unspecified; other: viridans, milleri)
- 2 entries of *Staphylococcus* (coagulase-negative (CoNS) and coagulase-positive (CoPS))
- 3 entries of *Trichomonas* (*Trichomonas vaginalis*, and its family and genus)
- 1 entry of *Candida* (*Candida krusei*), that is not (yet) in the Catalogue of Life
- 1 entry of *Blastocystis* (*Blastocystis hominis*), although it officially does not exist (Noel *et al.* 2005, PMID 15634993)
- 5 other 'undefined' entries (unknown, unknown Gram negatives, unknown Gram positives, unknown yeast and unknown fungus)
- 6 families under the Enterobacterales order, according to Adeolu *et al.* (2016, PMID 27620848), that are not (yet) in the Catalogue of Life
- 7,411 species from the DSMZ (Deutsche Sammlung von Mikroorganismen und Zellkulturen) since the DSMZ contain the latest taxonomic information based on recent publications

**Direct download:**

This data set is available as 'flat file' for use even without R - you can find the file here:

- `https://github.com/msberends/AMR/raw/master/data-raw/microorganisms.txt`

The file in R format (with preserved data structure) can be found here:

- `https://github.com/msberends/AMR/raw/master/data/microorganisms.rda`

#### About the records from DSMZ (see source)

Names of prokaryotes are defined as being validly published by the International Code of Nomenclature of Bacteria. Validly published are all names which are included in the Approved Lists of Bacterial Names and the names subsequently published in the International Journal of Systematic Bacteriology (IJSB) and, from January 2000, in the International Journal of Systematic and Evolutionary Microbiology (IJSEM) as original articles or in the validation lists. *(from* `https://www.dsmz.de/services/online-tools/prokaryotic-nomenclature-up-to-date`*)*

In February 2020, the DSMZ records were merged with the List of Prokaryotic names with Standing in Nomenclature (LPSN).

#### Catalogue of Life

This package contains the complete taxonomic tree of almost all microorganisms (~70,000 species) from the authoritative and comprehensive Catalogue of Life (CoL, `http://www.catalogueoflife.org`). The CoL is the most comprehensive and authoritative global index of species currently available. Nonetheless, we supplemented the CoL data with data from the List of Prokaryotic names with Standing in Nomenclature (LPSN, lpsn.dsmz.de). This supplementation is needed until the CoL+ project is finished, which we await.

Click here for more information about the included taxa. Check which versions of the CoL and LSPN were included in this package with `catalogue_of_life_version()`.

##### Reference data publicly available

All reference data sets (about microorganisms, antibiotics, R/SI interpretation, EUCAST rules, etc.) in this AMR package are publicly and freely available. We continually export our data sets to formats for use in R, SPSS, SAS, Stata and Excel. We also supply flat files that are machine-readable and suitable for input in any software program, such as laboratory information systems. Please find all download links on our website, which is automatically updated with every code change.

##### Read more on our website!

On our website `https://msberends.github.io/AMR/` you can find a comprehensive tutorial about how to conduct AMR analysis, the complete documentation of all functions and an example analysis using WHONET data. As we would like to better understand the backgrounds and needs of our users, please participate in our survey!

##### Source

Catalogue of Life: Annual Checklist (public online taxonomic database), `http://www.catalogueoflife.org` (check included annual version with `catalogue_of_life_version()`).

Parte, A.C. (2018). LPSN — List of Prokaryotic names with Standing in Nomenclature (bacterio.net), 20 years on. International Journal of Systematic and Evolutionary Microbiology, 68, 1825-1829; doi: 10.1099/ijsem.0.002786

Leibniz Institute DSMZ-German Collection of Microorganisms and Cell Cultures, Germany, Prokaryotic Nomenclature Up-to-Date, `https://www.dsmz.de/services/online-tools/prokaryotic-nomenclature-up-`[illegible] and `https://lpsn.dsmz.de` (check included version with `catalogue_of_life_version()`).

##### See Also

`as.mo()`, `mo_property()`, microorganisms.codes, intrinsic_resistant

---

`microorganisms.codes` *Data set with 5,583 common microorganism codes*

---

##### Description

A data set containing commonly used codes for microorganisms, from laboratory systems and WHONET. Define your own with `set_mo_source()`. They will all be searched when using `as.mo()` and consequently all the `mo_*` functions.

##### Usage

```
microorganisms.codes
```

##### Format

A data.frame with 5,583 observations and 2 variables:

- `code`
  Commonly used code of a microorganism
- `mo`
  ID of the microorganism in the microorganisms data set

##### Reference data publicly available

All reference data sets (about microorganisms, antibiotics, R/SI interpretation, EUCAST rules, etc.) in this `AMR` package are publicly and freely available. We continually export our data sets to formats for use in R, SPSS, SAS, Stata and Excel. We also supply flat files that are machine-readable and suitable for input in any software program, such as laboratory information systems. Please find all download links on our website, which is automatically updated with every code change.

##### Catalogue of Life

This package contains the complete taxonomic tree of almost all microorganisms (~70,000 species) from the authoritative and comprehensive Catalogue of Life (CoL, `http://www.catalogueoflife.org`). The CoL is the most comprehensive and authoritative global index of species currently available. Nonetheless, we supplemented the CoL data with data from the List of Prokaryotic names with Standing in Nomenclature (LPSN, lpsn.dsmz.de). This supplementation is needed until the CoL+ project is finished, which we await.

Click here for more information about the included taxa. Check which versions of the CoL and LSPN were included in this package with `catalogue_of_life_version()`.

##### Read more on our website!

On our website `https://msberends.github.io/AMR/` you can find a comprehensive tutorial about how to conduct AMR analysis, the complete documentation of all functions and an example analysis using WHONET data. As we would like to better understand the backgrounds and needs of our users, please participate in our survey!

##### See Also

`as.mo()` microorganisms

---

#### `microorganisms.old` *Data set with previously accepted taxonomic names*

---

##### Description

A data set containing old (previously valid or accepted) taxonomic names according to the Catalogue of Life. This data set is used internally by `as.mo()`.

##### Usage

```
microorganisms.old
```

##### Format

A data.frame with 12,708 observations and 4 variables:

- `fullname`
  Old full taxonomic name of the microorganism
- `fullname_new`
  New full taxonomic name of the microorganism
- `ref`
  Author(s) and year of concerning scientific publication
- `prevalence`
  Prevalence of the microorganism, see `as.mo()`

##### Catalogue of Life

This package contains the complete taxonomic tree of almost all microorganisms (~70,000 species) from the authoritative and comprehensive Catalogue of Life (CoL, http://www.catalogueoflife.org). The CoL is the most comprehensive and authoritative global index of species currently available. Nonetheless, we supplemented the CoL data with data from the List of Prokaryotic names with Standing in Nomenclature (LPSN, lpsn.dsmz.de). This supplementation is needed until the CoL+ project is finished, which we await.

Click here for more information about the included taxa. Check which versions of the CoL and LSPN were included in this package with `catalogue_of_life_version()`.

##### Reference data publicly available

All reference data sets (about microorganisms, antibiotics, R/SI interpretation, EUCAST rules, etc.) in this `AMR` package are publicly and freely available. We continually export our data sets to formats for use in R, SPSS, SAS, Stata and Excel. We also supply flat files that are machine-readable and suitable for input in any software program, such as laboratory information systems. Please find all download links on our website, which is automatically updated with every code change.

##### Read more on our website!

On our website https://msberends.github.io/AMR/ you can find a comprehensive tutorial about how to conduct AMR analysis, the complete documentation of all functions and an example analysis using WHONET data. As we would like to better understand the backgrounds and needs of our users, please participate in our survey!

##### Source

Catalogue of Life: Annual Checklist (public online taxonomic database), http://www.catalogueoflife.org (check included annual version with `catalogue_of_life_version()`).

Parte, A.C. (2018). LPSN — List of Prokaryotic names with Standing in Nomenclature (bacterio.net), 20 years on. International Journal of Systematic and Evolutionary Microbiology, 68, 1825-1829; doi: 10.1099/ijsem.0.002786

##### See Also

`as.mo()` `mo_property()` microorganisms

---

#### `mo_matching_score` *Calculate the matching score for microorganisms*

---

##### Description

This algorithm is used by `as.mo()` and all the `mo_*` functions to determine the most probable match of taxonomic records based on user input.

##### Usage

```
mo_matching_score(x, n)
```

##### Arguments

x Any user input value(s)

n A full taxonomic name, that exists in microorganisms$fullname

##### Matching score for microorganisms

With ambiguous user input in as.mo() and all the mo_* functions, the returned results are chosen based on their matching score using mo_matching_score(). This matching score $m$, is calculated as:

$$m_{(x,n)} = \frac{l_n - 0.5 \cdot \min \{ \, l_n \, \mathrm{lev}(x,n) }{l_n \cdot p_n \cdot k_n}$$

where:

- $x$ is the user input;
- $n$ is a taxonomic name (genus, species, and subspecies);
- $l_n$ is the length of $n$;
- lev is the Levenshtein distance function, which counts any insertion, deletion and substitution as 1 that is needed to change $x$ into $n$;
- $p_n$ is the human pathogenic prevalence group of $n$, as described below;
- $k_n$ is the taxonomic kingdom of $n$, set as Bacteria = 1, Fungi = 2, Protozoa = 3, Archaea = 4, others = 5.

The grouping into human pathogenic prevalence ($p$) is based on experience from several microbiological laboratories in the Netherlands in conjunction with international reports on pathogen prevalence. **Group 1** (most prevalent microorganisms) consists of all microorganisms where the taxonomic class is Gammaproteobacteria or where the taxonomic genus is *Enterococcus*, *Staphylococcus* or *Streptococcus*. This group consequently contains all common Gram-negative bacteria, such as *Pseudomonas* and *Legionella* and all species within the order Enterobacterales. **Group 2** consists of all microorganisms where the taxonomic phylum is Proteobacteria, Firmicutes, Actinobacteria or Sarcomastigophora, or where the taxonomic genus is *Absidia*, *Acremonium*, *Actinotignum*, *Alternaria*, *Anaerosalibacter*, *Apophysomyces*, *Arachnia*, *Aspergillus*, *Aureobacterium*, *Aureobasidium*, *Bacteroides*, *Basidiobolus*, *Beauveria*, *Blastocystis*, *Branhamella*, *Calymmatobacterium*, *Candida*, *Capnocytophaga*, *Catabacter*, *Chaetomium*, *Chryseobacterium*, *Chryseomonas*, *Chrysonilia*, *Cladophialophora*, *Cladosporium*, *Conidiobolus*, *Cryptococcus*, *Curvularia*, *Exophiala*, *Exserohilum*, *Flavobacterium*, *Fonsecaea*, *Fusarium*, *Fusobacterium*, *Hendersonula*, *Hypomyces*, *Koserella*, *Lelliottia*, *Leptosphaeria*, *Leptotrichia*, *Malassezia*, *Malbranchea*, *Mortierella*, *Mucor*, *Mycocentrospora*, *Mycoplasma*, *Nectria*, *Ochroconis*, *Oidiodendron*, *Phoma*, *Piedraia*, *Pithomyces*, *Pityrosporum*, *Prevotella*,*Pseudallescheria*, *Rhizomucor*, *Rhizopus*, *Rhodotorula*, *Scolecobasidium*, *Scopulariopsis*, *Scytalidium*,*Sporobolomyces*, *Stachybotrys*, *Stomatococcus*, *Treponema*, *Trichoderma*, *Trichophyton*, *Trichosporon*, *Tritirachium* or *Ureaplasma*. **Group 3** consists of all other microorganisms.

All matches are sorted descending on their matching score and for all user input values, the top match will be returned. This will lead to the effect that e.g., "E. coli" will return the microbial ID of *Escherichia coli* ($m = 0.688$, a highly prevalent microorganism found in humans) and not *Entamoeba coli* ($m = 0.079$, a less prevalent microorganism in humans), although the latter would alphabetically come first.

##### Stable lifecycle

The lifecycle of this function is **stable**. In a stable function, major changes are unlikely. This means that the unlying code will generally evolve by adding new arguments; removing arguments or changing the meaning of existing arguments will be avoided.

If the unlying code needs breaking changes, they will occur gradually. For example, a argument will be deprecated and first continue to work, but will emit an message informing you of the change. Next, typically after at least one newly released version on CRAN, the message will be transformed to an error.

##### Author(s)

Matthijs S. Berends

##### Examples

```
as.mo("E. coli")
mo_uncertainties()

mo_matching_score(x = "E. coli",
                  n = c("Escherichia coli", "Entamoeba coli"))
```

---

## mo_property    *Get properties of a microorganism*

---

##### Description

Use these functions to return a specific property of a microorganism based on the latest accepted taxonomy. All input values will be evaluated internally with `as.mo()`, which makes it possible to use microbial abbreviations, codes and names as input. Please see *Examples*.

##### Usage

```
mo_name(x, language = get_locale(), ...)

mo_fullname(x, language = get_locale(), ...)

mo_shortname(x, language = get_locale(), ...)

mo_subspecies(x, language = get_locale(), ...)

mo_species(x, language = get_locale(), ...)

mo_genus(x, language = get_locale(), ...)

mo_family(x, language = get_locale(), ...)

mo_order(x, language = get_locale(), ...)

mo_class(x, language = get_locale(), ...)

mo_phylum(x, language = get_locale(), ...)
```

```
mo_kingdom(x, language = get_locale(), ...)

mo_domain(x, language = get_locale(), ...)

mo_type(x, language = get_locale(), ...)

mo_gramstain(x, language = get_locale(), ...)

mo_is_gram_negative(x, language = get_locale(), ...)

mo_is_gram_positive(x, language = get_locale(), ...)

mo_is_intrinsic_resistant(x, ab, language = get_locale(), ...)

mo_snomed(x, language = get_locale(), ...)

mo_ref(x, language = get_locale(), ...)

mo_authors(x, language = get_locale(), ...)

mo_year(x, language = get_locale(), ...)

mo_rank(x, language = get_locale(), ...)

mo_taxonomy(x, language = get_locale(), ...)

mo_synonyms(x, language = get_locale(), ...)

mo_info(x, language = get_locale(), ...)

mo_url(x, open = FALSE, language = get_locale(), ...)

mo_property(x, property = "fullname", language = get_locale(), ...)
```

#### Arguments

| | |
|---|---|
| x | any character (vector) that can be coerced to a valid microorganism code with as.mo(). Can be left blank for auto-guessing the column containing microorganism codes when used inside dplyr verbs, such as filter(), mutate() and summarise(), please see *Examples*. |
| language | language of the returned text, defaults to system language (see get_locale()) and can be overwritten by setting the option AMR_locale, e.g. options(AMR_locale = "de"), see translate. Also used to translate text like "no growth". Use language = NULL or language = "" to prevent translation. |
| ... | other arguments passed on to as.mo(), such as 'allow_uncertain' and 'ignore_pattern' |
| ab | any (vector of) text that can be coerced to a valid antibiotic code with as.ab() |
| open | browse the URL using browseURL() |
| property | one of the column names of the microorganisms data set: "mo", "fullname", "kingdom", "phylum", "class", "order", "family", "genus", "species", "subspecies", "rank", "ref", "species_id", "source", "prevalence", "snomed", or must be "shortname" |

#### Details

All functions will return the most recently known taxonomic property according to the Catalogue of Life, except for `mo_ref()`, `mo_authors()` and `mo_year()`. Please refer to this example, knowing that *Escherichia blattae* was renamed to *Shimwellia blattae* in 2010:

- `mo_name("Escherichia blattae")` will return `"Shimwellia blattae"` (with a message about the renaming)
- `mo_ref("Escherichia blattae")` will return `"Burgess et al., 1973"` (with a message about the renaming)
- `mo_ref("Shimwellia blattae")` will return `"Priest et al., 2010"` (without a message)

The short name - `mo_shortname()` - almost always returns the first character of the genus and the full species, like `"E. coli"`. Exceptions are abbreviations of staphylococci (such as *"CoNS"*, Coagulase-Negative Staphylococci) and beta-haemolytic streptococci (such as *"GBS"*, Group B Streptococci). Please bear in mind that e.g. *E. coli* could mean *Escherichia coli* (kingdom of Bacteria) as well as *Entamoeba coli* (kingdom of Protozoa). Returning to the full name will be done using `as.mo()` internally, giving priority to bacteria and human pathogens, i.e. `"E. coli"` will be considered *Escherichia coli*. In other words, `mo_fullname(mo_shortname("Entamoeba coli"))` returns `"Escherichia coli"`.

Since the top-level of the taxonomy is sometimes referred to as 'kingdom' and sometimes as 'domain', the functions `mo_kingdom()` and `mo_domain()` return the exact same results.

The Gram stain - `mo_gramstain()` - will be determined based on the taxonomic kingdom and phylum. According to Cavalier-Smith (2002, PMID 11837318), who defined subkingdoms Negibacteria and Posibacteria, only these phyla are Posibacteria: Actinobacteria, Chloroflexi, Firmicutes and Tenericutes. These bacteria are considered Gram-positive - all other bacteria are considered Gram-negative. Species outside the kingdom of Bacteria will return a value `NA`. Functions `mo_is_gram_negative()` and `mo_is_gram_positive()` always return `TRUE` or `FALSE` (except when the input is `NA` or the MO code is `UNKNOWN`), thus always return `FALSE` for species outside the taxonomic kingdom of Bacteria.

Intrinsic resistance - `mo_is_intrinsic_resistant()` - will be determined based on the intrinsic_resistant data set, which is based on 'EUCAST Expert Rules' and 'EUCAST Intrinsic Resistance and Unusual Phenotypes' v3.2 from 2020. The `mo_is_intrinsic_resistant()` can be vectorised over arguments `x` (input for microorganisms) and over `ab` (input for antibiotics).

All output will be translated where possible.

The function `mo_url()` will return the direct URL to the online database entry, which also shows the scientific reference of the concerned species.

#### Value

- An integer in case of `mo_year()`
- A list in case of `mo_taxonomy()` and `mo_info()`
- A named character in case of `mo_url()`
- A double in case of `mo_snomed()`
- A character in all other cases

#### Stable lifecycle

The lifecycle of this function is **stable**. In a stable function, major changes are unlikely. This means that the unlying code will generally evolve by adding new arguments; removing arguments or changing the meaning of existing arguments will be avoided.

If the unlying code needs breaking changes, they will occur gradually. For example, a argument will be deprecated and first continue to work, but will emit an message informing you of the change. Next, typically after at least one newly released version on CRAN, the message will be transformed to an error.

##### Matching score for microorganisms

With ambiguous user input in `as.mo()` and all the `mo_*` functions, the returned results are chosen based on their matching score using `mo_matching_score()`. This matching score $m$, is calculated as:

$$m_{(x,n)} = \frac{l_n - 0.5 \cdot \min \{ \, l_{\,n} \, \text{lev}(x, n)}{l_n \cdot p_n \cdot k_n}$$

where:

- $x$ is the user input;
- $n$ is a taxonomic name (genus, species, and subspecies);
- $l_n$ is the length of $n$;
- lev is the Levenshtein distance function, which counts any insertion, deletion and substitution as 1 that is needed to change $x$ into $n$;
- $p_n$ is the human pathogenic prevalence group of $n$, as described below;
- $k_n$ is the taxonomic kingdom of $n$, set as Bacteria = 1, Fungi = 2, Protozoa = 3, Archaea = 4, others = 5.

The grouping into human pathogenic prevalence ($p$) is based on experience from several microbiological laboratories in the Netherlands in conjunction with international reports on pathogen prevalence. **Group 1** (most prevalent microorganisms) consists of all microorganisms where the taxonomic class is Gammaproteobacteria or where the taxonomic genus is *Enterococcus*, *Staphylococcus* or *Streptococcus*. This group consequently contains all common Gram-negative bacteria, such as *Pseudomonas* and *Legionella* and all species within the order Enterobacterales. **Group 2** consists of all microorganisms where the taxonomic phylum is Proteobacteria, Firmicutes, Actinobacteria or Sarcomastigophora, or where the taxonomic genus is *Absidia*, *Acremonium*, *Actinotignum*, *Alternaria*, *Anaerosalibacter*, *Apophysomyces*, *Arachnia*, *Aspergillus*, *Aureobacterium*, *Aureobasidium*, *Bacteroides*, *Basidiobolus*, *Beauveria*, *Blastocystis*, *Branhamella*, *Calymmatobacterium*, *Candida*, *Capnocytophaga*, *Catabacter*, *Chaetomium*, *Chryseobacterium*, *Chryseomonas*, *Chrysonilia*, *Cladophialophora*, *Cladosporium*, *Conidiobolus*, *Cryptococcus*, *Curvularia*, *Exophiala*, *Exserohilum*, *Flavobacterium*, *Fonsecaea*, *Fusarium*, *Fusobacterium*, *Hendersonula*, *Hypomyces*, *Koserella*, *Lelliottia*, *Leptosphaeria*, *Leptotrichia*, *Malassezia*, *Malbranchea*, *Mortierella*, *Mucor*, *Mycocentrospora*, *Mycoplasma*, *Nectria*, *Ochroconis*, *Oidiodendron*, *Phoma*, *Piedraia*, *Pithomyces*, *Pityrosporum*, *Prevotella,\Pseudallescheria*, *Rhizomucor*, *Rhizopus*, *Rhodotorula*, *Scolecobasidium*, *Scopulariopsis*, *Scytalidium,Sporobolomyces*, *Stachybotrys*, *Stomatococcus*, *Treponema*, *Trichoderma*, *Trichophyton*, *Trichosporon*, *Tritirachium* or *Ureaplasma*. **Group 3** consists of all other microorganisms.

All matches are sorted descending on their matching score and for all user input values, the top match will be returned. This will lead to the effect that e.g., `"E. coli"` will return the microbial ID of *Escherichia coli* ($m = 0.688$, a highly prevalent microorganism found in humans) and not *Entamoeba coli* ($m = 0.079$, a less prevalent microorganism in humans), although the latter would alphabetically come first.

##### Catalogue of Life

This package contains the complete taxonomic tree of almost all microorganisms (~70,000 species) from the authoritative and comprehensive Catalogue of Life (CoL, `http://www.catalogueoflife.org`). The CoL is the most comprehensive and authoritative global index of species currently available. Nonetheless, we supplemented the CoL data with data from the List of Prokaryotic names with Standing in Nomenclature (LPSN, lpsn.dsmz.de). This supplementation is needed until the CoL+ project is finished, which we await.

Click here for more information about the included taxa. Check which versions of the CoL and LSPN were included in this package with `catalogue_of_life_version()`.

##### Source

1. Becker K *et al.* **Coagulase-Negative Staphylococci**. 2014. Clin Microbiol Rev. 27(4): 870–926; doi: 10.1128/CMR.0010913
2. Becker K *et al.* **Implications of identifying the recently defined members of the *S. aureus* complex, *S. argenteus* and *S. schweitzeri*: A position paper of members of the ESCMID Study Group for staphylococci and Staphylococcal Diseases (ESGS).** 2019. Clin Microbiol Infect; doi: 10.1016/j.cmi.2019.02.028
3. Becker K *et al.* **Emergence of coagulase-negative staphylococci** 2020. Expert Rev Anti Infect Ther. 18(4):349-366; doi: 10.1080/14787210.2020.1730813
4. Lancefield RC **A serological differentiation of human and other groups of hemolytic streptococci**. 1933. J Exp Med. 57(4): 571–95; doi: 10.1084/jem.57.4.571
5. Catalogue of Life: Annual Checklist (public online taxonomic database), `http://www.catalogueoflife.org` (check included annual version with `catalogue_of_life_version()`).

##### Reference data publicly available

All reference data sets (about microorganisms, antibiotics, R/SI interpretation, EUCAST rules, etc.) in this `AMR` package are publicly and freely available. We continually export our data sets to formats for use in R, SPSS, SAS, Stata and Excel. We also supply flat files that are machine-readable and suitable for input in any software program, such as laboratory information systems. Please find all download links on our website, which is automatically updated with every code change.

##### Read more on our website!

On our website `https://msberends.github.io/AMR/` you can find a comprehensive tutorial about how to conduct AMR analysis, the complete documentation of all functions and an example analysis using WHONET data. As we would like to better understand the backgrounds and needs of our users, please participate in our survey!

##### See Also

microorganisms

##### Examples

```
# taxonomic tree -----------------------------------------------------------
mo_kingdom("E. coli")         # "Bacteria"
mo_phylum("E. coli")          # "Proteobacteria"
mo_class("E. coli")           # "Gammaproteobacteria"
mo_order("E. coli")           # "Enterobacterales"
mo_family("E. coli")          # "Enterobacteriaceae"
```

```
mo_genus("E. coli")           # "Escherichia"
mo_species("E. coli")         # "coli"
mo_subspecies("E. coli")      # ""

# colloquial properties -----------------------------------------------------
mo_name("E. coli")            # "Escherichia coli"
mo_fullname("E. coli")        # "Escherichia coli" - same as mo_name()
mo_shortname("E. coli")       # "E. coli"

# other properties ----------------------------------------------------------
mo_gramstain("E. coli")       # "Gram-negative"
mo_snomed("E. coli")          # 112283007, 116395006, ... (SNOMED codes)
mo_type("E. coli")            # "Bacteria" (equal to kingdom, but may be translated)
mo_rank("E. coli")            # "species"
mo_url("E. coli")             # get the direct url to the online database entry
mo_synonyms("E. coli")        # get previously accepted taxonomic names

# scientific reference ------------------------------------------------------
mo_ref("E. coli")             # "Castellani et al., 1919"
mo_authors("E. coli")         # "Castellani et al."
mo_year("E. coli")            # 1919

# abbreviations known in the field ------------------------------------------
mo_genus("MRSA")              # "Staphylococcus"
mo_species("MRSA")            # "aureus"
mo_shortname("VISA")          # "S. aureus"
mo_gramstain("VISA")          # "Gram-positive"

mo_genus("EHEC")              # "Escherichia"
mo_species("EHEC")            # "coli"

# known subspecies ----------------------------------------------------------
mo_name("doylei")             # "Campylobacter jejuni doylei"
mo_genus("doylei")            # "Campylobacter"
mo_species("doylei")          # "jejuni"
mo_subspecies("doylei")       # "doylei"

mo_fullname("K. pneu rh")     # "Klebsiella pneumoniae rhinoscleromatis"
mo_shortname("K. pneu rh")    # "K. pneumoniae"

# Becker classification, see ?as.mo ----------------------------------------
mo_fullname("S. epi")                     # "Staphylococcus epidermidis"
mo_fullname("S. epi", Becker = TRUE)      # "Coagulase-negative Staphylococcus (CoNS)"
mo_shortname("S. epi")                    # "S. epidermidis"
mo_shortname("S. epi", Becker = TRUE)     # "CoNS"

# Lancefield classification, see ?as.mo ------------------------------------
mo_fullname("S. pyo")                     # "Streptococcus pyogenes"
mo_fullname("S. pyo", Lancefield = TRUE)  # "Streptococcus group A"
mo_shortname("S. pyo")                    # "S. pyogenes"
mo_shortname("S. pyo", Lancefield = TRUE) # "GAS" (='Group A Streptococci')

# language support  ---------------------------------------------------------
mo_gramstain("E. coli", language = "de")  # "Gramnegativ"
mo_gramstain("E. coli", language = "nl")  # "Gram-negatief"
```

```
mo_gramstain("E. coli", language = "es")  # "Gram negativo"

# mo_type is equal to mo_kingdom, but mo_kingdom will remain official
mo_kingdom("E. coli")                     # "Bacteria" on a German system
mo_type("E. coli")                        # "Bakterien" on a German system
mo_type("E. coli")                        # "Bacteria" on an English system

mo_fullname("S. pyogenes",
            Lancefield = TRUE,
            language = "de")              # "Streptococcus Gruppe A"
mo_fullname("S. pyogenes",
            Lancefield = TRUE,
            language = "nl")              # "Streptococcus groep A"

# other --------------------------------------------------------------------

# gram stains and intrinsic resistance can also be used as a filter in dplyr verbs
if (require("dplyr")) {
  example_isolates %>%
    filter(mo_is_gram_positive())

  example_isolates %>%
    filter(mo_is_intrinsic_resistant(ab = "vanco"))
}

# get a list with the complete taxonomy (from kingdom to subspecies)
mo_taxonomy("E. coli")
# get a list with the taxonomy, the authors, Gram-stain and URL to the online database
mo_info("E. coli")
```

---

`mo_source` *User-defined reference data set for microorganisms*

---

#### Description

These functions can be used to predefine your own reference to be used in `as.mo()` and consequently all `mo_*` functions (such as `mo_genus()` and `mo_gramstain()`).

This is **the fastest way** to have your organisation (or analysis) specific codes picked up and translated by this package, since you don't have to bother about it again after setting it up once.

#### Usage

```
set_mo_source(
  path,
  destination = getOption("AMR_mo_source", "~/mo_source.rds")
)

get_mo_source(destination = getOption("AMR_mo_source", "~/mo_source.rds"))
```

##### Arguments

`path` location of your reference file, see Details. Can be `""`, `NULL` or `FALSE` to delete the reference file.

`destination` destination of the compressed data file, default to the user's home directory.

##### Details

The reference file can be a text file separated with commas (CSV) or tabs or pipes, an Excel file (either 'xls' or 'xlsx' format) or an R object file (extension '.rds'). To use an Excel file, you will need to have the `readxl` package installed.

`set_mo_source()` will check the file for validity: it must be a data.frame, must have a column named `"mo"` which contains values from `microorganisms$mo` and must have a reference column with your own defined values. If all tests pass, `set_mo_source()` will read the file into R and will ask to export it to `"~/mo_source.rds"`. The CRAN policy disallows packages to write to the file system, although '*exceptions may be allowed in interactive sessions if the package obtains confirmation from the user*'. For this reason, this function only works in interactive sessions so that the user can **specifically confirm and allow** that this file will be created. The destination of this file can be set with the `destination` argument and defaults to the user's home directory. It can also be set as an R option, using `options(AMR_mo_source = "my/location/file.rds")`.

The created compressed data file `"mo_source.rds"` will be used at default for MO determination (function `as.mo()` and consequently all mo_* functions like `mo_genus()` and `mo_gramstain()`). The location and timestamp of the original file will be saved as an attribute to the compressed data file.

The function `get_mo_source()` will return the data set by reading `"mo_source.rds"` with `readRDS()`. If the original file has changed (by checking the location and timestamp of the original file), it will call `set_mo_source()` to update the data file automatically if used in an interactive session.

Reading an Excel file (`.xlsx`) with only one row has a size of 8-9 kB. The compressed file created with `set_mo_source()` will then have a size of 0.1 kB and can be read by `get_mo_source()` in only a couple of microseconds (millionths of a second).

##### How to setup

Imagine this data on a sheet of an Excel file (mo codes were looked up in the microorganisms data set). The first column contains the organisation specific codes, the second column contains an MO code from this package:

```
  |         A          |       B      |
--|--------------------|--------------|
1 | Organisation XYZ   | mo           |
2 | lab_mo_ecoli       | B_ESCHR_COLI |
3 | lab_mo_kpneumoniae | B_KLBSL_PNMN |
4 |                    |              |
```

We save it as `"home/me/ourcodes.xlsx"`. Now we have to set it as a source:

```
set_mo_source("home/me/ourcodes.xlsx")
#> NOTE: Created mo_source file '/Users/me/mo_source.rds' (0.3 kB) from
#>       '/Users/me/Documents/ourcodes.xlsx' (9 kB), columns
#>       "Organisation XYZ" and "mo"
```

It has now created a file "~/mo_source.rds" with the contents of our Excel file. Only the first column with foreign values and the 'mo' column will be kept when creating the RDS file.

And now we can use it in our functions:

```
as.mo("lab_mo_ecoli")
#> Class <mo>
#> [1] B_ESCHR_COLI

mo_genus("lab_mo_kpneumoniae")
#> [1] "Klebsiella"

# other input values still work too
as.mo(c("Escherichia coli", "E. coli", "lab_mo_ecoli"))
#> NOTE: Translation to one microorganism was guessed with uncertainty.
#>       Use mo_uncertainties() to review it.
#> Class <mo>
#> [1] B_ESCHR_COLI B_ESCHR_COLI B_ESCHR_COLI
```

If we edit the Excel file by, let's say, adding row 4 like this:

```
  |         A          |       B      |
--|--------------------|--------------|
1 | Organisation XYZ   | mo           |
2 | lab_mo_ecoli       | B_ESCHR_COLI |
3 | lab_mo_kpneumoniae | B_KLBSL_PNMN |
4 | lab_Staph_aureus   | B_STPHY_AURS |
5 |                    |              |
```

...any new usage of an MO function in this package will update your data file:

```
as.mo("lab_mo_ecoli")
#> NOTE: Updated mo_source file '/Users/me/mo_source.rds' (0.3 kB) from
#>       '/Users/me/Documents/ourcodes.xlsx' (9 kB), columns
#>        "Organisation XYZ" and "mo"
#> Class <mo>
#> [1] B_ESCHR_COLI

mo_genus("lab_Staph_aureus")
#> [1] "Staphylococcus"
```

To delete the reference data file, just use `""`, `NULL` or `FALSE` as input for `set_mo_source()`:

```
set_mo_source(NULL)
#> Removed mo_source file '/Users/me/mo_source.rds'
```

If the original file (in the previous case an Excel file) is moved or deleted, the `mo_source.rds` file will be removed upon the next use of `as.mo()` or any `mo_*` function.

##### Stable lifecycle

The lifecycle of this function is **stable**. In a stable function, major changes are unlikely. This means that the unlying code will generally evolve by adding new arguments; removing arguments or changing the meaning of existing arguments will be avoided.

If the unlying code needs breaking changes, they will occur gradually. For example, a argument will be deprecated and first continue to work, but will emit an message informing you of the change. Next, typically after at least one newly released version on CRAN, the message will be transformed to an error.

#### Read more on our website!

On our website `https://msberends.github.io/AMR/` you can find a comprehensive tutorial about how to conduct AMR analysis, the complete documentation of all functions and an example analysis using WHONET data. As we would like to better understand the backgrounds and needs of our users, please participate in our survey!

---

`pca` *Principal Component Analysis (for AMR)*

---

#### Description

Performs a principal component analysis (PCA) based on a data set with automatic determination for afterwards plotting the groups and labels, and automatic filtering on only suitable (i.e. non-empty and numeric) variables.

#### Usage

```
pca(
  x,
  ...,
  retx = TRUE,
  center = TRUE,
  scale. = TRUE,
  tol = NULL,
  rank. = NULL
)
```

#### Arguments

| | |
|---|---|
| `x` | a data.frame containing numeric columns |
| `...` | columns of `x` to be selected for PCA, can be unquoted since it supports quasiquotation. |
| `retx` | a logical value indicating whether the rotated variables should be returned. |
| `center` | a logical value indicating whether the variables should be shifted to be zero centered. Alternately, a vector of length equal the number of columns of `x` can be supplied. The value is passed to `scale`. |
| `scale.` | a logical value indicating whether the variables should be scaled to have unit variance before the analysis takes place. The default is `FALSE` for consistency with S, but in general scaling is advisable. Alternatively, a vector of length equal the number of columns of `x` can be supplied. The value is passed to `scale`. |
| `tol` | a value indicating the magnitude below which components should be omitted. (Components are omitted if their standard deviations are less than or equal to `tol` times the standard deviation of the first component.) With the default null setting, no components are omitted (unless `rank.` is specified less than |

min(dim(x)).). Other settings for tol could be tol = 0 or tol = sqrt(.Machine$double.eps), which would omit essentially constant components.

rank. optionally, a number specifying the maximal rank, i.e., maximal number of principal components to be used. Can be set as alternative or in addition to tol, useful notably when the desired rank is considerably smaller than the dimensions of the matrix.

#### Details

The pca() function takes a data.frame as input and performs the actual PCA with the R function prcomp().

The result of the pca() function is a prcomp object, with an additional attribute non_numeric_cols which is a vector with the column names of all columns that do not contain numeric values. These are probably the groups and labels, and will be used by ggplot_pca().

#### Value

An object of classes pca and prcomp

#### Maturing lifecycle

The lifecycle of this function is **maturing**. The unlying code of a maturing function has been roughed out, but finer details might still change. Since this function needs wider usage and more extensive testing, you are very welcome to suggest changes at our repository or write us an email (see section 'Contact Us').

#### Examples

```
# `example_isolates` is a dataset available in the AMR package.
# See ?example_isolates.

if (require("dplyr")) {
  # calculate the resistance per group first
  resistance_data <- example_isolates %>%
    group_by(order = mo_order(mo),       # group on anything, like order
             genus = mo_genus(mo)) %>%   #  and genus as we do here
    summarise_if(is.rsi, resistance)     # then get resistance of all drugs

  # now conduct PCA for certain antimicrobial agents
  pca_result <- resistance_data %>%
    pca(AMC, CXM, CTX, CAZ, GEN, TOB, TMP, SXT)

  pca_result
  summary(pca_result)
  biplot(pca_result)
  ggplot_pca(pca_result) # a new and convenient plot function
}
```

`proportion` *Calculate microbial resistance*

#### Description

These functions can be used to calculate the (co-)resistance or susceptibility of microbial isolates (i.e. percentage of S, SI, I, IR or R). All functions support quasiquotation with pipes, can be used in `summarise()` from the `dplyr` package and also support grouped variables, please see *Examples*.

`resistance()` should be used to calculate resistance, `susceptibility()` should be used to calculate susceptibility.

#### Usage

```
resistance(..., minimum = 30, as_percent = FALSE, only_all_tested = FALSE)

susceptibility(..., minimum = 30, as_percent = FALSE, only_all_tested = FALSE)

proportion_R(..., minimum = 30, as_percent = FALSE, only_all_tested = FALSE)

proportion_IR(..., minimum = 30, as_percent = FALSE, only_all_tested = FALSE)

proportion_I(..., minimum = 30, as_percent = FALSE, only_all_tested = FALSE)

proportion_SI(..., minimum = 30, as_percent = FALSE, only_all_tested = FALSE)

proportion_S(..., minimum = 30, as_percent = FALSE, only_all_tested = FALSE)

proportion_df(
  data,
  translate_ab = "name",
  language = get_locale(),
  minimum = 30,
  as_percent = FALSE,
  combine_SI = TRUE,
  combine_IR = FALSE
)

rsi_df(
  data,
  translate_ab = "name",
  language = get_locale(),
  minimum = 30,
  as_percent = FALSE,
  combine_SI = TRUE,
  combine_IR = FALSE
)
```

#### Arguments

| | |
|---|---|
| ... | one or more vectors (or columns) with antibiotic interpretations. They will be transformed internally with as.rsi() if needed. Use multiple columns to calculate (the lack of) co-resistance: the probability where one of two drugs have a resistant or susceptible result. See Examples. |
| minimum | the minimum allowed number of available (tested) isolates. Any isolate count lower than minimum will return NA with a warning. The default number of 30 isolates is advised by the Clinical and Laboratory Standards Institute (CLSI) as best practice, see Source. |
| as_percent | a logical to indicate whether the output must be returned as a hundred fold with % sign (a character). A value of 0.123456 will then be returned as "12.3%". |
| only_all_tested | (for combination therapies, i.e. using more than one variable for ...): a logical to indicate that isolates must be tested for all antibiotics, see section *Combination therapy* below |
| data | a data.frame containing columns with class rsi (see as.rsi()) |
| translate_ab | a column name of the antibiotics data set to translate the antibiotic abbreviations to, using ab_property() |
| language | language of the returned text, defaults to system language (see get_locale()) and can also be set with getOption("AMR_locale"). Use language = NULL or language = "" to prevent translation. |
| combine_SI | a logical to indicate whether all values of S and I must be merged into one, so the output only consists of S+I vs. R (susceptible vs. resistant). This used to be the argument combine_IR, but this now follows the redefinition by EUCAST about the interpretation of I (increased exposure) in 2019, see section 'Interpretation of S, I and R' below. Default is TRUE. |
| combine_IR | a logical to indicate whether all values of I and R must be merged into one, so the output only consists of S vs. I+R (susceptible vs. non-susceptible). This is outdated, see argument combine_SI. |

#### Details

The function resistance() is equal to the function proportion_R(). The function susceptibility() is equal to the function proportion_SI().

**Remember that you should filter your table to let it contain only first isolates!** This is needed to exclude duplicates and to reduce selection bias. Use first_isolate() to determine them in your data set.

These functions are not meant to count isolates, but to calculate the proportion of resistance/susceptibility. Use the count() functions to count isolates. The function susceptibility() is essentially equal to count_susceptible() / count_all(). *Low counts can influence the outcome - the* proportion *functions may camouflage this, since they only return the proportion (albeit being dependent on the* minimum *argument).*

The function proportion_df() takes any variable from data that has an rsi class (created with as.rsi()) and calculates the proportions R, I and S. It also supports grouped variables. The function rsi_df() works exactly like proportion_df(), but adds the number of isolates.

#### Value

A double or, when as_percent = TRUE, a character.

##### Combination therapy

When using more than one variable for ... (= combination therapy), use `only_all_tested` to only count isolates that are tested for all antibiotics/variables that you test them for. See this example for two antibiotics, Drug A and Drug B, about how `susceptibility()` works to calculate the %SI:

| Drug A | Drug B | only_all_tested = FALSE<br>include as numerator | only_all_tested = FALSE<br>include as denominator | only_all_tested = TRUE<br>include as numerator | only_all_tested = TRUE<br>include as denominator |
|---|---|---|---|---|---|
| S or I | S or I | X | X | X | X |
| R | S or I | X | X | X | X |
| <NA> | S or I | X | X | - | - |
| S or I | R | X | X | X | X |
| R | R | - | X | - | X |
| <NA> | R | - | - | - | - |
| S or I | <NA> | X | X | - | - |
| R | <NA> | - | - | - | - |
| <NA> | <NA> | - | - | - | - |

Please note that, in combination therapies, for `only_all_tested = TRUE` applies that:

```
    count_S()    +   count_I()    +   count_R()    = count_all()
  proportion_S() + proportion_I() + proportion_R() = 1
```

and that, in combination therapies, for `only_all_tested = FALSE` applies that:

```
    count_S()    +   count_I()    +   count_R()    >= count_all()
  proportion_S() + proportion_I() + proportion_R() >= 1
```

Using `only_all_tested` has no impact when only using one antibiotic as input.

##### Stable lifecycle

The lifecycle of this function is **stable**. In a stable function, major changes are unlikely. This means that the unlying code will generally evolve by adding new arguments; removing arguments or changing the meaning of existing arguments will be avoided.

If the unlying code needs breaking changes, they will occur gradually. For example, a argument will be deprecated and first continue to work, but will emit an message informing you of the change. Next, typically after at least one newly released version on CRAN, the message will be transformed to an error.

##### Interpretation of R and S/I

In 2019, the European Committee on Antimicrobial Susceptibility Testing (EUCAST) has decided to change the definitions of susceptibility testing categories R and S/I as shown below (`https://www.eucast.org/newsiandr/`).

- **R = Resistant**
  A microorganism is categorised as *Resistant* when there is a high likelihood of therapeutic failure even when there is increased exposure. Exposure is a function of how the mode of administration, dose, dosing interval, infusion time, as well as distribution and excretion of the antimicrobial agent will influence the infecting organism at the site of infection.

- **S = Susceptible**
  A microorganism is categorised as *Susceptible, standard dosing regimen*, when there is a high likelihood of therapeutic success using a standard dosing regimen of the agent.
- **I = Increased exposure, but still susceptible**
  A microorganism is categorised as *Susceptible, Increased exposure* when there is a high likelihood of therapeutic success because exposure to the agent is increased by adjusting the dosing regimen or by its concentration at the site of infection.

This AMR package honours this new insight. Use `susceptibility()` (equal to `proportion_SI()`) to determine antimicrobial susceptibility and `count_susceptible()` (equal to `count_SI()`) to count susceptible isolates.

#### Read more on our website!

On our website `https://msberends.github.io/AMR/` you can find a comprehensive tutorial about how to conduct AMR analysis, the complete documentation of all functions and an example analysis using WHONET data. As we would like to better understand the backgrounds and needs of our users, please participate in our survey!

#### Source

**M39 Analysis and Presentation of Cumulative Antimicrobial Susceptibility Test Data, 4th Edition**, 2014, *Clinical and Laboratory Standards Institute (CLSI)*. `https://clsi.org/standards/products/microbiology/documents/m39/`.

#### See Also

`count()` to count resistant and susceptible isolates.

#### Examples

```
# example_isolates is a data set available in the AMR package.
?example_isolates

resistance(example_isolates$AMX)     # determines %R
susceptibility(example_isolates$AMX) # determines %S+I

# be more specific
proportion_S(example_isolates$AMX)
proportion_SI(example_isolates$AMX)
proportion_I(example_isolates$AMX)
proportion_IR(example_isolates$AMX)
proportion_R(example_isolates$AMX)

if (require("dplyr")) {
  example_isolates %>%
    group_by(hospital_id) %>%
    summarise(r = resistance(CIP),
              n = n_rsi(CIP)) # n_rsi works like n_distinct in dplyr, see ?n_rsi

  example_isolates %>%
    group_by(hospital_id) %>%
    summarise(R  = resistance(CIP, as_percent = TRUE),
              SI = susceptibility(CIP, as_percent = TRUE),
              n1 = count_all(CIP),  # the actual total; sum of all three
```

```
            n2 = n_rsi(CIP),      # same - analogous to n_distinct
            total = n())          # NOT the number of tested isolates!

  # Calculate co-resistance between amoxicillin/clav acid and gentamicin,
  # so we can see that combination therapy does a lot more than mono therapy:
  example_isolates %>% susceptibility(AMC)  # %SI = 76.3%
  example_isolates %>% count_all(AMC)       #   n = 1879

  example_isolates %>% susceptibility(GEN)  # %SI = 75.4%
  example_isolates %>% count_all(GEN)       #   n = 1855

  example_isolates %>% susceptibility(AMC, GEN) # %SI = 94.1%
  example_isolates %>% count_all(AMC, GEN)      #   n = 1939

  # See Details on how `only_all_tested` works. Example:
  example_isolates %>%
    summarise(numerator = count_susceptible(AMC, GEN),
              denominator = count_all(AMC, GEN),
              proportion = susceptibility(AMC, GEN))

  example_isolates %>%
    summarise(numerator = count_susceptible(AMC, GEN, only_all_tested = TRUE),
              denominator = count_all(AMC, GEN, only_all_tested = TRUE),
              proportion = susceptibility(AMC, GEN, only_all_tested = TRUE))

  example_isolates %>%
    group_by(hospital_id) %>%
    summarise(cipro_p = susceptibility(CIP, as_percent = TRUE),
              cipro_n = count_all(CIP),
              genta_p = susceptibility(GEN, as_percent = TRUE),
              genta_n = count_all(GEN),
              combination_p = susceptibility(CIP, GEN, as_percent = TRUE),
              combination_n = count_all(CIP, GEN))

  # Get proportions S/I/R immediately of all rsi columns
  example_isolates %>%
    select(AMX, CIP) %>%
    proportion_df(translate = FALSE)

  # It also supports grouping variables
  example_isolates %>%
    select(hospital_id, AMX, CIP) %>%
    group_by(hospital_id) %>%
    proportion_df(translate = FALSE)
}
```

---

`random` *Random MIC values/disk zones/RSI generation*

---

#### Description

These functions can be used for generating random MIC values and disk diffusion diameters, for AMR analysis practice.

#### Usage

```
random_mic(size, mo = NULL, ab = NULL, ...)

random_disk(size, mo = NULL, ab = NULL, ...)

random_rsi(size, prob_RSI = c(0.33, 0.33, 0.33), ...)
```

#### Arguments

| | |
|---|---|
| size | desired size of the returned vector |
| mo | any character that can be coerced to a valid microorganism code with as.mo() |
| ab | any character that can be coerced to a valid antimicrobial agent code with as.ab() |
| ... | extension for future versions, not used at the moment |
| prob_RSI | a vector of length 3: the probabilities for R (1st value), S (2nd value) and I (3rd value) |

#### Details

The base R function sample() is used for generating values.

Generated values are based on the latest EUCAST guideline implemented in the rsi_translation data set. To create specific generated values per bug or drug, set the mo and/or ab argument.

#### Value

class <mic> for random_mic() (see as.mic()) and class <disk> for random_disk() (see as.disk())

#### Maturing lifecycle

The lifecycle of this function is **maturing**. The unlying code of a maturing function has been roughed out, but finer details might still change. Since this function needs wider usage and more extensive testing, you are very welcome to suggest changes at our repository or write us an email (see section 'Contact Us').

#### Read more on our website!

On our website https://msberends.github.io/AMR/ you can find a comprehensive tutorial about how to conduct AMR analysis, the complete documentation of all functions and an example analysis using WHONET data. As we would like to better understand the backgrounds and needs of our users, please participate in our survey!

#### Examples

```
random_mic(100)
random_disk(100)
random_rsi(100)

# make the random generation more realistic by setting a bug and/or drug:
random_mic(100, "Klebsiella pneumoniae")                # range 0.0625-64
random_mic(100, "Klebsiella pneumoniae", "meropenem")   # range 0.0625-16
random_mic(100, "Streptococcus pneumoniae", "meropenem") # range 0.0625-4

random_disk(100, "Klebsiella pneumoniae")                 # range 11-50
```

```
random_disk(100, "Klebsiella pneumoniae", "ampicillin")    # range 6-14
random_disk(100, "Streptococcus pneumoniae", "ampicillin") # range 16-22
```

---

resistance_predict *Predict antimicrobial resistance*

---

##### Description

Create a prediction model to predict antimicrobial resistance for the next years on statistical solid ground. Standard errors (SE) will be returned as columns se_min and se_max. See *Examples* for a real live example.

##### Usage

```
resistance_predict(
  x,
  col_ab,
  col_date = NULL,
  year_min = NULL,
  year_max = NULL,
  year_every = 1,
  minimum = 30,
  model = NULL,
  I_as_S = TRUE,
  preserve_measurements = TRUE,
  info = interactive(),
  ...
)

rsi_predict(
  x,
  col_ab,
  col_date = NULL,
  year_min = NULL,
  year_max = NULL,
  year_every = 1,
  minimum = 30,
  model = NULL,
  I_as_S = TRUE,
  preserve_measurements = TRUE,
  info = interactive(),
  ...
)

## S3 method for class 'resistance_predict'
plot(x, main = paste("Resistance Prediction of", x_name), ...)

ggplot_rsi_predict(
  x,
  main = paste("Resistance Prediction of", x_name),
```

```
  ribbon = TRUE,
  ...
)
```

#### Arguments

| | |
|---|---|
| x | a data.frame containing isolates. Can be left blank when used inside dplyr verbs, such as `filter()`, `mutate()` and `summarise()`. |
| col_ab | column name of x containing antimicrobial interpretations ("R", "I" and "S") |
| col_date | column name of the date, will be used to calculate years if this column doesn't consist of years already, defaults to the first column of with a date class |
| year_min | lowest year to use in the prediction model, dafaults to the lowest year in col_date |
| year_max | highest year to use in the prediction model, defaults to 10 years after today |
| year_every | unit of sequence between lowest year found in the data and year_max |
| minimum | minimal amount of available isolates per year to include. Years containing less observations will be estimated by the model. |
| model | the statistical model of choice. This could be a generalised linear regression model with binomial distribution (i.e. using 'glm(..., family = binomial)", assuming that a period of zero resistance was followed by a period of increasing resistance leading slowly to more and more resistance. See Details for all valid options. |
| I_as_S | a logical to indicate whether values "I" should be treated as "S" (will otherwise be treated as "R"). The default, TRUE, follows the redefinition by EUCAST about the interpretation of I (increased exposure) in 2019, see section *Interpretation of S, I and R* below. |
| preserve_measurements | a logical to indicate whether predictions of years that are actually available in the data should be overwritten by the original data. The standard errors of those years will be NA. |
| info | a logical to indicate whether textual analysis should be printed with the name and `summary()` of the statistical model. |
| ... | arguments passed on to functions |
| main | title of the plot |
| ribbon | a logical to indicate whether a ribbon should be shown (default) or error bars |

#### Details

Valid options for the statistical model (argument model) are:

- "binomial" or "binom" or "logit": a generalised linear regression model with binomial distribution
- "loglin" or "poisson": a generalised log-linear regression model with poisson distribution
- "lin" or "linear": a linear regression model

#### Value

A data.frame with extra class `resistance_predict` with columns:

- year

- value, the same as estimated when preserve_measurements = FALSE, and a combination of observed and estimated otherwise
- se_min, the lower bound of the standard error with a minimum of 0 (so the standard error will never go below 0%)
- se_max the upper bound of the standard error with a maximum of 1 (so the standard error will never go above 100%)
- observations, the total number of available observations in that year, i.e. $S + I + R$
- observed, the original observed resistant percentages
- estimated, the estimated resistant percentages, calculated by the model

Furthermore, the model itself is available as an attribute: attributes(x)$model, please see *Examples*.

##### Maturing lifecycle

The lifecycle of this function is **maturing**. The unlying code of a maturing function has been roughed out, but finer details might still change. Since this function needs wider usage and more extensive testing, you are very welcome to suggest changes at our repository or write us an email (see section 'Contact Us').

##### Interpretation of R and S/I

In 2019, the European Committee on Antimicrobial Susceptibility Testing (EUCAST) has decided to change the definitions of susceptibility testing categories R and S/I as shown below (https://www.eucast.org/newsiandr/).

- **R = Resistant**
  A microorganism is categorised as *Resistant* when there is a high likelihood of therapeutic failure even when there is increased exposure. Exposure is a function of how the mode of administration, dose, dosing interval, infusion time, as well as distribution and excretion of the antimicrobial agent will influence the infecting organism at the site of infection.
- **S = Susceptible**
  A microorganism is categorised as *Susceptible, standard dosing regimen*, when there is a high likelihood of therapeutic success using a standard dosing regimen of the agent.
- **I = Increased exposure, but still susceptible**
  A microorganism is categorised as *Susceptible, Increased exposure* when there is a high likelihood of therapeutic success because exposure to the agent is increased by adjusting the dosing regimen or by its concentration at the site of infection.

This AMR package honours this new insight. Use susceptibility() (equal to proportion_SI()) to determine antimicrobial susceptibility and count_susceptible() (equal to count_SI()) to count susceptible isolates.

##### Read more on our website!

On our website https://msberends.github.io/AMR/ you can find a comprehensive tutorial about how to conduct AMR analysis, the complete documentation of all functions and an example analysis using WHONET data. As we would like to better understand the backgrounds and needs of our users, please participate in our survey!

#### See Also

The `proportion()` functions to calculate resistance

Models: `lm()` `glm()`

#### Examples

```
x <- resistance_predict(example_isolates,
                        col_ab = "AMX",
                        year_min = 2010,
                        model = "binomial")
plot(x)
if (require("ggplot2")) {
  ggplot_rsi_predict(x)
}

# using dplyr:
if (require("dplyr")) {
  x <- example_isolates %>%
    filter_first_isolate() %>%
    filter(mo_genus(mo) == "Staphylococcus") %>%
    resistance_predict("PEN", model = "binomial")
  plot(x)

  # get the model from the object
  mymodel <- attributes(x)$model
  summary(mymodel)
}

# create nice plots with ggplot2 yourself
if (require("dplyr") & require("ggplot2")) {

  data <- example_isolates %>%
    filter(mo == as.mo("E. coli")) %>%
    resistance_predict(col_ab = "AMX",
                       col_date = "date",
                       model = "binomial",
                       info = FALSE,
                       minimum = 15)

  ggplot(data,
         aes(x = year)) +
    geom_col(aes(y = value),
             fill = "grey75") +
    geom_errorbar(aes(ymin = se_min,
                      ymax = se_max),
                  colour = "grey50") +
    scale_y_continuous(limits = c(0, 1),
                       breaks = seq(0, 1, 0.1),
                       labels = paste0(seq(0, 100, 10), "%")) +
    labs(title = expression(paste("Forecast of Amoxicillin Resistance in ",
                                  italic("E. coli"))),
         y = "%R",
         x = "Year") +
    theme_minimal(base_size = 13)
}
```

rsi_translation *Data set for R/SI interpretation*

#### Description

Data set to interpret MIC and disk diffusion to R/SI values. Included guidelines are CLSI (2010-2019) and EUCAST (2011-2020). Use `as.rsi()` to transform MICs or disks measurements to R/SI values.

#### Usage

```
rsi_translation
```

#### Format

A data.frame with 18,650 observations and 10 variables:

- `guideline`
  Name of the guideline
- `method`
  Either "MIC" or "DISK"
- `site`
  Body site, e.g. "Oral" or "Respiratory"
- `mo`
  Microbial ID, see `as.mo()`
- `ab`
  Antibiotic ID, see `as.ab()`
- `ref_tbl`
  Info about where the guideline rule can be found
- `disk_dose`
  Dose of the used disk diffusion method
- `breakpoint_S`
  Lowest MIC value or highest number of millimetres that leads to "S"
- `breakpoint_R`
  Highest MIC value or lowest number of millimetres that leads to "R"
- `uti`
  A logical value (`TRUE`/`FALSE`) to indicate whether the rule applies to a urinary tract infection (UTI)

#### Details

The repository of this `AMR` package contains a file comprising this exact data set: `https://github.com/msberends/AMR/blob/master/data-raw/rsi_translation.txt`. This file **allows for machine reading EUCAST and CLSI guidelines**, which is almost impossible with the Excel and PDF files distributed by EUCAST and CLSI. The file is updated automatically.

##### Reference data publicly available

All reference data sets (about microorganisms, antibiotics, R/SI interpretation, EUCAST rules, etc.) in this AMR package are publicly and freely available. We continually export our data sets to formats for use in R, SPSS, SAS, Stata and Excel. We also supply flat files that are machine-readable and suitable for input in any software program, such as laboratory information systems. Please find all download links on our website, which is automatically updated with every code change.

##### Read more on our website!

On our website `https://msberends.github.io/AMR/` you can find a comprehensive tutorial about how to conduct AMR analysis, the complete documentation of all functions and an example analysis using WHONET data. As we would like to better understand the backgrounds and needs of our users, please participate in our survey!

##### See Also

intrinsic_resistant

---

#### skewness — *Skewness of the sample*

---

##### Description

Skewness is a measure of the asymmetry of the probability distribution of a real-valued random variable about its mean.

When negative ('left-skewed'): the left tail is longer; the mass of the distribution is concentrated on the right of a histogram. When positive ('right-skewed'): the right tail is longer; the mass of the distribution is concentrated on the left of a histogram. A normal distribution has a skewness of 0.

##### Usage

```
skewness(x, na.rm = FALSE)

## Default S3 method:
skewness(x, na.rm = FALSE)

## S3 method for class 'matrix'
skewness(x, na.rm = FALSE)

## S3 method for class 'data.frame'
skewness(x, na.rm = FALSE)
```

##### Arguments

| | |
|---|---|
| `x` | a vector of values, a matrix or a data.frame |
| `na.rm` | a logical value indicating whether NA values should be stripped before the computation proceeds |

##### Stable lifecycle

The lifecycle of this function is **stable**. In a stable function, major changes are unlikely. This means that the unlying code will generally evolve by adding new arguments; removing arguments or changing the meaning of existing arguments will be avoided.

If the unlying code needs breaking changes, they will occur gradually. For example, a argument will be deprecated and first continue to work, but will emit an message informing you of the change. Next, typically after at least one newly released version on CRAN, the message will be transformed to an error.

##### Read more on our website!

On our website `https://msberends.github.io/AMR/` you can find a comprehensive tutorial about how to conduct AMR analysis, the complete documentation of all functions and an example analysis using WHONET data. As we would like to better understand the backgrounds and needs of our users, please participate in our survey!

##### See Also

`kurtosis()`

---

`translate` *Translate strings from AMR package*

---

##### Description

For language-dependent output of AMR functions, like `mo_name()`, `mo_gramstain()`, `mo_type()` and `ab_name()`.

##### Usage

```
get_locale()
```

##### Details

Strings will be translated to foreign languages if they are defined in a local translation file. Additions to this file can be suggested at our repository. The file can be found here: `https://github.com/msberends/AMR/blob/master/data-raw/translations.tsv`. This file will be read by all functions where a translated output can be desired, like all `mo_*` functions (such as `mo_name()`, `mo_gramstain()`, `mo_type()`, etc.) and `ab_*` functions (such as `ab_name()`, `ab_group()`, etc.).

Currently supported languages are: Dutch, English, French, German, Italian, Portuguese, Spanish. Please note that currently not all these languages have translations available for all antimicrobial agents and colloquial microorganism names.

Please suggest your own translations by creating a new issue on our repository.

**Changing the default language:**

The system language will be used at default (as returned by `Sys.getenv("LANG")` or, if `LANG` is not set, `Sys.getlocale()`), if that language is supported. But the language to be used can be overwritten in two ways and will be checked in this order:

1. Setting the R option `AMR_locale`, e.g. by running `options(AMR_locale = "de")`
2. Setting the system variable `LANGUAGE` or `LANG`, e.g. by adding `LANGUAGE="de_DE.utf8"` to your `.Renviron` file in your home directory

So if the R option `AMR_locale` is set, the system variables `LANGUAGE` and `LANG` will be ignored.

##### Stable lifecycle

The lifecycle of this function is **stable**. In a stable function, major changes are unlikely. This means that the unlying code will generally evolve by adding new arguments; removing arguments or changing the meaning of existing arguments will be avoided.

If the unlying code needs breaking changes, they will occur gradually. For example, a argument will be deprecated and first continue to work, but will emit an message informing you of the change. Next, typically after at least one newly released version on CRAN, the message will be transformed to an error.

##### Read more on our website!

On our website `https://msberends.github.io/AMR/` you can find a comprehensive tutorial about how to conduct AMR analysis, the complete documentation of all functions and an example analysis using WHONET data. As we would like to better understand the backgrounds and needs of our users, please participate in our survey!

##### Examples

```
# The 'language' argument of below functions
# will be set automatically to your system language
# with get_locale()

# English
mo_name("CoNS", language = "en")
#> "Coagulase-negative Staphylococcus (CoNS)"

# German
mo_name("CoNS", language = "de")
#> "Koagulase-negative Staphylococcus (KNS)"

# Dutch
mo_name("CoNS", language = "nl")
#> "Coagulase-negatieve Staphylococcus (CNS)"

# Spanish
mo_name("CoNS", language = "es")
#> "Staphylococcus coagulasa negativo (SCN)"

# Italian
mo_name("CoNS", language = "it")
#> "Staphylococcus negativo coagulasi (CoNS)"

# Portuguese
mo_name("CoNS", language = "pt")
#> "Staphylococcus coagulase negativo (CoNS)"
```

---

| | |
|---|---|
| `WHOCC` | *WHOCC: WHO Collaborating Centre for Drug Statistics Methodology* |

---

##### Description

All antimicrobial drugs and their official names, ATC codes, ATC groups and defined daily dose (DDD) are included in this package, using the WHO Collaborating Centre for Drug Statistics Methodology.

##### WHOCC

This package contains **all ~550 antibiotic, antimycotic and antiviral drugs** and their Anatomical Therapeutic Chemical (ATC) codes, ATC groups and Defined Daily Dose (DDD) from the World Health Organization Collaborating Centre for Drug Statistics Methodology (WHOCC, https://www.whocc.no) and the Pharmaceuticals Community Register of the European Commission (http://ec.europa.eu/health/documents/community-register/html/atc.htm).

These have become the gold standard for international drug utilisation monitoring and research.

The WHOCC is located in Oslo at the Norwegian Institute of Public Health and funded by the Norwegian government. The European Commission is the executive of the European Union and promotes its general interest.

**NOTE: The WHOCC copyright does not allow use for commercial purposes, unlike any other info from this package.** See https://www.whocc.no/copyright_disclaimer/.

##### Read more on our website!

On our website https://msberends.github.io/AMR/ you can find a comprehensive tutorial about how to conduct AMR analysis, the complete documentation of all functions and an example analysis using WHONET data. As we would like to better understand the backgrounds and needs of our users, please participate in our survey!

##### Examples

```
as.ab("meropenem")
ab_name("J01DH02")

ab_tradenames("flucloxacillin")
```

---

`WHONET` *Data set with 500 isolates - WHONET example*

---

##### Description

This example data set has the exact same structure as an export file from WHONET. Such files can be used with this package, as this example data set shows. The antibiotic results are from our example_isolates data set. All patient names are created using online surname generators and are only in place for practice purposes.

##### Usage

```
WHONET
```

#### Format

A data.frame with 500 observations and 53 variables:

- Identification number
  ID of the sample
- Specimen number
  ID of the specimen
- `Organism`
  Name of the microorganism. Before analysis, you should transform this to a valid microbial class, using `as.mo()`.
- `Country`
  Country of origin
- `Laboratory`
  Name of laboratory
- Last name
  Fictitious last name of patient
- First name
  Fictitious initial of patient
- `Sex`
  Fictitious gender of patient
- `Age`
  Fictitious age of patient
- Age category
  Age group, can also be looked up using `age_groups()`
- Date of admission
  Date of hospital admission
- Specimen date
  Date when specimen was received at laboratory
- Specimen type
  Specimen type or group
- Specimen type (Numeric)
  Translation of `"Specimen type"`
- `Reason`
  Reason of request with Differential Diagnosis
- Isolate number
  ID of isolate
- Organism type
  Type of microorganism, can also be looked up using `mo_type()`
- `Serotype`
  Serotype of microorganism
- `Beta-lactamase`
  Microorganism produces beta-lactamase?
- `ESBL`
  Microorganism produces extended spectrum beta-lactamase?
- `Carbapenemase`
  Microorganism produces carbapenemase?
- MRSA screening test
  Microorganism is possible MRSA?

- Inducible clindamycin resistance
  Clindamycin can be induced?
- `Comment`
  Other comments
- Date of data entry
  Date this data was entered in WHONET
- `AMP_ND10:CIP_EE`
  28 different antibiotics. You can lookup the abbreviations in the antibiotics data set, or use e.g. `ab_name("AMP")` to get the official name immediately. Before analysis, you should transform this to a valid antibiotic class, using `as.rsi()`.

##### Reference data publicly available

All reference data sets (about microorganisms, antibiotics, R/SI interpretation, EUCAST rules, etc.) in this `AMR` package are publicly and freely available. We continually export our data sets to formats for use in R, SPSS, SAS, Stata and Excel. We also supply flat files that are machine-readable and suitable for input in any software program, such as laboratory information systems. Please find all download links on our website, which is automatically updated with every code change.

##### Read more on our website!

On our website `https://msberends.github.io/AMR/` you can find a comprehensive tutorial about how to conduct AMR analysis, the complete documentation of all functions and an example analysis using WHONET data. As we would like to better understand the backgrounds and needs of our users, please participate in our survey!

### Index

∗ **Becker**
as.mo, 22
∗ **Lancefield**
as.mo, 22
∗ **becker**
as.mo, 22
∗ **datasets**
antibiotics, 12
example_isolates, 50
example_isolates_unclean, 51
intrinsic_resistant, 72
microorganisms, 87
microorganisms.codes, 89
microorganisms.old, 90
rsi_translation, 114
WHONET, 118
∗ **guess**
as.mo, 22
∗ **lancefield**
as.mo, 22
∗ **mo**
as.mo, 22
%like% (like), 80
%like_case% (like), 80
3MRGN (mdro), 82
4MRGN (mdro), 82

ab, *18*
ab (as.ab), 17
ab_*, *4*, *12*, *18*, *116*
ab_atc (ab_property), 5
ab_atc_group1 (ab_property), 5
ab_atc_group2 (ab_property), 5
ab_cid (ab_property), 5
ab_cid(), *6*
ab_class (antibiotic_class_selectors), 15
ab_ddd (ab_property), 5
ab_ddd(), *6*
ab_from_text, 3
ab_from_text(), *19*
ab_group (ab_property), 5
ab_group(), *4*, *116*
ab_info (ab_property), 5
ab_info(), *6*
ab_loinc (ab_property), 5
ab_loinc(), *13*
ab_name (ab_property), 5
ab_name(), *4*, *50*, *68*, *116*
ab_name(AMP), *120*
ab_property, 5
ab_property(), *3*, *13*, *42*, *68*, *105*
ab_synonyms (ab_property), 5
ab_synonyms(), *6*
ab_tradenames (ab_property), 5
ab_tradenames(), *6*
ab_url (ab_property), 5
ab_url(), *6*
age, 8
age(), *9*, *10*
age_groups, 9
age_groups(), *8*, *119*
aminoglycosides (antibiotic_class_selectors), 15
aminoglycosides(), *16*
AMR, 11
anti_join_microorganisms (join), 73
antibiotic_class_selectors, 15
antibiotic_class_selectors(), *53*
antibiotics, *3*, *5–7*, *12*, 12, *13*, *16*, *17*, *19*, *37*, *42*, *50*, *52*, *68*, *70*, *71*, *105*, *120*
antivirals, *13*
antivirals (antibiotics), 12
as.ab, 17
as.ab(), *3*, *5*, *6*, *12*, *17*, *18*, *29*, *71*, *94*, *109*, *114*
as.character(), *81*
as.Date(), *62*
as.disk, 19
as.disk(), *30*, *32*, *109*
as.mic, 21
as.mic(), *30*, *32*, *109*
as.mo, 22
as.mo(), *24*, *25*, *29*, *32*, *37*, *39*, *45*, *50*, *51*, *55*, *73*, *75*, *83*, *87*, *89–96*, *99–101*, *109*, *114*, *119*

as.POSIXlt(), *8*
as.rsi, 27
as.rsi(), *20–22*, *29*, *30*, *42*, *46*, *50*, *51*, *67*, *68*, *105*, *114*, *120*
ATC *(*ab_property*)*, 5
atc_online_ddd *(*atc_online_property*)*, 33
atc_online_groups *(*atc_online_property*)*, 33
atc_online_property, 33
availability, 35

biplot(), *63*
BRMO *(*mdro*)*, 82
brmo *(*mdro*)*, 82
browseURL(), *94*
bug_drug_combinations, 36
bug_drug_combinations(), *37*

carbapenems *(*antibiotic_class_selectors*)*, 15
catalogue_of_life, 38
catalogue_of_life_version, 40
catalogue_of_life_version(), *25*, *26*, *38*, *40*, *88–91*, *97*
cephalosporins *(*antibiotic_class_selectors*)*, 15
cephalosporins_1st *(*antibiotic_class_selectors*)*, 15
cephalosporins_2nd *(*antibiotic_class_selectors*)*, 15
cephalosporins_3rd *(*antibiotic_class_selectors*)*, 15
cephalosporins_4th *(*antibiotic_class_selectors*)*, 15
cephalosporins_5th *(*antibiotic_class_selectors*)*, 15
character, *4*, *6*, *18*, *24*, *25*, *74*, *81*, *95*, *105*
chisq.test(), *58–60*
Click here, *26*, *38*, *40*, *88*, *90*, *91*, *97*
count, 41
count(), *105*, *107*
count_all *(*count*)*, 41
count_all(), *42*
count_df *(*count*)*, 41
count_df(), *42*, *68*
count_I *(*count*)*, 41
count_IR *(*count*)*, 41
count_R *(*count*)*, 41
count_R(), *42*
count_resistant *(*count*)*, 41
count_resistant(), *41*, *42*
count_S *(*count*)*, 41
count_SI *(*count*)*, 41
count_SI(), *31*, *42*, *43*, *86*, *107*, *112*
count_susceptible *(*count*)*, 41
count_susceptible(), *31*, *41–43*, *86*, *107*, *112*

data.frame, *13*, *19*, *23*, *24*, *27*, *29*, *35*, *37*, *42*, *47*, *50*, *51*, *55*, *67*, *71*, *72*, *75*, *79*, *83*, *87*, *89*, *90*, *100*, *102*, *103*, *105*, *111*, *114*, *115*, *119*
disk, *19*, *20*, *29*, *30*
disk *(*as.disk*)*, 19
double, *6*, *8*, *62*, *95*, *105*

EUCAST *(*eucast_rules*)*, 45
eucast_exceptional_phenotypes *(*mdro*)*, 82
eucast_rules, 45
eucast_rules(), *30*, *47*, *85*
example_isolates, 50, *118*
example_isolates_unclean, 51

facet_rsi *(*ggplot_rsi*)*, 66
facet_rsi(), *69*
factor, *9*, *21*, *31*, *84*
filter(), *55*, *62*, *83*, *94*, *111*
filter_1st_cephalosporins *(*filter_ab_class*)*, 52
filter_2nd_cephalosporins *(*filter_ab_class*)*, 52
filter_3rd_cephalosporins *(*filter_ab_class*)*, 52
filter_4th_cephalosporins *(*filter_ab_class*)*, 52
filter_5th_cephalosporins *(*filter_ab_class*)*, 52
filter_ab_class, 52
filter_ab_class(), *16*
filter_aminoglycosides *(*filter_ab_class*)*, 52
filter_aminoglycosides(), *53*
filter_carbapenems *(*filter_ab_class*)*, 52
filter_cephalosporins *(*filter_ab_class*)*, 52
filter_first_isolate *(*first_isolate*)*, 54
filter_first_isolate(), *55*, *56*
filter_first_weighted_isolate *(*first_isolate*)*, 54

filter_first_weighted_isolate(), 55, 56
filter_fluoroquinolones (filter_ab_class), 52
filter_glycopeptides (filter_ab_class), 52
filter_macrolides (filter_ab_class), 52
filter_penicillins (filter_ab_class), 52
filter_tetracyclines (filter_ab_class), 52
first_isolate, 54
first_isolate(), 55, 56, 62, 75–77, 105
fisher.test(), 59
fluoroquinolones (antibiotic_class_selectors), 15
format(), 36, 37
format.bug_drug_combinations (bug_drug_combinations), 36
full_join_microorganisms (join), 73

g.test, 58
g.test(), 58
geom_rsi (ggplot_rsi), 66
geom_rsi(), 68
get_episode, 61
get_episode(), 61, 62
get_locale (translate), 116
get_locale(), 6, 23, 37, 42, 68, 94, 105
get_mo_source (mo_source), 99
get_mo_source(), 23, 100
ggplot2, 66, 68
ggplot2::facet_wrap(), 69
ggplot2::geom_text(), 69
ggplot2::scale_fill_manual(), 69
ggplot2::scale_y_continuous(), 69
ggplot2::theme(), 69
ggplot_pca, 63
ggplot_pca(), 65, 103
ggplot_rsi, 66
ggplot_rsi(), 69
ggplot_rsi_predict (resistance_predict), 110
glm(), 113
glycopeptides (antibiotic_class_selectors), 15
grep(), 80, 81
guess_ab_col, 70
guess_ab_col(), 47, 75, 76, 84

inner_join (join), 73
inner_join_microorganisms (join), 73
integer, 6, 8, 20, 42, 95
intrinsic_resistant, 15, 29, 72, 89, 95, 115
is.ab (as.ab), 17
is.disk (as.disk), 19
is.mic (as.mic), 21
is.mo (as.mo), 22
is.rsi (as.rsi), 27
is.rsi.eligible(), 30
is_new_episode (get_episode), 61
is_new_episode(), 54, 56, 61, 62

join, 73

key_antibiotics, 74
key_antibiotics(), 55–57, 76, 77
key_antibiotics_equal (key_antibiotics), 74
key_antibiotics_equal(), 76, 77
kurtosis, 78
kurtosis(), 116

labels_rsi_count (ggplot_rsi), 66
labels_rsi_count(), 68, 69
left_join_microorganisms (join), 73
lifecycle, 4, 6, 8, 10, 18, 20, 21, 25, 31, 34, 36, 37, 42, 48, 53, 57, 60, 62, 65, 69, 71, 74, 77, 79, 79, 80, 81, 84, 93, 95, 101, 103, 106, 109, 112, 116, 117
like, 80
list, 3, 4, 6, 35, 40, 95
lm(), 113
logical, 9, 29, 46, 56, 62, 77, 80, 81, 83

macrolides (antibiotic_class_selectors), 15
matrix, 79, 115
MDR (mdro), 82
mdr_cmi2012 (mdro), 82
mdr_cmi2012(), 84
mdr_tb (mdro), 82
mdr_tb(), 84
mdro, 82
mdro(), 47, 83–85
merge(), 74
mic, 21, 29, 30
mic (as.mic), 21
microorganisms, 15, 24, 25, 27, 39–41, 50, 51, 73, 87, 89–91, 94, 97, 100
microorganisms.codes, 89, 89
microorganisms.old, 25, 90
microorganisms$fullname, 92
microorganisms$mo, 100
mo, 22, 23, 25, 29, 37, 45, 55, 73, 75, 83

mo (as.mo), 22
mo_*, *24*, *25*, *27*, *89*, *91*, *92*, *96*, *99*, *101*, *116*
mo_authors (mo_property), 93
mo_authors(), *95*
mo_class (mo_property), 93
mo_domain (mo_property), 93
mo_domain(), *95*
mo_failures (as.mo), 22
mo_failures(), *24*
mo_family (mo_property), 93
mo_fullname (mo_property), 93
mo_genus (mo_property), 93
mo_genus(), *27*, *99*, *100*
mo_gramstain (mo_property), 93
mo_gramstain(), *27*, *95*, *99*, *100*, *116*
mo_info (mo_property), 93
mo_info(), *95*
mo_is_gram_negative (mo_property), 93
mo_is_gram_negative(), *95*
mo_is_gram_positive (mo_property), 93
mo_is_gram_positive(), *95*
mo_is_intrinsic_resistant (mo_property), 93
mo_is_intrinsic_resistant(), *95*
mo_kingdom (mo_property), 93
mo_kingdom(), *95*
mo_matching_score, 91
mo_matching_score(), *25*, *92*, *96*
mo_name (mo_property), 93
mo_name(), *116*
mo_order (mo_property), 93
mo_phylum (mo_property), 93
mo_property, 93
mo_property(), *87*, *89*, *91*
mo_rank (mo_property), 93
mo_ref (mo_property), 93
mo_ref(), *95*
mo_renamed (as.mo), 22
mo_renamed(), *24*
mo_shortname (mo_property), 93
mo_shortname(), *37*, *95*
mo_snomed (mo_property), 93
mo_snomed(), *87*, *95*
mo_source, 99
mo_species (mo_property), 93
mo_subspecies (mo_property), 93
mo_synonyms (mo_property), 93
mo_taxonomy (mo_property), 93
mo_taxonomy(), *95*
mo_type (mo_property), 93
mo_type(), *116*, *119*
mo_uncertainties (as.mo), 22
mo_uncertainties(), *24*
mo_url (mo_property), 93
mo_url(), *95*
mo_year (mo_property), 93
mo_year(), *95*
mrgn (mdro), 82
mrgn(), *84*
mutate(), *55*, *62*, *83*, *94*, *111*

n_rsi (count), 41
n_rsi(), *42*

pca, 102, *103*
pca(), *64*, *65*, *103*
PDR (mdro), 82
penicillins (antibiotic_class_selectors), 15
plot.resistance_predict (resistance_predict), 110
portion (proportion), 104
prcomp, *103*
prcomp(), *64*, *103*
princomp, *64*
princomp(), *64*
proportion, 104
proportion(), *113*
proportion_*, *44*
proportion_df (proportion), 104
proportion_df(), *105*
proportion_I (proportion), 104
proportion_IR (proportion), 104
proportion_R (proportion), 104
proportion_R(), *105*
proportion_S (proportion), 104
proportion_SI (proportion), 104
proportion_SI(), *31*, *43*, *86*, *105*, *107*, *112*

random, 108
random_disk (random), 108
random_disk(), *109*
random_mic (random), 108
random_mic(), *109*
random_rsi (random), 108
readRDS(), *100*
resistance (proportion), 104
resistance(), *35*, *42*, *104*, *105*
resistance_predict, 110, *111*
right_join_microorganisms (join), 73
rsi, *28*, *31*, *42*, *50*, *67*, *68*, *105*
rsi (as.rsi), 27
rsi_df (proportion), 104
rsi_df(), *42*, *68*, *105*

rsi_predict (resistance_predict), 110
rsi_translation, 29, 109, 114

sample(), 109
scale, 102
scale_rsi_colours (ggplot_rsi), 66
scale_rsi_colours(), 69
scale_y_percent (ggplot_rsi), 66
scale_y_percent(), 69
semi_join_microorganisms (join), 73
set_mo_source (mo_source), 99
set_mo_source(), 23, 89, 100, 101
skewness, 115
skewness(), 79
summarise(), 55, 62, 83, 94, 111
summary(), 111
susceptibility (proportion), 104
susceptibility(), 31, 35, 42, 43, 86, 104–107, 112
Sys.getlocale(), 116

tetracyclines (antibiotic_class_selectors), 15
theme_rsi (ggplot_rsi), 66
theme_rsi(), 69
translate, 6, 94, 95, 116

utils::browseURL(), 6

variable grouping, 62
vector, 18, 24, 25

WHOCC, 117
WHONET, 118
write us an email (see section 'Contact Us'), 4, 65, 69, 80, 103, 109, 112

XDR (mdro), 82
