## Supplementary material for "AMR - An R Package for Working with Antimicrobial Resistance Data": Source package: CITATION

citHeader("To cite our AMR package in publications, please use (for now):")

citEntry(
  entry        = "Article",
  title        = "AMR - An R Package for Working with Antimicrobial Resistance Data",
  author       = "M S Berends and C F Luz and A W Friedrich and B N M Sinha and C J Albers and C Glasner",
  journal      = "bioRxiv",
  publisher    = "Cold Spring Harbor Laboratory",
  year         = 2019,
  url          = "https://doi.org/10.1101/810622",
  textVersion  = "Berends MS, Luz CF et al. (2019). AMR - An R Package for Working with Antimicrobial Resistance Data. bioRxiv, https://doi.org/10.1101/810622"
)

citFooter("The mentioned article is a preprinted version of a manuscript we sent to a journal. Many thanks for using our open-source method to work with microbial and antimicrobial data!")
