## Supplementary material for "AMR - An R Package for Working with Antimicrobial Resistance Data": Source package: datasets.html

Data sets for download / own use


### Data sets for download / own use

All reference data (about microorganisms, antibiotics, R/SI interpretation, EUCAST rules, etc.) in this `AMR` package are reliable, up-to-date and freely available. We continually export our data sets to formats for use in R, SPSS, SAS, Stata and Excel. We also supply tab separated files that are machine-readable and suitable for input in any software program, such as laboratory information systems.

On this page, we explain how to download them and how the structure of the data sets look like.

If you are reading this page from within R, please visit our website, which is automatically updated with every code change.

#### Microorganisms (currently accepted names)

A data set with 67,151 rows and 16 columns, containing the following column names:  
*‘mo’, ‘fullname’, ‘kingdom’, ‘phylum’, ‘class’, ‘order’, ‘family’, ‘genus’, ‘species’, ‘subspecies’, ‘rank’, ‘ref’, ‘species\_id’, ‘source’, ‘prevalence’, ‘snomed’*.

This data set is in R available as `microorganisms`, after you load the `AMR` package.

It was last updated on 3 September 2020 20:59:45 CEST. Find more info about the structure of this data set here.

**Direct download links:**

- Download as R file (2.7 MB)
- Download as Excel file (6.1 MB)
- Download as plain text file (13.3 MB)
- Download as SAS file (26.2 MB)
- Download as SPSS file (28.2 MB)
- Download as Stata file (25.2 MB)

##### Source

Our full taxonomy of microorganisms is based on the authoritative and comprehensive:

- Catalogue of Life (included version: 2019)
- List of Prokaryotic names with Standing in Nomenclature (LPSN, included version: May 2020)

##### Example content

Included (sub)species per taxonomic kingdom:

| Kingdom | Number of (sub)species |
| --- | --- |
| (unknown kingdom) | 1 |
| Animalia | 2,153 |
| Archaea | 697 |
| Bacteria | 19,244 |
| Chromista | 32,164 |
| Fungi | 9,582 |

Example rows when filtering on genus *Escherichia*:

| mo | fullname | kingdom | phylum | class | order | family | genus | species | subspecies | rank | ref | species\_id | source | prevalence | snomed |
| --- | --- | --- | --- | --- | --- | --- | --- | --- | --- | --- | --- | --- | --- | --- | --- |
| B\_ESCHR | Escherichia | Bacteria | Proteobacteria | Gammaproteobacteria | Enterobacterales | Enterobacteriaceae | Escherichia |  |  | genus |  | bc4fdde6867d5ecfc728000b0bfb49a3 | CoL | 1 | 64735005 |
| B\_ESCHR\_ALBR | Escherichia albertii | Bacteria | Proteobacteria | Gammaproteobacteria | Enterobacterales | Enterobacteriaceae | Escherichia | albertii |  | species | Huys et al., 2003 | 36618b1ed3b8b7e5a61f40eb9386e63c | CoL | 1 | 419388003 |
| B\_ESCHR\_COLI | Escherichia coli | Bacteria | Proteobacteria | Gammaproteobacteria | Enterobacterales | Enterobacteriaceae | Escherichia | coli |  | species | Castellani et al., 1919 | 3254b3db31bf16fdde669ac57bf8c4fe | CoL | 1 | 112283007 |
| B\_ESCHR\_FRGS | Escherichia fergusonii | Bacteria | Proteobacteria | Gammaproteobacteria | Enterobacterales | Enterobacteriaceae | Escherichia | fergusonii |  | species | Farmer et al., 1985 | 82d98b10c456ce5f4c8c515f4e1567e2 | CoL | 1 | 72461005 |
| B\_ESCHR\_HRMN | Escherichia hermannii | Bacteria | Proteobacteria | Gammaproteobacteria | Enterobacterales | Enterobacteriaceae | Escherichia | hermannii |  | species | Brenner et al., 1983 | b16086aee36e3b46b565510083ab4b65 | CoL | 1 | 85786000 |
| B\_ESCHR\_MRMT | Escherichia marmotae | Bacteria | Proteobacteria | Gammaproteobacteria | Enterobacterales | Enterobacteriaceae | Escherichia | marmotae |  | species | Liu et al., 2015 | 792928 | DSMZ | 1 |  |

#### Microorganisms (previously accepted names)

A data set with 12,708 rows and 4 columns, containing the following column names:  
*‘fullname’, ‘fullname\_new’, ‘ref’, ‘prevalence’*.

**Note:** remember that the ‘ref’ columns contains the scientific reference to the old taxonomic entries, i.e. of column *‘fullname’*. For the scientific reference of the new names, i.e. of column *‘fullname\_new’*, see the `microorganisms` data set.

This data set is in R available as `microorganisms.old`, after you load the `AMR` package.

It was last updated on 28 May 2020 11:17:56 CEST. Find more info about the structure of this data set here.

**Direct download links:**

- Download as R file (0.3 MB)
- Download as Excel file (0.4 MB)
- Download as plain text file (0.8 MB)
- Download as SAS file (1.9 MB)
- Download as SPSS file (1.9 MB)
- Download as Stata file (1.8 MB)

##### Source

This data set contains old, previously accepted taxonomic names. The data sources are the same as the `microorganisms` data set:

- Catalogue of Life (included version: 2019)
- List of Prokaryotic names with Standing in Nomenclature (LPSN, included version: May 2020)

##### Example content

Example rows when filtering on *Escherichia*:

| fullname | fullname\_new | ref | prevalence |
| --- | --- | --- | --- |
| Escherichia adecarboxylata | Leclercia adecarboxylata | Leclerc, 1962 | 1 |
| Escherichia blattae | Shimwellia blattae | Burgess et al., 1973 | 1 |
| Escherichia vulneris | Pseudescherichia vulneris | Brenner et al., 1983 | 1 |

#### Antibiotic agents

A data set with 455 rows and 14 columns, containing the following column names:  
*‘ab’, ‘atc’, ‘cid’, ‘name’, ‘group’, ‘atc\_group1’, ‘atc\_group2’, ‘abbreviations’, ‘synonyms’, ‘oral\_ddd’, ‘oral\_units’, ‘iv\_ddd’, ‘iv\_units’, ‘loinc’*.

This data set is in R available as `antibiotics`, after you load the `AMR` package.

It was last updated on 24 September 2020 00:50:35 CEST. Find more info about the structure of this data set here.

**Direct download links:**

- Download as R file (31 kB)
- Download as Excel file (65 kB)
- Download as plain text file (0.1 MB)
- Download as SAS file (1.8 MB)
- Download as SPSS file (1.3 MB)
- Download as Stata file (0.3 MB)

##### Source

This data set contains all EARS-Net and ATC codes gathered from WHO and WHONET, and all compound IDs from PubChem. It also contains all brand names (synonyms) as found on PubChem and Defined Daily Doses (DDDs) for oral and parenteral administration.

- ATC/DDD index from WHO Collaborating Centre for Drug Statistics Methodology (note: this may not be used for commercial purposes, but is frelly available from the WHO CC website for personal use)
- PubChem by the US National Library of Medicine
- WHONET software 2019

##### Example content

| ab | atc | cid | name | group | atc\_group1 | atc\_group2 | abbreviations | synonyms | oral\_ddd | oral\_units | iv\_ddd | iv\_units | loinc |
| --- | --- | --- | --- | --- | --- | --- | --- | --- | --- | --- | --- | --- | --- |
| AMK | J01GB06 | 37768 | Amikacin | Aminoglycosides | Aminoglycoside antibacterials | Other aminoglycosides | ak, ami, amik, … | amicacin, amikacillin, amikacin, … |  |  | 1.0 | g | 13546-7, 15098-7, 17798-0, … |
| AMX | J01CA04 | 33613 | Amoxicillin | Beta-lactams/penicillins | Beta-lactam antibacterials, penicillins | Penicillins with extended spectrum | ac, amox, amx | actimoxi, amoclen, amolin, … | 1.5 | g | 3.0 | g | 16365-9, 25274-2, 3344-9, … |
| AMC | J01CR02 | 23665637 | Amoxicillin/clavulanic acid | Beta-lactams/penicillins | Beta-lactam antibacterials, penicillins | Combinations of penicillins, incl. beta-lactamase inhibitors | a/c, amcl, aml, … | amocla, amoclan, amoclav, … | 1.5 | g | 3.0 | g |  |
| AMP | J01CA01 | 6249 | Ampicillin | Beta-lactams/penicillins | Beta-lactam antibacterials, penicillins | Penicillins with extended spectrum | am, amp, ampi | acillin, adobacillin, amblosin, … | 2.0 | g | 6.0 | g | 21066-6, 3355-5, 33562-0, … |
| AZM | J01FA10 | 447043 | Azithromycin | Macrolides/lincosamides | Macrolides, lincosamides and streptogramins | Macrolides | az, azi, azit, … | aritromicina, azasite, azenil, … | 0.3 | g | 0.5 | g | 16420-2, 25233-8 |
| CZO | J01DB04 | 33255 | Cefazolin | Cephalosporins (1st gen.) | Other beta-lactam antibacterials | First-generation cephalosporins | cfz, cfzl, cz, … | atirin, cefamezin, cefamezine, … |  |  | 3.0 | g | 16566-2, 25235-3, 3442-1, … |

#### Antiviral agents

A data set with 102 rows and 9 columns, containing the following column names:  
*‘atc’, ‘cid’, ‘name’, ‘atc\_group’, ‘synonyms’, ‘oral\_ddd’, ‘oral\_units’, ‘iv\_ddd’, ‘iv\_units’*.

This data set is in R available as `antivirals`, after you load the `AMR` package.

It was last updated on 29 August 2020 21:53:07 CEST. Find more info about the structure of this data set here.

**Direct download links:**

- Download as R file (5 kB)
- Download as Excel file (14 kB)
- Download as plain text file (16 kB)
- Download as SAS file (80 kB)
- Download as SPSS file (68 kB)
- Download as Stata file (67 kB)

##### Source

This data set contains all ATC codes gathered from WHO and all compound IDs from PubChem. It also contains all brand names (synonyms) as found on PubChem and Defined Daily Doses (DDDs) for oral and parenteral administration.

- ATC/DDD index from WHO Collaborating Centre for Drug Statistics Methodology (note: this may not be used for commercial purposes, but is frelly available from the WHO CC website for personal use)
- PubChem by the US National Library of Medicine

##### Example content

| atc | cid | name | atc\_group | synonyms | oral\_ddd | oral\_units | iv\_ddd | iv\_units |
| --- | --- | --- | --- | --- | --- | --- | --- | --- |
| J05AF06 | 441300 | Abacavir | Nucleoside and nucleotide reverse transcriptase inhibitors | Abacavir, Abacavir sulfate, Ziagen | 0.6 | g |  |  |
| J05AB01 | 135398513 | Aciclovir | Nucleosides and nucleotides excl. reverse transcriptase inhibitors | Acicloftal, Aciclovier, Aciclovir, … | 4.0 | g | 4 | g |
| J05AF08 | 60871 | Adefovir dipivoxil | Nucleoside and nucleotide reverse transcriptase inhibitors | Adefovir di ester, Adefovir dipivoxil, Adefovir Dipivoxil, … | 10.0 | mg |  |  |
| J05AE05 | 65016 | Amprenavir | Protease inhibitors | Agenerase, Amprenavir, Amprenavirum, … | 1.2 | g |  |  |
| J05AP06 | 16076883 | Asunaprevir | Antivirals for treatment of HCV infections | Asunaprevir, Sunvepra |  |  |  |  |
| J05AE08 | 148192 | Atazanavir | Protease inhibitors | Atazanavir, Atazanavir Base, Latazanavir, … | 0.3 | g |  |  |

#### Intrinsic bacterial resistance

A data set with 93,892 rows and 2 columns, containing the following column names:  
*‘microorganism’, ‘antibiotic’*.

This data set is in R available as `intrinsic_resistant`, after you load the `AMR` package.

It was last updated on 24 September 2020 00:50:35 CEST. Find more info about the structure of this data set here.

**Direct download links:**

- Download as R file (67 kB)
- Download as Excel file (0.9 MB)
- Download as plain text file (3.5 MB)
- Download as SAS file (7.1 MB)
- Download as SPSS file (7.9 MB)
- Download as Stata file (7 MB)

##### Source

This data set contains all defined intrinsic resistance by EUCAST of all bug-drug combinations, and is based on ‘’EUCAST Expert Rules’ and ‘EUCAST Intrinsic Resistance and Unusual Phenotypes’’, v3.2 from 2020.

##### Example content

Example rows when filtering on *Enterobacter cloacae*:

| microorganism | antibiotic |
| --- | --- |
| Enterobacter cloacae | Amoxicillin |
| Enterobacter cloacae | Amoxicillin/clavulanic acid |
| Enterobacter cloacae | Ampicillin |
| Enterobacter cloacae | Ampicillin/sulbactam |
| Enterobacter cloacae | Avoparcin |
| Enterobacter cloacae | Azithromycin |
| Enterobacter cloacae | Cefadroxil |
| Enterobacter cloacae | Cefazolin |
| Enterobacter cloacae | Cefoxitin |
| Enterobacter cloacae | Cephalexin |
| Enterobacter cloacae | Cephalothin |
| Enterobacter cloacae | Clarithromycin |
| Enterobacter cloacae | Clindamycin |
| Enterobacter cloacae | Cycloserine |
| Enterobacter cloacae | Dalbavancin |
| Enterobacter cloacae | Dirithromycin |
| Enterobacter cloacae | Erythromycin |
| Enterobacter cloacae | Flurithromycin |
| Enterobacter cloacae | Fusidic acid |
| Enterobacter cloacae | Josamycin |
| Enterobacter cloacae | Lincomycin |
| Enterobacter cloacae | Linezolid |
| Enterobacter cloacae | Midecamycin |
| Enterobacter cloacae | Miocamycin |
| Enterobacter cloacae | Norvancomycin |
| Enterobacter cloacae | Oleandomycin |
| Enterobacter cloacae | Oritavancin |
| Enterobacter cloacae | Benzylpenicillin |
| Enterobacter cloacae | Pirlimycin |
| Enterobacter cloacae | Pristinamycin |
| Enterobacter cloacae | Quinupristin/dalfopristin |
| Enterobacter cloacae | Ramoplanin |
| Enterobacter cloacae | Rifampicin |
| Enterobacter cloacae | Rokitamycin |
| Enterobacter cloacae | Roxithromycin |
| Enterobacter cloacae | Spiramycin |
| Enterobacter cloacae | Tedizolid |
| Enterobacter cloacae | Teicoplanin |
| Enterobacter cloacae | Teicoplanin-macromethod |
| Enterobacter cloacae | Telavancin |
| Enterobacter cloacae | Telithromycin |
| Enterobacter cloacae | Thiacetazone |
| Enterobacter cloacae | Troleandomycin |
| Enterobacter cloacae | Vancomycin |

#### Interpretation from MIC values / disk diameters to R/SI

A data set with 18,650 rows and 10 columns, containing the following column names:  
*‘guideline’, ‘method’, ‘site’, ‘mo’, ‘ab’, ‘ref\_tbl’, ‘disk\_dose’, ‘breakpoint\_S’, ‘breakpoint\_R’, ‘uti’*.

This data set is in R available as `rsi_translation`, after you load the `AMR` package.

It was last updated on 29 July 2020 13:12:34 CEST. Find more info about the structure of this data set here.

**Direct download links:**

- Download as R file (32 kB)
- Download as Excel file (0.6 MB)
- Download as plain text file (1.5 MB)
- Download as SAS file (3.2 MB)
- Download as SPSS file (3.4 MB)
- Download as Stata file (3 MB)

##### Source

This data set contains interpretation rules for MIC values and disk diffusion diameters. Included guidelines are CLSI (2010-2019) and EUCAST (2011-2020).

##### Example content

| guideline | method | site | mo | ab | ref\_tbl | disk\_dose | breakpoint\_S | breakpoint\_R | uti |
| --- | --- | --- | --- | --- | --- | --- | --- | --- | --- |
| EUCAST 2020 | DISK |  | Enterobacterales | Amoxicillin/clavulanic acid | Enterobacterales | 20-10ug | 19 | 19 | FALSE |
| EUCAST 2020 | DISK | UTI | Enterobacterales | Amoxicillin/clavulanic acid | Enterobacterales | 20-10ug | 16 | 16 | TRUE |
| EUCAST 2020 | MIC |  | Enterobacterales | Amoxicillin/clavulanic acid | Enterobacterales |  | 8 | 8 | FALSE |
| EUCAST 2020 | MIC | UTI | Enterobacterales | Amoxicillin/clavulanic acid | Enterobacterales |  | 32 | 32 | TRUE |
| EUCAST 2020 | MIC |  | Actinomyces | Amoxicillin/clavulanic acid | Anaerobes, Grampositive |  | 4 | 8 | FALSE |
| EUCAST 2020 | MIC |  | Bacteroides | Amoxicillin/clavulanic acid | Anaerobes, Gramnegative |  | 4 | 8 | FALSE |
