## Supplementary material for "AMR - An R Package for Working with Antimicrobial Resistance Data": Source package: datasets.R

```
## ----setup, include = FALSE, results = 'markup'-------------------------------
knitr::opts_chunk$set(
  warning = FALSE,
  collapse = TRUE,
  comment = "#",
  fig.width = 7.5,
  fig.height = 5
)

library(AMR)
library(dplyr)

options(knitr.kable.NA = '')

structure_txt <- function(dataset) {
  paste0("A data set with ",
         format(nrow(dataset), big.mark = ","), " rows and ", 
         ncol(dataset), " columns, containing the following column names:  \n*",
         paste0("'", colnames(dataset), "'", collapse = ", "), "*.")
}

download_txt <- function(filename) {
  msg <- paste0("It was last updated on ", 
                trimws(format(file.mtime(paste0("../data/", filename, ".rda")), "%e %B %Y %H:%M:%S %Z")), 
                ". Find more info about the structure of this data set [here](https://msberends.github.io/AMR/reference/", ifelse(filename == "antivirals", "antibiotics", filename), ".html).\n")
  github_base <- "https://github.com/msberends/AMR/raw/master/data-raw/"
  filename <- paste0("../data-raw/", filename)
  txt <- paste0(filename, ".txt")
  rds <- paste0(filename, ".rds")
  spss <- paste0(filename, ".sav")
  stata <- paste0(filename, ".dta")
  sas <- paste0(filename, ".sas")
  excel <- paste0(filename, ".xlsx")
  create_txt <- function(filename, type, software) {
    paste0("* Download as [", software, " file](", github_base, filename, ") (", AMR:::formatted_filesize(filename), ")  \n")
  }

  if (any(file.exists(rds),
          file.exists(excel),
          file.exists(txt),
          file.exists(sas),
          file.exists(spss),
          file.exists(stata))) {
    msg <- c(msg, "\n**Direct download links:**\n\n")
  }
  if (file.exists(rds)) msg <- c(msg, create_txt(rds, "rds", "R"))
  if (file.exists(excel)) msg <- c(msg, create_txt(excel, "xlsx", "Excel"))
  if (file.exists(txt)) msg <- c(msg, create_txt(txt, "txt", "plain text"))
  if (file.exists(sas)) msg <- c(msg, create_txt(sas, "sas", "SAS"))
  if (file.exists(spss)) msg <- c(msg, create_txt(spss, "sav", "SPSS"))
  if (file.exists(stata)) msg <- c(msg, create_txt(stata, "dta", "Stata"))
  paste0(msg, collapse = "")
}

print_df <- function(x, rows = 6) {
  x %>% 
    head(n = rows) %>% 
    mutate_all(function(x) {
      if (is.list(x)) {
        sapply(x, function(y) {
          if (length(y) > 3) {
            paste0(paste(y[1:3], collapse = ", "), ", ...")
          } else if (length(y) == 0 || all(is.na(y))) {
            ""
          } else {
            paste(y, collapse = ", ")
          }
        })
      } else {
        x
      }
    }) %>%
    knitr::kable(align = "c")
}


## ---- echo = FALSE------------------------------------------------------------
microorganisms %>% 
  pull(kingdom) %>% 
  table() %>% 
  as.data.frame() %>% 
  mutate(Freq = format(Freq, big.mark = ",")) %>% 
  setNames(c("Kingdom", "Number of (sub)species")) %>% 
  print_df()

## ---- echo = FALSE------------------------------------------------------------
microorganisms %>%
  filter(genus == "Escherichia") %>% 
  print_df()

## ---- echo = FALSE------------------------------------------------------------
microorganisms.old %>%
  filter(fullname %like% "^Escherichia") %>% 
  print_df()

## ---- echo = FALSE------------------------------------------------------------
antibiotics %>%
  filter(ab %in% colnames(example_isolates)) %>% 
  print_df()

## ---- echo = FALSE------------------------------------------------------------
antivirals %>%
  print_df()

## ---- echo = FALSE------------------------------------------------------------
intrinsic_resistant %>%
  filter(microorganism == "Enterobacter cloacae") %>% 
  print_df(rows = Inf)

## ---- echo = FALSE------------------------------------------------------------
rsi_translation %>% 
  mutate(ab = ab_name(ab), mo = mo_name(mo)) %>% 
  print_df()
```
