## Supplementary material for "AMR - An R Package for Working with Antimicrobial Resistance Data": Source package: NEWS.md

# AMR 1.5.0

*Note: the rules of 'EUCAST Clinical Breakpoints v11.0 (2021)' will be added in the next release, to be expected in February/March 2021.*

### New
* Functions `get_episode()` and `is_new_episode()` to determine (patient) episodes which are not necessarily based on microorganisms. The `get_episode()` function returns the index number of the episode per group, while the `is_new_episode()` function returns values `TRUE`/`FALSE` to indicate whether an item in a vector is the start of a new episode. They also support `dplyr`s grouping (i.e. using `group_by()`):
  ```r
  library(dplyr)
  example_isolates %>%
    group_by(patient_id, hospital_id) %>%
    filter(is_new_episode(date, episode_days = 60))
  ```
* Functions `mo_is_gram_negative()` and `mo_is_gram_positive()` as wrappers around `mo_gramstain()`. They always return `TRUE` or `FALSE` (except when the input is `NA` or the MO code is `UNKNOWN`), thus always return `FALSE` for species outside the taxonomic kingdom of Bacteria.
* Function `mo_is_intrinsic_resistant()` to test for intrinsic resistance, based on [EUCAST Intrinsic Resistance and Unusual Phenotypes v3.2](https://www.eucast.org/expert_rules_and_intrinsic_resistance/) from 2020.
* Functions `random_mic()`, `random_disk()` and `random_rsi()` for random value generation. The functions `random_mic()` and `random_disk()` take microorganism names and antibiotic names as input to make generation more realistic.

### Changed
* New argument `ampc_cephalosporin_resistance` in `eucast_rules()` to correct for AmpC de-repressed cephalosporin-resistant mutants
* Interpretation of antimicrobial resistance - `as.rsi()`:
  * Reference data used for `as.rsi()` can now be set by the user, using the `reference_data` argument. This allows for using own interpretation guidelines. The user-set data must have the same structure as `rsi_translation`.
  * Better determination of disk zones and MIC values when running `as.rsi()` on a data.frame
  * Fix for using `as.rsi()` on a data.frame in older R versions
  * `as.rsi()` on a data.frame will not print a message anymore if the values are already clean R/SI values
  * If using `as.rsi()` on MICs or disk diffusion while there is intrinsic antimicrobial resistance, a warning will be thrown to remind about this
  * Fix for using `as.rsi()` on a `data.frame` that only contains one column for antibiotic interpretations
* Some functions are now context-aware when used inside `dplyr` verbs, such as `filter()`, `mutate()` and `summarise()`. This means that then the data argument does not need to be set anymore. This is the case for the new functions:
  * `mo_is_gram_negative()`
  * `mo_is_gram_positive()`
  * `mo_is_intrinsic_resistant()`
  
  ... and for the existing functions:
  * `first_isolate()`,
  * `key_antibiotics()`,
  * `mdro()`,
  * `brmo()`,
  * `mrgn()`,
  * `mdr_tb()`,
  * `mdr_cmi2012()`,
  * `eucast_exceptional_phenotypes()`
  
  ```r
  # to select first isolates that are Gram-negative 
  # and view results of cephalosporins and aminoglycosides:
  library(dplyr)
  example_isolates %>%
    filter(first_isolate(), mo_is_gram_negative()) %>% 
    select(mo, cephalosporins(), aminoglycosides()) %>% 
    as_tibble()
  ```
* For antibiotic selection functions (such as `cephalosporins()`, `aminoglycosides()`) to select columns based on a certain antibiotic group, the dependency on the `tidyselect` package was removed, meaning that they can now also be used without the need to have this package installed and now also work in base R function calls (they rely on R 3.2 or later):
  ```r
  # above example in base R:
  example_isolates[which(first_isolate() & mo_is_gram_negative()),
                   c("mo", cephalosporins(), aminoglycosides())]
  ```
* For all function arguments in the code, it is now defined what the exact type of user input should be (inspired by the [`typed`](https://github.com/moodymudskipper/typed) package). If the user input for a certain function does not meet the requirements for a specific argument (such as the class or length), an informative error will be thrown. This makes the package more robust and the use of it more reproducible and reliable. In total, more than 420 arguments were defined.
* Fix for `set_mo_source()`, that previously would not remember the file location of the original file
* Deprecated function `p_symbol()` that not really fits the scope of this package. It will be removed in a future version. See [here](https://github.com/msberends/AMR/blob/v1.4.0/R/p_symbol.R) for the source code to preserve it.
* Updated coagulase-negative staphylococci determination with Becker *et al.* 2020 (PMID 32056452), meaning that the species *S. argensis*, *S. caeli*, *S. debuckii*, *S. edaphicus* and *S. pseudoxylosus* are now all considered CoNS
* Fix for using argument `reference_df` in `as.mo()` and `mo_*()` functions that contain old microbial codes (from previous package versions)
* Fixed a bug where `mo_uncertainties()` would not return the results based on the MO matching score
* Fixed a bug where `as.mo()` would not return results for known laboratory codes for microorganisms
* Fixed a bug where `as.ab()` would sometimes fail
* Better tibble printing for MIC values
* Fix for plotting MIC values with `plot()`
* Added `plot()` generic to class `<disk>`
* LA-MRSA and CA-MRSA are now recognised as an abbreviation for *Staphylococcus aureus*, meaning that e.g. `mo_genus("LA-MRSA")` will return `"Staphylococcus"` and `mo_is_gram_positive("LA-MRSA")` will return `TRUE`.
* Fix for printing class <mo> in tibbles when all values are `NA`
* Fix for `mo_shortname()` when the input contains `NA`
* If `as.mo()` takes more than 30 seconds, some suggestions will be done to improve speed

### Other
* All messages and warnings thrown by this package now break sentences on whole words
* More extensive unit tests
* Internal calls to `options()` were all removed in favour of a new internal environment `pkg_env`
* Improved internal type setting (among other things: replaced all `sapply()` calls with `vapply()`)
* Added CodeFactor as a continuous code review to this package: <https://www.codefactor.io/repository/github/msberends/amr/>
* Added Dr. Rogier Schade as contributor

# AMR 1.4.0

### New
* Support for 'EUCAST Expert Rules' / 'EUCAST Intrinsic Resistance and Unusual Phenotypes' version 3.2 of May 2020. With this addition to the previously implemented version 3.1 of 2016, the `eucast_rules()` function can now correct for more than 180 different antibiotics and the `mdro()` function can determine multidrug resistance based on more than 150 different antibiotics. All previously implemented versions of the EUCAST rules are now maintained and kept available in this package. The `eucast_rules()` function consequently gained the arguments `version_breakpoints` (at the moment defaults to v10.0, 2020) and `version_expertrules` (at the moment defaults to v3.2, 2020). The `example_isolates` data set now also reflects the change from v3.1 to v3.2. The `mdro()` function now accepts `guideline == "EUCAST3.1"` and `guideline == "EUCAST3.2"`.
* A new vignette and website page with info about all our public and freely available data sets, that can be downloaded as flat files or in formats for use in R, SPSS, SAS, Stata and Excel: https://msberends.github.io/AMR/articles/datasets.html
* Data set `intrinsic_resistant`. This data set contains all bug-drug combinations where the 'bug' is intrinsic resistant to the 'drug' according to the latest EUCAST insights. It contains just two columns: `microorganism` and `antibiotic`.

  Curious about which enterococci are actually intrinsic resistant to vancomycin?
  
  ```r
  library(AMR)
  library(dplyr)
  intrinsic_resistant %>%
    filter(antibiotic == "Vancomycin", microorganism %like% "Enterococcus") %>% 
    pull(microorganism)
  #> [1] "Enterococcus casseliflavus" "Enterococcus gallinarum"   
  ```
* Support for veterinary ATC codes
* Support for skimming classes `<rsi>`, `<mic>`, `<disk>` and `<mo>` with the `skimr` package 

### Changed
* Although advertised that this package should work under R 3.0.0, we still had a dependency on R 3.6.0. This is fixed, meaning that our package should now work under R 3.0.0.
* Improvements for `as.rsi()`:
  * Support for using `dplyr`'s `across()` to interpret MIC values or disk zone diameters, which also automatically determines the column with microorganism names or codes.
    ```r
    # until dplyr 1.0.0
    your_data %>% mutate_if(is.mic, as.rsi)
    your_data %>% mutate_if(is.disk, as.rsi)
  
    # since dplyr 1.0.0
    your_data %>% mutate(across(where(is.mic), as.rsi))
    your_data %>% mutate(across(where(is.disk), as.rsi))
    ```
  * Cleaning columns in a data.frame now allows you to specify those columns with tidy selection, e.g. `as.rsi(df, col1:col9)`
  * Big speed improvement for interpreting MIC values and disk zone diameters. When interpreting 5,000 MIC values of two antibiotics (10,000 values in total), our benchmarks showed a total run time going from 80.7-85.1 seconds to 1.8-2.0 seconds.
  * Added argument 'add_intrinsic_resistance' (defaults to `FALSE`), that considers intrinsic resistance according to EUCAST
  * Fixed a bug where in EUCAST rules the breakpoint for R would be interpreted as ">=" while this should have been "<"
* Added intelligent data cleaning to `as.disk()`, so numbers can also be extracted from text and decimal numbers will always be rounded up:
  ```r
  as.disk(c("disk zone: 23.4 mm", 23.4))
  #> Class <disk>
  #> [1] 24 24
  ```
* Improvements for `as.mo()`:
  * A completely new matching score for ambiguous user input, using `mo_matching_score()`. Any user input value that could mean more than one taxonomic entry is now considered 'uncertain'. Instead of a warning, a message will be thrown and the accompanying `mo_uncertainties()` has been changed completely; it now prints all possible candidates with their matching score.
  * Big speed improvement for already valid microorganism ID. This also means an significant speed improvement for using `mo_*` functions like `mo_name()` on microoganism IDs.
  * Added argument `ignore_pattern` to `as.mo()` which can also be given to `mo_*` functions like `mo_name()`, to exclude known non-relevant input from analysing. This can also be set with the option `AMR_ignore_pattern`.
* `get_locale()` now uses at default `Sys.getenv("LANG")` or, if `LANG` is not set, `Sys.getlocale()`. This can be overwritten by setting the option `AMR_locale`.
* Big speed improvement for `eucast_rules()`
* Overall speed improvement by tweaking joining functions
* Function `mo_shortname()` now returns the genus for input where the species is unknown
* BORSA is now recognised as an abbreviation for *Staphylococcus aureus*, meaning that e.g. `mo_genus("BORSA")` will return "Staphylococcus"
* Added a feature from AMR 1.1.0 and earlier again, but now without other package dependencies: `tibble` printing support for classes `<rsi>`, `<mic>`, `<disk>`, `<ab>` and `<mo>`. When using `tibble`s containing antimicrobial columns (class `<rsi>`), "S" will print in green, "I" will print in yellow and "R" will print in red. Microbial IDs (class `<mo>`) will emphasise on the genus and species, not on the kingdom.
* Names of antiviral agents in data set `antivirals` now have a starting capital letter, like it is the case in the `antibiotics` data set
* Updated the documentation of the `WHONET` data set to clarify that all patient names are fictitious
* Small `as.ab()` algorithm improvements
* Fix for combining MIC values with raw numbers, i.e. `c(as.mic(2), 2)` previously failed but now returns a valid MIC class
* `ggplot_rsi()` and `geom_rsi()` gained arguments `minimum` and `language`, to influence the internal use of `rsi_df()`
* Changes in the `antibiotics` data set:
  * Updated oral and parental DDDs from the WHOCC
  * Added abbreviation "piptazo" to 'Piperacillin/tazobactam' (TZP)
  * 'Penicillin G' (for intravenous use) is now named 'Benzylpenicillin' (code `PEN`)
  * 'Penicillin V' (for oral use, code `PNV`) was removed, since its actual entry 'Phenoxymethylpenicillin' (code `PHN`) already existed
  * The group name (`antibiotics$group`) of 'Linezolid' (`LNZ`), 'Cycloserine' (`CYC`), 'Tedizolid' (`TZD`) and 'Thiacetazone' (`THA`) is now "Oxazolidinones" instead of "Other antibacterials"
* Added support for using `unique()` on classes `<rsi>`, `<mic>`, `<disk>`, `<ab>` and `<mo>`
* Added argument `excess` to the `kurtosis()` function (defaults to `FALSE`), to return the *excess kurtosis*, defined as the kurtosis minus three.

### Other
* Removed functions `portion_R()`, `portion_S()` and `portion_I()` that were deprecated since version 0.9.0 (November 2019) and were replaced with `proportion_R()`, `proportion_S()` and `proportion_I()`
* Removed unnecessary references to the `base` package
* Added packages that could be useful for some functions to the `Suggests` field of the `DESCRIPTION` file

# AMR 1.3.0

### New
* Function `ab_from_text()` to retrieve antimicrobial drug names, doses and forms of administration from clinical texts in e.g. health care records, which also corrects for misspelling since it uses `as.ab()` internally
* [Tidyverse selection helpers](https://tidyselect.r-lib.org/reference/language.html) for antibiotic classes, that help to select the columns of antibiotics that are of a specific antibiotic class, without the need to define the columns or antibiotic abbreviations. They can be used in any function that allows selection helpers, like `dplyr::select()` and `tidyr::pivot_longer()`:
  ```r
  library(dplyr)
  
  # Columns 'IPM' and 'MEM' are in the example_isolates data set
  example_isolates %>% 
    select(carbapenems())
  #> Selecting carbapenems: `IPM` (imipenem), `MEM` (meropenem)
  ```
* Added `mo_domain()` as an alias to `mo_kingdom()`
* Added function `filter_penicillins()` to filter isolates on a specific result in any column with a name in the antimicrobial 'penicillins' class (more specific: ATC subgroup *Beta-lactam antibacterials, penicillins*)
* Added official antimicrobial names to all `filter_ab_class()` functions, such as `filter_aminoglycosides()`
* Added antibiotics code "FOX1" for cefoxitin screening (abbreviation "cfsc") to the `antibiotics` data set
* Added Monuril as trade name for fosfomycin
* Added argument `conserve_capped_values` to `as.rsi()` for interpreting MIC values - it makes sure that values starting with "<" (but not "<=") will always return "S" and values starting with ">" (but not ">=") will always return "R". The default behaviour of `as.rsi()` has not changed, so you need to specifically do `as.rsi(..., conserve_capped_values = TRUE)`.

### Changed
* Big speed improvement for using any function on microorganism codes from earlier package versions (prior to `AMR` v1.2.0), such as `as.mo()`, `mo_name()`, `first_isolate()`, `eucast_rules()`, `mdro()`, etc.

  As a consequence, very old microbial codes (from `AMR` v0.5.0 and lower) **are not supported anymore**. Use `as.mo()` on your microorganism names or codes to transform them to current abbreviations used in this package.
* Improvements for `susceptibility()` and `resistance()` and all `count_*()`, `proportion_*()` functions:
  * 95% speed improvement by using other base R functions for calculation
  * Using unexisting columns wil now return an error instead of dropping them silently
  * Using variables for column names (as well as selectors like `dplyr::all_of()`) now works again
* Improvements for `as.ab()`:
  * Dramatic improvement of the algorithm behind `as.ab()`, making many more input errors translatable, such as digitalised health care records, using too few or too many vowels or consonants and many more
  * Added progress bar
  * Fixed a bug where `as.ab()` would return an error on invalid input values
  * The `as.ab()` function will now throw a note if more than 1 antimicrobial drug could be retrieved from a single input value.
* Fixed a bug where `eucast_rules()` would not work on a tibble when the `tibble` or `dplyr` package was loaded
* Fixed a bug for CLSI 2019 guidelines (using `as.rsi()`), that also included results for animals. It now only contains interpretation guidelines for humans.
* All `*_join_microorganisms()` functions and `bug_drug_combinations()` now return the original data class (e.g. `tibble`s and `data.table`s)
* For functions `rsi_df()`, `proportion_df()` and `count_df()`:
  * Fixed a bug for using grouped versions
  * Fixed a bug where not all different antimicrobial results were added as rows
  * Fixed a bug when only calculating counts (`count_df()`) when all antibiotics in the data set have only `NA`s
* Improved auto-determination for columns of types `<mo>` and `<Date>`
* Fixed a bug in `bug_drug_combinations()` for when only one antibiotic was in the input data
* Changed the summary for class `<rsi>`, to highlight the %SI vs. %R
* Improved error handling, giving more useful info when functions return an error
* Any progress bar will now only show in interactive mode (i.e. not in R Markdown)
* Speed improvement for `mdro()` and `filter_ab_class()`
* New option `arrows_textangled` for `ggplot_pca()` to indicate whether the text at the end of the arrows should be angled (defaults to `TRUE`, as it was in previous versions)
* Added parenteral DDD to benzylpenicillin
* Fixed a bug where `as.mic()` could not handle dots without a leading zero (like `"<=.25`)

### Other
* Moved primary location of this project from GitLab to [GitHub](https://github.com/msberends/AMR), giving us native support for automated syntax checking without being dependent on external services such as AppVeyor and Travis CI.

# AMR 1.2.0

### Breaking 
* Removed code dependency on all other R packages, making this package fully independent of the development process of others. This is a major code change, but will probably not be noticeable by most users.

  Making this package independent of especially the tidyverse (e.g. packages `dplyr` and `tidyr`) tremendously increases sustainability on the long term, since tidyverse functions change quite often. Good for users, but hard for package maintainers. Most of our functions are replaced with versions that only rely on base R, which keeps this package fully functional for many years to come, without requiring a lot of maintenance to keep up with other packages anymore. Another upside it that this package can now be used with all versions of R since R-3.0.0 (April 2013). Our package is being used in settings where the resources are very limited. Fewer dependencies on newer software is helpful for such settings.
  
  Negative effects of this change are:
  * Function `freq()` that was borrowed from the `cleaner` package was removed. Use `cleaner::freq()`, or run `library("cleaner")` before you use `freq()`.
  * ~~Printing values of class `mo` or `rsi` in a tibble will no longer be in colour and printing `rsi` in a tibble will show the class `<ord>`, not `<rsi>` anymore. This is purely a visual effect.~~
  * ~~All functions from the `mo_*` family (like `mo_name()` and `mo_gramstain()`) are noticeably slower when running on hundreds of thousands of rows.~~
  * For developers: classes `mo` and `ab` now both also inherit class `character`, to support any data transformation. This change invalidates code that checks for class length == 1.

### Changed
* Taxonomy:
  * Updated the taxonomy of microorganisms to May 2020, using the Catalogue of Life (CoL), the Global Biodiversity Information Facility (GBIF) and the List of Prokaryotic names with Standing in Nomenclature (LPSN, hosted by DSMZ since February 2020). **Note:** a taxonomic update may always impact determination of first isolates (using `first_isolate()`), since some bacterial names might be renamed to other genera or other (sub)species. This is expected behaviour.
  * Removed the Catalogue of Life IDs (like 776351), since they now work with a species ID (hexadecimal string)
* EUCAST rules:
  * The `eucast_rules()` function no longer applies "other" rules at default that are made available by this package (like setting ampicillin = R when ampicillin + enzyme inhibitor = R). The default input value for `rules` is now `c("breakpoints", "expert")` instead of `"all"`, but this can be changed by the user. To return to the old behaviour, set `options(AMR.eucast_rules = "all")`.
  * Fixed a bug where checking antimicrobial results in the original data were not regarded as valid R/SI values
  * All "other" rules now apply for all drug combinations in the `antibiotics` data set these two rules:
    1. A drug **with** enzyme inhibitor will be set to S if the drug **without** enzyme inhibitor is S
    2. A drug **without** enzyme inhibitor will be set to R if the drug **with** enzyme inhibitor is R
    
    This works for all drug combinations, such as ampicillin/sulbactam, ceftazidime/avibactam, trimethoprim/sulfamethoxazole, etc.
  * Added official drug names to verbose output of `eucast_rules()`
* Added function `ab_url()` to return the direct URL of an antimicrobial agent from the official WHO website
* Improvements for algorithm in `as.ab()`, so that e.g. `as.ab("ampi sul")` and `ab_name("ampi sul")` work
* Functions `ab_atc()` and `ab_group()` now return `NA` if no antimicrobial agent could be found
* Small fix for some text input that could not be coerced as valid MIC values 
* Fix for interpretation of generic CLSI interpretation rules (thanks to Anthony Underwood)
* Fix for `set_mo_source()` to make sure that column `mo` will always be the second column
* Added abbreviation "cfsc" for Cefoxitin and "cfav" for Ceftazidime/avibactam

### Other
* Removed previously deprecated function `p.symbol()` - it was replaced with `p_symbol()`
* Removed function `read.4d()`, that was only useful for reading data from an old test database.

# AMR 1.1.0

### New
* Support for easy principal component analysis for AMR, using the new `pca()` function 
* Plotting biplots for principal component analysis using the new `ggplot_pca()` function

### Changed
* Improvements for the algorithm used by `as.mo()` (and consequently all `mo_*` functions, that use `as.mo()` internally):
  * Support for codes ending with `SPE` for species, like `"ESCSPE"` for *Escherichia coli*
  * Support for any encoding, which means that any language-specific character with accents can be used for input
  * Support for more arbitrary IDs used in laboratory information systems
  * Small fix for preventing viruses being treated as bacteria
  * Small fix for preventing contamination and lack of growth being treated as valid microorganisms
* Support for all abbreviations of antibiotics and antimycotics used by the Netherlands National Institute for Public Health and the Environment (Rijksinstituut voor Volksgezondheid en Milieu; RIVM)
* Added more abbreviations to the `antibiotics` data set
* Reloaded original EUCAST master tables from 2019 (2020 was already available). This seems more reliable than the data we used from WHONET.
* Added generic CLSI rules for R/SI interpretation using `as.rsi()` for years 2010-2019 (thanks to Anthony Underwood)

### Other
* Support for the upcoming `dplyr` version 1.0.0
* More robust assigning for classes `rsi` and `mic`

# AMR 1.0.1

### Changed
* Fixed important floating point error for some MIC comparisons in EUCAST 2020 guideline
* Interpretation from MIC values (and disk zones) to R/SI can now be used with `mutate_at()` of the `dplyr` package:
  ```r
  yourdata %>% 
    mutate_at(vars(antibiotic1:antibiotic25), as.rsi, mo = "E. coli")
    
  yourdata %>% 
    mutate_at(vars(antibiotic1:antibiotic25), as.rsi, mo = .$mybacteria)
  ```
* Added antibiotic abbreviations for a laboratory manufacturer (GLIMS) for cefuroxime, cefotaxime, ceftazidime, cefepime, cefoxitin and trimethoprim/sulfamethoxazole
* Added `uti` (as abbreviation of urinary tract infections) as argument to `as.rsi()`, so interpretation of MIC values and disk zones can be made dependent on isolates specifically from UTIs
* Info printing in functions `eucast_rules()`, `first_isolate()`, `mdro()` and `resistance_predict()` will now at default only print when R is in an interactive mode (i.e. not in RMarkdown)

# AMR 1.0.0

This software is now out of beta and considered stable. Nonetheless, this package will be developed continually.

### New
* Support for the newest [EUCAST Clinical Breakpoint Tables v.10.0](https://www.eucast.org/clinical_breakpoints/), valid from 1 January 2020. This affects translation of MIC and disk zones using `as.rsi()` and inferred resistance and susceptibility using `eucast_rules()`.
* The repository of this package now contains a clean version of the EUCAST and CLSI guidelines from 2011-2020 to translate MIC and disk diffusion values to R/SI: <https://github.com/msberends/AMR/blob/master/data-raw/rsi_translation.txt>. This **allows for machine reading these guidelines**, which is almost impossible with the Excel and PDF files distributed by EUCAST and CLSI. This file used to process the EUCAST Clinical Breakpoints Excel file [can be found here](https://github.com/msberends/AMR/blob/master/data-raw/read_EUCAST.R).
* Support for LOINC and SNOMED codes
  * Support for LOINC codes in the `antibiotics` data set. Use `ab_loinc()` to retrieve LOINC codes, or use a LOINC code for input in any `ab_*` function:
    ```r
    ab_loinc("ampicillin")
    #> [1] "21066-6" "3355-5"  "33562-0" "33919-2" "43883-8" "43884-6" "87604-5"
    ab_name("21066-6")
    #> [1] "Ampicillin"
    ab_atc("21066-6")
    #> [1] "J01CA01"
    ```
  * Support for SNOMED CT codes in the `microorganisms` data set. Use `mo_snomed()` to retrieve SNOMED codes, or use a SNOMED code for input in any `mo_*` function:
    ```r
    mo_snomed("S. aureus")
    #> [1] 115329001   3092008 113961008
    mo_name(115329001)
    #> [1] "Staphylococcus aureus"
    mo_gramstain(115329001)
    #> [1] "Gram-positive"
    ```

### Changes
* The `as.mo()` function previously wrote to the package folder to improve calculation speed for previously calculated results. This is no longer the case, to comply with CRAN policies. Consequently, the function `clear_mo_history()` was removed.
* Bugfix for some WHONET microorganism codes that were not interpreted correctly when using `as.rsi()`
* Improvements for the algorithm used by `as.mo()` (and consequently all `mo_*` functions, that use `as.mo()` internally):
  * Support for missing spaces, e.g. in `as.mo("Methicillin-resistant S.aureus")`
  * Better support for determination of *Salmonella* biovars
  * Speed improvements, especially for the *G. species* format (G for genus), like *E. coli* and *K pneumoniae*
  * Support for more common codes used in laboratory information systems
* Input values for `as.disk()` limited to a maximum of 50 millimeters
* Added a lifecycle state to every function, following [the lifecycle circle of the `tidyverse`](https://www.tidyverse.org/lifecycle)
* For in `as.ab()`: support for drugs starting with "co-" like co-amoxiclav, co-trimoxazole, co-trimazine and co-trimazole (thanks to Peter Dutey)
* Changes to the `antibiotics` data set (thanks to Peter Dutey):
  * Added more synonyms to colistin, imipenem and piperacillin/tazobactam
  * Moved synonyms Rifinah and Rimactazid from rifampicin (`RIF`) to rifampicin/isoniazid (`RFI`). Please note that [the combination rifampicin/isoniazid has no DDDs defined](https://www.whocc.no/atc_ddd_index/?code=J04AM02&showdescription=no), so e.g. `ab_ddd("Rimactazid")` will now return `NA`.
  * Moved synonyms Bactrimel and Cotrimazole from sulfamethoxazole (`SMX`) to trimethoprim/sulfamethoxazole (`SXT`)

### Other
* Add a `CITATION` file
* Full support for the upcoming R 4.0
* Removed unnecessary `AMR::` calls

# AMR 0.9.0

### Breaking
* Adopted Adeolu *et al.* (2016), [PMID 27620848](https:/pubmed.ncbi.nlm.nih.gov/27620848/) for the `microorganisms` data set, which means that the new order Enterobacterales now consists of a part of the existing family Enterobacteriaceae, but that this family has been split into other families as well (like *Morganellaceae* and *Yersiniaceae*). Although published in 2016, this information is not yet in the Catalogue of Life version of 2019. All MDRO determinations with `mdro()` will now use the Enterobacterales order for all guidelines before 2016 that were dependent on the Enterobacteriaceae family.
  * If you were dependent on the old Enterobacteriaceae family e.g. by using in your code:
    ```r
    if (mo_family(somebugs) == "Enterobacteriaceae") ...
    ```
    then please adjust this to:
    ```r
    if (mo_order(somebugs) == "Enterobacterales") ...
    ```

### New
* Functions `susceptibility()` and `resistance()` as aliases of `proportion_SI()` and `proportion_R()`, respectively. These functions were added to make it more clear that "I" should be considered susceptible and not resistant.
  ```r
  library(dplyr)
  example_isolates %>%
    group_by(bug = mo_name(mo)) %>% 
    summarise(amoxicillin = resistance(AMX),
              amox_clav   = resistance(AMC)) %>%
    filter(!is.na(amoxicillin) | !is.na(amox_clav))
  ```
* Support for a new MDRO guideline: Magiorakos AP, Srinivasan A *et al.* "Multidrug-resistant, extensively drug-resistant and pandrug-resistant bacteria: an international expert proposal for interim standard definitions for acquired resistance." Clinical Microbiology and Infection (2012).
  * This is now the new default guideline for the `mdro()` function
  * The new Verbose mode (`mdro(...., verbose = TRUE)`) returns an informative data set where the reason for MDRO determination is given for every isolate, and an list of the resistant antimicrobial agents
* Data set `antivirals`, containing all entries from the ATC J05 group with their DDDs for oral and parenteral treatment

### Changes
* Improvements to algorithm in `as.mo()`:
  * Now allows "ou" where "au" should have been used and vice versa
  * More intelligent way of coping with some consonants like "l" and "r"
  * Added a score (a certainty percentage) to `mo_uncertainties()`, that is calculated using the [Levenshtein distance](https://en.wikipedia.org/wiki/Levenshtein_distance):
    ```r
    as.mo(c("Stafylococcus aureus",
            "staphylokok aureuz"))
    #> Warning: 
    #> Results of two values were guessed with uncertainty. Use mo_uncertainties() to review them.
    #> Class 'mo'
    #> [1] B_STPHY_AURS B_STPHY_AURS
    
    mo_uncertainties()
    #> "Stafylococcus aureus" -> Staphylococcus aureus (B_STPHY_AURS, score: 95.2%)
    #> "staphylokok aureuz"   -> Staphylococcus aureus (B_STPHY_AURS, score: 85.7%)
    ```
* Removed previously deprecated function `as.atc()` - this function was replaced by `ab_atc()`
* Renamed all `portion_*` functions to `proportion_*`. All `portion_*` functions are still available as deprecated functions, and will return a warning when used.
* When running `as.rsi()` over a data set, it will now print the guideline that will be used if it is not specified by the user
* Improvements for `eucast_rules()`:
  * Fix where *Stenotrophomonas maltophilia* would always become ceftazidime R (following EUCAST v3.1)
  * Fix where *Leuconostoc* and *Pediococcus* would not always become glycopeptides R
  * non-EUCAST rules in `eucast_rules()` are now applied first and not as last anymore. This is to improve the dependency on certain antibiotics for the official EUCAST rules. Please see `?eucast_rules`.
* Fix for interpreting MIC values with `as.rsi()` where the input is `NA`
* Added "imi" and "imp" as allowed abbreviation for Imipenem (IPM)
* Fix for automatically determining columns with antibiotic results in `mdro()` and `eucast_rules()`
* Added ATC codes for ceftaroline, ceftobiprole and faropenem and fixed two typos in the `antibiotics` data set
* More robust way of determining valid MIC values
* Small changed to the `example_isolates` data set to better reflect reality
* Added more microorganisms codes from laboratory systems (esp. species of *Pseudescherichia* and *Rodentibacter*)
* Added Gram-stain to `mo_info()`

### Other
* Rewrote the complete documentation to markdown format, to be able to use the very latest version of the great [Roxygen2](https://roxygen2.r-lib.org/index.html), released in November 2019. This tremously improved the documentation quality, since the rewrite forced us to go over all texts again and make changes where needed.
* Change dependency on `clean` to `cleaner`, as this package was renamed accordingly upon CRAN request
* Added Dr. Sofia Ny as contributor

# AMR 0.8.0

### Breaking
* Determination of first isolates now **excludes** all 'unknown' microorganisms at default, i.e. microbial code `"UNKNOWN"`. They can be included with the new argument `include_unknown`:
  ```r
  first_isolate(..., include_unknown = TRUE)
  ```
  For WHONET users, this means that all records/isolates with organism code `"con"` (*contamination*) will be excluded at default, since `as.mo("con") = "UNKNOWN"`. The function always shows a note with the number of 'unknown' microorganisms that were included or excluded.
* For code consistency, classes `ab` and `mo` will now be preserved in any subsetting or assignment. For the sake of data integrity, this means that invalid assignments will now result in `NA`:
  ```r
  # how it works in base R:
  x <- factor("A")
  x[1] <- "B"
  #> Warning message:
  #> invalid factor level, NA generated
  
  # how it now works similarly for classes 'mo' and 'ab':
  x <- as.mo("E. coli")
  x[1] <- "testvalue"
  #> Warning message:
  #> invalid microorganism code, NA generated
  ```
  This is important, because a value like `"testvalue"` could never be understood by e.g. `mo_name()`, although the class would suggest a valid microbial code.
* Function `freq()` has moved to a new package, [`clean`](https://github.com/msberends/clean) ([CRAN link](https://cran.r-project.org/package=clean)), since creating frequency tables actually does not fit the scope of this package. The `freq()` function still works, since it is re-exported from the `clean` package (which will be installed automatically upon updating this `AMR` package).
* Renamed data set `septic_patients` to `example_isolates`

### New
* Function `bug_drug_combinations()` to quickly get a `data.frame` with the results of all bug-drug combinations in a data set. The column containing microorganism codes is guessed automatically and its input is transformed with `mo_shortname()` at default:
  ```r
  x <- bug_drug_combinations(example_isolates)
  #> NOTE: Using column `mo` as input for `col_mo`.
  x[1:4, ]
  #>             mo  ab S I R total
  #> 1 A. baumannii AMC 0 0 3     3
  #> 2 A. baumannii AMK 0 0 0     0
  #> 3 A. baumannii AMP 0 0 3     3
  #> 4 A. baumannii AMX 0 0 3     3
  #> NOTE: Use 'format()' on this result to get a publicable/printable format.

  # change the transformation with the FUN argument to anything you like:
  x <- bug_drug_combinations(example_isolates, FUN = mo_gramstain)
  #> NOTE: Using column `mo` as input for `col_mo`.
  x[1:4, ]
  #>              mo  ab   S  I   R total
  #> 1 Gram-negative AMC 469 89 174   732
  #> 2 Gram-negative AMK 251  0   2   253
  #> 3 Gram-negative AMP 227  0 405   632
  #> 4 Gram-negative AMX 227  0 405   632
  #> NOTE: Use 'format()' on this result to get a publicable/printable format.
  ```
  You can format this to a printable format, ready for reporting or exporting to e.g. Excel with the base R `format()` function:
  ```r
  format(x, combine_IR = FALSE)
  ```
* Additional way to calculate co-resistance, i.e. when using multiple antimicrobials as input for `portion_*` functions or `count_*` functions. This can be used to determine the empiric susceptibility of a combination therapy. A new argument `only_all_tested` (**which defaults to `FALSE`**) replaces the old `also_single_tested` and can be used to select one of the two methods to count isolates and calculate portions. The difference can be seen in this example table (which is also on the `portion` and `count` help pages), where the %SI is being determined:

  ```r
  # --------------------------------------------------------------------
  #                     only_all_tested = FALSE  only_all_tested = TRUE
  #                     -----------------------  -----------------------
  #  Drug A    Drug B   include as  include as   include as  include as
  #                     numerator   denominator  numerator   denominator
  # --------  --------  ----------  -----------  ----------  -----------
  #  S or I    S or I       X            X            X            X
  #    R       S or I       X            X            X            X
  #   <NA>     S or I       X            X            -            -
  #  S or I      R          X            X            X            X
  #    R         R          -            X            -            X
  #   <NA>       R          -            -            -            -
  #  S or I     <NA>        X            X            -            -
  #    R        <NA>        -            -            -            -
  #   <NA>      <NA>        -            -            -            -
  # --------------------------------------------------------------------
  ```
  
  Since this is a major change, usage of the old `also_single_tested` will throw an informative error that it has been replaced by `only_all_tested`.
* `tibble` printing support for classes `rsi`, `mic`, `disk`, `ab` `mo`. When using `tibble`s containing antimicrobial columns, values `S` will print in green, values `I` will print in yellow and values `R` will print in red. Microbial IDs (class `mo`) will emphasise on the genus and species, not on the kingdom.
  ```r
  # (run this on your own console, as this page does not support colour printing)
  library(dplyr)
  example_isolates %>%
    select(mo:AMC) %>% 
    as_tibble()
  ```

### Changed
* Many algorithm improvements for `as.mo()` (of which some led to additions to the `microorganisms` data set). Many thanks to all contributors that helped improving the algorithms.
  * Self-learning algorithm - the function now gains experience from previously determined microorganism IDs and learns from it (yielding 80-95% speed improvement for any guess after the first try)
  * Big improvement for misspelled input
  * These new trivial names known to the field are now understood: meningococcus, gonococcus, pneumococcus
  * Updated to the latest taxonomic data (updated to August 2019, from the International Journal of Systematic and Evolutionary Microbiology
  * Added support for Viridans Group Streptococci (VGS) and Milleri Group Streptococci (MGS)
  * Added support for *Blastocystis*
  * Added support for 5,000 new fungi
  * Added support for unknown yeasts and fungi
  * Changed most microorganism IDs to improve readability. For example, the old code `B_ENTRC_FAE` could have been both *E. faecalis* and *E. faecium*. Its new code is `B_ENTRC_FCLS` and *E. faecium* has become `B_ENTRC_FACM`. Also, the Latin character æ (ae) is now preserved at the start of each genus and species abbreviation. For example, the old code for *Aerococcus urinae* was `B_ARCCC_NAE`. This is now `B_AERCC_URIN`.
    **IMPORTANT:** Old microorganism IDs are still supported, but support will be dropped in a future version. Use `as.mo()` on your old codes to transform them to the new format. Using functions from the `mo_*` family (like `mo_name()` and `mo_gramstain()`) on old codes, will throw a warning.
* More intelligent guessing for `as.ab()`, including bidirectional language support
* Added support for the German national guideline (3MRGN/4MRGN) in the `mdro()` function, to determine multi-drug resistant organisms
* Function `eucast_rules()`:
  * Fixed a bug for *Yersinia pseudotuberculosis*
  * Added more informative errors and warnings
  * Printed info now distinguishes between added and changes values
  * Using Verbose mode (i.e. `eucast_rules(..., verbose = TRUE)`) returns more informative and readable output
  * Using factors as input now adds missing factors levels when the function changes antibiotic results
* Improved the internal auto-guessing function for determining antimicrobials in your data set (`AMR:::get_column_abx()`)
* Removed class `atc` - using `as.atc()` is now deprecated in favour of `ab_atc()` and this will return a character, not the `atc` class anymore
* Removed deprecated functions `abname()`, `ab_official()`, `atc_name()`, `atc_official()`, `atc_property()`, `atc_tradenames()`, `atc_trivial_nl()`
* Fix and speed improvement for `mo_shortname()`
* Fix for using `mo_*` functions where the coercion uncertainties and failures would not be available through `mo_uncertainties()` and `mo_failures()` anymore
* Deprecated the `country` argument of `mdro()` in favour of the already existing `guideline` argument to support multiple guidelines within one country
* The `name` of `RIF` is now Rifampicin instead of Rifampin
* The `antibiotics` data set is now sorted by name and all cephalosporins now have their generation between brackets
* Speed improvement for `guess_ab_col()` which is now 30 times faster for antibiotic abbreviations
* Improved `filter_ab_class()` to be more reliable and to support 5th generation cephalosporins
* Function `availability()` now uses `portion_R()` instead of `portion_IR()`, to comply with EUCAST insights
* Functions `age()` and `age_groups()` now have a `na.rm` argument to remove empty values
* Renamed function `p.symbol()` to `p_symbol()` (the former is now deprecated and will be removed in a future version)
* Using negative values for `x` in `age_groups()` will now introduce `NA`s and not return an error anymore
* Fix for determining the system's language
* Fix for `key_antibiotics()` on foreign systems
* Added 80 new LIS codes for microorganisms
* Relabeled the factor levels of `mdr_tb()`
* Added more MIC factor levels (`as.mic()`)

#### Other
* Added Prof. Dr. Casper Albers as doctoral advisor and added Dr. Judith Fonville, Eric Hazenberg, Dr. Bart Meijer, Dr. Dennis Souverein and Annick Lenglet as contributors
* Cleaned the coding style of every single syntax line in this package with the help of the `lintr` package

# AMR 0.7.1

#### New
* Function `rsi_df()` to transform a `data.frame` to a data set containing only the microbial interpretation (S, I, R), the antibiotic, the percentage of S/I/R and the number of available isolates. This is a convenient combination of the existing functions `count_df()` and `portion_df()` to immediately show resistance percentages and number of available isolates:
  ```r
  septic_patients %>%
    select(AMX, CIP) %>%
    rsi_df()
  #      antibiotic  interpretation      value  isolates
  # 1   Amoxicillin              SI  0.4442636       546
  # 2   Amoxicillin               R  0.5557364       683
  # 3 Ciprofloxacin              SI  0.8381831      1181
  # 4 Ciprofloxacin               R  0.1618169       228
  ```
* Support for all scientifically published pathotypes of *E. coli* to date (that we could find). Supported are: 

  * AIEC (Adherent-Invasive *E. coli*) 
  * ATEC (Atypical Entero-pathogenic *E. coli*) 
  * DAEC (Diffusely Adhering *E. coli*) 
  * EAEC (Entero-Aggresive *E. coli*) 
  * EHEC (Entero-Haemorrhagic *E. coli*) 
  * EIEC (Entero-Invasive *E. coli*) 
  * EPEC (Entero-Pathogenic *E. coli*) 
  * ETEC (Entero-Toxigenic *E. coli*) 
  * NMEC (Neonatal Meningitis‐causing *E. coli*) 
  * STEC (Shiga-toxin producing *E. coli*) 
  * UPEC (Uropathogenic *E. coli*)
  
  All these lead to the microbial ID of *E. coli*:
  ```r
  as.mo("UPEC")
  # B_ESCHR_COL
  mo_name("UPEC")
  # "Escherichia coli"
  mo_gramstain("EHEC")
  # "Gram-negative"
  ```
* Function `mo_info()` as an analogy to `ab_info()`. The `mo_info()` prints a list with the full taxonomy, authors, and the URL to the online database of a microorganism
* Function `mo_synonyms()` to get all previously accepted taxonomic names of a microorganism

#### Changed
* Column names of output `count_df()` and `portion_df()` are now lowercase
* Fixed bug in translation of microorganism names
* Fixed bug in determining taxonomic kingdoms
* Algorithm improvements for `as.ab()` and `as.mo()` to understand even more severely misspelled input
* Function `as.ab()` now allows spaces for coercing antibiotics names
* Added `ggplot2` methods for automatically determining the scale type of classes `mo` and `ab`
* Added names of object in the header in frequency tables, even when using pipes
* Prevented `"bacteria"` from getting coerced by `as.ab()` because Bacterial is a brand name of trimethoprim (TMP)
* Fixed a bug where setting an antibiotic would not work for `eucast_rules()` and `mdro()`
* Fixed a EUCAST rule for Staphylococci, where amikacin resistance would not be inferred from tobramycin
* Removed `latest_annual_release` from the `catalogue_of_life_version()` function
* Removed antibiotic code `PVM1` from the `antibiotics` data set as this was a duplicate of `PME`
* Fixed bug where not all old taxonomic names would be printed, when using a vector as input for `as.mo()`
* Manually added *Trichomonas vaginalis* from the kingdom of Protozoa, which is missing from the Catalogue of Life
* Small improvements to `plot()` and `barplot()` for MIC and RSI classes
* Allow Catalogue of Life IDs to be coerced by `as.mo()`

#### Other
* Fixed a note thrown by CRAN tests

# AMR 0.7.0

#### New
* Support for translation of disk diffusion and MIC values to RSI values (i.e. antimicrobial interpretations). Supported guidelines are EUCAST (2011 to 2019) and CLSI (2011 to 2019). Use `as.rsi()` on an MIC value (created with `as.mic()`), a disk diffusion value (created with the new `as.disk()`) or on a complete date set containing columns with MIC or disk diffusion values.
* Function `mo_name()` as alias of `mo_fullname()`
* Added guidelines of the WHO to determine multi-drug resistance (MDR) for TB (`mdr_tb()`) and added a new vignette about MDR. Read this tutorial [here on our website](https://msberends.gitlab.io/AMR/articles/MDR.html).

#### Changed
* Fixed a critical bug in `first_isolate()` where missing species would lead to incorrect FALSEs. This bug was not present in AMR v0.5.0, but was in v0.6.0 and v0.6.1.
* Fixed a bug in `eucast_rules()` where antibiotics from WHONET software would not be recognised
* Completely reworked the `antibiotics` data set:
  * All entries now have 3 different identifiers:
    * Column `ab` contains a human readable EARS-Net code, used by ECDC and WHO/WHONET - this is the primary identifier used in this package
    * Column `atc` contains the ATC code, used by WHO/WHOCC
    * Column `cid` contains the CID code (Compound ID), used by PubChem
  * Based on the Compound ID, almost 5,000 official brand names have been added from many different countries
  * All references to antibiotics in our package now use EARS-Net codes, like `AMX` for amoxicillin
  * Functions `atc_certe`, `ab_umcg` and `atc_trivial_nl` have been removed
  * All `atc_*` functions are superceded by `ab_*` functions
  * All output will be translated by using an included translation file which [can be viewed here](https://github.com/msberends/AMR/blob/master/data-raw/translations.tsv)
* Improvements to plotting AMR results with `ggplot_rsi()`:
  * New argument `colours` to set the bar colours
  * New arguments `title`, `subtitle`, `caption`, `x.title` and `y.title` to set titles and axis descriptions
* Improved intelligence of looking up antibiotic columns in a data set using `guess_ab_col()`
* Added ~5,000 more old taxonomic names to the `microorganisms.old` data set, which leads to better results finding when using the `as.mo()` function
* This package now honours the new EUCAST insight (2019) that S and I are but classified as susceptible, where I is defined as 'increased exposure' and not 'intermediate' anymore. For functions like `portion_df()` and `count_df()` this means that their new argument `combine_SI` is TRUE at default. Our plotting function `ggplot_rsi()` also reflects this change since it uses `count_df()` internally.
* The `age()` function gained a new argument `exact` to determine ages with decimals
* Removed deprecated functions `guess_mo()`, `guess_atc()`, `EUCAST_rules()`, `interpretive_reading()`, `rsi()`
* Frequency tables (`freq()`):
  * speed improvement for microbial IDs
  * fixed factor level names for R Markdown
  * when all values are unique it now shows a message instead of a warning
  * support for boxplots:
    ```r
    septic_patients %>% 
      freq(age) %>% 
      boxplot()
    # grouped boxplots:
    septic_patients %>% 
      group_by(hospital_id) %>% 
      freq(age) %>%
      boxplot()
    ```
* Removed all hardcoded EUCAST rules and replaced them with a new reference file which [can be viewed here](https://github.com/msberends/AMR/blob/master/data-raw/eucast_rules.tsv)
* Added ceftazidim intrinsic resistance to *Streptococci*
* Changed default settings for `age_groups()`, to let groups of fives and tens end with 100+ instead of 120+
* Fix for `freq()` for when all values are `NA`
* Fix for `first_isolate()` for when dates are missing
* Improved speed of `guess_ab_col()`
* Function `as.mo()` now gently interprets any number of whitespace characters (like tabs) as one space
* Function `as.mo()` now returns `UNKNOWN` for `"con"` (WHONET ID of 'contamination') and returns `NA` for `"xxx"`(WHONET ID of 'no growth')
* Small algorithm fix for `as.mo()`
* Removed viruses from data set `microorganisms.codes` and cleaned it up
* Fix for `mo_shortname()` where species would not be determined correctly

#### Other
* Support for R 3.6.0 and later by providing support for [staged install](https://developer.r-project.org/Blog/public/2019/02/14/staged-install/index.html)

# AMR 0.6.1

#### Changed
* Fixed a critical bug when using `eucast_rules()` with `verbose = TRUE`
* Coercion of microbial IDs are now written to the package namespace instead of the user's home folder, to comply with the CRAN policy

# AMR 0.6.0

**New website!**

We've got a new website: [https://msberends.gitlab.io/AMR](https://msberends.gitlab.io/AMR/) (built with the great [`pkgdown`](https://pkgdown.r-lib.org/))

* Contains the complete manual of this package and all of its functions with an explanation of their arguments
* Contains a comprehensive tutorial about how to conduct antimicrobial resistance analysis, import data from WHONET or SPSS and many more.

#### New
* **BREAKING**: removed deprecated functions, arguments and references to 'bactid'. Use `as.mo()` to identify an MO code.
* Catalogue of Life as a new taxonomic source for data about microorganisms, which also contains all ITIS data we used previously. The `microorganisms` data set now contains:
  * All ~55,000 (sub)species from the kingdoms of Archaea, Bacteria and Protozoa
  * All ~3,000 (sub)species from these orders of the kingdom of Fungi: Eurotiales, Onygenales, Pneumocystales, Saccharomycetales and Schizosaccharomycetales (covering at least like all species of *Aspergillus*, *Candida*, *Pneumocystis*, *Saccharomyces* and *Trichophyton*)
  * All ~2,000 (sub)species from ~100 other relevant genera, from the kingdoms of Animalia and Plantae (like *Strongyloides* and *Taenia*)
  * All ~15,000 previously accepted names of included (sub)species that have been taxonomically renamed
  * The responsible author(s) and year of scientific publication
  
    This data is updated annually - check the included version with the new function `catalogue_of_life_version()`.
  * Due to this change, some `mo` codes changed (e.g. *Streptococcus* changed from `B_STRPTC` to `B_STRPT`). A translation table is  used internally to support older microorganism IDs, so users will not notice this difference.
  * New function `mo_rank()` for the taxonomic rank (genus, species, infraspecies, etc.)
  * New function `mo_url()` to get the direct URL of a species from the Catalogue of Life
* Support for data from [WHONET](https://whonet.org/) and [EARS-Net](https://www.ecdc.europa.eu/en/about-us/partnerships-and-networks/disease-and-laboratory-networks/ears-net) (European Antimicrobial Resistance Surveillance Network):
  * Exported files from WHONET can be read and used in this package. For functions like `first_isolate()` and `eucast_rules()`, all arguments will be filled in automatically.
  * This package now knows all antibiotic abbrevations by EARS-Net (which are also being used by WHONET) - the `antibiotics` data set now contains a column `ears_net`.
  * The function `as.mo()` now knows all WHONET species abbreviations too, because almost 2,000 microbial abbreviations were added to the `microorganisms.codes` data set.
* New filters for antimicrobial classes. Use these functions to filter isolates on results in one of more antibiotics from a specific class:
  ```r
  filter_aminoglycosides()
  filter_carbapenems()
  filter_cephalosporins()
  filter_1st_cephalosporins()
  filter_2nd_cephalosporins()
  filter_3rd_cephalosporins()
  filter_4th_cephalosporins()
  filter_fluoroquinolones()
  filter_glycopeptides()
  filter_macrolides()
  filter_tetracyclines()
  ```
  The `antibiotics` data set will be searched, after which the input data will be checked for column names with a value in any abbreviations, codes or official names found in the `antibiotics` data set.
  For example:
  ```r
  septic_patients %>% filter_glycopeptides(result = "R")
  # Filtering on glycopeptide antibacterials: any of `vanc` or `teic` is R
  septic_patients %>% filter_glycopeptides(result = "R", scope = "all")
  # Filtering on glycopeptide antibacterials: all of `vanc` and `teic` is R
  ```
* All `ab_*` functions are deprecated and replaced by `atc_*` functions:
  ```r
  ab_property -> atc_property()
  ab_name -> atc_name()
  ab_official -> atc_official()
  ab_trivial_nl -> atc_trivial_nl()
  ab_certe -> atc_certe()
  ab_umcg -> atc_umcg()
  ab_tradenames -> atc_tradenames()
  ```
  These functions use `as.atc()` internally. The old `atc_property` has been renamed `atc_online_property()`. This is done for two reasons: firstly, not all ATC codes are of antibiotics (ab) but can also be of antivirals or antifungals. Secondly, the input must have class `atc` or must be coerable to this class. Properties of these classes should start with the same class name, analogous to `as.mo()` and e.g. `mo_genus`.
* New functions `set_mo_source()` and `get_mo_source()` to use your own predefined MO codes as input for `as.mo()` and consequently all `mo_*` functions
* Support for the upcoming [`dplyr`](https://dplyr.tidyverse.org) version 0.8.0
* New function `guess_ab_col()` to find an antibiotic column in a table
* New function `mo_failures()` to review values that could not be coerced to a valid MO code, using `as.mo()`. This latter function will now only show a maximum of 10 uncoerced values and will refer to `mo_failures()`.
* New function `mo_uncertainties()` to review values that could be coerced to a valid MO code using `as.mo()`, but with uncertainty.
* New function `mo_renamed()` to get a list of all returned values from `as.mo()` that have had taxonomic renaming
* New function `age()` to calculate the (patients) age in years
* New function `age_groups()` to split ages into custom or predefined groups (like children or elderly). This allows for easier demographic antimicrobial resistance analysis per age group.
* New function `ggplot_rsi_predict()` as well as the base R `plot()` function can now be used for resistance prediction calculated with `resistance_predict()`:
  ```r
  x <- resistance_predict(septic_patients, col_ab = "amox")
  plot(x)
  ggplot_rsi_predict(x)
  ```
* Functions `filter_first_isolate()` and `filter_first_weighted_isolate()` to shorten and fasten filtering on data sets with antimicrobial results, e.g.:
  ```r
  septic_patients %>% filter_first_isolate(...)
  # or
  filter_first_isolate(septic_patients, ...)
  ```
  is equal to:
  ```r
  septic_patients %>%
    mutate(only_firsts = first_isolate(septic_patients, ...)) %>%
    filter(only_firsts == TRUE) %>%
    select(-only_firsts)
  ```
* New function `availability()` to check the number of available (non-empty) results in a `data.frame`
* New vignettes about how to conduct AMR analysis, predict antimicrobial resistance, use the *G*-test and more. These are also available (and even easier readable) on our website: https://msberends.gitlab.io/AMR.

#### Changed
* Function `eucast_rules()`:
  * Updated EUCAST Clinical breakpoints to [version 9.0 of 1 January 2019](https://www.eucast.org/clinical_breakpoints/), the data set `septic_patients` now reflects these changes
  * Fixed a critical bug where some rules that depend on previous applied rules would not be applied adequately
  * Emphasised in manual that penicillin is meant as benzylpenicillin (ATC [J01CE01](https://www.whocc.no/atc_ddd_index/?code=J01CE01))
  * New info is returned when running this function, stating exactly what has been changed or added. Use `eucast_rules(..., verbose = TRUE)` to get a data set with all changed per bug and drug combination.
* Removed data sets `microorganisms.oldDT`, `microorganisms.prevDT`, `microorganisms.unprevDT` and `microorganismsDT` since they were no longer needed and only contained info already available in the `microorganisms` data set
* Added 65 antibiotics to the `antibiotics` data set, from the [Pharmaceuticals Community Register](http://ec.europa.eu/health/documents/community-register/html/atc.htm) of the European Commission
* Removed columns `atc_group1_nl` and `atc_group2_nl` from the `antibiotics` data set
* Functions `atc_ddd()` and `atc_groups()` have been renamed `atc_online_ddd()` and `atc_online_groups()`. The old functions are deprecated and will be removed in a future version.
* Function `guess_mo()` is now deprecated in favour of `as.mo()` and will be removed in future versions
* Function `guess_atc()` is now deprecated in favour of `as.atc()` and will be removed in future versions
* Improvements for `as.mo()`:
  * Now handles incorrect spelling, like `i` instead of `y` and `f` instead of `ph`:
    ```r
    # mo_fullname() uses as.mo() internally
    
    mo_fullname("Sthafilokockus aaureuz")
    #> [1] "Staphylococcus aureus"
    
    mo_fullname("S. klossi")
    #> [1] "Staphylococcus kloosii"
    ```
  * Uncertainty of the algorithm is now divided into four levels, 0 to 3, where the default `allow_uncertain = TRUE` is equal to uncertainty level 2. Run `?as.mo` for more info about these levels.
    ```r
    # equal:
    as.mo(..., allow_uncertain = TRUE)
    as.mo(..., allow_uncertain = 2)
    
    # also equal:
    as.mo(..., allow_uncertain = FALSE)
    as.mo(..., allow_uncertain = 0)
    ```
    Using `as.mo(..., allow_uncertain = 3)` could lead to very unreliable results.
  * Implemented the latest publication of Becker *et al.* (2019), for categorising coagulase-negative *Staphylococci*
  * All microbial IDs that found are now saved to a local file `~/.Rhistory_mo`. Use the new function `clean_mo_history()` to delete this file, which resets the algorithms.
  * Incoercible results will now be considered 'unknown', MO code `UNKNOWN`. On foreign systems, properties of these will be translated to all languages already previously supported: German, Dutch, French, Italian, Spanish and Portuguese:
    ```r
    mo_genus("qwerty", language = "es")
    # Warning: 
    # one unique value (^= 100.0%) could not be coerced and is considered 'unknown': "qwerty". Use mo_failures() to review it.
    #> [1] "(género desconocido)"
    ```
  * Fix for vector containing only empty values
  * Finds better results when input is in other languages
  * Better handling for subspecies
  * Better handling for *Salmonellae*, especially the 'city like' serovars like *Salmonella London*
  * Understanding of highly virulent *E. coli* strains like EIEC, EPEC and STEC
  * There will be looked for uncertain results at default - these results will be returned with an informative warning
  * Manual (help page) now contains more info about the algorithms
  * Progress bar will be shown when it takes more than 3 seconds to get results
  * Support for formatted console text
  * Console will return the percentage of uncoercable input
* Function `first_isolate()`:
  * Fixed a bug where distances between dates would not be calculated right - in the `septic_patients` data set this yielded a difference of 0.15% more isolates
  * Will now use a column named like "patid" for the patient ID (argument `col_patientid`), when this argument was left blank
  * Will now use a column named like "key(...)ab" or "key(...)antibiotics" for the key antibiotics (argument `col_keyantibiotics()`), when this argument was left blank
  * Removed argument `output_logical`, the function will now always return a logical value
  * Renamed argument `filter_specimen` to `specimen_group`, although using `filter_specimen` will still work
* A note to the manual pages of the `portion` functions, that low counts can influence the outcome and that the `portion` functions may camouflage this, since they only return the portion (albeit being dependent on the `minimum` argument)
* Merged data sets `microorganisms.certe` and `microorganisms.umcg` into `microorganisms.codes`
* Function `mo_taxonomy()` now contains the kingdom too
* Reduce false positives for `is.rsi.eligible()` using the new `threshold` argument
* New colours for `scale_rsi_colours()`
* Summaries of class `mo` will now return the top 3 and the unique count, e.g. using `summary(mo)`
* Small text updates to summaries of class `rsi` and `mic`
* Function `as.rsi()`:
  * Now gives a warning when inputting MIC values
  * Now accepts high and low resistance: `"HIGH S"` will return `S`
* Frequency tables (`freq()` function):
  * Support for tidyverse quasiquotation! Now you can create frequency tables of function outcomes:
    ```r
    # Determine genus of microorganisms (mo) in `septic_patients` data set:
    # OLD WAY
    septic_patients %>%
      mutate(genus = mo_genus(mo)) %>%
      freq(genus)
    # NEW WAY
    septic_patients %>% 
      freq(mo_genus(mo))
    
    # Even supports grouping variables:
    septic_patients %>%
      group_by(gender) %>% 
      freq(mo_genus(mo))
    ```
  * Header info is now available as a list, with the `header` function
  * The argument `header` is now set to `TRUE` at default, even for markdown
  * Added header info for class `mo` to show unique count of families, genera and species
  * Now honours the `decimal.mark` setting, which just like `format` defaults to `getOption("OutDec")`
  * The new `big.mark` argument will at default be `","` when `decimal.mark = "."` and `"."` otherwise
  * Fix for header text where all observations are `NA`
  * New argument `droplevels` to exclude empty factor levels when input is a factor
  * Factor levels will be in header when present in input data (maximum of 5)
  * Fix for using `select()` on frequency tables
* Function `scale_y_percent()` now contains the `limits` argument
* Automatic argument filling for `mdro()`, `key_antibiotics()` and `eucast_rules()`
* Updated examples for resistance prediction (`resistance_predict()` function)
* Fix for `as.mic()` to support more values ending in (several) zeroes
* if using different lengths of pattern and x in `%like%`, it will now return the call

#### Other
* Updated licence text to emphasise GPL 2.0 and that this is an R package.

# AMR 0.5.0

#### New
* Repository moved to GitLab
* Function `count_all` to get all available isolates (that like all `portion_*` and `count_*` functions also supports `summarise` and `group_by`), the old `n_rsi` is now an alias of `count_all`
* Function `get_locale` to determine language for language-dependent output for some `mo_*` functions. This is now the default value for their `language` argument, by which the system language will be used at default.
* Data sets `microorganismsDT`, `microorganisms.prevDT`, `microorganisms.unprevDT` and `microorganisms.oldDT` to improve the speed of `as.mo`. They are for reference only, since they are primarily for internal use of `as.mo`.
* Function `read.4D` to read from the 4D database of the MMB department of the UMCG
* Functions `mo_authors` and `mo_year` to get specific values about the scientific reference of a taxonomic entry

#### Changed
* Functions `MDRO`, `BRMO`, `MRGN` and `EUCAST_exceptional_phenotypes` were renamed to `mdro`, `brmo`, `mrgn` and `eucast_exceptional_phenotypes`
* `EUCAST_rules` was renamed to `eucast_rules`, the old function still exists as a deprecated function
* Big changes to the `eucast_rules` function:
  * Now also applies rules from the EUCAST 'Breakpoint tables for bacteria', version 8.1, 2018, https://www.eucast.org/clinical_breakpoints/ (see Source of the function)
  * New argument `rules` to specify which rules should be applied (expert rules, breakpoints, others or all)
  * New argument `verbose` which can be set to `TRUE` to get very specific messages about which columns and rows were affected
  * Better error handling when rules cannot be applied (i.e. new values could not be inserted)
  * The number of affected values will now only be measured once per row/column combination
  * Data set `septic_patients` now reflects these changes
  * Added argument `pipe` for piperacillin (J01CA12), also to the `mdro` function
  * Small fixes to EUCAST clinical breakpoint rules
* Added column `kingdom` to the microorganisms data set, and function `mo_kingdom` to look up values
* Tremendous speed improvement for `as.mo` (and subsequently all `mo_*` functions), as empty values wil be ignored *a priori*
* Fewer than 3 characters as input for `as.mo` will return NA
* Function `as.mo` (and all `mo_*` wrappers) now supports genus abbreviations with "species" attached
  ```r
  as.mo("E. species")        # B_ESCHR
  mo_fullname("E. spp.")     # "Escherichia species"
  as.mo("S. spp")            # B_STPHY
  mo_fullname("S. species")  # "Staphylococcus species"
  ```
* Added argument `combine_IR` (TRUE/FALSE) to functions `portion_df` and `count_df`, to indicate that all values of I and R must be merged into one, so the output only consists of S vs. IR (susceptible vs. non-susceptible)
* Fix for `portion_*(..., as_percent = TRUE)` when minimal number of isolates would not be met
* Added argument `also_single_tested` for `portion_*` and `count_*` functions to also include cases where not all antibiotics were tested but at least one of the tested antibiotics includes the target antimicribial interpretation, see `?portion`
* Using `portion_*` functions now throws a warning when total available isolate is below argument `minimum`
* Functions `as.mo`, `as.rsi`, `as.mic`, `as.atc` and `freq` will not set package name as attribute anymore
* Frequency tables - `freq()`:
  * Support for grouping variables, test with:
    ```r
    septic_patients %>% 
      group_by(hospital_id) %>% 
      freq(gender)
    ```
  * Support for (un)selecting columns:
    ```r
    septic_patients %>% 
      freq(hospital_id) %>% 
      select(-count, -cum_count) # only get item, percent, cum_percent
    ```
  * Check for `hms::is.hms`
  * Now prints in markdown at default in non-interactive sessions
  * No longer adds the factor level column and sorts factors on count again
  * Support for class `difftime`
  * New argument `na`, to choose which character to print for empty values
  * New argument `header` to turn the header info off (default when `markdown = TRUE`)
  * New argument `title` to manually setbthe title of the frequency table
* `first_isolate` now tries to find columns to use as input when arguments are left blank
* Improvements for MDRO algorithm (function `mdro`)
* Data set `septic_patients` is now a `data.frame`, not a tibble anymore
* Removed diacritics from all authors (columns `microorganisms$ref` and `microorganisms.old$ref`) to comply with CRAN policy to only allow ASCII characters
* Fix for `mo_property` not working properly
* Fix for `eucast_rules` where some Streptococci would become ceftazidime R in EUCAST rule 4.5
* Support for named vectors of class `mo`, useful for `top_freq()`
* `ggplot_rsi` and `scale_y_percent` have `breaks` argument
* AI improvements for `as.mo`:
  * `"CRS"` -> *Stenotrophomonas maltophilia*
  * `"CRSM"` -> *Stenotrophomonas maltophilia*
  * `"MSSA"` -> *Staphylococcus aureus*
  * `"MSSE"` -> *Staphylococcus epidermidis*
* Fix for `join` functions
* Speed improvement for `is.rsi.eligible`, now 15-20 times faster
* In `g.test`, when `sum(x)` is below 1000 or any of the expected values is below 5, Fisher's Exact Test will be suggested
* `ab_name` will try to fall back on `as.atc` when no results are found
* Removed the addin to view data sets
* Percentages will now will rounded more logically (e.g. in `freq` function)

#### Other
* New dependency on package `crayon`, to support formatted text in the console
* Dependency `tidyr` is now mandatory (went to `Import` field) since `portion_df` and `count_df` rely on it
* Updated vignettes to comply with README


# AMR 0.4.0

#### New
* The data set `microorganisms` now contains **all microbial taxonomic data from ITIS** (kingdoms Bacteria, Fungi and Protozoa), the Integrated Taxonomy Information System, available via https://itis.gov. The data set now contains more than 18,000 microorganisms with all known bacteria, fungi and protozoa according ITIS with genus, species, subspecies, family, order, class, phylum and subkingdom. The new data set `microorganisms.old` contains all previously known taxonomic names from those kingdoms.
* New functions based on the existing function `mo_property`:
  * Taxonomic names: `mo_phylum`, `mo_class`, `mo_order`, `mo_family`, `mo_genus`, `mo_species`, `mo_subspecies`
  * Semantic names: `mo_fullname`, `mo_shortname`
  * Microbial properties: `mo_type`, `mo_gramstain`
  * Author and year: `mo_ref`
  
  They also come with support for German, Dutch, French, Italian, Spanish and Portuguese:
  ```r
  mo_gramstain("E. coli")
  # [1] "Gram negative"
  mo_gramstain("E. coli", language = "de") # German
  # [1] "Gramnegativ"
  mo_gramstain("E. coli", language = "es") # Spanish
  # [1] "Gram negativo"
  mo_fullname("S. group A", language = "pt") # Portuguese
  # [1] "Streptococcus grupo A"
  ```
  
  Furthermore, former taxonomic names will give a note about the current taxonomic name:
  ```r
  mo_gramstain("Esc blattae")
  # Note: 'Escherichia blattae' (Burgess et al., 1973) was renamed 'Shimwellia blattae' (Priest and Barker, 2010)
  # [1] "Gram negative"
  ```
* Functions `count_R`, `count_IR`, `count_I`, `count_SI` and `count_S` to selectively count resistant or susceptible isolates
  * Extra function `count_df` (which works like `portion_df`) to get all counts of S, I and R of a data set with antibiotic columns, with support for grouped variables
* Function `is.rsi.eligible` to check for columns that have valid antimicrobial results, but do not have the `rsi` class yet. Transform the columns of your raw data with: `data %>% mutate_if(is.rsi.eligible, as.rsi)`
* Functions `as.mo` and `is.mo` as replacements for `as.bactid` and `is.bactid` (since the `microoganisms` data set not only contains bacteria). These last two functions are deprecated and will be removed in a future release. The `as.mo` function determines microbial IDs using intelligent rules:
  ```r
  as.mo("E. coli")
  # [1] B_ESCHR_COL
  as.mo("MRSA")
  # [1] B_STPHY_AUR
  as.mo("S group A")
  # [1] B_STRPTC_GRA
  ```
  And with great speed too - on a quite regular Linux server from 2007 it takes us less than 0.02 seconds to transform 25,000 items:
  ```r
  thousands_of_E_colis <- rep("E. coli", 25000)
  microbenchmark::microbenchmark(as.mo(thousands_of_E_colis), unit = "s")
  # Unit: seconds
  #         min       median         max  neval
  #  0.01817717  0.01843957  0.03878077    100
  ```
* Added argument `reference_df` for `as.mo`, so users can supply their own microbial IDs, name or codes as a reference table
* Renamed all previous references to `bactid` to `mo`, like:
  * Column names inputs of `EUCAST_rules`, `first_isolate` and `key_antibiotics`
  * Column names of datasets `microorganisms` and `septic_patients`
  * All old syntaxes will still work with this version, but will throw warnings
* Function `labels_rsi_count` to print datalabels on a RSI `ggplot2` model
* Functions `as.atc` and `is.atc` to transform/look up antibiotic ATC codes as defined by the WHO. The existing function `guess_atc` is now an alias of `as.atc`.

* Function `ab_property` and its aliases: `ab_name`, `ab_tradenames`, `ab_certe`, `ab_umcg` and `ab_trivial_nl`
* Introduction to AMR as a vignette
* Removed clipboard functions as it violated the CRAN policy
* Renamed `septic_patients$sex` to `septic_patients$gender`

#### Changed
* Added three antimicrobial agents to the `antibiotics` data set: Terbinafine (D01BA02), Rifaximin (A07AA11) and Isoconazole (D01AC05)
* Added 163 trade names to the `antibiotics` data set, it now contains 298 different trade names in total, e.g.:
  ```r
  ab_official("Bactroban")
  # [1] "Mupirocin"
  ab_name(c("Bactroban", "Amoxil", "Zithromax", "Floxapen"))
  # [1] "Mupirocin" "Amoxicillin" "Azithromycin" "Flucloxacillin"
  ab_atc(c("Bactroban", "Amoxil", "Zithromax", "Floxapen"))
  # [1] "R01AX06" "J01CA04" "J01FA10" "J01CF05"
  ```
* For `first_isolate`, rows will be ignored when there's no species available
* Function `ratio` is now deprecated and will be removed in a future release, as it is not really the scope of this package
* Fix for `as.mic` for values ending in zeroes after a real number
* Small fix where *B. fragilis* would not be found in the `microorganisms.umcg` data set
* Added `prevalence` column to the `microorganisms` data set
* Added arguments `minimum` and `as_percent` to `portion_df`
* Support for quasiquotation in the functions series `count_*` and `portions_*`, and `n_rsi`. This allows to check for more than 2 vectors or columns.
  ```r
  septic_patients %>% select(amox, cipr) %>% count_IR()
  # which is the same as:
  septic_patients %>% count_IR(amox, cipr)
  
  septic_patients %>% portion_S(amcl)
  septic_patients %>% portion_S(amcl, gent)
  septic_patients %>% portion_S(amcl, gent, pita)
  ```
* Edited `ggplot_rsi` and `geom_rsi` so they can cope with `count_df`. The new `fun` argument has value `portion_df` at default, but can be set to `count_df`.
* Fix for `ggplot_rsi` when the `ggplot2` package was not loaded
* Added datalabels function `labels_rsi_count` to `ggplot_rsi`
* Added possibility to set any argument to `geom_rsi` (and `ggplot_rsi`) so you can set your own preferences
* Fix for joins, where predefined suffices would not be honoured
* Added argument `quote` to the `freq` function
* Added generic function `diff` for frequency tables
* Added longest en shortest character length in the frequency table (`freq`) header of class `character`
* Support for types (classes) list and matrix for `freq`
  ```r
  my_matrix = with(septic_patients, matrix(c(age, gender), ncol = 2))
  freq(my_matrix)
  ```
  For lists, subsetting is possible:
  ```r
  my_list = list(age = septic_patients$age, gender = septic_patients$gender)
  my_list %>% freq(age)
  my_list %>% freq(gender)
  ```

#### Other
* More unit tests to ensure better integrity of functions

# AMR 0.3.0

#### New
* **BREAKING**: `rsi_df` was removed in favour of new functions `portion_R`, `portion_IR`, `portion_I`, `portion_SI` and `portion_S` to selectively calculate resistance or susceptibility. These functions are 20 to 30 times faster than the old `rsi` function. The old function still works, but is deprecated.
  * New function `portion_df` to get all portions of S, I and R of a data set with antibiotic columns, with support for grouped variables
* **BREAKING**: the methodology for determining first weighted isolates was changed. The antibiotics that are compared between isolates (call *key antibiotics*) to include more first isolates (afterwards called first *weighted* isolates) are now as follows:
  * Universal: amoxicillin, amoxicillin/clavlanic acid, cefuroxime, piperacillin/tazobactam, ciprofloxacin,  trimethoprim/sulfamethoxazole
  * Gram-positive: vancomycin, teicoplanin, tetracycline, erythromycin, oxacillin, rifampicin
  * Gram-negative: gentamicin, tobramycin, colistin, cefotaxime, ceftazidime, meropenem
* Support for `ggplot2`
  * New functions `geom_rsi`, `facet_rsi`, `scale_y_percent`, `scale_rsi_colours` and `theme_rsi`
  * New wrapper function `ggplot_rsi` to apply all above functions on a data set:
    * `septic_patients %>% select(tobr, gent) %>% ggplot_rsi` will show portions of S, I and R immediately in a pretty plot
    * Support for grouped variables, see `?ggplot_rsi`
* Determining bacterial ID:
  * New functions `as.bactid` and `is.bactid` to transform/ look up microbial ID's.
  * The existing function `guess_bactid` is now an alias of `as.bactid`
  * New Becker classification for *Staphylococcus* to categorise them into Coagulase Negative *Staphylococci* (CoNS) and Coagulase Positve *Staphylococci* (CoPS)
  * New Lancefield classification for *Streptococcus* to categorise them into Lancefield groups
* For convience, new descriptive statistical functions `kurtosis` and `skewness` that are lacking in base R - they are generic functions and have support for vectors, data.frames and matrices
* Function `g.test` to perform the Χ<sup>2</sup> distributed [*G*-test](https://en.wikipedia.org/wiki/G-test), which use is the same as `chisq.test`
* ~~Function `ratio` to transform a vector of values to a preset ratio~~
  * ~~For example: `ratio(c(10, 500, 10), ratio = "1:2:1")` would return `130, 260, 130`~~
* Support for Addins menu in RStudio to quickly insert `%in%` or `%like%` (and give them keyboard shortcuts), or to view the datasets that come with this package
* Function `p.symbol` to transform p values to their related symbols: `0 '***' 0.001 '**' 0.01 '*' 0.05 '.' 0.1 ' ' 1`
* Functions `clipboard_import` and `clipboard_export` as helper functions to quickly copy and paste from/to software like Excel and SPSS. These functions use the `clipr` package, but are a little altered to also support headless Linux servers (so you can use it in RStudio Server)
* New for frequency tables (function `freq`):
  * A vignette to explain its usage
  * Support for `rsi` (antimicrobial resistance) to use as input
  * Support for `table` to use as input: `freq(table(x, y))`
  * Support for existing functions `hist` and `plot` to use a frequency table as input: `hist(freq(df$age))`
  * Support for `as.vector`, `as.data.frame`, `as_tibble` and `format`
  * Support for quasiquotation: `freq(mydata, mycolumn)` is the same as `mydata %>% freq(mycolumn)`
  * Function `top_freq` function to return the top/below *n* items as vector
  * Header of frequency tables now also show Mean Absolute Deviaton (MAD) and Interquartile Range (IQR)
  * Possibility to globally set the default for the amount of items to print, with `options(max.print.freq = n)` where *n* is your preset value

#### Changed
* Improvements for forecasting with `resistance_predict` and added more examples
* More antibiotics added as arguments for EUCAST rules
* Updated version of the `septic_patients` data set to better reflect the reality
* Pretty printing for tibbles removed as it is not really the scope of this package
* Printing of `mic` and `rsi` classes now returns all values - use `freq` to check distributions
* Improved speed of key antibiotics comparison for determining first isolates
* Column names for the `key_antibiotics` function are now generic: 6 for broadspectrum ABs, 6 for Gram-positive specific and 6 for Gram-negative specific ABs
* Speed improvement for the `abname` function
* `%like%` now supports multiple patterns
* Frequency tables are now actual `data.frame`s with altered console printing to make it look like a frequency table. Because of this, the argument `toConsole` is not longer needed.
* Fix for `freq` where the class of an item would be lost
* Small translational improvements to the `septic_patients` dataset and the column `bactid` now has the new class `"bactid"`
* Small improvements to the `microorganisms` dataset (especially for *Salmonella*) and the column `bactid` now has the new class `"bactid"`
* Combined MIC/RSI values will now be coerced by the `rsi` and `mic` functions:
  * `as.rsi("<=0.002; S")` will return `S`
  * `as.mic("<=0.002; S")` will return `<=0.002`
* Now possible to coerce MIC values with a space between operator and value, i.e. `as.mic("<= 0.002")` now works
* Classes `rsi` and `mic` do not add the attribute `package.version` anymore
* Added `"groups"` option for `atc_property(..., property)`. It will return a vector of the ATC hierarchy as defined by the [WHO](https://www.whocc.no/atc/structure_and_principles/). The new function `atc_groups` is a convenient wrapper around this.
* Build-in host check for `atc_property` as it requires the host set by `url` to be responsive
* Improved `first_isolate` algorithm to exclude isolates where bacteria ID or genus is unavailable
* Fix for warning *hybrid evaluation forced for row_number* ([`924b62`](https://github.com/tidyverse/dplyr/commit/924b62)) from the `dplyr` package v0.7.5 and above
* Support for empty values and for 1 or 2 columns as input for `guess_bactid` (now called `as.bactid`)
  * So `yourdata %>% select(genus, species) %>% as.bactid()` now also works
* Other small fixes

#### Other
* Added integration tests (check if everything works as expected) for all releases of R 3.1 and higher
  * Linux and macOS: https://travis-ci.org/msberends/AMR
  * Windows: https://ci.appveyor.com/project/msberends/amr
* Added thesis advisors to DESCRIPTION file

# AMR 0.2.0

#### New
* Full support for Windows, Linux and macOS
* Full support for old R versions, only R-3.0.0 (April 2013) or later is needed (needed packages may have other dependencies)
* Function `n_rsi` to count cases where antibiotic test results were available, to be used in conjunction with `dplyr::summarise`, see ?rsi
* Function `guess_bactid` to **determine the ID** of a microorganism based on genus/species or known abbreviations like MRSA
* Function `guess_atc` to **determine the ATC** of an antibiotic based on name, trade name, or known abbreviations
* Function `freq` to create **frequency tables**, with additional info in a header
* Function `MDRO` to **determine Multi Drug Resistant Organisms (MDRO)** with support for country-specific guidelines.
  * [Exceptional resistances defined by EUCAST](https://www.eucast.org/expert_rules_and_intrinsic_resistance/) are also supported instead of countries alone
  * Functions `BRMO` and `MRGN` are wrappers for Dutch and German guidelines, respectively
* New algorithm to determine weighted isolates, can now be `"points"` or `"keyantibiotics"`, see `?first_isolate`
* New print format for `tibble`s and `data.table`s

#### Changed
* Fixed `rsi` class for vectors that contain only invalid antimicrobial interpretations
* Renamed dataset `ablist` to `antibiotics`
* Renamed dataset `bactlist` to `microorganisms`
* Added common abbreviations and trade names to the `antibiotics` dataset
* Added more microorganisms to the `microorganisms` dataset
* Added analysis examples on help page of dataset `septic_patients`
* Added support for character vector in `join` functions
* Added warnings when a join results in more rows after than before the join
* Altered `%like%` to make it case insensitive
* For arguments of functions `first_isolate` and `EUCAST_rules` column names are now case-insensitive
* Functions `as.rsi` and `as.mic` now add the package name and version as attributes

#### Other
* Expanded `README.md` with more examples
* Added [ORCID](https://orcid.org) of authors to DESCRIPTION file
* Added unit testing with the `testthat` package
* Added build tests for Linux and macOS using Travis CI (https://travis-ci.org/msberends/AMR)
* Added line coverage checking using CodeCov (https://codecov.io/gh/msberends/AMR/tree/master/R)

# AMR 0.1.1

* `EUCAST_rules` applies for amoxicillin even if ampicillin is missing
* Edited column names to comply with GLIMS, the laboratory information system
* Added more valid MIC values
* Renamed 'Daily Defined Dose' to 'Defined Daily Dose'
* Added barplots for `rsi` and `mic` classes

# AMR 0.1.0

* First submission to CRAN.
