## Supplementary material for "AMR - An R Package for Working with Antimicrobial Resistance Data": Source package: aa_helper_functions.R

```
# ==================================================================== #
# TITLE                                                                #
# Antimicrobial Resistance (AMR) Analysis for R                        #
#                                                                      #
# SOURCE                                                               #
# https://github.com/msberends/AMR                                     #
#                                                                      #
# LICENCE                                                              #
# (c) 2018-2021 Berends MS, Luz CF et al.                              #
# Developed at the University of Groningen, the Netherlands, in        #
# collaboration with non-profit organisations Certe Medical            #
# Diagnostics & Advice, and University Medical Center Groningen.       #
#                                                                      #
# This R package is free software; you can freely use and distribute   #
# it for both personal and commercial purposes under the terms of the  #
# GNU General Public License version 2.0 (GNU GPL-2), as published by  #
# the Free Software Foundation.                                        #
# We created this package for both routine data analysis and academic  #
# research and it was publicly released in the hope that it will be    #
# useful, but it comes WITHOUT ANY WARRANTY OR LIABILITY.              #
#                                                                      #
# Visit our website for the full manual and a complete tutorial about  #
# how to conduct AMR analysis: https://msberends.github.io/AMR/        #
# ==================================================================== #

# faster implementation of left_join than using merge() by poorman - we use match():
pm_left_join <- function(x, y, by = NULL, suffix = c(".x", ".y")) {
  if (is.null(by)) {
    by <- intersect(names(x), names(y))[1L]
    if (is.na(by)) {
      stop_("no common column found for pm_left_join()")
    }
    pm_join_message(by)
  } else if (!is.null(names(by))) {
    by <- unname(c(names(by), by))
  }
  if (length(by) == 1) {
    by <- rep(by, 2)
  }

  int_x <- colnames(x) %in% colnames(y) & colnames(x) != by[1]
  int_y <- colnames(y) %in% colnames(x) & colnames(y) != by[2]
  colnames(x)[int_x] <- paste0(colnames(x)[int_x], suffix[1L])
  colnames(y)[int_y] <- paste0(colnames(y)[int_y], suffix[2L])

  merged <- cbind(x,
                  y[match(x[, by[1], drop = TRUE],
                          y[, by[2], drop = TRUE]),
                    colnames(y)[!colnames(y) %in% colnames(x) & !colnames(y) == by[2]],
                    drop = FALSE])

  rownames(merged) <- NULL
  merged
}

quick_case_when <- function(...) {
  vectors <- list(...)
  split <- lapply(vectors, function(x) unlist(strsplit(paste(deparse(x), collapse = ""), "~", fixed = TRUE)))
  for (i in seq_len(length(vectors))) {
    if (eval(parse(text = split[[i]][1]), envir = parent.frame())) {
      return(eval(parse(text = split[[i]][2]), envir = parent.frame()))
    }
  }
  return(NA)
}

# No export, no Rd
addin_insert_in <- function() {
  import_fn("insertText", "rstudioapi")(" %in% ")
}

# No export, no Rd
addin_insert_like <- function() {
  import_fn("insertText", "rstudioapi")(" %like% ")
}

check_dataset_integrity <- function() {
  # check if user overwrote our data sets in their global environment
  data_in_pkg <- data(package = "AMR", envir = asNamespace("AMR"))$results[, "Item", drop = TRUE]
  data_in_globalenv <- ls(envir = globalenv())
  overwritten <- data_in_pkg[data_in_pkg %in% data_in_globalenv]
  # exception for example_isolates
  overwritten <- overwritten[overwritten != "example_isolates"]
  stop_if(length(overwritten) > 0,
          "the following data set is overwritten by your global environment and prevents the AMR package from working correctly:\n",
          paste0("'", overwritten, "'", collapse = ", "),
          ".\nPlease rename your object before using this function.", call = FALSE)
  # check if other packages did not overwrite our data sets
  tryCatch({
    check_microorganisms <- all(c("mo", "fullname", "kingdom", "phylum",
                                  "class", "order", "family", "genus",
                                  "species", "subspecies", "rank",
                                  "species_id", "source", "ref", "prevalence") %in% colnames(microorganisms),
                                na.rm = TRUE)
    check_antibiotics <- all(c("ab", "atc", "cid", "name", "group",
                               "atc_group1", "atc_group2", "abbreviations",
                               "synonyms", "oral_ddd", "oral_units",
                               "iv_ddd", "iv_units", "loinc") %in% colnames(antibiotics),
                             na.rm = TRUE)
  }, error = function(e) {
    # package not yet loaded
    require("AMR")
  })
  stop_if(!check_microorganisms | !check_antibiotics,
          "the data set `microorganisms` or `antibiotics` was overwritten in your environment because another package with the same object names was loaded _after_ the AMR package, preventing the AMR package from working correctly. Please load the AMR package last.")
  invisible(TRUE)
}

search_type_in_df <- function(x, type, info = TRUE) {
  meet_criteria(x, allow_class = "data.frame")
  meet_criteria(type, allow_class = "character", has_length = 1)
  
  # try to find columns based on type
  found <- NULL

  # remove attributes from other packages
  x <- as.data.frame(x, stringsAsFactors = FALSE)
  colnames(x) <- trimws(colnames(x))

  # -- mo
  if (type == "mo") {
    if (any(vapply(FUN.VALUE = logical(1), x, is.mo))) {
      found <- sort(colnames(x)[vapply(FUN.VALUE = logical(1), x, is.mo)])[1]
    } else if ("mo" %in% colnames(x) &
               suppressWarnings(
                 all(x$mo %in% c(NA,
                                 microorganisms$mo,
                                 microorganisms.translation$mo_old)))) {
      found <- "mo"
    } else if (any(colnames(x) %like% "^(mo|microorganism|organism|bacteria|ba[ck]terie)s?$")) {
      found <- sort(colnames(x)[colnames(x) %like% "^(mo|microorganism|organism|bacteria|ba[ck]terie)s?$"])[1]
    } else if (any(colnames(x) %like% "^(microorganism|organism|bacteria|ba[ck]terie)")) {
      found <- sort(colnames(x)[colnames(x) %like% "^(microorganism|organism|bacteria|ba[ck]terie)"])[1]
    } else if (any(colnames(x) %like% "species")) {
      found <- sort(colnames(x)[colnames(x) %like% "species"])[1]
    }

  }
  # -- key antibiotics
  if (type == "keyantibiotics") {
    if (any(colnames(x) %like% "^key.*(ab|antibiotics)")) {
      found <- sort(colnames(x)[colnames(x) %like% "^key.*(ab|antibiotics)"])[1]
    }
  }
  # -- date
  if (type == "date") {
    if (any(colnames(x) %like% "^(specimen date|specimen_date|spec_date)")) {
      # WHONET support
      found <- sort(colnames(x)[colnames(x) %like% "^(specimen date|specimen_date|spec_date)"])[1]
      if (!any(class(pm_pull(x, found)) %in% c("Date", "POSIXct"))) {
        stop(font_red(paste0("Found column '", font_bold(found), "' to be used as input for `col_", type,
                             "`, but this column contains no valid dates. Transform its values to valid dates first.")),
             call. = FALSE)
      }
    } else if (any(vapply(FUN.VALUE = logical(1), x, function(x) inherits(x, c("Date", "POSIXct"))))) {
      found <- sort(colnames(x)[vapply(FUN.VALUE = logical(1), x, function(x) inherits(x, c("Date", "POSIXct")))])[1]
    }
  }
  # -- patient id
  if (type == "patient_id") {
    if (any(colnames(x) %like% "^(identification |patient|patid)")) {
      found <- sort(colnames(x)[colnames(x) %like% "^(identification |patient|patid)"])[1]
    }
  }
  # -- specimen
  if (type == "specimen") {
    if (any(colnames(x) %like% "(specimen type|spec_type)")) {
      found <- sort(colnames(x)[colnames(x) %like% "(specimen type|spec_type)"])[1]
    } else if (any(colnames(x) %like% "^(specimen)")) {
      found <- sort(colnames(x)[colnames(x) %like% "^(specimen)"])[1]
    }
  }
  # -- UTI (urinary tract infection)
  if (type == "uti") {
    if (any(colnames(x) == "uti")) {
      found <- colnames(x)[colnames(x) == "uti"][1]
    } else if (any(colnames(x) %like% "(urine|urinary)")) {
      found <- sort(colnames(x)[colnames(x) %like% "(urine|urinary)"])[1]
    }
    if (!is.null(found)) {
      # this column should contain logicals
      if (!is.logical(x[, found, drop = TRUE])) {
        message_("Column '", font_bold(found), "' found as input for `col_", type,
                 "`, but this column does not contain 'logical' values (TRUE/FALSE) and was ignored.",
                 add_fn = font_red)
        found <- NULL
      }
    }
  }
  
  if (!is.null(found) & info == TRUE) {
    if (message_not_thrown_before(fn = paste0("search_", type))) {
      msg <- paste0("Using column '", font_bold(found), "' as input for `col_", type, "`.")
      if (type %in% c("keyantibiotics", "specimen")) {
        msg <- paste(msg, "Use", font_bold(paste0("col_", type), "= FALSE"), "to prevent this.")
      }
      message_(msg)
      remember_thrown_message(fn = paste0("search_", type))
    }
  }
  found
}

is_possibly_regex <- function(x) {
  tryCatch(vapply(FUN.VALUE = character(1), strsplit(x, ""),
                  function(y) any(y %in% c("$", "(", ")", "*", "+", "-", ".", "?", "[", "]", "^", "{", "|", "}", "\\"), na.rm = TRUE)),
           error = function(e) rep(TRUE, length(x)))
}

stop_ifnot_installed <- function(package) {
  # no "utils::installed.packages()" since it requires non-staged install since R 3.6.0
  # https://developer.r-project.org/Blog/public/2019/02/14/staged-install/index.html
  vapply(FUN.VALUE = character(1), package, function(pkg)
    tryCatch(get(".packageName", envir = asNamespace(pkg)),
             error = function(e) {
               if (package == "rstudioapi") {
                 stop("This function only works in RStudio.", call. = FALSE)
               } else if (pkg != "base") {
                 stop("This requires the '", pkg, "' package.",
                      "\nTry to install it with: install.packages(\"", pkg, "\")",
                      call. = FALSE)
               }
             }))
  return(invisible())
}

import_fn <- function(name, pkg, error_on_fail = TRUE) {
  if (isTRUE(error_on_fail)) {
    stop_ifnot_installed(pkg)
  }
  tryCatch(
    # don't use get() to avoid fetching non-API functions 
    getExportedValue(name = name, ns = asNamespace(pkg)),
    error = function(e) {
      if (isTRUE(error_on_fail)) {
        stop_("function ", name, "() is not an exported object from package '", pkg,
              "'. Please create an issue at https://github.com/msberends/AMR/issues. Many thanks!",
              call = FALSE)
      } else {
        return(NULL)
      }
    })
}

# this alternative wrapper to the message(), warning() and stop() functions:
# - wraps text to never break lines within words
# - ignores formatted text while wrapping
# - adds indentation dependent on the type of message (such as NOTE)
# - can add additional formatting functions like blue or bold text
word_wrap <- function(...,
                      add_fn = list(), 
                      as_note = FALSE,
                      width = 0.95 * getOption("width"),
                      extra_indent = 0) {
  msg <- paste0(c(...), collapse = "")
  
  if (isTRUE(as_note)) {
    msg <- paste0("NOTE: ", gsub("^note:? ?", "", msg, ignore.case = TRUE))
  }
  
  if (msg %like% "\n") {
    # run word_wraps() over every line here, bind them and return again
    return(paste0(vapply(FUN.VALUE = character(1),
                         trimws(unlist(strsplit(msg, "\n")), which = "right"),
                         word_wrap, 
                         add_fn = add_fn,
                         as_note = FALSE,
                         width = width, 
                         extra_indent = extra_indent),
                  collapse = "\n"))
  }
  
  # we need to correct for already applied style, that adds text like "\033[31m\"
  msg_stripped <- font_stripstyle(msg)
  # where are the spaces now?
  msg_stripped_wrapped <- paste0(strwrap(msg_stripped,
                                         simplify = TRUE,
                                         width = width),
                                 collapse = "\n")
  msg_stripped_wrapped <- paste0(unlist(strsplit(msg_stripped_wrapped, "(\n|\\*\\|\\*)")),
                                 collapse = "\n")
  msg_stripped_spaces <- which(unlist(strsplit(msg_stripped, "")) == " ")
  msg_stripped_wrapped_spaces <- which(unlist(strsplit(msg_stripped_wrapped, "")) != "\n")
  # so these are the indices of spaces that need to be replaced
  replace_spaces <- which(!msg_stripped_spaces %in% msg_stripped_wrapped_spaces)
  # put it together
  msg <- unlist(strsplit(msg, " "))
  msg[replace_spaces] <- paste0(msg[replace_spaces], "\n")
  msg <- paste0(msg, collapse = " ")
  msg <- gsub("\n ", "\n", msg, fixed = TRUE)
  
  if (msg_stripped %like% "^NOTE: ") {
    indentation <- 6 + extra_indent
  } else if (msg_stripped %like% "^=> ") {
    indentation <- 3 + extra_indent
  } else {
    indentation <- 0 + extra_indent
  }
  msg <- gsub("\n", paste0("\n", strrep(" ", indentation)), msg, fixed = TRUE)
  # remove trailing empty characters
  msg <- gsub("(\n| )+$", "", msg)
  
  if (length(add_fn) > 0) {
    if (!is.list(add_fn)) {
      add_fn <- list(add_fn)
    }
    for (i in seq_len(length(add_fn))) {
      msg <- add_fn[[i]](msg)
    }
  }
  
  # format backticks
  msg <- gsub("(`.+?`)", font_grey_bg("\\1"), msg)
  
  msg
}

message_ <- function(...,
                     appendLF = TRUE,
                     add_fn = list(font_blue),
                     as_note = TRUE) {
  message(word_wrap(..., 
                    add_fn = add_fn,
                    as_note = as_note),
          appendLF = appendLF)
}

warning_ <- function(...,
                     add_fn = list(),
                     immediate = FALSE,
                     call = TRUE) {
  warning(word_wrap(..., 
                    add_fn = add_fn,
                    as_note = FALSE),
          immediate. = immediate,
          call. = call)
}

# this alternative to the stop() function:
# - adds the function name where the error was thrown
# - wraps text to never break lines within words
stop_ <- function(..., call = TRUE) {
  msg <- paste0(c(...), collapse = "")
  if (!isFALSE(call)) {
    if (isTRUE(call)) {
      call <- as.character(sys.call(-1)[1])
    } else {
      # so you can go back more than 1 call, as used in rsi_calc(), that now throws a reference to e.g. n_rsi()
      call <- as.character(sys.call(call)[1])
    }
    msg <- paste0("in ", call, "(): ", msg)
  }
  msg <- word_wrap(msg, add_fn = list(), as_note = FALSE)
  stop(msg, call. = FALSE)
}

stop_if <- function(expr, ..., call = TRUE) {
  if (isTRUE(expr)) {
    if (isTRUE(call)) {
      call <- -1
    }
    if (!isFALSE(call)) {
      # since we're calling stop_(), which is another call
      call <- call - 1
    }
    stop_(..., call = call)
  }
}

stop_ifnot <- function(expr, ..., call = TRUE) {
  if (isFALSE(expr)) {
    if (isTRUE(call)) {
      call <- -1
    }
    if (!isFALSE(call)) {
      # since we're calling stop_(), which is another call
      call <- call - 1
    }
    stop_(..., call = call)
  }
}

"%or%" <- function(x, y) {
  if (is.null(x) | is.null(y)) {
    if (is.null(x)) {
      return(y)
    } else {
      return(x)
    }
  }
  ifelse(!is.na(x),
         x,
         ifelse(!is.na(y), y, NA))
}

class_integrity_check <- function(value, type, check_vector) {
  if (!all(value[!is.na(value)] %in% check_vector)) {
    warning_(paste0("invalid ", type, ", NA generated"), call = FALSE)
    value[!value %in% check_vector] <- NA
  }
  value
}

# transforms data set to data.frame with only ASCII values, to comply with CRAN policies
dataset_UTF8_to_ASCII <- function(df) {
  trans <- function(vect) {
    iconv(vect, from = "UTF-8", to = "ASCII//TRANSLIT")
  }
  df <- as.data.frame(df, stringsAsFactors = FALSE)
  for (i in seq_len(NCOL(df))) {
    col <- df[, i]
    if (is.list(col)) {
      col <- lapply(col, function(j) trans(j))
      df[, i] <- list(col)
    } else {
      if (is.factor(col)) {
        levels(col) <- trans(levels(col))
      } else if (is.character(col)) {
        col <- trans(col)
      } else {
        col
      }
      df[, i] <- col
    }
  }
  df
}

# for eucast_rules() and mdro(), creates markdown output with URLs and names
create_ab_documentation <- function(ab) {
  ab_names <- ab_name(ab, language = NULL, tolower = TRUE)
  ab <- ab[order(ab_names)]
  ab_names <- ab_names[order(ab_names)]
  atcs <- ab_atc(ab)
  atcs[!is.na(atcs)] <- paste0("[", atcs[!is.na(atcs)], "](", ab_url(ab[!is.na(atcs)]), ")")
  atcs[is.na(atcs)] <- "no ATC code"
  out <- paste0(ab_names, " (`", ab, "`, ", atcs, ")", collapse = ", ")
  substr(out, 1, 1) <- toupper(substr(out, 1, 1))
  out
}

# a check for every single argument in all functions
meet_criteria <- function(object,
                          allow_class = NULL,
                          has_length = NULL,
                          looks_like = NULL,
                          is_in = NULL,
                          contains_column_class = NULL,
                          allow_NULL = FALSE,
                          allow_NA = FALSE,
                          ignore.case = FALSE,
                          .call_depth = 0) { # depth in calling

  obj_name <- deparse(substitute(object))
  call_depth <- -2 - abs(.call_depth)

  if (is.null(object)) {
    stop_if(allow_NULL == FALSE, "argument `", obj_name, "` must not be NULL", call = call_depth)
    return(invisible())
  }
  if (is.null(dim(object)) && length(object) == 1 && suppressWarnings(is.na(object))) { # suppressWarnings for functions
    stop_if(allow_NA == FALSE, "argument `", obj_name, "` must not be NA", call = call_depth)
    return(invisible())
  }

  vector_or <- function(v, quotes) {
    if (length(v) == 1) {
      return(paste0(ifelse(quotes, '"', ""), v, ifelse(quotes, '"', "")))
    }
    # all commas except for last item, so will become '"val1", "val2", "val3" or "val4"'
    paste0(paste0(ifelse(quotes, '"', ""), v[seq_len(length(v) - 1)], ifelse(quotes, '"', ""), collapse = ", "),
           " or ", paste0(ifelse(quotes, '"', ""), v[length(v)], ifelse(quotes, '"', "")))
  }

  if (!is.null(allow_class)) {
    stop_ifnot(inherits(object, allow_class), "argument `", obj_name,
               "` must ", # ifelse(allow_NULL, "be NULL or must ", ""),
               "be of class ", vector_or(allow_class, quotes = TRUE),
               ", not \"", paste(class(object), collapse = "/"), "\"",
               call = call_depth)
    # check data.frames for data
    if (inherits(object, "data.frame")) {
      stop_if(any(dim(object) == 0),
              "the data provided in argument `", obj_name,
              "` must contain rows and columns (current dimensions: ",
              paste(dim(object), collapse = "x"), ")",
              call = call_depth)
    }
  }
  if (!is.null(has_length)) {
    stop_ifnot(length(object) %in% has_length, "argument `", obj_name,
               "` must ", # ifelse(allow_NULL, "be NULL or must ", ""),
               "be of length ", vector_or(has_length, quotes = FALSE),
               ", not ", length(object),
               call = call_depth)
  }
  if (!is.null(looks_like)) {
    stop_ifnot(object %like% looks_like, "argument `", obj_name,
               "` must ", # ifelse(allow_NULL, "be NULL or must ", ""),
               "resemble the regular expression \"", looks_like, "\"",
               call = call_depth)
  }
  if (!is.null(is_in)) {
    if (ignore.case == TRUE) {
      object <- tolower(object)
      is_in <- tolower(is_in)
    }
    stop_ifnot(all(object %in% is_in, na.rm = TRUE), "argument `", obj_name,
               "` must be ",
               ifelse(!is.null(has_length) && length(has_length) == 1 && has_length == 1, "one of: ", ""),
               vector_or(is_in, quotes = TRUE),
               ", not ", paste0("\"", object, "\"", collapse = "/"), "",
               call = call_depth)
  }
  if (!is.null(contains_column_class)) {
    stop_ifnot(any(vapply(FUN.VALUE = logical(1), 
                          object,
                          function(col, columns_class = contains_column_class) {
                            inherits(col, columns_class)
                          }), na.rm = TRUE),
               "the data provided in argument `", obj_name,
               "` must contain at least one column of class <", contains_column_class, ">. ",
               "See ?as.", contains_column_class, ".",
               call = call_depth)
  }
  return(invisible())
}

get_current_data <- function(arg_name, call) {
  if (as.double(R.Version()$major) + (as.double(R.Version()$minor) / 10) < 3.2) {
    if (is.na(arg_name)) {
      warning_("this function can only be used in R >= 3.2", call = call)
      return(data.frame())
    } else {
      stop_("argument `", arg_name, "` is missing with no default", call = call)
    }
  }
  
  # try a (base R) method, by going over the complete system call stack with sys.frames()
  not_set <- TRUE
  frms <- lapply(sys.frames(), function(el) {
    if (".Generic" %in% names(el)) {
      if (tryCatch(not_set == TRUE && ".data" %in% names(el) && is.data.frame(el$`.data`), error = function(e) FALSE)) {
        # dplyr? - an element `.data` will be in the system call stack
        not_set <<- FALSE
        el$`.data`
      } else if (tryCatch(not_set == TRUE && any(c("x", "xx") %in% names(el)), error = function(e) FALSE)) {
        # otherwise try base R:
        # an element `x` will be in this environment for only cols, e.g. `example_isolates[, carbapenems()]`
        # an element `xx` will be in this environment for rows + cols, e.g. `example_isolates[c(1:3), carbapenems()]`
        if (tryCatch(is.data.frame(el$xx), error = function(e) FALSE)) {
          not_set <<- FALSE
          el$xx
        } else if (tryCatch(is.data.frame(el$x))) {
          not_set <<- FALSE
          el$x
        } else {
          NULL
        }
      } else {
        NULL
      }
    } else {
      NULL
    }
  })
  vars_df <- tryCatch(frms[[which(!vapply(FUN.VALUE = logical(1), frms, is.null))]], error = function(e) NULL)
  if (is.data.frame(vars_df)) {
    return(vars_df)
  }
  
  # nothing worked, so:
  if (is.na(arg_name)) {
    stop_("this function must be used inside valid dplyr selection verbs or inside a data.frame call", 
          call = call)
  } else {
    stop_("argument `", arg_name, "` is missing with no default ",
          "or function not used inside a valid dplyr verb", 
          call = call)
  }
}

unique_call_id <- function(entire_session = FALSE) {
  if (entire_session == TRUE) {
    c(envir = "session",
      call = "session")
  } else {
    # combination of environment ID (like "0x7fed4ee8c848")
    # and highest system call
    c(envir = gsub("<environment: (.*)>", "\\1", utils::capture.output(sys.frames()[[1]])),
      call = paste0(deparse(sys.calls()[[1]]), collapse = ""))
  }
}

remember_thrown_message <- function(fn, entire_session = FALSE) {
  # this is to prevent that messages/notes will be printed for every dplyr group
  # e.g. this would show a msg 4 times: example_isolates %>% group_by(hospital_id) %>% filter(mo_is_gram_negative())
  assign(x = paste0("thrown_msg_", fn),
         value = unique_call_id(entire_session = entire_session),
         envir = pkg_env)
}

message_not_thrown_before <- function(fn, entire_session = FALSE) {
  is.null(pkg_env[[paste0("thrown_msg_", fn)]]) || !identical(pkg_env[[paste0("thrown_msg_", fn)]], unique_call_id(entire_session))
}

reset_all_thrown_messages <- function() {
  # for unit tests, where the environment and highest system call do not change
  pkg_env_contents <- ls(envir = pkg_env)
  rm(list = pkg_env_contents[pkg_env_contents %like% "^thrown_msg_"],
     envir = pkg_env)
}

has_colour <- function() {
  # this is a base R version of crayon::has_color, but disables colours on emacs
  
  if (Sys.getenv("EMACS") != "" || Sys.getenv("INSIDE_EMACS") != "") {
    # disable on emacs, which only supports 8 colours
    return(FALSE)
  }
  enabled <- getOption("crayon.enabled")
  if (!is.null(enabled)) {
    return(isTRUE(enabled))
  }
  rstudio_with_ansi_support <- function(x) {
    if (Sys.getenv("RSTUDIO", "") == "") {
      return(FALSE)
    }
    if ((cols <- Sys.getenv("RSTUDIO_CONSOLE_COLOR", "")) != "" && !is.na(as.numeric(cols))) {
      return(TRUE)
    }
    tryCatch(get("isAvailable", envir = asNamespace("rstudioapi"))(), error = function(e) return(FALSE)) &&
      tryCatch(get("hasFun", envir = asNamespace("rstudioapi"))("getConsoleHasColor"), error = function(e) return(FALSE))
  }
  if (rstudio_with_ansi_support() && sink.number() == 0) {
    return(TRUE)
  }
  if (!isatty(stdout())) {
    return(FALSE)
  }
  if (tolower(Sys.info()["sysname"]) == "windows") {
    if (Sys.getenv("ConEmuANSI") == "ON") {
      return(TRUE)
    }
    if (Sys.getenv("CMDER_ROOT") != "") {
      return(TRUE)
    }
    return(FALSE)
  }
  if ("COLORTERM" %in% names(Sys.getenv())) {
    return(TRUE)
  }
  if (Sys.getenv("TERM") == "dumb") {
    return(FALSE)
  }
  grepl(pattern = "^screen|^xterm|^vt100|color|ansi|cygwin|linux",
        x = Sys.getenv("TERM"),
        ignore.case = TRUE,
        perl = TRUE)
}

# set colours if console has_colour()
try_colour <- function(..., before, after, collapse = " ") {
  txt <- paste0(unlist(list(...)), collapse = collapse)
  if (isTRUE(has_colour())) {
    if (is.null(collapse)) {
      paste0(before, txt, after, collapse = NULL)
    } else {
      paste0(before, txt, after, collapse = "")
    }
  } else {
    txt
  }
}
font_black <- function(..., collapse = " ") {
  try_colour(..., before = "\033[38;5;232m", after = "\033[39m", collapse = collapse)
}
font_blue <- function(..., collapse = " ") {
  try_colour(..., before = "\033[34m", after = "\033[39m", collapse = collapse)
}
font_green <- function(..., collapse = " ") {
  try_colour(..., before = "\033[32m", after = "\033[39m", collapse = collapse)
}
font_magenta <- function(..., collapse = " ") {
  try_colour(..., before = "\033[35m", after = "\033[39m", collapse = collapse)
}
font_red <- function(..., collapse = " ") {
  try_colour(..., before = "\033[31m", after = "\033[39m", collapse = collapse)
}
font_silver <- function(..., collapse = " ") {
  try_colour(..., before = "\033[90m", after = "\033[39m", collapse = collapse)
}
font_white <- function(..., collapse = " ") {
  try_colour(..., before = "\033[37m", after = "\033[39m", collapse = collapse)
}
font_yellow <- function(..., collapse = " ") {
  try_colour(..., before = "\033[33m", after = "\033[39m", collapse = collapse)
}
font_subtle <- function(..., collapse = " ") {
  try_colour(..., before = "\033[38;5;246m", after = "\033[39m", collapse = collapse)
}
font_grey <- function(..., collapse = " ") {
  try_colour(..., before = "\033[38;5;249m", after = "\033[39m", collapse = collapse)
}
font_grey_bg <- function(..., collapse = " ") {
  try_colour(..., before = "\033[48;5;254m", after = "\033[49m", collapse = collapse)
}
font_green_bg <- function(..., collapse = " ") {
  try_colour(..., before = "\033[42m", after = "\033[49m", collapse = collapse)
}
font_rsi_R_bg <- function(..., collapse = " ") {
  try_colour(..., before = "\033[48;5;202m", after = "\033[49m", collapse = collapse)
}
font_rsi_S_bg <- function(..., collapse = " ") {
  try_colour(..., before = "\033[48;5;76m", after = "\033[49m", collapse = collapse)
}
font_rsi_I_bg <- function(..., collapse = " ") {
  try_colour(..., before = "\033[48;5;148m", after = "\033[49m", collapse = collapse)
}
font_red_bg <- function(..., collapse = " ") {
  try_colour(..., before = "\033[41m", after = "\033[49m", collapse = collapse)
}
font_yellow_bg <- function(..., collapse = " ") {
  try_colour(..., before = "\033[43m", after = "\033[49m", collapse = collapse)
}
font_na <- function(..., collapse = " ") {
  font_red(..., collapse = collapse)
}
font_bold <- function(..., collapse = " ") {
  try_colour(..., before = "\033[1m", after = "\033[22m", collapse = collapse)
}
font_italic <- function(..., collapse = " ") {
  try_colour(..., before = "\033[3m", after = "\033[23m", collapse = collapse)
}
font_underline <- function(..., collapse = " ") {
  try_colour(..., before = "\033[4m", after = "\033[24m", collapse = collapse)
}
font_stripstyle <- function(x) {
  # from crayon:::ansi_regex
  gsub("(?:(?:\\x{001b}\\[)|\\x{009b})(?:(?:[0-9]{1,3})?(?:(?:;[0-9]{0,3})*)?[A-M|f-m])|\\x{001b}[A-M]", "", x, perl = TRUE)
}

progress_ticker <- function(n = 1, n_min = 0, ...) {
  if (!interactive() || n < n_min) {
    pb <- list()
    pb$tick <- function() {
      invisible()
    }
    pb$kill <- function() {
      invisible()
    }
    set_clean_class(pb, new_class = "txtProgressBar")
  } else if (n >= n_min) {
    pb <- utils::txtProgressBar(max = n, style = 3)
    pb$tick <- function() {
      pb$up(pb$getVal() + 1)
    }
    pb
  }
}

set_clean_class <- function(x, new_class) {
  # return the object with only the new class and no additional attributes where possible
  if (is.null(x)) {
    x <- NA_character_
  }
  if (is.factor(x)) {
    # keep only levels and remove all other attributes
    lvls <- levels(x)
    attributes(x) <- NULL
    levels(x) <- lvls
  } else if (!is.list(x) && !is.function(x)) {
    attributes(x) <- NULL
  }
  class(x) <- new_class
  x
}

formatted_filesize <- function(...) {
  size_kb <- file.size(...) / 1024
  if (size_kb < 1) {
    paste(round(size_kb, 1), "kB")
  } else if (size_kb < 100) {
    paste(round(size_kb, 0), "kB")
  } else {
    paste(round(size_kb / 1024, 1), "MB")
  }
}

create_pillar_column <- function(x, ...) {
  new_pillar_shaft_simple <- import_fn("new_pillar_shaft_simple", "pillar", error_on_fail = FALSE)
  if (!is.null(new_pillar_shaft_simple)) {
    new_pillar_shaft_simple(x, ...)
  } else {
    # does not exist in package 'pillar' anymore
    structure(list(x),
              class = "pillar_shaft_simple",
              ...)
  }
}

# copied from vctrs::s3_register by their permission:
# https://github.com/r-lib/vctrs/blob/05968ce8e669f73213e3e894b5f4424af4f46316/R/register-s3.R
s3_register <- function(generic, class, method = NULL) {
  stopifnot(is.character(generic), length(generic) == 1)
  stopifnot(is.character(class), length(class) == 1)
  pieces <- strsplit(generic, "::")[[1]]
  stopifnot(length(pieces) == 2)
  package <- pieces[[1]]
  generic <- pieces[[2]]
  caller <- parent.frame()
  get_method_env <- function() {
    top <- topenv(caller)
    if (isNamespace(top)) {
      asNamespace(environmentName(top))
    }
    else {
      caller
    }
  }
  get_method <- function(method, env) {
    if (is.null(method)) {
      get(paste0(generic, ".", class), envir = get_method_env())
    }
    else {
      method
    }
  }
  method_fn <- get_method(method)
  stopifnot(is.function(method_fn))
  setHook(packageEvent(package, "onLoad"), function(...) {
    ns <- asNamespace(package)
    method_fn <- get_method(method)
    registerS3method(generic, class, method_fn, envir = ns)
  })
  if (!isNamespaceLoaded(package)) {
    return(invisible())
  }
  envir <- asNamespace(package)
  if (exists(generic, envir)) {
    registerS3method(generic, class, method_fn, envir = envir)
  }
  invisible()
}

# works exactly like round(), but rounds `round2(44.55, 1)` to 44.6 instead of 44.5
# and adds decimal zeroes until `digits` is reached when force_zero = TRUE
round2 <- function(x, digits = 0, force_zero = TRUE) {
  x <- as.double(x)
  # https://stackoverflow.com/a/12688836/4575331
  val <- (trunc((abs(x) * 10 ^ digits) + 0.5) / 10 ^ digits) * sign(x)
  if (digits > 0 & force_zero == TRUE) {
    values_trans <- val[val != as.integer(val) & !is.na(val)]
    val[val != as.integer(val) & !is.na(val)] <- paste0(values_trans,
                                                        strrep("0",
                                                               max(0,
                                                                   digits - nchar(
                                                                     format(
                                                                       as.double(
                                                                         gsub(".*[.](.*)$",
                                                                              "\\1",
                                                                              values_trans)),
                                                                       scientific = FALSE)))))
  }
  as.double(val)
}


# percentage from our other package: 'cleaner'
percentage <- function(x, digits = NULL, ...) {

  # getdecimalplaces() function
  getdecimalplaces <- function(x, minimum = 0, maximum = 3) {
    if (maximum < minimum) {
      maximum <- minimum
    }
    if (minimum > maximum) {
      minimum <- maximum
    }
    max_places <- max(unlist(lapply(strsplit(sub("0+$", "",
                                                 as.character(x * 100)), ".", fixed = TRUE),
                                    function(y) ifelse(length(y) == 2, nchar(y[2]), 0))), na.rm = TRUE)
    max(min(max_places,
            maximum, na.rm = TRUE),
        minimum, na.rm = TRUE)
  }

  # format_percentage() function
  format_percentage <- function(x, digits = NULL, ...) {
    if (is.null(digits)) {
      digits <- getdecimalplaces(x)
    }

    # round right: percentage(0.4455) and format(as.percentage(0.4455), 1) should return "44.6%", not "44.5%"
    x_formatted <- format(round2(as.double(x), digits = digits + 2) * 100,
                          scientific = FALSE,
                          digits = digits,
                          nsmall = digits,
                          ...)
    x_formatted <- paste0(x_formatted, "%")
    x_formatted[!grepl(pattern = "^[0-9.,e-]+$", x = x)] <- NA_character_
    x_formatted
  }

  # the actual working part
  x <- as.double(x)
  if (is.null(digits)) {
    # max one digit if undefined
    digits <- getdecimalplaces(x, minimum = 0, maximum = 1)
  }
  format_percentage(structure(.Data = as.double(x),
                              class = c("percentage", "numeric")),
                    digits = digits, ...)
}

# prevent dependency on package 'backports'
# these functions were not available in previous versions of R (last checked: R 4.0.3)
# see here for the full list: https://github.com/r-lib/backports
strrep <- function(x, times) {
  x <- as.character(x)
  if (length(x) == 0L)
    return(x)
  unlist(.mapply(function(x, times) {
    if (is.na(x) || is.na(times))
      return(NA_character_)
    if (times <= 0L)
      return("")
    paste0(replicate(times, x), collapse = "")
  }, list(x = x, times = times), MoreArgs = list()), use.names = FALSE)
}
trimws <- function(x, which = c("both", "left", "right")) {
  which <- match.arg(which)
  mysub <- function(re, x) sub(re, "", x, perl = TRUE)
  if (which == "left")
    return(mysub("^[ \t\r\n]+", x))
  if (which == "right")
    return(mysub("[ \t\r\n]+$", x))
  mysub("[ \t\r\n]+$", mysub("^[ \t\r\n]+", x))
}
isFALSE <- function(x) {
  is.logical(x) && length(x) == 1L && !is.na(x) && !x
}
deparse1 <- function(expr, collapse = " ", width.cutoff = 500L, ...) {
  paste(deparse(expr, width.cutoff, ...), collapse = collapse)
}
file.size <- function(...) {
  file.info(...)$size
}
file.mtime <- function(...) {
  file.info(...)$mtime
}
str2lang <- function(s) {
  stopifnot(length(s) == 1L)
  ex <- parse(text = s, keep.source = FALSE)
  stopifnot(length(ex) == 1L)
  ex[[1L]]
}
isNamespaceLoaded <- function(pkg) {
  pkg %in% loadedNamespaces()
}
lengths <- function(x, use.names = TRUE) {
  vapply(x, length, FUN.VALUE = NA_integer_, USE.NAMES = use.names)
}
```
