## Supplementary material for "AMR - An R Package for Working with Antimicrobial Resistance Data": Source package: aa_helper_pm_functions.R

```
# ==================================================================== #
# TITLE                                                                #
# Antimicrobial Resistance (AMR) Analysis for R                        #
#                                                                      #
# SOURCE                                                               #
# https://github.com/msberends/AMR                                     #
#                                                                      #
# LICENCE                                                              #
# (c) 2018-2021 Berends MS, Luz CF et al.                              #
# Developed at the University of Groningen, the Netherlands, in        #
# collaboration with non-profit organisations Certe Medical            #
# Diagnostics & Advice, and University Medical Center Groningen.       # 
#                                                                      #
# This R package is free software; you can freely use and distribute   #
# it for both personal and commercial purposes under the terms of the  #
# GNU General Public License version 2.0 (GNU GPL-2), as published by  #
# the Free Software Foundation.                                        #
# We created this package for both routine data analysis and academic  #
# research and it was publicly released in the hope that it will be    #
# useful, but it comes WITHOUT ANY WARRANTY OR LIABILITY.              #
#                                                                      #
# Visit our website for the full manual and a complete tutorial about  #
# how to conduct AMR analysis: https://msberends.github.io/AMR/        #
# ==================================================================== #

# ------------------------------------------------
# THIS FILE WAS CREATED AUTOMATICALLY!
# Source file: data-raw/reproduction_of_poorman.R
# ------------------------------------------------

# poorman: a package to replace all dplyr functions with base R so we can lose dependency on dplyr.
# These functions were downloaded from https://github.com/nathaneastwood/poorman,
# from this commit: https://github.com/nathaneastwood/poorman/tree/52eb6947e0b4430cd588976ed8820013eddf955f.
#
# All functions are prefixed with 'pm_' to make it obvious that they are dplyr substitutes.
#
# All code below was released under MIT license, that permits 'free of charge, to any person obtaining a 
# copy of the software and associated documentation files (the "Software"), to deal in the Software
# without restriction, including without limitation the rights to use, copy, modify, merge, publish,
# distribute, sublicense, and/or sell copies of the Software, and to permit persons to whom the Software 
# is furnished to do so', given that a copyright notice is given in the software.
#
# Copyright notice on 19 September 2020, the day this code was downloaded, as found on
# https://github.com/nathaneastwood/poorman/blob/52eb6947e0b4430cd588976ed8820013eddf955f/LICENSE:
# YEAR: 2020
# COPYRIGHT HOLDER: Nathan Eastwood

pm_arrange <- function(.data, ...) {
  pm_check_is_dataframe(.data)
  if ("grouped_data" %in% class(.data)) {
    pm_arrange.grouped_data(.data, ...)
  } else {
    pm_arrange.default(.data, ...)
  }
}

pm_arrange.default <- function(.data, ...) {
  pm_context$setup(.data)
  on.exit(pm_context$clean(), add = TRUE)
  rows <- eval(substitute(order(...)), envir = pm_context$.data)
  .data[rows, , drop = FALSE]
}

pm_arrange.grouped_data <- function(.data, ...) {
  pm_apply_grouped_function("pm_arrange", .data, drop = TRUE, ...)
}
pm_between <- function(x, left, right) {
  if (!is.null(attr(x, "class")) && !inherits(x, c("Date", "POSIXct"))) {
    warning("`pm_between()` called on numeric vector with S3 class")
  }
  if (!is.double(x)) x <- as.numeric(x)
  x >= as.numeric(left) & x <= as.numeric(right)
}
pm_context <- new.env()

# Data
pm_context$setup <- function(.data) pm_context$.data <- .data
pm_context$get_data <- function() pm_context$.data
pm_context$get_nrow <- function() nrow(pm_context$.data)
pm_context$get_colnames <- function() colnames(pm_context$.data)
pm_context$clean <- function() rm(list = c(".data"), envir = pm_context)


pm_n <- function() {
  pm_check_group_pm_context("`pm_n()`")
  pm_context$get_nrow()
}

pm_cur_data <- function() {
  pm_check_group_pm_context("`pm_cur_data()`")
  data <- pm_context$get_data()
  data[, !(colnames(data) %in% pm_get_groups(data)), drop = FALSE]
}

pm_cur_group <- function() {
  pm_check_group_pm_context("`pm_cur_group()`")
  data <- pm_context$get_data()
  res <- data[1L, pm_get_groups(data), drop = FALSE]
  rownames(res) <- NULL
  res
}

pm_cur_group_id <- function() {
  pm_check_group_pm_context("`pm_cur_group_id()`")
  data <- pm_context$get_data()
  res <- data[1L, pm_get_groups(data), drop = FALSE]
  details <- pm_get_group_details(data)
  details[, ".group_id"] <- seq_len(nrow(details))
  res <- suppressMessages(pm_semi_join(details, res))
  list(res[, ".group_id"])
}

pm_cur_group_rows <- function() {
  pm_check_group_pm_context("`pm_cur_group_rows()`")
  data <- pm_context$get_data()
  res <- data[1L, pm_get_groups(data), drop = FALSE]
  res <- suppressMessages(pm_semi_join(pm_get_group_details(data), res))
  unlist(res[, ".rows"])
}

pm_check_group_pm_context <- function(fn) {
  if (is.null(pm_context$.data)) {
    stop(fn, " must only be used inside poorman verbs")
  }
}
pm_count <- function(x, ..., wt = NULL, sort = FALSE, name = NULL) {
  pm_groups <- pm_get_groups(x)
  if (!missing(...)) x <- pm_group_by(x, ..., .add = TRUE)
  wt <- pm_deparse_var(wt)
  res <- do.call(pm_tally, list(x, wt, sort, name))
  if (length(pm_groups) > 0L) res <- do.call(pm_group_by, list(res, as.name(pm_groups)))
  res
}

pm_tally <- function(x, wt = NULL, sort = FALSE, name = NULL) {
  name <- pm_check_name(x, name)
  wt <- pm_deparse_var(wt)
  res <- do.call(pm_summarise, pm_set_names(list(x, pm_tally_n(x, wt)), c(".data", name)))
  res <- pm_ungroup(res)
  if (isTRUE(sort)) res <- do.call(pm_arrange, list(res, call("pm_desc", as.name(name))))
  rownames(res) <- NULL
  res
}

pm_add_count <- function(x, ..., wt = NULL, sort = FALSE, name = NULL) {
  name <- pm_check_name(x, name)
  row_names <- rownames(x)
  wt <- pm_deparse_var(wt)
  if (!missing(...)) x <- pm_group_by(x, ..., .add = TRUE)
  res <- do.call(pm_add_tally, list(x, wt, sort, name))
  res[row_names, ]
}

pm_add_tally <- function(x, wt = NULL, sort = FALSE, name = NULL) {
  wt <- pm_deparse_var(wt)
  pm_n <- pm_tally_n(x, wt)
  name <- pm_check_name(x, name)
  res <- do.call(pm_mutate, pm_set_names(list(x, pm_n), c(".data", name)))

  if (isTRUE(sort)) {
    do.call(pm_arrange, list(res, call("pm_desc", as.name(name))))
  } else {
    res
  }
}

pm_tally_n <- function(x, wt) {
  if (is.null(wt) && "pm_n" %in% colnames(x)) {
    message("Using `pm_n` as weighting variable")
    wt <- "pm_n"
  }
  pm_context$setup(.data = x)
  on.exit(pm_context$clean(), add = TRUE)
  if (is.null(wt)) {
    call("pm_n")
  } else {
    call("sum", as.name(wt), na.rm = TRUE)
  }
}

pm_check_name <- function(df, name) {
  if (is.null(name)) {
    if ("pm_n" %in% colnames(df)) {
      stop(
        "Column 'pm_n' is already present in output\n",
        "* Use `name = \"new_name\"` to pick a new name"
      )
    }
    return("pm_n")
  }

  if (!is.character(name) || length(name) != 1) {
    stop("`name` must be a single string")
  }

  name
}
pm_desc <- function(x) -xtfrm(x)
pm_distinct <- function(.data, ...) {
  pm_check_is_dataframe(.data)
  if ("grouped_data" %in% class(.data)) {
    pm_distinct.grouped_data(.data, ...)
  } else {
    pm_distinct.default(.data, ...)
  }
}

pm_distinct.default <- function(.data, ..., .keep_all = FALSE) {
  if (ncol(.data) == 0L) return(.data[1, ])
  cols <- pm_deparse_dots(...)
  col_names <- names(cols)
  col_len <- length(cols)
  if (is.null(col_names) && col_len > 0L) names(cols) <- cols
  if (col_len == 0L) {
    res <- .data
  } else {
    res <- pm_mutate(.data, ...)
    col_names <- names(cols)
    res <- if (!is.null(col_names)) {
      zero_names <- nchar(col_names) == 0L
      if (any(zero_names)) {
        names(cols)[zero_names] <- cols[zero_names]
        col_names <- names(cols)
      }
      suppressMessages(pm_select(res, col_names))
    } else {
      suppressMessages(pm_select(res, cols))
    }
  }
  res <- unique(res)
  if (isTRUE(.keep_all)) {
    res <- cbind(res, .data[rownames(res), setdiff(colnames(.data), colnames(res)), drop = FALSE])
  }
  common_cols <- c(intersect(colnames(.data), colnames(res)), setdiff(col_names, colnames(.data)))
  if (length(common_cols) > 0L) res[, common_cols, drop = FALSE] else res
}

pm_distinct.grouped_data <- function(.data, ..., .keep_all = FALSE) {
  pm_apply_grouped_function("pm_distinct", .data, drop = TRUE, ..., .keep_all = .keep_all)
}
pm_eval_env <- new.env()
pm_filter <- function(.data, ...) {
  pm_check_is_dataframe(.data)
  if ("grouped_data" %in% class(.data)) {
    pm_filter.grouped_data(.data, ...)
  } else {
    pm_filter.default(.data, ...)
  }
}

pm_filter.default <- function(.data, ...) {
  conditions <- pm_dotdotdot(...)
  cond_class <- vapply(conditions, typeof, NA_character_)
  if (any(cond_class != "language")) stop("Conditions must be logical vectors")
  pm_context$setup(.data)
  on.exit(pm_context$clean(), add = TRUE)
  pm_eval_env$env <- parent.frame()
  on.exit(rm(list = "env", envir = pm_eval_env), add = TRUE)
  rows <- lapply(
    conditions,
    function(cond, frame) eval(cond, pm_context$.data, frame),
    frame = pm_eval_env$env
  )
  rows <- Reduce("&", rows)
  .data[rows & !is.na(rows), ]
}

pm_filter.grouped_data <- function(.data, ...) {
  rows <- rownames(.data)
  res <- pm_apply_grouped_function("pm_filter", .data, drop = TRUE, ...)
  res[rows[rows %in% rownames(res)], ]
}
pm_group_by <- function(.data, ..., .add = FALSE) {
  pm_check_is_dataframe(.data)
  pre_groups <- pm_get_groups(.data)
  pm_groups <- pm_deparse_dots(...)
  if (isTRUE(.add)) pm_groups <- unique(c(pre_groups, pm_groups))
  unknown <- !(pm_groups %in% colnames(.data))
  if (any(unknown)) stop("Invalid pm_groups: ", pm_groups[unknown])
  class(.data) <- c("grouped_data", class(.data))
  pm_set_groups(.data, pm_groups)
}

pm_ungroup <- function(x, ...) {
  pm_check_is_dataframe(x)
  rm_groups <- pm_deparse_dots(...)
  pm_groups <- pm_get_groups(x)
  if (length(rm_groups) == 0L) rm_groups <- pm_groups
  x <- pm_set_groups(x, pm_groups[!(pm_groups %in% rm_groups)])
  if (length(attr(x, "pm_groups")) == 0L) {
    attr(x, "pm_groups") <- NULL
    class(x) <- class(x)[!(class(x) %in% "grouped_data")]
  }
  x
}

pm_set_groups <- function(x, pm_groups) {
  attr(x, "pm_groups") <- if (is.null(pm_groups) || length(pm_groups) == 0L) {
    NULL
  } else {
    pm_group_data_worker(x, pm_groups)
  }
  x
}

pm_get_groups <- function(x) {
  pm_groups <- attr(x, "pm_groups", exact = TRUE)
  if (is.null(pm_groups)) character(0) else colnames(pm_groups)[!colnames(pm_groups) %in% c(".group_id", ".rows")]
}

pm_get_group_details <- function(x) {
  pm_groups <- attr(x, "pm_groups", exact = TRUE)
  if (is.null(pm_groups)) character(0) else pm_groups
}

pm_has_groups <- function(x) {
  pm_groups <- pm_get_groups(x)
  if (length(pm_groups) == 0L) FALSE else TRUE
}

pm_apply_grouped_function <- function(fn, .data, drop = FALSE, ...) {
  pm_groups <- pm_get_groups(.data)
  grouped <- pm_split_into_groups(.data, pm_groups, drop)
  res <- do.call(rbind, unname(lapply(grouped, fn, ...)))
  if (any(pm_groups %in% colnames(res))) {
    class(res) <- c("grouped_data", class(res))
    res <- pm_set_groups(res, pm_groups[pm_groups %in% colnames(res)])
  }
  res
}

pm_print.grouped_data <- function(x, ..., digits = NULL, quote = FALSE, right = TRUE, row.names = TRUE, max = NULL) {
  class(x) <- "data.frame"
  print(x, ..., digits = digits, quote = quote, right = right, row.names = row.names, max = max)
  cat("\nGroups: ", paste(pm_get_groups(x), collapse = ", "), "\n\n")
}

pm_group_data <- function(.data) {
  if (!pm_has_groups(.data)) return(data.frame(.rows = I(list(seq_len(nrow(.data))))))
  pm_groups <- pm_get_groups(.data)
  pm_group_data_worker(.data, pm_groups)
}

pm_group_data_worker <- function(.data, pm_groups) {
  res <- unique(.data[, pm_groups, drop = FALSE])
  class(res) <- "data.frame"
  nrow_res <- nrow(res)
  rows <- rep(list(NA), nrow_res)
  for (i in seq_len(nrow_res)) {
    rows[[i]] <- which(interaction(.data[, pm_groups]) %in% interaction(res[i, pm_groups]))
  }
  res$`.rows` <- rows
  res <- res[do.call(order, lapply(pm_groups, function(x) res[, x])), , drop = FALSE]
  rownames(res) <- NULL
  res
}

pm_group_rows <- function(.data) {
  pm_group_data(.data)[[".rows"]]
}

pm_group_indices <- function(.data) {
  if (!pm_has_groups(.data)) return(rep(1L, nrow(.data)))
  pm_groups <- pm_get_groups(.data)
  res <- unique(.data[, pm_groups, drop = FALSE])
  res <- res[do.call(order, lapply(pm_groups, function(x) res[, x])), , drop = FALSE]
  class(res) <- "data.frame"
  nrow_data <- nrow(.data)
  rows <- rep(NA, nrow_data)
  for (i in seq_len(nrow_data)) {
    rows[i] <- which(interaction(res[, pm_groups]) %in% interaction(.data[i, pm_groups]))
  }
  rows
}

pm_group_vars <- function(x) {
  pm_get_groups(x)
}

pm_groups <- function(x) {
  lapply(pm_get_groups(x), as.symbol)
}

pm_group_size <- function(x) {
  lengths(pm_group_rows(x))
}

pm_n_groups <- function(x) {
  nrow(pm_group_data(x))
}
# pm_group_split <- function(.data, ..., .keep = TRUE) {
#   dots_len <- ...length() > 0L
#   if (pm_has_groups(.data) && isTRUE(dots_len)) {
#     warning("... is ignored in pm_group_split(<grouped_df>), please use pm_group_by(..., .add = TRUE) %pm>% pm_group_split()")
#   }
#   if (!pm_has_groups(.data) && isTRUE(dots_len)) {
#     .data <- pm_group_by(.data, ...)
#   }
#   if (!pm_has_groups(.data) && isFALSE(dots_len)) {
#     return(list(.data))
#   }
#   pm_context$setup(.data)
#   on.exit(pm_context$clean(), add = TRUE)
#   pm_groups <- pm_get_groups(.data)
#   attr(pm_context$.data, "pm_groups") <- NULL
#   res <- pm_split_into_groups(pm_context$.data, pm_groups)
#   names(res) <- NULL
#   if (isFALSE(.keep)) {
#     res <- lapply(res, function(x) x[, !colnames(x) %in% pm_groups])
#   }
#   any_empty <- unlist(lapply(res, function(x) !(nrow(x) == 0L)))
#   res[any_empty]
# }

pm_group_keys <- function(.data) {
  pm_groups <- pm_get_groups(.data)
  pm_context$setup(.data)
  res <- pm_context$.data[, pm_context$get_colnames() %in% pm_groups, drop = FALSE]
  res <- res[!duplicated(res), , drop = FALSE]
  if (nrow(res) == 0L) return(res)
  class(res) <- "data.frame"
  res <- res[do.call(order, lapply(pm_groups, function(x) res[, x])), , drop = FALSE]
  rownames(res) <- NULL
  res
}

pm_split_into_groups <- function(.data, pm_groups, drop = FALSE, ...) {
  class(.data) <- "data.frame"
  group_factors <- lapply(pm_groups, function(x, .data) as.factor(.data[, x]), .data)
  split(x = .data, f = group_factors, drop = drop, ...)
}
pm_if_else <- function(condition, true, false, missing = NULL) {
  if (!is.logical(condition)) stop("`condition` must be a logical vector.")
  cls_true <- class(true)
  cls_false <- class(false)
  cls_missing <- class(missing)
  if (!identical(cls_true, cls_false)) {
    stop("The class of `true` <", class(true), "> is not the same as the class of `false` <", class(false), ">")
  }
  if (!is.null(missing) && !identical(cls_true, cls_missing)) {
    stop("`missing` must be a ", cls_true, " vector, not a ", cls_missing, " vector.")
  }
  res <- ifelse(condition, true, false)
  if (!is.null(missing)) res[is.na(res)] <- missing
  attributes(res) <- attributes(true)
  res
}

pm_anti_join <- function(x, y, by = NULL) {
  pm_filter_join_worker(x, y, by, type = "anti")
}

pm_semi_join <- function(x, y, by = NULL) {
  pm_filter_join_worker(x, y, by, type = "semi")
}

pm_filter_join_worker <- function(x, y, by = NULL, type = c("anti", "semi")) {
  type <- match.arg(type, choices = c("anti", "semi"), several.ok = FALSE)
  if (is.null(by)) {
    by <- intersect(names(x), names(y))
    pm_join_message(by)
  }
  rows <- interaction(x[, by]) %in% interaction(y[, by])
  if (type == "anti") rows <- !rows
  res <- x[rows, , drop = FALSE]
  rownames(res) <- NULL
  res
}

pm_inner_join <- function(x, y, by = NULL, suffix = c(".x", ".y")) {
  pm_join_worker(x = x, y = y, by = by, suffix = suffix, sort = FALSE)
}

# pm_left_join <- function(x, y, by = NULL, suffix = c(".x", ".y")) {
#   pm_join_worker(x = x, y = y, by = by, suffix = suffix, all.x = TRUE)
# }

pm_right_join <- function(x, y, by = NULL, suffix = c(".x", ".y")) {
  pm_join_worker(x = x, y = y, by = by, suffix = suffix, all.y = TRUE)
}

pm_full_join <- function(x, y, by = NULL, suffix = c(".x", ".y")) {
  pm_join_worker(x = x, y = y, by = by, suffix = suffix, all = TRUE)
}

pm_join_worker <- function(x, y, by = NULL, suffix = c(".x", ".y"), ...) {
  x[, ".join_id"] <- seq_len(nrow(x))
  if (is.null(by)) {
    by <- intersect(names(x), names(y))
    pm_join_message(by)
    merged <- merge(x = x, y = y, by = by, suffixes = suffix, ...)[, union(names(x), names(y))]
  } else if (is.null(names(by))) {
    merged <- merge(x = x, y = y, by = by, suffixes = suffix, ...)
  } else {
    merged <- merge(x = x, y = y, by.x = names(by), by.y = by, suffixes = suffix, ...)
  }
  merged <- merged[order(merged[, ".join_id"]), colnames(merged) != ".join_id"]
  rownames(merged) <- NULL
  merged
}

pm_join_message <- function(by) {
  if (length(by) > 1L) {
    message("Joining, by = c(\"", paste0(by, collapse = "\", \""), "\")\n", sep = "")
  } else {
    message("Joining, by = \"", by, "\"\n", sep = "")
  }
}
pm_lag <- function(x, pm_n = 1L, default = NA) {
  if (inherits(x, "ts")) stop("`x` must be a vector, not a `ts` object, do you want `stats::pm_lag()`?")
  if (length(pm_n) != 1L || !is.numeric(pm_n) || pm_n < 0L) stop("`pm_n` must be a nonnegative integer scalar")
  if (pm_n == 0L) return(x)
  tryCatch(
    storage.mode(default) <- typeof(x),
    warning = function(w) {
      stop("Cannot convert `default` <", typeof(default), "> to `x` <", typeof(x), ">")
    }
  )
  xlen <- length(x)
  pm_n <- pmin(pm_n, xlen)
  res <- c(rep(default, pm_n), x[seq_len(xlen - pm_n)])
  attributes(res) <- attributes(x)
  res
}

pm_lead <- function(x, pm_n = 1L, default = NA) {
  if (length(pm_n) != 1L || !is.numeric(pm_n) || pm_n < 0L) stop("pm_n must be a nonnegative integer scalar")
  if (pm_n == 0L) return(x)
  tryCatch(
    storage.mode(default) <- typeof(x),
    warning = function(w) {
      stop("Cannot convert `default` <", typeof(default), "> to `x` <", typeof(x), ">")
    }
  )
  xlen <- length(x)
  pm_n <- pmin(pm_n, xlen)
  res <- c(x[-seq_len(pm_n)], rep(default, pm_n))
  attributes(res) <- attributes(x)
  res
}
pm_mutate <- function(.data, ...) {
  pm_check_is_dataframe(.data)
  if ("grouped_data" %in% class(.data)) {
    pm_mutate.grouped_data(.data, ...)
  } else {
    pm_mutate.default(.data, ...)
  }
}

pm_mutate.default <- function(.data, ...) {
  conditions <- pm_dotdotdot(..., .impute_names = TRUE)
  .data[, setdiff(names(conditions), names(.data))] <- NA
  pm_context$setup(.data)
  on.exit(pm_context$clean(), add = TRUE)
  for (i in seq_along(conditions)) {
    pm_context$.data[, names(conditions)[i]] <- eval(conditions[[i]], envir = pm_context$.data)
  }
  pm_context$.data
}

pm_mutate.grouped_data <- function(.data, ...) {
  rows <- rownames(.data)
  res <- pm_apply_grouped_function("pm_mutate", .data, drop = TRUE, ...)
  res[rows, ]
}
pm_n_distinct <- function(..., na.rm = FALSE) {
  res <- c(...)
  if (is.list(res)) return(nrow(unique(as.data.frame(res, stringsAsFactors = FALSE))))
  if (isTRUE(na.rm)) res <- res[!is.na(res)]
  length(unique(res))
}
pm_na_if <- function(x, y) {
  y_len <- length(y)
  x_len <- length(x)
  if (!(y_len %in% c(1L, x_len))) stop("`y` must be length ", x_len, " (same as `x`) or 1, not ", y_len)
  x[x == y] <- NA
  x
}
pm_near <- function(x, y, tol = .Machine$double.eps^0.5) {
  abs(x - y) < tol
}
`%pm>%` <- function(lhs, rhs) {
  lhs <- substitute(lhs)
  rhs <- substitute(rhs)
  eval(as.call(c(rhs[[1L]], lhs, as.list(rhs[-1L]))), envir = parent.frame())
}
pm_pull <- function(.data, var = -1) {
  var_deparse <- pm_deparse_var(var)
  col_names <- colnames(.data)
  if (!(var_deparse %in% col_names) & grepl("^[[:digit:]]+L|[[:digit:]]", var_deparse)) {
    var <- as.integer(gsub("L", "", var_deparse))
    var <- pm_if_else(var < 1L, rev(col_names)[abs(var)], col_names[var])
  } else if (var_deparse %in% col_names) {
    var <- var_deparse
  }
  .data[, var]
}
pm_set_names <- function(object = nm, nm) {
  names(object) <- nm
  object
}

pm_vec_head <- function(x, pm_n = 6L, ...) {
  stopifnot(length(pm_n) == 1L)
  pm_n <- if (pm_n < 0L) max(length(x) + pm_n, 0L) else min(pm_n, length(x))
  x[seq_len(pm_n)]
}
pm_relocate <- function(.data, ..., .before = NULL, .after = NULL) {
  pm_check_is_dataframe(.data)
  data_names <- colnames(.data)
  col_pos <- pm_select_positions(.data, ...)

  .before <- pm_deparse_var(.before)
  .after <- pm_deparse_var(.after)
  has_before <- !is.null(.before)
  has_after <- !is.null(.after)

  if (has_before && has_after) {
    stop("You must supply only one of `.before` and `.after`")
  } else if (has_before) {
    pm_where <- min(match(.before, data_names))
    col_pos <- c(setdiff(col_pos, pm_where), pm_where)
  } else if (has_after) {
    pm_where <- max(match(.after, data_names))
    col_pos <- c(pm_where, setdiff(col_pos, pm_where))
  } else {
    pm_where <- 1L
    col_pos <- union(col_pos, pm_where)
  }
  lhs <- setdiff(seq(1L, pm_where - 1L), col_pos)
  rhs <- setdiff(seq(pm_where + 1L, ncol(.data)), col_pos)
  col_pos <- unique(c(lhs, col_pos, rhs))
  col_pos <- col_pos[col_pos <= length(data_names)]

  res <- .data[col_pos]
  if (pm_has_groups(.data)) res <- pm_set_groups(res, pm_get_groups(.data))
  res
}
pm_rename <- function(.data, ...) {
  pm_check_is_dataframe(.data)
  new_names <- names(pm_deparse_dots(...))
  if (length(new_names) == 0L) {
    warning("You didn't give any new names")
    return(.data)
  }
  col_pos <- pm_select_positions(.data, ...)
  old_names <- colnames(.data)[col_pos]
  new_names_zero <- nchar(new_names) == 0L
  if (any(new_names_zero)) {
    warning("You didn't provide new names for: ", paste0("`", old_names[new_names_zero], collapse = ", "), "`")
    new_names[new_names_zero] <- old_names[new_names_zero]
  }
  colnames(.data)[col_pos] <- new_names
  .data
}

pm_rename_with <- function(.data, .fn, .cols = pm_everything(), ...) {
  if (!is.function(.fn)) stop("`", .fn, "` is not a valid function")
  grouped <- inherits(.data, "grouped_data")
  if (grouped) grp_pos <- which(colnames(.data) %in% pm_group_vars(.data))
  col_pos <- eval(substitute(pm_select_positions(.data, .cols)))
  cols <- colnames(.data)[col_pos]
  new_cols <- .fn(cols, ...)
  if (any(duplicated(new_cols))) {
    stop("New names must be unique however `", deparse(substitute(.fn)), "` returns duplicate column names")
  }
  colnames(.data)[col_pos] <- new_cols
  if (grouped) .data <- pm_set_groups(.data, colnames(.data)[grp_pos])
  .data
}
pm_replace_with <- function(x, i, val, arg_name) {
  if (is.null(val)) return(x)
  pm_check_length(val, x, arg_name)
  pm_check_type(val, x, arg_name)
  pm_check_class(val, x, arg_name)
  i[is.na(i)] <- FALSE
  if (length(val) == 1L) {
    x[i] <- val
  }
  else {
    x[i] <- val[i]
  }
  x
}

pm_check_length <- function(x, y, arg_name) {
  length_x <- length(x)
  length_y <- length(y)
  if (all(length_x %in% c(1L, length_y))) return()
  if (length_y == 1) {
    stop(arg_name, " must be length 1, not ", paste(length_x, sep = ", "))
  } else {
    stop(arg_name, " must be length ", length_y, " or 1, not ", length_x)
  }
}

pm_check_type <- function(x, y, arg_name) {
  x_type <- typeof(x)
  y_type <- typeof(y)
  if (identical(x_type, y_type)) return()
  stop(arg_name, " must be `", y_type, "`, not `", x_type, "`")
}

pm_check_class <- function(x, y, arg_name) {
  if (!is.object(x)) return()
  exp_classes <- class(y)
  out_classes <- class(x)
  if (identical(out_classes, exp_classes)) return()
  stop(arg_name, " must have class `", exp_classes, "`, not class `", out_classes, "`")
}
pm_rownames_to_column <- function(.data, var = "rowname") {
  pm_check_is_dataframe(.data)
  col_names <- colnames(.data)
  if (var %in% col_names) stop("Column `", var, "` already exists in `.data`")
  .data[, var] <- rownames(.data)
  rownames(.data) <- NULL
  .data[, c(var, setdiff(col_names, var))]
}
pm_starts_with <- function(match, ignore.case = TRUE, vars = pm_peek_vars()) {
  grep(pattern = paste0("^", paste0(match, collapse = "|^")), x = vars, ignore.case = ignore.case)
}

pm_ends_with <- function(match, ignore.case = TRUE, vars = pm_peek_vars()) {
  grep(pattern = paste0(paste0(match, collapse = "$|"), "$"), x = vars, ignore.case = ignore.case)
}

pm_contains <- function(match, ignore.case = TRUE, vars = pm_peek_vars()) {
  pm_matches <- lapply(
    match,
    function(x) {
      if (isTRUE(ignore.case)) {
        match_u <- toupper(x)
        match_l <- tolower(x)
        pos_u <- grep(pattern = match_u, x = toupper(vars), fixed = TRUE)
        pos_l <- grep(pattern = match_l, x = tolower(vars), fixed = TRUE)
        unique(c(pos_l, pos_u))
      } else {
        grep(pattern = x, x = vars, fixed = TRUE)
      }
    }
  )
  unique(unlist(pm_matches))
}

pm_matches <- function(match, ignore.case = TRUE, perl = FALSE, vars = pm_peek_vars()) {
  grep(pattern = match, x = vars, ignore.case = ignore.case, perl = perl)
}

pm_num_range <- function(prefix, range, width = NULL, vars = pm_peek_vars()) {
  if (!is.null(width)) {
    range <- sprintf(paste0("%0", width, "d"), range)
  }
  find <- paste0(prefix, range)
  if (any(duplicated(vars))) {
    stop("Column names must be unique")
  } else {
    x <- match(find, vars)
    x[!is.na(x)]
  }
}

pm_all_of <- function(x, vars = pm_peek_vars()) {
  x_ <- !x %in% vars
  if (any(x_)) {
    which_x_ <- which(x_)
    if (length(which_x_) == 1L) {
      stop("The column ", x[which_x_], " does not exist.")
    } else {
      stop("The columns ", paste(x[which_x_], collapse = ", "), " do not exist.")
    }
  } else {
    which(vars %in% x)
  }
}

pm_any_of <- function(x, vars = pm_peek_vars()) {
  which(vars %in% x)
}

pm_everything <- function(vars = pm_peek_vars()) {
  seq_along(vars)
}

pm_last_col <- function(offset = 0L, vars = pm_peek_vars()) {
  if (!pm_is_wholenumber(offset)) stop("`offset` must be an integer")
  pm_n <- length(vars)
  if (offset && pm_n <= offset) {
    stop("`offset` must be smaller than the number of `vars`")
  } else if (pm_n == 0) {
    stop("Can't pm_select last column when `vars` is empty")
  } else {
    pm_n - offset
  }
}

pm_peek_vars <- function() {
  pm_select_env$get_colnames()
}
pm_select_positions <- function(.data, ..., .group_pos = FALSE) {
  cols <- pm_dotdotdot(...)
  pm_select_env$setup(.data = .data, calling_frame = parent.frame(2L))
  on.exit(pm_select_env$clean(), add = TRUE)
  data_names <- pm_select_env$get_colnames()
  pos <- unlist(lapply(cols, pm_eval_expr))
  col_len <- pm_select_env$get_ncol()
  if (any(pos > col_len)) {
    oor <- pos[which(pos > col_len)]
    oor_len <- length(oor)
    stop(
      "Location", if (oor_len > 1) "s " else " ", pm_collapse_to_sentence(oor),
      if (oor_len > 1) " don't " else " doesn't ", "exist. There are only ", col_len, " columns."
    )
  }
  if (isTRUE(.group_pos)) {
    pm_groups <- pm_get_groups(.data)
    missing_groups <- !(pm_groups %in% cols)
    if (any(missing_groups)) {
      sel_missing <- pm_groups[missing_groups]
      message("Adding missing grouping variables: `", paste(sel_missing, collapse = "`, `"), "`")
      readd <- match(sel_missing, data_names)
      if (length(names(cols)) > 0L) names(readd) <- data_names[readd]
      pos <- c(readd, pos)
    }
  }
  pos[!duplicated(pos)]
}

pm_eval_expr <- function(x) {
  type <- typeof(x)
  switch(
    type,
    "integer" = x,
    "double" = as.integer(x),
    "character" = pm_select_char(x),
    "symbol" = pm_select_symbol(x),
    "language" = pm_eval_call(x),
    stop("Expressions of type <", typeof(x), "> cannot be evaluated for use when subsetting.")
  )
}

pm_select_char <- function(expr) {
  pos <- match(expr, pm_select_env$get_colnames())
  if (is.na(pos)) stop("Column `", expr, "` does not exist")
  pos
}

pm_select_symbol <- function(expr) {
  expr_name <- as.character(expr)
  if (grepl("^is\\.", expr_name) && pm_is_function(expr)) {
    stop(
      "Predicate functions must be wrapped in `pm_where()`.\n\n",
      sprintf("  data %%pm>%% pm_select(pm_where(%s))", expr_name)
    )
  }
  res <- try(pm_select_char(as.character(expr)), silent = TRUE)
  if (inherits(res, "try-error")) {
    res <- tryCatch(
      unlist(lapply(eval(expr, envir = pm_select_env$calling_frame), pm_eval_expr)),
      error = function(e) stop("Column ", expr, " does not exist.")
    )
  }
  res
}

pm_eval_call <- function(x) {
  type <- as.character(x[[1]])
  switch(
    type,
    `:` = pm_select_seq(x),
    `!` = pm_select_negate(x),
    `-` = pm_select_minus(x),
    `c` = pm_select_c(x),
    `(` = pm_select_bracket(x),
    pm_select_pm_context(x)
  )
}

pm_select_seq <- function(expr) {
  x <- pm_eval_expr(expr[[2]])
  y <- pm_eval_expr(expr[[3]])
  x:y
}

pm_select_negate <- function(expr) {
  x <- if (pm_is_negated_colon(expr)) {
    expr <- call(":", expr[[2]][[2]], expr[[2]][[3]][[2]])
    pm_eval_expr(expr)
  } else {
    pm_eval_expr(expr[[2]])
  }
  x * -1L
}

pm_is_negated_colon <- function(expr) {
  expr[[1]] == "!" && length(expr[[2]]) > 1L && expr[[2]][[1]] == ":" && expr[[2]][[3]][[1]] == "!"
}

pm_select_minus <- function(expr) {
  x <- pm_eval_expr(expr[[2]])
  x * -1L
}

pm_select_c <- function(expr) {
  lst_expr <- as.list(expr)
  lst_expr[[1]] <- NULL
  unlist(lapply(lst_expr, pm_eval_expr))
}

pm_select_bracket <- function(expr) {
  pm_eval_expr(expr[[2]])
}

pm_select_pm_context <- function(expr) {
  eval(expr, envir = pm_select_env$.data)
}

pm_select_env <- new.env()
pm_select_env$setup <- function(.data, calling_frame) {
  pm_select_env$.data <- .data
  pm_select_env$calling_frame <- calling_frame
}
pm_select_env$clean <- function() {
  rm(list = c(".data", "calling_frame"), envir = pm_select_env)
}
pm_select_env$get_colnames <- function() colnames(pm_select_env$.data)
pm_select_env$get_nrow <- function() nrow(pm_select_env$.data)
pm_select_env$get_ncol <- function() ncol(pm_select_env$.data)

pm_select <- function(.data, ...) {
  col_pos <- pm_select_positions(.data, ..., .group_pos = TRUE)
  map_names <- names(col_pos)
  map_names_length <- nchar(map_names)
  if (any(map_names_length == 0L)) {
    no_new_names <- which(map_names_length == 0L)
    map_names[no_new_names] <- colnames(.data)[no_new_names]
  }
  res <- .data[, col_pos, drop = FALSE]
  if (!is.null(map_names) && all(col_pos > 0L)) colnames(res) <- map_names
  if (pm_has_groups(.data)) res <- pm_set_groups(res, pm_get_groups(.data))
  res
}
pm_summarise <- function(.data, ...) {
  pm_check_is_dataframe(.data)
  if ("grouped_data" %in% class(.data)) {
    pm_summarise.grouped_data(.data, ...)
  } else {
    pm_summarise.default(.data, ...)
  }
}

pm_summarise.default <- function(.data, ...) {
  fns <- pm_dotdotdot(...)
  pm_context$setup(.data)
  on.exit(pm_context$clean(), add = TRUE)
  pm_groups_exist <- pm_has_groups(pm_context$.data)
  if (pm_groups_exist) {
    group <- unique(pm_context$.data[, pm_get_groups(pm_context$.data), drop = FALSE])
  }
  res <- lapply(
    fns,
    function(x) {
      x_res <- do.call(with, list(pm_context$.data, x))
      if (is.list(x_res)) I(x_res) else x_res
    }
  )
  res <- as.data.frame(res)
  fn_names <- names(fns)
  colnames(res) <- if (is.null(fn_names)) fns else fn_names
  if (pm_groups_exist) res <- cbind(group, res, row.names = NULL)
  res
}

pm_summarise.grouped_data <- function(.data, ...) {
  pm_groups <- pm_get_groups(.data)
  res <- pm_apply_grouped_function("pm_summarise", .data, drop = TRUE, ...)
  res <- res[do.call(order, lapply(pm_groups, function(x) res[, x])), ]
  rownames(res) <- NULL
  res
}

pm_transmute <- function(.data, ...) {
  pm_check_is_dataframe(.data)
  if ("grouped_data" %in% class(.data)) {
    pm_transmute.grouped_data(.data, ...)
  } else {
    pm_transmute.default(.data, ...)
  }
}

pm_transmute.default <- function(.data, ...) {
  conditions <- pm_deparse_dots(...)
  mutated <- pm_mutate(.data, ...)
  mutated[, names(conditions), drop = FALSE]
}

pm_transmute.grouped_data <- function(.data, ...) {
  rows <- rownames(.data)
  res <- pm_apply_grouped_function("pm_transmute", .data, drop = TRUE, ...)
  res[rows, ]
}
pm_dotdotdot <- function(..., .impute_names = FALSE) {
  dots <- eval(substitute(alist(...)))
  if (isTRUE(.impute_names)) {
    pm_deparse_dots <- lapply(dots, deparse)
    names_dots <- names(dots)
    unnamed <- if (is.null(names_dots)) rep(TRUE, length(dots)) else nchar(names_dots) == 0L
    names(dots)[unnamed] <- pm_deparse_dots[unnamed]
  }
  dots
}

pm_deparse_dots <- function(...) {
  vapply(substitute(...()), deparse, NA_character_)
}

pm_deparse_var <- function(var, frame = if (is.null(pm_eval_env$env)) parent.frame() else pm_eval_env$env) {
  sub_var <- eval(substitute(substitute(var)), frame)
  if (is.symbol(sub_var)) var <- as.character(sub_var)
  var
}

pm_check_is_dataframe <- function(.data) {
  parent_fn <- all.names(sys.call(-1L), max.names = 1L)
  if (!is.data.frame(.data)) stop(parent_fn, " must be given a data.frame")
  invisible()
}

pm_is_wholenumber <- function(x) {
  x %% 1L == 0L
}

pm_seq2 <- function (from, to) {
  if (length(from) != 1) stop("`from` must be length one")
  if (length(to) != 1) stop("`to` must be length one")
  if (from > to) integer() else seq.int(from, to)
}

pm_is_function <- function(x, frame) {
  res <- tryCatch(
    is.function(x),
    warning = function(w) FALSE,
    error = function(e) FALSE
  )
  if (isTRUE(res)) return(res)
  res <- tryCatch(
    is.function(eval(x)),
    warning = function(w) FALSE,
    error = function(e) FALSE
  )
  if (isTRUE(res)) return(res)
  res <- tryCatch(
    is.function(eval(as.symbol(deparse(substitute(x))))),
    warning = function(w) FALSE,
    error = function(e) FALSE
  )
  if (isTRUE(res)) return(res)
  FALSE
}

pm_collapse_to_sentence <- function(x) {
  len_x <- length(x)
  if (len_x == 0L) {
    stop("Length of `x` is 0")
  } else if (len_x == 1L) {
    as.character(x)
  } else if (len_x == 2L) {
    paste(x, collapse = " and ")
  } else {
    paste(paste(x[1:(len_x - 1)], collapse = ", "), x[len_x], sep = " and ")
  }
}
pm_where <- function(fn) {
  if (!pm_is_function(fn)) {
    stop(pm_deparse_var(fn), " is not a valid predicate function.")
  }
  preds <- unlist(lapply(
    pm_select_env$.data,
    function(x, fn) {
      do.call("fn", list(x))
    },
    fn
  ))
  if (!is.logical(preds)) stop("`pm_where()` must be used with functions that return `TRUE` or `FALSE`.")
  data_cols <- pm_select_env$get_colnames()
  cols <- data_cols[preds]
  which(data_cols %in% cols)
}

pm_cume_dist <- function(x) {
  rank(x, ties.method = "max", na.last = "keep") / sum(!is.na(x))
}

pm_dense_rank <- function(x) {
  match(x, sort(unique(x)))
}

pm_min_rank <- function(x) {
  rank(x, ties.method = "min", na.last = "keep")
}

pm_ntile <- function(x = pm_row_number(), pm_n) {
  if (!missing(x)) x <- pm_row_number(x)
  len <- length(x) - sum(is.na(x))
  pm_n <- as.integer(floor(pm_n))
  if (len == 0L) {
    rep(NA_integer_, length(x))
  } else {
    pm_n_larger <- as.integer(len %% pm_n)
    pm_n_smaller <- as.integer(pm_n - pm_n_larger)
    size <- len / pm_n
    larger_size <- as.integer(ceiling(size))
    smaller_size <- as.integer(floor(size))
    larger_threshold <- larger_size * pm_n_larger
    bins <- pm_if_else(
      x <= larger_threshold,
      (x + (larger_size - 1L)) / larger_size,
      (x + (-larger_threshold + smaller_size - 1L)) / smaller_size + pm_n_larger
    )
    as.integer(floor(bins))
  }
}

pm_percent_rank <- function(x) {
  (pm_min_rank(x) - 1) / (sum(!is.na(x)) - 1)
}

pm_row_number <- function(x) {
  if (missing(x)) seq_len(pm_n()) else rank(x, ties.method = "first", na.last = "keep")
}
```
