## Supplementary material for "AMR - An R Package for Working with Antimicrobial Resistance Data": Source package: ab.R

```
# ==================================================================== #
# TITLE                                                                #
# Antimicrobial Resistance (AMR) Analysis for R                        #
#                                                                      #
# SOURCE                                                               #
# https://github.com/msberends/AMR                                     #
#                                                                      #
# LICENCE                                                              #
# (c) 2018-2021 Berends MS, Luz CF et al.                              #
# Developed at the University of Groningen, the Netherlands, in        #
# collaboration with non-profit organisations Certe Medical            #
# Diagnostics & Advice, and University Medical Center Groningen.       # 
#                                                                      #
# This R package is free software; you can freely use and distribute   #
# it for both personal and commercial purposes under the terms of the  #
# GNU General Public License version 2.0 (GNU GPL-2), as published by  #
# the Free Software Foundation.                                        #
# We created this package for both routine data analysis and academic  #
# research and it was publicly released in the hope that it will be    #
# useful, but it comes WITHOUT ANY WARRANTY OR LIABILITY.              #
#                                                                      #
# Visit our website for the full manual and a complete tutorial about  #
# how to conduct AMR analysis: https://msberends.github.io/AMR/        #
# ==================================================================== #

#' Transform input to an antibiotic ID
#'
#' Use this function to determine the antibiotic code of one or more antibiotics. The data set [antibiotics] will be searched for abbreviations, official names and synonyms (brand names).
#' @inheritSection lifecycle Stable lifecycle
#' @param x character vector to determine to antibiotic ID
#' @param flag_multiple_results logical to indicate whether a note should be printed to the console that probably more than one antibiotic code or name can be retrieved from a single input value.
#' @param info logical to indicate whether a progress bar should be printed
#' @param ... arguments passed on to internal functions
#' @rdname as.ab
#' @inheritSection WHOCC WHOCC
#' @details All entries in the [antibiotics] data set have three different identifiers: a human readable EARS-Net code (column `ab`, used by ECDC and WHONET), an ATC code (column `atc`, used by WHO), and a CID code (column `cid`, Compound ID, used by PubChem). The data set contains more than 5,000 official brand names from many different countries, as found in PubChem.
#' 
#' All these properties will be searched for the user input. The [as.ab()] can correct for different forms of misspelling:
#' 
#'  * Wrong spelling of drug names (such as "tobramicin" or "gentamycin"), which corrects for most audible similarities such as f/ph, x/ks, c/z/s, t/th, etc.
#'  * Too few or too many vowels or consonants
#'  * Switching two characters (such as "mreopenem", often the case in clinical data, when doctors typed too fast)
#'  * Digitalised paper records, leaving artefacts like 0/o/O (zero and O's), B/8, n/r, etc.
#'
#' Use the [`ab_*`][ab_property()] functions to get properties based on the returned antibiotic ID, see Examples.
#' 
#' Note: the [as.ab()] and [`ab_*`][ab_property()] functions may use very long regular expression to match brand names of antimicrobial agents. This may fail on some systems.
#' @section Source:
#' World Health Organization (WHO) Collaborating Centre for Drug Statistics Methodology: \url{https://www.whocc.no/atc_ddd_index/}
#'
#' WHONET 2019 software: \url{http://www.whonet.org/software.html}
#'
#' European Commission Public Health PHARMACEUTICALS - COMMUNITY REGISTER: \url{http://ec.europa.eu/health/documents/community-register/html/atc.htm}
#' @aliases ab
#' @return A [character] [vector] with additional class [`ab`]
#' @seealso 
#' * [antibiotics] for the [data.frame] that is being used to determine ATCs
#' * [ab_from_text()] for a function to retrieve antimicrobial drugs from clinical text (from health care records)
#' @inheritSection AMR Reference data publicly available
#' @inheritSection AMR Read more on our website!
#' @export
#' @examples
#' # these examples all return "ERY", the ID of erythromycin:
#' as.ab("J01FA01")
#' as.ab("J 01 FA 01")
#' as.ab("Erythromycin")
#' as.ab("eryt")
#' as.ab("   eryt 123")
#' as.ab("ERYT")
#' as.ab("ERY")
#' as.ab("eritromicine") # spelled wrong, yet works
#' as.ab("Erythrocin")   # trade name
#' as.ab("Romycin")      # trade name
#' 
#' # spelling from different languages and dyslexia are no problem
#' ab_atc("ceftriaxon")
#' ab_atc("cephtriaxone")     # small spelling error
#' ab_atc("cephthriaxone")    # or a bit more severe
#' ab_atc("seephthriaaksone") # and even this works
#'
#' # use ab_* functions to get a specific properties (see ?ab_property);
#' # they use as.ab() internally:
#' ab_name("J01FA01")    # "Erythromycin"
#' ab_name("eryt")       # "Erythromycin"
as.ab <- function(x, flag_multiple_results = TRUE, info = TRUE, ...) {
  meet_criteria(x, allow_class = c("character", "numeric", "integer", "factor"), allow_NA = TRUE)
  meet_criteria(flag_multiple_results, allow_class = "logical", has_length = 1)
  meet_criteria(info, allow_class = "logical", has_length = 1)
  
  check_dataset_integrity()
  
  if (is.ab(x)) {
    return(x)
  }
  
  initial_search <- is.null(list(...)$initial_search)
  already_regex <- isTRUE(list(...)$already_regex)
  
  if (all(toupper(x) %in% antibiotics$ab)) {
    # valid AB code, but not yet right class
    return(set_clean_class(toupper(x),
                           new_class = c("ab", "character")))
  }
  
  x_bak <- x
  x <- toupper(x)
  # remove diacritics
  x <- iconv(x, from = "UTF-8", to = "ASCII//TRANSLIT")
  x <- gsub('"', "", x, fixed = TRUE)
  x <- gsub("(specimen|specimen date|specimen_date|spec_date)", "", x, ignore.case = TRUE, perl = TRUE)
  x_bak_clean <- x
  if (already_regex == FALSE) {
    x_bak_clean <- generalise_antibiotic_name(x_bak_clean)
  }
  
  x <- unique(x_bak_clean) # this means that every x is in fact generalise_antibiotic_name(x)
  x_new <- rep(NA_character_, length(x))
  x_unknown <- character(0)
  
  note_if_more_than_one_found <- function(found, index, from_text) {
    if (initial_search == TRUE & isTRUE(length(from_text) > 1)) {
      abnames <- ab_name(from_text, tolower = TRUE, initial_search = FALSE)
      if (ab_name(found[1L], language = NULL) %like% "clavulanic acid") {
        abnames <- abnames[!abnames == "clavulanic acid"]
      }
      if (length(abnames) > 1) {
        message_("More than one result was found for item ", index, ": ",
                 paste0(abnames, collapse = ", "))
      }
    }
    found[1L]
  }
  
  if (initial_search == TRUE) {
    progress <- progress_ticker(n = length(x), n_min = ifelse(isTRUE(info), 25, length(x) + 1)) # start if n >= 25
    on.exit(close(progress))
  }
  
  for (i in seq_len(length(x))) {
    if (initial_search == TRUE) {
      progress$tick()
    }
    
    if (is.na(x[i]) | is.null(x[i])) {
      next
    }
    if (identical(x[i], "") |
        # no short names:
        nchar(x[i]) <= 2 |
        # prevent "bacteria" from coercing to TMP, since Bacterial is a brand name of it:
        identical(tolower(x[i]), "bacteria")) {
      x_unknown <- c(x_unknown, x_bak[x[i] == x_bak_clean][1])
      next
    }
    
    if (isTRUE(flag_multiple_results) & x[i] %like% "[ ]") {
      from_text <- tryCatch(suppressWarnings(ab_from_text(x[i], initial_search = FALSE, translate_ab = FALSE)[[1]]),
                            error = function(e) character(0))
    } else {
      from_text <- character(0)
    }
    
    # exact name
    found <- antibiotics[which(AB_lookup$generalised_name == x[i]), ]$ab
    if (length(found) > 0) {
      x_new[i] <- found[1L]
      next
    }
    
    # exact AB code
    found <- antibiotics[which(antibiotics$ab == x[i]), ]$ab
    if (length(found) > 0) {
      x_new[i] <- note_if_more_than_one_found(found, i, from_text)
      next
    }
    
    # exact ATC code
    found <- antibiotics[which(antibiotics$atc == x[i]), ]$ab
    if (length(found) > 0) {
      x_new[i] <- note_if_more_than_one_found(found, i, from_text)
      next
    }
    
    # exact CID code
    found <- antibiotics[which(antibiotics$cid == x[i]), ]$ab
    if (length(found) > 0) {
      x_new[i] <- note_if_more_than_one_found(found, i, from_text)
      next
    }
    
    # exact LOINC code
    loinc_found <- unlist(lapply(AB_lookup$generalised_loinc,
                                 function(s) x[i] %in% s))
    found <- antibiotics$ab[loinc_found == TRUE]
    if (length(found) > 0) {
      x_new[i] <- note_if_more_than_one_found(found, i, from_text)
      next
    }
    
    # exact synonym
    synonym_found <- unlist(lapply(AB_lookup$generalised_synonyms,
                                   function(s) x[i] %in% s))
    found <- antibiotics$ab[synonym_found == TRUE]
    if (length(found) > 0) {
      x_new[i] <- note_if_more_than_one_found(found, i, from_text)
      next
    }
    
    # exact abbreviation
    abbr_found <- unlist(lapply(AB_lookup$generalised_abbreviations,
                                function(s) x[i] %in% s))
    found <- antibiotics$ab[abbr_found == TRUE]
    if (length(found) > 0) {
      x_new[i] <- note_if_more_than_one_found(found, i, from_text)
      next
    }
    
    # allow characters that resemble others, but only continue when having more than 3 characters
    if (nchar(x[i]) <= 3) {
      x_unknown <- c(x_unknown, x_bak[x[i] == x_bak_clean][1])
      next
    }
    x_spelling <- x[i]
    if (already_regex == FALSE) {
      x_spelling <- gsub("[IY]+", "[IY]+", x_spelling, perl = TRUE)
      x_spelling <- gsub("(C|K|Q|QU|S|Z|X|KS)+", "(C|K|Q|QU|S|Z|X|KS)+", x_spelling, perl = TRUE)
      x_spelling <- gsub("(PH|F|V)+", "(PH|F|V)+", x_spelling, perl = TRUE)
      x_spelling <- gsub("(TH|T)+", "(TH|T)+", x_spelling, perl = TRUE)
      x_spelling <- gsub("A+", "A+", x_spelling, perl = TRUE)
      x_spelling <- gsub("E+", "E+", x_spelling, perl = TRUE)
      x_spelling <- gsub("O+", "O+", x_spelling, perl = TRUE)
      # allow any ending of -in/-ine and -im/-ime
      x_spelling <- gsub("(\\[IY\\]\\+(N|M)|\\[IY\\]\\+(N|M)E\\+)$", "[IY]+(N|M)E*", x_spelling, perl = TRUE)
      # allow any ending of -ol/-ole
      x_spelling <- gsub("(O\\+L|O\\+LE\\+)$", "O+LE*", x_spelling, perl = TRUE)
      # allow any ending of -on/-one
      x_spelling <- gsub("(O\\+N|O\\+NE\\+)$", "O+NE*", x_spelling, perl = TRUE)
      # replace multiple same characters to single one with '+', like "ll" -> "l+"
      x_spelling <- gsub("(.)\\1+", "\\1+", x_spelling, perl = TRUE)
      # replace spaces and slashes with a possibility on both
      x_spelling <- gsub("[ /]", "( .*|.*/)", x_spelling, perl = TRUE)
      # correct for digital reading text (OCR)
      x_spelling <- gsub("[NRD8B]", "[NRD8B]", x_spelling, perl = TRUE)
      x_spelling <- gsub("(O|0)", "(O|0)+", x_spelling, perl = TRUE)
      x_spelling <- gsub("++", "+", x_spelling, fixed = TRUE)
    }
    
    # try if name starts with it
    found <- antibiotics[which(AB_lookup$generalised_name %like% paste0("^", x_spelling)), ]$ab
    if (length(found) > 0) {
      x_new[i] <- note_if_more_than_one_found(found, i, from_text)
      next
    }
    # try if name ends with it
    found <- antibiotics[which(AB_lookup$generalised_name %like% paste0(x_spelling, "$")), ]$ab
    if (nchar(x[i]) >= 4 & length(found) > 0) {
      x_new[i] <- note_if_more_than_one_found(found, i, from_text)
      next
    }
    
    # and try if any synonym starts with it
    synonym_found <- unlist(lapply(AB_lookup$generalised_synonyms,
                                   function(s) any(s %like% paste0("^", x_spelling))))
    found <- antibiotics$ab[synonym_found == TRUE]
    if (length(found) > 0) {
      x_new[i] <- note_if_more_than_one_found(found, i, from_text)
      next
    }
    
    # INITIAL SEARCH - More uncertain results ----
    
    if (initial_search == TRUE) {
      # only run on first try
      
      # try by removing all spaces
      if (x[i] %like% " ") {
        found <- suppressWarnings(as.ab(gsub(" +", "", x[i], perl = TRUE), initial_search = FALSE))
        if (length(found) > 0 & !is.na(found)) {
          x_new[i] <- note_if_more_than_one_found(found, i, from_text)
          next
        }
      }
      
      # try by removing all spaces and numbers
      if (x[i] %like% " " | x[i] %like% "[0-9]") {
        found <- suppressWarnings(as.ab(gsub("[ 0-9]", "", x[i], perl = TRUE), initial_search = FALSE))
        if (length(found) > 0 & !is.na(found)) {
          x_new[i] <- note_if_more_than_one_found(found, i, from_text)
          next
        }
      }
      
      # transform back from other languages and try again
      x_translated <- paste(lapply(strsplit(x[i], "[^A-Z0-9]"),
                                   function(y) {
                                     for (i in seq_len(length(y))) {
                                       y[i] <- ifelse(tolower(y[i]) %in% tolower(translations_file$replacement),
                                                      translations_file[which(tolower(translations_file$replacement) == tolower(y[i]) &
                                                                                !isFALSE(translations_file$fixed)), "pattern"],
                                                      y[i])
                                     }
                                     generalise_antibiotic_name(y)
                                   })[[1]],
                            collapse = "/")
      x_translated_guess <- suppressWarnings(as.ab(x_translated, initial_search = FALSE))
      if (!is.na(x_translated_guess)) {
        x_new[i] <- x_translated_guess
        next
      }
      
      # now also try to coerce brandname combinations like "Amoxy/clavulanic acid"
      x_translated <- paste(lapply(strsplit(x_translated, "[^A-Z0-9 ]"),
                                   function(y) {
                                     for (i in seq_len(length(y))) {
                                       y_name <- suppressWarnings(ab_name(y[i], language = NULL, initial_search = FALSE))
                                       y[i] <- ifelse(!is.na(y_name),
                                                      y_name,
                                                      y[i])
                                     }
                                     generalise_antibiotic_name(y)
                                   })[[1]],
                            collapse = "/")
      x_translated_guess <- suppressWarnings(as.ab(x_translated, initial_search = FALSE))
      if (!is.na(x_translated_guess)) {
        x_new[i] <- x_translated_guess
        next
      }
      
      # try by removing all trailing capitals
      if (x[i] %like_case% "[a-z]+[A-Z]+$") {
        found <- suppressWarnings(as.ab(gsub("[A-Z]+$", "", x[i], perl = TRUE), initial_search = FALSE))
        if (!is.na(found)) {
          x_new[i] <- note_if_more_than_one_found(found, i, from_text)
          next
        }
      }
      
      # keep only letters
      found <- suppressWarnings(as.ab(gsub("[^A-Z]", "", x[i], perl = TRUE), initial_search = FALSE))
      if (!is.na(found)) {
        x_new[i] <- note_if_more_than_one_found(found, i, from_text)
        next
      }
      
      # try from a bigger text, like from a health care record, see ?ab_from_text
      # already calculated above if flag_multiple_results = TRUE
      if (isTRUE(flag_multiple_results)) {
        found <- from_text[1L]
      } else {
        found <- tryCatch(suppressWarnings(ab_from_text(x[i], initial_search = FALSE, translate_ab = FALSE)[[1]][1L]),
                          error = function(e) NA_character_)
      }
      if (!is.na(found)) {
        x_new[i] <- note_if_more_than_one_found(found, i, from_text)
        next
      }
      
      # first 5 except for cephalosporins, then first 7 (those cephalosporins all start quite the same!)
      found <- suppressWarnings(as.ab(substr(x[i], 1, 5), initial_search = FALSE))
      if (!is.na(found) && !ab_group(found, initial_search = FALSE) %like% "cephalosporins") {
        x_new[i] <- note_if_more_than_one_found(found, i, from_text)
        next
      }
      found <- suppressWarnings(as.ab(substr(x[i], 1, 7), initial_search = FALSE))
      if (!is.na(found)) {
        x_new[i] <- note_if_more_than_one_found(found, i, from_text)
        next
      }
      
      # make all consonants facultative
      search_str <- gsub("([BCDFGHJKLMNPQRSTVWXZ])", "\\1*", x[i], perl = TRUE)
      found <- suppressWarnings(as.ab(search_str, initial_search = FALSE, already_regex = TRUE))
      # keep at least 4 normal characters
      if (nchar(gsub(".\\*", "", search_str, perl = TRUE)) < 4) {
        found <- NA
      }
      if (!is.na(found)) {
        x_new[i] <- note_if_more_than_one_found(found, i, from_text)
        next
      }
      
      # make all vowels facultative
      search_str <- gsub("([AEIOUY])", "\\1*", x[i], perl = TRUE)
      found <- suppressWarnings(as.ab(search_str, initial_search = FALSE, already_regex = TRUE))
      # keep at least 5 normal characters
      if (nchar(gsub(".\\*", "", search_str, perl = TRUE)) < 5) {
        found <- NA
      }
      if (!is.na(found)) {
        x_new[i] <- note_if_more_than_one_found(found, i, from_text)
        next
      }
      
      # allow misspelling of vowels
      x_spelling <- gsub("A+", "[AEIOU]+", x_spelling, fixed = TRUE)
      x_spelling <- gsub("E+", "[AEIOU]+", x_spelling, fixed = TRUE)
      x_spelling <- gsub("I+", "[AEIOU]+", x_spelling, fixed = TRUE)
      x_spelling <- gsub("O+", "[AEIOU]+", x_spelling, fixed = TRUE)
      x_spelling <- gsub("U+", "[AEIOU]+", x_spelling, fixed = TRUE)
      found <- suppressWarnings(as.ab(x_spelling, initial_search = FALSE, already_regex = TRUE))
      if (!is.na(found)) {
        x_new[i] <- note_if_more_than_one_found(found, i, from_text)
        next
      }
      
      # try with switched character, like "mreopenem"
      for (j in seq_len(nchar(x[i]))) {
        x_switched <- paste0(
          # beginning part:
          substr(x[i], 1, j - 1),
          # here is the switching of 2 characters:
          substr(x[i], j + 1, j + 1), 
          substr(x[i], j, j), 
          # ending part:
          substr(x[i], j + 2, nchar(x[i])))
        found <- suppressWarnings(as.ab(x_switched, initial_search = FALSE))
        if (!is.na(found)) {
          break
        }
      }
      if (!is.na(found)) {
        x_new[i] <- found[1L]
        next
      }
      
    } # end of initial_search = TRUE
    
    # not found
    x_unknown <- c(x_unknown, x_bak[x[i] == x_bak_clean][1])
  }
  
  if (initial_search == TRUE) {
    close(progress)
  }
  
  # take failed ATC codes apart from rest
  x_unknown_ATCs <- x_unknown[x_unknown %like% "[A-Z][0-9][0-9][A-Z][A-Z][0-9][0-9]"]
  x_unknown <- x_unknown[!x_unknown %in% x_unknown_ATCs]
  if (length(x_unknown_ATCs) > 0) {
    warning_("These ATC codes are not (yet) in the antibiotics data set: ",
             paste('"', sort(unique(x_unknown_ATCs)), '"', sep = "", collapse = ", "),
             ".",
             call = FALSE)
  }
  
  if (length(x_unknown) > 0) {
    warning_("These values could not be coerced to a valid antimicrobial ID: ",
             paste('"', sort(unique(x_unknown)), '"', sep = "", collapse = ", "),
             ".",
             call = FALSE)
  }
  
  x_result <- data.frame(x = x_bak_clean, stringsAsFactors = FALSE) %pm>%
    pm_left_join(data.frame(x = x, x_new = x_new, stringsAsFactors = FALSE), by = "x") %pm>%
   pm_pull(x_new)
  
  if (length(x_result) == 0) {
    x_result <- NA_character_
  }
  
  set_clean_class(x_result,
                  new_class = c("ab", "character"))
}

#' @rdname as.ab
#' @export
is.ab <- function(x) {
  inherits(x, "ab")
}

# will be exported using s3_register() in R/zzz.R
pillar_shaft.ab <- function(x, ...) {
  out <- trimws(format(x))
  out[is.na(x)] <- font_na(NA)
  create_pillar_column(out, align = "left", min_width = 4)
}

# will be exported using s3_register() in R/zzz.R
type_sum.ab <- function(x, ...) {
  "ab"
}

#' @method print ab
#' @export
#' @noRd
print.ab <- function(x, ...) {
  cat("Class <ab>\n")
  print(as.character(x), quote = FALSE)
}

#' @method as.data.frame ab
#' @export
#' @noRd
as.data.frame.ab <- function(x, ...) {
  nm <- deparse1(substitute(x))
  if (!"nm" %in% names(list(...))) {
    as.data.frame.vector(as.ab(x), ..., nm = nm)
  } else {
    as.data.frame.vector(as.ab(x), ...)
  }
}
#' @method [ ab
#' @export
#' @noRd
"[.ab" <- function(x, ...) {
  y <- NextMethod()
  attributes(y) <- attributes(x)
  y
}
#' @method [[ ab
#' @export
#' @noRd
"[[.ab" <- function(x, ...) {
  y <- NextMethod()
  attributes(y) <- attributes(x)
  y
}
#' @method [<- ab
#' @export
#' @noRd
"[<-.ab" <- function(i, j, ..., value) {
  y <- NextMethod()
  attributes(y) <- attributes(i)
  class_integrity_check(y, "antimicrobial code", antibiotics$ab)
}
#' @method [[<- ab
#' @export
#' @noRd
"[[<-.ab" <- function(i, j, ..., value) {
  y <- NextMethod()
  attributes(y) <- attributes(i)
  class_integrity_check(y, "antimicrobial code", antibiotics$ab)
}
#' @method c ab
#' @export
#' @noRd
c.ab <- function(x, ...) {
  y <- NextMethod()
  attributes(y) <- attributes(x)
  class_integrity_check(y, "antimicrobial code", antibiotics$ab)
}

#' @method unique ab
#' @export
#' @noRd
unique.ab <- function(x, incomparables = FALSE, ...) {
  y <- NextMethod()
  attributes(y) <- attributes(x)
  y
}

generalise_antibiotic_name <- function(x) {
  x <- toupper(x)
  # remove suffices
  x <- gsub("_(MIC|RSI|DIS[CK])$", "", x, perl = TRUE)
  # remove disk concentrations, like LVX_NM -> LVX
  x <- gsub("_[A-Z]{2}[0-9_.]{0,3}$", "", x, perl = TRUE)
  # remove part between brackets if that's followed by another string
  x <- gsub("(.*)+ [(].*[)]", "\\1", x)
  # keep only max 1 space
  x <- trimws2(gsub(" +", " ", x, perl = TRUE))
  # non-character, space or number should be a slash
  x <- gsub("[^A-Z0-9 -]", "/", x, perl = TRUE)
  # spaces around non-characters must be removed: amox + clav -> amox/clav
  x <- gsub("(.*[A-Z0-9]) ([^A-Z0-9].*)", "\\1\\2", x, perl = TRUE)
  x <- gsub("(.*[^A-Z0-9]) ([A-Z0-9].*)", "\\1\\2", x, perl = TRUE)
  # remove hyphen after a starting "co"
  x <- gsub("^CO-", "CO", x, perl = TRUE)
  # replace operators with a space
  x <- gsub("(/| AND | WITH | W/|[+]|[-])+", " ", x, perl = TRUE)
  x
}
```
