## Supplementary material for "AMR - An R Package for Working with Antimicrobial Resistance Data": Source package: ab_class_selectors.R

```
# ==================================================================== #
# TITLE                                                                #
# Antimicrobial Resistance (AMR) Analysis for R                        #
#                                                                      #
# SOURCE                                                               #
# https://github.com/msberends/AMR                                     #
#                                                                      #
# LICENCE                                                              #
# (c) 2018-2021 Berends MS, Luz CF et al.                              #
# Developed at the University of Groningen, the Netherlands, in        #
# collaboration with non-profit organisations Certe Medical            #
# Diagnostics & Advice, and University Medical Center Groningen.       # 
#                                                                      #
# This R package is free software; you can freely use and distribute   #
# it for both personal and commercial purposes under the terms of the  #
# GNU General Public License version 2.0 (GNU GPL-2), as published by  #
# the Free Software Foundation.                                        #
# We created this package for both routine data analysis and academic  #
# research and it was publicly released in the hope that it will be    #
# useful, but it comes WITHOUT ANY WARRANTY OR LIABILITY.              #
#                                                                      #
# Visit our website for the full manual and a complete tutorial about  #
# how to conduct AMR analysis: https://msberends.github.io/AMR/        #
# ==================================================================== #

#' Antibiotic class selectors
#' 
#' These functions help to select the columns of antibiotics that are of a specific antibiotic class, without the need to define the columns or antibiotic abbreviations.
#' @inheritParams filter_ab_class 
#' @details \strong{\Sexpr{ifelse(as.double(R.Version()$major) + (as.double(R.Version()$minor) / 10) < 3.2, paste0("NOTE: THESE FUNCTIONS DO NOT WORK ON YOUR CURRENT R VERSION. These functions require R version 3.2 or later - you have ", R.version.string, "."), "")}}
#' 
#' All columns will be searched for known antibiotic names, abbreviations, brand names and codes (ATC, EARS-Net, WHO, etc.) in the [antibiotics] data set. This means that a selector like e.g. [aminoglycosides()] will pick up column names like 'gen', 'genta', 'J01GB03', 'tobra', 'Tobracin', etc.
#' @rdname antibiotic_class_selectors
#' @seealso [filter_ab_class()] for the `filter()` equivalent.
#' @name antibiotic_class_selectors
#' @export
#' @inheritSection AMR Reference data publicly available
#' @inheritSection AMR Read more on our website!
#' @examples 
#' # `example_isolates` is a dataset available in the AMR package.
#' # See ?example_isolates.
#' 
#' # this will select columns 'IPM' (imipenem) and 'MEM' (meropenem):
#' example_isolates[, c(carbapenems())]
#' # this will select columns 'mo', 'AMK', 'GEN', 'KAN' and 'TOB':
#' example_isolates[, c("mo", aminoglycosides())]
#' 
#' if (require("dplyr")) {
#' 
#'   # this will select columns 'IPM' (imipenem) and 'MEM' (meropenem):
#'   example_isolates %>% 
#'     select(carbapenems())
#'     
#'   # this will select columns 'mo', 'AMK', 'GEN', 'KAN' and 'TOB':
#'   example_isolates %>% 
#'     select(mo, aminoglycosides())
#'     
#'   # this will select columns 'mo' and all antimycobacterial drugs ('RIF'):
#'   example_isolates %>% 
#'     select(mo, ab_class("mycobact"))
#'     
#'     
#'   # get bug/drug combinations for only macrolides in Gram-positives:
#'   example_isolates %>% 
#'     filter(mo_is_gram_positive()) %>% 
#'     select(mo, macrolides()) %>% 
#'     bug_drug_combinations() %>%
#'     format()
#'     
#'     
#'   data.frame(some_column = "some_value",
#'              J01CA01 = "S") %>%   # ATC code of ampicillin
#'     select(penicillins())         # only the 'J01CA01' column will be selected
#'     
#'     
#'   # with dplyr 1.0.0 and higher (that adds 'across()'), this is equal:
#'   # (though the row names on the first are more correct)
#'   example_isolates %>% filter_carbapenems("R", "all")
#'   example_isolates %>% filter(across(carbapenems(), ~. == "R"))
#' }
ab_class <- function(ab_class) {
  ab_selector(ab_class, function_name = "ab_class")
}

#' @rdname antibiotic_class_selectors
#' @export
aminoglycosides <- function() {
  ab_selector("aminoglycoside", function_name = "aminoglycosides")
}

#' @rdname antibiotic_class_selectors
#' @export
carbapenems <- function() {
  ab_selector("carbapenem", function_name = "carbapenems")
}

#' @rdname antibiotic_class_selectors
#' @export
cephalosporins <- function() {
  ab_selector("cephalosporin", function_name = "cephalosporins")
}

#' @rdname antibiotic_class_selectors
#' @export
cephalosporins_1st <- function() {
  ab_selector("cephalosporins.*1", function_name = "cephalosporins_1st")
}

#' @rdname antibiotic_class_selectors
#' @export
cephalosporins_2nd <- function() {
  ab_selector("cephalosporins.*2", function_name = "cephalosporins_2nd")
}

#' @rdname antibiotic_class_selectors
#' @export
cephalosporins_3rd <- function() {
  ab_selector("cephalosporins.*3", function_name = "cephalosporins_3rd")
}

#' @rdname antibiotic_class_selectors
#' @export
cephalosporins_4th <- function() {
  ab_selector("cephalosporins.*4", function_name = "cephalosporins_4th")
}

#' @rdname antibiotic_class_selectors
#' @export
cephalosporins_5th <- function() {
  ab_selector("cephalosporins.*5", function_name = "cephalosporins_5th")
}

#' @rdname antibiotic_class_selectors
#' @export
fluoroquinolones <- function() {
  ab_selector("fluoroquinolone", function_name = "fluoroquinolones")
}

#' @rdname antibiotic_class_selectors
#' @export
glycopeptides <- function() {
  ab_selector("glycopeptide", function_name = "glycopeptides")
}

#' @rdname antibiotic_class_selectors
#' @export
macrolides <- function() {
  ab_selector("macrolide", function_name = "macrolides")
}

#' @rdname antibiotic_class_selectors
#' @export
penicillins <- function() {
  ab_selector("penicillin", function_name = "penicillins")
}

#' @rdname antibiotic_class_selectors
#' @export
tetracyclines <- function() {
  ab_selector("tetracycline", function_name = "tetracyclines")
}

ab_selector <- function(ab_class, function_name) {
  meet_criteria(ab_class, allow_class = "character", has_length = 1, .call_depth = 1)
  meet_criteria(function_name, allow_class = "character", has_length = 1, .call_depth = 1)
  
  if (as.double(R.Version()$major) + (as.double(R.Version()$minor) / 10) < 3.2) {
    warning_("antibiotic class selectors such as ", function_name, 
             "() require R version 3.2 or later - you have ", R.version.string,
             call = FALSE)
    return(NULL)
  }
  
  vars_df <- get_current_data(arg_name = NA, call = -3)
  ab_in_data <- get_column_abx(vars_df, info = FALSE)
  
  if (length(ab_in_data) == 0) {
    message_("No antimicrobial agents found.")
    return(NULL)
  }
  
  ab_reference <- subset(antibiotics,
                         group %like% ab_class | 
                           atc_group1 %like% ab_class | 
                           atc_group2 %like% ab_class)
  ab_group <- find_ab_group(ab_class)
  if (ab_group == "") {
    ab_group <- paste0("'", ab_class, "'")
    examples <- ""
  } else {
    examples <- paste0(" (such as ", find_ab_names(ab_class, 2), ")")
  }
  # get the columns with a group names in the chosen ab class
  agents <- ab_in_data[names(ab_in_data) %in% ab_reference$ab]
  if (message_not_thrown_before(function_name)) {
    if (length(agents) == 0) {
      message_("No antimicrobial agents of class ", ab_group, " found", examples, ".")
    } else {
      agents_formatted <- paste0("column '", font_bold(agents, collapse = NULL), "'")
      agents_names <- ab_name(names(agents), tolower = TRUE, language = NULL)
      agents_formatted[agents != agents_names] <- paste0(agents_formatted[agents != agents_names],
                                                         " (", agents_names[agents != agents_names], ")")
      message_("Selecting ", ab_group, ": ", paste(agents_formatted, collapse = ", "),
               as_note = FALSE,
               extra_indent = nchar(paste0("Selecting ", ab_group, ": ")))
    }
   remember_thrown_message(function_name)
 }
  unname(agents)
}
```
