## Supplementary material for "AMR - An R Package for Working with Antimicrobial Resistance Data": Source package: ab_from_text.R

```
# ==================================================================== #
# TITLE                                                                #
# Antimicrobial Resistance (AMR) Analysis for R                        #
#                                                                      #
# SOURCE                                                               #
# https://github.com/msberends/AMR                                     #
#                                                                      #
# LICENCE                                                              #
# (c) 2018-2021 Berends MS, Luz CF et al.                              #
# Developed at the University of Groningen, the Netherlands, in        #
# collaboration with non-profit organisations Certe Medical            #
# Diagnostics & Advice, and University Medical Center Groningen.       # 
#                                                                      #
# This R package is free software; you can freely use and distribute   #
# it for both personal and commercial purposes under the terms of the  #
# GNU General Public License version 2.0 (GNU GPL-2), as published by  #
# the Free Software Foundation.                                        #
# We created this package for both routine data analysis and academic  #
# research and it was publicly released in the hope that it will be    #
# useful, but it comes WITHOUT ANY WARRANTY OR LIABILITY.              #
#                                                                      #
# Visit our website for the full manual and a complete tutorial about  #
# how to conduct AMR analysis: https://msberends.github.io/AMR/        #
# ==================================================================== #

#' Retrieve antimicrobial drug names and doses from clinical text
#' 
#' Use this function on e.g. clinical texts from health care records. It returns a [list] with all antimicrobial drugs, doses and forms of administration found in the texts.
#' @inheritSection lifecycle Maturing lifecycle
#' @param text text to analyse
#' @param type type of property to search for, either `"drug"`, `"dose"` or `"administration"`, see *Examples*
#' @param collapse character to pass on to `paste(..., collapse = ...)` to only return one character per element of `text`, see *Examples*
#' @param translate_ab if `type = "drug"`: a column name of the [antibiotics] data set to translate the antibiotic abbreviations to, using [ab_property()]. Defaults to `FALSE`. Using `TRUE` is equal to using "name".
#' @param thorough_search logical to indicate whether the input must be extensively searched for misspelling and other faulty input values. Setting this to `TRUE` will take considerably more time than when using `FALSE`. At default, it will turn `TRUE` when all input elements contain a maximum of three words.
#' @param ... arguments passed on to [as.ab()]
#' @details This function is also internally used by [as.ab()], although it then only searches for the first drug name and will throw a note if more drug names could have been returned. Note: the [as.ab()] function may use very long regular expression to match brand names of antimicrobial agents. This may fail on some systems.
#' 
#' ## Argument `type`
#' At default, the function will search for antimicrobial drug names. All text elements will be searched for official names, ATC codes and brand names. As it uses [as.ab()] internally, it will correct for misspelling.
#' 
#' With `type = "dose"` (or similar, like "dosing", "doses"), all text elements will be searched for numeric values that are higher than 100 and do not resemble years. The output will be numeric. It supports any unit (g, mg, IE, etc.) and multiple values in one clinical text, see *Examples*.
#' 
#' With `type = "administration"` (or abbreviations, like "admin", "adm"), all text elements will be searched for a form of drug administration. It supports the following forms (including common abbreviations): buccal, implant, inhalation, instillation, intravenous, nasal, oral, parenteral, rectal, sublingual, transdermal and vaginal. Abbreviations for oral (such as 'po', 'per os') will become "oral", all values for intravenous (such as 'iv', 'intraven') will become "iv". It supports multiple values in one clinical text, see *Examples*.
#' 
#' ## Argument `collapse`
#' Without using `collapse`, this function will return a [list]. This can be convenient to use e.g. inside a `mutate()`):\cr
#' `df %>% mutate(abx = ab_from_text(clinical_text))` 
#' 
#' The returned AB codes can be transformed to official names, groups, etc. with all [`ab_*`][ab_property()] functions such as [ab_name()] and [ab_group()], or by using the `translate_ab` argument.
#' 
#' With using `collapse`, this function will return a [character]:\cr
#' `df %>% mutate(abx = ab_from_text(clinical_text, collapse = "|"))` 
#' @export
#' @return A [list], or a  [character] if `collapse` is not `NULL`
#' @inheritSection AMR Read more on our website!
#' @examples 
#' # mind the bad spelling of amoxicillin in this line, 
#' # straight from a true health care record:
#' ab_from_text("28/03/2020 regular amoxicilliin 500mg po tds")
#' 
#' ab_from_text("500 mg amoxi po and 400mg cipro iv")
#' ab_from_text("500 mg amoxi po and 400mg cipro iv", type = "dose")
#' ab_from_text("500 mg amoxi po and 400mg cipro iv", type = "admin")
#' 
#' ab_from_text("500 mg amoxi po and 400mg cipro iv", collapse = ", ")
#' \donttest{
#' # if you want to know which antibiotic groups were administered, do e.g.:
#' abx <- ab_from_text("500 mg amoxi po and 400mg cipro iv")
#' ab_group(abx[[1]])
#' 
#' if (require("dplyr")) {
#'   tibble(clinical_text = c("given 400mg cipro and 500 mg amox",
#'                            "started on doxy iv today")) %>% 
#'     mutate(abx_codes = ab_from_text(clinical_text),
#'            abx_doses = ab_from_text(clinical_text, type = "doses"),
#'            abx_admin = ab_from_text(clinical_text, type = "admin"),
#'            abx_coll = ab_from_text(clinical_text, collapse = "|"),
#'            abx_coll_names = ab_from_text(clinical_text,
#'                                          collapse = "|",
#'                                          translate_ab = "name"),
#'            abx_coll_doses = ab_from_text(clinical_text,
#'                                          type = "doses",
#'                                          collapse = "|"),
#'            abx_coll_admin = ab_from_text(clinical_text,
#'                                          type = "admin",
#'                                          collapse = "|"))
#' 
#' }
#' }
ab_from_text <- function(text,
                         type = c("drug", "dose", "administration"),
                         collapse = NULL,
                         translate_ab = FALSE,
                         thorough_search = NULL,
                         ...) {
  if (missing(type)) {
    type <- type[1L]
  }
  
  meet_criteria(text)
  meet_criteria(type, allow_class = "character", has_length = 1)
  meet_criteria(collapse, has_length = 1, allow_NULL = TRUE)
  meet_criteria(translate_ab, allow_NULL = FALSE) # get_translate_ab() will be more informative about what's allowed
  meet_criteria(thorough_search, allow_class = "logical", has_length = 1, allow_NULL = TRUE)

  type <- tolower(trimws(type))
  
  text <- tolower(as.character(text))
  text_split_all <- strsplit(text, "[ ;.,:\\|]")
  progress <- progress_ticker(n = length(text_split_all), n_min = 5)
  on.exit(close(progress))
  
  if (type %like% "(drug|ab|anti)") {
    
    translate_ab <- get_translate_ab(translate_ab)
    
    if (isTRUE(thorough_search) | 
        (isTRUE(is.null(thorough_search)) & max(vapply(FUN.VALUE = double(1), text_split_all, length), na.rm = TRUE) <= 3)) {
      text_split_all <- text_split_all[nchar(text_split_all) >= 4 & grepl("[a-z]+", text_split_all)]
      result <- lapply(text_split_all, function(text_split) {
        progress$tick()
        suppressWarnings(
          out <- as.ab(text_split, ...)
        )
      })
      
    } else {
      # no thorough search
      abbr <- unlist(antibiotics$abbreviations)
      abbr <- abbr[nchar(abbr) >= 4]
      names_atc <- substr(c(antibiotics$name, antibiotics$atc), 1, 5)
      synonyms <- unlist(antibiotics$synonyms)
      synonyms <- synonyms[nchar(synonyms) >= 4]
      # regular expression must not be too long, so split synonyms in two:
      synonyms_part1 <- synonyms[seq_len(0.5 * length(synonyms))]
      synonyms_part2 <- synonyms[!synonyms %in% synonyms_part1]
      to_regex <- function(x) {
        paste0("^(",
               paste0(unique(gsub("[^a-z0-9]+", "", sort(tolower(x)))), collapse = "|"),
               ").*")
      }
      result <- lapply(text_split_all, function(text_split) {
        progress$tick()
        suppressWarnings(
          out <- as.ab(unique(c(text_split[text_split %like_case% to_regex(abbr)],
                                text_split[text_split %like_case% to_regex(names_atc)],
                                text_split[text_split %like_case% to_regex(synonyms_part1)],
                                text_split[text_split %like_case% to_regex(synonyms_part2)])
          ),
          ...)
        )
      })
    }
    
    close(progress)
    
    result <- lapply(result, function(out) {
      out <- out[!is.na(out)]
      if (length(out) == 0) {
        as.ab(NA)
      } else {
        if (!isFALSE(translate_ab)) {
          out <- ab_property(out, property = translate_ab, initial_search = FALSE)
        }
        out
      }
      
    })
    
  } else if (type %like% "dos") {
    text_split_all <- strsplit(text, " ")
    result <- lapply(text_split_all, function(text_split) {
      text_split <- text_split[text_split %like% "^[0-9]{2,}(/[0-9]+)?[a-z]*$"]
      # only left part of "/", like 500 in  "500/125"
      text_split <-  gsub("/.*", "", text_split)
      text_split <- gsub(",", ".", text_split, fixed = TRUE) # foreign system using comma as decimal sep
      text_split <- as.double(gsub("[^0-9.]", "", text_split))
      # minimal 100 units/mg and no years that unlikely doses
      text_split <- text_split[text_split >= 100 & !text_split %in% c(1951:1999, 2001:2049)]
      
      if (length(text_split) > 0) {
        text_split
      } else {
        NA_real_
      }
    })
    
  } else if (type %like% "adm") {
    result <- lapply(text_split_all, function(text_split) {
      text_split <- text_split[text_split %like% "(^iv$|intraven|^po$|per os|oral|implant|inhal|instill|nasal|paren|rectal|sublingual|buccal|trans.*dermal|vaginal)"]
      if (length(text_split) > 0) {
        text_split <- gsub("(^po$|.*per os.*)", "oral", text_split)
        text_split <- gsub("(^iv$|.*intraven.*)", "iv", text_split)
        text_split
      } else {
        NA_character_
      }
    })
    
  } else {
    stop_("`type` must be either 'drug', 'dose' or 'administration'")
  }
  
  # collapse text if needed
  if (!is.null(collapse)) {
    result <- vapply(FUN.VALUE = character(1), result, function(x) {
      if (length(x) == 1 & all(is.na(x))) {
        NA_character_
      } else {
        paste0(x, collapse = collapse)
      }
    })
  }
  
  result
  
}
```
