## Supplementary material for "AMR - An R Package for Working with Antimicrobial Resistance Data": Source package: ab_property.R

```
# ==================================================================== #
# TITLE                                                                #
# Antimicrobial Resistance (AMR) Analysis for R                        #
#                                                                      #
# SOURCE                                                               #
# https://github.com/msberends/AMR                                     #
#                                                                      #
# LICENCE                                                              #
# (c) 2018-2021 Berends MS, Luz CF et al.                              #
# Developed at the University of Groningen, the Netherlands, in        #
# collaboration with non-profit organisations Certe Medical            #
# Diagnostics & Advice, and University Medical Center Groningen.       # 
#                                                                      #
# This R package is free software; you can freely use and distribute   #
# it for both personal and commercial purposes under the terms of the  #
# GNU General Public License version 2.0 (GNU GPL-2), as published by  #
# the Free Software Foundation.                                        #
# We created this package for both routine data analysis and academic  #
# research and it was publicly released in the hope that it will be    #
# useful, but it comes WITHOUT ANY WARRANTY OR LIABILITY.              #
#                                                                      #
# Visit our website for the full manual and a complete tutorial about  #
# how to conduct AMR analysis: https://msberends.github.io/AMR/        #
# ==================================================================== #

#' Get properties of an antibiotic
#'
#' Use these functions to return a specific property of an antibiotic from the [antibiotics] data set. All input values will be evaluated internally with [as.ab()].
#' @inheritSection lifecycle Stable lifecycle
#' @param x any (vector of) text that can be coerced to a valid antibiotic code with [as.ab()]
#' @param tolower logical to indicate whether the first character of every output should be transformed to a lower case character. This will lead to e.g. "polymyxin B" and not "polymyxin b".
#' @param property one of the column names of one of the [antibiotics] data set
#' @param language language of the returned text, defaults to system language (see [get_locale()]) and can also be set with `getOption("AMR_locale")`. Use `language = NULL` or `language = ""` to prevent translation.
#' @param administration way of administration, either `"oral"` or `"iv"`
#' @param units a logical to indicate whether the units instead of the DDDs itself must be returned, see Examples
#' @param open browse the URL using [utils::browseURL()]
#' @param ... other arguments passed on to [as.ab()]
#' @details All output will be [translate]d where possible.
#' 
#' The function [ab_url()] will return the direct URL to the official WHO website. A warning will be returned if the required ATC code is not available.
#' @inheritSection as.ab Source
#' @rdname ab_property
#' @name ab_property
#' @return 
#' - An [integer] in case of [ab_cid()]
#' - A named [list] in case of [ab_info()] and multiple [ab_synonyms()]/[ab_tradenames()]
#' - A [double] in case of [ab_ddd()]
#' - A [character] in all other cases
#' @export
#' @seealso [antibiotics]
#' @inheritSection AMR Reference data publicly available
#' @inheritSection AMR Read more on our website!
#' @examples
#' # all properties:
#' ab_name("AMX")       # "Amoxicillin"
#' ab_atc("AMX")        # J01CA04 (ATC code from the WHO)
#' ab_cid("AMX")        # 33613 (Compound ID from PubChem)
#' ab_synonyms("AMX")   # a list with brand names of amoxicillin
#' ab_tradenames("AMX") # same
#' ab_group("AMX")      # "Beta-lactams/penicillins"
#' ab_atc_group1("AMX") # "Beta-lactam antibacterials, penicillins"
#' ab_atc_group2("AMX") # "Penicillins with extended spectrum"
#' ab_url("AMX")        # link to the official WHO page
#'
#' # smart lowercase tranformation
#' ab_name(x = c("AMC", "PLB"))  # "Amoxicillin/clavulanic acid" "Polymyxin B"
#' ab_name(x = c("AMC", "PLB"),
#'         tolower = TRUE)       # "amoxicillin/clavulanic acid" "polymyxin B"
#'
#' # defined daily doses (DDD)
#' ab_ddd("AMX", "oral")               #  1
#' ab_ddd("AMX", "oral", units = TRUE) # "g"
#' ab_ddd("AMX", "iv")                 #  1
#' ab_ddd("AMX", "iv", units = TRUE)   # "g"
#'
#' ab_info("AMX")       # all properties as a list
#'
#' # all ab_* functions use as.ab() internally, so you can go from 'any' to 'any':
#' ab_atc("AMP")           # ATC code of AMP (ampicillin)
#' ab_group("J01CA01")     # Drug group of ampicillins ATC code
#' ab_loinc("ampicillin")  # LOINC codes of ampicillin
#' ab_name("21066-6")      # "Ampicillin" (using LOINC)
#' ab_name(6249)           # "Ampicillin" (using CID)
#' ab_name("J01CA01")      # "Ampicillin" (using ATC)
#' 
#' # spelling from different languages and dyslexia are no problem
#' ab_atc("ceftriaxon")
#' ab_atc("cephtriaxone")
#' ab_atc("cephthriaxone")
#' ab_atc("seephthriaaksone")
ab_name <- function(x, language = get_locale(), tolower = FALSE, ...) {
  meet_criteria(x, allow_NA = TRUE)
  meet_criteria(language, has_length = 1, is_in = c(LANGUAGES_SUPPORTED, ""), allow_NULL = TRUE, allow_NA = TRUE)
  meet_criteria(tolower, allow_class = "logical", has_length = 1)
  
  x <- translate_AMR(ab_validate(x = x, property = "name", ...), language = language)
  if (tolower == TRUE) {
    # use perl to only transform the first character
    # as we want "polymyxin B", not "polymyxin b"
    x <- gsub("^([A-Z])", "\\L\\1", x, perl = TRUE)
  }
  x
}

#' @rdname ab_property
#' @aliases ATC
#' @export
ab_atc <- function(x, ...) {
  meet_criteria(x, allow_NA = TRUE)
  ab_validate(x = x, property = "atc", ...)
}

#' @rdname ab_property
#' @export
ab_cid <- function(x, ...) {
  meet_criteria(x, allow_NA = TRUE)
  ab_validate(x = x, property = "cid", ...)
}

#' @rdname ab_property
#' @export
ab_synonyms <- function(x, ...) {
  meet_criteria(x, allow_NA = TRUE)
  syns <- ab_validate(x = x, property = "synonyms", ...)
  names(syns) <- x
  if (length(syns) == 1) {
    unname(unlist(syns))
  } else {
    syns
  }
}

#' @rdname ab_property
#' @export
ab_tradenames <- function(x, ...) {
  meet_criteria(x, allow_NA = TRUE)
  ab_synonyms(x, ...)
}

#' @rdname ab_property
#' @export
ab_group <- function(x, language = get_locale(), ...) {
  meet_criteria(x, allow_NA = TRUE)
  meet_criteria(language, has_length = 1, is_in = c(LANGUAGES_SUPPORTED, ""), allow_NULL = TRUE, allow_NA = TRUE)
  translate_AMR(ab_validate(x = x, property = "group", ...), language = language)
}

#' @rdname ab_property
#' @export
ab_atc_group1 <- function(x, language = get_locale(), ...) {
  meet_criteria(x, allow_NA = TRUE)
  meet_criteria(language, has_length = 1, is_in = c(LANGUAGES_SUPPORTED, ""), allow_NULL = TRUE, allow_NA = TRUE)
  translate_AMR(ab_validate(x = x, property = "atc_group1", ...), language = language)
}

#' @rdname ab_property
#' @export
ab_atc_group2 <- function(x, language = get_locale(), ...) {
  meet_criteria(x, allow_NA = TRUE)
  meet_criteria(language, has_length = 1, is_in = c(LANGUAGES_SUPPORTED, ""), allow_NULL = TRUE, allow_NA = TRUE)
  translate_AMR(ab_validate(x = x, property = "atc_group2", ...), language = language)
}

#' @rdname ab_property
#' @export
ab_loinc <- function(x, ...) {
  meet_criteria(x, allow_NA = TRUE)
  loincs <- ab_validate(x = x, property = "loinc", ...)
  names(loincs) <- x
  if (length(loincs) == 1) {
    unname(unlist(loincs))
  } else {
    loincs
  }
}

#' @rdname ab_property
#' @export
ab_ddd <- function(x, administration = "oral", units = FALSE, ...) {
  meet_criteria(x, allow_NA = TRUE)
  meet_criteria(administration, is_in = c("oral", "iv"), has_length = 1)
  meet_criteria(units, allow_class = "logical", has_length = 1)

  ddd_prop <- administration
  if (units == TRUE) {
    ddd_prop <- paste0(ddd_prop, "_units")
  } else {
    ddd_prop <- paste0(ddd_prop, "_ddd")
  }
  ab_validate(x = x, property = ddd_prop, ...)
}

#' @rdname ab_property
#' @export
ab_info <- function(x, language = get_locale(), ...) {
  meet_criteria(x, allow_NA = TRUE)
  meet_criteria(language, has_length = 1, is_in = c(LANGUAGES_SUPPORTED, ""), allow_NULL = TRUE, allow_NA = TRUE)
  
  x <- as.ab(x, ...)
  list(ab = as.character(x),
             atc = ab_atc(x),
             cid = ab_cid(x),
             name = ab_name(x, language = language),
             group = ab_group(x, language = language),
             atc_group1 = ab_atc_group1(x, language = language),
             atc_group2 = ab_atc_group2(x, language = language),
             tradenames = ab_tradenames(x),
             ddd = list(oral = list(amount = ab_ddd(x, administration = "oral", units = FALSE),
                                    units = ab_ddd(x, administration = "oral", units = TRUE)),
                        iv = list(amount = ab_ddd(x, administration = "iv", units = FALSE),
                                  units = ab_ddd(x, administration = "iv", units = TRUE))))
}


#' @rdname ab_property
#' @export
ab_url <- function(x, open = FALSE, ...) {
  meet_criteria(x, allow_NA = TRUE)
  meet_criteria(open, allow_class = "logical", has_length = 1)
  
  ab <- as.ab(x = x, ... = ...)
  u <- paste0("https://www.whocc.no/atc_ddd_index/?code=", ab_atc(ab), "&showdescription=no")
  u[is.na(ab_atc(ab))] <- NA_character_
  names(u) <- ab_name(ab)
  
  NAs <- ab_name(ab, tolower = TRUE, language = NULL)[!is.na(ab) & is.na(ab_atc(ab))]
  if (length(NAs) > 0) {
    warning_("No ATC code available for ", paste0(NAs, collapse = ", "), ".")
  }
  
  if (open == TRUE) {
    if (length(u) > 1 & !is.na(u[1L])) {
      warning_("Only the first URL will be opened, as `browseURL()` only suports one string.")
    }
    if (!is.na(u[1L])) {
      utils::browseURL(u[1L])
    }
  }
  u
}

#' @rdname ab_property
#' @export
ab_property <- function(x, property = "name", language = get_locale(), ...) {
  meet_criteria(x, allow_NA = TRUE)
  meet_criteria(property, is_in = colnames(antibiotics), has_length = 1)
  meet_criteria(language, is_in = c(LANGUAGES_SUPPORTED, ""), has_length = 1, allow_NULL = TRUE, allow_NA = TRUE)
  translate_AMR(ab_validate(x = x, property = property, ...), language = language)
}

ab_validate <- function(x, property, ...) {
  
  check_dataset_integrity()
  
  # try to catch an error when inputting an invalid argument
  # so the 'call.' can be set to FALSE
  tryCatch(x[1L] %in% antibiotics[1, property],
           error = function(e) stop(e$message, call. = FALSE))
  x_bak <- x
  if (!all(x %in% antibiotics[, property])) {
    x <- data.frame(ab = as.ab(x, ...), stringsAsFactors = FALSE) %pm>%
      pm_left_join(antibiotics, by = "ab") %pm>%
      pm_pull(property)
  }
  if (property == "ab") {
    return(set_clean_class(x, new_class = c("ab", "character")))
  } else if (property == "cid") {
    return(as.integer(x))
  } else if (property %like% "ddd") {
    return(as.double(x))
  } else {
    x[is.na(x) & !is.na(x_bak)] <- NA
    return(x)
  }
}
```
