## Supplementary material for "AMR - An R Package for Working with Antimicrobial Resistance Data": Source package: age.R

```
# ==================================================================== #
# TITLE                                                                #
# Antimicrobial Resistance (AMR) Analysis for R                        #
#                                                                      #
# SOURCE                                                               #
# https://github.com/msberends/AMR                                     #
#                                                                      #
# LICENCE                                                              #
# (c) 2018-2021 Berends MS, Luz CF et al.                              #
# Developed at the University of Groningen, the Netherlands, in        #
# collaboration with non-profit organisations Certe Medical            #
# Diagnostics & Advice, and University Medical Center Groningen.       # 
#                                                                      #
# This R package is free software; you can freely use and distribute   #
# it for both personal and commercial purposes under the terms of the  #
# GNU General Public License version 2.0 (GNU GPL-2), as published by  #
# the Free Software Foundation.                                        #
# We created this package for both routine data analysis and academic  #
# research and it was publicly released in the hope that it will be    #
# useful, but it comes WITHOUT ANY WARRANTY OR LIABILITY.              #
#                                                                      #
# Visit our website for the full manual and a complete tutorial about  #
# how to conduct AMR analysis: https://msberends.github.io/AMR/        #
# ==================================================================== #

#' Age in years of individuals
#'
#' Calculates age in years based on a reference date, which is the sytem date at default.
#' @inheritSection lifecycle Stable lifecycle
#' @param x date(s), will be coerced with [as.POSIXlt()]
#' @param reference reference date(s) (defaults to today), will be coerced with [as.POSIXlt()]
#' @param exact a logical to indicate whether age calculation should be exact, i.e. with decimals. It divides the number of days of [year-to-date](https://en.wikipedia.org/wiki/Year-to-date) (YTD) of `x` by the number of days in the year of `reference` (either 365 or 366).
#' @param na.rm a logical to indicate whether missing values should be removed
#' @param ... arguments passed on to [as.POSIXlt()], such as `origin`
#' @details Ages below 0 will be returned as `NA` with a warning. Ages above 120 will only give a warning.
#' @return An [integer] (no decimals) if `exact = FALSE`, a [double] (with decimals) otherwise
#' @seealso To split ages into groups, use the [age_groups()] function.
#' @inheritSection AMR Read more on our website!
#' @export
#' @examples
#' # 10 random birth dates
#' df <- data.frame(birth_date = Sys.Date() - runif(10) * 25000)
#' # add ages
#' df$age <- age(df$birth_date)
#' # add exact ages
#' df$age_exact <- age(df$birth_date, exact = TRUE)
#'
#' df
age <- function(x, reference = Sys.Date(), exact = FALSE, na.rm = FALSE, ...) {
  meet_criteria(x, allow_class = c("character", "Date", "POSIXt"))
  meet_criteria(reference, allow_class = c("character", "Date", "POSIXt"))
  meet_criteria(exact, allow_class = "logical", has_length = 1)
  meet_criteria(na.rm, allow_class = "logical", has_length = 1)
  
  if (length(x) != length(reference)) {
    stop_if(length(reference) != 1, "`x` and `reference` must be of same length, or `reference` must be of length 1.")
    reference <- rep(reference, length(x))
  }
  x <- as.POSIXlt(x, ...)
  reference <- as.POSIXlt(reference, ...)
  
  # from https://stackoverflow.com/a/25450756/4575331
  years_gap <- reference$year - x$year
  ages <- ifelse(reference$mon < x$mon | (reference$mon == x$mon & reference$mday < x$mday),
                 as.integer(years_gap - 1),
                 as.integer(years_gap))
  
  # add decimals
  if (exact == TRUE) {
    # get dates of `x` when `x` would have the year of `reference`
    x_in_reference_year <- as.POSIXlt(paste0(format(reference, "%Y"), format(x, "-%m-%d")))
    # get differences in days
    n_days_x_rest <- as.double(difftime(reference, x_in_reference_year, units = "days"))
    # get numbers of days the years of `reference` has for a reliable denominator
    n_days_reference_year <- as.POSIXlt(paste0(format(reference, "%Y"), "-12-31"))$yday + 1
    # add decimal parts of year
    mod <- n_days_x_rest / n_days_reference_year
    # negative mods are cases where `x_in_reference_year` > `reference` - so 'add' a year
    mod[mod < 0] <- 1 + mod[mod < 0]
    # and finally add to ages
    ages <- ages + mod
  }
  
  if (any(ages < 0, na.rm = TRUE)) {
    ages[ages < 0] <- NA
    warning_("NAs introduced for ages below 0.", call = TRUE)
  }
  if (any(ages > 120, na.rm = TRUE)) {
    warning_("Some ages are above 120.", call = TRUE)
  }
  
  if (isTRUE(na.rm)) {
    ages <- ages[!is.na(ages)]
  }
  
  ages
}

#' Split ages into age groups
#'
#' Split ages into age groups defined by the `split` argument. This allows for easier demographic (antimicrobial resistance) analysis.
#' @inheritSection lifecycle Stable lifecycle
#' @param x age, e.g. calculated with [age()]
#' @param split_at values to split `x` at, defaults to age groups 0-11, 12-24, 25-54, 55-74 and 75+. See Details.
#' @param na.rm a [logical] to indicate whether missing values should be removed
#' @details To split ages, the input for the `split_at` argument can be:
#' 
#' * A numeric vector. A value of e.g. `c(10, 20)` will split `x` on 0-9, 10-19 and 20+. A value of only `50` will split `x` on 0-49 and 50+.
#'   The default is to split on young children (0-11), youth (12-24), young adults (25-54), middle-aged adults (55-74) and elderly (75+).
#' * A character:
#'   - `"children"` or `"kids"`, equivalent of: `c(0, 1, 2, 4, 6, 13, 18)`. This will split on 0, 1, 2-3, 4-5, 6-12, 13-17 and 18+.
#'   - `"elderly"` or `"seniors"`, equivalent of: `c(65, 75, 85)`. This will split on 0-64, 65-74, 75-84, 85+.
#'   - `"fives"`, equivalent of: `1:20 * 5`. This will split on 0-4, 5-9, ..., 95-99, 100+.
#'   - `"tens"`, equivalent of: `1:10 * 10`. This will split on 0-9, 10-19, ..., 90-99, 100+.
#' @return Ordered [factor]
#' @seealso To determine ages, based on one or more reference dates, use the [age()] function.
#' @export
#' @inheritSection AMR Read more on our website!
#' @examples
#' ages <- c(3, 8, 16, 54, 31, 76, 101, 43, 21)
#'
#' # split into 0-49 and 50+
#' age_groups(ages, 50)
#'
#' # split into 0-19, 20-49 and 50+
#' age_groups(ages, c(20, 50))
#'
#' # split into groups of ten years
#' age_groups(ages, 1:10 * 10)
#' age_groups(ages, split_at = "tens")
#'
#' # split into groups of five years
#' age_groups(ages, 1:20 * 5)
#' age_groups(ages, split_at = "fives")
#'
#' # split specifically for children
#' age_groups(ages, c(1, 2, 4, 6, 13, 17))
#' age_groups(ages, "children")
#'
#' \donttest{
#' # resistance of ciprofloxacin per age group
#' if (require("dplyr")) {
#'   example_isolates %>%
#'     filter_first_isolate() %>%
#'     filter(mo == as.mo("E. coli")) %>%
#'     group_by(age_group = age_groups(age)) %>%
#'     select(age_group, CIP) %>%
#'     ggplot_rsi(x = "age_group", minimum = 0)
#' }
#' }
age_groups <- function(x, split_at = c(12, 25, 55, 75), na.rm = FALSE) {
  meet_criteria(x, allow_class = c("numeric", "integer"))
  meet_criteria(split_at, allow_class = c("numeric", "integer", "character"))
  meet_criteria(na.rm, allow_class = "logical", has_length = 1) 
  
  if (any(x < 0, na.rm = TRUE)) {
    x[x < 0] <- NA
    warning_("NAs introduced for ages below 0.", call = TRUE)
  }
  if (is.character(split_at)) {
    split_at <- split_at[1L]
    if (split_at %like% "^(child|kid|junior)") {
      split_at <- c(0, 1, 2, 4, 6, 13, 18)
    } else if (split_at %like% "^(elder|senior)") {
      split_at <-  c(65, 75, 85)
    } else if (split_at %like% "^five") {
      split_at <- 1:20 * 5
    } else if (split_at %like% "^ten") {
      split_at <- 1:10 * 10
    }
  }
  split_at <- sort(unique(as.integer(split_at)))
  if (!split_at[1] == 0) {
    # add base number 0
    split_at <- c(0, split_at)
  }
  split_at <- split_at[!is.na(split_at)]
  stop_if(length(split_at) == 1, "invalid value for `split_at`") # only 0 is available
  
  # turn input values to 'split_at' indices
  y <- x
  lbls <- split_at
  for (i in seq_len(length(split_at))) {
    y[x >= split_at[i]] <- i
    # create labels
    lbls[i - 1] <- paste0(unique(c(split_at[i - 1], split_at[i] - 1)), collapse = "-")
  }
  
  # last category
  lbls[length(lbls)] <- paste0(split_at[length(split_at)], "+")
  
  agegroups <- factor(lbls[y], levels = lbls, ordered = TRUE)
  
  if (isTRUE(na.rm)) {
    agegroups <- agegroups[!is.na(agegroups)]
  }
  
  agegroups
}
```
