## Supplementary material for "AMR - An R Package for Working with Antimicrobial Resistance Data": Source package: atc_online.R

```
# ==================================================================== #
# TITLE                                                                #
# Antimicrobial Resistance (AMR) Analysis for R                        #
#                                                                      #
# SOURCE                                                               #
# https://github.com/msberends/AMR                                     #
#                                                                      #
# LICENCE                                                              #
# (c) 2018-2021 Berends MS, Luz CF et al.                              #
# Developed at the University of Groningen, the Netherlands, in        #
# collaboration with non-profit organisations Certe Medical            #
# Diagnostics & Advice, and University Medical Center Groningen.       # 
#                                                                      #
# This R package is free software; you can freely use and distribute   #
# it for both personal and commercial purposes under the terms of the  #
# GNU General Public License version 2.0 (GNU GPL-2), as published by  #
# the Free Software Foundation.                                        #
# We created this package for both routine data analysis and academic  #
# research and it was publicly released in the hope that it will be    #
# useful, but it comes WITHOUT ANY WARRANTY OR LIABILITY.              #
#                                                                      #
# Visit our website for the full manual and a complete tutorial about  #
# how to conduct AMR analysis: https://msberends.github.io/AMR/        #
# ==================================================================== #

#' Get ATC properties from WHOCC website
#'
#' Gets data from the WHO to determine properties of an ATC (e.g. an antibiotic), such as the name, defined daily dose (DDD) or standard unit.
#' @inheritSection lifecycle Stable lifecycle
#' @param atc_code a character or character vector with ATC code(s) of antibiotic(s)
#' @param property property of an ATC code. Valid values are `"ATC"`, `"Name"`, `"DDD"`, `"U"` (`"unit"`), `"Adm.R"`, `"Note"` and `groups`. For this last option, all hierarchical groups of an ATC code will be returned, see Examples.
#' @param administration type of administration when using `property = "Adm.R"`, see Details
#' @param url url of website of the WHOCC. The sign `%s` can be used as a placeholder for ATC codes.
#' @param url_vet url of website of the WHOCC for veterinary medicine. The sign `%s` can be used as a placeholder for ATC_vet codes (that all start with "Q").
#' @param ... arguments to pass on to `atc_property`
#' @details
#' Options for argument `administration`:
#' 
#' - `"Implant"` = Implant
#' - `"Inhal"` = Inhalation
#' - `"Instill"` = Instillation
#' - `"N"` = nasal
#' - `"O"` = oral
#' - `"P"` = parenteral
#' - `"R"` = rectal
#' - `"SL"` = sublingual/buccal
#' - `"TD"` = transdermal
#' - `"V"` = vaginal
#'
#' Abbreviations of return values when using `property = "U"` (unit):
#' 
#' - `"g"` = gram
#' - `"mg"` = milligram
#' - `"mcg"`` = microgram
#' - `"U"` = unit
#' - `"TU"` = thousand units
#' - `"MU"` = million units
#' - `"mmol"` = millimole
#' - `"ml"` = milliliter (e.g. eyedrops)
#' 
#' **N.B. This function requires an internet connection and only works if the following packages are installed: `curl`, `rvest`, `xml2`.**
#' @export
#' @rdname atc_online
#' @inheritSection AMR Read more on our website!
#' @source <https://www.whocc.no/atc_ddd_alterations__cumulative/ddd_alterations/abbrevations/>
#' @examples
#' \donttest{
#' # oral DDD (Defined Daily Dose) of amoxicillin
#' atc_online_property("J01CA04", "DDD", "O")
#' 
#' # parenteral DDD (Defined Daily Dose) of amoxicillin
#' atc_online_property("J01CA04", "DDD", "P")
#'
#' atc_online_property("J01CA04", property = "groups") # search hierarchical groups of amoxicillin
#' }
atc_online_property <- function(atc_code,
                                property,
                                administration = "O",
                                url = "https://www.whocc.no/atc_ddd_index/?code=%s&showdescription=no",
                                url_vet = "https://www.whocc.no/atcvet/atcvet_index/?code=%s&showdescription=no") {
  meet_criteria(atc_code, allow_class = "character")
  meet_criteria(property, allow_class = "character", has_length = 1, is_in = c("ATC", "Name", "DDD", "U", "Adm.R", "Note", "groups"), ignore.case = TRUE)
  meet_criteria(administration, allow_class = "character", has_length = 1)
  meet_criteria(url, allow_class = "character", has_length = 1, looks_like = "https?://")
  meet_criteria(url_vet, allow_class = "character", has_length = 1, looks_like = "https?://")
  
  has_internet <- import_fn("has_internet", "curl")
  html_attr <- import_fn("html_attr", "rvest")
  html_children <- import_fn("html_children", "rvest")
  html_node <- import_fn("html_node", "rvest")
  html_nodes <- import_fn("html_nodes", "rvest")
  html_table <- import_fn("html_table", "rvest")
  html_text <- import_fn("html_text", "rvest")
  read_html <- import_fn("read_html", "xml2")
  
  check_dataset_integrity()
  
  if (!all(atc_code %in% antibiotics)) {
    atc_code <- as.character(ab_atc(atc_code))
  }
  
  if (!has_internet()) {
    message_("There appears to be no internet connection, returning NA.",
             add_fn = font_red,
             as_note = FALSE)
    return(rep(NA, length(atc_code)))
  }
  
  # also allow unit as property
  if (property %like% "unit") {
    property <- "U"
  }
  
  property <- tolower(property)
  if (property == "ddd") {
    returnvalue <- rep(NA_real_, length(atc_code))
  } else if (property == "groups") {
    returnvalue <- list()
  } else {
    returnvalue <- rep(NA_character_, length(atc_code))
  }
  
  progress <- progress_ticker(n = length(atc_code), 3)
  on.exit(close(progress))
  
  for (i in seq_len(length(atc_code))) {
    
    progress$tick()

    if (atc_code[i] %like% "^Q") {
      # veterinary drugs, ATC_vet codes start with a "Q"
      atc_url <- url_vet
    } else {
      atc_url <- url
    }
    atc_url <- sub("%s", atc_code[i], atc_url, fixed = TRUE)
    
    if (property == "groups") {
      tbl <- read_html(atc_url) %pm>%
        html_node("#content") %pm>%
        html_children() %pm>%
        html_node("a")
      
      # get URLS of items
      hrefs <- tbl %pm>% html_attr("href")
      # get text of items
      texts <- tbl %pm>% html_text()
      # select only text items where URL like "code="
      texts <- texts[grepl("?code=", tolower(hrefs), fixed = TRUE)]
      # last one is antibiotics, skip it
      texts <- texts[seq_len(length(texts)) - 1]
      returnvalue <- c(list(texts), returnvalue)
      
    } else {
      tbl <- read_html(atc_url) %pm>%
        html_nodes("table") %pm>%
        html_table(header = TRUE) %pm>%
        as.data.frame(stringsAsFactors = FALSE)
      
      # case insensitive column names
      colnames(tbl) <- gsub("^atc.*", "atc", tolower(colnames(tbl)))
      
      if (length(tbl) == 0) {
        warning_("ATC not found: ", atc_code[i], ". Please check ", atc_url, ".", call = FALSE)
        returnvalue[i] <- NA
        next
      }
      
      if (property %in% c("atc", "name")) {
        # ATC and name are only in first row
        returnvalue[i] <- tbl[1, property]
      } else {
        if (!"adm.r" %in% colnames(tbl) | is.na(tbl[1, "adm.r"])) {
          returnvalue[i] <- NA
          next
        } else {
          for (j in seq_len(nrow(tbl))) {
            if (tbl[j, "adm.r"] == administration) {
              returnvalue[i] <- tbl[j, property]
            }
          }
        }
      }
    }
  }
  
  if (property == "groups" & length(returnvalue) == 1) {
    returnvalue <- returnvalue[[1]]
  }
  
  returnvalue
}

#' @rdname atc_online
#' @export
atc_online_groups <- function(atc_code, ...) {
  meet_criteria(atc_code, allow_class = "character")
  atc_online_property(atc_code = atc_code, property = "groups", ...)
}

#' @rdname atc_online
#' @export
atc_online_ddd <- function(atc_code, ...) {
  meet_criteria(atc_code, allow_class = "character")
  atc_online_property(atc_code = atc_code, property = "ddd", ...)
}
```
