## Supplementary material for "AMR - An R Package for Working with Antimicrobial Resistance Data": Source package: availability.R

```
# ==================================================================== #
# TITLE                                                                #
# Antimicrobial Resistance (AMR) Analysis for R                        #
#                                                                      #
# SOURCE                                                               #
# https://github.com/msberends/AMR                                     #
#                                                                      #
# LICENCE                                                              #
# (c) 2018-2021 Berends MS, Luz CF et al.                              #
# Developed at the University of Groningen, the Netherlands, in        #
# collaboration with non-profit organisations Certe Medical            #
# Diagnostics & Advice, and University Medical Center Groningen.       # 
#                                                                      #
# This R package is free software; you can freely use and distribute   #
# it for both personal and commercial purposes under the terms of the  #
# GNU General Public License version 2.0 (GNU GPL-2), as published by  #
# the Free Software Foundation.                                        #
# We created this package for both routine data analysis and academic  #
# research and it was publicly released in the hope that it will be    #
# useful, but it comes WITHOUT ANY WARRANTY OR LIABILITY.              #
#                                                                      #
# Visit our website for the full manual and a complete tutorial about  #
# how to conduct AMR analysis: https://msberends.github.io/AMR/        #
# ==================================================================== #

#' Check availability of columns
#'
#' Easy check for data availability of all columns in a data set. This makes it easy to get an idea of which antimicrobial combinations can be used for calculation with e.g. [susceptibility()] and [resistance()].
#' @inheritSection lifecycle Stable lifecycle
#' @param tbl a [data.frame] or [list]
#' @param width number of characters to present the visual availability, defaults to filling the width of the console
#' @details The function returns a [data.frame] with columns `"resistant"` and `"visual_resistance"`. The values in that columns are calculated with [resistance()].
#' @return [data.frame] with column names of `tbl` as row names
#' @inheritSection AMR Read more on our website!
#' @export
#' @examples
#' availability(example_isolates)
#'
#' if (require("dplyr")) {
#'   example_isolates %>%
#'     filter(mo == as.mo("E. coli")) %>%
#'     select_if(is.rsi) %>%
#'     availability()
#' }
availability <- function(tbl, width = NULL) {
  meet_criteria(tbl, allow_class = "data.frame")
  meet_criteria(width, allow_class = "numeric", allow_NULL = TRUE)
  
  x <- vapply(FUN.VALUE = double(1), tbl, function(x) {
    1 - sum(is.na(x)) / length(x) 
  })
  n <- vapply(FUN.VALUE = double(1), tbl, function(x) length(x[!is.na(x)]))
  R <- vapply(FUN.VALUE = double(1), tbl, function(x) ifelse(is.rsi(x), resistance(x, minimum = 0), NA_real_))
  R_print <- character(length(R))
  R_print[!is.na(R)] <- percentage(R[!is.na(R)])
  R_print[is.na(R)] <- ""
  
  if (is.null(width)) {
    width <- options()$width -
      (max(nchar(colnames(tbl))) +
         # count col
         8 +
         # available % column
         10 +
         # resistant % column
         10 +
         # extra margin
         5)
    width <- width / 2
  }
  
  if (length(R[is.na(R)]) == ncol(tbl)) {
    width <- width * 2 + 10
  }
  
  x_chars_R <- strrep("#", round(width * R, digits = 2))
  x_chars_SI <- strrep("-", width - nchar(x_chars_R))
  vis_resistance <- paste0("|", x_chars_R, x_chars_SI, "|")
  vis_resistance[is.na(R)] <- ""
  
  x_chars <- strrep("#", round(x, digits = 2) / (1 / width))
  x_chars_empty <- strrep("-", width - nchar(x_chars))
  
  df <- data.frame(count = n,
                   available = percentage(x),
                   visual_availabilty = paste0("|", x_chars, x_chars_empty, "|"),
                   resistant = R_print,
                   visual_resistance = vis_resistance,
                   stringsAsFactors = FALSE)
  if (length(R[is.na(R)]) == ncol(tbl)) {
    df[, 1:3]
  } else {
    df
  }
}
```
