## Supplementary material for "AMR - An R Package for Working with Antimicrobial Resistance Data": Source package: bug_drug_combinations.R

```
# ==================================================================== #
# TITLE                                                                #
# Antimicrobial Resistance (AMR) Analysis for R                        #
#                                                                      #
# SOURCE                                                               #
# https://github.com/msberends/AMR                                     #
#                                                                      #
# LICENCE                                                              #
# (c) 2018-2021 Berends MS, Luz CF et al.                              #
# Developed at the University of Groningen, the Netherlands, in        #
# collaboration with non-profit organisations Certe Medical            #
# Diagnostics & Advice, and University Medical Center Groningen.       # 
#                                                                      #
# This R package is free software; you can freely use and distribute   #
# it for both personal and commercial purposes under the terms of the  #
# GNU General Public License version 2.0 (GNU GPL-2), as published by  #
# the Free Software Foundation.                                        #
# We created this package for both routine data analysis and academic  #
# research and it was publicly released in the hope that it will be    #
# useful, but it comes WITHOUT ANY WARRANTY OR LIABILITY.              #
#                                                                      #
# Visit our website for the full manual and a complete tutorial about  #
# how to conduct AMR analysis: https://msberends.github.io/AMR/        #
# ==================================================================== #

#' Determine bug-drug combinations
#' 
#' Determine antimicrobial resistance (AMR) of all bug-drug combinations in your data set where at least 30 (default) isolates are available per species. Use [format()] on the result to prettify it to a publicable/printable format, see Examples.
#' @inheritSection lifecycle Stable lifecycle
#' @inheritParams eucast_rules
#' @param combine_IR logical to indicate whether values R and I should be summed
#' @param add_ab_group logical to indicate where the group of the antimicrobials must be included as a first column
#' @param remove_intrinsic_resistant logical to indicate that rows and columns with 100% resistance for all tested antimicrobials must be removed from the table
#' @param FUN function to call on the `mo` column to transform the microorganism IDs, defaults to [mo_shortname()] 
#' @param translate_ab character of length 1 containing column names of the [antibiotics] data set
#' @param ... arguments passed on to `FUN`
#' @inheritParams rsi_df
#' @inheritParams base::formatC
#' @details The function [format()] calculates the resistance per bug-drug combination. Use `combine_IR = FALSE` (default) to test R vs. S+I and `combine_IR = TRUE` to test R+I vs. S. 
#' @export
#' @rdname bug_drug_combinations
#' @return The function [bug_drug_combinations()] returns a [data.frame] with columns "mo", "ab", "S", "I", "R" and "total".
#' @source \strong{M39 Analysis and Presentation of Cumulative Antimicrobial Susceptibility Test Data, 4th Edition}, 2014, *Clinical and Laboratory Standards Institute (CLSI)*. <https://clsi.org/standards/products/microbiology/documents/m39/>.
#' @inheritSection AMR Read more on our website!
#' @examples 
#' \donttest{
#' x <- bug_drug_combinations(example_isolates)
#' x
#' format(x, translate_ab = "name (atc)")
#' 
#' # Use FUN to change to transformation of microorganism codes
#' bug_drug_combinations(example_isolates, 
#'                       FUN = mo_gramstain)
#'                            
#' bug_drug_combinations(example_isolates,
#'                       FUN = function(x) ifelse(x == as.mo("E. coli"),
#'                                                "E. coli",
#'                                                "Others"))
#' }
bug_drug_combinations <- function(x, 
                                  col_mo = NULL, 
                                  FUN = mo_shortname,
                                  ...) {
  meet_criteria(x, allow_class = "data.frame", contains_column_class = "rsi")
  meet_criteria(col_mo, allow_class = "character", is_in = colnames(x), has_length = 1, allow_NULL = TRUE)
  meet_criteria(FUN, allow_class = "function", has_length = 1)

  # try to find columns based on type
  # -- mo
  if (is.null(col_mo)) {
    col_mo <- search_type_in_df(x = x, type = "mo")
  }
  stop_if(is.null(col_mo), "`col_mo` must be set")
  
  x_class <- class(x)
  x <- as.data.frame(x, stringsAsFactors = FALSE)
  x[, col_mo] <- FUN(x[, col_mo, drop = TRUE], ...)
  x <- x[, c(col_mo, names(which(vapply(FUN.VALUE = logical(1), x, is.rsi)))), drop = FALSE]
  
  unique_mo <- sort(unique(x[, col_mo, drop = TRUE]))
  
  out <- data.frame(mo = character(0),
                    ab = character(0),
                    S = integer(0),
                    I = integer(0),
                    R = integer(0),
                    total = integer(0),
                    stringsAsFactors = FALSE)
  
  for (i in seq_len(length(unique_mo))) {
    # filter on MO group and only select R/SI columns
    x_mo_filter <- x[which(x[, col_mo, drop = TRUE] == unique_mo[i]), names(which(vapply(FUN.VALUE = logical(1), x, is.rsi))), drop = FALSE]
    # turn and merge everything
    pivot <- lapply(x_mo_filter, function(x) {
      m <- as.matrix(table(x))
      data.frame(S = m["S", ], I = m["I", ], R = m["R", ], stringsAsFactors = FALSE)
    })
    merged <- do.call(rbind, pivot)
    out_group <- data.frame(mo = unique_mo[i],
                            ab = rownames(merged),
                            S = merged$S,
                            I = merged$I,
                            R = merged$R,
                            total = merged$S + merged$I + merged$R,
                            stringsAsFactors = FALSE)
    out <- rbind(out, out_group, stringsAsFactors = FALSE)
  }
  
  set_clean_class(out,
                  new_class = c("bug_drug_combinations", x_class))
}

#' @method format bug_drug_combinations
#' @export
#' @rdname bug_drug_combinations
format.bug_drug_combinations <- function(x,
                                         translate_ab = "name (ab, atc)",
                                         language = get_locale(),
                                         minimum = 30,
                                         combine_SI = TRUE,
                                         combine_IR = FALSE,
                                         add_ab_group = TRUE,
                                         remove_intrinsic_resistant = FALSE,
                                         decimal.mark = getOption("OutDec"),
                                         big.mark = ifelse(decimal.mark == ",", ".", ","),
                                         ...) {
  meet_criteria(x, allow_class = "data.frame")
  meet_criteria(translate_ab, allow_class = c("character", "logical"), has_length = 1, allow_NA = TRUE)
  meet_criteria(language, has_length = 1, is_in = c(LANGUAGES_SUPPORTED, ""), allow_NULL = TRUE, allow_NA = TRUE)
  meet_criteria(minimum, allow_class = c("numeric", "integer"), has_length = 1)
  meet_criteria(combine_SI, allow_class = "logical", has_length = 1)
  meet_criteria(combine_IR, allow_class = "logical", has_length = 1)
  meet_criteria(add_ab_group, allow_class = "logical", has_length = 1)
  meet_criteria(remove_intrinsic_resistant, allow_class = "logical", has_length = 1)
  meet_criteria(decimal.mark, allow_class = "character", has_length = 1)
  meet_criteria(big.mark, allow_class = "character", has_length = 1)
  
  x <- as.data.frame(x, stringsAsFactors = FALSE)
  x <- subset(x, total >= minimum)
  
  if (remove_intrinsic_resistant == TRUE) {
    x <- subset(x, R != total)
  }
  if (combine_SI == TRUE | combine_IR == FALSE) {
    x$isolates <- x$R
  } else {
    x$isolates <- x$R + x$I
  }
  
  give_ab_name <- function(ab, format, language) {
    format <- tolower(format)
    ab_txt <- rep(format, length(ab))
    for (i in seq_len(length(ab_txt))) {
      ab_txt[i] <- gsub("ab", as.character(as.ab(ab[i])), ab_txt[i])
      ab_txt[i] <- gsub("cid", ab_cid(ab[i]), ab_txt[i])
      ab_txt[i] <- gsub("group", ab_group(ab[i], language = language), ab_txt[i])
      ab_txt[i] <- gsub("atc_group1", ab_atc_group1(ab[i], language = language), ab_txt[i])
      ab_txt[i] <- gsub("atc_group2", ab_atc_group2(ab[i], language = language), ab_txt[i])
      ab_txt[i] <- gsub("atc", ab_atc(ab[i]), ab_txt[i])
      ab_txt[i] <- gsub("name", ab_name(ab[i], language = language), ab_txt[i])
      ab_txt[i]
    }
    ab_txt
  }
  
  remove_NAs <- function(.data) {
    cols <- colnames(.data)
    .data <- as.data.frame(lapply(.data, function(x) ifelse(is.na(x), "", x)),
                           stringsAsFactors = FALSE)
    colnames(.data) <- cols
    .data
  }
  
  create_var <- function(.data, ...) {
    dots <- list(...)
    for (i in seq_len(length(dots))) {
      .data[, names(dots)[i]] <- dots[[i]]
    }
    .data
  }
  
  y <- x %pm>%
    create_var(ab = as.ab(x$ab),
               ab_txt = give_ab_name(ab = x$ab, format = translate_ab, language = language)) %pm>%
    pm_group_by(ab, ab_txt, mo) %pm>% 
    pm_summarise(isolates = sum(isolates, na.rm = TRUE),
              total = sum(total, na.rm = TRUE)) %pm>% 
    pm_ungroup()
  
  y <- y %pm>% 
    create_var(txt = paste0(percentage(y$isolates / y$total, decimal.mark = decimal.mark, big.mark = big.mark), 
                            " (", trimws(format(y$isolates, big.mark = big.mark)), "/",
                            trimws(format(y$total, big.mark = big.mark)), ")")) %pm>% 
    pm_select(ab, ab_txt, mo, txt) %pm>%
    pm_arrange(mo)
  
  # replace tidyr::pivot_wider() from here
  for (i in unique(y$mo)) {
    mo_group <- y[which(y$mo == i), c("ab", "txt")]
    colnames(mo_group) <- c("ab", i)
    rownames(mo_group) <- NULL
    y <- y %pm>% 
      pm_left_join(mo_group, by = "ab")
  }
  y <- y %pm>% 
    pm_distinct(ab, .keep_all = TRUE) %pm>% 
    pm_select(-mo, -txt) %pm>% 
    # replace tidyr::pivot_wider() until here
    remove_NAs()

  select_ab_vars <- function(.data) {
    .data[, c("ab_group", "ab_txt", colnames(.data)[!colnames(.data) %in% c("ab_group", "ab_txt", "ab")])]
  }
  
  y <- y %pm>% 
    create_var(ab_group = ab_group(y$ab, language = language)) %pm>% 
    select_ab_vars() %pm>% 
    pm_arrange(ab_group, ab_txt)
  y <- y %pm>% 
    create_var(ab_group = ifelse(y$ab_group != pm_lag(y$ab_group) | is.na(pm_lag(y$ab_group)), y$ab_group, ""))
  
  if (add_ab_group == FALSE) {
    y <- y %pm>% 
      pm_select(-ab_group) %pm>%
      pm_rename("Drug" = ab_txt)
    colnames(y)[1] <- translate_AMR(colnames(y)[1], language, only_unknown = FALSE)
  } else {
    y <- y %pm>% 
      pm_rename("Group" = ab_group,
                "Drug" = ab_txt)
  }
  
  if (!is.null(language)) {
    colnames(y) <- translate_AMR(colnames(y), language, only_unknown = FALSE)
  }
  
  if (remove_intrinsic_resistant == TRUE) {
    y <- y[, !vapply(FUN.VALUE = logical(1), y, function(col) all(col %like% "100", na.rm = TRUE) & !any(is.na(col))), drop = FALSE]
  }
  
  rownames(y) <- NULL
  y
}

#' @method print bug_drug_combinations
#' @export
print.bug_drug_combinations <- function(x, ...) {
  x_class <- class(x)
  print(set_clean_class(x, 
                        new_class = x_class[x_class != "bug_drug_combinations"]),
        ...)
  message_("Use 'format()' on this result to get a publishable/printable format.", as_note = FALSE)
}
```
