## Supplementary material for "AMR - An R Package for Working with Antimicrobial Resistance Data": Source package: catalogue_of_life.R

```
# ==================================================================== #
# TITLE                                                                #
# Antimicrobial Resistance (AMR) Analysis for R                        #
#                                                                      #
# SOURCE                                                               #
# https://github.com/msberends/AMR                                     #
#                                                                      #
# LICENCE                                                              #
# (c) 2018-2021 Berends MS, Luz CF et al.                              #
# Developed at the University of Groningen, the Netherlands, in        #
# collaboration with non-profit organisations Certe Medical            #
# Diagnostics & Advice, and University Medical Center Groningen.       # 
#                                                                      #
# This R package is free software; you can freely use and distribute   #
# it for both personal and commercial purposes under the terms of the  #
# GNU General Public License version 2.0 (GNU GPL-2), as published by  #
# the Free Software Foundation.                                        #
# We created this package for both routine data analysis and academic  #
# research and it was publicly released in the hope that it will be    #
# useful, but it comes WITHOUT ANY WARRANTY OR LIABILITY.              #
#                                                                      #
# Visit our website for the full manual and a complete tutorial about  #
# how to conduct AMR analysis: https://msberends.github.io/AMR/        #
# ==================================================================== #

format_included_data_number <- function(data) {
  if (is.data.frame(data)) {
    n <- nrow(data)
  } else {
    n <- length(unique(data))
  }
  if (n > 10000) {
    rounder <- -3 # round on thousands
  } else if (n > 1000) {
    rounder <- -2 # round on hundreds
  } else {
    rounder <- -1 # round on tens
  }
  paste0("~", format(round(n, rounder), decimal.mark = ".", big.mark = ","))
}

#' The Catalogue of Life
#'
#' This package contains the complete taxonomic tree of almost all microorganisms from the authoritative and comprehensive Catalogue of Life.
#' @section Catalogue of Life:
#' \if{html}{\figure{logo_col.png}{options: height=40px style=margin-bottom:5px} \cr}
#' This package contains the complete taxonomic tree of almost all microorganisms (~70,000 species) from the authoritative and comprehensive Catalogue of Life (CoL, <http://www.catalogueoflife.org>). The CoL is the most comprehensive and authoritative global index of species currently available. Nonetheless, we supplemented the CoL data with data from the List of Prokaryotic names with Standing in Nomenclature (LPSN, [lpsn.dsmz.de](https://lpsn.dsmz.de)). This supplementation is needed until the [CoL+ project](https://github.com/CatalogueOfLife/general) is finished, which we await.
#'
#' [Click here][catalogue_of_life] for more information about the included taxa. Check which versions of the CoL and LSPN were included in this package with [catalogue_of_life_version()].
#' @section Included taxa:
#' Included are:
#' - All `r format_included_data_number(microorganisms[which(microorganisms$kingdom %in% c("Archeae", "Bacteria", "Chromista", "Protozoa")), ])` (sub)species from the kingdoms of Archaea, Bacteria, Chromista and Protozoa
#' - All `r format_included_data_number(microorganisms[which(microorganisms$kingdom == "Fungi" & microorganisms$order %in% c("Eurotiales", "Microascales", "Mucorales", "Onygenales", "Pneumocystales", "Saccharomycetales", "Schizosaccharomycetales", "Tremellales")), ])` (sub)species from these orders of the kingdom of Fungi: Eurotiales, Microascales, Mucorales, Onygenales, Pneumocystales, Saccharomycetales, Schizosaccharomycetales and Tremellales, as well as `r format_included_data_number(microorganisms[which(microorganisms$kingdom == "Fungi" & !microorganisms$order %in% c("Eurotiales", "Microascales", "Mucorales", "Onygenales", "Pneumocystales", "Saccharomycetales", "Schizosaccharomycetales", "Tremellales")), ])` other fungal (sub)species. The kingdom of Fungi is a very large taxon with almost 300,000 different (sub)species, of which most are not microbial (but rather macroscopic, like mushrooms). Because of this, not all fungi fit the scope of this package and including everything would tremendously slow down our algorithms too. By only including the aforementioned taxonomic orders, the most relevant fungi are covered (such as all species of *Aspergillus*, *Candida*, *Cryptococcus*, *Histplasma*, *Pneumocystis*, *Saccharomyces* and *Trichophyton*).
#' - All `r format_included_data_number(microorganisms[which(microorganisms$kingdom == "Animalia"), ])` (sub)species from `r format_included_data_number(microorganisms[which(microorganisms$kingdom == "Animalia"), "genus"])` other relevant genera from the kingdom of Animalia (such as *Strongyloides* and *Taenia*)
#' - All `r format_included_data_number(microorganisms.old)` previously accepted names of all included (sub)species (these were taxonomically renamed)
#' - The complete taxonomic tree of all included (sub)species: from kingdom to subspecies
#' - The responsible author(s) and year of scientific publication
#'
#' The Catalogue of Life (<http://www.catalogueoflife.org>) is the most comprehensive and authoritative global index of species currently available. It holds essential information on the names, relationships and distributions of over 1.9 million species. The Catalogue of Life is used to support the major biodiversity and conservation information services such as the Global Biodiversity Information Facility (GBIF), Encyclopedia of Life (EoL) and the International Union for Conservation of Nature Red List. It is recognised by the Convention on Biological Diversity as a significant component of the Global Taxonomy Initiative and a contribution to Target 1 of the Global Strategy for Plant Conservation.
#'
#' The syntax used to transform the original data to a cleansed \R format, can be found here: <https://github.com/msberends/AMR/blob/master/data-raw/reproduction_of_microorganisms.R>.
#' @inheritSection AMR Read more on our website!
#' @name catalogue_of_life
#' @rdname catalogue_of_life
#' @seealso Data set [microorganisms] for the actual data. \cr
#' Function [as.mo()] to use the data for intelligent determination of microorganisms.
#' @examples
#' # Get version info of included data set
#' catalogue_of_life_version()
#'
#'
#' # Get a note when a species was renamed
#' mo_shortname("Chlamydophila psittaci")
#' # Note: 'Chlamydophila psittaci' (Everett et al., 1999) was renamed back to
#' #       'Chlamydia psittaci' (Page, 1968)
#' #> [1] "C. psittaci"
#'
#' # Get any property from the entire taxonomic tree for all included species
#' mo_class("E. coli")
#' #> [1] "Gammaproteobacteria"
#'
#' mo_family("E. coli")
#' #> [1] "Enterobacteriaceae"
#'
#' mo_gramstain("E. coli") # based on kingdom and phylum, see ?mo_gramstain
#' #> [1] "Gram-negative"
#'
#' mo_ref("E. coli")
#' #> [1] "Castellani et al., 1919"
#'
#' # Do not get mistaken - this package is about microorganisms
#' mo_kingdom("C. elegans")
#' #> [1] "Fungi"                 # Fungi?!
#' mo_name("C. elegans")
#' #> [1] "Cladosporium elegans"  # Because a microorganism was found
NULL

#' Version info of included Catalogue of Life
#'
#' This function returns information about the included data from the Catalogue of Life.
#' @seealso [microorganisms]
#' @details For DSMZ, see [microorganisms].
#' @return a [list], which prints in pretty format
#' @inheritSection catalogue_of_life Catalogue of Life
#' @inheritSection AMR Read more on our website!
#' @export
catalogue_of_life_version <- function() {
  
  check_dataset_integrity()
  
  # see the `catalogue_of_life` list in R/data.R
  lst <- list(catalogue_of_life =
                list(version = gsub("{year}", catalogue_of_life$year, catalogue_of_life$version, fixed = TRUE),
                     url = gsub("{year}", catalogue_of_life$year, catalogue_of_life$url_CoL, fixed = TRUE),
                     n = nrow(pm_filter(microorganisms, source == "CoL"))),
              deutsche_sammlung_von_mikroorganismen_und_zellkulturen =
                list(version = "Prokaryotic Nomenclature Up-to-Date from DSMZ",
                     url = catalogue_of_life$url_DSMZ,
                     yearmonth = catalogue_of_life$yearmonth_DSMZ,
                     n = nrow(pm_filter(microorganisms, source == "DSMZ"))),
              total_included =
                list(
                  n_total_species = nrow(microorganisms),
                  n_total_synonyms = nrow(microorganisms.old)))
  
  set_clean_class(lst,
                  new_class = c("catalogue_of_life_version", "list"))
}

#' @method print catalogue_of_life_version
#' @export
#' @noRd
print.catalogue_of_life_version <- function(x, ...) {
  lst <- x
  cat(paste0(font_bold("Included in this AMR package are:\n\n"),
             font_underline(lst$catalogue_of_life$version), "\n",
             "  Available at: ", lst$catalogue_of_life$url, "\n",
             "  Number of included species: ", format(lst$catalogue_of_life$n, big.mark = ","), "\n",
             font_underline(paste0(lst$deutsche_sammlung_von_mikroorganismen_und_zellkulturen$version, " (",
                                   lst$deutsche_sammlung_von_mikroorganismen_und_zellkulturen$yearmonth, ")")), "\n",
             "  Available at: ", lst$deutsche_sammlung_von_mikroorganismen_und_zellkulturen$url, "\n",
             "  Number of included species: ", format(lst$deutsche_sammlung_von_mikroorganismen_und_zellkulturen$n, big.mark = ","), "\n\n",
             "=> Total number of species included:  ", format(lst$total_included$n_total_species, big.mark = ","), "\n",
             "=> Total number of synonyms included: ", format(lst$total_included$n_total_synonyms, big.mark = ","), "\n\n",
             "See for more info ?microorganisms and ?catalogue_of_life.\n"))
}
```
