## Supplementary material for "AMR - An R Package for Working with Antimicrobial Resistance Data": Source package: count.R

```
# ==================================================================== #
# TITLE                                                                #
# Antimicrobial Resistance (AMR) Analysis for R                        #
#                                                                      #
# SOURCE                                                               #
# https://github.com/msberends/AMR                                     #
#                                                                      #
# LICENCE                                                              #
# (c) 2018-2021 Berends MS, Luz CF et al.                              #
# Developed at the University of Groningen, the Netherlands, in        #
# collaboration with non-profit organisations Certe Medical            #
# Diagnostics & Advice, and University Medical Center Groningen.       # 
#                                                                      #
# This R package is free software; you can freely use and distribute   #
# it for both personal and commercial purposes under the terms of the  #
# GNU General Public License version 2.0 (GNU GPL-2), as published by  #
# the Free Software Foundation.                                        #
# We created this package for both routine data analysis and academic  #
# research and it was publicly released in the hope that it will be    #
# useful, but it comes WITHOUT ANY WARRANTY OR LIABILITY.              #
#                                                                      #
# Visit our website for the full manual and a complete tutorial about  #
# how to conduct AMR analysis: https://msberends.github.io/AMR/        #
# ==================================================================== #

#' Count available isolates
#'
#' @description These functions can be used to count resistant/susceptible microbial isolates. All functions support quasiquotation with pipes, can be used in `summarise()` from the `dplyr` package and also support grouped variables, please see *Examples*.
#'
#' [count_resistant()] should be used to count resistant isolates, [count_susceptible()] should be used to count susceptible isolates.
#' @inheritSection lifecycle Stable lifecycle
#' @param ... one or more vectors (or columns) with antibiotic interpretations. They will be transformed internally with [as.rsi()] if needed.
#' @inheritParams proportion
#' @inheritSection as.rsi Interpretation of R and S/I
#' @details These functions are meant to count isolates. Use the [resistance()]/[susceptibility()] functions to calculate microbial resistance/susceptibility.
#' 
#' The function [count_resistant()] is equal to the function [count_R()]. The function [count_susceptible()] is equal to the function [count_SI()].
#'
#' The function [n_rsi()] is an alias of [count_all()]. They can be used to count all available isolates, i.e. where all input antibiotics have an available result (S, I or R). Their use is equal to `n_distinct()`. Their function is equal to `count_susceptible(...) + count_resistant(...)`.
#'
#' The function [count_df()] takes any variable from `data` that has an [`rsi`] class (created with [as.rsi()]) and counts the number of S's, I's and R's. It also supports grouped variables. The function [rsi_df()] works exactly like [count_df()], but adds the percentage of S, I and R.
#' @inheritSection proportion Combination therapy
#' @seealso [`proportion_*`][proportion] to calculate microbial resistance and susceptibility.
#' @return An [integer]
#' @rdname count
#' @name count
#' @export
#' @inheritSection AMR Read more on our website!
#' @examples
#' # example_isolates is a data set available in the AMR package.
#' ?example_isolates
#' 
#' count_resistant(example_isolates$AMX)   # counts "R"
#' count_susceptible(example_isolates$AMX) # counts "S" and "I"
#' count_all(example_isolates$AMX)         # counts "S", "I" and "R"
#'
#' # be more specific
#' count_S(example_isolates$AMX)
#' count_SI(example_isolates$AMX)
#' count_I(example_isolates$AMX)
#' count_IR(example_isolates$AMX)
#' count_R(example_isolates$AMX)
#'
#' # Count all available isolates
#' count_all(example_isolates$AMX)
#' n_rsi(example_isolates$AMX)
#'
#' # n_rsi() is an alias of count_all().
#' # Since it counts all available isolates, you can
#' # calculate back to count e.g. susceptible isolates.
#' # These results are the same:
#' count_susceptible(example_isolates$AMX)
#' susceptibility(example_isolates$AMX) * n_rsi(example_isolates$AMX)
#'
#' 
#' if (require("dplyr")) {
#'   example_isolates %>%
#'     group_by(hospital_id) %>%
#'     summarise(R  = count_R(CIP),
#'               I  = count_I(CIP),
#'               S  = count_S(CIP),
#'               n1 = count_all(CIP),  # the actual total; sum of all three
#'               n2 = n_rsi(CIP),      # same - analogous to n_distinct
#'               total = n())          # NOT the number of tested isolates!
#'  
#'   # Count co-resistance between amoxicillin/clav acid and gentamicin,
#'   # so we can see that combination therapy does a lot more than mono therapy.
#'   # Please mind that `susceptibility()` calculates percentages right away instead.
#'   example_isolates %>% count_susceptible(AMC) # 1433
#'   example_isolates %>% count_all(AMC)         # 1879
#'  
#'   example_isolates %>% count_susceptible(GEN) # 1399
#'   example_isolates %>% count_all(GEN)         # 1855
#'  
#'   example_isolates %>% count_susceptible(AMC, GEN) # 1764
#'   example_isolates %>% count_all(AMC, GEN)         # 1936
#'  
#'   # Get number of S+I vs. R immediately of selected columns
#'   example_isolates %>%
#'     select(AMX, CIP) %>%
#'     count_df(translate = FALSE)
#'  
#'   # It also supports grouping variables
#'   example_isolates %>%
#'     select(hospital_id, AMX, CIP) %>%
#'     group_by(hospital_id) %>%
#'     count_df(translate = FALSE)
#' }
count_resistant <- function(..., only_all_tested = FALSE) {
  rsi_calc(...,
           ab_result = "R",
           only_all_tested = only_all_tested,
           only_count = TRUE)
}

#' @rdname count
#' @export
count_susceptible <- function(..., only_all_tested = FALSE) {
  rsi_calc(...,
           ab_result = c("S", "I"),
           only_all_tested = only_all_tested,
           only_count = TRUE)
}

#' @rdname count
#' @export
count_R <- function(..., only_all_tested = FALSE) {
  rsi_calc(...,
           ab_result = "R",
           only_all_tested = only_all_tested,
           only_count = TRUE)
}

#' @rdname count
#' @export
count_IR <- function(..., only_all_tested = FALSE) {
  if (message_not_thrown_before("count_IR")) {
    warning_("Using count_IR() is discouraged; use count_resistant() instead to not consider \"I\" being resistant.", call = FALSE)
    remember_thrown_message("count_IR")
  }
  rsi_calc(...,
           ab_result = c("I", "R"),
           only_all_tested = only_all_tested,
           only_count = TRUE)
}

#' @rdname count
#' @export
count_I <- function(..., only_all_tested = FALSE) {
  rsi_calc(...,
           ab_result = "I",
           only_all_tested = only_all_tested,
           only_count = TRUE)
}

#' @rdname count
#' @export
count_SI <- function(..., only_all_tested = FALSE) {
  rsi_calc(...,
           ab_result = c("S", "I"),
           only_all_tested = only_all_tested,
           only_count = TRUE)
}

#' @rdname count
#' @export
count_S <- function(..., only_all_tested = FALSE) {
  if (message_not_thrown_before("count_S")) {
    warning_("Using count_S() is discouraged; use count_susceptible() instead to also consider \"I\" being susceptible.", call = FALSE)
    remember_thrown_message("count_S")
  }
  rsi_calc(...,
           ab_result = "S",
           only_all_tested = only_all_tested,
           only_count = TRUE)
}

#' @rdname count
#' @export
count_all <- function(..., only_all_tested = FALSE) {
  rsi_calc(...,
           ab_result = c("S", "I", "R"),
           only_all_tested = only_all_tested,
           only_count = TRUE)
}

#' @rdname count
#' @export
n_rsi <- count_all

#' @rdname count
#' @export
count_df <- function(data,
                     translate_ab = "name",
                     language = get_locale(),
                     combine_SI = TRUE,
                     combine_IR = FALSE) {
  rsi_calc_df(type = "count",
              data = data,
              translate_ab = translate_ab,
              language = language,
              combine_SI = combine_SI,
              combine_IR = combine_IR,
              combine_SI_missing = missing(combine_SI))
}
```
