## Supplementary material for "AMR - An R Package for Working with Antimicrobial Resistance Data": Source package: data.R

```
# ==================================================================== #
# TITLE                                                                #
# Antimicrobial Resistance (AMR) Analysis for R                        #
#                                                                      #
# SOURCE                                                               #
# https://github.com/msberends/AMR                                     #
#                                                                      #
# LICENCE                                                              #
# (c) 2018-2021 Berends MS, Luz CF et al.                              #
# Developed at the University of Groningen, the Netherlands, in        #
# collaboration with non-profit organisations Certe Medical            #
# Diagnostics & Advice, and University Medical Center Groningen.       # 
#                                                                      #
# This R package is free software; you can freely use and distribute   #
# it for both personal and commercial purposes under the terms of the  #
# GNU General Public License version 2.0 (GNU GPL-2), as published by  #
# the Free Software Foundation.                                        #
# We created this package for both routine data analysis and academic  #
# research and it was publicly released in the hope that it will be    #
# useful, but it comes WITHOUT ANY WARRANTY OR LIABILITY.              #
#                                                                      #
# Visit our website for the full manual and a complete tutorial about  #
# how to conduct AMR analysis: https://msberends.github.io/AMR/        #
# ==================================================================== #

#' Data sets with `r format(nrow(antibiotics) + nrow(antivirals), big.mark = ",")` antimicrobials
#'
#' Two data sets containing all antibiotics/antimycotics and antivirals. Use [as.ab()] or one of the [`ab_*`][ab_property()] functions to retrieve values from the [antibiotics] data set. Three identifiers are included in this data set: an antibiotic ID (`ab`, primarily used in this package) as defined by WHONET/EARS-Net, an ATC code (`atc`) as defined by the WHO, and a Compound ID (`cid`) as found in PubChem. Other properties in this data set are derived from one or more of these codes.
#' @format
#' ### For the [antibiotics] data set: a [data.frame] with `r nrow(antibiotics)` observations and `r ncol(antibiotics)` variables:
#' - `ab`\cr Antibiotic ID as used in this package (such as `AMC`), using the official EARS-Net (European Antimicrobial Resistance Surveillance Network) codes where available
#' - `atc`\cr ATC code (Anatomical Therapeutic Chemical) as defined by the WHOCC, like `J01CR02`
#' - `cid`\cr Compound ID as found in PubChem
#' - `name`\cr Official name as used by WHONET/EARS-Net or the WHO
#' - `group`\cr A short and concise group name, based on WHONET and WHOCC definitions
#' - `atc_group1`\cr Official pharmacological subgroup (3rd level ATC code) as defined by the WHOCC, like `"Macrolides, lincosamides and streptogramins"`
#' - `atc_group2`\cr Official chemical subgroup (4th level ATC code) as defined by the WHOCC, like `"Macrolides"`
#' - `abbr`\cr List of abbreviations as used in many countries, also for antibiotic susceptibility testing (AST)
#' - `synonyms`\cr Synonyms (often trade names) of a drug, as found in PubChem based on their compound ID
#' - `oral_ddd`\cr Defined Daily Dose (DDD), oral treatment
#' - `oral_units`\cr Units of `oral_ddd`
#' - `iv_ddd`\cr Defined Daily Dose (DDD), parenteral treatment
#' - `iv_units`\cr Units of `iv_ddd`
#' - `loinc`\cr All LOINC codes (Logical Observation Identifiers Names and Codes) associated with the name of the antimicrobial agent. Use [ab_loinc()] to retrieve them quickly, see [ab_property()].
#' 
#' ### For the [antivirals] data set: a [data.frame] with `r nrow(antivirals)` observations and `r ncol(antivirals)` variables:
#' - `atc`\cr ATC code (Anatomical Therapeutic Chemical) as defined by the WHOCC
#' - `cid`\cr Compound ID as found in PubChem
#' - `name`\cr Official name as used by WHONET/EARS-Net or the WHO
#' - `atc_group`\cr Official pharmacological subgroup (3rd level ATC code) as defined by the WHOCC
#' - `synonyms`\cr Synonyms (often trade names) of a drug, as found in PubChem based on their compound ID
#' - `oral_ddd`\cr Defined Daily Dose (DDD), oral treatment
#' - `oral_units`\cr Units of `oral_ddd`
#' - `iv_ddd`\cr Defined Daily Dose (DDD), parenteral treatment
#' - `iv_units`\cr Units of `iv_ddd`
#' @details Properties that are based on an ATC code are only available when an ATC is available. These properties are: `atc_group1`, `atc_group2`, `oral_ddd`, `oral_units`, `iv_ddd` and `iv_units`.
#'
#' Synonyms (i.e. trade names) are derived from the Compound ID (`cid`) and consequently only available where a CID is available.
#' 
#' ### Direct download
#' These data sets are available as 'flat files' for use even without \R - you can find the files here:
#' 
#' * <https://github.com/msberends/AMR/raw/master/data-raw/antibiotics.txt>
#' * <https://github.com/msberends/AMR/raw/master/data-raw/antivirals.txt>
#' 
#' Files in \R format (with preserved data structure) can be found here:
#' 
#' * <https://github.com/msberends/AMR/raw/master/data/antibiotics.rda>
#' * <https://github.com/msberends/AMR/raw/master/data/antivirals.rda>
#' @source World Health Organization (WHO) Collaborating Centre for Drug Statistics Methodology (WHOCC): <https://www.whocc.no/atc_ddd_index/>
#'
#' WHONET 2019 software: <http://www.whonet.org/software.html>
#'
#' European Commission Public Health PHARMACEUTICALS - COMMUNITY REGISTER: <http://ec.europa.eu/health/documents/community-register/html/atc.htm>
#' @inheritSection AMR Reference data publicly available
#' @inheritSection WHOCC WHOCC
#' @inheritSection AMR Read more on our website!
#' @seealso [microorganisms], [intrinsic_resistant]
"antibiotics"

#' @rdname antibiotics
"antivirals"

#' Data set with `r format(nrow(microorganisms), big.mark = ",")` microorganisms
#'
#' A data set containing the microbial taxonomy of six kingdoms from the Catalogue of Life. MO codes can be looked up using [as.mo()].
#' @inheritSection catalogue_of_life Catalogue of Life
#' @format A [data.frame] with `r format(nrow(microorganisms), big.mark = ",")` observations and `r ncol(microorganisms)` variables:
#' - `mo`\cr ID of microorganism as used by this package
#' - `fullname`\cr Full name, like `"Escherichia coli"`
#' - `kingdom`, `phylum`, `class`, `order`, `family`, `genus`, `species`, `subspecies`\cr Taxonomic rank of the microorganism
#' - `rank`\cr Text of the taxonomic rank of the microorganism, like `"species"` or `"genus"`
#' - `ref`\cr Author(s) and year of concerning scientific publication
#' - `species_id`\cr ID of the species as used by the Catalogue of Life
#' - `source`\cr Either "CoL", "DSMZ" (see Source) or "manually added"
#' - `prevalence`\cr Prevalence of the microorganism, see [as.mo()]
#' - `snomed`\cr SNOMED code of the microorganism. Use [mo_snomed()] to retrieve it quickly, see [mo_property()].
#' @details Manually added were:
#' - 11 entries of *Streptococcus* (beta-haemolytic: groups A, B, C, D, F, G, H, K and unspecified; other: viridans, milleri)
#' - 2 entries of *Staphylococcus* (coagulase-negative (CoNS) and coagulase-positive (CoPS))
#' - 3 entries of *Trichomonas* (*Trichomonas vaginalis*, and its family and genus)
#' - 1 entry of *Candida* (*Candida krusei*), that is not (yet) in the Catalogue of Life
#' - 1 entry of *Blastocystis* (*Blastocystis hominis*), although it officially does not exist (Noel *et al.* 2005, PMID 15634993)
#' - 5 other 'undefined' entries (unknown, unknown Gram negatives, unknown Gram positives, unknown yeast and unknown fungus)
#' - 6 families under the Enterobacterales order, according to Adeolu *et al.* (2016, PMID 27620848), that are not (yet) in the Catalogue of Life
#' - `r format(nrow(subset(microorganisms, source == "DSMZ")), big.mark = ",")` species from the DSMZ (Deutsche Sammlung von Mikroorganismen und Zellkulturen) since the DSMZ contain the latest taxonomic information based on recent publications
#' 
#' ### Direct download
#' This data set is available as 'flat file' for use even without \R - you can find the file here:
#' 
#' * <https://github.com/msberends/AMR/raw/master/data-raw/microorganisms.txt>
#' 
#' The file in \R format (with preserved data structure) can be found here:
#' 
#' * <https://github.com/msberends/AMR/raw/master/data/microorganisms.rda>
#' @section About the records from DSMZ (see source):
#' Names of prokaryotes are defined as being validly published by the International Code of Nomenclature of Bacteria. Validly published are all names which are included in the Approved Lists of Bacterial Names and the names subsequently published in the International Journal of Systematic Bacteriology (IJSB) and, from January 2000, in the International Journal of Systematic and Evolutionary Microbiology (IJSEM) as original articles or in the validation lists.
#' *(from <https://www.dsmz.de/services/online-tools/prokaryotic-nomenclature-up-to-date>)*
#' 
#' In February 2020, the DSMZ records were merged with the List of Prokaryotic names with Standing in Nomenclature (LPSN).
#' @source Catalogue of Life: Annual Checklist (public online taxonomic database), <http://www.catalogueoflife.org> (check included annual version with [catalogue_of_life_version()]).
#' 
#' Parte, A.C. (2018). LPSN — List of Prokaryotic names with Standing in Nomenclature (bacterio.net), 20 years on. International Journal of Systematic and Evolutionary Microbiology, 68, 1825-1829; \doi{10.1099/ijsem.0.002786}
#'
#' Leibniz Institute DSMZ-German Collection of Microorganisms and Cell Cultures, Germany, Prokaryotic Nomenclature Up-to-Date, <https://www.dsmz.de/services/online-tools/prokaryotic-nomenclature-up-to-date> and <https://lpsn.dsmz.de> (check included version with [catalogue_of_life_version()]).
#' @inheritSection AMR Reference data publicly available
#' @inheritSection AMR Read more on our website!
#' @seealso [as.mo()], [mo_property()], [microorganisms.codes], [intrinsic_resistant]
"microorganisms"

catalogue_of_life <- list(
  year = 2019,
  version = "Catalogue of Life: {year} Annual Checklist",
  url_CoL = "http://www.catalogueoflife.org/col/",
  url_DSMZ = "https://lpsn.dsmz.de",
  yearmonth_DSMZ = "May 2020"
)

#' Data set with previously accepted taxonomic names
#'
#' A data set containing old (previously valid or accepted) taxonomic names according to the Catalogue of Life. This data set is used internally by [as.mo()].
#' @inheritSection catalogue_of_life Catalogue of Life
#' @format A [data.frame] with `r format(nrow(microorganisms.old), big.mark = ",")` observations and `r ncol(microorganisms.old)` variables:
#' - `fullname`\cr Old full taxonomic name of the microorganism
#' - `fullname_new`\cr New full taxonomic name of the microorganism
#' - `ref`\cr Author(s) and year of concerning scientific publication
#' - `prevalence`\cr Prevalence of the microorganism, see [as.mo()]
#' @source Catalogue of Life: Annual Checklist (public online taxonomic database), <http://www.catalogueoflife.org> (check included annual version with [catalogue_of_life_version()]).
#' 
#' Parte, A.C. (2018). LPSN — List of Prokaryotic names with Standing in Nomenclature (bacterio.net), 20 years on. International Journal of Systematic and Evolutionary Microbiology, 68, 1825-1829; \doi{10.1099/ijsem.0.002786}
#' @inheritSection AMR Reference data publicly available
#' @inheritSection AMR Read more on our website!
#' @seealso [as.mo()] [mo_property()] [microorganisms]
"microorganisms.old"

#' Data set with `r format(nrow(microorganisms.codes), big.mark = ",")` common microorganism codes
#'
#' A data set containing commonly used codes for microorganisms, from laboratory systems and WHONET. Define your own with [set_mo_source()]. They will all be searched when using [as.mo()] and consequently all the [`mo_*`][mo_property()] functions.
#' @format A [data.frame] with `r format(nrow(microorganisms.codes), big.mark = ",")` observations and `r ncol(microorganisms.codes)` variables:
#' - `code`\cr Commonly used code of a microorganism
#' - `mo`\cr ID of the microorganism in the [microorganisms] data set
#' @inheritSection AMR Reference data publicly available
#' @inheritSection catalogue_of_life Catalogue of Life
#' @inheritSection AMR Read more on our website!
#' @seealso [as.mo()] [microorganisms]
"microorganisms.codes"

#' Data set with `r format(nrow(example_isolates), big.mark = ",")` example isolates
#'
#' A data set containing `r format(nrow(example_isolates), big.mark = ",")` microbial isolates with their full antibiograms. The data set reflects reality and can be used to practice AMR analysis. For examples, please read [the tutorial on our website](https://msberends.github.io/AMR/articles/AMR.html).
#' @format A [data.frame] with `r format(nrow(example_isolates), big.mark = ",")` observations and `r ncol(example_isolates)` variables:
#' - `date`\cr date of receipt at the laboratory
#' - `hospital_id`\cr ID of the hospital, from A to D
#' - `ward_icu`\cr logical to determine if ward is an intensive care unit
#' - `ward_clinical`\cr logical to determine if ward is a regular clinical ward
#' - `ward_outpatient`\cr logical to determine if ward is an outpatient clinic
#' - `age`\cr age of the patient
#' - `gender`\cr gender of the patient
#' - `patient_id`\cr ID of the patient
#' - `mo`\cr ID of microorganism created with [as.mo()], see also [microorganisms]
#' - `PEN:RIF`\cr `r sum(vapply(FUN.VALUE = logical(1), example_isolates, is.rsi))` different antibiotics with class [`rsi`] (see [as.rsi()]); these column names occur in the [antibiotics] data set and can be translated with [ab_name()]
#' @inheritSection AMR Reference data publicly available
#' @inheritSection AMR Read more on our website!
"example_isolates"

#' Data set with unclean data
#'
#' A data set containing `r format(nrow(example_isolates_unclean), big.mark = ",")` microbial isolates that are not cleaned up and consequently not ready for AMR analysis. This data set can be used for practice.
#' @format A [data.frame] with `r format(nrow(example_isolates_unclean), big.mark = ",")` observations and `r ncol(example_isolates_unclean)` variables:
#' - `patient_id`\cr ID of the patient
#' - `date`\cr date of receipt at the laboratory
#' - `hospital`\cr ID of the hospital, from A to C
#' - `bacteria`\cr info about microorganism that can be transformed with [as.mo()], see also [microorganisms]
#' - `AMX:GEN`\cr 4 different antibiotics that have to be transformed with [as.rsi()]
#' @inheritSection AMR Reference data publicly available
#' @inheritSection AMR Read more on our website!
"example_isolates_unclean"

#' Data set with `r format(nrow(WHONET), big.mark = ",")` isolates - WHONET example
#'
#' This example data set has the exact same structure as an export file from WHONET. Such files can be used with this package, as this example data set shows. The antibiotic results are from our [example_isolates] data set. All patient names are created using online surname generators and are only in place for practice purposes.
#' @format A [data.frame] with `r format(nrow(WHONET), big.mark = ",")` observations and `r ncol(WHONET)` variables:
#' - `Identification number`\cr ID of the sample
#' - `Specimen number`\cr ID of the specimen
#' - `Organism`\cr Name of the microorganism. Before analysis, you should transform this to a valid microbial class, using [as.mo()].
#' - `Country`\cr Country of origin
#' - `Laboratory`\cr Name of laboratory
#' - `Last name`\cr Fictitious last name of patient
#' - `First name`\cr Fictitious initial of patient
#' - `Sex`\cr Fictitious gender of patient
#' - `Age`\cr Fictitious age of patient
#' - `Age category`\cr Age group, can also be looked up using [age_groups()]
#' - `Date of admission`\cr Date of hospital admission
#' - `Specimen date`\cr Date when specimen was received at laboratory
#' - `Specimen type`\cr Specimen type or group
#' - `Specimen type (Numeric)`\cr Translation of `"Specimen type"`
#' - `Reason`\cr Reason of request with Differential Diagnosis
#' - `Isolate number`\cr ID of isolate
#' - `Organism type`\cr Type of microorganism, can also be looked up using [mo_type()]
#' - `Serotype`\cr Serotype of microorganism
#' - `Beta-lactamase`\cr Microorganism produces beta-lactamase?
#' - `ESBL`\cr Microorganism produces extended spectrum beta-lactamase?
#' - `Carbapenemase`\cr Microorganism produces carbapenemase?
#' - `MRSA screening test`\cr Microorganism is possible MRSA?
#' - `Inducible clindamycin resistance`\cr Clindamycin can be induced?
#' - `Comment`\cr Other comments
#' - `Date of data entry`\cr Date this data was entered in WHONET
#' - `AMP_ND10:CIP_EE`\cr `r sum(vapply(FUN.VALUE = logical(1), WHONET, is.rsi))` different antibiotics. You can lookup the abbreviations in the [antibiotics] data set, or use e.g. [`ab_name("AMP")`][ab_name()] to get the official name immediately. Before analysis, you should transform this to a valid antibiotic class, using [as.rsi()].
#' @inheritSection AMR Reference data publicly available
#' @inheritSection AMR Read more on our website!
"WHONET"

#' Data set for R/SI interpretation
#'
#' Data set to interpret MIC and disk diffusion to R/SI values. Included guidelines are CLSI (`r min(as.integer(gsub("[^0-9]", "", subset(rsi_translation, guideline %like% "CLSI")$guideline)))`-`r max(as.integer(gsub("[^0-9]", "", subset(rsi_translation, guideline %like% "CLSI")$guideline)))`) and EUCAST (`r min(as.integer(gsub("[^0-9]", "", subset(rsi_translation, guideline %like% "EUCAST")$guideline)))`-`r max(as.integer(gsub("[^0-9]", "", subset(rsi_translation, guideline %like% "EUCAST")$guideline)))`). Use [as.rsi()] to transform MICs or disks measurements to R/SI values.
#' @format A [data.frame] with `r format(nrow(rsi_translation), big.mark = ",")` observations and `r ncol(rsi_translation)` variables:
#' - `guideline`\cr Name of the guideline
#' - `method`\cr Either "MIC" or "DISK"
#' - `site`\cr Body site, e.g. "Oral" or "Respiratory"
#' - `mo`\cr Microbial ID, see [as.mo()]
#' - `ab`\cr Antibiotic ID, see [as.ab()]
#' - `ref_tbl`\cr Info about where the guideline rule can be found
#' - `disk_dose`\cr Dose of the used disk diffusion method
#' - `breakpoint_S`\cr Lowest MIC value or highest number of millimetres that leads to "S"
#' - `breakpoint_R`\cr Highest MIC value or lowest number of millimetres that leads to "R"
#' - `uti`\cr A logical value (`TRUE`/`FALSE`) to indicate whether the rule applies to a urinary tract infection (UTI)
#' @details The repository of this `AMR` package contains a file comprising this exact data set: <https://github.com/msberends/AMR/blob/master/data-raw/rsi_translation.txt>. This file **allows for machine reading EUCAST and CLSI guidelines**, which is almost impossible with the Excel and PDF files distributed by EUCAST and CLSI. The file is updated automatically.
#' @inheritSection AMR Reference data publicly available
#' @inheritSection AMR Read more on our website!
#' @seealso [intrinsic_resistant]
"rsi_translation"

#' Data set with bacterial intrinsic resistance
#'
#' Data set containing defined intrinsic resistance by EUCAST of all bug-drug combinations.
#' @format A [data.frame] with `r format(nrow(intrinsic_resistant), big.mark = ",")` observations and `r ncol(intrinsic_resistant)` variables:
#' - `microorganism`\cr Name of the microorganism
#' - `antibiotic`\cr Name of the antibiotic drug
#' @details The repository of this `AMR` package contains a file comprising this exact data set: <https://github.com/msberends/AMR/blob/master/data-raw/intrinsic_resistant.txt>. This file **allows for machine reading EUCAST guidelines about intrinsic resistance**, which is almost impossible with the Excel and PDF files distributed by EUCAST. The file is updated automatically.
#' 
#' This data set is based on `r format_eucast_version_nr(3.2)`.
#' @inheritSection AMR Reference data publicly available
#' @inheritSection AMR Read more on our website!
#' @examples
#' if (require("dplyr")) {
#'   intrinsic_resistant %>%
#'     filter(antibiotic == "Vancomycin", microorganism %like% "Enterococcus") %>% 
#'     pull(microorganism)
#'   # [1] "Enterococcus casseliflavus" "Enterococcus gallinarum"
#' }
"intrinsic_resistant"
```
