## Supplementary material for "AMR - An R Package for Working with Antimicrobial Resistance Data": Source package: deprecated.R

```
# ==================================================================== #
# TITLE                                                                #
# Antimicrobial Resistance (AMR) Analysis for R                        #
#                                                                      #
# SOURCE                                                               #
# https://github.com/msberends/AMR                                     #
#                                                                      #
# LICENCE                                                              #
# (c) 2018-2021 Berends MS, Luz CF et al.                              #
# Developed at the University of Groningen, the Netherlands, in        #
# collaboration with non-profit organisations Certe Medical            #
# Diagnostics & Advice, and University Medical Center Groningen.       # 
#                                                                      #
# This R package is free software; you can freely use and distribute   #
# it for both personal and commercial purposes under the terms of the  #
# GNU General Public License version 2.0 (GNU GPL-2), as published by  #
# the Free Software Foundation.                                        #
# We created this package for both routine data analysis and academic  #
# research and it was publicly released in the hope that it will be    #
# useful, but it comes WITHOUT ANY WARRANTY OR LIABILITY.              #
#                                                                      #
# Visit our website for the full manual and a complete tutorial about  #
# how to conduct AMR analysis: https://msberends.github.io/AMR/        #
# ==================================================================== #

#' Deprecated functions
#'
#' These functions are so-called '[Deprecated]'. They will be removed in a future release. Using the functions will give a warning with the name of the function it has been replaced by (if there is one).
#' @inheritSection lifecycle Retired lifecycle
#' @inheritSection AMR Read more on our website!
#' @keywords internal
#' @name AMR-deprecated
#' @export
p_symbol <- function(p, emptychar = " ") {
  .Deprecated(package = "AMR", new = "cleaner::p_symbol")
  
  p <- as.double(p)
  s <- rep(NA_character_, length(p))
  
  s[p <= 1] <- emptychar
  s[p <= 0.100] <- "."
  s[p <= 0.050] <- "*"
  s[p <= 0.010] <- "**"
  s[p <= 0.001] <- "***"
  
  s
}
```
