## Supplementary material for "AMR - An R Package for Working with Antimicrobial Resistance Data": Source package: disk.R

```
# ==================================================================== #
# TITLE                                                                #
# Antimicrobial Resistance (AMR) Analysis for R                        #
#                                                                      #
# SOURCE                                                               #
# https://github.com/msberends/AMR                                     #
#                                                                      #
# LICENCE                                                              #
# (c) 2018-2021 Berends MS, Luz CF et al.                              #
# Developed at the University of Groningen, the Netherlands, in        #
# collaboration with non-profit organisations Certe Medical            #
# Diagnostics & Advice, and University Medical Center Groningen.       # 
#                                                                      #
# This R package is free software; you can freely use and distribute   #
# it for both personal and commercial purposes under the terms of the  #
# GNU General Public License version 2.0 (GNU GPL-2), as published by  #
# the Free Software Foundation.                                        #
# We created this package for both routine data analysis and academic  #
# research and it was publicly released in the hope that it will be    #
# useful, but it comes WITHOUT ANY WARRANTY OR LIABILITY.              #
#                                                                      #
# Visit our website for the full manual and a complete tutorial about  #
# how to conduct AMR analysis: https://msberends.github.io/AMR/        #
# ==================================================================== #

#' Transform input to disk diffusion diameters
#'
#' This transforms a vector to a new class [`disk`], which is a disk diffusion growth zone size (around an antibiotic disk) in millimetres between 6 and 50.
#' @inheritSection lifecycle Stable lifecycle
#' @rdname as.disk
#' @param x vector
#' @param na.rm a logical indicating whether missing values should be removed
#' @details Interpret disk values as RSI values with [as.rsi()]. It supports guidelines from EUCAST and CLSI.
#' @return An [integer] with additional class [`disk`]
#' @aliases disk
#' @export
#' @seealso [as.rsi()]
#' @inheritSection AMR Read more on our website!
#' @examples
#' \donttest{
#' # transform existing disk zones to the `disk` class
#' df <- data.frame(microorganism = "E. coli",
#'                  AMP = 20,
#'                  CIP = 14,
#'                  GEN = 18,
#'                  TOB = 16)
#' df[, 2:5] <- lapply(df[, 2:5], as.disk)
#' # same with dplyr:
#' # df %>% mutate(across(AMP:TOB, as.disk))
#' 
#' # interpret disk values, see ?as.rsi
#' as.rsi(x = as.disk(18),
#'        mo = "Strep pneu",  # `mo` will be coerced with as.mo()
#'        ab = "ampicillin",  # and `ab` with as.ab()
#'        guideline = "EUCAST")
#'        
#' as.rsi(df)
#' }
as.disk <- function(x, na.rm = FALSE) {
  meet_criteria(x, allow_class = c("disk", "character", "numeric", "integer"), allow_NA = TRUE)
  meet_criteria(na.rm, allow_class = "logical", has_length = 1)
  
  if (!is.disk(x)) {
    x <- x %pm>% unlist()
    if (na.rm == TRUE) {
      x <- x[!is.na(x)]
    }
    x.bak <- x
    
    na_before <- length(x[is.na(x)])
    
    # heavily based on cleaner::clean_double():
    clean_double2 <- function(x, remove = "[^0-9.,-]", fixed = FALSE) {
      x <- gsub(",", ".", x)
      # remove ending dot/comma
      x <- gsub("[,.]$", "", x)
      # only keep last dot/comma
      reverse <- function(x) vapply(FUN.VALUE = character(1), lapply(strsplit(x, NULL), rev), paste, collapse = "")
      x <- sub("{{dot}}", ".", 
               gsub(".", "",
                    reverse(sub(".", "}}tod{{",
                                reverse(x), 
                                fixed = TRUE)),
                    fixed = TRUE), 
               fixed = TRUE)
      x_clean <- gsub(remove, "", x, ignore.case = TRUE, fixed = fixed)
      # remove everything that is not a number or dot
      as.numeric(gsub("[^0-9.]+", "", x_clean))
    }
    
    # round up and make it an integer
    x <- as.integer(ceiling(clean_double2(x)))
    
    # disks can never be less than 6 mm (size of smallest disk) or more than 50 mm
    x[x < 6 | x > 50] <- NA_integer_
    na_after <- length(x[is.na(x)])
    
    if (na_before != na_after) {
      list_missing <- x.bak[is.na(x) & !is.na(x.bak)] %pm>%
        unique() %pm>%
        sort()
      list_missing <- paste0('"', list_missing, '"', collapse = ", ")
      warning_(na_after - na_before, " results truncated (",
               round(((na_after - na_before) / length(x)) * 100),
               "%) that were invalid disk zones: ",
               list_missing, call = FALSE)
    }
  }
  set_clean_class(as.integer(x),
                  new_class = c("disk", "integer"))
}

all_valid_disks <- function(x) {
  if (!inherits(x, c("disk", "character", "numeric", "integer"))) {
    return(FALSE)
  }
  x_disk <- tryCatch(suppressWarnings(as.disk(x[!is.na(x)])),
                     error = function(e) NA)
  !any(is.na(x_disk)) && !all(is.na(x))
}

#' @rdname as.disk
#' @export
is.disk <- function(x) {
  inherits(x, "disk")
}

# will be exported using s3_register() in R/zzz.R
pillar_shaft.disk <- function(x, ...) {
  out <- trimws(format(x))
  out[is.na(x)] <- font_na(NA)
  create_pillar_column(out, align = "right", width = 2)
}

# will be exported using s3_register() in R/zzz.R
type_sum.disk <- function(x, ...) {
  "disk"
}

#' @method print disk
#' @export
#' @noRd
print.disk <- function(x, ...) {
  cat("Class <disk>\n")
  print(as.integer(x), quote = FALSE)
}

#' @method plot disk
#' @export
#' @importFrom graphics barplot axis
#' @rdname plot
plot.disk <- function(x,
                      main = paste("Disk zones values of", deparse(substitute(x))),
                      ylab = "Frequency",
                      xlab = "Disk diffusion (mm)",
                      axes = FALSE,
                      ...) {
  meet_criteria(main, allow_class = "character", has_length = 1)
  meet_criteria(ylab, allow_class = "character", has_length = 1)
  meet_criteria(xlab, allow_class = "character", has_length = 1)
  meet_criteria(axes, allow_class = "logical", has_length = 1)
  
  barplot(table(x),
          ylab = ylab,
          xlab = xlab,
          axes = axes,
          main = main,
          ...)
  axis(2, seq(0, max(table(x))))
}

#' @method [ disk
#' @export
#' @noRd
"[.disk" <- function(x, ...) {
  y <- NextMethod()
  attributes(y) <- attributes(x)
  y
}
#' @method [[ disk
#' @export
#' @noRd
"[[.disk" <- function(x, ...) {
  y <- NextMethod()
  attributes(y) <- attributes(x)
  y
}
#' @method [<- disk
#' @export
#' @noRd
"[<-.disk" <- function(i, j, ..., value) {
  value <- as.disk(value)
  y <- NextMethod()
  attributes(y) <- attributes(i)
  y
}
#' @method [[<- disk
#' @export
#' @noRd
"[[<-.disk" <- function(i, j, ..., value) {
  value <- as.disk(value)
  y <- NextMethod()
  attributes(y) <- attributes(i)
  y
}
#' @method c disk
#' @export
#' @noRd
c.disk <- function(x, ...) {
  y <- NextMethod()
  y <- as.disk(y)
  attributes(y) <- attributes(x)
  y
}

#' @method unique disk
#' @export
#' @noRd
unique.disk <- function(x, incomparables = FALSE, ...) {
  y <- NextMethod()
  attributes(y) <- attributes(x)
  y
}

# will be exported using s3_register() in R/zzz.R
get_skimmers.disk <- function(column) {
  skimr::sfl(
    skim_type = "disk",
    min = ~min(as.double(.), na.rm = TRUE),
    max = ~max(as.double(.), na.rm = TRUE),
    median = ~stats::median(as.double(.), na.rm = TRUE),
    n_unique = ~pm_n_distinct(., na.rm = TRUE),
    hist = ~skimr::inline_hist(stats::na.omit(as.double(.)))
  )
}
```
