## Supplementary material for "AMR - An R Package for Working with Antimicrobial Resistance Data": Source package: episode.R

```
# ==================================================================== #
# TITLE                                                                #
# Antimicrobial Resistance (AMR) Analysis for R                        #
#                                                                      #
# SOURCE                                                               #
# https://github.com/msberends/AMR                                     #
#                                                                      #
# LICENCE                                                              #
# (c) 2018-2021 Berends MS, Luz CF et al.                              #
# Developed at the University of Groningen, the Netherlands, in        #
# collaboration with non-profit organisations Certe Medical            #
# Diagnostics & Advice, and University Medical Center Groningen.       # 
#                                                                      #
# This R package is free software; you can freely use and distribute   #
# it for both personal and commercial purposes under the terms of the  #
# GNU General Public License version 2.0 (GNU GPL-2), as published by  #
# the Free Software Foundation.                                        #
# We created this package for both routine data analysis and academic  #
# research and it was publicly released in the hope that it will be    #
# useful, but it comes WITHOUT ANY WARRANTY OR LIABILITY.              #
#                                                                      #
# Visit our website for the full manual and a complete tutorial about  #
# how to conduct AMR analysis: https://msberends.github.io/AMR/        #
# ==================================================================== #

#' Determine (new) episodes for patients
#' 
#' These functions determine which items in a vector can be considered (the start of) a new episode, based on the argument `episode_days`. This can be used to determine clinical episodes for any epidemiological analysis. The [get_episode()] function returns the index number of the episode per group, while the [is_new_episode()] function returns values `TRUE`/`FALSE` to indicate whether an item in a vector is the start of a new episode.
#' @inheritSection lifecycle Stable lifecycle
#' @param x vector of dates (class `Date` or `POSIXt`)
#' @param episode_days length of the required episode in days, please see *Details*
#' @param ... arguments passed on to [as.Date()]
#' @details 
#' Dates are first sorted from old to new. The oldest date will mark the start of the first episode. After this date, the next date will be marked that is at least `episode_days` days later than the start of the first episode. From that second marked date on, the next date will be marked that is at least `episode_days` days later than the start of the second episode which will be the start of the third episode, and so on. Before the vector is being returned, the original order will be restored.
#' 
#' The [first_isolate()] function is a wrapper around the [is_new_episode()] function, but is more efficient for data sets containing microorganism codes or names.
#' 
#' The `dplyr` package is not required for these functions to work, but these functions support [variable grouping][dplyr::group_by()] and work conveniently inside `dplyr` verbs such as [`filter()`][dplyr::filter()], [`mutate()`][dplyr::mutate()] and [`summarise()`][dplyr::summarise()].
#' @return 
#' * [get_episode()]: a [double] vector
#' * [is_new_episode()]: a [logical] vector
#' @seealso [first_isolate()]
#' @rdname get_episode
#' @export
#' @inheritSection AMR Read more on our website!
#' @examples
#' # `example_isolates` is a dataset available in the AMR package.
#' # See ?example_isolates.
#' 
#' get_episode(example_isolates$date, episode_days = 60)
#' is_new_episode(example_isolates$date, episode_days = 60)
#' 
#' # filter on results from the third 60-day episode only, using base R
#' example_isolates[which(get_episode(example_isolates$date, 60) == 3), ]
#' 
#' \donttest{
#' if (require("dplyr")) {
#'   # is_new_episode() can also be used in dplyr verbs to determine patient
#'   # episodes based on any (combination of) grouping variables:
#'   example_isolates %>%
#'     mutate(condition = sample(x = c("A", "B", "C"), 
#'                               size = 2000,
#'                               replace = TRUE)) %>% 
#'     group_by(condition) %>%
#'     mutate(new_episode = is_new_episode(date, 365))
#'     
#'   example_isolates %>%
#'     group_by(hospital_id, patient_id) %>%
#'     transmute(date, 
#'               patient_id,
#'               new_index = get_episode(date, 60),
#'               new_logical = is_new_episode(date, 60))
#'   
#'   
#'   example_isolates %>%
#'     group_by(hospital_id) %>% 
#'     summarise(patients = n_distinct(patient_id),
#'               n_episodes_365 = sum(is_new_episode(date, episode_days = 365)),
#'               n_episodes_60  = sum(is_new_episode(date, episode_days = 60)),
#'               n_episodes_30  = sum(is_new_episode(date, episode_days = 30)))
#'     
#'     
#'   # grouping on patients and microorganisms leads to the same results
#'   # as first_isolate():
#'   x <- example_isolates %>%
#'     filter(first_isolate(., include_unknown = TRUE))
#'     
#'   y <- example_isolates %>%
#'     group_by(patient_id, mo) %>%
#'     filter(is_new_episode(date, 365))
#'
#'   identical(x$patient_id, y$patient_id)
#'   
#'   # but is_new_episode() has a lot more flexibility than first_isolate(),
#'   # since you can now group on anything that seems relevant:
#'   example_isolates %>%
#'     group_by(patient_id, mo, hospital_id, ward_icu) %>%
#'     mutate(flag_episode = is_new_episode(date, 365))
#' }
#' }
get_episode <- function(x, episode_days, ...) {
  meet_criteria(x, allow_class = c("Date", "POSIXt"))
  meet_criteria(episode_days, allow_class = c("numeric", "double", "integer"), has_length = 1)
  
  exec_episode(type = "sequential",
               x = x, 
               episode_days = episode_days,
               ... = ...)
}

#' @rdname get_episode
#' @export
is_new_episode <- function(x, episode_days, ...) {
  meet_criteria(x, allow_class = c("Date", "POSIXt"))
  meet_criteria(episode_days, allow_class = c("numeric", "double", "integer"), has_length = 1)
  
  exec_episode(type = "logical",
               x = x, 
               episode_days = episode_days,
               ... = ...)
}

exec_episode <- function(type, x, episode_days, ...) {
  x <- as.double(as.Date(x, ...)) # as.Date() for POSIX classes
  if (length(x) == 1) {
    if (type == "logical") {
      return(TRUE)
    } else if (type == "sequential") {
      return(1)
    }
  } else if (length(x) == 2) {
    if (max(x) - min(x) >= episode_days) {
      if (type == "logical") {
        return(c(TRUE, TRUE))
      } else if (type == "sequential") {
        return(c(1, 2))
      }
    } else {
      if (type == "logical") {
        return(c(TRUE, FALSE))
      } else if (type == "sequential") {
        return(c(1, 1))
      }
    }
  }
  
  # I asked on StackOverflow:
  # https://stackoverflow.com/questions/42122245/filter-one-row-every-year
  exec <- function(x, episode_days) {
    indices <- integer()
    start <- x[1]
    ind <- 1
    indices[1] <- 1
    for (i in 2:length(x)) {
      if (isTRUE((x[i] - start) >= episode_days)) {
        ind <- ind + 1
        if (type == "logical") {
          indices[ind] <- i
        }
        start <- x[i]
      }
      if (type == "sequential") {
        indices[i] <- ind
      }
    }
    if (type == "logical") {
      result <- rep(FALSE, length(x))
      result[indices] <- TRUE
      result
    } else if (type == "sequential") {
      indices
    }
  }
  
  df <- data.frame(x = x,
                   y = seq_len(length(x))) %pm>%
    pm_arrange(x)
  df$new <- exec(df$x, episode_days)
  df %pm>%
    pm_arrange(y) %pm>%
    pm_pull(new)
}
```
