## Supplementary material for "AMR - An R Package for Working with Antimicrobial Resistance Data": Source package: eucast_rules.R

```
# ==================================================================== #
# TITLE                                                                #
# Antimicrobial Resistance (AMR) Analysis for R                        #
#                                                                      #
# SOURCE                                                               #
# https://github.com/msberends/AMR                                     #
#                                                                      #
# LICENCE                                                              #
# (c) 2018-2021 Berends MS, Luz CF et al.                              #
# Developed at the University of Groningen, the Netherlands, in        #
# collaboration with non-profit organisations Certe Medical            #
# Diagnostics & Advice, and University Medical Center Groningen.       # 
#                                                                      #
# This R package is free software; you can freely use and distribute   #
# it for both personal and commercial purposes under the terms of the  #
# GNU General Public License version 2.0 (GNU GPL-2), as published by  #
# the Free Software Foundation.                                        #
# We created this package for both routine data analysis and academic  #
# research and it was publicly released in the hope that it will be    #
# useful, but it comes WITHOUT ANY WARRANTY OR LIABILITY.              #
#                                                                      #
# Visit our website for the full manual and a complete tutorial about  #
# how to conduct AMR analysis: https://msberends.github.io/AMR/        #
# ==================================================================== #

# add new version numbers here, and add the rules themselves to "data-raw/eucast_rules.tsv"
# (running "data-raw/internals.R" will process that TSV file)
EUCAST_VERSION_BREAKPOINTS <- list("10.0" = list(version_txt = "v10.0",
                                                 year = 2020, 
                                                 title = "'EUCAST Clinical Breakpoints'",
                                                 url = "https://www.eucast.org/clinical_breakpoints/"))
EUCAST_VERSION_EXPERT_RULES <- list("3.1" = list(version_txt = "v3.1",
                                                 year = 2016, 
                                                 title = "'EUCAST Expert Rules, Intrinsic Resistance and Exceptional Phenotypes'",
                                                 url = "https://www.eucast.org/expert_rules_and_intrinsic_resistance/"),
                                    "3.2" = list(version_txt = "v3.2",
                                                 year = 2020, 
                                                 title = "'EUCAST Expert Rules' and 'EUCAST Intrinsic Resistance and Unusual Phenotypes'",
                                                 url = "https://www.eucast.org/expert_rules_and_intrinsic_resistance/"))

format_eucast_version_nr <- function(version, markdown = TRUE) {
  # for documentation - adds title, version number, year and url in markdown language
  lst <- c(EUCAST_VERSION_BREAKPOINTS, EUCAST_VERSION_EXPERT_RULES)
  version <- format(version, nsmall = 1)
  if (markdown == TRUE) {
    paste0("[", lst[[version]]$title, " ", lst[[version]]$version_txt, "](", lst[[version]]$url, ")",
           " from ", lst[[version]]$year)
  } else {
    paste0(lst[[version]]$title, " ", lst[[version]]$version_txt,
           " from ", lst[[version]]$year)
  }
}

#' Apply EUCAST rules
#' 
#' @description
#' Apply rules for clinical breakpoints and intrinsic resistance as defined by the European Committee on Antimicrobial Susceptibility Testing (EUCAST, <https://eucast.org>), see *Source*.
#' 
#' To improve the interpretation of the antibiogram before EUCAST rules are applied, some non-EUCAST rules can applied at default, see Details.
#' @inheritSection lifecycle Stable lifecycle
#' @param x data with antibiotic columns, such as `amox`, `AMX` and `AMC`
#' @param info a logical to indicate whether progress should be printed to the console, defaults to only print while in interactive sessions
#' @param rules a character vector that specifies which rules should be applied. Must be one or more of `"breakpoints"`, `"expert"`, `"other"`, `"all"`, and defaults to `c("breakpoints", "expert")`. The default value can be set to another value, e.g. using `options(AMR_eucastrules = "all")`.
#' @param verbose a [logical] to turn Verbose mode on and off (default is off). In Verbose mode, the function does not apply rules to the data, but instead returns a data set in logbook form with extensive info about which rows and columns would be effected and in which way. Using Verbose mode takes a lot more time.
#' @param version_breakpoints the version number to use for the EUCAST Clinical Breakpoints guideline. Currently supported: `r paste0(names(EUCAST_VERSION_BREAKPOINTS), collapse = ", ")`.
#' @param version_expertrules the version number to use for the EUCAST Expert Rules and Intrinsic Resistance guideline. Currently supported: `r paste0(names(EUCAST_VERSION_EXPERT_RULES), collapse = ", ")`.
#' @param ampc_cephalosporin_resistance a character value that should be applied for AmpC de-repressed cephalosporin-resistant mutants, defaults to `NA`. Currently only works when `version_expertrules` is `3.2`; '*EUCAST Expert Rules v3.2 on Enterobacterales*' states that susceptible (S) results of cefotaxime, ceftriaxone and ceftazidime should be reported with a note, or results should be suppressed (emptied) for these agents. A value of `NA` for this argument will remove results for these agents, while e.g. a value of `"R"` will make the results for these agents resistant. Use `NULL` to not alter the results for AmpC de-repressed cephalosporin-resistant mutants. \cr For *EUCAST Expert Rules* v3.2, this rule applies to: *`r gsub("[)(^]", "", gsub("|", ", ", eucast_rules_file[which(eucast_rules_file$reference.version == 3.2 & eucast_rules_file$reference.rule %like% "ampc"), "this_value"][1], fixed = TRUE))`*.
#' 
#' @param ... column name of an antibiotic, please see section *Antibiotics* below
#' @inheritParams first_isolate
#' @details
#' **Note:** This function does not translate MIC values to RSI values. Use [as.rsi()] for that. \cr
#' **Note:** When ampicillin (AMP, J01CA01) is not available but amoxicillin (AMX, J01CA04) is, the latter will be used for all rules where there is a dependency on ampicillin. These drugs are interchangeable when it comes to expression of antimicrobial resistance.
#'
#' The file containing all EUCAST rules is located here: <https://github.com/msberends/AMR/blob/master/data-raw/eucast_rules.tsv>.
#' 
#' ## 'Other' rules
#' 
#' Before further processing, two non-EUCAST rules about drug combinations can be applied to improve the efficacy of the EUCAST rules, and the reliability of your data (analysis). These rules are:
#' 
#' 1. A drug **with** enzyme inhibitor will be set to S if the same drug **without** enzyme inhibitor is S
#' 2. A drug **without** enzyme inhibitor will be set to R if the same drug **with** enzyme inhibitor is R
#' 
#' Important examples include amoxicillin and amoxicillin/clavulanic acid, and trimethoprim and trimethoprim/sulfamethoxazole. Needless to say, for these rules to work, both drugs must be available in the data set.
#' 
#' Since these rules are not officially approved by EUCAST, they are not applied at default. To use these rules, include `"other"` to the `rules` argument, or use `eucast_rules(..., rules = "all")`. You can also set the option `AMR_eucastrules`, i.e. run `options(AMR_eucastrules = "all")`.
#' @section Antibiotics:
#' To define antibiotics column names, leave as it is to determine it automatically with [guess_ab_col()] or input a text (case-insensitive), or use `NULL` to skip a column (e.g. `TIC = NULL` to skip ticarcillin). Manually defined but non-existing columns will be skipped with a warning.
#'
#' The following antibiotics are used for the functions [eucast_rules()] and [mdro()]. These are shown below in the format 'name (`antimicrobial ID`, [ATC code](https://www.whocc.no/atc/structure_and_principles/))', sorted alphabetically:
#'
#' `r create_ab_documentation(c("AMC", "AMK", "AMP", "AMX", "ATM", "AVO", "AZL", "AZM", "BAM", "BPR", "CAC", "CAT", "CAZ", "CCP", "CCV", "CCX", "CDC", "CDR", "CDZ", "CEC", "CED", "CEI", "CEM", "CEP", "CFM", "CFM1", "CFP", "CFR", "CFS", "CFZ", "CHE", "CHL", "CID", "CIP", "CLI", "CLR", "CMX", "CMZ", "CND", "COL", "CPD", "CPI", "CPL", "CPM", "CPO", "CPR", "CPT", "CPX", "CRB", "CRD", "CRN", "CRO", "CSL", "CTB", "CTC", "CTF", "CTL", "CTS", "CTT", "CTX", "CTZ", "CXM", "CYC", "CZA", "CZD", "CZO", "CZP", "CZX", "DAL", "DAP", "DIR", "DIT", "DIX", "DIZ", "DKB", "DOR", "DOX", "ENX", "EPC", "ERY", "ETP", "FEP", "FLC", "FLE", "FLR1", "FOS", "FOV", "FOX", "FOX1", "FUS", "GAT", "GEM", "GEN", "GRX", "HAP", "HET", "IPM", "ISE", "JOS", "KAN", "LEX", "LIN", "LNZ", "LOM", "LOR", "LTM", "LVX", "MAN", "MCM", "MEC", "MEM", "MEV", "MEZ", "MFX", "MID", "MNO", "MTM", "NAL", "NEO", "NET", "NIT", "NOR", "NOV", "NVA", "OFX", "OLE", "ORI", "OXA", "PAZ", "PEF", "PEN", "PHN", "PIP", "PLB", "PME", "PRI", "PRL", "PRU", "PVM", "QDA", "RAM", "RFL", "RID", "RIF", "ROK", "RST", "RXT", "SAM", "SBC", "SDI", "SDM", "SIS", "SLF", "SLF1", "SLF10", "SLF11", "SLF12", "SLF13", "SLF2", "SLF3", "SLF4", "SLF5", "SLF6", "SLF7", "SLF8", "SLF9", "SLT1", "SLT2", "SLT3", "SLT4", "SLT5", "SMX", "SPI", "SPX", "STR", "STR1", "SUD", "SUT", "SXT", "SZO", "TAL", "TCC", "TCM", "TCY", "TEC", "TEM", "TGC", "THA", "TIC", "TIO", "TLT", "TLV", "TMP", "TMX", "TOB", "TRL", "TVA", "TZD", "TZP", "VAN"))`
#' @aliases EUCAST
#' @rdname eucast_rules
#' @export
#' @return The input of `x`, possibly with edited values of antibiotics. Or, if `verbose = TRUE`, a [data.frame] with all original and new values of the affected bug-drug combinations.
#' @source
#' - EUCAST Expert Rules. Version 2.0, 2012.\cr
#'   Leclercq et al. **EUCAST expert rules in antimicrobial susceptibility testing.** *Clin Microbiol Infect.* 2013;19(2):141-60; \doi{https://doi.org/10.1111/j.1469-0691.2011.03703.x}
#' - EUCAST Expert Rules, Intrinsic Resistance and Exceptional Phenotypes Tables. Version 3.1, 2016. [(link)](https://www.eucast.org/fileadmin/src/media/PDFs/EUCAST_files/Expert_Rules/Expert_rules_intrinsic_exceptional_V3.1.pdf)
#' - EUCAST Intrinsic Resistance and Unusual Phenotypes. Version 3.2, 2020. [(link)](https://www.eucast.org/fileadmin/src/media/PDFs/EUCAST_files/Expert_Rules/2020/Intrinsic_Resistance_and_Unusual_Phenotypes_Tables_v3.2_20200225.pdf)
#' - EUCAST Breakpoint tables for interpretation of MICs and zone diameters. Version 9.0, 2019. [(link)](https://www.eucast.org/fileadmin/src/media/PDFs/EUCAST_files/Breakpoint_tables/v_9.0_Breakpoint_Tables.xlsx)
#' - EUCAST Breakpoint tables for interpretation of MICs and zone diameters. Version 10.0, 2020. [(link)](https://www.eucast.org/fileadmin/src/media/PDFs/EUCAST_files/Breakpoint_tables/v_10.0_Breakpoint_Tables.xlsx)
#' @inheritSection AMR Reference data publicly available
#' @inheritSection AMR Read more on our website!
#' @examples
#' \donttest{
#' a <- data.frame(mo = c("Staphylococcus aureus",
#'                        "Enterococcus faecalis",
#'                        "Escherichia coli",
#'                        "Klebsiella pneumoniae",
#'                        "Pseudomonas aeruginosa"),
#'                 VAN = "-",       # Vancomycin
#'                 AMX = "-",       # Amoxicillin
#'                 COL = "-",       # Colistin
#'                 CAZ = "-",       # Ceftazidime
#'                 CXM = "-",       # Cefuroxime
#'                 PEN = "S",       # Benzylpenicillin
#'                 FOX = "S",       # Cefoxitin
#'                 stringsAsFactors = FALSE)
#'
#' a
#' #                       mo  VAN  AMX  COL  CAZ  CXM  PEN  FOX
#' # 1  Staphylococcus aureus    -    -    -    -    -    S    S
#' # 2  Enterococcus faecalis    -    -    -    -    -    S    S
#' # 3       Escherichia coli    -    -    -    -    -    S    S
#' # 4  Klebsiella pneumoniae    -    -    -    -    -    S    S
#' # 5 Pseudomonas aeruginosa    -    -    -    -    -    S    S
#'
#'
#' # apply EUCAST rules: some results wil be changed
#' b <- eucast_rules(a)
#'
#' b
#' #                       mo  VAN  AMX  COL  CAZ  CXM  PEN  FOX
#' # 1  Staphylococcus aureus    -    S    R    R    S    S    S
#' # 2  Enterococcus faecalis    -    -    R    R    R    S    R
#' # 3       Escherichia coli    R    -    -    -    -    R    S
#' # 4  Klebsiella pneumoniae    R    R    -    -    -    R    S
#' # 5 Pseudomonas aeruginosa    R    R    -    -    R    R    R
#'
#'
#' # do not apply EUCAST rules, but rather get a data.frame
#' # containing all details about the transformations:
#' c <- eucast_rules(a, verbose = TRUE)
#' }
eucast_rules <- function(x,
                         col_mo = NULL,
                         info = interactive(),
                         rules = getOption("AMR_eucastrules", default = c("breakpoints", "expert")),
                         verbose = FALSE,
                         version_breakpoints = 10.0,
                         version_expertrules = 3.2,
                         ampc_cephalosporin_resistance = NA,
                         ...) {
  meet_criteria(x, allow_class = "data.frame")
  meet_criteria(col_mo, allow_class = "character", has_length = 1, is_in = colnames(x), allow_NULL = TRUE)
  meet_criteria(info, allow_class = "logical", has_length = 1)
  meet_criteria(rules, allow_class = "character", has_length = c(1, 2, 3, 4), is_in = c("breakpoints", "expert", "other", "all"))
  meet_criteria(verbose, allow_class = "logical", has_length = 1)
  meet_criteria(version_breakpoints, allow_class = "numeric", has_length = 1)
  meet_criteria(version_expertrules, allow_class = "numeric", has_length = 1)
  meet_criteria(ampc_cephalosporin_resistance, allow_class = c("rsi", "character"), has_length = 1, allow_NA = TRUE, allow_NULL = TRUE, is_in = c("R", "S", "I"))
  
  x_deparsed <- deparse(substitute(x))
  if (length(x_deparsed) > 1 || !all(x_deparsed %like% "[a-z]+")) {
    x_deparsed <- "your_data"
  }
  
  check_dataset_integrity()
  
  version_breakpoints <- as.double(gsub("[^0-9.]+", "", version_breakpoints))
  version_expertrules <- as.double(gsub("[^0-9.]+", "", version_expertrules))
  stop_ifnot(version_breakpoints %in% as.double(names(EUCAST_VERSION_BREAKPOINTS)),
             "EUCAST version ", version_breakpoints, " for clinical breakpoints not found")
  stop_ifnot(version_expertrules %in% as.double(names(EUCAST_VERSION_EXPERT_RULES)),
             "EUCAST version ", version_expertrules, " for expert rules/intrinsic resistance not found")
  breakpoints_info <- EUCAST_VERSION_BREAKPOINTS[[which(as.double(names(EUCAST_VERSION_BREAKPOINTS)) == version_breakpoints)]]
  expertrules_info <- EUCAST_VERSION_EXPERT_RULES[[which(as.double(names(EUCAST_VERSION_EXPERT_RULES)) == version_expertrules)]]
  
  # support old setting (until AMR v1.3.0)
  if (missing(rules) & !is.null(getOption("AMR.eucast_rules", default = NULL))) {
    rules <- getOption("AMR.eucast_rules")
  }
  
  if (interactive() & verbose == TRUE & info == TRUE) {
    txt <- paste0("WARNING: In Verbose mode, the eucast_rules() function does not apply rules to the data, but instead returns a data set in logbook form with extensive info about which rows and columns would be effected and in which way.",
                  "\n\nThis may overwrite your existing data if you use e.g.:",
                  "\ndata <- eucast_rules(data, verbose = TRUE)\n\nDo you want to continue?")
    showQuestion <- import_fn("showQuestion", "rstudioapi", error_on_fail = FALSE)
    if (!is.null(showQuestion)) {
      q_continue <- showQuestion("Using verbose = TRUE with eucast_rules()", txt)
    } else {
      q_continue <- utils::menu(choices = c("OK", "Cancel"), graphics = FALSE, title = txt)
    }
    if (q_continue %in% c(FALSE, 2)) {
      message_("Cancelled, returning original data", add_fn = font_red, as_note = FALSE)
      return(x)
    }
  }
  
  # try to find columns based on type
  # -- mo
  if (is.null(col_mo)) {
    col_mo <- search_type_in_df(x = x, type = "mo", info = info)
  }
  stop_if(is.null(col_mo), "`col_mo` must be set")
  
  decimal.mark <- getOption("OutDec")
  big.mark <- ifelse(decimal.mark != ",", ",", ".")
  formatnr <- function(x, big = big.mark, dec = decimal.mark) {
    trimws(format(x, big.mark = big, decimal.mark = dec))
  }
  
  warned <- FALSE
  warn_lacking_rsi_class <- character(0)
  txt_ok <- function(n_added, n_changed, warned = FALSE) {
    if (warned == FALSE) {
      if (n_added + n_changed == 0) {
        cat(font_subtle(" (no changes)\n"))
      } else {
        # opening
        cat(font_grey(" ("))
        # additions
        if (n_added > 0) {
          if (n_added == 1) {
            cat(font_green("1 value added"))
          } else {
            cat(font_green(formatnr(n_added), "values added"))
          }
        }
        # separator
        if (n_added > 0 & n_changed > 0) {
          cat(font_grey(", "))
        }
        # changes
        if (n_changed > 0) {
          if (n_changed == 1) {
            cat(font_blue("1 value changed"))
          } else {
            cat(font_blue(formatnr(n_changed), "values changed"))
          }
        } 
        # closing
        cat(font_grey(")\n"))
      }
      warned <<- FALSE
    }
  }
  
  cols_ab <- get_column_abx(x = x,
                            soft_dependencies = c("AMC",
                                                  "AMP",
                                                  "AMX",
                                                  "CIP",
                                                  "ERY",
                                                  "FOX1",
                                                  "GEN",
                                                  "MFX",
                                                  "NAL",
                                                  "NOR",
                                                  "PEN",
                                                  "PIP",
                                                  "TCY",
                                                  "TIC",
                                                  "TOB"),
                            hard_dependencies = NULL,
                            verbose = verbose,
                            info = info,
                            ...)
  
  AMC <- cols_ab["AMC"]
  AMK <- cols_ab["AMK"]
  AMP <- cols_ab["AMP"]
  AMX <- cols_ab["AMX"]
  ATM <- cols_ab["ATM"]
  AVO <- cols_ab["AVO"]
  AZL <- cols_ab["AZL"]
  AZM <- cols_ab["AZM"]
  BAM <- cols_ab["BAM"]
  BPR <- cols_ab["BPR"]
  CAC <- cols_ab["CAC"]
  CAT <- cols_ab["CAT"]
  CAZ <- cols_ab["CAZ"]
  CCP <- cols_ab["CCP"]
  CCV <- cols_ab["CCV"]
  CCX <- cols_ab["CCX"]
  CDC <- cols_ab["CDC"]
  CDR <- cols_ab["CDR"]
  CDZ <- cols_ab["CDZ"]
  CEC <- cols_ab["CEC"]
  CED <- cols_ab["CED"]
  CEI <- cols_ab["CEI"]
  CEM <- cols_ab["CEM"]
  CEP <- cols_ab["CEP"]
  CFM <- cols_ab["CFM"]
  CFM1 <- cols_ab["CFM1"]
  CFP <- cols_ab["CFP"]
  CFR <- cols_ab["CFR"]
  CFS <- cols_ab["CFS"]
  CFZ <- cols_ab["CFZ"]
  CHE <- cols_ab["CHE"]
  CHL <- cols_ab["CHL"]
  CID <- cols_ab["CID"]
  CIP <- cols_ab["CIP"]
  CLI <- cols_ab["CLI"]
  CLR <- cols_ab["CLR"]
  CMX <- cols_ab["CMX"]
  CMZ <- cols_ab["CMZ"]
  CND <- cols_ab["CND"]
  COL <- cols_ab["COL"]
  CPD <- cols_ab["CPD"]
  CPI <- cols_ab["CPI"]
  CPL <- cols_ab["CPL"]
  CPM <- cols_ab["CPM"]
  CPO <- cols_ab["CPO"]
  CPR <- cols_ab["CPR"]
  CPT <- cols_ab["CPT"]
  CPX <- cols_ab["CPX"]
  CRB <- cols_ab["CRB"]
  CRD <- cols_ab["CRD"]
  CRN <- cols_ab["CRN"]
  CRO <- cols_ab["CRO"]
  CSL <- cols_ab["CSL"]
  CTB <- cols_ab["CTB"]
  CTC <- cols_ab["CTC"]
  CTF <- cols_ab["CTF"]
  CTL <- cols_ab["CTL"]
  CTS <- cols_ab["CTS"]
  CTT <- cols_ab["CTT"]
  CTX <- cols_ab["CTX"]
  CTZ <- cols_ab["CTZ"]
  CXM <- cols_ab["CXM"]
  CYC <- cols_ab["CYC"]
  CZA <- cols_ab["CZA"]
  CZD <- cols_ab["CZD"]
  CZO <- cols_ab["CZO"]
  CZP <- cols_ab["CZP"]
  CZX <- cols_ab["CZX"]
  DAL <- cols_ab["DAL"]
  DAP <- cols_ab["DAP"]
  DIR <- cols_ab["DIR"]
  DIT <- cols_ab["DIT"]
  DIX <- cols_ab["DIX"]
  DIZ <- cols_ab["DIZ"]
  DKB <- cols_ab["DKB"]
  DOR <- cols_ab["DOR"]
  DOX <- cols_ab["DOX"]
  ENX <- cols_ab["ENX"]
  EPC <- cols_ab["EPC"]
  ERY <- cols_ab["ERY"]
  ETP <- cols_ab["ETP"]
  FEP <- cols_ab["FEP"]
  FLC <- cols_ab["FLC"]
  FLE <- cols_ab["FLE"]
  FLR1 <- cols_ab["FLR1"]
  FOS <- cols_ab["FOS"]
  FOV <- cols_ab["FOV"]
  FOX <- cols_ab["FOX"]
  FOX1 <- cols_ab["FOX1"]
  FUS <- cols_ab["FUS"]
  GAT <- cols_ab["GAT"]
  GEM <- cols_ab["GEM"]
  GEN <- cols_ab["GEN"]
  GRX <- cols_ab["GRX"]
  HAP <- cols_ab["HAP"]
  HET <- cols_ab["HET"]
  IPM <- cols_ab["IPM"]
  ISE <- cols_ab["ISE"]
  JOS <- cols_ab["JOS"]
  KAN <- cols_ab["KAN"]
  LEX <- cols_ab["LEX"]
  LIN <- cols_ab["LIN"]
  LNZ <- cols_ab["LNZ"]
  LOM <- cols_ab["LOM"]
  LOR <- cols_ab["LOR"]
  LTM <- cols_ab["LTM"]
  LVX <- cols_ab["LVX"]
  MAN <- cols_ab["MAN"]
  MCM <- cols_ab["MCM"]
  MEC <- cols_ab["MEC"]
  MEM <- cols_ab["MEM"]
  MEV <- cols_ab["MEV"]
  MEZ <- cols_ab["MEZ"]
  MFX <- cols_ab["MFX"]
  MID <- cols_ab["MID"]
  MNO <- cols_ab["MNO"]
  MTM <- cols_ab["MTM"]
  NAL <- cols_ab["NAL"]
  NEO <- cols_ab["NEO"]
  NET <- cols_ab["NET"]
  NIT <- cols_ab["NIT"]
  NOR <- cols_ab["NOR"]
  NOV <- cols_ab["NOV"]
  NVA <- cols_ab["NVA"]
  OFX <- cols_ab["OFX"]
  OLE <- cols_ab["OLE"]
  ORI <- cols_ab["ORI"]
  OXA <- cols_ab["OXA"]
  PAZ <- cols_ab["PAZ"]
  PEF <- cols_ab["PEF"]
  PEN <- cols_ab["PEN"]
  PHN <- cols_ab["PHN"]
  PIP <- cols_ab["PIP"]
  PLB <- cols_ab["PLB"]
  PME <- cols_ab["PME"]
  PRI <- cols_ab["PRI"]
  PRL <- cols_ab["PRL"]
  PRU <- cols_ab["PRU"]
  PVM <- cols_ab["PVM"]
  QDA <- cols_ab["QDA"]
  RAM <- cols_ab["RAM"]
  RFL <- cols_ab["RFL"]
  RID <- cols_ab["RID"]
  RIF <- cols_ab["RIF"]
  ROK <- cols_ab["ROK"]
  RST <- cols_ab["RST"]
  RXT <- cols_ab["RXT"]
  SAM <- cols_ab["SAM"]
  SBC <- cols_ab["SBC"]
  SDI <- cols_ab["SDI"]
  SDM <- cols_ab["SDM"]
  SIS <- cols_ab["SIS"]
  SLF <- cols_ab["SLF"]
  SLF1 <- cols_ab["SLF1"]
  SLF10 <- cols_ab["SLF10"]
  SLF11 <- cols_ab["SLF11"]
  SLF12 <- cols_ab["SLF12"]
  SLF13 <- cols_ab["SLF13"]
  SLF2 <- cols_ab["SLF2"]
  SLF3 <- cols_ab["SLF3"]
  SLF4 <- cols_ab["SLF4"]
  SLF5 <- cols_ab["SLF5"]
  SLF6 <- cols_ab["SLF6"]
  SLF7 <- cols_ab["SLF7"]
  SLF8 <- cols_ab["SLF8"]
  SLF9 <- cols_ab["SLF9"]
  SLT1 <- cols_ab["SLT1"]
  SLT2 <- cols_ab["SLT2"]
  SLT3 <- cols_ab["SLT3"]
  SLT4 <- cols_ab["SLT4"]
  SLT5 <- cols_ab["SLT5"]
  SMX <- cols_ab["SMX"]
  SPI <- cols_ab["SPI"]
  SPX <- cols_ab["SPX"]
  STR <- cols_ab["STR"]
  STR1 <- cols_ab["STR1"]
  SUD <- cols_ab["SUD"]
  SUT <- cols_ab["SUT"]
  SXT <- cols_ab["SXT"]
  SZO <- cols_ab["SZO"]
  TAL <- cols_ab["TAL"]
  TCC <- cols_ab["TCC"]
  TCM <- cols_ab["TCM"]
  TCY <- cols_ab["TCY"]
  TEC <- cols_ab["TEC"]
  TEM <- cols_ab["TEM"]
  TGC <- cols_ab["TGC"]
  THA <- cols_ab["THA"]
  TIC <- cols_ab["TIC"]
  TIO <- cols_ab["TIO"]
  TLT <- cols_ab["TLT"]
  TLV <- cols_ab["TLV"]
  TMP <- cols_ab["TMP"]
  TMX <- cols_ab["TMX"]
  TOB <- cols_ab["TOB"]
  TRL <- cols_ab["TRL"]
  TVA <- cols_ab["TVA"]
  TZD <- cols_ab["TZD"]
  TZP <- cols_ab["TZP"]
  VAN <- cols_ab["VAN"]
  
  ab_missing <- function(ab) {
    all(ab %in% c(NULL, NA))
  }
  
  if (ab_missing(AMP) & !ab_missing(AMX)) {
    # ampicillin column is missing, but amoxicillin is available
    if (info == TRUE) {
      message_("Using column '", font_bold(AMX), "' as input for ampicillin since many EUCAST rules depend on it.")
    }
    AMP <- AMX
  }
  
  # data preparation ----
  if (info == TRUE & NROW(x) > 10000) {
    message_("Preparing data...", appendLF = FALSE, as_note = FALSE)
  }
  
  # nolint start
  # antibiotic classes ----
  aminoglycosides <- c(AMK, DKB, GEN, ISE, KAN, NEO, NET, RST, SIS, STR, STR1, TOB)
  aminopenicillins <- c(AMP, AMX)
  carbapenems <- c(DOR, ETP, IPM, MEM, MEV)
  cephalosporins <- c(CDZ, CCP, CAC, CEC, CFR, RID, MAN, CTZ, CZD, CZO, CDR, DIT, FEP, CAT, CFM, CMX, CMZ, DIZ, CID, CFP, CSL, CND, CTX, CTT, CTF, FOX, CPM, CPO, CPD, CPR, CRD, CFS, CPT, CAZ, CCV, CTL, CTB, CZX, BPR, CFM1, CEI, CRO, CXM, LEX, CEP, HAP, CED, LTM, LOR)
  cephalosporins_1st <- c(CAC, CFR, RID, CTZ, CZD, CZO, CRD, CTL, LEX, CEP, HAP, CED)
  cephalosporins_2nd <- c(CEC, MAN, CMZ, CID, CND, CTT, CTF, FOX, CPR, CXM, LOR)
  cephalosporins_3rd <- c(CDZ, CCP, CCX, CDR, DIT, DIX, CAT, CPI, CFM, CMX, DIZ, CFP, CSL, CTX, CTC, CTS, CHE, FOV, CFZ, CPM, CPD, CPX, CDC, CFS, CAZ, CZA, CCV, CEM, CPL, CTB, TIO, CZX, CZP, CRO, LTM)
  cephalosporins_except_CAZ <- cephalosporins[cephalosporins != ifelse(is.null(CAZ), "", CAZ)]
  fluoroquinolones <- c(CIP, ENX, FLE, GAT, GEM, GRX, LVX, LOM, MFX, NOR, OFX, PAZ, PEF, PRU, RFL, SPX, TMX, TVA)
  glycopeptides <- c(AVO, NVA, RAM, TEC, TCM, VAN) # dalba/orita/tela are in lipoglycopeptides
  lincosamides <- c(CLI, LIN, PRL)
  lipoglycopeptides <- c(DAL, ORI, TLV)
  macrolides <- c(AZM, CLR, DIR, ERY, FLR1, JOS, MID, MCM, OLE, ROK, RXT, SPI, TLT, TRL)
  oxazolidinones <- c(CYC, LNZ, THA, TZD)
  polymyxins <- c(PLB, COL)
  streptogramins <- c(QDA, PRI)
  tetracyclines <- c(DOX, MNO, TCY) # since EUCAST v3.1 tigecycline (TGC) is set apart
  ureidopenicillins <- c(PIP, TZP, AZL, MEZ)
  all_betalactams <- c(aminopenicillins, cephalosporins, carbapenems, ureidopenicillins, AMC, OXA, FLC, PEN)
  # nolint end
  
  # Some helper functions ---------------------------------------------------
  get_antibiotic_columns <- function(x, df) {
    x <- trimws(unlist(strsplit(x, ",", fixed = TRUE)))
    y <- character(0)
    for (i in seq_len(length(x))) {
      if (is.function(get(x[i]))) {
        stop("Column ", x[i], " is also a function. Please create an issue on github.com/msberends/AMR/issues.")
      }
      y <- c(y, tryCatch(get(x[i]), error = function(e) ""))
    }
    y[y != "" & y %in% colnames(df)]
  }
  markup_italics_where_needed <- function(x) {
    # returns names found in family, genus or species as italics
    if (!has_colour()) {
      return(x)
    }
    x <- unlist(strsplit(x, " "))
    ind <- gsub("[)(:]", "", x) %in% c(MO_lookup[which(MO_lookup$rank %in% c("family", "genus")), ]$fullname,
                                       MO_lookup[which(MO_lookup$rank == "species"), ]$species)
    x[ind] <- font_italic(x[ind], collapse = NULL)
    paste(x, collapse = " ")
  }
  get_antibiotic_names <- function(x) {
    x <- x %pm>%
      strsplit(",") %pm>%
      unlist() %pm>%
      trimws() %pm>%
      vapply(FUN.VALUE = character(1), function(x) if (x %in% antibiotics$ab) ab_name(x, language = NULL, tolower = TRUE) else x) %pm>%
      sort() %pm>%
      paste(collapse = ", ")
    x <- gsub("_", " ", x, fixed = TRUE)
    x <- gsub("except CAZ", paste("except", ab_name("CAZ", language = NULL, tolower = TRUE)), x, fixed = TRUE)
    x <- gsub("cephalosporins (1st|2nd|3rd|4th|5th)", "cephalosporins (\\1 gen.)", x)
    x
  }
  format_antibiotic_names <- function(ab_names, ab_results) {
    ab_names <- trimws(unlist(strsplit(ab_names, ",")))
    ab_results <- trimws(unlist(strsplit(ab_results, ",")))
    if (length(ab_results) == 1) {
      if (length(ab_names) == 1) {
        # like FOX S
        x <- paste(ab_names, "is")
      } else if (length(ab_names) == 2) {
        # like PEN,FOX S
        x <- paste(paste0(ab_names, collapse = " and "), "are both")
      } else {
        # like PEN,FOX,GEN S (although dependency on > 2 ABx does not exist at the moment)
        # nolint start
        # x <- paste(paste0(ab_names, collapse = " and "), "are all")
        # nolint end
      }
      return(paste0(x, " '", ab_results, "'"))
    } else {
      if (length(ab_names) == 2) {
        # like PEN,FOX S,R
        paste0(ab_names[1], " is '", ab_results[1], "' and ", 
               ab_names[2], " is '", ab_results[2], "'")
      } else {
        # like PEN,FOX,GEN S,R,R (although dependency on > 2 ABx does not exist at the moment)
        paste0(ab_names[1], " is '", ab_results[1], "' and ", 
               ab_names[2], " is '", ab_results[2], "' and ", 
               ab_names[3], " is '", ab_results[3], "'")
      }
    }
  }
  as.rsi_no_warning <- function(x) {
    if (is.rsi(x)) {
      return(x)
    }
    suppressWarnings(as.rsi(x))
  }
  
  # Preparing the data ------------------------------------------------------
  
  verbose_info <- data.frame(rowid = character(0),
                             col = character(0),
                             mo_fullname = character(0),
                             old = as.rsi(character(0)),
                             new = as.rsi(character(0)),
                             rule = character(0),
                             rule_group = character(0),
                             rule_name = character(0),
                             rule_source = character(0),
                             stringsAsFactors = FALSE)
  
  old_cols <- colnames(x)
  old_attributes <- attributes(x)
  x <- as.data.frame(x, stringsAsFactors = FALSE) # no tibbles, data.tables, etc.
  rownames(x) <- NULL # will later be restored with old_attributes
  # create unique row IDs - combination of the MO and all ABx columns (so they will only run once per unique combination)
  x$`.rowid` <- vapply(FUN.VALUE = character(1),
                       as.list(as.data.frame(t(x[, c(col_mo, cols_ab), drop = FALSE]),
                                             stringsAsFactors = FALSE)),
                       function(x) {
                         x[is.na(x)] <- "."
                         paste0(x, collapse = "")
                       })
  
  # save original table, with the new .rowid column
  x.bak <- x
  # keep only unique rows for MO and ABx
  x <- x %pm>% 
    pm_arrange(`.rowid`) %pm>% 
    # big speed gain! only analyse unique rows:
    pm_distinct(`.rowid`, .keep_all = TRUE) %pm>% 
    as.data.frame(stringsAsFactors = FALSE)
  x[, col_mo] <- as.mo(as.character(x[, col_mo, drop = TRUE]))
  x <- x %pm>%
    left_join_microorganisms(by = col_mo, suffix = c("_oldcols", ""))
  x$gramstain <- mo_gramstain(x[, col_mo, drop = TRUE], language = NULL)
  x$genus_species <- paste(x$genus, x$species)
  if (info == TRUE & NROW(x) > 10000) {
    message_(" OK.", add_fn = list(font_green, font_bold), as_note = FALSE)
  }
  
  if (any(x$genus == "Staphylococcus", na.rm = TRUE)) {
    all_staph <- MO_lookup[which(MO_lookup$genus == "Staphylococcus"), ]
    all_staph$CNS_CPS <- suppressWarnings(mo_name(all_staph$mo, Becker = "all", language = NULL))
  }
  if (any(x$genus == "Streptococcus", na.rm = TRUE)) {
    all_strep <- MO_lookup[which(MO_lookup$genus == "Streptococcus"), ]
    all_strep$Lancefield <- suppressWarnings(mo_name(all_strep$mo, Lancefield = TRUE, language = NULL))
  }
  
  n_added <- 0
  n_changed <- 0
  
  # Other rules: enzyme inhibitors ------------------------------------------
  if (any(c("all", "other") %in% rules)) {
    if (info == TRUE) {
      cat("\n")
      cat(word_wrap(
        font_bold(paste0("Rules by this AMR package (",
                         font_red(paste0("v", utils::packageDescription("AMR")$Version, ", ", 
                                         format(as.Date(utils::packageDescription("AMR")$Date), format = "%Y"))), "), see ?eucast_rules\n"))))
    }
    
    ab_enzyme <- subset(antibiotics, name %like% "/")[, c("ab", "name")]
    ab_enzyme$base_name <- gsub("^([a-zA-Z0-9]+).*", "\\1", ab_enzyme$name)
    ab_enzyme$base_ab <- as.ab(ab_enzyme$base_name)
    for (i in seq_len(nrow(ab_enzyme))) {
      if (all(c(ab_enzyme[i, ]$ab, ab_enzyme[i, ]$base_ab) %in% names(cols_ab), na.rm = TRUE)) {
        ab_name_base <- ab_name(cols_ab[ab_enzyme[i, ]$base_ab], language = NULL, tolower = TRUE)
        ab_name_enzyme <- ab_name(cols_ab[ab_enzyme[i, ]$ab], language = NULL, tolower = TRUE)
        
        # Set base to R where base + enzyme inhibitor is R ----
        rule_current <- paste0("Set ", ab_name_base, " (", cols_ab[ab_enzyme[i, ]$base_ab], ") = R where ",
                               ab_name_enzyme, " (", cols_ab[ab_enzyme[i, ]$ab], ") = R")
        if (info == TRUE) {
          cat(word_wrap(rule_current))
          cat("\n")
        }
        run_changes <- edit_rsi(x = x,
                                col_mo = col_mo,
                                to = "R",
                                rule = c(rule_current, "Other rules", "",
                                         paste0("Non-EUCAST: AMR package v", utils::packageDescription("AMR")$Version)),
                                rows = which(as.rsi_no_warning(x[, cols_ab[ab_enzyme[i, ]$ab]]) == "R"),
                                cols = cols_ab[ab_enzyme[i, ]$base_ab],
                                last_verbose_info = verbose_info,
                                original_data = x.bak,
                                warned = warned,
                                info = info)
        n_added <- n_added + run_changes$added
        n_changed <- n_changed + run_changes$changed
        verbose_info <- run_changes$verbose_info
        x <- run_changes$output
        warn_lacking_rsi_class <- c(warn_lacking_rsi_class, run_changes$rsi_warn)
        # Print number of new changes
        if (info == TRUE) {
          # print only on last one of rules in this group
          txt_ok(n_added = n_added, n_changed = n_changed, warned = warned)
          # and reset counters
          n_added <- 0
          n_changed <- 0
        }
        
        # Set base + enzyme inhibitor to S where base is S ----
        rule_current <- paste0("Set ", ab_name_enzyme, " (", cols_ab[ab_enzyme[i, ]$ab], ") = S where ",
                               ab_name_base, " (", cols_ab[ab_enzyme[i, ]$base_ab], ") = S")
        if (info == TRUE) {
          cat(word_wrap(rule_current))
          cat("\n")
        }
        run_changes <- edit_rsi(x = x,
                                col_mo = col_mo,
                                to = "S",
                                rule = c(rule_current, "Other rules", "", 
                                         paste0("Non-EUCAST: AMR package v", utils::packageDescription("AMR")$Version)),
                                rows = which(as.rsi_no_warning(x[, cols_ab[ab_enzyme[i, ]$base_ab]]) == "S"),
                                cols = cols_ab[ab_enzyme[i, ]$ab],
                                last_verbose_info = verbose_info,
                                original_data = x.bak,
                                warned = warned,
                                info = info)
        n_added <- n_added + run_changes$added
        n_changed <- n_changed + run_changes$changed
        verbose_info <- run_changes$verbose_info
        x <- run_changes$output
        warn_lacking_rsi_class <- c(warn_lacking_rsi_class, run_changes$rsi_warn)
        # Print number of new changes
        if (info == TRUE) {
          # print only on last one of rules in this group
          txt_ok(n_added = n_added, n_changed = n_changed, warned = warned)
          # and reset counters
          n_added <- 0
          n_changed <- 0
        }
      }
    }
    
  } else {
    if (info == TRUE) {
      message_("\n\nSkipping inheritance rules defined by this package, such as setting trimethoprim (TMP) = R where trimethoprim/sulfamethoxazole (SXT) = R.",
               as_note = FALSE,
               add_fn = font_red)
      message_("Use eucast_rules(..., rules = \"all\") to also apply those rules.",
               as_note = FALSE,
               add_fn = font_red)
    }
  }
  
  # Official EUCAST rules ---------------------------------------------------
  eucast_notification_shown <- FALSE
  if (!is.null(list(...)$eucast_rules_df)) {
    # this allows: eucast_rules(x, eucast_rules_df = AMR:::eucast_rules_file %>% filter(is.na(have_these_values)))
    eucast_rules_df <- list(...)$eucast_rules_df
  } else {
    # otherwise internal data file, created in data-raw/internals.R
    eucast_rules_df <- eucast_rules_file
  }
  
  # filter on user-set guideline versions ----
  if (any(c("all", "breakpoints") %in% rules)) {
    eucast_rules_df <- subset(eucast_rules_df,
                              !reference.rule_group %like% "breakpoint" |
                                (reference.rule_group %like% "breakpoint" & reference.version == version_breakpoints))
  }
  if (any(c("all", "expert") %in% rules)) {
    eucast_rules_df <- subset(eucast_rules_df,
                              !reference.rule_group %like% "expert" |
                                (reference.rule_group %like% "expert" & reference.version == version_expertrules))
  }
  # filter out AmpC de-repressed cephalosporin-resistant mutants ----
  if (is.null(ampc_cephalosporin_resistance)) {
    eucast_rules_df <- subset(eucast_rules_df,
                              !reference.rule %like% "ampc")
  } else {
    eucast_rules_df[which(eucast_rules_df$reference.rule %like% "ampc"), "to_value"] <- as.character(ampc_cephalosporin_resistance)
  }
  
  for (i in seq_len(nrow(eucast_rules_df))) {
    
    rule_previous <- eucast_rules_df[max(1, i - 1), "reference.rule", drop = TRUE]
    rule_current <- eucast_rules_df[i, "reference.rule", drop = TRUE]
    rule_next <- eucast_rules_df[min(nrow(eucast_rules_df), i + 1), "reference.rule", drop = TRUE]
    rule_group_previous <- eucast_rules_df[max(1, i - 1), "reference.rule_group", drop = TRUE]
    rule_group_current <- eucast_rules_df[i, "reference.rule_group", drop = TRUE]
    if (isFALSE(info) | isFALSE(verbose)) {
      rule_text <- ""
    } else {
      if (is.na(eucast_rules_df[i, "and_these_antibiotics", drop = TRUE])) {
        rule_text <- paste0("always report as '", eucast_rules_df[i, "to_value", drop = TRUE], "': ", get_antibiotic_names(eucast_rules_df[i, "then_change_these_antibiotics", drop = TRUE]))
      } else {
        rule_text <- paste0("report as '", eucast_rules_df[i, "to_value", drop = TRUE], "' when ",
                            format_antibiotic_names(ab_names = get_antibiotic_names(eucast_rules_df[i, "and_these_antibiotics", drop = TRUE]),
                                                    ab_results = eucast_rules_df[i, "have_these_values", drop = TRUE]), ": ",
                            get_antibiotic_names(eucast_rules_df[i, "then_change_these_antibiotics", drop = TRUE]))
      }
    }
    if (i == 1) {
      rule_previous <- ""
      rule_group_previous <- ""
    }
    if (i == nrow(eucast_rules_df)) {
      rule_next <- ""
    }
    
    # don't apply rules if user doesn't want to apply them
    if (rule_group_current %like% "breakpoint" & !any(c("all", "breakpoints") %in% rules)) {
      next
    }
    if (rule_group_current %like% "expert" & !any(c("all", "expert") %in% rules)) {
      next
    }
    
    if (info == TRUE) {
      # Print EUCAST intro ------------------------------------------------------
      if (!rule_group_current %like% "other" & eucast_notification_shown == FALSE) {
        cat(
          paste0("\n", font_grey(strrep("-", 0.95 * options()$width)), "\n",
                 word_wrap("Rules by the ", font_bold("European Committee on Antimicrobial Susceptibility Testing (EUCAST)")), "\n", 
                 font_blue("https://eucast.org/"), "\n"))
        eucast_notification_shown <- TRUE
      }
      
      # Print rule (group) ------------------------------------------------------
      if (rule_group_current != rule_group_previous) {
        # is new rule group, one of Breakpoints, Expert Rules and Other
        cat(font_bold(
          ifelse(
            rule_group_current %like% "breakpoint",
            paste0("\n", 
                   word_wrap(
                     breakpoints_info$title, " (",
                     font_red(paste0(breakpoints_info$version_txt, ", ", breakpoints_info$year)), ")\n")),
            ifelse(
              rule_group_current %like% "expert",
              paste0("\n", 
                     word_wrap(
                       expertrules_info$title, " (",
                       font_red(paste0(expertrules_info$version_txt, ", ", expertrules_info$year)), ")\n")),
              ""))), "\n")
      }
      # Print rule  -------------------------------------------------------------
      if (rule_current != rule_previous) {
        # is new rule within group, print its name
        cat(markup_italics_where_needed(word_wrap(rule_current, 
                                                  width = getOption("width") - 30,
                                                  extra_indent = 4)))
        warned <- FALSE
      }
    }
    
    # Get rule from file ------------------------------------------------------
    if_mo_property <- trimws(eucast_rules_df[i, "if_mo_property", drop = TRUE])
    like_is_one_of <- trimws(eucast_rules_df[i, "like.is.one_of", drop = TRUE])
    mo_value <- trimws(eucast_rules_df[i, "this_value", drop = TRUE])
    
    # be sure to comprise all coagulase-negative/-positive staphylococci when they are mentioned
    if (mo_value %like% "coagulase" && any(x$genus == "Staphylococcus", na.rm = TRUE)) {
      if (mo_value %like% "negative") {
        eucast_rules_df[i, "this_value"] <- paste0("^(", paste0(all_staph[which(all_staph$CNS_CPS %like% "negative"),
                                                                          "fullname", 
                                                                          drop = TRUE],
                                                                collapse = "|"),
                                                   ")$")
      } else {
        eucast_rules_df[i, "this_value"] <- paste0("^(", paste0(all_staph[which(all_staph$CNS_CPS %like% "positive"),
                                                                          "fullname", 
                                                                          drop = TRUE],
                                                                collapse = "|"),
                                                   ")$")
      }
      like_is_one_of <- "like"
    }
    # be sure to comprise all beta-haemolytic Streptococci (Lancefield groups A, B, C and G) when they are mentioned
    if (mo_value %like% "group [ABCG]" && any(x$genus == "Streptococcus", na.rm = TRUE)) {
      eucast_rules_df[i, "this_value"] <- paste0("^(", paste0(all_strep[which(all_strep$Lancefield %like% "group [ABCG]"),
                                                                        "fullname", 
                                                                        drop = TRUE],
                                                              collapse = "|"),
                                                 ")$")
      like_is_one_of <- "like"
    }
    
    if (like_is_one_of == "is") {
      # so e.g. 'Enterococcus' will turn into '^Enterococcus$'
      mo_value <- paste0("^", mo_value, "$")
    } else if (like_is_one_of == "one_of") {
      # so 'Clostridium, Actinomyces, ...' will turn into '^(Clostridium|Actinomyces|...)$'
      mo_value <- paste0("^(",
                         paste(trimws(unlist(strsplit(mo_value, ",", fixed = TRUE))),
                               collapse = "|"),
                         ")$")
    } else if (like_is_one_of != "like") {
      stop("invalid value for column 'like.is.one_of'", call. = FALSE)
    }
    
    source_antibiotics <- eucast_rules_df[i, "and_these_antibiotics", drop = TRUE]
    source_value <- trimws(unlist(strsplit(eucast_rules_df[i, "have_these_values", drop = TRUE], ",", fixed = TRUE)))
    target_antibiotics <- eucast_rules_df[i, "then_change_these_antibiotics", drop = TRUE]
    target_value <- eucast_rules_df[i, "to_value", drop = TRUE]
    
    if (is.na(source_antibiotics)) {
      rows <- tryCatch(which(x[, if_mo_property, drop = TRUE] %like_perl% mo_value),
                       error = function(e) integer(0))
    } else {
      source_antibiotics <- get_antibiotic_columns(source_antibiotics, x)
      if (length(source_value) == 1 & length(source_antibiotics) > 1) {
        source_value <- rep(source_value, length(source_antibiotics))
      }
      if (length(source_antibiotics) == 0) {
        rows <- integer(0)
      } else if (length(source_antibiotics) == 1) {
        rows <-  tryCatch(which(x[, if_mo_property, drop = TRUE] %like_perl% mo_value
                                & as.rsi_no_warning(x[, source_antibiotics[1L]]) == source_value[1L]),
                          error = function(e) integer(0))
      } else if (length(source_antibiotics) == 2) {
        rows <-  tryCatch(which(x[, if_mo_property, drop = TRUE] %like_perl% mo_value
                                & as.rsi_no_warning(x[, source_antibiotics[1L]]) == source_value[1L]
                                & as.rsi_no_warning(x[, source_antibiotics[2L]]) == source_value[2L]),
                          error = function(e) integer(0))
        # nolint start
      # } else if (length(source_antibiotics) == 3) {
      #   rows <-  tryCatch(which(x[, if_mo_property, drop = TRUE] %like_perl% mo_value
      #                           & as.rsi_no_warning(x[, source_antibiotics[1L]]) == source_value[1L]
      #                           & as.rsi_no_warning(x[, source_antibiotics[2L]]) == source_value[2L]
      #                           & as.rsi_no_warning(x[, source_antibiotics[3L]]) == source_value[3L]),
      #                     error = function(e) integer(0))
        # nolint end
      } else {
        stop_("only 2 antibiotics supported for source_antibiotics")
      }
    }
    
    cols <- get_antibiotic_columns(target_antibiotics, x)
    
    # Apply rule on data ------------------------------------------------------
    # this will return the unique number of changes
    run_changes <- edit_rsi(x = x,
                            col_mo = col_mo,
                            to = target_value,
                            rule = c(rule_text, rule_group_current, rule_current, 
                                     ifelse(rule_group_current %like% "breakpoint",
                                            paste0(breakpoints_info$title, " ", breakpoints_info$version_txt, ", ", breakpoints_info$year),
                                            paste0(expertrules_info$title, " ", expertrules_info$version_txt, ", ", expertrules_info$year))),
                            rows = rows,
                            cols = cols,
                            last_verbose_info = verbose_info,
                            original_data = x.bak,
                            warned = warned,
                            info = info)
    n_added <- n_added + run_changes$added
    n_changed <- n_changed + run_changes$changed
    verbose_info <- run_changes$verbose_info
    x <- run_changes$output
    warn_lacking_rsi_class <- c(warn_lacking_rsi_class, run_changes$rsi_warn)
    # Print number of new changes ---------------------------------------------
    if (info == TRUE & rule_next != rule_current) {
      # print only on last one of rules in this group
      txt_ok(n_added = n_added, n_changed = n_changed, warned = warned)
      # and reset counters
      n_added <- 0
      n_changed <- 0
    }
  }
  
  # Print overview ----------------------------------------------------------
  if (info == TRUE | verbose == TRUE) {
    verbose_info <- x.bak %pm>%
      pm_mutate(row = pm_row_number()) %pm>%
      pm_select(`.rowid`, row) %pm>%
      pm_right_join(verbose_info,
                    by = c(".rowid" = "rowid")) %pm>% 
      pm_select(-`.rowid`) %pm>% 
      pm_select(row, pm_everything()) %pm>% 
      pm_filter(!is.na(new) | is.na(new) & !is.na(old)) %pm>%
      pm_arrange(row, rule_group, rule_name, col)
    rownames(verbose_info) <- NULL
  }
  
  if (info == TRUE) {
    
    if (verbose == TRUE) {
      wouldve <- "would have "
    } else {
      wouldve <- ""
    }
    
    cat(paste0("\n", font_grey(strrep("-", 0.95 * options()$width)), "\n"))
    cat(word_wrap(paste0("The rules ", paste0(wouldve, "affected "),
                         font_bold(formatnr(pm_n_distinct(verbose_info$row)),
                                   "out of", formatnr(nrow(x.bak)),
                                   "rows"), 
                         ", making a total of ",
                         font_bold(formatnr(nrow(verbose_info)), "edits\n"))))
    
    total_n_added <- verbose_info %pm>% pm_filter(is.na(old)) %pm>% nrow()
    total_n_changed <- verbose_info %pm>% pm_filter(!is.na(old)) %pm>% nrow()
    
    # print added values
    if (total_n_added == 0) {
      colour <- cat # is function
    } else {
      colour <- font_green # is function
    }
    cat(colour(paste0("=> ", wouldve, "added ",
                      font_bold(formatnr(verbose_info %pm>%
                                           pm_filter(is.na(old)) %pm>%
                                           nrow()), "test results"),
                      "\n")))
    if (total_n_added > 0) {
      added_summary <- verbose_info %pm>%
        pm_filter(is.na(old)) %pm>%
        pm_count(new, name = "n")
      cat(paste("   -", 
                paste0(formatnr(added_summary$n), " test result", ifelse(added_summary$n > 1, "s", ""), 
                       " added as ", paste0('"', added_summary$new, '"')), collapse = "\n"))
    }
    
    # print changed values
    if (total_n_changed == 0) {
      colour <- cat # is function
    } else {
      colour <- font_blue # is function
    }
    if (total_n_added + total_n_changed > 0) {
      cat("\n")
    }
    cat(colour(paste0("=> ", wouldve, "changed ",
                      font_bold(formatnr(verbose_info %pm>%
                                           pm_filter(!is.na(old)) %pm>%
                                           nrow()), "test results"),
                      "\n")))
    if (total_n_changed > 0) {
      changed_summary <- verbose_info %pm>%
        pm_filter(!is.na(old)) %pm>%
        pm_mutate(new = ifelse(is.na(new), "NA", new)) %pm>%
        pm_count(old, new, name = "n")
      cat(paste("   -", 
                paste0(formatnr(changed_summary$n), " test result", ifelse(changed_summary$n > 1, "s", ""), " changed from ", 
                       paste0('"', changed_summary$old, '"'), " to ", paste0('"', changed_summary$new, '"')), collapse = "\n"))
      cat("\n")
    }
    
    cat(paste0(font_grey(strrep("-", 0.95 * options()$width)), "\n"))
    
    if (verbose == FALSE & total_n_added + total_n_changed > 0) {
      cat("\n", word_wrap("Use ", font_bold("eucast_rules(..., verbose = TRUE)"), " (on your original data) to get a data.frame with all specified edits instead."), "\n\n", sep = "")
    } else if (verbose == TRUE) {
      cat("\n", word_wrap("Used 'Verbose mode' (", font_bold("verbose = TRUE"), "), which returns a data.frame with all specified edits.\nUse ", font_bold("verbose = FALSE"), " to apply the rules on your data."), "\n\n", sep = "")
    }
  }
  
  if (length(warn_lacking_rsi_class) > 0) {
    warn_lacking_rsi_class <- unique(warn_lacking_rsi_class)
    warning_("Not all columns with antimicrobial results are of class <rsi>. Transform them on beforehand, with e.g.:\n",
             "  ", x_deparsed, " %>% mutate_if(is.rsi.eligible, as.rsi)\n",
             "  ", x_deparsed, " %>% as.rsi(", ifelse(length(warn_lacking_rsi_class) == 1, 
                                                      warn_lacking_rsi_class,
                                                      paste0(warn_lacking_rsi_class[1], ":", warn_lacking_rsi_class[length(warn_lacking_rsi_class)])), 
             ")",
             call = FALSE)
  }
  
  # Return data set ---------------------------------------------------------
  if (verbose == TRUE) {
    verbose_info
  } else {
    # x was analysed with only unique rows, so join everything together again
    x <- x[, c(cols_ab, ".rowid"), drop = FALSE]
    x.bak <- x.bak[, setdiff(colnames(x.bak), cols_ab), drop = FALSE]
    x.bak <- x.bak %pm>% 
      pm_left_join(x, by = ".rowid")
    x.bak <- x.bak[, old_cols, drop = FALSE]
    # reset original attributes
    attributes(x.bak) <- old_attributes
    x.bak
  }
}

# helper function for editing the table ----
edit_rsi <- function(x, 
                     col_mo,
                     to, 
                     rule, 
                     rows,
                     cols,
                     last_verbose_info, 
                     original_data,
                     warned,
                     info) {
  cols <- unique(cols[!is.na(cols) & !is.null(cols)])
  
  # for Verbose Mode, keep track of all changes and return them
  track_changes <- list(added = 0,
                        changed = 0,
                        output = x,
                        verbose_info = last_verbose_info,
                        rsi_warn = character(0))
  
  txt_error <- function() {
    if (info == TRUE) cat("", font_red_bg(font_white(" ERROR ")), "\n\n") 
  }
  txt_warning <- function() {
    if (warned == FALSE) {
      if (info == TRUE) cat("", font_yellow_bg(font_black(" WARNING ")))
    }
    warned <<- TRUE 
  }
  
  if (length(rows) > 0 & length(cols) > 0) {
    new_edits <- x
    if (any(!vapply(FUN.VALUE = logical(1), x[, cols, drop = FALSE], is.rsi), na.rm = TRUE)) {
      track_changes$rsi_warn <- cols[!vapply(FUN.VALUE = logical(1), x[, cols, drop = FALSE], is.rsi)]
    }
    tryCatch(
      # insert into original table
      new_edits[rows, cols] <- to,
      warning = function(w) {
        if (w$message %like% "invalid factor level") {
          xyz <- vapply(FUN.VALUE = logical(1), cols, function(col) {
            new_edits[, col] <<- factor(x = as.character(pm_pull(new_edits, col)),
                                        levels = unique(c(to, levels(pm_pull(new_edits, col)))))
            TRUE
          })
          suppressWarnings(new_edits[rows, cols] <<- to)
          warning_('Value "', to, '" added to the factor levels of column(s) `', paste(cols, collapse = "`, `"), "` because this value was not an existing factor level. A better way is to use as.rsi() on beforehand on antimicrobial columns to guarantee the right structure.", call = FALSE)
          txt_warning()
          warned <- FALSE
        } else {
          warning_(w$message, call = FALSE)
          txt_warning()
          cat("\n") # txt_warning() does not append a "\n" on itself
        }
      },
      error = function(e) {
        txt_error()
        stop(paste0("In row(s) ", paste(rows[1:min(length(rows), 10)], collapse = ","), 
                    ifelse(length(rows) > 10, "...", ""),
                    " while writing value '", to, 
                    "' to column(s) `", paste(cols, collapse = "`, `"), 
                    "`:\n", e$message),
             call. = FALSE)
      }
    )
    
    track_changes$output <- new_edits
    if (isTRUE(info) && !isTRUE(all.equal(x, track_changes$output))) {
      get_original_rows <- function(rowids) {
        as.integer(rownames(original_data[which(original_data$.rowid %in% rowids), , drop = FALSE]))
      }
      for (i in seq_len(length(cols))) {
        verbose_new <- data.frame(rowid = new_edits[rows, ".rowid", drop = TRUE],
                                  col = cols[i],
                                  mo_fullname = new_edits[rows, "fullname", drop = TRUE],
                                  old = x[rows, cols[i], drop = TRUE],
                                  new = to,
                                  rule = font_stripstyle(rule[1]),
                                  rule_group = font_stripstyle(rule[2]),
                                  rule_name = font_stripstyle(rule[3]),
                                  rule_source = font_stripstyle(rule[4]),
                                  stringsAsFactors = FALSE)
        colnames(verbose_new) <- c("rowid", "col", "mo_fullname", "old", "new",
                                   "rule", "rule_group", "rule_name", "rule_source")
        verbose_new <- verbose_new %pm>% pm_filter(old != new | is.na(old) | is.na(new) & !is.na(old))
        # save changes to data set 'verbose_info'
        track_changes$verbose_info <- rbind(track_changes$verbose_info,
                                            verbose_new,
                                            stringsAsFactors = FALSE)
        # count adds and changes
        track_changes$added <- track_changes$added + verbose_new %pm>%
          pm_filter(is.na(old)) %pm>%
          pm_pull(rowid) %pm>% 
          get_original_rows() %pm>% 
          length()
        track_changes$changed <- track_changes$changed + verbose_new %pm>%
          pm_filter(!is.na(old)) %pm>%
          pm_pull(rowid) %pm>% 
          get_original_rows() %pm>% 
          length()
      }
    }
  }
  return(track_changes)
}
```
