## Supplementary material for "AMR - An R Package for Working with Antimicrobial Resistance Data": Source package: filter_ab_class.R

```
# ==================================================================== #
# TITLE                                                                #
# Antimicrobial Resistance (AMR) Analysis for R                        #
#                                                                      #
# SOURCE                                                               #
# https://github.com/msberends/AMR                                     #
#                                                                      #
# LICENCE                                                              #
# (c) 2018-2021 Berends MS, Luz CF et al.                              #
# Developed at the University of Groningen, the Netherlands, in        #
# collaboration with non-profit organisations Certe Medical            #
# Diagnostics & Advice, and University Medical Center Groningen.       # 
#                                                                      #
# This R package is free software; you can freely use and distribute   #
# it for both personal and commercial purposes under the terms of the  #
# GNU General Public License version 2.0 (GNU GPL-2), as published by  #
# the Free Software Foundation.                                        #
# We created this package for both routine data analysis and academic  #
# research and it was publicly released in the hope that it will be    #
# useful, but it comes WITHOUT ANY WARRANTY OR LIABILITY.              #
#                                                                      #
# Visit our website for the full manual and a complete tutorial about  #
# how to conduct AMR analysis: https://msberends.github.io/AMR/        #
# ==================================================================== #

#' Filter isolates on result in antimicrobial class
#'
#' Filter isolates on results in specific antimicrobial classes. This makes it easy to filter on isolates that were tested for e.g. any aminoglycoside, or to filter on carbapenem-resistant isolates without the need to specify the drugs.
#' @inheritSection lifecycle Stable lifecycle
#' @param x a data set
#' @param ab_class an antimicrobial class, like `"carbapenems"`. The columns `group`, `atc_group1` and `atc_group2` of the [antibiotics] data set will be searched (case-insensitive) for this value.
#' @param result an antibiotic result: S, I or R (or a combination of more of them)
#' @param scope the scope to check which variables to check, can be `"any"` (default) or `"all"`
#' @param ... previously used when this package still depended on the `dplyr` package, now ignored
#' @details All columns of `x` will be searched for known antibiotic names, abbreviations, brand names and codes (ATC, EARS-Net, WHO, etc.). This means that a filter function like e.g. [filter_aminoglycosides()] will include column names like 'gen', 'genta', 'J01GB03', 'tobra', 'Tobracin', etc.
#' @rdname filter_ab_class
#' @seealso [antibiotic_class_selectors()] for the `select()` equivalent.
#' @export
#' @examples
#' filter_aminoglycosides(example_isolates)
#' 
#' \donttest{
#' if (require("dplyr")) {
#'
#'   # filter on isolates that have any result for any aminoglycoside
#'   example_isolates %>% filter_aminoglycosides()
#'   example_isolates %>% filter_ab_class("aminoglycoside")
#' 
#'   # this is essentially the same as (but without determination of column names):
#'   example_isolates %>%
#'     filter_at(.vars = vars(c("GEN", "TOB", "AMK", "KAN")),
#'               .vars_predicate = any_vars(. %in% c("S", "I", "R")))
#'
#'
#'   # filter on isolates that show resistance to ANY aminoglycoside
#'   example_isolates %>% filter_aminoglycosides("R", "any")
#'  
#'   # filter on isolates that show resistance to ALL aminoglycosides
#'   example_isolates %>% filter_aminoglycosides("R", "all")
#'  
#'   # filter on isolates that show resistance to
#'   # any aminoglycoside and any fluoroquinolone
#'   example_isolates %>%
#'     filter_aminoglycosides("R") %>%
#'     filter_fluoroquinolones("R")
#'  
#'   # filter on isolates that show resistance to
#'   # all aminoglycosides and all fluoroquinolones
#'   example_isolates %>%
#'     filter_aminoglycosides("R", "all") %>%
#'     filter_fluoroquinolones("R", "all")
#'   
#'   # with dplyr 1.0.0 and higher (that adds 'across()'), this is equal:
#'   # (though the row names on the first are more correct)
#'   example_isolates %>% filter_carbapenems("R", "all")
#'   example_isolates %>% filter(across(carbapenems(), ~. == "R"))
#' }
#' }
filter_ab_class <- function(x,
                            ab_class,
                            result = NULL,
                            scope = "any",
                            ...) {
  .call_depth <- list(...)$`.call_depth`
  if (is.null(.call_depth)) {
    .call_depth <- 0
  }
  meet_criteria(x, allow_class = "data.frame", .call_depth = .call_depth)
  meet_criteria(ab_class, allow_class = "character", has_length = 1, .call_depth = .call_depth)
  meet_criteria(result, allow_class = "character", has_length = c(1, 2, 3), allow_NULL = TRUE, .call_depth = .call_depth)
  meet_criteria(scope, allow_class = "character", has_length = 1, is_in = c("all", "any"), .call_depth = .call_depth)

  check_dataset_integrity()

  # save to return later
  x_class <- class(x)
  x.bak <- x
  x <- as.data.frame(x, stringsAsFactors = FALSE)
  
  if (is.null(result)) {
    result <- c("S", "I", "R")
  }
  # make result = "SI" works too:
  result <- unlist(strsplit(result, ""))
  
  stop_ifnot(all(result %in% c("S", "I", "R")), "`result` must be one or more of: 'S', 'I', 'R'")
  stop_ifnot(all(scope %in% c("any", "all")), "`scope` must be one of: 'any', 'all'")
  
  # get all columns in data with names that resemble antibiotics
  ab_in_data <- get_column_abx(x, info = FALSE)
  if (length(ab_in_data) == 0) {
    message_("No columns with class <rsi> found (see ?as.rsi), data left unchanged.")
    return(x.bak)
  }
  # get reference data
  ab_class.bak <- ab_class
  ab_class <- gsub("[^a-zA-Z0-9]+", ".*", ab_class)
  ab_class <- gsub("(ph|f)", "(ph|f)", ab_class)
  ab_class <- gsub("(t|th)", "(t|th)", ab_class)
  ab_reference <- subset(antibiotics,
                         group %like% ab_class | 
                           atc_group1 %like% ab_class | 
                           atc_group2 %like% ab_class)
  ab_group <- find_ab_group(ab_class)
  if (ab_group == "") {
    message_("Unknown antimicrobial class '", ab_class.bak, "', data left unchanged.")
    return(x.bak)
  }
  # get the columns with a group names in the chosen ab class
  agents <- ab_in_data[names(ab_in_data) %in% ab_reference$ab]
  if (length(agents) == 0) {
    message_("NOTE: no antimicrobial agents of class ", ab_group, 
             " found (such as ", find_ab_names(ab_class, 2), 
             "), data left unchanged.")
    return(x.bak)
  }
  
  if (length(result) == 1) {
    operator <- " is "
  } else {
    operator <- " is one of "
  }
  if (scope == "any") {
    scope_txt <- " or "
    scope_fn <- any
  } else {
    scope_txt <- " and "
    scope_fn <- all
    if (length(agents) > 1) {
      operator <- gsub("is", "are", operator)
    }
  }
  if (length(agents) > 1) {
    scope <- paste(scope, "of columns ")
  } else {
    scope <- "column "
  }
  
  # sort columns on official name
  agents <- agents[order(ab_name(names(agents), language = NULL))]
  
  message_("Filtering on ", ab_group, ": ", scope,
           paste(paste0("`", font_bold(agents, collapse = NULL),
                        "` (", ab_name(names(agents), tolower = TRUE, language = NULL), ")"),
                 collapse = scope_txt),
           operator, toString(result), as_note = FALSE)
  x_transposed <- as.list(as.data.frame(t(x[, agents, drop = FALSE]), stringsAsFactors = FALSE))
  filtered <- vapply(FUN.VALUE = logical(1), x_transposed, function(y) scope_fn(y %in% result, na.rm = TRUE))
  x <- x[which(filtered), , drop = FALSE]
  class(x) <- x_class
  x
}

#' @rdname filter_ab_class
#' @export
filter_aminoglycosides <- function(x,
                                   result = NULL,
                                   scope = "any",
                                   ...) {
  filter_ab_class(x = x,
                  ab_class = "aminoglycoside",
                  result = result,
                  scope = scope,
                  .call_depth = 1,
                  ...)
}

#' @rdname filter_ab_class
#' @export
filter_carbapenems <- function(x,
                               result = NULL,
                               scope = "any",
                               ...) {
  filter_ab_class(x = x,
                  ab_class = "carbapenem",
                  result = result,
                  scope = scope,
                  .call_depth = 1,
                  ...)
}

#' @rdname filter_ab_class
#' @export
filter_cephalosporins <- function(x,
                                  result = NULL,
                                  scope = "any",
                                  ...) {
  filter_ab_class(x = x,
                  ab_class = "cephalosporin",
                  result = result,
                  scope = scope,
                  .call_depth = 1,
                  ...)
}

#' @rdname filter_ab_class
#' @export
filter_1st_cephalosporins <- function(x,
                                      result = NULL,
                                      scope = "any",
                                      ...) {
  filter_ab_class(x = x,
                  ab_class = "cephalosporins (1st gen.)",
                  result = result,
                  scope = scope,
                  .call_depth = 1,
                  ...)
}

#' @rdname filter_ab_class
#' @export
filter_2nd_cephalosporins <- function(x,
                                      result = NULL,
                                      scope = "any",
                                      ...) {
  filter_ab_class(x = x,
                  ab_class = "cephalosporins (2nd gen.)",
                  result = result,
                  scope = scope,
                  .call_depth = 1,
                  ...)
}

#' @rdname filter_ab_class
#' @export
filter_3rd_cephalosporins <- function(x,
                                      result = NULL,
                                      scope = "any",
                                      ...) {
  filter_ab_class(x = x,
                  ab_class = "cephalosporins (3rd gen.)",
                  result = result,
                  scope = scope,
                  .call_depth = 1,
                  ...)
}

#' @rdname filter_ab_class
#' @export
filter_4th_cephalosporins <- function(x,
                                      result = NULL,
                                      scope = "any",
                                      ...) {
  filter_ab_class(x = x,
                  ab_class = "cephalosporins (4th gen.)",
                  result = result,
                  scope = scope,
                  .call_depth = 1,
                  ...)
}

#' @rdname filter_ab_class
#' @export
filter_5th_cephalosporins <- function(x,
                                      result = NULL,
                                      scope = "any",
                                      ...) {
  filter_ab_class(x = x,
                  ab_class = "cephalosporins (5th gen.)",
                  result = result,
                  scope = scope,
                  .call_depth = 1,
                  ...)
}

#' @rdname filter_ab_class
#' @export
filter_fluoroquinolones <- function(x,
                                    result = NULL,
                                    scope = "any",
                                    ...) {
  filter_ab_class(x = x,
                  ab_class = "fluoroquinolone",
                  result = result,
                  scope = scope,
                  .call_depth = 1,
                  ...)
}

#' @rdname filter_ab_class
#' @export
filter_glycopeptides <- function(x,
                                 result = NULL,
                                 scope = "any",
                                 ...) {
  filter_ab_class(x = x,
                  ab_class = "glycopeptide",
                  result = result,
                  scope = scope,
                  .call_depth = 1,
                  ...)
}

#' @rdname filter_ab_class
#' @export
filter_macrolides <- function(x,
                              result = NULL,
                              scope = "any",
                              ...) {
  filter_ab_class(x = x,
                  ab_class = "macrolide",
                  result = result,
                  scope = scope,
                  .call_depth = 1,
                  ...)
}

#' @rdname filter_ab_class
#' @export
filter_penicillins <- function(x,
                               result = NULL,
                               scope = "any",
                               ...) {
  filter_ab_class(x = x,
                  ab_class = "penicillin",
                  result = result,
                  scope = scope,
                  .call_depth = 1,
                  ...)
}

#' @rdname filter_ab_class
#' @export
filter_tetracyclines <- function(x,
                                 result = NULL,
                                 scope = "any",
                                 ...) {
  filter_ab_class(x = x,
                  ab_class = "tetracycline",
                  result = result,
                  scope = scope,
                  .call_depth = 1,
                  ...)
}

find_ab_group <- function(ab_class) {
  ab_class <- gsub("[^a-zA-Z0-9]", ".*", ab_class)
  ifelse(ab_class %in% c("aminoglycoside",
                         "carbapenem",
                         "cephalosporin",
                         "fluoroquinolone",
                         "glycopeptide",
                         "macrolide",
                         "tetracycline"),
         paste0(ab_class, "s"),
         antibiotics %pm>%
           subset(group %like% ab_class | 
                    atc_group1 %like% ab_class | 
                    atc_group2 %like% ab_class) %pm>%
          pm_pull(group) %pm>%
           unique() %pm>%
           tolower() %pm>%
           sort() %pm>% 
           paste(collapse = "/")
  )
}

find_ab_names <- function(ab_group, n = 3) {
  ab_group <- gsub("[^a-zA-Z0-9]", ".*", ab_group)
  drugs <- antibiotics[which(antibiotics$group %like% ab_group & !antibiotics$ab %like% "[0-9]$"), ]$name
  paste0(sort(ab_name(sample(drugs, size = min(n, length(drugs)), replace = FALSE),
                      tolower = TRUE, language = NULL)), 
         collapse = ", ")
}
```
