## Supplementary material for "AMR - An R Package for Working with Antimicrobial Resistance Data": Source package: first_isolate.R

````
# ==================================================================== #
# TITLE                                                                #
# Antimicrobial Resistance (AMR) Analysis for R                        #
#                                                                      #
# SOURCE                                                               #
# https://github.com/msberends/AMR                                     #
#                                                                      #
# LICENCE                                                              #
# (c) 2018-2021 Berends MS, Luz CF et al.                              #
# Developed at the University of Groningen, the Netherlands, in        #
# collaboration with non-profit organisations Certe Medical            #
# Diagnostics & Advice, and University Medical Center Groningen.       # 
#                                                                      #
# This R package is free software; you can freely use and distribute   #
# it for both personal and commercial purposes under the terms of the  #
# GNU General Public License version 2.0 (GNU GPL-2), as published by  #
# the Free Software Foundation.                                        #
# We created this package for both routine data analysis and academic  #
# research and it was publicly released in the hope that it will be    #
# useful, but it comes WITHOUT ANY WARRANTY OR LIABILITY.              #
#                                                                      #
# Visit our website for the full manual and a complete tutorial about  #
# how to conduct AMR analysis: https://msberends.github.io/AMR/        #
# ==================================================================== #

#' Determine first (weighted) isolates
#'
#' Determine first (weighted) isolates of all microorganisms of every patient per episode and (if needed) per specimen type. To determine patient episodes not necessarily based on microorganisms, use [is_new_episode()] that also supports grouping with the `dplyr` package.
#' @inheritSection lifecycle Stable lifecycle
#' @param x a [data.frame] containing isolates. Can be left blank when used inside `dplyr` verbs, such as [`filter()`][dplyr::filter()], [`mutate()`][dplyr::mutate()] and [`summarise()`][dplyr::summarise()].
#' @param col_date column name of the result date (or date that is was received on the lab), defaults to the first column with a date class
#' @param col_patient_id column name of the unique IDs of the patients, defaults to the first column that starts with 'patient' or 'patid' (case insensitive)
#' @param col_mo column name of the IDs of the microorganisms (see [as.mo()]), defaults to the first column of class [`mo`]. Values will be coerced using [as.mo()].
#' @param col_testcode column name of the test codes. Use `col_testcode = NULL` to **not** exclude certain test codes (such as test codes for screening). In that case `testcodes_exclude` will be ignored.
#' @param col_specimen column name of the specimen type or group
#' @param col_icu column name of the logicals (`TRUE`/`FALSE`) whether a ward or department is an Intensive Care Unit (ICU)
#' @param col_keyantibiotics column name of the key antibiotics to determine first *weighted* isolates, see [key_antibiotics()]. Defaults to the first column that starts with 'key' followed by 'ab' or 'antibiotics' (case insensitive). Use `col_keyantibiotics = FALSE` to prevent this.
#' @param episode_days episode in days after which a genus/species combination will be determined as 'first isolate' again. The default of 365 days is based on the guideline by CLSI, see Source. 
#' @param testcodes_exclude character vector with test codes that should be excluded (case-insensitive)
#' @param icu_exclude logical whether ICU isolates should be excluded (rows with value `TRUE` in the column set with `col_icu`)
#' @param specimen_group value in the column set with `col_specimen` to filter on
#' @param type type to determine weighed isolates; can be `"keyantibiotics"` or `"points"`, see Details
#' @param ignore_I logical to determine whether antibiotic interpretations with `"I"` will be ignored when `type = "keyantibiotics"`, see Details
#' @param points_threshold points until the comparison of key antibiotics will lead to inclusion of an isolate when `type = "points"`, see Details
#' @param info print progress
#' @param include_unknown logical to determine whether 'unknown' microorganisms should be included too, i.e. microbial code `"UNKNOWN"`, which defaults to `FALSE`. For WHONET users, this means that all records with organism code `"con"` (*contamination*) will be excluded at default. Isolates with a microbial ID of `NA` will always be excluded as first isolate.
#' @param ... arguments passed on to [first_isolate()] when using [filter_first_isolate()], or arguments passed on to [key_antibiotics()] when using [filter_first_weighted_isolate()]
#' @details 
#' These functions are context-aware when used inside `dplyr` verbs, such as `filter()`, `mutate()` and `summarise()`. This means that then the `x` argument can be left blank, please see *Examples*.
#' 
#' The [first_isolate()] function is a wrapper around the [is_new_episode()] function, but more efficient for data sets containing microorganism codes or names.
#' 
#' All isolates with a microbial ID of `NA` will be excluded as first isolate.
#' 
#' ### Why this is so important
#' To conduct an analysis of antimicrobial resistance, you should only include the first isolate of every patient per episode [(Hindler *et al.* 2007)](https://pubmed.ncbi.nlm.nih.gov/17304462/). If you would not do this, you could easily get an overestimate or underestimate of the resistance of an antibiotic. Imagine that a patient was admitted with an MRSA and that it was found in 5 different blood cultures the following week. The resistance percentage of oxacillin of all *S. aureus* isolates would be overestimated, because you included this MRSA more than once. It would be [selection bias](https://en.wikipedia.org/wiki/Selection_bias).
#'
#' ### `filter_*()` shortcuts
#'
#' The functions [filter_first_isolate()] and [filter_first_weighted_isolate()] are helper functions to quickly filter on first isolates.
#' 
#' The function [filter_first_isolate()] is essentially equal to either:
#' 
#' ```
#'   x[first_isolate(x, ...), ]
#'   
#'   x %>% filter(first_isolate(...))
#' ```
#' 
#' The function [filter_first_weighted_isolate()] is essentially equal to:
#' 
#' ```
#'   x %>%
#'     mutate(keyab = key_antibiotics(.)) %>%
#'     mutate(only_weighted_firsts = first_isolate(x,
#'                                                 col_keyantibiotics = "keyab", ...)) %>%
#'     filter(only_weighted_firsts == TRUE) %>%
#'     select(-only_weighted_firsts, -keyab)
#' ```
#' @section Key antibiotics:
#' There are two ways to determine whether isolates can be included as first *weighted* isolates which will give generally the same results:
#'
#' 1. Using `type = "keyantibiotics"` and argument `ignore_I`
#' 
#'    Any difference from S to R (or vice versa) will (re)select an isolate as a first weighted isolate. With `ignore_I = FALSE`, also differences from I to S|R (or vice versa) will lead to this. This is a reliable method and 30-35 times faster than method 2. Read more about this in the [key_antibiotics()] function.
#'    
#' 2. Using `type = "points"` and argument `points_threshold`
#' 
#'    A difference from I to S|R (or vice versa) means 0.5 points, a difference from S to R (or vice versa) means 1 point. When the sum of points exceeds `points_threshold`, which default to `2`, an isolate will be (re)selected as a first weighted isolate.
#' @rdname first_isolate
#' @seealso [key_antibiotics()]
#' @export
#' @return A [`logical`] vector
#' @source Methodology of this function is strictly based on:
#' 
#' **M39 Analysis and Presentation of Cumulative Antimicrobial Susceptibility Test Data, 4th Edition**, 2014, *Clinical and Laboratory Standards Institute (CLSI)*. <https://clsi.org/standards/products/microbiology/documents/m39/>.
#' @inheritSection AMR Read more on our website!
#' @examples
#' # `example_isolates` is a dataset available in the AMR package.
#' # See ?example_isolates.
#' 
#' # basic filtering on first isolates
#' example_isolates[first_isolate(example_isolates), ]
#' 
#' # filtering based on isolates ----------------------------------------------
#' \donttest{
#' if (require("dplyr")) {
#'   # filter on first isolates:
#'   example_isolates %>%
#'     mutate(first_isolate = first_isolate(.)) %>%
#'     filter(first_isolate == TRUE)
#'  
#'   # short-hand versions:
#'   example_isolates %>%
#'     filter(first_isolate())
#'   example_isolates %>%
#'     filter_first_isolate()
#'     
#'   example_isolates %>%
#'     filter_first_weighted_isolate()
#'   
#'   # now let's see if first isolates matter:
#'   A <- example_isolates %>%
#'     group_by(hospital_id) %>%
#'     summarise(count = n_rsi(GEN),            # gentamicin availability
#'               resistance = resistance(GEN))  # gentamicin resistance
#'  
#'   B <- example_isolates %>%
#'     filter_first_weighted_isolate() %>%      # the 1st isolate filter
#'     group_by(hospital_id) %>%
#'     summarise(count = n_rsi(GEN),            # gentamicin availability
#'               resistance = resistance(GEN))  # gentamicin resistance
#'  
#'   # Have a look at A and B.
#'   # B is more reliable because every isolate is counted only once.
#'   # Gentamicin resistance in hospital D appears to be 3.7% higher than
#'   # when you (erroneously) would have used all isolates for analysis.
#' }
#' }
first_isolate <- function(x,
                          col_date = NULL,
                          col_patient_id = NULL,
                          col_mo = NULL,
                          col_testcode = NULL,
                          col_specimen = NULL,
                          col_icu = NULL,
                          col_keyantibiotics = NULL,
                          episode_days = 365,
                          testcodes_exclude = NULL,
                          icu_exclude = FALSE,
                          specimen_group = NULL,
                          type = "keyantibiotics",
                          ignore_I = TRUE,
                          points_threshold = 2,
                          info = interactive(),
                          include_unknown = FALSE,
                          ...) {
  if (missing(x)) {
    x <- get_current_data(arg_name = "x", call = -2)
  }
  meet_criteria(x, allow_class = "data.frame") # also checks dimensions to be >0
  meet_criteria(col_date, allow_class = "character", has_length = 1, allow_NULL = TRUE, is_in = colnames(x))
  meet_criteria(col_patient_id, allow_class = "character", has_length = 1, allow_NULL = TRUE, is_in = colnames(x))
  meet_criteria(col_mo, allow_class = "character", has_length = 1, allow_NULL = TRUE, is_in = colnames(x))
  meet_criteria(col_testcode, allow_class = "character", has_length = 1, allow_NULL = TRUE, is_in = colnames(x))
  if (isFALSE(col_specimen)) {
    col_specimen <- NULL
  }
  meet_criteria(col_specimen, allow_class = "character", has_length = 1, allow_NULL = TRUE, is_in = colnames(x))
  meet_criteria(col_icu, allow_class = "character", has_length = 1, allow_NULL = TRUE, is_in = colnames(x))
  if (isFALSE(col_keyantibiotics)) {
    col_keyantibiotics <- NULL
  }
  meet_criteria(col_keyantibiotics, allow_class = "character", has_length = 1, allow_NULL = TRUE, is_in = colnames(x))
  meet_criteria(episode_days, allow_class = c("numeric", "integer"), has_length = 1)
  meet_criteria(testcodes_exclude, allow_class = "character", allow_NULL = TRUE)
  meet_criteria(icu_exclude, allow_class = "logical", has_length = 1)
  meet_criteria(specimen_group, allow_class = "character", has_length = 1, allow_NULL = TRUE)
  meet_criteria(type, allow_class = "character", has_length = 1)
  meet_criteria(ignore_I, allow_class = "logical", has_length = 1)
  meet_criteria(points_threshold, allow_class = c("numeric", "integer"), has_length = 1)
  meet_criteria(info, allow_class = "logical", has_length = 1)
  meet_criteria(include_unknown, allow_class = "logical", has_length = 1)
  
  dots <- unlist(list(...))
  if (length(dots) != 0) {
    # backwards compatibility with old arguments
    dots.names <- dots %pm>% names()
    if ("filter_specimen" %in% dots.names) {
      specimen_group <- dots[which(dots.names == "filter_specimen")]
    }
    if ("tbl" %in% dots.names) {
      x <- dots[which(dots.names == "tbl")]
    }
  }
  
  # remove data.table, grouping from tibbles, etc.
  x <- as.data.frame(x, stringsAsFactors = FALSE)
  
 # try to find columns based on type
  # -- mo
  if (is.null(col_mo)) {
    col_mo <- search_type_in_df(x = x, type = "mo")
    stop_if(is.null(col_mo), "`col_mo` must be set")
    stop_ifnot(col_mo %in% colnames(x), "column '", col_mo, "' (`col_mo`) not found")
  }
  
  # -- date
  if (is.null(col_date)) {
    col_date <- search_type_in_df(x = x, type = "date")
    stop_if(is.null(col_date), "`col_date` must be set")
  }
  
  # -- patient id
  if (is.null(col_patient_id)) {
    if (all(c("First name", "Last name", "Sex") %in% colnames(x))) {
      # WHONET support
      x$patient_id <- paste(x$`First name`, x$`Last name`, x$Sex)
      col_patient_id <- "patient_id"
      message_("Using combined columns '", font_bold("First name"), "', '", font_bold("Last name"), "' and '", font_bold("Sex"), "' as input for `col_patient_id`")
    } else {
      col_patient_id <- search_type_in_df(x = x, type = "patient_id")
    }
    stop_if(is.null(col_patient_id), "`col_patient_id` must be set")
  }
  
  # -- key antibiotics
  if (is.null(col_keyantibiotics)) {
    col_keyantibiotics <- search_type_in_df(x = x, type = "keyantibiotics")
  }

  # -- specimen
  if (is.null(col_specimen) & !is.null(specimen_group)) {
    col_specimen <- search_type_in_df(x = x, type = "specimen")
  }
  
  # check if columns exist
  check_columns_existance <- function(column, tblname = x) {
    if (!is.null(column)) {
      stop_ifnot(column %in% colnames(tblname),
                 "Column '", column, "' not found.", call = FALSE)
    }
  }
  
  check_columns_existance(col_date)
  check_columns_existance(col_patient_id)
  check_columns_existance(col_mo)
  check_columns_existance(col_testcode)
  check_columns_existance(col_icu)
  check_columns_existance(col_keyantibiotics)
  
  # convert dates to Date
  dates <- as.Date(x[, col_date, drop = TRUE])
  dates[is.na(dates)] <- as.Date("1970-01-01")
  x[, col_date] <- dates
  
  # create original row index
  x$newvar_row_index <- seq_len(nrow(x))
  x$newvar_mo <- x[, col_mo, drop = TRUE]
  x$newvar_genus_species <- paste(mo_genus(x$newvar_mo), mo_species(x$newvar_mo))
  x$newvar_date <- x[, col_date, drop = TRUE]
  x$newvar_patient_id <- x[, col_patient_id, drop = TRUE]
  
  if (is.null(col_testcode)) {
    testcodes_exclude <- NULL
  }
  # remove testcodes
  if (!is.null(testcodes_exclude) & info == TRUE) {
    message_("[Criterion] Exclude test codes: ", toString(paste0("'", testcodes_exclude, "'")),
             add_fn = font_black,
             as_note = FALSE)
  }
  
  if (is.null(col_specimen)) {
    specimen_group <- NULL
  }
  
  # filter on specimen group and keyantibiotics when they are filled in
  if (!is.null(specimen_group)) {
    check_columns_existance(col_specimen, x)
    if (info == TRUE) {
      message_("[Criterion] Exclude other than specimen group '", specimen_group, "'",
               add_fn = font_black,
               as_note = FALSE)
    }
  }
  if (!is.null(col_keyantibiotics)) {
    x$newvar_key_ab <- x[, col_keyantibiotics, drop = TRUE]
  }
  
  if (is.null(testcodes_exclude)) {
    testcodes_exclude <- ""
  }
  
  # arrange data to the right sorting
  if (is.null(specimen_group)) {
    x <- x[order(x$newvar_patient_id, 
                 x$newvar_genus_species,
                 x$newvar_date), ]
    rownames(x) <- NULL
    row.start <- 1
    row.end <- nrow(x)
  } else {
    # filtering on specimen and only analyse these rows to save time
    x <- x[order(pm_pull(x, col_specimen),
                 x$newvar_patient_id, 
                 x$newvar_genus_species,
                 x$newvar_date), ]
    rownames(x) <- NULL
    suppressWarnings(
      row.start <- which(x %pm>% pm_pull(col_specimen) == specimen_group) %pm>% min(na.rm = TRUE)
    )
    suppressWarnings(
      row.end <- which(x %pm>% pm_pull(col_specimen) == specimen_group) %pm>% max(na.rm = TRUE)
    )
  }
  
  # speed up - return immediately if obvious
  if (abs(row.start) == Inf | abs(row.end) == Inf) {
    if (info == TRUE) {
      message_("=> Found ", font_bold("no isolates"),
               add_fn = font_black, 
               as_note = FALSE)
    }
    return(rep(FALSE, nrow(x)))
  }
  if (row.start == row.end) {
    if (info == TRUE) {
      message_("=> Found ", font_bold("1 isolate"), ", as the data only contained 1 row", 
               add_fn = font_black,
               as_note = FALSE)
    }
    return(TRUE)
  }
  if (length(c(row.start:row.end)) == pm_n_distinct(x[c(row.start:row.end), col_mo, drop = TRUE])) {
    if (info == TRUE) {
      message_("=> Found ", font_bold(paste(length(c(row.start:row.end)), "isolates")),
               ", as all isolates were different microorganisms",
               add_fn = font_black,
               as_note = FALSE)
    }
    return(rep(TRUE, length(c(row.start:row.end))))
  }
  
  # did find some isolates - add new index numbers of rows
  x$newvar_row_index_sorted <- seq_len(nrow(x))
  
  scope.size <- nrow(x[which(x$newvar_row_index_sorted %in% c(row.start + 1:row.end) &
                               !is.na(x$newvar_mo)), , drop = FALSE])
  
  # Analysis of first isolate ----
  x$other_pat_or_mo <- ifelse(x$newvar_patient_id == pm_lag(x$newvar_patient_id) &
                                x$newvar_genus_species == pm_lag(x$newvar_genus_species),
                              FALSE,
                              TRUE)
  x$episode_group <- paste(x$newvar_patient_id, x$newvar_genus_species)
  x$more_than_episode_ago <- unlist(lapply(unique(x$episode_group),
                                           function(g,
                                                    df = x,
                                                    days = episode_days) {
                                             is_new_episode(x = df[which(df$episode_group == g), ]$newvar_date,
                                                            episode_days = days)
                                           }))
  
  weighted.notice <- ""
  if (!is.null(col_keyantibiotics)) {
    weighted.notice <- "weighted "
    if (info == TRUE) {
      if (type == "keyantibiotics") {
        message_("[Criterion] Base inclusion on key antibiotics, ",
                 ifelse(ignore_I == FALSE, "not ", ""),
                 "ignoring I",
                 add_fn = font_black,
                 as_note = FALSE)
      }
      if (type == "points") {
        message_("[Criterion] Base inclusion on key antibiotics, using points threshold of "
                 , points_threshold,
                 add_fn = font_black,
                 as_note = FALSE)
      }
    }
    type_param <- type
    
    x$other_key_ab <- !key_antibiotics_equal(y = x$newvar_key_ab,
                                             z = pm_lag(x$newvar_key_ab),
                                             type = type_param,
                                             ignore_I = ignore_I,
                                             points_threshold = points_threshold,
                                             info = info)
    # with key antibiotics
    x$newvar_first_isolate <- pm_if_else(x$newvar_row_index_sorted >= row.start &
                                           x$newvar_row_index_sorted <= row.end &
                                           x$newvar_genus_species != "" & 
                                           (x$other_pat_or_mo | x$more_than_episode_ago | x$other_key_ab),
                                         TRUE,
                                         FALSE)
    
  } else {
    # no key antibiotics
    x$newvar_first_isolate <- pm_if_else(x$newvar_row_index_sorted >= row.start &
                                           x$newvar_row_index_sorted <= row.end &
                                           x$newvar_genus_species != "" & 
                                           (x$other_pat_or_mo | x$more_than_episode_ago),
                                         TRUE,
                                         FALSE)
  }
  
  # first one as TRUE
  x[row.start, "newvar_first_isolate"] <- TRUE
  # no tests that should be included, or ICU
  if (!is.null(col_testcode)) {
    x[which(x[, col_testcode] %in% tolower(testcodes_exclude)), "newvar_first_isolate"] <- FALSE
  }
  if (!is.null(col_icu)) {
    if (icu_exclude == TRUE) {
      message_("[Criterion] Exclude isolates from ICU.",
               add_fn = font_black,
               as_note = FALSE)
      x[which(as.logical(x[, col_icu, drop = TRUE])), "newvar_first_isolate"] <- FALSE
    } else {
      message_("[Criterion] Include isolates from ICU.",
               add_fn = font_black,
               as_note = FALSE)
    }
  }
  
  decimal.mark <- getOption("OutDec")
  big.mark <- ifelse(decimal.mark != ",", ",", ".")
  
  # handle empty microorganisms
  if (any(x$newvar_mo == "UNKNOWN", na.rm = TRUE) & info == TRUE) {
    message_(ifelse(include_unknown == TRUE, "Included ", "Excluded "), 
             format(sum(x$newvar_mo == "UNKNOWN", na.rm = TRUE),
                    decimal.mark = decimal.mark, big.mark = big.mark), 
             " isolates with a microbial ID 'UNKNOWN' (column '", font_bold(col_mo), "')")
  }
  x[which(x$newvar_mo == "UNKNOWN"), "newvar_first_isolate"] <- include_unknown
  
  # exclude all NAs
  if (any(is.na(x$newvar_mo)) & info == TRUE) {
    message_("Excluded ", format(sum(is.na(x$newvar_mo), na.rm = TRUE),
                                 decimal.mark = decimal.mark, big.mark = big.mark), 
             " isolates with a microbial ID 'NA' (column '", font_bold(col_mo), "')")
  }
  x[which(is.na(x$newvar_mo)), "newvar_first_isolate"] <- FALSE
  
  # arrange back according to original sorting again
  x <- x[order(x$newvar_row_index), ]
  rownames(x) <- NULL
  
  if (info == TRUE) {
    n_found <- sum(x$newvar_first_isolate, na.rm = TRUE)
    p_found_total <- percentage(n_found / nrow(x[which(!is.na(x$newvar_mo)), , drop = FALSE]), digits = 1)
    p_found_scope <- percentage(n_found / scope.size, digits = 1)
    if (!p_found_total %like% "[.]") {
      p_found_total <- gsub("%", ".0%", p_found_total, fixed = TRUE)
    }
    if (!p_found_scope %like% "[.]") {
      p_found_scope <- gsub("%", ".0%", p_found_scope, fixed = TRUE)
    }
    # mark up number of found
    n_found <- format(n_found, big.mark = big.mark, decimal.mark = decimal.mark)
    if (p_found_total != p_found_scope) {
      msg_txt <- paste0("=> Found ",
                        font_bold(paste0(n_found, " first ", weighted.notice, "isolates")),
                        " (", p_found_scope, " within scope and ", p_found_total, " of total where a microbial ID was available)")
    } else {
      msg_txt <- paste0("=> Found ",
                        font_bold(paste0(n_found, " first ", weighted.notice, "isolates")),
                        " (", p_found_total, " of total where a microbial ID was available)")
    }
    message_(msg_txt, add_fn = font_black, as_note = FALSE)
  }
  
  x$newvar_first_isolate
  
}

#' @rdname first_isolate
#' @export
filter_first_isolate <- function(x,
                                 col_date = NULL,
                                 col_patient_id = NULL,
                                 col_mo = NULL,
                                 ...) {
  meet_criteria(x, allow_class = "data.frame")
  meet_criteria(col_date, allow_class = "character", has_length = 1, allow_NULL = TRUE, is_in = colnames(x))
  meet_criteria(col_patient_id, allow_class = "character", has_length = 1, allow_NULL = TRUE, is_in = colnames(x))
  meet_criteria(col_mo, allow_class = "character", has_length = 1, allow_NULL = TRUE, is_in = colnames(x))
  subset(x, first_isolate(x = x,
                          col_date = col_date,
                          col_patient_id = col_patient_id,
                          col_mo = col_mo,
                          ...))
}

#' @rdname first_isolate
#' @export
filter_first_weighted_isolate <- function(x,
                                          col_date = NULL,
                                          col_patient_id = NULL,
                                          col_mo = NULL,
                                          col_keyantibiotics = NULL,
                                          ...) {
  meet_criteria(x, allow_class = "data.frame")
  meet_criteria(col_date, allow_class = "character", has_length = 1, allow_NULL = TRUE, is_in = colnames(x))
  meet_criteria(col_patient_id, allow_class = "character", has_length = 1, allow_NULL = TRUE, is_in = colnames(x))
  meet_criteria(col_mo, allow_class = "character", has_length = 1, allow_NULL = TRUE, is_in = colnames(x))
  meet_criteria(col_keyantibiotics, allow_class = "character", has_length = 1, allow_NULL = TRUE, is_in = colnames(x))
  y <- x
  if (is.null(col_keyantibiotics)) {
    # first try to look for it
    col_keyantibiotics <- search_type_in_df(x = x, type = "keyantibiotics")
    # still NULL? Then create it since we are calling filter_first_WEIGHTED_isolate()
    if (is.null(col_keyantibiotics)) {
      y$keyab <- suppressMessages(key_antibiotics(x,
                                                  col_mo = col_mo,
                                                  ...))
      col_keyantibiotics <- "keyab"
    }
  }
  
  subset(x, first_isolate(x = y,
                          col_date = col_date,
                          col_patient_id = col_patient_id))
}
````
