## Supplementary material for "AMR - An R Package for Working with Antimicrobial Resistance Data": Source package: g.test.R

````
# ==================================================================== #
# TITLE                                                                #
# Antimicrobial Resistance (AMR) Analysis for R                        #
#                                                                      #
# SOURCE                                                               #
# https://github.com/msberends/AMR                                     #
#                                                                      #
# LICENCE                                                              #
# (c) 2018-2021 Berends MS, Luz CF et al.                              #
# Developed at the University of Groningen, the Netherlands, in        #
# collaboration with non-profit organisations Certe Medical            #
# Diagnostics & Advice, and University Medical Center Groningen.       # 
#                                                                      #
# This R package is free software; you can freely use and distribute   #
# it for both personal and commercial purposes under the terms of the  #
# GNU General Public License version 2.0 (GNU GPL-2), as published by  #
# the Free Software Foundation.                                        #
# We created this package for both routine data analysis and academic  #
# research and it was publicly released in the hope that it will be    #
# useful, but it comes WITHOUT ANY WARRANTY OR LIABILITY.              #
#                                                                      #
# Visit our website for the full manual and a complete tutorial about  #
# how to conduct AMR analysis: https://msberends.github.io/AMR/        #
# ==================================================================== #

#' *G*-test for Count Data
#'
#' [g.test()] performs chi-squared contingency table tests and goodness-of-fit tests, just like [chisq.test()] but is more reliable (1). A *G*-test can be used to see whether the number of observations in each category fits a theoretical expectation (called a ***G*-test of goodness-of-fit**), or to see whether the proportions of one variable are different for different values of the other variable (called a ***G*-test of independence**).
#' @inheritSection lifecycle Questioning lifecycle
#' @inherit stats::chisq.test params return
#' @details If `x` is a matrix with one row or column, or if `x` is a vector and `y` is not given, then a *goodness-of-fit test* is performed (`x` is treated as a one-dimensional contingency table). The entries of `x` must be non-negative integers. In this case, the hypothesis tested is whether the population probabilities equal those in `p`, or are all equal if `p` is not given.
#'
#' If `x` is a matrix with at least two rows and columns, it is taken as a two-dimensional contingency table: the entries of `x` must be non-negative integers.  Otherwise, `x` and `y` must be vectors or factors of the same length; cases with missing values are removed, the objects are coerced to factors, and the contingency table is computed from these.  Then Pearson's chi-squared test is performed of the null hypothesis that the joint distribution of the cell counts in a 2-dimensional contingency table is the product of the row and column marginals.
#'
#' The p-value is computed from the asymptotic chi-squared distribution of the test statistic.
#'
#' In the contingency table case simulation is done by random sampling from the set of all contingency tables with given marginals, and works only if the marginals are strictly positive. Note that this is not the usual sampling situation assumed for a chi-squared test (such as the *G*-test) but rather that for Fisher's exact test.
#'
#' In the goodness-of-fit case simulation is done by random sampling from the discrete distribution specified by `p`, each sample being of size `n = sum(x)`. This simulation is done in \R and may be slow.
#'   
#' ## *G*-test of goodness-of-fit (likelihood ratio test)
#' Use the *G*-test of goodness-of-fit when you have one nominal variable with two or more values (such as male and female, or red, pink and white flowers). You compare the observed counts of numbers of observations in each category with the expected counts, which you calculate using some kind of theoretical expectation (such as a 1:1 sex ratio or a 1:2:1 ratio in a genetic cross).
#'
#' If the expected number of observations in any category is too small, the *G*-test may give inaccurate results, and you should use an exact test instead ([fisher.test()]).
#'
#' The *G*-test of goodness-of-fit is an alternative to the chi-square test of goodness-of-fit ([chisq.test()]); each of these tests has some advantages and some disadvantages, and the results of the two tests are usually very similar.
#'
#' ## *G*-test of independence
#' Use the *G*-test of independence when you have two nominal variables, each with two or more possible values. You want to know whether the proportions for one variable are different among values of the other variable.
#'
#' It is also possible to do a *G*-test of independence with more than two nominal variables. For example, Jackson et al. (2013) also had data for children under 3, so you could do an analysis of old vs. young, thigh vs. arm, and reaction vs. no reaction, all analyzed together.
#'
#' Fisher's exact test ([fisher.test()]) is an **exact** test, where the *G*-test is still only an **approximation**. For any 2x2 table, Fisher's Exact test may be slower but will still run in seconds, even if the sum of your observations is multiple millions.
#'
#' The *G*-test of independence is an alternative to the chi-square test of independence ([chisq.test()]), and they will give approximately the same results.
#'
#' ## How the test works
#' Unlike the exact test of goodness-of-fit ([fisher.test()]), the *G*-test does not directly calculate the probability of obtaining the observed results or something more extreme. Instead, like almost all statistical tests, the *G*-test has an intermediate step; it uses the data to calculate a test statistic that measures how far the observed data are from the null expectation. You then use a mathematical relationship, in this case the chi-square distribution, to estimate the probability of obtaining that value of the test statistic.
#'
#' The *G*-test uses the log of the ratio of two likelihoods as the test statistic, which is why it is also called a likelihood ratio test or log-likelihood ratio test. The formula to calculate a *G*-statistic is:
#' 
#' \eqn{G = 2 * sum(x * log(x / E))}
#' 
#' where `E` are the expected values. Since this is chi-square distributed, the p value can be calculated in \R with:
#' ```
#' p <- stats::pchisq(G, df, lower.tail = FALSE)
#' ```
#' where `df` are the degrees of freedom.
#'
#' If there are more than two categories and you want to find out which ones are significantly different from their null expectation, you can use the same method of testing each category vs. the sum of all categories, with the Bonferroni correction. You use *G*-tests for each category, of course.
#' @seealso [chisq.test()]
#' @references 1. McDonald, J.H. 2014. **Handbook of Biological Statistics (3rd ed.)**. Sparky House Publishing, Baltimore, Maryland. <http://www.biostathandbook.com/gtestgof.html>.
#' @source The code for this function is identical to that of [chisq.test()], except that:
#' - The calculation of the statistic was changed to \eqn{2 * sum(x * log(x / E))}
#' - Yates' continuity correction was removed as it does not apply to a *G*-test
#' - The possibility to simulate p values with `simulate.p.value` was removed
#' @export
#' @importFrom stats pchisq complete.cases
#' @inheritSection AMR Read more on our website!
#' @examples
#' # = EXAMPLE 1 =
#' # Shivrain et al. (2006) crossed clearfield rice (which are resistant
#' # to the herbicide imazethapyr) with red rice (which are susceptible to
#' # imazethapyr). They then crossed the hybrid offspring and examined the
#' # F2 generation, where they found 772 resistant plants, 1611 moderately
#' # resistant plants, and 737 susceptible plants. If resistance is controlled
#' # by a single gene with two co-dominant alleles, you would expect a 1:2:1
#' # ratio.
#'
#' x <- c(772, 1611, 737)
#' G <- g.test(x, p = c(1, 2, 1) / 4)
#' # G$p.value = 0.12574.
#'
#' # There is no significant difference from a 1:2:1 ratio.
#' # Meaning: resistance controlled by a single gene with two co-dominant
#' # alleles, is plausible.
#'
#'
#' # = EXAMPLE 2 =
#' # Red crossbills (Loxia curvirostra) have the tip of the upper bill either
#' # right or left of the lower bill, which helps them extract seeds from pine
#' # cones. Some have hypothesized that frequency-dependent selection would
#' # keep the number of right and left-billed birds at a 1:1 ratio. Groth (1992)
#' # observed 1752 right-billed and 1895 left-billed crossbills.
#'
#' x <- c(1752, 1895)
#' g.test(x)
#' # p = 0.01787343
#'
#' # There is a significant difference from a 1:1 ratio.
#' # Meaning: there are significantly more left-billed birds.
#'
g.test <- function(x,
                   y = NULL,
                   # correct = TRUE,
                   p = rep(1 / length(x), length(x)),
                   rescale.p = FALSE) {
  DNAME <- deparse(substitute(x))
  if (is.data.frame(x))
    x <- as.matrix(x)
  if (is.matrix(x)) {
    if (min(dim(x)) == 1L)
      x <- as.vector(x)
  }
  if (!is.matrix(x) && !is.null(y)) {
    if (length(x) != length(y))
      stop("'x' and 'y' must have the same length")
    DNAME2 <- deparse(substitute(y))
    xname <- if (length(DNAME) > 1L || nchar(DNAME, "w") >
                 30)
      ""
    else DNAME
    yname <- if (length(DNAME2) > 1L || nchar(DNAME2, "w") >
                 30)
      ""
    else DNAME2
    OK <- complete.cases(x, y)
    x <- factor(x[OK])
    y <- factor(y[OK])
    if ((nlevels(x) < 2L) || (nlevels(y) < 2L))
      stop("'x' and 'y' must have at least 2 levels")
    x <- table(x, y)
    names(dimnames(x)) <- c(xname, yname)
    DNAME <- paste(paste(DNAME, collapse = "\n"), "and",
                   paste(DNAME2, collapse = "\n"))
  }
  if (any(x < 0) || any(is.na((x)))) # this last one was anyNA, but only introduced in R 3.1.0
    stop("all entries of 'x' must be nonnegative and finite")
  if ((n <- sum(x)) == 0)
    stop("at least one entry of 'x' must be positive")
  
  
  if (is.matrix(x)) {
    METHOD <- "G-test of independence"
    nr <- as.integer(nrow(x))
    nc <- as.integer(ncol(x))
    if (is.na(nr) || is.na(nc) || is.na(nr * nc))
      stop("invalid nrow(x) or ncol(x)", domain = NA)
    # add fisher.test suggestion
    if (nr == 2 && nc == 2)
      warning("`fisher.test()` is always more reliable for 2x2 tables and although much slower, often only takes seconds.")
    sr <- rowSums(x)
    sc <- colSums(x)
    E <- outer(sr, sc, "*") / n
    v <- function(r, c, n) c * r * (n - r) * (n - c) / n ^ 3
    V <- outer(sr, sc, v, n)
    dimnames(E) <- dimnames(x)
    
    STATISTIC <- 2 * sum(x * log(x / E)) # sum((abs(x - E) - YATES)^2/E) for chisq.test
    PARAMETER <- (nr - 1L) * (nc - 1L)
    PVAL <- pchisq(STATISTIC, PARAMETER, lower.tail = FALSE)
    
  }
  else {
    if (length(dim(x)) > 2L)
      stop("invalid 'x'")
    if (length(x) == 1L)
      stop("'x' must at least have 2 elements")
    if (length(x) != length(p))
      stop("'x' and 'p' must have the same number of elements")
    if (any(p < 0))
      stop("probabilities must be non-negative.")
    if (abs(sum(p) - 1) > sqrt(.Machine$double.eps)) {
      if (rescale.p)
        p <- p / sum(p)
      else stop("probabilities must sum to 1.")
    }
    METHOD <- "G-test of goodness-of-fit (likelihood ratio test)"
    E <- n * p
    V <- n * p * (1 - p)
    STATISTIC <- 2 * sum(x * log(x / E)) # sum((x - E)^2/E) for chisq.test
    names(E) <- names(x)
    
    PARAMETER <- length(x) - 1
    PVAL <- pchisq(STATISTIC, PARAMETER, lower.tail = FALSE)
    
  }
  names(STATISTIC) <- "X-squared"
  names(PARAMETER) <- "df"
  if (any(E < 5) && is.finite(PARAMETER))
    warning("G-statistic approximation may be incorrect due to E < 5")
  
  structure(list(statistic = STATISTIC, argument = PARAMETER,
                 p.value = PVAL, method = METHOD, data.name = DNAME,
                 observed = x, expected = E, residuals = (x - E) / sqrt(E),
                 stdres = (x - E) / sqrt(V)), class = "htest")
}
````
