## Supplementary material for "AMR - An R Package for Working with Antimicrobial Resistance Data": Source package: ggplot_pca.R

```
# ==================================================================== #
# TITLE                                                                #
# Antimicrobial Resistance (AMR) Analysis for R                        #
#                                                                      #
# SOURCE                                                               #
# https://github.com/msberends/AMR                                     #
#                                                                      #
# LICENCE                                                              #
# (c) 2018-2021 Berends MS, Luz CF et al.                              #
# Developed at the University of Groningen, the Netherlands, in        #
# collaboration with non-profit organisations Certe Medical            #
# Diagnostics & Advice, and University Medical Center Groningen.       # 
#                                                                      #
# This R package is free software; you can freely use and distribute   #
# it for both personal and commercial purposes under the terms of the  #
# GNU General Public License version 2.0 (GNU GPL-2), as published by  #
# the Free Software Foundation.                                        #
# We created this package for both routine data analysis and academic  #
# research and it was publicly released in the hope that it will be    #
# useful, but it comes WITHOUT ANY WARRANTY OR LIABILITY.              #
#                                                                      #
# Visit our website for the full manual and a complete tutorial about  #
# how to conduct AMR analysis: https://msberends.github.io/AMR/        #
# ==================================================================== #

#' PCA biplot with `ggplot2`
#'
#' Produces a `ggplot2` variant of a so-called [biplot](https://en.wikipedia.org/wiki/Biplot) for PCA (principal component analysis), but is more flexible and more appealing than the base \R [biplot()] function.
#' @inheritSection lifecycle Maturing lifecycle
#' @param x an object returned by [pca()], [prcomp()] or [princomp()]
#' @inheritParams stats::biplot.prcomp
#' @param labels an optional vector of labels for the observations. If set, the labels will be placed below their respective points. When using the [pca()] function as input for `x`, this will be determined automatically based on the attribute `non_numeric_cols`, see [pca()].
#' @param labels_textsize the size of the text used for the labels
#' @param labels_text_placement adjustment factor the placement of the variable names (`>=1` means further away from the arrow head)
#' @param groups an optional vector of groups for the labels, with the same length as `labels`. If set, the points and labels will be coloured according to these groups. When using the [pca()] function as input for `x`, this will be determined automatically based on the attribute `non_numeric_cols`, see [pca()].
#' @param ellipse a logical to indicate whether a normal data ellipse should be drawn for each group (set with `groups`)
#' @param ellipse_prob statistical size of the ellipse in normal probability
#' @param ellipse_size the size of the ellipse line
#' @param ellipse_alpha the alpha (transparency) of the ellipse line
#' @param points_size the size of the points
#' @param points_alpha the alpha (transparency) of the points
#' @param arrows a logical to indicate whether arrows should be drawn
#' @param arrows_textsize the size of the text for variable names
#' @param arrows_colour the colour of the arrow and their text
#' @param arrows_size the size (thickness) of the arrow lines
#' @param arrows_textsize the size of the text at the end of the arrows
#' @param arrows_textangled a logical whether the text at the end of the arrows should be angled
#' @param arrows_alpha the alpha (transparency) of the arrows and their text
#' @param base_textsize the text size for all plot elements except the labels and arrows
#' @param ... Arguments passed on to functions
#' @source The [ggplot_pca()] function is based on the `ggbiplot()` function from the `ggbiplot` package by Vince Vu, as found on GitHub: <https://github.com/vqv/ggbiplot> (retrieved: 2 March 2020, their latest commit: [`7325e88`](https://github.com/vqv/ggbiplot/commit/7325e880485bea4c07465a0304c470608fffb5d9); 12 February 2015).
#' 
#' As per their GPL-2 licence that demands documentation of code changes, the changes made based on the source code were: 
#' 1. Rewritten code to remove the dependency on packages `plyr`, `scales` and `grid`
#' 2. Parametrised more options, like arrow and ellipse settings
#' 3. Hardened all input possibilities by defining the exact type of user input for every argument
#' 4. Added total amount of explained variance as a caption in the plot
#' 5. Cleaned all syntax based on the `lintr` package, fixed grammatical errors and added integrity checks
#' 6. Updated documentation
#' @details The colours for labels and points can be changed by adding another scale layer for colour, like `scale_colour_viridis_d()` or `scale_colour_brewer()`.
#' @rdname ggplot_pca
#' @export
#' @examples 
#' # `example_isolates` is a dataset available in the AMR package.
#' # See ?example_isolates.
#'
#' # See ?pca for more info about Principal Component Analysis (PCA).
#' if (require("dplyr")) {
#'   pca_model <- example_isolates %>% 
#'     filter(mo_genus(mo) == "Staphylococcus") %>% 
#'     group_by(species = mo_shortname(mo)) %>%
#'     summarise_if (is.rsi, resistance) %>%
#'     pca(FLC, AMC, CXM, GEN, TOB, TMP, SXT, CIP, TEC, TCY, ERY)
#'     
#'   # old (base R)
#'   biplot(pca_model)
#'   
#'   # new 
#'   ggplot_pca(pca_model)
#'   
#'   if (require("ggplot2")) {
#'     ggplot_pca(pca_model) +
#'       scale_colour_viridis_d() +
#'       labs(title = "Title here")
#'   }
#' }
ggplot_pca <- function(x,
                       choices = 1:2,
                       scale = 1,
                       pc.biplot = TRUE,
                       labels = NULL,
                       labels_textsize = 3,
                       labels_text_placement = 1.5,
                       groups = NULL,
                       ellipse = TRUE,
                       ellipse_prob = 0.68,
                       ellipse_size = 0.5,
                       ellipse_alpha = 0.5,
                       points_size = 2,
                       points_alpha = 0.25,
                       arrows = TRUE,
                       arrows_colour = "darkblue",
                       arrows_size = 0.5,
                       arrows_textsize = 3,
                       arrows_textangled = TRUE,
                       arrows_alpha = 0.75,
                       base_textsize = 10,
                       ...) {
  
  stop_ifnot_installed("ggplot2")
  meet_criteria(x, allow_class = c("prcomp", "princomp", "PCA", "lda"))
  meet_criteria(choices, allow_class = c("numeric", "integer"), has_length = 2)
  meet_criteria(scale, allow_class = c("numeric", "integer", "logical"), has_length = 1)
  meet_criteria(pc.biplot, allow_class = "logical", has_length = 1)
  meet_criteria(labels, allow_class = "character", allow_NULL = TRUE)
  meet_criteria(labels_textsize, allow_class = c("numeric", "integer"), has_length = 1)
  meet_criteria(labels_text_placement, allow_class = c("numeric", "integer"), has_length = 1)
  meet_criteria(groups, allow_class = "character", allow_NULL = TRUE)
  meet_criteria(ellipse, allow_class = "logical", has_length = 1)
  meet_criteria(ellipse_prob, allow_class = c("numeric", "integer"), has_length = 1)
  meet_criteria(ellipse_size, allow_class = c("numeric", "integer"), has_length = 1)
  meet_criteria(ellipse_alpha, allow_class = c("numeric", "integer"), has_length = 1)
  meet_criteria(points_size, allow_class = c("numeric", "integer"), has_length = 1)
  meet_criteria(points_alpha, allow_class = c("numeric", "integer"), has_length = 1)
  meet_criteria(arrows, allow_class = "logical", has_length = 1)
  meet_criteria(arrows_colour, allow_class = "character", has_length = 1)
  meet_criteria(arrows_size, allow_class = c("numeric", "integer"), has_length = 1)
  meet_criteria(arrows_textsize, allow_class = c("numeric", "integer"), has_length = 1)
  meet_criteria(arrows_textangled, allow_class = "logical", has_length = 1)
  meet_criteria(arrows_alpha, allow_class = c("numeric", "integer"), has_length = 1)
  meet_criteria(base_textsize, allow_class = c("numeric", "integer"), has_length = 1)
  
  calculations <- pca_calculations(pca_model = x,
                                   groups = groups, 
                                   groups_missing = missing(groups),
                                   labels = labels,
                                   labels_missing = missing(labels),
                                   choices = choices,
                                   scale = scale,
                                   pc.biplot = pc.biplot,
                                   ellipse_prob = ellipse_prob,
                                   labels_text_placement = labels_text_placement)
  choices <- calculations$choices
  df.u <- calculations$df.u
  df.v <- calculations$df.v
  ell <- calculations$ell
  groups <- calculations$groups
  group_name <- calculations$group_name
  labels <- calculations$labels
  
  # Append the proportion of explained variance to the axis labels
  if ((1 - as.integer(scale)) == 0) {
    u.axis.labs <- paste0("Standardised PC", choices)
  } else {
    u.axis.labs <- paste0("PC", choices)
  }
  u.axis.labs <- paste0(u.axis.labs,
                        paste0("\n(explained var: ", 
                               percentage(x$sdev[choices] ^ 2 / sum(x$sdev ^ 2)),
                               ")"))
  
  # Score Labels
  if (!is.null(labels)) {
    df.u$labels <- labels
  }
  
  # Grouping variable
  if (!is.null(groups)) {
    df.u$groups <- groups
  }
  
  # Base plot
  g <- ggplot2::ggplot(data = df.u, 
                       ggplot2::aes(x = xvar, y = yvar)) + 
    ggplot2::xlab(u.axis.labs[1]) + 
    ggplot2::ylab(u.axis.labs[2]) + 
    ggplot2::expand_limits(x = c(-1.15, 1.15),
                           y = c(-1.15, 1.15))
  
  # Draw either labels or points
  if (!is.null(df.u$labels)) {
    if (!is.null(df.u$groups)) {
      g <- g + ggplot2::geom_point(ggplot2::aes(colour = groups),
                                   alpha = points_alpha, 
                                   size = points_size) +
        ggplot2::geom_text(ggplot2::aes(label = labels, colour = groups),
                           nudge_y = -0.05,
                           size = labels_textsize) +
        ggplot2::labs(colour = group_name)
    } else {
      g <- g + ggplot2::geom_point(alpha = points_alpha, 
                                   size = points_size) +
        ggplot2::geom_text(ggplot2::aes(label = labels),
                           nudge_y = -0.05,
                           size = labels_textsize)      
    }
  } else {
    if (!is.null(df.u$groups)) {
      g <- g + ggplot2::geom_point(ggplot2::aes(colour = groups),
                                   alpha = points_alpha,
                                   size = points_size) +
        ggplot2::labs(colour = group_name)
    } else {
      g <- g + ggplot2::geom_point(alpha = points_alpha,
                                   size = points_size)      
    }
  }
  
  # Overlay a concentration ellipse if there are groups
  if (!is.null(df.u$groups) & !is.null(ell) & isTRUE(ellipse)) {
    g <- g + ggplot2::geom_path(data = ell, 
                                ggplot2::aes(colour = groups, group = groups),
                                size = ellipse_size,
                                alpha = points_alpha)
  }
  
  # Label the variable axes
  if (arrows == TRUE) {
    g <- g + ggplot2::geom_segment(data = df.v,
                                   ggplot2::aes(x = 0, y = 0, xend = xvar, yend = yvar),
                                   arrow = ggplot2::arrow(length = ggplot2::unit(0.5, "picas"),
                                                          angle = 20,
                                                          ends = "last",
                                                          type = "open"), 
                                   colour = arrows_colour, 
                                   size = arrows_size,
                                   alpha = arrows_alpha)
    if (arrows_textangled == TRUE) {
      g <- g + ggplot2::geom_text(data = df.v, 
                                  ggplot2::aes(label = varname, x = xvar, y = yvar, angle = angle, hjust = hjust), 
                                  colour = arrows_colour,
                                  size = arrows_textsize,
                                  alpha = arrows_alpha)
    } else {
      g <- g + ggplot2::geom_text(data = df.v, 
                                  ggplot2::aes(label = varname, x = xvar, y = yvar, hjust = hjust), 
                                  colour = arrows_colour,
                                  size = arrows_textsize,
                                  alpha = arrows_alpha)
    }
  }
  
  # Add caption label about total explained variance
  g <- g + ggplot2::labs(caption = paste0("Total explained variance: ",
                                          percentage(sum(x$sdev[choices] ^ 2 / sum(x$sdev ^ 2)))))
  
  # mark-up nicely
  g <- g + ggplot2::theme_minimal(base_size = base_textsize) +
    ggplot2::theme(panel.grid.major = ggplot2::element_line(colour = "grey85"),
                   panel.grid.minor = ggplot2::element_blank(),
                   # centre title and subtitle
                   plot.title = ggplot2::element_text(hjust = 0.5),
                   plot.subtitle = ggplot2::element_text(hjust = 0.5))
  
  g
}

#' @importFrom stats qchisq var
pca_calculations <- function(pca_model,
                             groups = NULL,
                             groups_missing = TRUE,
                             labels = NULL,
                             labels_missing = TRUE,
                             choices = 1:2,
                             scale = 1,
                             pc.biplot = TRUE,
                             ellipse_prob = 0.68,
                             labels_text_placement = 1.5) {
  
  non_numeric_cols <- attributes(pca_model)$non_numeric_cols
  if (groups_missing) {
    groups <- tryCatch(non_numeric_cols[[1]],
                       error = function(e) NULL)
    group_name <- tryCatch(colnames(non_numeric_cols[1]),
                           error = function(e) NULL)
  }
  if (labels_missing) {
    labels <- tryCatch(non_numeric_cols[[2]],
                       error = function(e) NULL)
  }
  if (!is.null(groups) & is.null(labels)) {
    # turn them around
    labels <- groups
    groups <- NULL
    group_name <- NULL
  }
  
  # Recover the SVD
  if (inherits(pca_model, "prcomp")) {
    nobs.factor <- sqrt(nrow(pca_model$x) - 1)
    d <- pca_model$sdev
    u <- sweep(pca_model$x, 2, 1 / (d * nobs.factor), FUN = "*")
    v <- pca_model$rotation
  } else if (inherits(pca_model, "princomp")) {
    nobs.factor <- sqrt(pca_model$n.obs)
    d <- pca_model$sdev
    u <- sweep(pca_model$scores, 2, 1 / (d * nobs.factor), FUN = "*")
    v <- pca_model$loadings
  } else if (inherits(pca_model, "PCA")) {
    nobs.factor <- sqrt(nrow(pca_model$call$X))
    d <- unlist(sqrt(pca_model$eig)[1])
    u <- sweep(pca_model$ind$coord, 2, 1 / (d * nobs.factor), FUN = "*")
    v <- sweep(pca_model$var$coord, 2, sqrt(pca_model$eig[seq_len(ncol(pca_model$var$coord)), 1]), FUN = "/")
  } else if (inherits(pca_model, "lda")) {
    nobs.factor <- sqrt(pca_model$N)
    d <- pca_model$svd
    u <- predict(pca_model)$x / nobs.factor
    v <- pca_model$scaling
  } else {
    stop("Expected an object of class prcomp, princomp, PCA, or lda")
  }
  
  # Scores
  choices <- pmin(choices, ncol(u))
  obs.scale <- 1 - as.integer(scale)
  df.u <- as.data.frame(sweep(u[, choices], 2, d[choices] ^ obs.scale, FUN = "*"),
                        stringsAsFactors = FALSE)
  
  # Directions
  v <- sweep(v, 2, d ^ as.integer(scale), FUN = "*")
  df.v <- as.data.frame(v[, choices],
                        stringsAsFactors = FALSE)
  
  names(df.u) <- c("xvar", "yvar")
  names(df.v) <- names(df.u)
  
  if (isTRUE(pc.biplot)) {
    df.u <- df.u * nobs.factor
  }
  
  # Scale the radius of the correlation circle so that it corresponds to 
  # a data ellipse for the standardized PC scores
  circle_prob <- 0.69
  r <- sqrt(qchisq(circle_prob, df = 2)) * prod(colMeans(df.u ^ 2)) ^ (0.25)
  
  # Scale directions
  v.scale <- rowSums(v ^ 2)
  df.v <- r * df.v / sqrt(max(v.scale))
  
  # Grouping variable
  if (!is.null(groups)) {
    df.u$groups <- groups
  }
  
  df.v$varname <- rownames(v)
  
  # Variables for text label placement
  df.v$angle <- with(df.v, (180 / pi) * atan(yvar / xvar))
  df.v$hjust <- with(df.v, (1 - labels_text_placement * sign(xvar)) / 2)
  
  if (!is.null(df.u$groups)) {
    theta <- c(seq(-pi, pi, length = 50), seq(pi, -pi, length = 50))
    circle <- cbind(cos(theta), sin(theta))
    
    df.groups <- lapply(unique(df.u$groups), function(g, df = df.u) {
      x <- df[which(df$groups == g), , drop = FALSE]
      if (nrow(x) <= 2) {
        return(data.frame(X1 = numeric(0),
                          X2 = numeric(0),
                          groups = character(0),
                          stringsAsFactors = FALSE))
      }
      sigma <- var(cbind(x$xvar, x$yvar))
      mu <- c(mean(x$xvar), mean(x$yvar))
      ed <- sqrt(qchisq(ellipse_prob, df = 2))
      data.frame(sweep(circle %*% chol(sigma) * ed,
                       MARGIN = 2,
                       STATS = mu, 
                       FUN = "+"), 
                 groups = x$groups[1],
                 stringsAsFactors = FALSE)
    })
    ell <- do.call(rbind, df.groups)
    if (NROW(ell) == 0) {
      ell <- NULL
    } else {
      names(ell)[1:2] <- c("xvar", "yvar")
    }
  } else {
    ell <- NULL
  }
  
  list(choices = choices,
       df.u = df.u,
       df.v = df.v,
       ell = ell,
       groups = groups,
       group_name = group_name,
       labels = labels
  )
}
```
