## Supplementary material for "AMR - An R Package for Working with Antimicrobial Resistance Data": Source package: ggplot_rsi.R

```
# ==================================================================== #
# TITLE                                                                #
# Antimicrobial Resistance (AMR) Analysis for R                        #
#                                                                      #
# SOURCE                                                               #
# https://github.com/msberends/AMR                                     #
#                                                                      #
# LICENCE                                                              #
# (c) 2018-2021 Berends MS, Luz CF et al.                              #
# Developed at the University of Groningen, the Netherlands, in        #
# collaboration with non-profit organisations Certe Medical            #
# Diagnostics & Advice, and University Medical Center Groningen.       # 
#                                                                      #
# This R package is free software; you can freely use and distribute   #
# it for both personal and commercial purposes under the terms of the  #
# GNU General Public License version 2.0 (GNU GPL-2), as published by  #
# the Free Software Foundation.                                        #
# We created this package for both routine data analysis and academic  #
# research and it was publicly released in the hope that it will be    #
# useful, but it comes WITHOUT ANY WARRANTY OR LIABILITY.              #
#                                                                      #
# Visit our website for the full manual and a complete tutorial about  #
# how to conduct AMR analysis: https://msberends.github.io/AMR/        #
# ==================================================================== #

#' AMR plots with `ggplot2`
#'
#' Use these functions to create bar plots for antimicrobial resistance analysis. All functions rely on [ggplot2][ggplot2::ggplot()] functions.
#' @inheritSection lifecycle Maturing lifecycle
#' @param data a [data.frame] with column(s) of class [`rsi`] (see [as.rsi()])
#' @param position position adjustment of bars, either `"fill"`, `"stack"` or `"dodge"`
#' @param x variable to show on x axis, either `"antibiotic"` (default) or `"interpretation"` or a grouping variable
#' @param fill variable to categorise using the plots legend, either `"antibiotic"` (default) or `"interpretation"` or a grouping variable
#' @param breaks numeric vector of positions
#' @param limits numeric vector of length two providing limits of the scale, use `NA` to refer to the existing minimum or maximum
#' @param facet variable to split plots by, either `"interpretation"` (default) or `"antibiotic"` or a grouping variable
#' @inheritParams proportion
#' @param nrow (when using `facet`) number of rows
#' @param colours a named vector with colours for the bars. The names must be one or more of: S, SI, I, IR, R or be `FALSE` to use default [ggplot2][ggplot2::ggplot()] colours.
#' @param datalabels show datalabels using [labels_rsi_count()]
#' @param datalabels.size size of the datalabels
#' @param datalabels.colour colour of the datalabels
#' @param title text to show as title of the plot
#' @param subtitle text to show as subtitle of the plot
#' @param caption text to show as caption of the plot
#' @param x.title text to show as x axis description
#' @param y.title text to show as y axis description
#' @param ... other arguments passed on to [geom_rsi()]
#' @details At default, the names of antibiotics will be shown on the plots using [ab_name()]. This can be set with the `translate_ab` argument. See [count_df()].
#'
#' ## The functions
#' [geom_rsi()] will take any variable from the data that has an [`rsi`] class (created with [as.rsi()]) using [rsi_df()] and will plot bars with the percentage R, I and S. The default behaviour is to have the bars stacked and to have the different antibiotics on the x axis.
#'
#' [facet_rsi()] creates 2d plots (at default based on S/I/R) using [ggplot2::facet_wrap()].
#'
#' [scale_y_percent()] transforms the y axis to a 0 to 100% range using [ggplot2::scale_y_continuous()].
#'
#' [scale_rsi_colours()] sets colours to the bars: pastel blue for S, pastel turquoise for I and pastel red for R, using [ggplot2::scale_fill_manual()].
#'
#' [theme_rsi()] is a [ggplot2 theme][[ggplot2::theme()] with minimal distraction.
#'
#' [labels_rsi_count()] print datalabels on the bars with percentage and amount of isolates using [ggplot2::geom_text()].
#'
#' [ggplot_rsi()] is a wrapper around all above functions that uses data as first input. This makes it possible to use this function after a pipe (`%>%`). See Examples.
#' @rdname ggplot_rsi
#' @export
#' @inheritSection AMR Read more on our website!
#' @examples
#' if (require("ggplot2") & require("dplyr")) {
#'  
#'   # get antimicrobial results for drugs against a UTI:
#'   ggplot(example_isolates %>% select(AMX, NIT, FOS, TMP, CIP)) +
#'     geom_rsi()
#'  
#'   # prettify the plot using some additional functions:
#'   df <- example_isolates %>% select(AMX, NIT, FOS, TMP, CIP)
#'   ggplot(df) +
#'     geom_rsi() +
#'     scale_y_percent() +
#'     scale_rsi_colours() +
#'     labels_rsi_count() +
#'     theme_rsi()
#'  
#'   # or better yet, simplify this using the wrapper function - a single command:
#'   example_isolates %>%
#'     select(AMX, NIT, FOS, TMP, CIP) %>%
#'     ggplot_rsi()
#'  
#'   # get only proportions and no counts:
#'   example_isolates %>%
#'     select(AMX, NIT, FOS, TMP, CIP) %>%
#'     ggplot_rsi(datalabels = FALSE)
#'  
#'   # add other ggplot2 arguments as you like:
#'   example_isolates %>%
#'     select(AMX, NIT, FOS, TMP, CIP) %>%
#'     ggplot_rsi(width = 0.5,
#'                colour = "black",
#'                size = 1,
#'                linetype = 2,
#'                alpha = 0.25)
#'  
#'   example_isolates %>%
#'     select(AMX) %>%
#'     ggplot_rsi(colours = c(SI = "yellow"))
#'   
#' }
#'   
#' \donttest{
#' # resistance of ciprofloxacine per age group
#' example_isolates %>%
#'   mutate(first_isolate = first_isolate(.)) %>%
#'   filter(first_isolate == TRUE,
#'          mo == as.mo("E. coli")) %>%
#'   # age_groups() is also a function in this AMR package:
#'   group_by(age_group = age_groups(age)) %>%
#'   select(age_group,
#'          CIP) %>%
#'   ggplot_rsi(x = "age_group")
#'   
#' # for colourblind mode, use divergent colours from the viridis package:
#' example_isolates %>%
#'   select(AMX, NIT, FOS, TMP, CIP) %>%
#'   ggplot_rsi() + 
#'   scale_fill_viridis_d()
#' # a shorter version which also adjusts data label colours:
#' example_isolates %>%
#'   select(AMX, NIT, FOS, TMP, CIP) %>%
#'   ggplot_rsi(colours = FALSE)
#'
#'
#' # it also supports groups (don't forget to use the group var on `x` or `facet`):
#' example_isolates %>%
#'   select(hospital_id, AMX, NIT, FOS, TMP, CIP) %>%
#'   group_by(hospital_id) %>%
#'   ggplot_rsi(x = "hospital_id",
#'              facet = "antibiotic",
#'              nrow = 1,
#'              title = "AMR of Anti-UTI Drugs Per Hospital",
#'              x.title = "Hospital",
#'              datalabels = FALSE)
#' }
ggplot_rsi <- function(data,
                       position = NULL,
                       x = "antibiotic",
                       fill = "interpretation",
                       # params = list(),
                       facet = NULL,
                       breaks = seq(0, 1, 0.1),
                       limits = NULL,
                       translate_ab = "name",
                       combine_SI = TRUE,
                       combine_IR = FALSE,
                       minimum = 30,
                       language = get_locale(),
                       nrow = NULL,
                       colours = c(S = "#61a8ff",
                                   SI = "#61a8ff",
                                   I = "#61f7ff",
                                   IR = "#ff6961",
                                   R = "#ff6961"),
                       datalabels = TRUE,
                       datalabels.size = 2.5,
                       datalabels.colour = "grey15",
                       title = NULL,
                       subtitle = NULL,
                       caption = NULL,
                       x.title = "Antimicrobial",
                       y.title = "Proportion",
                       ...) {
  
  stop_ifnot_installed("ggplot2")
  meet_criteria(data, allow_class = "data.frame", contains_column_class = "rsi")
  meet_criteria(position, allow_class = "character", has_length = 1, is_in = c("fill", "stack", "dodge"), allow_NULL = TRUE)
  meet_criteria(x, allow_class = "character", has_length = 1)
  meet_criteria(fill, allow_class = "character", has_length = 1)
  meet_criteria(facet, allow_class = "character", has_length = 1, allow_NULL = TRUE)
  meet_criteria(breaks, allow_class = c("numeric", "integer"))
  meet_criteria(limits, allow_class = c("numeric", "integer"), has_length = 2, allow_NULL = TRUE, allow_NA = TRUE)
  meet_criteria(translate_ab, allow_class = c("character", "logical"), has_length = 1, allow_NA = TRUE)
  meet_criteria(combine_SI, allow_class = "logical", has_length = 1)
  meet_criteria(combine_IR, allow_class = "logical", has_length = 1)
  meet_criteria(minimum, allow_class = c("numeric", "integer"), has_length = 1)
  meet_criteria(language, has_length = 1, is_in = c(LANGUAGES_SUPPORTED, ""), allow_NULL = TRUE, allow_NA = TRUE)
  meet_criteria(nrow, allow_class = c("numeric", "integer"), has_length = 1, allow_NULL = TRUE)
  meet_criteria(colours, allow_class = c("character", "logical"))
  meet_criteria(datalabels, allow_class = "logical", has_length = 1)
  meet_criteria(datalabels.size, allow_class = c("numeric", "integer"), has_length = 1)
  meet_criteria(datalabels.colour, allow_class = "character", has_length = 1)
  meet_criteria(title, allow_class = "character", has_length = 1, allow_NULL = TRUE)
  meet_criteria(subtitle, allow_class = "character", has_length = 1, allow_NULL = TRUE)
  meet_criteria(caption, allow_class = "character", has_length = 1, allow_NULL = TRUE)
  meet_criteria(x.title, allow_class = "character", has_length = 1, allow_NULL = TRUE)
  meet_criteria(y.title, allow_class = "character", has_length = 1, allow_NULL = TRUE)

  # we work with aes_string later on
  x_deparse <- deparse(substitute(x))
  if (x_deparse != "x") {
    x <- x_deparse
  }
  if (x %like% '".*"') {
    x <- substr(x, 2, nchar(x) - 1)
  }
  facet_deparse <- deparse(substitute(facet))
  if (facet_deparse != "facet") {
    facet <- facet_deparse
  }
  if (facet %like% '".*"') {
    facet <- substr(facet, 2, nchar(facet) - 1)
  }
  if (facet %in% c("NULL", "")) {
    facet <- NULL
  }
  
  if (is.null(position)) {
    position <- "fill"
  }
  
  p <- ggplot2::ggplot(data = data) +
    geom_rsi(position = position, x = x, fill = fill, translate_ab = translate_ab,
             minimum = minimum, language = language,
             combine_SI = combine_SI, combine_IR = combine_IR, ...) +
    theme_rsi()
  
  if (fill == "interpretation") {
    # set RSI colours
    if (isFALSE(colours) & missing(datalabels.colour)) {
      # set datalabel colour to middle grey
      datalabels.colour <- "grey50"
    }
    p <- p + scale_rsi_colours(colours = colours)
  }
  
  if (identical(position, "fill")) {
    # proportions, so use y scale with percentage
    p <- p + scale_y_percent(breaks = breaks, limits = limits)
  }
  
  if (datalabels == TRUE) {
    p <- p + labels_rsi_count(position = position,
                              x = x,
                              translate_ab = translate_ab,
                              minimum = minimum,
                              language = language,
                              combine_SI = combine_SI,
                              combine_IR = combine_IR,
                              datalabels.size = datalabels.size,
                              datalabels.colour = datalabels.colour)
  }
  
  if (!is.null(facet)) {
    p <- p + facet_rsi(facet = facet, nrow = nrow)
  }
  
  p <- p + ggplot2::labs(title = title,
                         subtitle = subtitle,
                         caption = caption,
                         x = x.title,
                         y = y.title)
  
  p
}

#' @rdname ggplot_rsi
#' @export
geom_rsi <- function(position = NULL,
                     x = c("antibiotic", "interpretation"),
                     fill = "interpretation",
                     translate_ab = "name",
                     minimum = 30,
                     language = get_locale(),
                     combine_SI = TRUE,
                     combine_IR = FALSE,
                     ...)  {
  x <- x[1]
  stop_ifnot_installed("ggplot2")
  stop_if(is.data.frame(position), "`position` is invalid. Did you accidentally use '%>%' instead of '+'?")
  meet_criteria(position, allow_class = "character", has_length = 1, is_in = c("fill", "stack", "dodge"), allow_NULL = TRUE)
  meet_criteria(x, allow_class = "character", has_length = 1)
  meet_criteria(fill, allow_class = "character", has_length = 1)
  meet_criteria(translate_ab, allow_class = c("character", "logical"), has_length = 1, allow_NA = TRUE)
  meet_criteria(minimum, allow_class = c("numeric", "integer"), has_length = 1)
  meet_criteria(language, has_length = 1, is_in = c(LANGUAGES_SUPPORTED, ""), allow_NULL = TRUE, allow_NA = TRUE)
  meet_criteria(combine_SI, allow_class = "logical", has_length = 1)
  meet_criteria(combine_IR, allow_class = "logical", has_length = 1)
  
  y <- "value"
  if (missing(position) | is.null(position)) {
    position <- "fill"
  }
  
  if (identical(position, "fill")) {
    position <- ggplot2::position_fill(vjust = 0.5, reverse = TRUE)
  }
  
  # we work with aes_string later on
  x_deparse <- deparse(substitute(x))
  if (x_deparse != "x") {
    x <- x_deparse
  }
  if (x %like% '".*"') {
    x <- substr(x, 2, nchar(x) - 1)
  }
  
  if (tolower(x) %in% tolower(c("ab", "abx", "antibiotics"))) {
    x <- "antibiotic"
  } else if (tolower(x) %in% tolower(c("SIR", "RSI", "interpretations", "result"))) {
    x <- "interpretation"
  }
  
  ggplot2::layer(geom = "bar", stat = "identity", position = position,
                 mapping = ggplot2::aes_string(x = x, y = y, fill = fill),
                 params = list(...), data = function(x) {
                   rsi_df(data = x,
                          translate_ab = translate_ab,
                          language = language,
                          minimum = minimum,
                          combine_SI = combine_SI,
                          combine_IR = combine_IR)
                 })
  
}

#' @rdname ggplot_rsi
#' @export
facet_rsi <- function(facet = c("interpretation", "antibiotic"), nrow = NULL) {
  facet <- facet[1]
  stop_ifnot_installed("ggplot2")
  meet_criteria(facet, allow_class = "character", has_length = 1)
  meet_criteria(nrow, allow_class = c("numeric", "integer"), has_length = 1, allow_NULL = TRUE)
  
  # we work with aes_string later on
  facet_deparse <- deparse(substitute(facet))
  if (facet_deparse != "facet") {
    facet <- facet_deparse
  }
  if (facet %like% '".*"') {
    facet <- substr(facet, 2, nchar(facet) - 1)
  }
  
  if (tolower(facet) %in% tolower(c("SIR", "RSI", "interpretations", "result"))) {
    facet <- "interpretation"
  } else if (tolower(facet) %in% tolower(c("ab", "abx", "antibiotics"))) {
    facet <- "antibiotic"
  }
  
  ggplot2::facet_wrap(facets = facet, scales = "free_x", nrow = nrow)
}

#' @rdname ggplot_rsi
#' @export
scale_y_percent <- function(breaks = seq(0, 1, 0.1), limits = NULL) {
  stop_ifnot_installed("ggplot2")
  meet_criteria(breaks, allow_class = c("numeric", "integer"))
  meet_criteria(limits, allow_class = c("numeric", "integer"), has_length = 2, allow_NULL = TRUE, allow_NA = TRUE)
  
  if (all(breaks[breaks != 0] > 1)) {
    breaks <- breaks / 100
  }
  ggplot2::scale_y_continuous(breaks = breaks,
                              labels = percentage(breaks),
                              limits = limits)
}

#' @rdname ggplot_rsi
#' @export
scale_rsi_colours <- function(colours = c(S = "#61a8ff",
                                          SI = "#61a8ff",
                                          I = "#61f7ff",
                                          IR = "#ff6961",
                                          R = "#ff6961")) {
  stop_ifnot_installed("ggplot2")
  meet_criteria(colours, allow_class = c("character", "logical"))
  
  # previous colour: palette = "RdYlGn"
  # previous colours: values = c("#b22222", "#ae9c20", "#7cfc00")
  
  if (!identical(colours, FALSE)) {
    original_cols <- c(S = "#61a8ff",
                       SI = "#61a8ff",
                       I = "#61f7ff",
                       IR = "#ff6961",
                       R = "#ff6961")
    colours <- replace(original_cols, names(colours), colours)
    ggplot2::scale_fill_manual(values = colours)
  }
}

#' @rdname ggplot_rsi
#' @export
theme_rsi <- function() {
  stop_ifnot_installed("ggplot2")
  ggplot2::theme_minimal(base_size = 10) +
    ggplot2::theme(panel.grid.major.x = ggplot2::element_blank(),
                   panel.grid.minor = ggplot2::element_blank(),
                   panel.grid.major.y = ggplot2::element_line(colour = "grey75"),
                   # center title and subtitle
                   plot.title = ggplot2::element_text(hjust = 0.5),
                   plot.subtitle = ggplot2::element_text(hjust = 0.5))
}

#' @rdname ggplot_rsi
#' @export
labels_rsi_count <- function(position = NULL,
                             x = "antibiotic",
                             translate_ab = "name",
                             minimum = 30,
                             language = get_locale(),
                             combine_SI = TRUE,
                             combine_IR = FALSE,
                             datalabels.size = 3,
                             datalabels.colour = "grey15") {
  stop_ifnot_installed("ggplot2")
  meet_criteria(position, allow_class = "character", has_length = 1, is_in = c("fill", "stack", "dodge"), allow_NULL = TRUE)
  meet_criteria(x, allow_class = "character", has_length = 1)
  meet_criteria(translate_ab, allow_class = c("character", "logical"), has_length = 1, allow_NA = TRUE)
  meet_criteria(minimum, allow_class = c("numeric", "integer"), has_length = 1)
  meet_criteria(language, has_length = 1, is_in = c(LANGUAGES_SUPPORTED, ""), allow_NULL = TRUE, allow_NA = TRUE)
  meet_criteria(combine_SI, allow_class = "logical", has_length = 1)
  meet_criteria(combine_IR, allow_class = "logical", has_length = 1)
  meet_criteria(datalabels.size, allow_class = c("numeric", "integer"), has_length = 1)
  meet_criteria(datalabels.colour, allow_class = "character", has_length = 1)
  
  if (is.null(position)) {
    position <- "fill"
  }
  if (identical(position, "fill")) {
    position <- ggplot2::position_fill(vjust = 0.5, reverse = TRUE)
  }
  x_name <- x
  ggplot2::geom_text(mapping = ggplot2::aes_string(label = "lbl",
                                                   x = x,
                                                   y = "value"),
                     position = position,
                     inherit.aes = FALSE,
                     size = datalabels.size,
                     colour = datalabels.colour,
                     lineheight = 0.75,
                     data = function(x) {
                       transformed <- rsi_df(data = x,
                                             translate_ab = translate_ab,
                                             combine_SI = combine_SI,
                                             combine_IR = combine_IR,
                                             minimum = minimum,
                                             language = language)
                       transformed$gr <- transformed[, x_name, drop = TRUE]
                       transformed %pm>% 
                         pm_group_by(gr) %pm>% 
                         pm_mutate(lbl = paste0("n=", isolates)) %pm>% 
                         pm_ungroup() %pm>% 
                         pm_select(-gr)
                     })
}
```
