## Supplementary material for "AMR - An R Package for Working with Antimicrobial Resistance Data": Source package: globals.R

```
# ==================================================================== #
# TITLE                                                                #
# Antimicrobial Resistance (AMR) Analysis for R                        #
#                                                                      #
# SOURCE                                                               #
# https://github.com/msberends/AMR                                     #
#                                                                      #
# LICENCE                                                              #
# (c) 2018-2021 Berends MS, Luz CF et al.                              #
# Developed at the University of Groningen, the Netherlands, in        #
# collaboration with non-profit organisations Certe Medical            #
# Diagnostics & Advice, and University Medical Center Groningen.       # 
#                                                                      #
# This R package is free software; you can freely use and distribute   #
# it for both personal and commercial purposes under the terms of the  #
# GNU General Public License version 2.0 (GNU GPL-2), as published by  #
# the Free Software Foundation.                                        #
# We created this package for both routine data analysis and academic  #
# research and it was publicly released in the hope that it will be    #
# useful, but it comes WITHOUT ANY WARRANTY OR LIABILITY.              #
#                                                                      #
# Visit our website for the full manual and a complete tutorial about  #
# how to conduct AMR analysis: https://msberends.github.io/AMR/        #
# ==================================================================== #

globalVariables(c(".rowid",
                  "ab",
                  "ab_txt",
                  "angle",
                  "antibiotic",
                  "antibiotics",
                  "atc_group1",
                  "atc_group2",
                  "code",
                  "data",
                  "fullname",
                  "fullname_lower",
                  "g_species",
                  "genus",
                  "gr",
                  "group",
                  "guideline",
                  "hjust",
                  "input",
                  "intrinsic_resistant",
                  "isolates",
                  "lang",
                  "language",
                  "lookup",
                  "method",
                  "microorganism",
                  "microorganisms",
                  "microorganisms.codes",
                  "microorganisms.old",
                  "mo",
                  "name",
                  "new",
                  "observations",
                  "old",
                  "old_name",
                  "pattern",
                  "R",
                  "reference.rule",
                  "reference.rule_group",
                  "reference.version",
                  "rsi_translation",
                  "rowid",
                  "rule_group",
                  "rule_name",
                  "se_max",
                  "se_min",
                  "species",
                  "species_id",
                  "total",
                  "txt",
                  "value",
                  "varname",
                  "xvar",
                  "y",
                  "year",
                  "yvar"))
```
