## Supplementary material for "AMR - An R Package for Working with Antimicrobial Resistance Data": Source package: guess_ab_col.R

```
# ==================================================================== #
# TITLE                                                                #
# Antimicrobial Resistance (AMR) Analysis for R                        #
#                                                                      #
# SOURCE                                                               #
# https://github.com/msberends/AMR                                     #
#                                                                      #
# LICENCE                                                              #
# (c) 2018-2021 Berends MS, Luz CF et al.                              #
# Developed at the University of Groningen, the Netherlands, in        #
# collaboration with non-profit organisations Certe Medical            #
# Diagnostics & Advice, and University Medical Center Groningen.       # 
#                                                                      #
# This R package is free software; you can freely use and distribute   #
# it for both personal and commercial purposes under the terms of the  #
# GNU General Public License version 2.0 (GNU GPL-2), as published by  #
# the Free Software Foundation.                                        #
# We created this package for both routine data analysis and academic  #
# research and it was publicly released in the hope that it will be    #
# useful, but it comes WITHOUT ANY WARRANTY OR LIABILITY.              #
#                                                                      #
# Visit our website for the full manual and a complete tutorial about  #
# how to conduct AMR analysis: https://msberends.github.io/AMR/        #
# ==================================================================== #

#' Guess antibiotic column
#'
#' This tries to find a column name in a data set based on information from the [antibiotics] data set. Also supports WHONET abbreviations.
#' @inheritSection lifecycle Stable lifecycle
#' @param x a [data.frame]
#' @param search_string a text to search `x` for, will be checked with [as.ab()] if this value is not a column in `x`
#' @param verbose a logical to indicate whether additional info should be printed
#' @details You can look for an antibiotic (trade) name or abbreviation and it will search `x` and the [antibiotics] data set for any column containing a name or code of that antibiotic. **Longer columns names take precedence over shorter column names.**
#' @return A column name of `x`, or `NULL` when no result is found.
#' @export
#' @inheritSection AMR Read more on our website!
#' @examples
#' df <- data.frame(amox = "S",
#'                  tetr = "R")
#'
#' guess_ab_col(df, "amoxicillin")
#' # [1] "amox"
#' guess_ab_col(df, "J01AA07") # ATC code of tetracycline
#' # [1] "tetr"
#'
#' guess_ab_col(df, "J01AA07", verbose = TRUE)
#' # NOTE: Using column 'tetr' as input for J01AA07 (tetracycline).
#' # [1] "tetr"
#'
#' # WHONET codes
#' df <- data.frame(AMP_ND10 = "R",
#'                  AMC_ED20 = "S")
#' guess_ab_col(df, "ampicillin")
#' # [1] "AMP_ND10"
#' guess_ab_col(df, "J01CR02")
#' # [1] "AMC_ED20"
#' guess_ab_col(df, as.ab("augmentin"))
#' # [1] "AMC_ED20"
#'
#' # Longer names take precendence:
#' df <- data.frame(AMP_ED2 = "S",
#'                  AMP_ED20 = "S")
#' guess_ab_col(df, "ampicillin")
#' # [1] "AMP_ED20"
guess_ab_col <- function(x = NULL, search_string = NULL, verbose = FALSE) {
  meet_criteria(x, allow_class = "data.frame", allow_NULL = TRUE)
  meet_criteria(search_string, allow_class = "character", has_length = 1, allow_NULL = TRUE)
  meet_criteria(verbose, allow_class = "logical", has_length = 1)
  
  if (is.null(x) & is.null(search_string)) {
    return(as.name("guess_ab_col"))
  }
  
  if (search_string %in% colnames(x)) {
    ab_result <- search_string
  } else {
    search_string.ab <- suppressWarnings(as.ab(search_string))
    if (search_string.ab %in% colnames(x)) {
      ab_result <- colnames(x)[colnames(x) == search_string.ab][1L]
      
    } else if (any(tolower(colnames(x)) %in% tolower(unlist(ab_property(search_string.ab, "abbreviations", language = NULL))))) {
      ab_result <- colnames(x)[tolower(colnames(x)) %in% tolower(unlist(ab_property(search_string.ab, "abbreviations", language = NULL)))][1L]
      
    } else {
      # sort colnames on length - longest first
      cols <- colnames(x[, x %pm>% colnames() %pm>% nchar() %pm>% order() %pm>% rev()])
      df_trans <- data.frame(cols = cols,
                             abs = suppressWarnings(as.ab(cols)),
                             stringsAsFactors = FALSE)
      ab_result <- df_trans[which(df_trans$abs == search_string.ab), "cols"]
      ab_result <- ab_result[!is.na(ab_result)][1L]
    }
  }
  
  if (length(ab_result) == 0) {
    if (verbose == TRUE) {
      message_("No column found as input for ", search_string,
               " (", ab_name(search_string, language = NULL, tolower = TRUE), ").",
               add_fn = font_black,
               as_note = FALSE)
    }
    return(NULL)
  } else {
    if (verbose == TRUE) {
      message_("Using column '", font_bold(ab_result), "' as input for ", search_string,
               " (", ab_name(search_string, language = NULL, tolower = TRUE), ").")
    }
    return(ab_result)
  }
}

get_column_abx <- function(x,
                           soft_dependencies = NULL,
                           hard_dependencies = NULL,
                           verbose = FALSE,
                           info = TRUE,
                           ...) {
  meet_criteria(x, allow_class = "data.frame")
  meet_criteria(soft_dependencies, allow_class = "character", allow_NULL = TRUE)
  meet_criteria(hard_dependencies, allow_class = "character", allow_NULL = TRUE)
  meet_criteria(verbose, allow_class = "logical", has_length = 1)
  meet_criteria(info, allow_class = "logical", has_length = 1)
  
  if (info == TRUE) {
    message_("Auto-guessing columns suitable for analysis", appendLF = FALSE)
  }
  
  x <- as.data.frame(x, stringsAsFactors = FALSE)
  if (NROW(x) > 10000) {
    # only test maximum of 10,000 values per column
    if (info == TRUE) {
      message_(" (using only ", font_bold("the first 10,000 rows"), ")...",
               appendLF = FALSE, 
               as_note = FALSE)
    }
    x <- x[1:10000, , drop = FALSE]
  } else if (info == TRUE) {
    message_("...", appendLF = FALSE, as_note = FALSE)
  }
  x_bak <- x
  # only check columns that are a valid AB code, ATC code, name, abbreviation or synonym,
  # or already have the <rsi> class (as.rsi) 
  # and that they have no more than 50% invalid values
  vectr_antibiotics <- unique(toupper(unlist(antibiotics[, c("ab", "atc", "name", "abbreviations", "synonyms")])))
  vectr_antibiotics <- vectr_antibiotics[!is.na(vectr_antibiotics) & nchar(vectr_antibiotics) >= 3]
  x_columns <- vapply(FUN.VALUE = character(1), colnames(x), function(col, df = x_bak) {
    if (toupper(col) %in% vectr_antibiotics || 
        is.rsi(as.data.frame(df, stringsAsFactors = FALSE)[, col, drop = TRUE]) ||
        is.rsi.eligible(as.data.frame(df, stringsAsFactors = FALSE)[, col, drop = TRUE],
                        threshold = 0.5)) {
      return(col)
    } else {
      return(NA_character_)
    }
  })
  x_columns <- x_columns[!is.na(x_columns)]
  x <- x[, x_columns, drop = FALSE] # without drop = TRUE, x will become a vector when x_columns is length 1
  
  df_trans <- data.frame(colnames = colnames(x),
                         abcode = suppressWarnings(as.ab(colnames(x), info = FALSE)),
                         stringsAsFactors = FALSE)
  df_trans <- df_trans[!is.na(df_trans$abcode), , drop = FALSE]
  x <- as.character(df_trans$colnames)
  names(x) <- df_trans$abcode
  
  # add from self-defined dots (...):
  # such as get_column_abx(example_isolates %pm>% rename(thisone = AMX), amox = "thisone")
  dots <- list(...)
  if (length(dots) > 0) {
    newnames <- suppressWarnings(as.ab(names(dots), info = FALSE))
    if (any(is.na(newnames))) {
      warning_("Invalid antibiotic reference(s): ", toString(names(dots)[is.na(newnames)]),
               call = FALSE,
               immediate = TRUE)
    }
    # turn all NULLs to NAs
    dots <- unlist(lapply(dots, function(x) if (is.null(x)) NA else x))
    names(dots) <- newnames
    dots <- dots[!is.na(names(dots))]
    # merge, but overwrite automatically determined ones by 'dots'
    x <- c(x[!x %in% dots & !names(x) %in% names(dots)], dots)
    # delete NAs, this will make e.g. eucast_rules(... TMP = NULL) work to prevent TMP from being used
    x <- x[!is.na(x)]
  }
  
  if (length(x) == 0) {
    if (info == TRUE) {
      message_("No columns found.")
    }
    return(x)
  }
  
  # sort on name
  x <- x[order(names(x), x)]
  duplicates <- c(x[duplicated(x)], x[duplicated(names(x))]) 
  duplicates <- duplicates[unique(names(duplicates))]
  x <- c(x[!names(x) %in% names(duplicates)], duplicates)
  x <- x[order(names(x), x)]
  
  # succeeded with auto-guessing
  if (info == TRUE) {
    message_(" OK.", add_fn = list(font_green, font_bold), as_note = FALSE)
  }
  
  for (i in seq_len(length(x))) {
    if (info == TRUE & verbose == TRUE & !names(x[i]) %in% names(duplicates)) {
      message_("Using column '", font_bold(x[i]), "' as input for ", names(x)[i],
               " (", ab_name(names(x)[i], tolower = TRUE, language = NULL), ").")
    }
    if (info == TRUE & names(x[i]) %in% names(duplicates)) {
      warning_(paste0("Using column '", font_bold(x[i]), "' as input for ", names(x)[i],
                      " (", ab_name(names(x)[i], tolower = TRUE, language = NULL),
                      "), although it was matched for multiple antibiotics or columns."),
               add_fn = font_red,
               call = FALSE, 
               immediate = verbose)
    }
  }
  
  
  if (!is.null(hard_dependencies)) {
    hard_dependencies <- unique(hard_dependencies)
    if (!all(hard_dependencies %in% names(x))) {
      # missing a hard dependency will return NA and consequently the data will not be analysed
      missing <- hard_dependencies[!hard_dependencies %in% names(x)]
      generate_warning_abs_missing(missing, any = FALSE)
      return(NA)
    }
  }
  if (!is.null(soft_dependencies)) {
    soft_dependencies <- unique(soft_dependencies)
    if (info == TRUE & !all(soft_dependencies %in% names(x))) {
      # missing a soft dependency may lower the reliability
      missing <- soft_dependencies[!soft_dependencies %in% names(x)]
      missing_msg <- paste(paste0(ab_name(missing, tolower = TRUE, language = NULL), 
                                  " (", font_bold(missing, collapse = NULL), ")"), 
                           collapse = ", ")
      message_("Reliability would be improved if these antimicrobial results would be available too: ",
               missing_msg)
    }
  }
  x
}

generate_warning_abs_missing <- function(missing, any = FALSE) {
  missing <- paste0(missing, " (", ab_name(missing, tolower = TRUE, language = NULL), ")")
  if (any == TRUE) {
    any_txt <- c(" any of", "is")
  } else {
    any_txt <- c("", "are")
  }
  warning_(paste0("Introducing NAs since", any_txt[1], " these antimicrobials ", any_txt[2], " required: ",
                  paste(missing, collapse = ", ")),
           immediate = TRUE,
           call = FALSE)
}
```
