## Supplementary material for "AMR - An R Package for Working with Antimicrobial Resistance Data": Source package: join_microorganisms.R

```
# ==================================================================== #
# TITLE                                                                #
# Antimicrobial Resistance (AMR) Analysis for R                        #
#                                                                      #
# SOURCE                                                               #
# https://github.com/msberends/AMR                                     #
#                                                                      #
# LICENCE                                                              #
# (c) 2018-2021 Berends MS, Luz CF et al.                              #
# Developed at the University of Groningen, the Netherlands, in        #
# collaboration with non-profit organisations Certe Medical            #
# Diagnostics & Advice, and University Medical Center Groningen.       # 
#                                                                      #
# This R package is free software; you can freely use and distribute   #
# it for both personal and commercial purposes under the terms of the  #
# GNU General Public License version 2.0 (GNU GPL-2), as published by  #
# the Free Software Foundation.                                        #
# We created this package for both routine data analysis and academic  #
# research and it was publicly released in the hope that it will be    #
# useful, but it comes WITHOUT ANY WARRANTY OR LIABILITY.              #
#                                                                      #
# Visit our website for the full manual and a complete tutorial about  #
# how to conduct AMR analysis: https://msberends.github.io/AMR/        #
# ==================================================================== #

#' Join [microorganisms] to a data set
#'
#' Join the data set [microorganisms] easily to an existing table or character vector.
#' @inheritSection lifecycle Stable lifecycle
#' @rdname join
#' @name join
#' @aliases join inner_join
#' @param x existing table to join, or character vector
#' @param by a variable to join by - if left empty will search for a column with class [`mo`] (created with [as.mo()]) or will be `"mo"` if that column name exists in `x`, could otherwise be a column name of `x` with values that exist in `microorganisms$mo` (such as `by = "bacteria_id"`), or another column in [microorganisms] (but then it should be named, like `by = c("bacteria_id" = "fullname")`)
#' @param suffix if there are non-joined duplicate variables in `x` and `y`, these suffixes will be added to the output to disambiguate them. Should be a character vector of length 2.
#' @param ... ignored
#' @details **Note:** As opposed to the `join()` functions of `dplyr`, [character] vectors are supported and at default existing columns will get a suffix `"2"` and the newly joined columns will not get a suffix. 
#' 
#' If the `dplyr` package is installed, their join functions will be used. Otherwise, the much slower [merge()] function from base R will be used.
#' @inheritSection AMR Read more on our website!
#' @export
#' @examples
#' left_join_microorganisms(as.mo("K. pneumoniae"))
#' left_join_microorganisms("B_KLBSL_PNE")
#'
#' \donttest{
#' if (require("dplyr")) {
#'   example_isolates %>%
#'     left_join_microorganisms() %>% 
#'     colnames()
#'  
#'   df <- data.frame(date = seq(from = as.Date("2018-01-01"),
#'                               to = as.Date("2018-01-07"),
#'                               by = 1),
#'                    bacteria = as.mo(c("S. aureus", "MRSA", "MSSA", "STAAUR",
#'                                       "E. coli", "E. coli", "E. coli")),
#'                    stringsAsFactors = FALSE)
#'   colnames(df)
#'   df_joined <- left_join_microorganisms(df, "bacteria")
#'   colnames(df_joined)
#' }
#' }
inner_join_microorganisms <- function(x, by = NULL, suffix = c("2", ""), ...) {
  meet_criteria(x, allow_class = c("data.frame", "character"))
  meet_criteria(by, allow_class = "character", allow_NULL = TRUE)
  meet_criteria(suffix, allow_class = "character", has_length = 2)
  
  check_dataset_integrity()
  x <- check_groups_before_join(x, "inner_join_microorganisms")
  checked <- joins_check_df(x, by)
  x_class <- get_prejoined_class(x)
  x <- checked$x
  by <- checked$by
  # use dplyr if available - it's much faster
  dplyr_inner <- import_fn("inner_join", "dplyr", error_on_fail = FALSE)
  if (!is.null(dplyr_inner)) {
    join <- suppressWarnings(
      dplyr_inner(x = x, y = microorganisms, by = by, suffix = suffix, ...)
    )
  } else {
    join <- suppressWarnings(
      pm_inner_join(x = x, y = microorganisms, by = by, suffix = suffix, ...)
    )
  }
  if (NROW(join) > NROW(x)) {
    warning_("The newly joined tbl contains ", nrow(join) - nrow(x), " rows more that its original.")
  }
  class(join) <- x_class
  join
}

#' @rdname join
#' @export
left_join_microorganisms <- function(x, by = NULL, suffix = c("2", ""), ...) {
  meet_criteria(x, allow_class = c("data.frame", "character"))
  meet_criteria(by, allow_class = "character", allow_NULL = TRUE)
  meet_criteria(suffix, allow_class = "character", has_length = 2)
  
  check_dataset_integrity()
  x <- check_groups_before_join(x, "left_join_microorganisms")
  checked <- joins_check_df(x, by)
  x_class <- get_prejoined_class(x)
  x <- checked$x
  by <- checked$by
  # use dplyr if available - it's much faster
  dplyr_left <- import_fn("left_join", "dplyr", error_on_fail = FALSE)
  if (!is.null(dplyr_left)) {
    join <- suppressWarnings(
      dplyr_left(x = x, y = microorganisms, by = by, suffix = suffix, ...)
    )
  } else {
    join <- suppressWarnings(
      pm_left_join(x = x, y = microorganisms, by = by, suffix = suffix, ...)
    )
  }
  if (NROW(join) > NROW(x)) {
    warning_("The newly joined tbl contains ", nrow(join) - nrow(x), " rows more that its original.")
  }
  class(join) <- x_class
  join
}

#' @rdname join
#' @export
right_join_microorganisms <- function(x, by = NULL, suffix = c("2", ""), ...) {
  meet_criteria(x, allow_class = c("data.frame", "character"))
  meet_criteria(by, allow_class = "character", allow_NULL = TRUE)
  meet_criteria(suffix, allow_class = "character", has_length = 2)
  
  check_dataset_integrity()
  x <- check_groups_before_join(x, "right_join_microorganisms")
  checked <- joins_check_df(x, by)
  x_class <- get_prejoined_class(x)
  x <- checked$x
  by <- checked$by
  # use dplyr if available - it's much faster
  dplyr_right <- import_fn("right_join", "dplyr", error_on_fail = FALSE)
  if (!is.null(dplyr_right)) {
    join <- suppressWarnings(
      dplyr_right(x = x, y = microorganisms, by = by, suffix = suffix, ...)
    )
  } else {
    join <- suppressWarnings(
      pm_right_join(x = x, y = microorganisms, by = by, suffix = suffix, ...)
    )
  }
  if (NROW(join) > NROW(x)) {
    warning_("The newly joined tbl contains ", nrow(join) - nrow(x), " rows more that its original.")
  }
  class(join) <- x_class
  join
}

#' @rdname join
#' @export
full_join_microorganisms <- function(x, by = NULL, suffix = c("2", ""), ...) {
  meet_criteria(x, allow_class = c("data.frame", "character"))
  meet_criteria(by, allow_class = "character", allow_NULL = TRUE)
  meet_criteria(suffix, allow_class = "character", has_length = 2)
  
  check_dataset_integrity()
  x <- check_groups_before_join(x, "full_join_microorganisms")
  checked <- joins_check_df(x, by)
  x_class <- get_prejoined_class(x)
  x <- checked$x
  by <- checked$by
  # use dplyr if available - it's much faster
  dplyr_full <- import_fn("full_join", "dplyr", error_on_fail = FALSE)
  if (!is.null(dplyr_full)) {
    join <- suppressWarnings(
      dplyr_full(x = x, y = microorganisms, by = by, suffix = suffix, ...)
    )
  } else {
    join <- suppressWarnings(
      pm_full_join(x = x, y = microorganisms, by = by, suffix = suffix, ...)
    )
  }
  if (NROW(join) > NROW(x)) {
    warning_("The newly joined tbl contains ", nrow(join) - nrow(x), " rows more that its original.")
  }
  class(join) <- x_class
  join
}

#' @rdname join
#' @export
semi_join_microorganisms <- function(x, by = NULL, ...) {
  meet_criteria(x, allow_class = c("data.frame", "character"))
  meet_criteria(by, allow_class = "character", allow_NULL = TRUE)
  
  check_dataset_integrity()
  x <- check_groups_before_join(x, "semi_join_microorganisms")
  x_class <- get_prejoined_class(x)
  checked <- joins_check_df(x, by)
  x <- checked$x
  by <- checked$by
  # use dplyr if available - it's much faster
  dplyr_semi <- import_fn("semi_join", "dplyr", error_on_fail = FALSE)
  if (!is.null(dplyr_semi)) {
    join <- suppressWarnings(
      dplyr_semi(x = x, y = microorganisms, by = by, ...)
    )
  } else {
    join <- suppressWarnings(
      pm_semi_join(x = x, y = microorganisms, by = by, ...)
    )
  }
  class(join) <- x_class
  join
}

#' @rdname join
#' @export
anti_join_microorganisms <- function(x, by = NULL, ...) {
  meet_criteria(x, allow_class = c("data.frame", "character"))
  meet_criteria(by, allow_class = "character", allow_NULL = TRUE)
  
  check_dataset_integrity()
  x <- check_groups_before_join(x, "anti_join_microorganisms")
  checked <- joins_check_df(x, by)
  x_class <- get_prejoined_class(x)
  x <- checked$x
  by <- checked$by
  # use dplyr if available - it's much faster
  dplyr_anti <- import_fn("anti_join", "dplyr", error_on_fail = FALSE)
  if (!is.null(dplyr_anti)) {
    join <- suppressWarnings(
      dplyr_anti(x = x, y = microorganisms, by = by, ...)
    )
  } else {
    join <- suppressWarnings(
      pm_anti_join(x = x, y = microorganisms, by = by, ...)
    )
  }
  class(join) <- x_class
  join
}

joins_check_df <- function(x, by) {
  if (!any(class(x) %in% c("data.frame", "matrix"))) {
    x <- data.frame(mo = as.mo(x), stringsAsFactors = FALSE)
    if (is.null(by)) {
      by <- "mo"
    }
  }
  x <- as.data.frame(x, stringsAsFactors = FALSE)
  if (is.null(by)) {
    # search for column with class `mo` and return first one found
    by <- colnames(x)[lapply(x, is.mo) == TRUE][1]
    if (is.na(by)) {
      if ("mo" %in% colnames(x)) {
        by <- "mo"
        x[, "mo"] <- as.mo(x[, "mo"])
      } else {
        stop("Cannot join - no column found with name 'mo' or with class <mo>.", call. = FALSE)
      }
    }
    message_('Joining, by = "', by, '"', add_fn = font_black, as_note = FALSE) # message same as dplyr::join functions
  }
  if (is.null(names(by))) {
    joinby <- colnames(microorganisms)[1]
    names(joinby) <- by
  } else {
    joinby <- by
  }
  list(x = x,
       by = joinby)
}

get_prejoined_class <- function(x) {
  if (is.data.frame(x)) {
    class(x)
  } else {
    "data.frame"
  }
}

check_groups_before_join <- function(x, fn) {
  if (is.data.frame(x) && !is.null(attributes(x)$groups)) {
    x <- pm_ungroup(x)
    attr(x, "groups") <- NULL
    class(x) <- class(x)[!class(x) %like% "group"]
    warning_("Groups are dropped, since the ", fn, "() function relies on merge() from base R.", call = FALSE)
  }
  x
}
```
