## Supplementary material for "AMR - An R Package for Working with Antimicrobial Resistance Data": Source package: key_antibiotics.R

```
# ==================================================================== #
# TITLE                                                                #
# Antimicrobial Resistance (AMR) Analysis for R                        #
#                                                                      #
# SOURCE                                                               #
# https://github.com/msberends/AMR                                     #
#                                                                      #
# LICENCE                                                              #
# (c) 2018-2021 Berends MS, Luz CF et al.                              #
# Developed at the University of Groningen, the Netherlands, in        #
# collaboration with non-profit organisations Certe Medical            #
# Diagnostics & Advice, and University Medical Center Groningen.       # 
#                                                                      #
# This R package is free software; you can freely use and distribute   #
# it for both personal and commercial purposes under the terms of the  #
# GNU General Public License version 2.0 (GNU GPL-2), as published by  #
# the Free Software Foundation.                                        #
# We created this package for both routine data analysis and academic  #
# research and it was publicly released in the hope that it will be    #
# useful, but it comes WITHOUT ANY WARRANTY OR LIABILITY.              #
#                                                                      #
# Visit our website for the full manual and a complete tutorial about  #
# how to conduct AMR analysis: https://msberends.github.io/AMR/        #
# ==================================================================== #

#' Key antibiotics for first *weighted* isolates
#'
#' These function can be used to determine first isolates (see [first_isolate()]). Using key antibiotics to determine first isolates is more reliable than without key antibiotics. These selected isolates can then be called first *weighted* isolates.
#' @inheritSection lifecycle Stable lifecycle
#' @param x a [data.frame] with antibiotics columns, like `AMX` or `amox`. Can be left blank when used inside `dplyr` verbs, such as `filter()`, `mutate()` and `summarise()`.
#' @param y,z character vectors to compare
#' @inheritParams first_isolate
#' @param universal_1,universal_2,universal_3,universal_4,universal_5,universal_6 column names of **broad-spectrum** antibiotics, case-insensitive. See details for which antibiotics will be used at default (which are guessed with [guess_ab_col()]).
#' @param GramPos_1,GramPos_2,GramPos_3,GramPos_4,GramPos_5,GramPos_6 column names of antibiotics for **Gram-positives**, case-insensitive. See details for which antibiotics will be used at default (which are guessed with [guess_ab_col()]).
#' @param GramNeg_1,GramNeg_2,GramNeg_3,GramNeg_4,GramNeg_5,GramNeg_6 column names of antibiotics for **Gram-negatives**, case-insensitive. See details for which antibiotics will be used at default (which are guessed with [guess_ab_col()]).
#' @param warnings give a warning about missing antibiotic columns (they will be ignored)
#' @param ... other arguments passed on to functions
#' @details 
#' The [key_antibiotics()] function is context-aware when used inside `dplyr` verbs, such as `filter()`, `mutate()` and `summarise()`. This means that then the `x` argument can be left blank, please see *Examples*.
#' 
#' The function [key_antibiotics()] returns a character vector with 12 antibiotic results for every isolate. These isolates can then be compared using [key_antibiotics_equal()], to check if two isolates have generally the same antibiogram. Missing and invalid values are replaced with a dot (`"."`) by [key_antibiotics()] and ignored by [key_antibiotics_equal()].
#' 
#' The [first_isolate()] function only uses this function on the same microbial species from the same patient. Using this, e.g. an MRSA will be included after a susceptible *S. aureus* (MSSA) is found within the same patient episode. Without key antibiotic comparison it would not. See [first_isolate()] for more info.
#'
#' At default, the antibiotics that are used for **Gram-positive bacteria** are:
#' - Amoxicillin
#' - Amoxicillin/clavulanic acid
#' - Cefuroxime
#' - Piperacillin/tazobactam
#' - Ciprofloxacin
#' - Trimethoprim/sulfamethoxazole
#' - Vancomycin
#' - Teicoplanin
#' - Tetracycline
#' - Erythromycin
#' - Oxacillin
#' - Rifampin
#'
#' At default the antibiotics that are used for **Gram-negative bacteria** are:
#' - Amoxicillin
#' - Amoxicillin/clavulanic acid
#' - Cefuroxime
#' - Piperacillin/tazobactam
#' - Ciprofloxacin
#' - Trimethoprim/sulfamethoxazole
#' - Gentamicin
#' - Tobramycin
#' - Colistin
#' - Cefotaxime
#' - Ceftazidime
#' - Meropenem
#' 
#' The function [key_antibiotics_equal()] checks the characters returned by [key_antibiotics()] for equality, and returns a [`logical`] vector.
#' @inheritSection first_isolate Key antibiotics
#' @rdname key_antibiotics
#' @export
#' @seealso [first_isolate()]
#' @inheritSection AMR Read more on our website!
#' @examples
#' # `example_isolates` is a dataset available in the AMR package.
#' # See ?example_isolates.
#' 
#' # output of the `key_antibiotics()` function could be like this:
#' strainA <- "SSSRR.S.R..S"
#' strainB <- "SSSIRSSSRSSS"
#'
#' # those strings can be compared with:
#' key_antibiotics_equal(strainA, strainB)
#' # TRUE, because I is ignored (as well as missing values)
#'
#' key_antibiotics_equal(strainA, strainB, ignore_I = FALSE)
#' # FALSE, because I is not ignored and so the 4th character differs
#'
#' \donttest{
#' if (require("dplyr")) {
#'   # set key antibiotics to a new variable
#'   my_patients <- example_isolates %>%
#'     mutate(keyab = key_antibiotics()) %>% # no need to define `x`
#'     mutate(
#'       # now calculate first isolates
#'       first_regular = first_isolate(col_keyantibiotics = FALSE),
#'       # and first WEIGHTED isolates
#'       first_weighted = first_isolate(col_keyantibiotics = "keyab")
#'     )
#'  
#'   # Check the difference, in this data set it results in a lot more isolates:
#'   sum(my_patients$first_regular, na.rm = TRUE)
#'   sum(my_patients$first_weighted, na.rm = TRUE)
#' }
#' }
key_antibiotics <- function(x,
                            col_mo = NULL,
                            universal_1 = guess_ab_col(x, "amoxicillin"),
                            universal_2 = guess_ab_col(x, "amoxicillin/clavulanic acid"),
                            universal_3 = guess_ab_col(x, "cefuroxime"),
                            universal_4 = guess_ab_col(x, "piperacillin/tazobactam"),
                            universal_5 = guess_ab_col(x, "ciprofloxacin"),
                            universal_6 = guess_ab_col(x, "trimethoprim/sulfamethoxazole"),
                            GramPos_1 = guess_ab_col(x, "vancomycin"),
                            GramPos_2 = guess_ab_col(x, "teicoplanin"),
                            GramPos_3 = guess_ab_col(x, "tetracycline"),
                            GramPos_4 = guess_ab_col(x, "erythromycin"),
                            GramPos_5 = guess_ab_col(x, "oxacillin"),
                            GramPos_6 = guess_ab_col(x, "rifampin"),
                            GramNeg_1 = guess_ab_col(x, "gentamicin"),
                            GramNeg_2 = guess_ab_col(x, "tobramycin"),
                            GramNeg_3 = guess_ab_col(x, "colistin"),
                            GramNeg_4 = guess_ab_col(x, "cefotaxime"),
                            GramNeg_5 = guess_ab_col(x, "ceftazidime"),
                            GramNeg_6 = guess_ab_col(x, "meropenem"),
                            warnings = TRUE,
                            ...) {
  if (missing(x)) {
    x <- get_current_data(arg_name = "x", call = -2)
  }
  meet_criteria(x, allow_class = "data.frame")
  meet_criteria(col_mo, allow_class = "character", has_length = 1, allow_NULL = TRUE, allow_NA = TRUE)
  meet_criteria(universal_1, allow_class = "character", has_length = 1, allow_NULL = TRUE, allow_NA = TRUE)
  meet_criteria(universal_2, allow_class = "character", has_length = 1, allow_NULL = TRUE, allow_NA = TRUE)
  meet_criteria(universal_3, allow_class = "character", has_length = 1, allow_NULL = TRUE, allow_NA = TRUE)
  meet_criteria(universal_4, allow_class = "character", has_length = 1, allow_NULL = TRUE, allow_NA = TRUE)
  meet_criteria(universal_5, allow_class = "character", has_length = 1, allow_NULL = TRUE, allow_NA = TRUE)
  meet_criteria(universal_6, allow_class = "character", has_length = 1, allow_NULL = TRUE, allow_NA = TRUE)
  meet_criteria(GramPos_1, allow_class = "character", has_length = 1, allow_NULL = TRUE, allow_NA = TRUE)
  meet_criteria(GramPos_2, allow_class = "character", has_length = 1, allow_NULL = TRUE, allow_NA = TRUE)
  meet_criteria(GramPos_3, allow_class = "character", has_length = 1, allow_NULL = TRUE, allow_NA = TRUE)
  meet_criteria(GramPos_4, allow_class = "character", has_length = 1, allow_NULL = TRUE, allow_NA = TRUE)
  meet_criteria(GramPos_5, allow_class = "character", has_length = 1, allow_NULL = TRUE, allow_NA = TRUE)
  meet_criteria(GramPos_6, allow_class = "character", has_length = 1, allow_NULL = TRUE, allow_NA = TRUE)
  meet_criteria(GramNeg_1, allow_class = "character", has_length = 1, allow_NULL = TRUE, allow_NA = TRUE)
  meet_criteria(GramNeg_2, allow_class = "character", has_length = 1, allow_NULL = TRUE, allow_NA = TRUE)
  meet_criteria(GramNeg_3, allow_class = "character", has_length = 1, allow_NULL = TRUE, allow_NA = TRUE)
  meet_criteria(GramNeg_4, allow_class = "character", has_length = 1, allow_NULL = TRUE, allow_NA = TRUE)
  meet_criteria(GramNeg_5, allow_class = "character", has_length = 1, allow_NULL = TRUE, allow_NA = TRUE)
  meet_criteria(GramNeg_6, allow_class = "character", has_length = 1, allow_NULL = TRUE, allow_NA = TRUE)
  meet_criteria(warnings, allow_class = "logical", has_length = 1)
  
  dots <- unlist(list(...))
  if (length(dots) != 0) {
    # backwards compatibility with old arguments
    dots.names <- dots %pm>% names()
    if ("info" %in% dots.names) {
      warnings <- dots[which(dots.names == "info")]
    }
  }
  
  # try to find columns based on type
  # -- mo
  if (is.null(col_mo)) {
    col_mo <- search_type_in_df(x = x, type = "mo")
  }
  stop_if(is.null(col_mo), "`col_mo` must be set")
  
  # check columns
  col.list <- c(universal_1, universal_2, universal_3, universal_4, universal_5, universal_6,
                GramPos_1, GramPos_2, GramPos_3, GramPos_4, GramPos_5, GramPos_6,
                GramNeg_1, GramNeg_2, GramNeg_3, GramNeg_4, GramNeg_5, GramNeg_6)
  check_available_columns <- function(x, col.list, warnings = TRUE) {
    # check columns
    col.list <- col.list[!is.na(col.list) & !is.null(col.list)]
    names(col.list) <- col.list
    col.list.bak <- col.list
    # are they available as upper case or lower case then?
    for (i in seq_len(length(col.list))) {
      if (is.null(col.list[i]) | isTRUE(is.na(col.list[i]))) {
        col.list[i] <- NA
      } else if (toupper(col.list[i]) %in% colnames(x)) {
        col.list[i] <- toupper(col.list[i])
      } else if (tolower(col.list[i]) %in% colnames(x)) {
        col.list[i] <- tolower(col.list[i])
      } else if (!col.list[i] %in% colnames(x)) {
        col.list[i] <- NA
      }
    }
    if (!all(col.list %in% colnames(x))) {
      if (warnings == TRUE) {
        warning_("Some columns do not exist and will be ignored: ",
                 col.list.bak[!(col.list %in% colnames(x))] %pm>% toString(),
                 ".\nTHIS MAY STRONGLY INFLUENCE THE OUTCOME.",
                 immediate = TRUE,
                 call = FALSE)
      }
    }
    col.list
  }
  
  col.list <- check_available_columns(x = x, col.list = col.list, warnings = warnings)
  universal_1 <- col.list[universal_1]
  universal_2 <- col.list[universal_2]
  universal_3 <- col.list[universal_3]
  universal_4 <- col.list[universal_4]
  universal_5 <- col.list[universal_5]
  universal_6 <- col.list[universal_6]
  GramPos_1 <- col.list[GramPos_1]
  GramPos_2 <- col.list[GramPos_2]
  GramPos_3 <- col.list[GramPos_3]
  GramPos_4 <- col.list[GramPos_4]
  GramPos_5 <- col.list[GramPos_5]
  GramPos_6 <- col.list[GramPos_6]
  GramNeg_1 <- col.list[GramNeg_1]
  GramNeg_2 <- col.list[GramNeg_2]
  GramNeg_3 <- col.list[GramNeg_3]
  GramNeg_4 <- col.list[GramNeg_4]
  GramNeg_5 <- col.list[GramNeg_5]
  GramNeg_6 <- col.list[GramNeg_6]
  
  universal <- c(universal_1, universal_2, universal_3,
                 universal_4, universal_5, universal_6)
  
  gram_positive <- c(universal,
                     GramPos_1, GramPos_2, GramPos_3,
                     GramPos_4, GramPos_5, GramPos_6)
  gram_positive <- gram_positive[!is.null(gram_positive)]
  gram_positive <- gram_positive[!is.na(gram_positive)]
  if (length(gram_positive) < 12) {
    warning_("Only using ", length(gram_positive), " different antibiotics as key antibiotics for Gram-positives. See ?key_antibiotics.", call = FALSE)
  }
  
  gram_negative <- c(universal,
                     GramNeg_1, GramNeg_2, GramNeg_3,
                     GramNeg_4, GramNeg_5, GramNeg_6)
  gram_negative <- gram_negative[!is.null(gram_negative)]
  gram_negative <- gram_negative[!is.na(gram_negative)]
  if (length(gram_negative) < 12) {
    warning_("Only using ", length(gram_negative), " different antibiotics as key antibiotics for Gram-negatives. See ?key_antibiotics.", call = FALSE)
  }
  
  x <- as.data.frame(x, stringsAsFactors = FALSE)
  x[, col_mo] <- as.mo(x[, col_mo, drop = TRUE])
  x$gramstain <- mo_gramstain(x[, col_mo, drop = TRUE], language = NULL)
  x$key_ab <- NA_character_
  
  # Gram +
  x$key_ab <- pm_if_else(x$gramstain == "Gram-positive",
                      tryCatch(apply(X = x[, gram_positive],
                                     MARGIN = 1,
                                     FUN = function(x) paste(x, collapse = "")),
                               error = function(e) paste0(rep(".", 12), collapse = "")),
                      x$key_ab)
  
  # Gram -
  x$key_ab <- pm_if_else(x$gramstain == "Gram-negative",
                      tryCatch(apply(X = x[, gram_negative],
                                     MARGIN = 1,
                                     FUN = function(x) paste(x, collapse = "")),
                               error = function(e) paste0(rep(".", 12), collapse = "")),
                      x$key_ab)
  
  # format
  key_abs <- toupper(gsub("[^SIR]", ".", gsub("(NA|NULL)", ".", x$key_ab)))
  
  if (pm_n_distinct(key_abs) == 1) {
    warning_("No distinct key antibiotics determined.", call = FALSE)
  }
  
  key_abs
  
}

#' @rdname key_antibiotics
#' @export
key_antibiotics_equal <- function(y,
                                  z,
                                  type = c("keyantibiotics", "points"),
                                  ignore_I = TRUE,
                                  points_threshold = 2,
                                  info = FALSE) {
  meet_criteria(y, allow_class = "character")
  meet_criteria(z, allow_class = "character")
  meet_criteria(type, allow_class = "character", has_length = c(1, 2))
  meet_criteria(ignore_I, allow_class = "logical", has_length = 1)
  meet_criteria(points_threshold, allow_class = c("numeric", "integer"), has_length = 1)
  meet_criteria(info, allow_class = "logical", has_length = 1)
  
  stop_ifnot(length(y) == length(z), "length of `y` and `z` must be equal")
  # y is active row, z is lag
  x <- y
  y <- z
  
  type <- type[1]
  
  # only show progress bar on points or when at least 5000 isolates
  info_needed <- info == TRUE & (type == "points" | length(x) > 5000)
  
  result <- logical(length(x))
  
  if (info_needed == TRUE) {
    p <- progress_ticker(length(x))
    on.exit(close(p))
  }
  
  for (i in seq_len(length(x))) {
    
    if (info_needed == TRUE) {
      p$tick()
    }
    
    if (is.na(x[i])) {
      x[i] <- ""
    }
    if (is.na(y[i])) {
      y[i] <- ""
    }
    
    if (x[i] == y[i]) {
      
      result[i] <- TRUE
      
    } else if (nchar(x[i]) != nchar(y[i])) {
      
      result[i] <- FALSE
      
    } else {
      
      x_split <- strsplit(x[i], "")[[1]]
      y_split <- strsplit(y[i], "")[[1]]
      
      if (type == "keyantibiotics") {
        
        if (ignore_I == TRUE) {
          x_split[x_split == "I"] <- "."
          y_split[y_split == "I"] <- "."
        }
        
        y_split[x_split == "."] <- "."
        x_split[y_split == "."] <- "."
        
        result[i] <- all(x_split == y_split)
        
      } else if (type == "points") {
        # count points for every single character:
        # - no change is 0 points
        # - I <-> S|R is 0.5 point
        # - S|R <-> R|S is 1 point
        # use the levels of as.rsi (S = 1, I = 2, R = 3)
        
        suppressWarnings(x_split <- x_split %pm>% as.rsi() %pm>% as.double())
        suppressWarnings(y_split <- y_split %pm>% as.rsi() %pm>% as.double())
        
        points <- (x_split - y_split) %pm>% abs() %pm>% sum(na.rm = TRUE) / 2
        result[i] <- points >= points_threshold
        
      } else {
        stop("`", type, '` is not a valid value for type, must be "points" or "keyantibiotics". See ?key_antibiotics')
      }
    }
  }
  if (info_needed == TRUE) {
    close(p)
  }
  result
}
```
