## Supplementary material for "AMR - An R Package for Working with Antimicrobial Resistance Data": Source package: kurtosis.R

```
# ==================================================================== #
# TITLE                                                                #
# Antimicrobial Resistance (AMR) Analysis for R                        #
#                                                                      #
# SOURCE                                                               #
# https://github.com/msberends/AMR                                     #
#                                                                      #
# LICENCE                                                              #
# (c) 2018-2021 Berends MS, Luz CF et al.                              #
# Developed at the University of Groningen, the Netherlands, in        #
# collaboration with non-profit organisations Certe Medical            #
# Diagnostics & Advice, and University Medical Center Groningen.       # 
#                                                                      #
# This R package is free software; you can freely use and distribute   #
# it for both personal and commercial purposes under the terms of the  #
# GNU General Public License version 2.0 (GNU GPL-2), as published by  #
# the Free Software Foundation.                                        #
# We created this package for both routine data analysis and academic  #
# research and it was publicly released in the hope that it will be    #
# useful, but it comes WITHOUT ANY WARRANTY OR LIABILITY.              #
#                                                                      #
# Visit our website for the full manual and a complete tutorial about  #
# how to conduct AMR analysis: https://msberends.github.io/AMR/        #
# ==================================================================== #

#' Kurtosis of the sample
#'
#' @description Kurtosis is a measure of the "tailedness" of the probability distribution of a real-valued random variable. A normal distribution has a kurtosis of 3 and a excess kurtosis of 0.
#' @inheritSection lifecycle Stable lifecycle
#' @param x a vector of values, a [matrix] or a [data.frame]
#' @param na.rm a logical to indicate whether `NA` values should be stripped before the computation proceeds
#' @param excess a logical to indicate whether the *excess kurtosis* should be returned, defined as the kurtosis minus 3.
#' @seealso [skewness()]
#' @rdname kurtosis
#' @inheritSection AMR Read more on our website!
#' @export
kurtosis <- function(x, na.rm = FALSE, excess = FALSE) {
  meet_criteria(na.rm, allow_class = "logical", has_length = 1)
  meet_criteria(excess, allow_class = "logical", has_length = 1)
  UseMethod("kurtosis")
}

#' @method kurtosis default
#' @rdname kurtosis
#' @export
kurtosis.default <- function(x, na.rm = FALSE, excess = FALSE) {
  meet_criteria(na.rm, allow_class = "logical", has_length = 1)
  meet_criteria(excess, allow_class = "logical", has_length = 1)
  x <- as.vector(x)
  if (na.rm == TRUE) {
    x <- x[!is.na(x)]
  }
  n <- length(x)
  k <- n * sum((x - mean(x, na.rm = na.rm))^4, na.rm = na.rm) /
    (sum((x - mean(x, na.rm = na.rm))^2, na.rm = na.rm)^2)
  k - ifelse(excess, 3, 0)
}

#' @method kurtosis matrix
#' @rdname kurtosis
#' @export
kurtosis.matrix <- function(x, na.rm = FALSE, excess = FALSE) {
  meet_criteria(na.rm, allow_class = "logical", has_length = 1)
  meet_criteria(excess, allow_class = "logical", has_length = 1)
  apply(x, 2, kurtosis.default, na.rm = na.rm, excess = excess)
}

#' @method kurtosis data.frame
#' @rdname kurtosis
#' @export
kurtosis.data.frame <- function(x, na.rm = FALSE, excess = FALSE) {
  meet_criteria(na.rm, allow_class = "logical", has_length = 1)
  meet_criteria(excess, allow_class = "logical", has_length = 1)
  vapply(FUN.VALUE = double(1), x, kurtosis.default, na.rm = na.rm, excess = excess)
}
```
