## Supplementary material for "AMR - An R Package for Working with Antimicrobial Resistance Data": Source package: lifecycle.R

```
# ==================================================================== #
# TITLE                                                                #
# Antimicrobial Resistance (AMR) Analysis for R                        #
#                                                                      #
# SOURCE                                                               #
# https://github.com/msberends/AMR                                     #
#                                                                      #
# LICENCE                                                              #
# (c) 2018-2021 Berends MS, Luz CF et al.                              #
# Developed at the University of Groningen, the Netherlands, in        #
# collaboration with non-profit organisations Certe Medical            #
# Diagnostics & Advice, and University Medical Center Groningen.       # 
#                                                                      #
# This R package is free software; you can freely use and distribute   #
# it for both personal and commercial purposes under the terms of the  #
# GNU General Public License version 2.0 (GNU GPL-2), as published by  #
# the Free Software Foundation.                                        #
# We created this package for both routine data analysis and academic  #
# research and it was publicly released in the hope that it will be    #
# useful, but it comes WITHOUT ANY WARRANTY OR LIABILITY.              #
#                                                                      #
# Visit our website for the full manual and a complete tutorial about  #
# how to conduct AMR analysis: https://msberends.github.io/AMR/        #
# ==================================================================== #

###############
# NOTE TO SELF: could also have done this with the 'lifecycle' package, but why add a package dependency for such an easy job??
###############

#' Lifecycles of functions in the `AMR` package
#' @name lifecycle
#' @rdname lifecycle
#' @description Functions in this `AMR` package are categorised using [the lifecycle circle of the Tidyverse as found on www.tidyverse.org/lifecycle](https://www.Tidyverse.org/lifecycle).
#' 
#' \if{html}{\figure{lifecycle_tidyverse.svg}{options: height=200px style=margin-bottom:5px} \cr}
#' This page contains a section for every lifecycle (with text borrowed from the aforementioned Tidyverse website), so they can be used in the manual pages of the functions. 
#' @section Experimental lifecycle:
#' \if{html}{\figure{lifecycle_experimental.svg}{options: style=margin-bottom:5px} \cr}
#' The [lifecycle][AMR::lifecycle] of this function is **experimental**. An experimental function is in early stages of development. The unlying code might be changing frequently. Experimental functions might be removed without deprecation, so you are generally best off waiting until a function is more mature before you use it in production code. Experimental functions are only available in development versions of this `AMR` package and will thus not be included in releases that are submitted to CRAN, since such functions have not yet matured enough.
#' @section Maturing lifecycle:
#' \if{html}{\figure{lifecycle_maturing.svg}{options: style=margin-bottom:5px} \cr}
#' The [lifecycle][AMR::lifecycle] of this function is **maturing**. The unlying code of a maturing function has been roughed out, but finer details might still change. Since this function needs wider usage and more extensive testing, you are very welcome [to suggest changes at our repository](https://github.com/msberends/AMR/issues) or [write us an email (see section 'Contact Us')][AMR::AMR].
#' @section Stable lifecycle:
#' \if{html}{\figure{lifecycle_stable.svg}{options: style=margin-bottom:5px} \cr}
#' The [lifecycle][AMR::lifecycle] of this function is **stable**. In a stable function, major changes are unlikely. This means that the unlying code will generally evolve by adding new arguments; removing arguments or changing the meaning of existing arguments will be avoided.
#' 
#' If the unlying code needs breaking changes, they will occur gradually. For example, a argument will be deprecated and first continue to work, but will emit an message informing you of the change. Next, typically after at least one newly released version on CRAN, the message will be transformed to an error.
#' @section Retired lifecycle:
#' \if{html}{\figure{lifecycle_retired.svg}{options: style=margin-bottom:5px} \cr}
#' The [lifecycle][AMR::lifecycle] of this function is **retired**. A retired function is no longer under active development, and (if appropiate) a better alternative is available. No new arguments will be added, and only the most critical bugs will be fixed. In a future version, this function will be removed.
#' @section Questioning lifecycle:
#' \if{html}{\figure{lifecycle_questioning.svg}{options: style=margin-bottom:5px} \cr}
#' The [lifecycle][AMR::lifecycle] of this function is **questioning**. This function might be no longer be optimal approach, or is it questionable whether this function should be in this `AMR` package at all.
NULL
```
