## Supplementary material for "AMR - An R Package for Working with Antimicrobial Resistance Data": Source package: like.R

```
# ==================================================================== #
# TITLE                                                                #
# Antimicrobial Resistance (AMR) Analysis for R                        #
#                                                                      #
# SOURCE                                                               #
# https://github.com/msberends/AMR                                     #
#                                                                      #
# LICENCE                                                              #
# (c) 2018-2021 Berends MS, Luz CF et al.                              #
# Developed at the University of Groningen, the Netherlands, in        #
# collaboration with non-profit organisations Certe Medical            #
# Diagnostics & Advice, and University Medical Center Groningen.       # 
#                                                                      #
# This R package is free software; you can freely use and distribute   #
# it for both personal and commercial purposes under the terms of the  #
# GNU General Public License version 2.0 (GNU GPL-2), as published by  #
# the Free Software Foundation.                                        #
# We created this package for both routine data analysis and academic  #
# research and it was publicly released in the hope that it will be    #
# useful, but it comes WITHOUT ANY WARRANTY OR LIABILITY.              #
#                                                                      #
# Visit our website for the full manual and a complete tutorial about  #
# how to conduct AMR analysis: https://msberends.github.io/AMR/        #
# ==================================================================== #

#' Pattern matching with keyboard shortcut
#'
#' Convenient wrapper around [grep()] to match a pattern: `x %like% pattern`. It always returns a [`logical`] vector and is always case-insensitive (use `x %like_case% pattern` for case-sensitive matching). Also, `pattern` can be as long as `x` to compare items of each index in both vectors, or they both can have the same length to iterate over all cases.
#' @inheritSection lifecycle Stable lifecycle
#' @param x a character vector where matches are sought, or an object which can be coerced by [as.character()] to a character vector.
#' @param pattern a character string containing a regular expression (or [character] string for `fixed = TRUE`) to be matched in the given character vector. Coerced by [as.character()] to a character string if possible.  If a [character] vector of length 2 or more is supplied, the first element is used with a warning.
#' @param ignore.case if `FALSE`, the pattern matching is *case sensitive* and if `TRUE`, case is ignored during matching.
#' @return A [`logical`] vector
#' @name like
#' @rdname like
#' @export
#' @details
#' The `%like%` function:
#' * Is case-insensitive (use `%like_case%` for case-sensitive matching)
#' * Supports multiple patterns
#' * Checks if `pattern` is a regular expression and sets `fixed = TRUE` if not, to greatly improve speed
#' * Tries again with `perl = TRUE` if regex fails
#' 
#' Using RStudio? The text `%like%` can also be directly inserted in your code from the Addins menu and can have its own Keyboard Shortcut like `Ctrl+Shift+L` or `Cmd+Shift+L` (see `Tools` > `Modify Keyboard Shortcuts...`).
#' @source Idea from the [`like` function from the `data.table` package](https://github.com/Rdatatable/data.table/blob/master/R/like.R)
#' @seealso [grep()]
#' @inheritSection AMR Read more on our website!
#' @examples
#' # simple test
#' a <- "This is a test"
#' b <- "TEST"
#' a %like% b
#' #> TRUE
#' b %like% a
#' #> FALSE
#'
#' # also supports multiple patterns, length must be equal to x
#' a <- c("Test case", "Something different", "Yet another thing")
#' b <- c(     "case",           "diff",      "yet")
#' a %like% b
#' #> TRUE TRUE TRUE
#'
#' # get isolates whose name start with 'Ent' or 'ent'
#' \donttest{
#' if (require("dplyr")) {
#'   example_isolates %>%
#'     filter(mo_name(mo) %like% "^ent")
#' }
#' }
like <- function(x, pattern, ignore.case = TRUE) {
  meet_criteria(x, allow_NA = TRUE)
  meet_criteria(pattern, allow_NA = FALSE)
  meet_criteria(ignore.case, allow_class = "logical", has_length = 1)

  # set to fixed if no regex found
  fixed <- !any(is_possibly_regex(pattern))
  if (ignore.case == TRUE) {
    # set here, otherwise if fixed = TRUE, this warning will be thrown: argument `ignore.case = TRUE` will be ignored
    x <- tolower(x)
    pattern <- tolower(pattern)
  }
  
  if (length(pattern) > 1 & length(x) == 1) {
    x <- rep(x, length(pattern))
  }
  
  if (all(is.na(x))) {
    return(rep(FALSE, length(x)))
  }
    
  if (length(pattern) > 1) {
    res <- vector(length = length(pattern))
    if (length(x) != length(pattern)) {
      if (length(x) == 1) {
        x <- rep(x, length(pattern))
      }
      # return TRUE for every 'x' that matches any 'pattern', FALSE otherwise
      for (i in seq_len(length(res))) {
        if (is.factor(x[i])) {
          res[i] <- as.integer(x[i]) %in% grep(pattern[i], levels(x[i]), ignore.case = FALSE, fixed = fixed)
        } else {
          res[i] <- grepl(pattern[i], x[i], ignore.case = FALSE, fixed = fixed)
        }
      }
      res <- vapply(FUN.VALUE = logical(1), pattern, function(pttrn) grepl(pttrn, x, ignore.case = FALSE, fixed = fixed))
      res2 <- as.logical(rowSums(res))
      # get only first item of every hit in pattern
      res2[duplicated(res)] <- FALSE
      res2[rowSums(res) == 0] <- NA
      return(res2)
    } else {
      # x and pattern are of same length, so items with each other
      for (i in seq_len(length(res))) {
        if (is.factor(x[i])) {
          res[i] <- as.integer(x[i]) %in% grep(pattern[i], levels(x[i]), ignore.case = FALSE, fixed = fixed)
        } else {
          res[i] <- grepl(pattern[i], x[i], ignore.case = FALSE, fixed = fixed)
        }
      }
      return(res)
    }
  }
  
  # the regular way how grepl works; just one pattern against one or more x
  if (is.factor(x)) {
    as.integer(x) %in% grep(pattern, levels(x), ignore.case = FALSE, fixed = fixed)
  } else {
    tryCatch(grepl(pattern, x, ignore.case = FALSE, fixed = fixed),
             error = function(e) {
               if (grepl("invalid reg(ular )?exp", e$message, ignore.case = TRUE)) {
                 # try with perl = TRUE:
                 return(grepl(pattern = pattern, 
                                    x = x,
                                    ignore.case = FALSE, 
                                    fixed = fixed,
                                    perl = TRUE))
               } else {
                 # stop otherwise
                 stop(e$message)
               }
             })
  }
}

#' @rdname like
#' @export
"%like%" <- function(x, pattern) {
  meet_criteria(x, allow_NA = TRUE)
  meet_criteria(pattern, allow_NA = FALSE)
  like(x, pattern, ignore.case = TRUE)
}

#' @rdname like
#' @export
"%like_case%" <- function(x, pattern) {
  meet_criteria(x, allow_NA = TRUE)
  meet_criteria(pattern, allow_NA = FALSE)
  like(x, pattern, ignore.case = FALSE)
}

"%like_perl%" <- function(x, pattern) {
  meet_criteria(x, allow_NA = TRUE)
  meet_criteria(pattern, allow_NA = FALSE)
  # convenient for e.g. matching all Klebsiella and Raoultella, but not 
  # K. aerogenes: fullname %like_perl% "^(Klebsiella(?! aerogenes)|Raoultella)"
  grepl(x = tolower(x),
        pattern = tolower(pattern),
        perl = TRUE,
        fixed = FALSE,
        ignore.case = TRUE)
}
```
