## Supplementary material for "AMR - An R Package for Working with Antimicrobial Resistance Data": Source package: mdro.R

```
# ==================================================================== #
# TITLE                                                                #
# Antimicrobial Resistance (AMR) Analysis for R                        #
#                                                                      #
# SOURCE                                                               #
# https://github.com/msberends/AMR                                     #
#                                                                      #
# LICENCE                                                              #
# (c) 2018-2021 Berends MS, Luz CF et al.                              #
# Developed at the University of Groningen, the Netherlands, in        #
# collaboration with non-profit organisations Certe Medical            #
# Diagnostics & Advice, and University Medical Center Groningen.       # 
#                                                                      #
# This R package is free software; you can freely use and distribute   #
# it for both personal and commercial purposes under the terms of the  #
# GNU General Public License version 2.0 (GNU GPL-2), as published by  #
# the Free Software Foundation.                                        #
# We created this package for both routine data analysis and academic  #
# research and it was publicly released in the hope that it will be    #
# useful, but it comes WITHOUT ANY WARRANTY OR LIABILITY.              #
#                                                                      #
# Visit our website for the full manual and a complete tutorial about  #
# how to conduct AMR analysis: https://msberends.github.io/AMR/        #
# ==================================================================== #

#' Determine multidrug-resistant organisms (MDRO)
#'
#' Determine which isolates are multidrug-resistant organisms (MDRO) according to international and national guidelines.
#' @inheritSection lifecycle Stable lifecycle
#' @param x a [data.frame] with antibiotics columns, like `AMX` or `amox`. Can be left blank when used inside `dplyr` verbs, such as [`filter()`][dplyr::filter()], [`mutate()`][dplyr::mutate()] and [`summarise()`][dplyr::summarise()].
#' @param guideline a specific guideline to follow. When left empty, the publication by Magiorakos *et al.* (2012, Clinical Microbiology and Infection) will be followed, please see *Details*.
#' @inheritParams eucast_rules
#' @param pct_required_classes minimal required percentage of antimicrobial classes that must be available per isolate, rounded down. For example, with the default guideline, 17 antimicrobial classes must be available for *S. aureus*. Setting this `pct_required_classes` argument to `0.5` (default) means that for every *S. aureus* isolate at least 8 different classes must be available. Any lower number of available classes will return `NA` for that isolate.
#' @param combine_SI a [logical] to indicate whether all values of S and I must be merged into one, so resistance is only considered when isolates are R, not I. As this is the default behaviour of the [mdro()] function, it follows the redefinition by EUCAST about the interpretation of I (increased exposure) in 2019, see section 'Interpretation of S, I and R' below. When using `combine_SI = FALSE`, resistance is considered when isolates are R or I.
#' @param verbose a logical to turn Verbose mode on and off (default is off). In Verbose mode, the function does not return the MDRO results, but instead returns a data set in logbook form with extensive info about which isolates would be MDRO-positive, or why they are not.
#' @inheritSection eucast_rules Antibiotics
#' @details 
#' These functions are context-aware when used inside `dplyr` verbs, such as `filter()`, `mutate()` and `summarise()`. This means that then the `x` argument can be left blank, please see *Examples*.
#' 
#' For the `pct_required_classes` argument, values above 1 will be divided by 100. This is to support both fractions (`0.75` or `3/4`) and percentages (`75`).
#' 
#' Currently supported guidelines are (case-insensitive):
#' 
#' * `guideline = "CMI2012"` (default)
#'
#'   Magiorakos AP, Srinivasan A *et al.* "Multidrug-resistant, extensively drug-resistant and pandrug-resistant bacteria: an international expert proposal for interim standard definitions for acquired resistance." Clinical Microbiology and Infection (2012) ([link](https://www.clinicalmicrobiologyandinfection.com/article/S1198-743X(14)61632-3/fulltext))
#' 
#' * `guideline = "EUCAST3.2"` (or simply `guideline = "EUCAST"`)
#'
#'   The European international guideline - EUCAST Expert Rules Version 3.2 "Intrinsic Resistance and Unusual Phenotypes" ([link](https://www.eucast.org/fileadmin/src/media/PDFs/EUCAST_files/Expert_Rules/2020/Intrinsic_Resistance_and_Unusual_Phenotypes_Tables_v3.2_20200225.pdf))
#' 
#' * `guideline = "EUCAST3.1"`
#'
#'   The European international guideline - EUCAST Expert Rules Version 3.1 "Intrinsic Resistance and Exceptional Phenotypes Tables" ([link](https://www.eucast.org/fileadmin/src/media/PDFs/EUCAST_files/Expert_Rules/Expert_rules_intrinsic_exceptional_V3.1.pdf))
#' 
#' * `guideline = "TB"`
#'
#'   The international guideline for multi-drug resistant tuberculosis - World Health Organization "Companion handbook to the WHO guidelines for the programmatic management of drug-resistant tuberculosis" ([link](https://www.who.int/tb/publications/pmdt_companionhandbook/en/))
#' 
#' * `guideline = "MRGN"`
#'
#'   The German national guideline - Mueller et al. (2015) Antimicrobial Resistance and Infection Control 4:7; \doi{10.1186/s13756-015-0047-6}
#' 
#' * `guideline = "BRMO"`
#'
#'   The Dutch national guideline - Rijksinstituut voor Volksgezondheid en Milieu "WIP-richtlijn BRMO (Bijzonder Resistente Micro-Organismen) (ZKH)" ([link](https://www.rivm.nl/wip-richtlijn-brmo-bijzonder-resistente-micro-organismen-zkh))
#' 
#' Please suggest your own (country-specific) guidelines by letting us know: <https://github.com/msberends/AMR/issues/new>.
#' 
#' **Note:** Every test that involves the Enterobacteriaceae family, will internally be performed using its newly named *order* Enterobacterales, since the Enterobacteriaceae family has been taxonomically reclassified by Adeolu *et al.* in 2016. Before that, Enterobacteriaceae was the only family under the Enterobacteriales (with an i) order. All species under the old Enterobacteriaceae family are still under the new Enterobacterales (without an i) order, but divided into multiple families. The way tests are performed now by this [mdro()] function makes sure that results from before 2016 and after 2016 are identical.
#' @inheritSection as.rsi Interpretation of R and S/I
#' @return
#' - CMI 2012 paper - function [mdr_cmi2012()] or [mdro()]:\cr
#'   Ordered [factor] with levels `Negative` < `Multi-drug-resistant (MDR)` < `Extensively drug-resistant (XDR)` < `Pandrug-resistant (PDR)`
#' - TB guideline - function [mdr_tb()] or [`mdro(..., guideline = "TB")`][mdro()]:\cr
#'   Ordered [factor] with levels `Negative` < `Mono-resistant` < `Poly-resistant` < `Multi-drug-resistant` < `Extensively drug-resistant`
#' - German guideline - function [mrgn()] or [`mdro(..., guideline = "MRGN")`][mdro()]:\cr
#'   Ordered [factor] with levels `Negative` < `3MRGN` < `4MRGN`
#' - Everything else:\cr
#'   Ordered [factor] with levels `Negative` < `Positive, unconfirmed` < `Positive`. The value `"Positive, unconfirmed"` means that, according to the guideline, it is not entirely sure if the isolate is multi-drug resistant and this should be confirmed with additional (e.g. molecular) tests
#' @rdname mdro
#' @aliases MDR XDR PDR BRMO 3MRGN 4MRGN
#' @export
#' @inheritSection AMR Read more on our website!
#' @source
#' Please see *Details* for the list of publications used for this function.
#' @examples
#' mdro(example_isolates, guideline = "EUCAST")
#' 
#' \donttest{
#' if (require("dplyr")) {
#'   example_isolates %>%
#'     mdro() %>%
#'     table()
#'   
#'   # no need to define `x` when used inside dplyr verbs:
#'   example_isolates %>%
#'     mutate(MDRO = mdro(),
#'            EUCAST = eucast_exceptional_phenotypes(),
#'            BRMO = brmo(),
#'            MRGN = mrgn())
#' }
#' }
mdro <- function(x,
                 guideline = "CMI2012",
                 col_mo = NULL,
                 info = interactive(),
                 pct_required_classes = 0.5,
                 combine_SI = TRUE,
                 verbose = FALSE,
                 ...) {
  if (missing(x)) {
    x <- get_current_data(arg_name = "x", call = -2)
  }
  meet_criteria(x, allow_class = "data.frame")
  meet_criteria(guideline, allow_class = "character", has_length = 1, allow_NULL = TRUE)
  meet_criteria(col_mo, allow_class = "character", has_length = 1, is_in = colnames(x), allow_NULL = TRUE)
  meet_criteria(info, allow_class = "logical", has_length = 1)
  meet_criteria(pct_required_classes, allow_class = "numeric", has_length = 1)
  meet_criteria(combine_SI, allow_class = "logical", has_length = 1)
  meet_criteria(verbose, allow_class = "logical", has_length = 1)
  
  check_dataset_integrity()
  
  if (interactive() & verbose == TRUE & info == TRUE) {
    txt <- paste0("WARNING: In Verbose mode, the mdro() function does not return the MDRO results, but instead returns a data set in logbook form with extensive info about which isolates would be MDRO-positive, or why they are not.",
                  "\n\nThis may overwrite your existing data if you use e.g.:",
                  "\ndata <- mdro(data, verbose = TRUE)\n\nDo you want to continue?")
    showQuestion <- import_fn("showQuestion", "rstudioapi", error_on_fail = FALSE)
    if (!is.null(showQuestion)) {
      q_continue <- showQuestion("Using verbose = TRUE with mdro()", txt)
    } else {
      q_continue <- utils::menu(choices = c("OK", "Cancel"), graphics = FALSE, title = txt)
    }
    if (q_continue %in% c(FALSE, 2)) {
      message_("Cancelled, returning original data", add_fn = font_red, as_note = FALSE)
      return(x)
    }
  }
  
  stop_ifnot(is.data.frame(x), "`x` must be a data.frame")
  stop_if(any(dim(x) == 0), "`x` must contain rows and columns")
  
  # force regular data.frame, not a tibble or data.table
  x <- as.data.frame(x, stringsAsFactors = FALSE)
  
  stop_ifnot(is.numeric(pct_required_classes), "`pct_required_classes` must be numeric")
  if (pct_required_classes > 1) {
    # allow pct_required_classes = 75 -> pct_required_classes = 0.75
    pct_required_classes <- pct_required_classes / 100
  }
  
  if (!is.null(list(...)$country)) {
    warning_("Using `country` is deprecated, use `guideline` instead. Please see ?mdro.", call = FALSE)
    guideline <- list(...)$country
  }

    guideline.bak <- guideline
  guideline <- tolower(gsub("[^a-zA-Z0-9.]+", "", guideline))
  if (is.null(guideline)) {
    # default to the paper by Magiorakos et al. (2012)
    guideline <- "cmi2012"
  }
  if (guideline == "eucast") {
    # turn into latest EUCAST guideline
    guideline <- "eucast3.2"
  }
  if (guideline == "nl") {
    guideline <- "brmo"
  }
  if (guideline == "de") {
    guideline <- "mrgn"
  }
  stop_ifnot(guideline %in% c("brmo", "mrgn", "eucast3.1", "eucast3.2", "tb", "cmi2012"),
             "invalid guideline: ", guideline.bak)
  guideline <- list(code = guideline)
  
  # try to find columns based on type
  # -- mo
  if (is.null(col_mo)) {
    col_mo <- search_type_in_df(x = x, type = "mo", info = info)
  }
  if (is.null(col_mo) & guideline$code == "tb") {
    message_("No column found as input for `col_mo`, ",
             font_bold("assuming all records contain", font_italic("Mycobacterium tuberculosis.")))
             x$mo <- as.mo("Mycobacterium tuberculosis")
             col_mo <- "mo"
  }
  stop_if(is.null(col_mo), "`col_mo` must be set")
  stop_ifnot(col_mo %in% colnames(x), "column '", col_mo, "' (`col_mo`) not found")
  
  if (guideline$code == "cmi2012") {
    guideline$name <- "Multidrug-resistant, extensively drug-resistant and pandrug-resistant bacteria: an international expert proposal for interim standard definitions for acquired resistance."
    guideline$author <- "Magiorakos AP, Srinivasan A, Carey RB, ..., Vatopoulos A, Weber JT, Monnet DL"
    guideline$version <- NA
    guideline$source_url <- "Clinical Microbiology and Infection 18:3, 2012; doi: 10.1111/j.1469-0691.2011.03570.x"
    guideline$type <- "MDRs/XDRs/PDRs"
    
  } else if (guideline$code == "eucast3.1") {
    guideline$name <- "EUCAST Expert Rules, \"Intrinsic Resistance and Exceptional Phenotypes Tables\""
    guideline$author <- "EUCAST (European Committee on Antimicrobial Susceptibility Testing)"
    guideline$version <- "3.1, 2016"
    guideline$source_url <- "https://www.eucast.org/fileadmin/src/media/PDFs/EUCAST_files/Expert_Rules/Expert_rules_intrinsic_exceptional_V3.1.pdf"
    guideline$type <- "EUCAST Exceptional Phenotypes"
    
  } else if (guideline$code == "eucast3.2") {
    guideline$name <- "EUCAST Expert Rules, \"Intrinsic Resistance and Unusual Phenotypes\""
    guideline$author <- "EUCAST (European Committee on Antimicrobial Susceptibility Testing)"
    guideline$version <- "3.2, 2020"
    guideline$source_url <- "https://www.eucast.org/fileadmin/src/media/PDFs/EUCAST_files/Expert_Rules/2020/Intrinsic_Resistance_and_Unusual_Phenotypes_Tables_v3.2_20200225.pdf"
    guideline$type <- "EUCAST Unusual Phenotypes"
    
  } else if (guideline$code == "tb") {
    guideline$name <- "Companion handbook to the WHO guidelines for the programmatic management of drug-resistant tuberculosis"
    guideline$author <- "WHO (World Health Organization)"
    guideline$version <- "WHO/HTM/TB/2014.11, 2014"
    guideline$source_url <- "https://www.who.int/tb/publications/pmdt_companionhandbook/en/"
    guideline$type <- "MDR-TB's"
    
    # support per country:
  } else if (guideline$code == "mrgn") {
    guideline$name <- "Cross-border comparison of the Dutch and German guidelines on multidrug-resistant Gram-negative microorganisms"
    guideline$author <- "M\u00fcller J, Voss A, K\u00f6ck R, ..., Kern WV, Wendt C, Friedrich AW"
    guideline$version <- NA
    guideline$source_url <- "Antimicrobial Resistance and Infection Control 4:7, 2015; doi: 10.1186/s13756-015-0047-6"
    guideline$type <- "MRGNs"
    
  } else if (guideline$code == "brmo") {
    guideline$name <- "WIP-Richtlijn Bijzonder Resistente Micro-organismen (BRMO)"
    guideline$author <- "RIVM (Rijksinstituut voor de Volksgezondheid)"
    guideline$version <- "Revision as of December 2017"
    guideline$source_url <- "https://www.rivm.nl/Documenten_en_publicaties/Professioneel_Praktisch/Richtlijnen/Infectieziekten/WIP_Richtlijnen/WIP_Richtlijnen/Ziekenhuizen/WIP_richtlijn_BRMO_Bijzonder_Resistente_Micro_Organismen_ZKH"
    guideline$type <- "BRMOs"
  } else {
    stop("This guideline is currently unsupported: ", guideline$code, call. = FALSE)
  }
  
  if (guideline$code == "cmi2012") {
    cols_ab <- get_column_abx(x = x,
                              soft_dependencies = c(
                                # table 1 (S aureus):
                                "GEN",
                                "RIF",
                                "CPT",
                                "OXA",
                                "CIP",
                                "MFX",
                                "SXT",
                                "FUS",
                                "VAN",
                                "TEC",
                                "TLV",
                                "TGC",
                                "CLI",
                                "DAP",
                                "ERY",
                                "LNZ",
                                "CHL",
                                "FOS",
                                "QDA",
                                "TCY",
                                "DOX",
                                "MNO",
                                # table 2 (Enterococcus)
                                "GEH",
                                "STH",
                                "IPM",
                                "MEM",
                                "DOR",
                                "CIP",
                                "LVX",
                                "MFX",
                                "VAN",
                                "TEC",
                                "TGC",
                                "DAP",
                                "LNZ",
                                "AMP",
                                "QDA",
                                "DOX",
                                "MNO",
                                # table 3 (Enterobacteriaceae)
                                "GEN",
                                "TOB",
                                "AMK",
                                "NET",
                                "CPT",
                                "TCC",
                                "TZP",
                                "ETP",
                                "IPM",
                                "MEM",
                                "DOR",
                                "CZO",
                                "CXM",
                                "CTX",
                                "CAZ",
                                "FEP",
                                "FOX",
                                "CTT",
                                "CIP",
                                "SXT",
                                "TGC",
                                "ATM",
                                "AMP",
                                "AMC",
                                "SAM",
                                "CHL",
                                "FOS",
                                "COL",
                                "TCY",
                                "DOX",
                                "MNO",
                                # table 4 (Pseudomonas)
                                "GEN",
                                "TOB",
                                "AMK",
                                "NET",
                                "IPM",
                                "MEM",
                                "DOR",
                                "CAZ",
                                "FEP",
                                "CIP",
                                "LVX",
                                "TCC",
                                "TZP",
                                "ATM",
                                "FOS",
                                "COL",
                                "PLB",
                                # table 5 (Acinetobacter)
                                "GEN",
                                "TOB",
                                "AMK",
                                "NET",
                                "IPM",
                                "MEM",
                                "DOR",
                                "CIP",
                                "LVX",
                                "TZP",
                                "TCC",
                                "CTX",
                                "CRO",
                                "CAZ",
                                "FEP",
                                "SXT",
                                "SAM",
                                "COL",
                                "PLB",
                                "TCY",
                                "DOX",
                                "MNO"),
                              verbose = verbose,
                              info = info,
                              ...)
  } else if (guideline$code == "eucast3.2") {
    cols_ab <- get_column_abx(x = x,
                              soft_dependencies = c("AMP",
                                                    "AMX",
                                                    "CIP",
                                                    "DAL",
                                                    "DAP",
                                                    "ERV",
                                                    "FDX",
                                                    "GEN",
                                                    "LNZ",
                                                    "MEM",
                                                    "MTR",
                                                    "OMC",
                                                    "ORI",
                                                    "PEN",
                                                    "QDA",
                                                    "RIF",
                                                    "TEC",
                                                    "TGC",
                                                    "TLV",
                                                    "TOB",
                                                    "TZD",
                                                    "VAN"),
                              info = info,
                              verbose = verbose,
                              ...)
  } else if (guideline$code == "tb") {
    cols_ab <- get_column_abx(x = x,
                              soft_dependencies = c("CAP",
                                                    "ETH",
                                                    "GAT",
                                                    "INH",
                                                    "PZA",
                                                    "RIF",
                                                    "RIB",
                                                    "RFP"),
                              info = info,
                              verbose = verbose,
                              ...)
  } else if (guideline$code == "mrgn") {
    cols_ab <- get_column_abx(x = x,
                              soft_dependencies = c("PIP",
                                                    "CTX",
                                                    "CAZ",
                                                    "IPM",
                                                    "MEM",
                                                    "CIP"),
                              verbose = verbose,
                              info = info,
                              ...)
  } else {
    cols_ab <- get_column_abx(x = x,
                              verbose = verbose,
                              info = info,
                              ...)
  }
  
  # nolint start
  AMC <- cols_ab["AMC"]
  AMK <- cols_ab["AMK"]
  AMP <- cols_ab["AMP"]
  AMX <- cols_ab["AMX"]
  ATM <- cols_ab["ATM"]
  AZL <- cols_ab["AZL"]
  AZM <- cols_ab["AZM"]
  BPR <- cols_ab["BPR"]
  CAC <- cols_ab["CAC"]
  CAT <- cols_ab["CAT"]
  CAZ <- cols_ab["CAZ"]
  CCV <- cols_ab["CCV"]
  CDR <- cols_ab["CDR"]
  CDZ <- cols_ab["CDZ"]
  CEC <- cols_ab["CEC"]
  CED <- cols_ab["CED"]
  CEI <- cols_ab["CEI"]
  CEP <- cols_ab["CEP"]
  CFM <- cols_ab["CFM"]
  CFM1 <- cols_ab["CFM1"]
  CFP <- cols_ab["CFP"]
  CFR <- cols_ab["CFR"]
  CFS <- cols_ab["CFS"]
  CHL <- cols_ab["CHL"]
  CID <- cols_ab["CID"]
  CIP <- cols_ab["CIP"]
  CLI <- cols_ab["CLI"]
  CLR <- cols_ab["CLR"]
  CMX <- cols_ab["CMX"]
  CMZ <- cols_ab["CMZ"]
  CND <- cols_ab["CND"]
  COL <- cols_ab["COL"]
  CPD <- cols_ab["CPD"]
  CPM <- cols_ab["CPM"]
  CPO <- cols_ab["CPO"]
  CPR <- cols_ab["CPR"]
  CPT <- cols_ab["CPT"]
  CRD <- cols_ab["CRD"]
  CRO <- cols_ab["CRO"]
  CSL <- cols_ab["CSL"]
  CTB <- cols_ab["CTB"]
  CTF <- cols_ab["CTF"]
  CTL <- cols_ab["CTL"]
  CTT <- cols_ab["CTT"]
  CTX <- cols_ab["CTX"]
  CTZ <- cols_ab["CTZ"]
  CXM <- cols_ab["CXM"]
  CZD <- cols_ab["CZD"]
  CZO <- cols_ab["CZO"]
  CZX <- cols_ab["CZX"]
  DAL <- cols_ab["DAL"]
  DAP <- cols_ab["DAP"]
  DIT <- cols_ab["DIT"]
  DIZ <- cols_ab["DIZ"]
  DOR <- cols_ab["DOR"]
  DOX <- cols_ab["DOX"]
  ENX <- cols_ab["ENX"]
  ERV <- cols_ab["ERV"]
  ERY <- cols_ab["ERY"]
  ETP <- cols_ab["ETP"]
  FDX <- cols_ab["FDX"]
  FEP <- cols_ab["FEP"]
  FLC <- cols_ab["FLC"]
  FLE <- cols_ab["FLE"]
  FOS <- cols_ab["FOS"]
  FOX <- cols_ab["FOX"]
  FUS <- cols_ab["FUS"]
  GAT <- cols_ab["GAT"]
  GEH <- cols_ab["GEH"]
  GEM <- cols_ab["GEM"]
  GEN <- cols_ab["GEN"]
  GRX <- cols_ab["GRX"]
  HAP <- cols_ab["HAP"]
  IPM <- cols_ab["IPM"]
  KAN <- cols_ab["KAN"]
  LEX <- cols_ab["LEX"]
  LIN <- cols_ab["LIN"]
  LNZ <- cols_ab["LNZ"]
  LOM <- cols_ab["LOM"]
  LOR <- cols_ab["LOR"]
  LTM <- cols_ab["LTM"]
  LVX <- cols_ab["LVX"]
  MAN <- cols_ab["MAN"]
  MEM <- cols_ab["MEM"]
  MEV <- cols_ab["MEV"]
  MEZ <- cols_ab["MEZ"]
  MFX <- cols_ab["MFX"]
  MNO <- cols_ab["MNO"]
  MTR <- cols_ab["MTR"]
  NAL <- cols_ab["NAL"]
  NEO <- cols_ab["NEO"]
  NET <- cols_ab["NET"]
  NIT <- cols_ab["NIT"]
  NOR <- cols_ab["NOR"]
  NOV <- cols_ab["NOV"]
  OFX <- cols_ab["OFX"]
  OMC <- cols_ab["OMC"]
  ORI <- cols_ab["ORI"]
  OXA <- cols_ab["OXA"]
  PAZ <- cols_ab["PAZ"]
  PEF <- cols_ab["PEF"]
  PEN <- cols_ab["PEN"]
  PIP <- cols_ab["PIP"]
  PLB <- cols_ab["PLB"]
  PRI <- cols_ab["PRI"]
  PRU <- cols_ab["PRU"]
  QDA <- cols_ab["QDA"]
  RFL <- cols_ab["RFL"]
  RID <- cols_ab["RID"]
  RIF <- cols_ab["RIF"]
  RXT <- cols_ab["RXT"]
  SAM <- cols_ab["SAM"]
  SIS <- cols_ab["SIS"]
  SPT <- cols_ab["SPT"]
  SPX <- cols_ab["SPX"]
  STH <- cols_ab["STH"]
  SXT <- cols_ab["SXT"]
  TCC <- cols_ab["TCC"]
  TCY <- cols_ab["TCY"]
  TEC <- cols_ab["TEC"]
  TGC <- cols_ab["TGC"]
  TIC <- cols_ab["TIC"]
  TLV <- cols_ab["TLV"]
  TMP <- cols_ab["TMP"]
  TMX <- cols_ab["TMX"]
  TOB <- cols_ab["TOB"]
  TVA <- cols_ab["TVA"]
  TZD <- cols_ab["TZD"]
  TZP <- cols_ab["TZP"]
  VAN <- cols_ab["VAN"]
  # additional for TB
  CAP <- cols_ab["CAP"]
  ETH <- cols_ab["ETH"]
  GAT <- cols_ab["GAT"]
  INH <- cols_ab["INH"]
  PZA <- cols_ab["PZA"]
  RIF <- cols_ab["RIF"]
  RIB <- cols_ab["RIB"]
  RFP <- cols_ab["RFP"]
  abx_tb <- c(CAP, ETH, GAT, INH, PZA, RIF, RIB, RFP)
  abx_tb <- abx_tb[!is.na(abx_tb)]
  stop_if(guideline$code == "tb" & length(abx_tb) == 0, "no antimycobacterials found in data set")
  # nolint end
  
  if (combine_SI == TRUE) {
    search_result <- "R"
  } else {
    search_result <- c("R", "I")
  }
  
  if (info == TRUE) {
    if (combine_SI == TRUE) {
      cat(font_red("\nOnly results with 'R' are considered as resistance. Use `combine_SI = FALSE` to also consider 'I' as resistance.\n"))
    } else {
      cat(font_red("\nResults with 'R' or 'I' are considered as resistance. Use `combine_SI = TRUE` to only consider 'R' as resistance.\n"))
    }
    cat("\n", word_wrap("Determining multidrug-resistant organisms (MDRO), according to:"), "\n",
        word_wrap(paste0(font_bold("Guideline: "), font_italic(guideline$name)), extra_indent = 11, as_note = FALSE), "\n",
        word_wrap(paste0(font_bold("Author(s): "), guideline$author), extra_indent = 11, as_note = FALSE), "\n",
        ifelse(!is.na(guideline$version),
               paste0(word_wrap(paste0(font_bold("Version:   "), guideline$version), extra_indent = 11, as_note = FALSE), "\n"),
               ""),
        paste0(font_bold("Source:    "), guideline$source_url),
        "\n\n", sep = "")
  }
  
  ab_missing <- function(ab) {
    isTRUE(ab %in% c(NULL, NA)) | length(ab) == 0
  }
  ab_NA <- function(x) {
    x[!is.na(x)]
  }
  
  # antibiotic classes
  # nolint start
  aminoglycosides <- c(TOB, GEN)
  cephalosporins <- c(CDZ, CAC, CEC, CFR, RID, MAN, CTZ, CZD, CZO, CDR, DIT, FEP, CAT, CFM, CMX, CMZ, DIZ, CID, CFP, CSL, CND, CTX, CTT, CTF, FOX, CPM, CPO, CPD, CPR, CRD, CFS, CPT, CAZ, CCV, CTL, CTB, CZX, BPR, CFM1, CEI, CRO, CXM, LEX, CEP, HAP, CED, LTM, LOR)
  cephalosporins_1st <- c(CAC, CFR, RID, CTZ, CZD, CZO, CRD, CTL, LEX, CEP, HAP, CED)
  cephalosporins_2nd <- c(CEC, MAN, CMZ, CID, CND, CTT, CTF, FOX, CPR, CXM, LOR)
  cephalosporins_3rd <- c(CDZ, CDR, DIT, CAT, CFM, CMX, DIZ, CFP, CSL, CTX, CPM, CPD, CFS, CAZ, CCV, CTB, CZX, CRO, LTM)
  carbapenems <- c(DOR, ETP, IPM, MEM, MEV)
  fluoroquinolones <- c(CIP, ENX, FLE, GAT, GEM, GRX, LVX, LOM, MFX, NOR, OFX, PAZ, PEF, PRU, RFL, SPX, TMX, TVA)
  # nolint end
  
  # helper function for editing the table
  trans_tbl <- function(to, rows, cols, any_all) {
    cols <- cols[!ab_missing(cols)]
    cols <- cols[!is.na(cols)]
    if (length(rows) > 0 & length(cols) > 0) {
      x[, cols] <- as.data.frame(lapply(x[, cols, drop = FALSE],
                                        function(col) as.rsi(col)), 
                                 stringsAsFactors = FALSE)
      x[rows, "columns_nonsusceptible"] <<- vapply(FUN.VALUE = character(1),
                                                   rows, 
                                                   function(row, group_vct = cols) {
                                                     cols_nonsus <- vapply(FUN.VALUE = logical(1), x[row, group_vct, drop = FALSE], 
                                                                           function(y) y %in% search_result)
                                                     paste(sort(c(unlist(strsplit(x[row, "columns_nonsusceptible", drop = TRUE], ", ")),
                                                                  names(cols_nonsus)[cols_nonsus])), 
                                                           collapse = ", ")
                                                   })
      
      if (any_all == "any") {
        search_function <- any
      } else if (any_all == "all") {
        search_function <- all
      }
      x_transposed <- as.list(as.data.frame(t(x[, cols, drop = FALSE]),
                                            stringsAsFactors = FALSE))
      row_filter <- vapply(FUN.VALUE = logical(1), x_transposed, function(y) search_function(y %in% search_result, na.rm = TRUE))
      row_filter <- x[which(row_filter), "row_number", drop = TRUE]
      rows <- rows[rows %in% row_filter]
      x[rows, "MDRO"] <<- to
      x[rows, "reason"] <<- paste0(any_all, " of the required antibiotics ", ifelse(any_all == "any", "is", "are"), " R")
    }
  }
  trans_tbl2 <- function(txt, rows, lst) {
    if (info == TRUE) {
      message_(txt, "...", appendLF = FALSE, as_note = FALSE)
    }
    if (length(rows) > 0) {
      # function specific for the CMI paper of 2012 (Magiorakos et al.)
      lst_vector <- unlist(lst)[!is.na(unlist(lst))]
      x[, lst_vector] <- as.data.frame(lapply(x[, lst_vector, drop = FALSE],
                                              function(col) as.rsi(col)),
                                       stringsAsFactors = FALSE)
      x[rows, "classes_in_guideline"] <<- length(lst)
      x[rows, "classes_available"] <<- vapply(FUN.VALUE = double(1),
                                              rows, 
                                              function(row, group_tbl = lst) {
                                                sum(vapply(FUN.VALUE = logical(1),
                                                           group_tbl, 
                                                           function(group) any(unlist(x[row, group[!is.na(group)], drop = TRUE]) %in% c("S", "I", "R"))))
                                              })
      
      if (verbose == TRUE) {
        x[rows, "columns_nonsusceptible"] <<- vapply(FUN.VALUE = character(1),
                                                     rows, 
                                                     function(row, group_vct = lst_vector) {
                                                       cols_nonsus <- vapply(FUN.VALUE = logical(1), x[row, group_vct, drop = FALSE], function(y) y %in% search_result)
                                                       paste(sort(names(cols_nonsus)[cols_nonsus]), collapse = ", ")
                                                     })
      }
      x[rows, "classes_affected"] <<- vapply(FUN.VALUE = double(1),
                                             rows, 
                                             function(row, group_tbl = lst) {
                                               sum(vapply(FUN.VALUE = logical(1),
                                                          group_tbl, 
                                                          function(group) {
                                                            any(unlist(x[row, group[!is.na(group)], drop = TRUE]) %in% search_result, na.rm = TRUE)
                                                          }),
                                                   na.rm = TRUE) 
                                             })
      # for PDR; all agents are R (or I if combine_SI = FALSE)
      x_transposed <- as.list(as.data.frame(t(x[rows, lst_vector, drop = FALSE]),
                                            stringsAsFactors = FALSE))
      row_filter <- vapply(FUN.VALUE = logical(1), x_transposed, function(y) all(y %in% search_result, na.rm = TRUE))
      x[which(row_filter), "classes_affected"] <<- 999
    }
    
    if (info == TRUE) {
      message_(" OK.", add_fn = list(font_green, font_bold), as_note = FALSE)
    }
  }
  
  x[, col_mo] <- as.mo(as.character(x[, col_mo, drop = TRUE]))
  # join to microorganisms data set
  x <- left_join_microorganisms(x, by = col_mo)
  x$MDRO <- ifelse(!is.na(x$genus), 1, NA_integer_)
  x$row_number <- seq_len(nrow(x))
  x$reason <- paste0("not covered by ", toupper(guideline$code), " guideline")
  x$columns_nonsusceptible <- ""
  
  if (guideline$code == "cmi2012") {
    # CMI, 2012 ---------------------------------------------------------------
    # Non-susceptible = R and I
    # (see header 'Approaches to Creating Definitions for MDR, XDR and PDR' in paper)
    
    # take amoxicillin if ampicillin is unavailable
    if (is.na(AMP) & !is.na(AMX)) {
      if (verbose == TRUE) {
        message_("Filling ampicillin (AMP) results with amoxicillin (AMX) results")
      }
      AMP <- AMX
    }
    # take ceftriaxone if cefotaxime is unavailable and vice versa
    if (is.na(CRO) & !is.na(CTX)) {
      if (verbose == TRUE) {
        message_("Filling ceftriaxone (CRO) results with cefotaxime (CTX) results")
      }
      CRO <- CTX
    }
    if (is.na(CTX) & !is.na(CRO)) {
      if (verbose == TRUE) {
        message_("Filling cefotaxime (CTX) results with ceftriaxone (CRO) results")
      }
      CTX <- CRO
    }
    
    # intrinsic resistant must not be considered for the determination of MDR,
    # so let's just remove them, meticulously following the paper
    x[which(x$genus == "Enterococcus" & x$species == "faecium"), ab_NA(IPM)] <- NA
    x[which(x$genus == "Enterococcus" & x$species == "faecalis"), ab_NA(QDA)] <- NA
    x[which((x$genus == "Providencia" & x$species == "rettgeri")
            | (x$genus == "Providencia" & x$species == "stuartii")), ab_NA(c(GEN, TOB, NET))] <- NA
    x[which(x$genus == "Escherichia" & x$species == "hermannii"), ab_NA(c(TCC, TZP))] <- NA
    x[which((x$genus == "Citrobacter" & x$species == "freundii")
            | (x$genus == "Enterobacter" & x$species == "aerogenes")
            | (x$genus == "Enterobacter" & x$species == "cloacae")
            | (x$genus == "Hafnia" & x$species == "alvei")
            | (x$genus == "Morganella" & x$species == "morganii")
            | (x$genus == "Proteus" & x$species == "penneri")
            | (x$genus == "Proteus" & x$species == "vulgaris")
            | (x$genus == "Serratia" & x$species == "marcescens")), ab_NA(CZO)] <- NA
    x[which((x$genus == "Morganella" & x$species == "morganii")
            | (x$genus == "Proteus" & x$species == "penneri")
            | (x$genus == "Proteus" & x$species == "vulgaris")
            | (x$genus == "Serratia" & x$species == "marcescens")), ab_NA(CXM)] <- NA
    x[which((x$genus == "Morganella" & x$species == "morganii")
            | (x$genus == "Proteus" & x$species == "mirabilis")
            | (x$genus == "Proteus" & x$species == "penneri")
            | (x$genus == "Proteus" & x$species == "vulgaris")
            | (x$genus == "Providencia" & x$species == "rettgeri")
            | (x$genus == "Providencia" & x$species == "stuartii")), ab_NA(TGC)] <- NA
    x[which((x$genus == "Citrobacter" & x$species == "koseri")
            | (x$genus == "Citrobacter" & x$species == "freundii")
            | (x$genus == "Enterobacter" & x$species == "aerogenes")
            | (x$genus == "Enterobacter" & x$species == "cloacae")
            | (x$genus == "Escherichia" & x$species == "hermannii")
            | (x$genus == "Hafnia" & x$species == "alvei")
            | (x$genus == "Klebsiella")
            | (x$genus == "Morganella" & x$species == "morganii")
            | (x$genus == "Proteus" & x$species == "penneri")
            | (x$genus == "Proteus" & x$species == "vulgaris")
            | (x$genus == "Providencia" & x$species == "rettgeri")
            | (x$genus == "Providencia" & x$species == "stuartii")
            | (x$genus == "Serratia" & x$species == "marcescens")), ab_NA(AMP)] <- NA
    x[which((x$genus == "Citrobacter" & x$species == "freundii")
            | (x$genus == "Enterobacter" & x$species == "aerogenes")
            | (x$genus == "Enterobacter" & x$species == "cloacae")
            | (x$genus == "Hafnia" & x$species == "alvei")
            | (x$genus == "Morganella" & x$species == "morganii")
            | (x$genus == "Providencia" & x$species == "rettgeri")
            | (x$genus == "Providencia" & x$species == "stuartii")
            | (x$genus == "Serratia" & x$species == "marcescens")), ab_NA(AMC)] <- NA
    x[which((x$genus == "Citrobacter" & x$species == "freundii")
            | (x$genus == "Citrobacter" & x$species == "koseri")
            | (x$genus == "Enterobacter" & x$species == "aerogenes")
            | (x$genus == "Enterobacter" & x$species == "cloacae")
            | (x$genus == "Hafnia" & x$species == "alvei")
            | (x$genus == "Providencia" & x$species == "rettgeri")
            | (x$genus == "Serratia" & x$species == "marcescens")), ab_NA(SAM)] <- NA
    x[which((x$genus == "Morganella" & x$species == "morganii")
            | (x$genus == "Proteus" & x$species == "mirabilis")
            | (x$genus == "Proteus" & x$species == "penneri")
            | (x$genus == "Proteus" & x$species == "vulgaris")
            | (x$genus == "Providencia" & x$species == "rettgeri")
            | (x$genus == "Providencia" & x$species == "stuartii")
            | (x$genus == "Serratia" & x$species == "marcescens")), ab_NA(COL)] <- NA
    x[which((x$genus == "Morganella" & x$species == "morganii")
            | (x$genus == "Proteus" & x$species == "mirabilis")
            | (x$genus == "Proteus" & x$species == "penneri")
            | (x$genus == "Proteus" & x$species == "vulgaris")
            | (x$genus == "Providencia" & x$species == "rettgeri")
            | (x$genus == "Providencia" & x$species == "stuartii")), ab_NA(TCY)] <- NA
    x[which((x$genus == "Morganella" & x$species == "morganii")
            | (x$genus == "Proteus" & x$species == "penneri")
            | (x$genus == "Proteus" & x$species == "vulgaris")
            | (x$genus == "Providencia" & x$species == "rettgeri")
            | (x$genus == "Providencia" & x$species == "stuartii")), ab_NA(c(DOX, MNO))] <- NA
    
    x$classes_in_guideline <- NA_integer_
    x$classes_available <- NA_integer_
    x$classes_affected <- NA_integer_
    
    # now add the MDR levels to the data
    trans_tbl(2,
              which(x$genus == "Staphylococcus" & x$species == "aureus"),
              c(OXA, FOX),
              "any")
    trans_tbl2(paste("Table 1 -", font_italic("Staphylococcus aureus")),
               which(x$genus == "Staphylococcus" & x$species == "aureus"),
               list(GEN,
                    RIF,
                    CPT,
                    c(OXA, FOX),
                    c(CIP, MFX),
                    SXT,
                    FUS,
                    c(VAN, TEC, TLV),
                    TGC,
                    CLI,
                    DAP,
                    ERY,
                    LNZ,
                    CHL,
                    FOS,
                    QDA,
                    c(TCY, DOX, MNO)))
    trans_tbl2(paste("Table 2 -", font_italic("Enterococcus"), "spp."),
               which(x$genus == "Enterococcus"),
               list(GEH,
                    STH,
                    c(IPM, MEM, DOR),
                    c(CIP, LVX, MFX),
                    c(VAN, TEC),
                    TGC,
                    DAP,
                    LNZ,
                    AMP,
                    QDA,
                    c(DOX, MNO)))
    trans_tbl2(paste0("Table 3 - ", font_italic("Enterobacteriaceae")),
               # this new order was previously 'Enterobacteriales' and contained only the family 'Enterobacteriaceae':
               which(x$order == "Enterobacterales"),
               list(c(GEN, TOB, AMK, NET),
                    CPT,
                    c(TCC, TZP),
                    c(ETP, IPM, MEM, DOR),
                    CZO,
                    CXM,
                    c(CTX, CAZ, FEP),
                    c(FOX, CTT),
                    CIP,
                    SXT,
                    TGC,
                    ATM,
                    AMP,
                    c(AMC, SAM),
                    CHL,
                    FOS,
                    COL,
                    c(TCY, DOX, MNO)))
    trans_tbl2(paste("Table 4 -", font_italic("Pseudomonas aeruginosa")),
               which(x$genus == "Pseudomonas" & x$species == "aeruginosa"),
               list(c(GEN, TOB, AMK, NET),
                    c(IPM, MEM, DOR),
                    c(CAZ, FEP),
                    c(CIP, LVX),
                    c(TCC, TZP),
                    ATM,
                    FOS,
                    c(COL, PLB)))
    trans_tbl2(paste("Table 5 -", font_italic("Acinetobacter"), "spp."),
               which(x$genus == "Acinetobacter"),
               list(c(GEN, TOB, AMK, NET),
                    c(IPM, MEM, DOR),
                    c(CIP, LVX),
                    c(TZP, TCC),
                    c(CTX, CRO, CAZ, FEP),
                    SXT,
                    SAM,
                    c(COL, PLB),
                    c(TCY, DOX, MNO)))
    
    # now set MDROs: 
    # MDR (=2): >=3 classes affected
    x[which(x$classes_affected >= 3), "MDRO"] <- 2
    if (verbose == TRUE) {
      x[which(x$classes_affected >= 3), "reason"] <- paste0("at least 3 classes contain R or I: ", x$classes_affected[which(x$classes_affected >= 3)],
                                                            " out of ", x$classes_available[which(x$classes_affected >= 3)], " available classes")
    }
    
    # XDR (=3): all but <=2 classes affected
    x[which((x$classes_in_guideline - x$classes_affected) <= 2), "MDRO"] <- 3
    if (verbose == TRUE) {
      x[which(x$MDRO == 3), "reason"] <- paste0("less than 3 classes remain susceptible (", x$classes_in_guideline[which((x$classes_in_guideline - x$classes_affected) <= 2)] - x$classes_affected[which(x$MDRO == 3)],
                                                " out of ", x$classes_in_guideline[which(x$MDRO == 3)], " classes)")
    }
    
    # PDR (=4): all agents are R 
    x[which(x$classes_affected == 999 & x$classes_in_guideline == x$classes_available), "MDRO"] <- 4
    if (verbose == TRUE) {
      x[which(x$MDRO == 4), "reason"] <- paste("all antibiotics in all", x$classes_in_guideline[which(x$MDRO == 4)], "classes were tested R or I")
    }
    
    # not enough classes available
    x[which(x$MDRO %in% c(1, 3) & x$classes_available < floor(x$classes_in_guideline * pct_required_classes)), "MDRO"] <- -1
    if (verbose == TRUE) {
      x[which(x$MDRO == -1), "reason"] <- paste0("not enough classes available: ", x$classes_available[which(x$MDRO == -1)], 
                                                 " of required ", (floor(x$classes_in_guideline * pct_required_classes))[which(x$MDRO == -1)], 
                                                 " (~", percentage(pct_required_classes), " of ", x$classes_in_guideline[which(x$MDRO == -1)], ")")
    }
    
    # add antibiotic names of resistant ones to verbose output
    
  }
  
  if (guideline$code == "eucast3.1") {
    # EUCAST 3.1 --------------------------------------------------------------
    # Table 5
    trans_tbl(3,
              which(x$order == "Enterobacterales"
                    | (x$genus == "Pseudomonas" & x$species == "aeruginosa")
                    | x$genus == "Acinetobacter"),
              COL,
              "all")
    trans_tbl(3,
              which(x$genus == "Salmonella" & x$species == "Typhi"),
              c(carbapenems, fluoroquinolones),
              "any")
    trans_tbl(3,
              which(x$genus == "Haemophilus" & x$species == "influenzae"),
              c(cephalosporins_3rd, carbapenems, fluoroquinolones),
              "any")
    trans_tbl(3,
              which(x$genus == "Moraxella" & x$species == "catarrhalis"),
              c(cephalosporins_3rd, fluoroquinolones),
              "any")
    trans_tbl(3,
              which(x$genus == "Neisseria" & x$species == "meningitidis"),
              c(cephalosporins_3rd, fluoroquinolones),
              "any")
    trans_tbl(3,
              which(x$genus == "Neisseria" & x$species == "gonorrhoeae"),
              AZM,
              "any")
    # Table 6
    trans_tbl(3,
              which(x$fullname %like% "^(Coagulase-negative|Staphylococcus (aureus|epidermidis|hominis|haemolyticus|intermedius|pseudointermedius))"),
              c(VAN, TEC, DAP, LNZ, QDA, TGC),
              "any")
    trans_tbl(3,
              which(x$genus == "Corynebacterium"),
              c(VAN, TEC, DAP, LNZ, QDA, TGC),
              "any")
    trans_tbl(3,
              which(x$genus == "Streptococcus" & x$species == "pneumoniae"),
              c(carbapenems, VAN, TEC, DAP, LNZ, QDA, TGC, RIF),
              "any")
    trans_tbl(3, # Sr. groups A/B/C/G
              which(x$fullname %like% "^Streptococcus (group (A|B|C|G)|pyogenes|agalactiae|equisimilis|equi|zooepidemicus|dysgalactiae|anginosus)"),
              c(PEN, cephalosporins, VAN, TEC, DAP, LNZ, QDA, TGC),
              "any")
    trans_tbl(3,
              which(x$genus == "Enterococcus"),
              c(DAP, LNZ, TGC, TEC),
              "any")
    trans_tbl(3,
              which(x$genus == "Enterococcus" & x$species == "faecalis"),
              c(AMP, AMX),
              "any")
    # Table 7
    trans_tbl(3,
              which(x$genus == "Bacteroides"),
              MTR,
              "any")
    trans_tbl(3,
              which(x$genus == "Clostridium" & x$species == "difficile"),
              c(MTR, VAN),
              "any")
  }
  
  if (guideline$code == "eucast3.2") {
    # EUCAST 3.2 --------------------------------------------------------------
    # Table 6
    trans_tbl(3,
              which((x$order == "Enterobacterales" &
                       !x$family == "Morganellaceae" & 
                       !(x$genus == "Serratia" & x$species == "marcescens"))
                    | (x$genus == "Pseudomonas" & x$species == "aeruginosa")
                    | x$genus == "Acinetobacter"),
              COL,
              "all")
    trans_tbl(3,
              which(x$genus == "Salmonella" & x$species == "Typhi"),
              c(carbapenems),
              "any")
    trans_tbl(3,
              which(x$genus == "Haemophilus" & x$species == "influenzae"),
              c(cephalosporins_3rd, carbapenems, fluoroquinolones),
              "any")
    trans_tbl(3,
              which(x$genus == "Moraxella" & x$species == "catarrhalis"),
              c(cephalosporins_3rd, fluoroquinolones),
              "any")
    trans_tbl(3,
              which(x$genus == "Neisseria" & x$species == "meningitidis"),
              c(cephalosporins_3rd, fluoroquinolones),
              "any")
    trans_tbl(3,
              which(x$genus == "Neisseria" & x$species == "gonorrhoeae"),
              SPT,
              "any")
    # Table 7
    trans_tbl(3,
              which(x$genus == "Staphylococcus" & x$species == "aureus"),
              c(VAN, TEC, TLV, DAL, ORI, DAP, LNZ, TZD, QDA, TGC, ERV, OMC),
              "any")
    trans_tbl(3,
              which(x$mo %in% MO_CONS), # coagulase-negative Staphylococcus
              c(VAN, TLV, DAL, ORI, DAP, LNZ, TZD, QDA, TGC, ERV, OMC),
              "any")
    trans_tbl(3,
              which(x$genus == "Corynebacterium"),
              c(VAN, TEC, TLV, DAL, ORI, DAP, LNZ, TZD, QDA, TGC),
              "any")
    trans_tbl(3,
              which(x$genus == "Streptococcus" & x$species == "pneumoniae"),
              c(carbapenems, VAN, TEC, TLV, DAL, ORI, DAP, LNZ, TZD, QDA, TGC, ERV, OMC, RIF),
              "any")
    streps <- MO_lookup[which(MO_lookup$genus == "Streptococcus"), "mo", drop = TRUE]
    streps_ABCG <- streps[as.mo(streps, Lancefield = TRUE) %in% c("B_STRPT_GRPA", "B_STRPT_GRPB", "B_STRPT_GRPC", "B_STRPT_GRPG")]
    trans_tbl(3, # Sr. groups A/B/C/G
              which(x$mo %in% streps_ABCG),
              c(PEN, cephalosporins, VAN, TEC, TLV, DAL, ORI, DAP, LNZ, TZD, QDA, TGC, ERV, OMC),
              "any")
    trans_tbl(3,
              which(x$genus == "Enterococcus"),
              c(DAP, LNZ, TGC, ERV, OMC, TEC),
              "any")
    trans_tbl(3,
              which(x$genus == "Enterococcus" & x$species == "faecalis"),
              c(AMP, AMX),
              "any")
    # Table 8
    trans_tbl(3,
              which(x$genus == "Bacteroides"),
              MTR,
              "any")
    trans_tbl(3,
              which(x$genus == "Clostridium" & x$species == "difficile"),
              c(MTR, VAN, FDX),
              "any")
  }
  
  if (guideline$code == "mrgn") {
    # Germany -----------------------------------------------------------------
    CTX_or_CAZ <- CTX %or% CAZ
    IPM_or_MEM <- IPM %or% MEM
    x$missing <- NA_character_
    if (is.na(PIP)) PIP <- "missing"
    if (is.na(CTX_or_CAZ)) CTX_or_CAZ <- "missing"
    if (is.na(IPM_or_MEM)) IPM_or_MEM <- "missing"
    if (is.na(IPM)) IPM <- "missing"
    if (is.na(MEM)) MEM <- "missing"
    if (is.na(CIP)) CIP <- "missing"
    
    # Table 1
    x[which((x$order == "Enterobacterales" |  # following in fact the old Enterobacteriaceae classification
               (x$genus == "Acinetobacter" & x$species ==  "baumannii")) &
              x[, PIP] == "R" &
              x[, CTX_or_CAZ] == "R" &
              x[, IPM_or_MEM] == "S" &
              x[, CIP] == "R"),
      "MDRO"] <- 2 # 2 = 3MRGN
    
    x[which((x$order == "Enterobacterales" |  # following in fact the old Enterobacteriaceae classification
               (x$genus == "Acinetobacter" & x$species ==  "baumannii")) &
              x[, PIP] == "R" &
              x[, CTX_or_CAZ] == "R" &
              x[, IPM_or_MEM] == "R" &
              x[, CIP] == "R"),
      "MDRO"] <- 3 # 3 = 4MRGN, overwrites 3MRGN if applicable
    
    x[which((x$order == "Enterobacterales" |  # following in fact the old Enterobacteriaceae classification
               (x$genus == "Acinetobacter" & x$species ==  "baumannii")) &
              x[, IPM] == "R" | x[, MEM] == "R"),
      "MDRO"] <- 3 # 3 = 4MRGN, always when imipenem or meropenem is R
    
    x[which(x$genus == "Pseudomonas" & x$species == "aeruginosa" &
              (x[, PIP] == "S") +
              (x[, CTX_or_CAZ] == "S") +
              (x[, IPM_or_MEM] == "S") +
              (x[, CIP] == "S") == 1),
      "MDRO"] <- 2 # 2 = 3MRGN, if only 1 group is S
    
    x[which((x$genus == "Pseudomonas" & x$species == "aeruginosa") &
              x[, PIP] == "R" &
              x[, CTX_or_CAZ] == "R" &
              x[, IPM_or_MEM] == "R" &
              x[, CIP] == "R"),
      "MDRO"] <- 3 # 3 = 4MRGN
  }
  
  if (guideline$code == "brmo") {
    # Netherlands -------------------------------------------------------------
    aminoglycosides <- aminoglycosides[!is.na(aminoglycosides)]
    fluoroquinolones <- fluoroquinolones[!is.na(fluoroquinolones)]
    carbapenems <- carbapenems[!is.na(carbapenems)]
    amino <- AMX %or% AMP
    third <- CAZ %or% CTX
    ESBLs <- c(amino, third)
    ESBLs <- ESBLs[!is.na(ESBLs)]
    if (length(ESBLs) != 2) {
      ESBLs <- character(0)
    }
    
    # Table 1
    trans_tbl(3,
              which(x$order == "Enterobacterales"), # following in fact the old Enterobacteriaceae classification
              c(aminoglycosides, fluoroquinolones),
              "all")
    
    trans_tbl(2,
              which(x$order == "Enterobacterales"), # following in fact the old Enterobacteriaceae classification
              carbapenems,
              "any")
    
    trans_tbl(2,
              which(x$order == "Enterobacterales"), # following in fact the old Enterobacteriaceae classification
              ESBLs,
              "all")
    
    # Table 2
    trans_tbl(2,
              which(x$genus == "Acinetobacter"),
              c(carbapenems),
              "any")
    trans_tbl(3,
              which(x$genus == "Acinetobacter"),
              c(aminoglycosides, fluoroquinolones),
              "all")
    
    trans_tbl(3,
              which(x$genus == "Stenotrophomonas" & x$species == "maltophilia"),
              SXT,
              "all")
    
    if (!ab_missing(MEM) & !ab_missing(IPM)
        & !ab_missing(GEN) & !ab_missing(TOB)
        & !ab_missing(CIP)
        & !ab_missing(CAZ)
        & !ab_missing(TZP)) {
      x$psae <- 0
      x[which(x[, MEM] == "R" | x[, IPM] == "R"), "psae"] <- 1 + x[which(x[, MEM] == "R" | x[, IPM] == "R"), "psae"]
      x[which(x[, GEN] == "R" & x[, TOB] == "R"), "psae"] <- 1 + x[which(x[, GEN] == "R" & x[, TOB] == "R"), "psae"]
      x[which(x[, CIP] == "R"), "psae"] <- 1 + x[which(x[, CIP] == "R"), "psae"]
      x[which(x[, CAZ] == "R"), "psae"] <- 1 + x[which(x[, CAZ] == "R"), "psae"]
      x[which(x[, TZP] == "R"), "psae"] <- 1 + x[which(x[, TZP] == "R"), "psae"]
    } else {
      x$psae <- 0
    }
    x[which(
      x$genus == "Pseudomonas" & x$species == "aeruginosa"
      & x$psae >= 3), "MDRO"] <- 3
    
    # Table 3
    trans_tbl(3,
              which(x$genus == "Streptococcus" & x$species == "pneumoniae"),
              PEN,
              "all")
    trans_tbl(3,
              which(x$genus == "Streptococcus" & x$species == "pneumoniae"),
              VAN,
              "all")
    trans_tbl(3,
              which(x$genus == "Enterococcus" & x$species == "faecium"),
              c(PEN, VAN),
              "all")
  }
  
  if (guideline$code == "tb") {
    # Tuberculosis ------------------------------------------------------------
    prepare_drug <- function(ab) {
      # returns vector values of drug
      # if `ab` is a column name, looks up the values in `x`
      if (length(ab) == 1 & is.character(ab)) {
        if (ab %in% colnames(x)) {
          ab <- x[, ab, drop = TRUE]
        }
      }
      ab <- as.character(as.rsi(ab))
      ab[is.na(ab)] <- ""
      ab
    }
    drug_is_R <- function(ab) {
      # returns logical vector
      ab <- prepare_drug(ab)
      if (length(ab) == 0) {
        rep(FALSE, NROW(x))
      } else if (length(ab) == 1) {
        rep(ab, NROW(x)) == "R"
      } else {
        ab == "R"
      }
    }
    drug_is_not_R <- function(ab) {
      # returns logical vector
      ab <- prepare_drug(ab)
      if (length(ab) == 0) {
        rep(TRUE, NROW(x))
      } else if (length(ab) == 1) {
        rep(ab, NROW(x)) != "R"
      } else {
        ab != "R"
      }
    }
    
    x$mono_count <- 0
    x[drug_is_R(INH), "mono_count"] <- x[drug_is_R(INH), "mono_count", drop = TRUE] + 1
    x[drug_is_R(RIF), "mono_count"] <- x[drug_is_R(RIF), "mono_count", drop = TRUE] + 1
    x[drug_is_R(ETH), "mono_count"] <- x[drug_is_R(ETH), "mono_count", drop = TRUE] + 1
    x[drug_is_R(PZA), "mono_count"] <- x[drug_is_R(PZA), "mono_count", drop = TRUE] + 1
    x[drug_is_R(RIB), "mono_count"] <- x[drug_is_R(RIB), "mono_count", drop = TRUE] + 1
    x[drug_is_R(RFP), "mono_count"] <- x[drug_is_R(RFP), "mono_count", drop = TRUE] + 1
    
    x$mono <- x$mono_count > 0
    x$poly <- x$mono_count > 1 & drug_is_not_R(RIF) & drug_is_not_R(INH)
    x$mdr <- drug_is_R(RIF) & drug_is_R(INH)
    x$xdr <- drug_is_R(LVX) | drug_is_R(MFX) | drug_is_R(GAT)
    x$second <- drug_is_R(CAP) | drug_is_R(KAN) | drug_is_R(AMK)
    x$xdr <- x$mdr & x$xdr & x$second
    x$MDRO <- ifelse(x$xdr, 5,
                     ifelse(x$mdr, 4,
                            ifelse(x$poly, 3,
                                   ifelse(x$mono, 2,
                                          1))))
    # keep all real TB, make other species NA
    x$MDRO <- ifelse(x$fullname == "Mycobacterium tuberculosis", x$MDRO, NA_real_)
  }
  
  if (info == TRUE) {
    if (sum(!is.na(x$MDRO) == 0)) {
      cat(font_bold(paste0("=> Found 0 MDROs since no isolates are covered by the guideline")))
    } else {
      cat(font_bold(paste0("=> Found ", sum(x$MDRO %in% c(2:5), na.rm = TRUE), " ", guideline$type, " out of ", sum(!is.na(x$MDRO)), 
                           " isolates (", trimws(percentage(sum(x$MDRO %in% c(2:5), na.rm = TRUE) / sum(!is.na(x$MDRO)))), ")\n")))
    }
  }
  
  # some more info on negative results
  if (verbose == TRUE) {
    if (guideline$code == "cmi2012") {
      x[which(x$MDRO == 1 & !is.na(x$classes_affected)), "reason"] <- paste0(x$classes_affected[which(x$MDRO == 1 & !is.na(x$classes_affected))], " of ", x$classes_available[which(x$MDRO == 1 & !is.na(x$classes_affected))], " available classes contain R or I (3 required for MDR)")
    } else {
      x[which(x$MDRO == 1), "reason"] <- "too few antibiotics are R"
    }
  }
  
  # Results ----
  if (guideline$code == "cmi2012") {
    if (any(x$MDRO == -1, na.rm = TRUE)) {
      warning_("NA introduced for isolates where the available percentage of antimicrobial classes was below ",
              percentage(pct_required_classes), " (set with `pct_required_classes`)", call = FALSE)
      # set these -1s to NA
      x[which(x$MDRO == -1), "MDRO"] <- NA_integer_
    }
    x$MDRO <- factor(x = x$MDRO,
                     levels = 1:4,
                     labels = c("Negative", "Multi-drug-resistant (MDR)", 
                                "Extensively drug-resistant (XDR)", "Pandrug-resistant (PDR)"),
                     ordered = TRUE)
  } else if (guideline$code == "tb") {
    x$MDRO <- factor(x = x$MDRO,
                     levels = 1:5,
                     labels = c("Negative", "Mono-resistant", "Poly-resistant", 
                                "Multi-drug-resistant", "Extensively drug-resistant"),
                     ordered = TRUE)
  } else if (guideline$code == "mrgn") {
    x$MDRO <- factor(x = x$MDRO,
                     levels = 1:3,
                     labels = c("Negative", "3MRGN", "4MRGN"),
                     ordered = TRUE)
  } else {
    x$MDRO <- factor(x = x$MDRO,
                     levels = 1:3,
                     labels = c("Negative", "Positive, unconfirmed", "Positive"),
                     ordered = TRUE)
  }
  
  if (verbose == TRUE) {
    x[, c("row_number",
          col_mo,
          "MDRO",
          "reason",
          "columns_nonsusceptible")]
    #x
  } else {
    x$MDRO
  }
  
}

#' @rdname mdro
#' @export
brmo <- function(x, guideline = "BRMO", ...) {
  if (missing(x)) {
    x <- get_current_data(arg_name = "x", call = -2)
  }
  meet_criteria(x, allow_class = "data.frame")
  meet_criteria(guideline, allow_class = "character", has_length = 1)
  mdro(x, guideline = "BRMO", ...)
}

#' @rdname mdro
#' @export
mrgn <- function(x, guideline = "MRGN", ...) {
  if (missing(x)) {
    x <- get_current_data(arg_name = "x", call = -2)
  }
  meet_criteria(x, allow_class = "data.frame")
  meet_criteria(guideline, allow_class = "character", has_length = 1)
  mdro(x = x, guideline = "MRGN", ...)
}

#' @rdname mdro
#' @export
mdr_tb <- function(x, guideline = "TB", ...) {
  if (missing(x)) {
    x <- get_current_data(arg_name = "x", call = -2)
  }
  meet_criteria(x, allow_class = "data.frame")
  meet_criteria(guideline, allow_class = "character", has_length = 1)
  mdro(x = x, guideline = "TB", ...)
}

#' @rdname mdro
#' @export
mdr_cmi2012 <- function(x, guideline = "CMI2012", ...) {
  if (missing(x)) {
    x <- get_current_data(arg_name = "x", call = -2)
  }
  meet_criteria(x, allow_class = "data.frame")
  meet_criteria(guideline, allow_class = "character", has_length = 1)
  mdro(x = x, guideline = "CMI2012", ...)
}

#' @rdname mdro
#' @export
eucast_exceptional_phenotypes <- function(x, guideline = "EUCAST", ...) {
  if (missing(x)) {
    x <- get_current_data(arg_name = "x", call = -2)
  }
  meet_criteria(x, allow_class = "data.frame")
  meet_criteria(guideline, allow_class = "character", has_length = 1)
  mdro(x = x, guideline = "EUCAST", ...)
}
```
