## Supplementary material for "AMR - An R Package for Working with Antimicrobial Resistance Data": Source package: mic.R

```
# ==================================================================== #
# TITLE                                                                #
# Antimicrobial Resistance (AMR) Analysis for R                        #
#                                                                      #
# SOURCE                                                               #
# https://github.com/msberends/AMR                                     #
#                                                                      #
# LICENCE                                                              #
# (c) 2018-2021 Berends MS, Luz CF et al.                              #
# Developed at the University of Groningen, the Netherlands, in        #
# collaboration with non-profit organisations Certe Medical            #
# Diagnostics & Advice, and University Medical Center Groningen.       # 
#                                                                      #
# This R package is free software; you can freely use and distribute   #
# it for both personal and commercial purposes under the terms of the  #
# GNU General Public License version 2.0 (GNU GPL-2), as published by  #
# the Free Software Foundation.                                        #
# We created this package for both routine data analysis and academic  #
# research and it was publicly released in the hope that it will be    #
# useful, but it comes WITHOUT ANY WARRANTY OR LIABILITY.              #
#                                                                      #
# Visit our website for the full manual and a complete tutorial about  #
# how to conduct AMR analysis: https://msberends.github.io/AMR/        #
# ==================================================================== #

#' Transform input to minimum inhibitory concentrations (MIC)
#'
#' This transforms a vector to a new class [`mic`], which is an ordered [factor] with valid minimum inhibitory concentrations (MIC) as levels. Invalid MIC values will be translated as `NA` with a warning.
#' @inheritSection lifecycle Stable lifecycle
#' @rdname as.mic
#' @param x vector
#' @param na.rm a logical indicating whether missing values should be removed
#' @details To interpret MIC values as RSI values, use [as.rsi()] on MIC values. It supports guidelines from EUCAST and CLSI.
#' @return Ordered [factor] with additional class [`mic`]
#' @aliases mic
#' @export
#' @seealso [as.rsi()]
#' @inheritSection AMR Read more on our website!
#' @examples
#' mic_data <- as.mic(c(">=32", "1.0", "1", "1.00", 8, "<=0.128", "8", "16", "16"))
#' is.mic(mic_data)
#'
#' # this can also coerce combined MIC/RSI values:
#' as.mic("<=0.002; S") # will return <=0.002
#'
#' # interpret MIC values
#' as.rsi(x = as.mic(2),
#'        mo = as.mo("S. pneumoniae"),
#'        ab = "AMX",
#'        guideline = "EUCAST")
#' as.rsi(x = as.mic(4),
#'        mo = as.mo("S. pneumoniae"),
#'        ab = "AMX",
#'        guideline = "EUCAST")
#'
#' plot(mic_data)
#' barplot(mic_data)
as.mic <- function(x, na.rm = FALSE) {
  meet_criteria(x, allow_class = c("mic", "character", "numeric", "integer"), allow_NA = TRUE)
  meet_criteria(na.rm, allow_class = "logical", has_length = 1)
  
  if (is.mic(x)) {
    x
  } else {
    x <- x %pm>% unlist()
    if (na.rm == TRUE) {
      x <- x[!is.na(x)]
    }
    x.bak <- x
    
    # comma to period
    x <- gsub(",", ".", x, fixed = TRUE)
    # transform Unicode for >= and <=
    x <- gsub("\u2264", "<=", x, fixed = TRUE)
    x <- gsub("\u2265", ">=", x, fixed = TRUE)
    # remove space between operator and number ("<= 0.002" -> "<=0.002")
    x <- gsub("(<|=|>) +", "\\1", x)
    # transform => to >= and =< to <=
    x <- gsub("=<", "<=", x, fixed = TRUE)
    x <- gsub("=>", ">=", x, fixed = TRUE)
    # dots without a leading zero must start with 0
    x <- gsub("([^0-9]|^)[.]", "\\10.", x)
    # values like "<=0.2560.512" should be 0.512
    x <- gsub(".*[.].*[.]", "0.", x)
    # remove ending .0
    x <- gsub("[.]+0$", "", x)
    # remove all after last digit
    x <- gsub("[^0-9]+$", "", x)
    # keep only one zero before dot
    x <- gsub("0+[.]", "0.", x)
    # starting 00 is probably 0.0 if there's no dot yet
    x[!x %like% "[.]"] <- gsub("^00", "0.0", x[!x %like% "[.]"])
    # remove last zeroes
    x <- gsub("([.].?)0+$", "\\1", x)
    x <- gsub("(.*[.])0+$", "\\10", x)
    # remove ending .0 again
    x[x %like% "[.]"] <- gsub("0+$", "", x[x %like% "[.]"])
    # never end with dot
    x <- gsub("[.]$", "", x)
    # force to be character
    x <- as.character(x)
    # trim it
    x <- trimws(x)
    
    ## previously unempty values now empty - should return a warning later on
    x[x.bak != "" & x == ""] <- "invalid"
    
    # these are allowed MIC values and will become factor levels
    ops <- c("<", "<=", "", ">=", ">")
    lvls <- c(c(t(vapply(FUN.VALUE = character(9), ops, function(x) paste0(x, "0.00", 1:9)))),
              unique(c(t(vapply(FUN.VALUE = character(104), ops, function(x) paste0(x, sort(as.double(paste0("0.0", 
                                                                                 sort(c(1:99, 125, 128, 256, 512, 625)))))))))),
              unique(c(t(vapply(FUN.VALUE = character(103), ops, function(x) paste0(x, sort(as.double(paste0("0.", 
                                                                                 c(1:99, 125, 128, 256, 512))))))))),
              c(t(vapply(FUN.VALUE = character(10), ops, function(x) paste0(x, sort(c(1:9, 1.5)))))),
              c(t(vapply(FUN.VALUE = character(45), ops, function(x) paste0(x, c(10:98)[9:98 %% 2 == TRUE])))),
              c(t(vapply(FUN.VALUE = character(15), ops, function(x) paste0(x, sort(c(2 ^ c(7:10), 80 * c(2:12))))))))
    
    na_before <- x[is.na(x) | x == ""] %pm>% length()
    x[!x %in% lvls] <- NA
    na_after <- x[is.na(x) | x == ""] %pm>% length()
    
    if (na_before != na_after) {
      list_missing <- x.bak[is.na(x) & !is.na(x.bak) & x.bak != ""] %pm>%
        unique() %pm>%
        sort()
      list_missing <- paste0('"', list_missing, '"', collapse = ", ")
      warning_(na_after - na_before, " results truncated (",
               round(((na_after - na_before) / length(x)) * 100),
               "%) that were invalid MICs: ",
               list_missing, call = FALSE)
    }
    
    set_clean_class(factor(x, levels = lvls, ordered = TRUE),
                    new_class =  c("mic", "ordered", "factor"))
  }
}

all_valid_mics <- function(x) {
  if (!inherits(x, c("mic", "character", "factor", "numeric", "integer"))) {
    return(FALSE)
  }
  x_mic <- tryCatch(suppressWarnings(as.mic(x[!is.na(x)])),
                    error = function(e) NA)
  !any(is.na(x_mic)) && !all(is.na(x))
}

#' @rdname as.mic
#' @export
is.mic <- function(x) {
  inherits(x, "mic")
}

#' @method as.double mic
#' @export
#' @noRd
as.double.mic <- function(x, ...) {
  as.double(gsub("[<=>]+", "", as.character(x)))
}

#' @method as.integer mic
#' @export
#' @noRd
as.integer.mic <- function(x, ...) {
  as.integer(gsub("[<=>]+", "", as.character(x)))
}

#' @method as.numeric mic
#' @export
#' @noRd
as.numeric.mic <- function(x, ...) {
  as.numeric(gsub("[<=>]+", "", as.character(x)))
}

#' @method droplevels mic
#' @export
#' @noRd
droplevels.mic <- function(x, exclude = if (any(is.na(levels(x)))) NULL else NA, ...) {
  x <- droplevels.factor(x, exclude = exclude, ...)
  class(x) <- c("mic", "ordered", "factor")
  x
}

# will be exported using s3_register() in R/zzz.R
pillar_shaft.mic <- function(x, ...) {
  crude_numbers <- as.double(x)
  operators <- gsub("[^<=>]+", "", as.character(x))
  pasted <- trimws(paste0(operators, trimws(format(crude_numbers))))
  out <- pasted
  out[is.na(x)] <- font_na(NA)
  out <- gsub("(<|=|>)", font_silver("\\1"), out)
  create_pillar_column(out, align = "right", width = max(nchar(pasted)))
}

# will be exported using s3_register() in R/zzz.R
type_sum.mic <- function(x, ...) {
  "mic"
}

#' @method print mic
#' @export
#' @noRd
print.mic <- function(x, ...) {
  cat("Class <mic>\n")
  print(as.character(x), quote = FALSE)
}

#' @method summary mic
#' @export
#' @noRd
summary.mic <- function(object, ...) {
  x <- object
  n_total <- length(x)
  x <- x[!is.na(x)]
  n <- length(x)
  value <- c("Class" = "mic",
             "<NA>" = n_total - n,
             "Min." = as.character(sort(x)[1]),
             "Max." = as.character(sort(x)[n]))
  class(value) <- c("summaryDefault", "table")
  value
}

#' @method plot mic
#' @export
#' @importFrom graphics barplot axis
#' @rdname plot
plot.mic <- function(x,
                     main = paste("MIC values of", deparse(substitute(x))),
                     ylab = "Frequency",
                     xlab = "MIC value",
                     axes = FALSE,
                     ...) {
  meet_criteria(main, allow_class = "character", has_length = 1)
  meet_criteria(ylab, allow_class = "character", has_length = 1)
  meet_criteria(xlab, allow_class = "character", has_length = 1)
  meet_criteria(axes, allow_class = "logical", has_length = 1)
  
  barplot(table(as.double(x)),
          ylab = ylab,
          xlab = xlab,
          axes = axes,
          main = main,
          ...)
  axis(2, seq(0, max(table(as.double(x)))))
}

#' @method barplot mic
#' @export
#' @importFrom graphics barplot axis
#' @rdname plot
barplot.mic <- function(height,
                        main = paste("MIC values of", deparse(substitute(height))),
                        ylab = "Frequency",
                        xlab = "MIC value",
                        axes = FALSE,
                        ...) {
  meet_criteria(main, allow_class = "character", has_length = 1)
  meet_criteria(ylab, allow_class = "character", has_length = 1)
  meet_criteria(xlab, allow_class = "character", has_length = 1)
  meet_criteria(axes, allow_class = "logical", has_length = 1)
  
  barplot(table(as.double(height)),
          ylab = ylab,
          xlab = xlab,
          axes = axes,
          main = main,
          ...)
  axis(2, seq(0, max(table(as.double(height)))))
}

#' @method [ mic
#' @export
#' @noRd
"[.mic" <- function(x, ...) {
  y <- NextMethod()
  attributes(y) <- attributes(x)
  y
}
#' @method [[ mic
#' @export
#' @noRd
"[[.mic" <- function(x, ...) {
  y <- NextMethod()
  attributes(y) <- attributes(x)
  y
}
#' @method [<- mic
#' @export
#' @noRd
"[<-.mic" <- function(i, j, ..., value) {
  value <- as.mic(value)
  y <- NextMethod()
  attributes(y) <- attributes(i)
  y
}
#' @method [[<- mic
#' @export
#' @noRd
"[[<-.mic" <- function(i, j, ..., value) {
  value <- as.mic(value)
  y <- NextMethod()
  attributes(y) <- attributes(i)
  y
}
#' @method c mic
#' @export
#' @noRd
c.mic <- function(x, ...) {
  y <- unlist(lapply(list(...), as.character))
  x <- as.character(x)
  as.mic(c(x, y))
}

#' @method unique mic
#' @export
#' @noRd
unique.mic <- function(x, incomparables = FALSE, ...) {
  y <- NextMethod()
  attributes(y) <- attributes(x)
  y
}

# will be exported using s3_register() in R/zzz.R
get_skimmers.mic <- function(column) {
  skimr::sfl(
    skim_type = "mic",
    min = ~as.character(sort(stats::na.omit(.))[1]),
    max = ~as.character(sort(stats::na.omit(.))[length(stats::na.omit(.))]),
    median = ~as.character(stats::na.omit(.)[as.double(stats::na.omit(.)) == median(as.double(stats::na.omit(.)))])[1],
    n_unique = ~pm_n_distinct(., na.rm = TRUE),
    hist_log2 = ~skimr::inline_hist(log2(as.double(stats::na.omit(.))))
  )
}
```
