## Supplementary material for "AMR - An R Package for Working with Antimicrobial Resistance Data": Source package: mo.R

````
# ==================================================================== #
# TITLE                                                                #
# Antimicrobial Resistance (AMR) Analysis for R                        #
#                                                                      #
# SOURCE                                                               #
# https://github.com/msberends/AMR                                     #
#                                                                      #
# LICENCE                                                              #
# (c) 2018-2021 Berends MS, Luz CF et al.                              #
# Developed at the University of Groningen, the Netherlands, in        #
# collaboration with non-profit organisations Certe Medical            #
# Diagnostics & Advice, and University Medical Center Groningen.       #
#                                                                      #
# This R package is free software; you can freely use and distribute   #
# it for both personal and commercial purposes under the terms of the  #
# GNU General Public License version 2.0 (GNU GPL-2), as published by  #
# the Free Software Foundation.                                        #
# We created this package for both routine data analysis and academic  #
# research and it was publicly released in the hope that it will be    #
# useful, but it comes WITHOUT ANY WARRANTY OR LIABILITY.              #
#                                                                      #
# Visit our website for the full manual and a complete tutorial about  #
# how to conduct AMR analysis: https://msberends.github.io/AMR/        #
# ==================================================================== #

#' Transform input to a microorganism ID
#'
#' Use this function to determine a valid microorganism ID ([`mo`]). Determination is done using intelligent rules and the complete taxonomic kingdoms Bacteria, Chromista, Protozoa, Archaea and most microbial species from the kingdom Fungi (see Source). The input can be almost anything: a full name (like `"Staphylococcus aureus"`), an abbreviated name (such as `"S. aureus"`), an abbreviation known in the field (such as `"MRSA"`), or just a genus. Please see *Examples*.
#' @inheritSection lifecycle Stable lifecycle
#' @param x a character vector or a [data.frame] with one or two columns
#' @param Becker a logical to indicate whether staphylococci should be categorised into coagulase-negative staphylococci ("CoNS") and coagulase-positive staphylococci ("CoPS") instead of their own species, according to Karsten Becker *et al.* (1,2,3).
#'
#' This excludes *Staphylococcus aureus* at default, use `Becker = "all"` to also categorise *S. aureus* as "CoPS".
#' @param Lancefield a logical to indicate whether beta-haemolytic *Streptococci* should be categorised into Lancefield groups instead of their own species, according to Rebecca C. Lancefield (4). These *Streptococci* will be categorised in their first group, e.g. *Streptococcus dysgalactiae* will be group C, although officially it was also categorised into groups G and L.
#'
#' This excludes *Enterococci* at default (who are in group D), use `Lancefield = "all"` to also categorise all *Enterococci* as group D.
#' @param allow_uncertain a number between `0` (or `"none"`) and `3` (or `"all"`), or `TRUE` (= `2`) or `FALSE` (= `0`) to indicate whether the input should be checked for less probable results, please see *Details*
#' @param reference_df a [data.frame] to be used for extra reference when translating `x` to a valid [`mo`]. See [set_mo_source()] and [get_mo_source()] to automate the usage of your own codes (e.g. used in your analysis or organisation).
#' @param ignore_pattern a regular expression (case-insensitive) of which all matches in `x` must return `NA`. This can be convenient to exclude known non-relevant input and can also be set with the option `AMR_ignore_pattern`, e.g. `options(AMR_ignore_pattern = "(not reported|contaminated flora)")`.
#' @param language language to translate text like "no growth", which defaults to the system language (see [get_locale()])
#' @param ... other arguments passed on to functions
#' @rdname as.mo
#' @aliases mo
#' @keywords mo Becker becker Lancefield lancefield guess
#' @details
#' ## General info
#'
#' A microorganism ID from this package (class: [`mo`]) is human readable and typically looks like these examples:
#' ```
#'   Code               Full name
#'   ---------------    --------------------------------------
#'   B_KLBSL            Klebsiella
#'   B_KLBSL_PNMN       Klebsiella pneumoniae
#'   B_KLBSL_PNMN_RHNS  Klebsiella pneumoniae rhinoscleromatis
#'   |   |    |    |
#'   |   |    |    |
#'   |   |    |    \---> subspecies, a 4-5 letter acronym
#'   |   |    \----> species, a 4-5 letter acronym
#'   |   \----> genus, a 5-7 letter acronym
#'   \----> taxonomic kingdom: A (Archaea), AN (Animalia), B (Bacteria),
#'                             C (Chromista), F (Fungi), P (Protozoa)
#' ```
#'
#' Values that cannot be coerced will be considered 'unknown' and will get the MO code `UNKNOWN`.
#'
#' Use the [`mo_*`][mo_property()] functions to get properties based on the returned code, see Examples.
#'
#' The algorithm uses data from the Catalogue of Life (see below) and from one other source (see [microorganisms]).
#'
#' The [as.mo()] function uses several coercion rules for fast and logical results. It assesses the input matching criteria in the following order:
#'
#' 1. Human pathogenic prevalence: the function  starts with more prevalent microorganisms, followed by less prevalent ones;
#' 2. Taxonomic kingdom: the function starts with determining Bacteria, then Fungi, then Protozoa, then others;
#' 3. Breakdown of input values to identify possible matches.
#'
#' This will lead to the effect that e.g. `"E. coli"` (a microorganism highly prevalent in humans) will return the microbial ID of *Escherichia coli* and not *Entamoeba coli* (a microorganism less prevalent in humans), although the latter would alphabetically come first.
#'
#' ## Coping with uncertain results
#'
#' In addition, the [as.mo()] function can differentiate four levels of uncertainty to guess valid results:
#' - Uncertainty level 0: no additional rules are applied;
#' - Uncertainty level 1: allow previously accepted (but now invalid) taxonomic names and minor spelling errors;
#' - Uncertainty level 2: allow all of level 1, strip values between brackets, inverse the words of the input, strip off text elements from the end keeping at least two elements;
#' - Uncertainty level 3: allow all of level 1 and 2, strip off text elements from the end, allow any part of a taxonomic name.
#'
#' The level of uncertainty can be set using the argument `allow_uncertain`. The default is `allow_uncertain = TRUE`, which is equal to uncertainty level 2. Using `allow_uncertain = FALSE` is equal to uncertainty level 0 and will skip all rules. You can also use e.g. `as.mo(..., allow_uncertain = 1)` to only allow up to level 1 uncertainty.
#'
#' With the default setting (`allow_uncertain = TRUE`, level 2), below examples will lead to valid results:
#' - `"Streptococcus group B (known as S. agalactiae)"`. The text between brackets will be removed and a warning will be thrown that the result *Streptococcus group B* (``r as.mo("Streptococcus group B")``) needs review.
#' - `"S. aureus - please mind: MRSA"`. The last word will be stripped, after which the function will try to find a match. If it does not, the second last word will be stripped, etc. Again, a warning will be thrown that the result *Staphylococcus aureus* (``r as.mo("Staphylococcus aureus")``) needs review.
#' - `"Fluoroquinolone-resistant Neisseria gonorrhoeae"`. The first word will be stripped, after which the function will try to find a match. A warning will be thrown that the result *Neisseria gonorrhoeae* (``r as.mo("Neisseria gonorrhoeae")``) needs review.
#'
#' There are three helper functions that can be run after using the [as.mo()] function:
#' - Use [mo_uncertainties()] to get a [data.frame] that prints in a pretty format with all taxonomic names that were guessed. The output contains the matching score for all matches (see *Background on matching score*).
#' - Use [mo_failures()] to get a [character] [vector] with all values that could not be coerced to a valid value.
#' - Use [mo_renamed()] to get a [data.frame] with all values that could be coerced based on old, previously accepted taxonomic names.
#'
#' ## Microbial prevalence of pathogens in humans
#'
#' The intelligent rules consider the prevalence of microorganisms in humans grouped into three groups, which is available as the `prevalence` columns in the [microorganisms] and [microorganisms.old] data sets. The grouping into human pathogenic prevalence is explained in the section *Matching score for microorganisms* below.
#' @inheritSection mo_matching_score Matching score for microorganisms
#' @inheritSection catalogue_of_life Catalogue of Life
#  (source as a section here, so it can be inherited by other man pages:)
#' @section Source:
#' 1. Becker K *et al.* **Coagulase-Negative Staphylococci**. 2014. Clin Microbiol Rev. 27(4): 870–926; \doi{10.1128/CMR.00109-13}
#' 2. Becker K *et al.* **Implications of identifying the recently defined members of the *S. aureus* complex, *S. argenteus* and *S. schweitzeri*: A position paper of members of the ESCMID Study Group for staphylococci and Staphylococcal Diseases (ESGS).** 2019. Clin Microbiol Infect; \doi{10.1016/j.cmi.2019.02.028}
#' 3. Becker K *et al.* **Emergence of coagulase-negative staphylococci** 2020. Expert Rev Anti Infect Ther. 18(4):349-366; \doi{10.1080/14787210.2020.1730813}
#' 4. Lancefield RC **A serological differentiation of human and other groups of hemolytic streptococci**. 1933. J Exp Med. 57(4): 571–95; \doi{10.1084/jem.57.4.571}
#' 5. Catalogue of Life: Annual Checklist (public online taxonomic database), <http://www.catalogueoflife.org> (check included annual version with [catalogue_of_life_version()]).
#' @export
#' @return A [character] [vector] with additional class [`mo`]
#' @seealso [microorganisms] for the [data.frame] that is being used to determine ID's.
#'
#' The [`mo_*`][mo_property()] functions (such as [mo_genus()], [mo_gramstain()]) to get properties based on the returned code.
#' @inheritSection AMR Reference data publicly available
#' @inheritSection AMR Read more on our website!
#' @examples
#' \donttest{
#' # These examples all return "B_STPHY_AURS", the ID of S. aureus:
#' as.mo("sau") # WHONET code
#' as.mo("stau")
#' as.mo("STAU")
#' as.mo("staaur")
#' as.mo("S. aureus")
#' as.mo("S aureus")
#' as.mo("Staphylococcus aureus")
#' as.mo("Staphylococcus aureus (MRSA)")
#' as.mo("Zthafilokkoockus oureuz") # handles incorrect spelling
#' as.mo("MRSA")    # Methicillin Resistant S. aureus
#' as.mo("VISA")    # Vancomycin Intermediate S. aureus
#' as.mo("VRSA")    # Vancomycin Resistant S. aureus
#' as.mo(115329001) # SNOMED CT code
#'
#' # Dyslexia is no problem - these all work:
#' as.mo("Ureaplasma urealyticum")
#' as.mo("Ureaplasma urealyticus")
#' as.mo("Ureaplasmium urealytica")
#' as.mo("Ureaplazma urealitycium")
#'
#' as.mo("Streptococcus group A")
#' as.mo("GAS") # Group A Streptococci
#' as.mo("GBS") # Group B Streptococci
#'
#' as.mo("S. epidermidis")                 # will remain species: B_STPHY_EPDR
#' as.mo("S. epidermidis", Becker = TRUE)  # will not remain species: B_STPHY_CONS
#'
#' as.mo("S. pyogenes")                    # will remain species: B_STRPT_PYGN
#' as.mo("S. pyogenes", Lancefield = TRUE) # will not remain species: B_STRPT_GRPA
#'
#' # All mo_* functions use as.mo() internally too (see ?mo_property):
#' mo_genus("E. coli")                           # returns "Escherichia"
#' mo_gramstain("E. coli")                       # returns "Gram negative"
#' mo_is_intrinsic_resistant("E. coli", "vanco") # returns TRUE
#' }
as.mo <- function(x,
                  Becker = FALSE,
                  Lancefield = FALSE,
                  allow_uncertain = TRUE,
                  reference_df = get_mo_source(),
                  ignore_pattern = getOption("AMR_ignore_pattern"),
                  language = get_locale(),
                  ...) {
  meet_criteria(x, allow_class = c("mo", "data.frame", "list", "character", "numeric", "integer", "factor"), allow_NA = TRUE)
  meet_criteria(Becker, allow_class = c("logical", "character"), has_length = 1)
  meet_criteria(Lancefield, allow_class = c("logical", "character"), has_length = 1)
  meet_criteria(allow_uncertain, allow_class = c("logical", "numeric", "integer"), has_length = 1)
  meet_criteria(reference_df, allow_class = "data.frame", allow_NULL = TRUE)
  meet_criteria(ignore_pattern, allow_class = "character", has_length = 1, allow_NULL = TRUE)
  meet_criteria(language, has_length = 1, is_in = c(LANGUAGES_SUPPORTED, ""), allow_NULL = TRUE, allow_NA = TRUE)

  check_dataset_integrity()

  if (tryCatch(all(x[!is.na(x)] %in% MO_lookup$mo)
               & isFALSE(Becker)
               & isFALSE(Lancefield), error = function(e) FALSE)) {
    # don't look into valid MO codes, just return them
    # is.mo() won't work - MO codes might change between package versions
    return(set_clean_class(x, new_class = c("mo", "character")))
  }

  if (tryCatch(all(x == "" | gsub(".*(unknown ).*", "unknown name", tolower(x), perl = TRUE) %in% MO_lookup$fullname_lower, na.rm = TRUE)
               & isFALSE(Becker)
               & isFALSE(Lancefield), error = function(e) FALSE)) {
    # to improve speed, special case for taxonomically correct full names (case-insensitive)
    return(MO_lookup[match(gsub(".*(unknown ).*", "unknown name", tolower(x), perl = TRUE), MO_lookup$fullname_lower), "mo", drop = TRUE])
  }

  # start off with replaced language-specific non-ASCII characters with ASCII characters
  x <- parse_and_convert(x)
  # replace mo codes used in older package versions
  x <- replace_old_mo_codes(x, property = "mo")
  # ignore cases that match the ignore pattern
  x <- replace_ignore_pattern(x, ignore_pattern)

  # WHONET: xxx = no growth
  x[tolower(as.character(paste0(x, ""))) %in% c("", "xxx", "na", "nan")] <- NA_character_
  # Laboratory systems: remove (translated) entries like "no growth", etc.
  x[trimws2(x) %like% translate_AMR("no .*growth", language = language)] <- NA_character_
  x[trimws2(x) %like% paste0("^(", translate_AMR("no|not", language = language), ") [a-z]+")] <- "UNKNOWN"
  uncertainty_level <- translate_allow_uncertain(allow_uncertain)

  if (!is.null(reference_df)
      && check_validity_mo_source(reference_df)
      && isFALSE(Becker)
      && isFALSE(Lancefield)
      && all(x %in% unlist(reference_df), na.rm = TRUE)) {

    reference_df <- repair_reference_df(reference_df)
    suppressWarnings(
      y <- data.frame(x = x, stringsAsFactors = FALSE) %pm>%
        pm_left_join(reference_df, by = "x") %pm>%
        pm_pull(mo) 
    )

  } else if (all(x[!is.na(x)] %in% MO_lookup$mo)
             & isFALSE(Becker)
             & isFALSE(Lancefield)) {
    y <- x

  } else {
    # will be checked for mo class in validation and uses exec_as.mo internally if necessary
    y <- mo_validate(x = x, property = "mo",
                     Becker = Becker, Lancefield = Lancefield,
                     allow_uncertain = uncertainty_level,
                     reference_df = reference_df,
                     ignore_pattern = ignore_pattern,
                     language = language,
                     ...)
  }

  set_clean_class(y,
                  new_class = c("mo", "character"))
}

#' @rdname as.mo
#' @export
is.mo <- function(x) {
  inherits(x, "mo")
}

# param property a column name of microorganisms
# param initial_search logical - is FALSE when coming from uncertain tries, which uses exec_as.mo internally too
# param dyslexia_mode logical - also check for characters that resemble others
# param debug logical - show different lookup texts while searching
# param reference_data_to_use data.frame - the data set to check for
# param actual_uncertainty - (only for initial_search = FALSE) the actual uncertainty level used in the function for score calculation (sometimes passed as 2 or 3 by uncertain_fn())
# param actual_input - (only for initial_search = FALSE) the actual, original input
# param language - used for translating "no growth", etc.
exec_as.mo <- function(x,
                       Becker = FALSE,
                       Lancefield = FALSE,
                       allow_uncertain = TRUE,
                       reference_df = get_mo_source(),
                       property = "mo",
                       initial_search = TRUE,
                       dyslexia_mode = FALSE,
                       debug = FALSE,
                       ignore_pattern = getOption("AMR_ignore_pattern"),
                       reference_data_to_use = MO_lookup,
                       actual_uncertainty = 1,
                       actual_input = NULL,
                       language = get_locale()) {
  meet_criteria(x, allow_class = c("mo", "data.frame", "list", "character", "numeric", "integer", "factor"), allow_NA = TRUE)
  meet_criteria(Becker, allow_class = c("logical", "character"), has_length = 1)
  meet_criteria(Lancefield, allow_class = c("logical", "character"), has_length = 1)
  meet_criteria(allow_uncertain, allow_class = c("logical", "numeric", "integer"), has_length = 1)
  meet_criteria(reference_df, allow_class = "data.frame", allow_NULL = TRUE)
  meet_criteria(property, allow_class = "character", has_length = 1, is_in = colnames(microorganisms))
  meet_criteria(initial_search, allow_class = "logical", has_length = 1)
  meet_criteria(dyslexia_mode, allow_class = "logical", has_length = 1)
  meet_criteria(debug, allow_class = "logical", has_length = 1)
  meet_criteria(ignore_pattern, allow_class = "character", has_length = 1, allow_NULL = TRUE)
  meet_criteria(reference_data_to_use, allow_class = "data.frame")
  meet_criteria(actual_uncertainty, allow_class = "numeric", has_length = 1)
  meet_criteria(actual_input, allow_class = "character", allow_NULL = TRUE)
  meet_criteria(language, has_length = 1, is_in = c(LANGUAGES_SUPPORTED, ""), allow_NULL = TRUE, allow_NA = TRUE)

  check_dataset_integrity()
  
  lookup <- function(needle,
                     column = property,
                     haystack = reference_data_to_use,
                     n = 1,
                     debug_mode = debug,
                     initial = initial_search,
                     uncertainty = actual_uncertainty,
                     input_actual = actual_input) {

    if (!is.null(input_actual)) {
      input <- input_actual
    } else {
      input <- tryCatch(x_backup[i], error = function(e) "")
    }

    # `column` can be NULL for all columns, or a selection
    # returns a character (vector) - if `column` > length 1 then with columns as names
    if (isTRUE(debug_mode)) {
      cat(font_silver("looking up: ", substitute(needle), collapse = ""))
    }
    if (length(column) == 1) {
      res_df <- haystack[which(eval(substitute(needle), envir = haystack, enclos = parent.frame())), , drop = FALSE]
      if (NROW(res_df) > 1 & uncertainty != -1) {
        # sort the findings on matching score
        scores <- mo_matching_score(x = input,
                                    n = res_df[, "fullname", drop = TRUE])
        res_df <- res_df[order(scores, decreasing = TRUE), , drop = FALSE]
      }
      res <- as.character(res_df[, column, drop = TRUE])
      if (length(res) == 0) {
        if (isTRUE(debug_mode)) {
          cat(font_red(" (no match)\n"))
        }
        NA_character_
      } else {
        if (isTRUE(debug_mode)) {
          cat(font_green(paste0(" **MATCH** (", NROW(res_df), " results)\n")))
        }
        if ((length(res) > n | uncertainty > 1) & uncertainty != -1) {
          # save the other possible results as well, but not for forced certain results (then uncertainty == -1)
          uncertainties <<- rbind(uncertainties,
                                  format_uncertainty_as_df(uncertainty_level = uncertainty,
                                                           input = input,
                                                           result_mo = res_df[1, "mo", drop = TRUE],
                                                           candidates = as.character(res_df[, "fullname", drop = TRUE])),
                                  stringsAsFactors = FALSE)
        }
        res[seq_len(min(n, length(res)))]
      }
    } else {
      if (isTRUE(debug_mode)) {
        cat("\n")
      }
      if (is.null(column)) {
        column <- names(haystack)
      }
      res <- haystack[which(eval(substitute(needle), envir = haystack, enclos = parent.frame())), , drop = FALSE]
      res <- res[seq_len(min(n, nrow(res))), column, drop = TRUE]
      if (NROW(res) == 0) {
        res <- rep(NA_character_, length(column))
      }
      res <- as.character(res)
      names(res) <- column
      res
    }
  }

  # start off with replaced language-specific non-ASCII characters with ASCII characters
  x <- parse_and_convert(x)
  # replace mo codes used in older package versions
  x <- replace_old_mo_codes(x, property)
  # ignore cases that match the ignore pattern
  x <- replace_ignore_pattern(x, ignore_pattern)

  # WHONET: xxx = no growth
  x[tolower(as.character(paste0(x, ""))) %in% c("", "xxx", "na", "nan")] <- NA_character_
  # Laboratory systems: remove (translated) entries like "no growth", etc.
  x[trimws2(x) %like% translate_AMR("no .*growth", language = language)] <- NA_character_
  x[trimws2(x) %like% paste0("^(", translate_AMR("no|not", language = language), ") [a-z]+")] <- "UNKNOWN"

  if (initial_search == TRUE) {
    # keep track of time - give some hints to improve speed if it takes a long time
    start_time <- Sys.time()
    
    pkg_env$mo_failures <- NULL
    pkg_env$mo_uncertainties <- NULL
    pkg_env$mo_renamed <- NULL
  }
  pkg_env$mo_renamed_last_run <- NULL

  failures <- character(0)
  uncertainty_level <- translate_allow_uncertain(allow_uncertain)
  uncertainties <- data.frame(uncertainty = integer(0),
                              input = character(0),
                              fullname = character(0),
                              renamed_to = character(0),
                              mo = character(0),
                              candidates = character(0),
                              stringsAsFactors = FALSE)

  x_input <- x
  # already strip leading and trailing spaces
  x <- trimws(x)
  # only check the uniques, which is way faster
  x <- unique(x)
  # remove empty values (to later fill them in again with NAs)
  # ("xxx" is WHONET code for 'no growth')
  x <- x[!is.na(x)
         & !is.null(x)
         & !identical(x, "")
         & !identical(x, "xxx")]

  # defined df to check for
  if (!is.null(reference_df)) {
    check_validity_mo_source(reference_df)
    reference_df <- repair_reference_df(reference_df)
  }

  # all empty
  if (all(identical(trimws(x_input), "") | is.na(x_input) | length(x) == 0)) {
    if (property == "mo") {
      return(set_clean_class(rep(NA_character_, length(x_input)),
                             new_class = c("mo", "character")))
    } else {
      return(rep(NA_character_, length(x_input)))
    }

  } else if (all(x %in% reference_df[, 1][[1]])) {
    # all in reference df
    colnames(reference_df)[1] <- "x"
    suppressWarnings(
      x <- MO_lookup[match(reference_df[match(x, reference_df$x), "mo", drop = TRUE], MO_lookup$mo), property, drop = TRUE]
    )

  } else if (all(x %in% reference_data_to_use$mo)) {
    x <- MO_lookup[match(x, MO_lookup$mo), property, drop = TRUE]

  } else if (all(tolower(x) %in% reference_data_to_use$fullname_lower)) {
    # we need special treatment for very prevalent full names, they are likely!
    # e.g. as.mo("Staphylococcus aureus")
    x <- MO_lookup[match(tolower(x), MO_lookup$fullname_lower), property, drop = TRUE]

  } else if (all(x %in% reference_data_to_use$fullname)) {
    # we need special treatment for very prevalent full names, they are likely!
    # e.g. as.mo("Staphylococcus aureus")
    x <- MO_lookup[match(x, MO_lookup$fullname), property, drop = TRUE]

  } else if (all(toupper(x) %in% microorganisms.codes$code)) {
    # commonly used MO codes
    x <- MO_lookup[match(microorganisms.codes[match(toupper(x),
                                                    microorganisms.codes$code),
                                              "mo",
                                              drop = TRUE],
                         MO_lookup$mo),
                   property,
                   drop = TRUE]

  } else if (!all(x %in% microorganisms[, property])) {

    strip_whitespace <- function(x, dyslexia_mode) {
      # all whitespaces (tab, new lines, etc.) should be one space
      # and spaces before and after should be left blank
      trimmed <- trimws2(x)
      # also, make sure the trailing and leading characters are a-z or 0-9
      # in case of non-regex
      if (dyslexia_mode == FALSE) {
        trimmed <- gsub("^[^a-zA-Z0-9)(]+", "", trimmed, perl = TRUE)
        trimmed <- gsub("[^a-zA-Z0-9)(]+$", "", trimmed, perl = TRUE)
      }
      trimmed
    }

    x_backup_untouched <- x
    x <- strip_whitespace(x, dyslexia_mode)
    x_backup <- x

    # from here on case-insensitive
    x <- tolower(x)

    x_backup[grepl("^(fungus|fungi)$", x)] <- "F_FUNGUS" # will otherwise become the kingdom

    # remove spp and species
    x <- gsub(" +(spp.?|ssp.?|sp.? |ss ?.?|subsp.?|subspecies|biovar |serovar |species)", " ", x, perl = TRUE)
    x <- gsub("(spp.?|subsp.?|subspecies|biovar|serovar|species)", "", x, perl = TRUE)
    x <- gsub("^([a-z]{2,4})(spe.?)$", "\\1", x, perl = TRUE) # when ending in SPE instead of SPP and preceded by 2-4 characters
    x <- strip_whitespace(x, dyslexia_mode)

    x_backup_without_spp <- x
    x_species <- paste(x, "species")
    # translate to English for supported languages of mo_property
    x <- gsub("(gruppe|groep|grupo|gruppo|groupe)", "group", x, perl = TRUE)
    # no groups and complexes as ending
    x <- gsub("(complex|group)$", "", x, perl = TRUE)
    x <- gsub("((an)?aero+b)[a-z]*", "", x, perl = TRUE)
    x <- gsub("^atyp[a-z]*", "", x, perl = TRUE)
    x <- gsub("(vergroen)[a-z]*", "viridans", x, perl = TRUE)
    x <- gsub("[a-z]*diff?erent[a-z]*", "", x, perl = TRUE)
    x <- gsub("(hefe|gist|gisten|levadura|lievito|fermento|levure)[a-z]*", "yeast", x, perl = TRUE)
    x <- gsub("(schimmels?|mofo|molde|stampo|moisissure|fungi)[a-z]*", "fungus", x, perl = TRUE)
    x <- gsub("fungus[ph|f]rya", "fungiphrya", x, perl = TRUE)
    # no contamination
    x <- gsub("(contamination|kontamination|mengflora|contaminaci.n|contamina..o)", "", x, perl = TRUE)
    # remove non-text in case of "E. coli" except dots and spaces
    x <- trimws(gsub("[^.a-zA-Z0-9/ \\-]+", " ", x, perl = TRUE))
    # but make sure that dots are followed by a space
    x <- gsub("[.] ?", ". ", x, perl = TRUE)
    # replace minus by a space
    x <- gsub("-+", " ", x, perl = TRUE)
    # replace hemolytic by haemolytic
    x <- gsub("ha?emoly", "haemoly", x, perl = TRUE)
    # place minus back in streptococci
    x <- gsub("(alpha|beta|gamma).?ha?emoly", "\\1-haemoly", x, perl = TRUE)
    # remove genus as first word
    x <- gsub("^genus ", "", x, perl = TRUE)
    # remove 'uncertain'-like texts
    x <- trimws(gsub("(uncertain|susp[ie]c[a-z]+|verdacht)", "", x, perl = TRUE))
    # allow characters that resemble others = dyslexia_mode ----
    if (dyslexia_mode == TRUE) {
      x <- tolower(x)
      x <- gsub("[iy]+", "[iy]+", x, perl = TRUE)
      x <- gsub("(c|k|q|qu|s|z|x|ks)+", "(c|k|q|qu|s|z|x|ks)+", x, perl = TRUE)
      x <- gsub("(ph|hp|f|v)+", "(ph|hp|f|v)+", x, perl = TRUE)
      x <- gsub("(th|ht|t)+", "(th|ht|t)+", x, perl = TRUE)
      x <- gsub("a+", "a+", x, perl = TRUE)
      x <- gsub("u+", "u+", x, perl = TRUE)
      # allow any ending of -um, -us, -ium, -icum, -ius, -icus, -ica, -ia and -a (needs perl for the negative backward lookup):
      x <- gsub("(u\\+\\(c\\|k\\|q\\|qu\\+\\|s\\|z\\|x\\|ks\\)\\+)(?![a-z])",
                "(u[s|m]|[iy][ck]?u[ms]|[iy]?[ck]?a)", x, perl = TRUE)
      x <- gsub("(\\[iy\\]\\+\\(c\\|k\\|q\\|qu\\+\\|s\\|z\\|x\\|ks\\)\\+a\\+)(?![a-z])",
                "(u[s|m]|[iy][ck]?u[ms]|[iy]?[ck]?a)", x, perl = TRUE)
      x <- gsub("(\\[iy\\]\\+u\\+m)(?![a-z])",
                "(u[s|m]|[iy][ck]?u[ms]|[iy]?[ck]?a)", x, perl = TRUE)
      x <- gsub("(\\[iy\\]\\+a\\+)(?![a-z])",
                "([iy]*a+|[iy]+a*)", x, perl = TRUE)
      x <- gsub("e+", "e+", x, perl = TRUE)
      x <- gsub("o+", "o+", x, perl = TRUE)
      x <- gsub("(.)\\1+", "\\1+", x, perl = TRUE)
      # allow multiplication of all other consonants
      x <- gsub("([bdgjlnrw]+)", "\\1+", x, perl = TRUE)
      # allow ending in -en or -us
      x <- gsub("e\\+n(?![a-z[])", "(e+n|u+(c|k|q|qu|s|z|x|ks)+)", x, perl = TRUE)
      # if the input is longer than 10 characters, allow any forgotten consonant between all characters, as some might just have forgotten one...
      # this will allow "Pasteurella damatis" to be correctly read as "Pasteurella dagmatis".
      consonants <- paste(letters[!letters %in% c("a", "e", "i", "o", "u")], collapse = "")
      x[nchar(x_backup_without_spp) > 10] <- gsub("[+]", paste0("+[", consonants, "]?"), x[nchar(x_backup_without_spp) > 10])
      # allow au and ou after all these regex implementations
      x <- gsub("a+[bcdfghjklmnpqrstvwxyz]?u+[bcdfghjklmnpqrstvwxyz]?", "(a+u+|o+u+)[bcdfghjklmnpqrstvwxyz]?", x, fixed = TRUE)
      x <- gsub("o+[bcdfghjklmnpqrstvwxyz]?u+[bcdfghjklmnpqrstvwxyz]?", "(a+u+|o+u+)[bcdfghjklmnpqrstvwxyz]?", x, fixed = TRUE)
    }
    x <- strip_whitespace(x, dyslexia_mode)
    # make sure to remove regex overkill (will lead to errors)
    x <- gsub("++", "+", x, fixed = TRUE)
    x <- gsub("?+", "?", x, fixed = TRUE)

    x_trimmed <- x
    x_trimmed_species <- paste(x_trimmed, "species")
    x_trimmed_without_group <- gsub(" gro.u.p$", "", x_trimmed, perl = TRUE)
    # remove last part from "-" or "/"
    x_trimmed_without_group <- gsub("(.*)[-/].*", "\\1", x_trimmed_without_group)
    # replace space and dot by regex sign
    x_withspaces <- gsub("[ .]+", ".* ", x, perl = TRUE)
    x <- gsub("[ .]+", ".*", x, perl = TRUE)
    # add start en stop regex
    x <- paste0("^", x, "$")

    x_withspaces_start_only <- paste0("^", x_withspaces)
    x_withspaces_end_only <- paste0(x_withspaces, "$")
    x_withspaces_start_end <- paste0("^", x_withspaces, "$")

    if (isTRUE(debug)) {
      cat(paste0(font_blue("x"), '                       "', x, '"\n'))
      cat(paste0(font_blue("x_species"), '               "', x_species, '"\n'))
      cat(paste0(font_blue("x_withspaces_start_only"), ' "', x_withspaces_start_only, '"\n'))
      cat(paste0(font_blue("x_withspaces_end_only"), '   "', x_withspaces_end_only, '"\n'))
      cat(paste0(font_blue("x_withspaces_start_end"), '  "', x_withspaces_start_end, '"\n'))
      cat(paste0(font_blue("x_backup"), '                "', x_backup, '"\n'))
      cat(paste0(font_blue("x_backup_without_spp"), '    "', x_backup_without_spp, '"\n'))
      cat(paste0(font_blue("x_trimmed"), '               "', x_trimmed, '"\n'))
      cat(paste0(font_blue("x_trimmed_species"), '       "', x_trimmed_species, '"\n'))
      cat(paste0(font_blue("x_trimmed_without_group"), ' "', x_trimmed_without_group, '"\n'))
    }

    if (initial_search == TRUE) {
      progress <- progress_ticker(n = length(x), n_min = 25) # start if n >= 25
      on.exit(close(progress))
    }

    for (i in seq_len(length(x))) {

      if (initial_search == TRUE) {
        progress$tick()
      }

      # valid MO code ----
      found <- lookup(mo == toupper(x_backup[i]))
      if (!is.na(found)) {
        x[i] <- found[1L]
        next
      }

      # valid fullname ----
      found <- lookup(fullname_lower %in% gsub("[^a-zA-Z0-9_. -]", "", tolower(c(x_backup[i], x_backup_without_spp[i])), perl = TRUE))
      # added the gsub() for "(unknown fungus)", since fullname_lower does not contain brackets
      if (!is.na(found)) {
        x[i] <- found[1L]
        next
      }

      # old fullname ----
      found <- lookup(fullname_lower %in% tolower(c(x_backup[i], x_backup_without_spp[i])),
                      column = NULL, # all columns
                      haystack = MO.old_lookup)
      if (!all(is.na(found))) {
        # when property is "ref" (which is the case in mo_ref, mo_authors and mo_year), return the old value, so:
        # mo_ref() of "Chlamydia psittaci" will be "Page, 1968" (with warning)
        # mo_ref() of "Chlamydophila psittaci" will be "Everett et al., 1999"
        if (property == "ref") {
          x[i] <- found["ref"]
        } else {
          x[i] <- lookup(fullname == found["fullname_new"], haystack = MO_lookup)
        }
        pkg_env$mo_renamed_last_run <- found["fullname"]
        was_renamed(name_old = found["fullname"],
                    name_new = lookup(fullname == found["fullname_new"], "fullname", haystack = MO_lookup),
                    ref_old = found["ref"],
                    ref_new = lookup(fullname == found["fullname_new"], "ref", haystack = MO_lookup),
                    mo = lookup(fullname == found["fullname_new"], "mo", haystack = MO_lookup))
        next
      }

      if (x_backup[i] %like_case% "\\(unknown [a-z]+\\)" | tolower(x_backup_without_spp[i]) %in% c("other", "none", "unknown")) {
        # empty and nonsense values, ignore without warning
        x[i] <- lookup(mo == "UNKNOWN")
        next
      }

      # exact SNOMED code ----
      if (x_backup[i] %like% "^[0-9]+$") {
        snomed_found <- unlist(lapply(reference_data_to_use$snomed,
                                      function(s) if (x_backup[i] %in% s) {
                                        TRUE
                                      } else {
                                        FALSE
                                      }))
        if (sum(snomed_found, na.rm = TRUE) > 0) {
          found <- reference_data_to_use[snomed_found == TRUE, property][[1]]
          if (!is.na(found)) {
            x[i] <- found[1L]
            next
          }
        }
      }

      # very probable: is G. species ----
      found <- lookup(g_species %in% gsub("[^a-z0-9/ \\-]+", "",
                                          tolower(c(x_backup[i], x_backup_without_spp[i])), perl = TRUE))
      if (!is.na(found)) {
        x[i] <- found[1L]
        next
      }

      # WHONET and other common LIS codes ----
      found <- microorganisms.codes[which(microorganisms.codes$code %in% toupper(c(x_backup_untouched[i], x_backup[i], x_backup_without_spp[i]))), "mo", drop = TRUE][1L]
      if (!is.na(found)) {
        x[i] <- lookup(mo == found)
        next
      }

      # user-defined reference ----
      if (!is.null(reference_df)) {
        if (x_backup[i] %in% reference_df[, 1]) {
          # already checked integrity of reference_df, all MOs are valid
          ref_mo <- reference_df[reference_df[, 1] == x_backup[i], "mo"][[1L]]
          x[i] <- lookup(mo == ref_mo)
          next
        }
      }

      # WHONET: xxx = no growth
      if (tolower(as.character(paste0(x_backup_without_spp[i], ""))) %in% c("", "xxx", "na", "nan")) {
        x[i] <- NA_character_
        next
      }

      # check for very small input, but ignore the O antigens of E. coli
      if (nchar(gsub("[^a-zA-Z]", "", x_trimmed[i])) < 3
          & !toupper(x_backup_without_spp[i]) %like_case% "O?(26|103|104|104|111|121|145|157)") {
        # fewer than 3 chars and not looked for species, add as failure
        x[i] <- lookup(mo == "UNKNOWN")
        if (initial_search == TRUE) {
          failures <- c(failures, x_backup[i])
        }
        next
      }

      if (x_backup_without_spp[i] %like_case% "(virus|viridae)") {
        # there is no fullname like virus or viridae, so don't try to coerce it
        x[i] <- NA_character_
        next
      }

      # translate known trivial abbreviations to genus + species ----
      if (toupper(x_backup_without_spp[i]) %in% c("MRSA", "MSSA", "VISA", "VRSA", "BORSA")
          | x_backup_without_spp[i] %like_case% "(^| )(mrsa|mssa|visa|vrsa|borsa|la-?mrsa|ca-?mrsa)( |$)") {
        x[i] <- lookup(fullname == "Staphylococcus aureus", uncertainty = -1)
        next
      }
      if (toupper(x_backup_without_spp[i]) %in% c("MRSE", "MSSE")
          | x_backup_without_spp[i] %like_case% "(^| )(mrse|msse)( |$)") {
        x[i] <- lookup(fullname == "Staphylococcus epidermidis", uncertainty = -1)
        next
      }
      if (toupper(x_backup_without_spp[i]) == "VRE"
          | x_backup_without_spp[i] %like_case% "(^| )vre "
          | x_backup_without_spp[i] %like_case% "(enterococci|enterokok|enterococo)[a-z]*?$")  {
        x[i] <- lookup(genus == "Enterococcus", uncertainty = -1)
        next
      }
      # support for:
      # - AIEC (Adherent-Invasive E. coli)
      # - ATEC (Atypical Entero-pathogenic E. coli)
      # - DAEC (Diffusely Adhering E. coli)
      # - EAEC (Entero-Aggresive E. coli)
      # - EHEC (Entero-Haemorrhagic E. coli)
      # - EIEC (Entero-Invasive E. coli)
      # - EPEC (Entero-Pathogenic E. coli)
      # - ETEC (Entero-Toxigenic E. coli)
      # - NMEC (Neonatal Meningitis‐causing E. coli)
      # - STEC (Shiga-toxin producing E. coli)
      # - UPEC (Uropathogenic E. coli)
      if (toupper(x_backup_without_spp[i]) %in% c("AIEC", "ATEC", "DAEC", "EAEC", "EHEC", "EIEC", "EPEC", "ETEC", "NMEC", "STEC", "UPEC")
          # also support O-antigens of E. coli: O26, O103, O104, O111, O121, O145, O157
          | x_backup_without_spp[i] %like_case% "o?(26|103|104|111|121|145|157)") {
        x[i] <- lookup(fullname == "Escherichia coli", uncertainty = -1)
        next
      }
      if (toupper(x_backup_without_spp[i]) == "MRPA"
          | x_backup_without_spp[i] %like_case% "(^| )mrpa( |$)") {
        # multi resistant P. aeruginosa
        x[i] <- lookup(fullname == "Pseudomonas aeruginosa", uncertainty = -1)
        next
      }
      if (toupper(x_backup_without_spp[i]) == "CRSM") {
        # co-trim resistant S. maltophilia
        x[i] <- lookup(fullname == "Stenotrophomonas maltophilia", uncertainty = -1)
        next
      }
      if (toupper(x_backup_without_spp[i]) %in% c("PISP", "PRSP", "VISP", "VRSP")
          | x_backup_without_spp[i] %like_case% "(^| )(pisp|prsp|visp|vrsp)( |$)") {
        # peni I, peni R, vanco I, vanco R: S. pneumoniae
        x[i] <- lookup(fullname == "Streptococcus pneumoniae", uncertainty = -1)
        next
      }
      if (x_backup_without_spp[i] %like_case% "^g[abcdfghk]s$") {
        # Streptococci, like GBS = Group B Streptococci (B_STRPT_GRPB)
        x[i] <- lookup(mo == toupper(gsub("g([abcdfghk])s",
                                          "B_STRPT_GRP\\1",
                                          x_backup_without_spp[i])), uncertainty = -1)
        next
      }
      if (x_backup_without_spp[i] %like_case% "(streptococ|streptokok).* [abcdfghk]$") {
        # Streptococci in different languages, like "estreptococos grupo B"
        x[i] <- lookup(mo == toupper(gsub(".*(streptococ|streptokok|estreptococ).* ([abcdfghk])$",
                                          "B_STRPT_GRP\\2",
                                          x_backup_without_spp[i])), uncertainty = -1)
        next
      }
      if (x_backup_without_spp[i] %like_case% "group [abcdfghk] (streptococ|streptokok|estreptococ)") {
        # Streptococci in different languages, like "Group A Streptococci"
        x[i] <- lookup(mo == toupper(gsub(".*group ([abcdfghk]) (streptococ|streptokok|estreptococ).*",
                                          "B_STRPT_GRP\\1",
                                          x_backup_without_spp[i])), uncertainty = -1)
        next
      }
      if (x_backup_without_spp[i] %like_case% "haemoly.*strep") {
        # Haemolytic streptococci in different languages
        x[i] <- lookup(mo == "B_STRPT_HAEM", uncertainty = -1)
        next
      }
      # CoNS/CoPS in different languages (support for German, Dutch, Spanish, Portuguese) ----
      if (x_backup_without_spp[i] %like_case% "[ck]oagulas[ea] negatie?[vf]"
          | x_trimmed[i] %like_case% "[ck]oagulas[ea] negatie?[vf]"
          | x_backup_without_spp[i] %like_case% "[ck]o?ns[^a-z]?$") {
        # coerce S. coagulase negative
        x[i] <- lookup(mo == "B_STPHY_CONS", uncertainty = -1)
        next
      }
      if (x_backup_without_spp[i] %like_case% "[ck]oagulas[ea] positie?[vf]"
          | x_trimmed[i] %like_case% "[ck]oagulas[ea] positie?[vf]"
          | x_backup_without_spp[i] %like_case% "[ck]o?ps[^a-z]?$") {
        # coerce S. coagulase positive
        x[i] <- lookup(mo == "B_STPHY_COPS", uncertainty = -1)
        next
      }
      # streptococcal groups: milleri and viridans
      if (x_trimmed[i] %like_case% "strepto.* mil+er+i"
          | x_backup_without_spp[i] %like_case% "strepto.* mil+er+i"
          | x_backup_without_spp[i] %like_case% "mgs[^a-z]?$") {
        # Milleri Group Streptococcus (MGS)
        x[i] <- lookup(mo == "B_STRPT_MILL", uncertainty = -1)
        next
      }
      if (x_trimmed[i] %like_case% "strepto.* viridans"
          | x_backup_without_spp[i] %like_case% "strepto.* viridans"
          | x_backup_without_spp[i] %like_case% "vgs[^a-z]?$") {
        # Viridans Group Streptococcus (VGS)
        x[i] <- lookup(mo == "B_STRPT_VIRI", uncertainty = -1)
        next
      }
      if (x_backup_without_spp[i] %like_case% "gram[ -]?neg.*"
          | x_backup_without_spp[i] %like_case% "negatie?[vf]"
          | x_trimmed[i] %like_case% "gram[ -]?neg.*") {
        # coerce Gram negatives
        x[i] <- lookup(mo == "B_GRAMN", uncertainty = -1)
        next
      }
      if (x_backup_without_spp[i] %like_case% "gram[ -]?pos.*"
          | x_backup_without_spp[i] %like_case% "positie?[vf]"
          | x_trimmed[i] %like_case% "gram[ -]?pos.*") {
        # coerce Gram positives
        x[i] <- lookup(mo == "B_GRAMP", uncertainty = -1)
        next
      }
      if (x_backup_without_spp[i] %like_case% "mycoba[ck]teri.[nm]?$") {
        # coerce mycobacteria in multiple languages
        x[i] <- lookup(genus == "Mycobacterium", uncertainty = -1)
        next
      }

      if (x_backup_without_spp[i] %like_case% "salmonella [a-z]+ ?.*") {
        if (x_backup_without_spp[i] %like_case% "salmonella group") {
          # Salmonella Group A to Z, just return S. species for now
          x[i] <- lookup(genus == "Salmonella", uncertainty = -1)
          next
        } else if (grepl("[sS]almonella [A-Z][a-z]+ ?.*", x_backup[i], ignore.case = FALSE) &
                   !x_backup[i] %like% "t[iy](ph|f)[iy]") {
          # Salmonella with capital letter species like "Salmonella Goettingen" - they're all S. enterica
          # except for S. typhi, S. paratyphi, S. typhimurium
          x[i] <- lookup(fullname == "Salmonella enterica", uncertainty = -1)
          uncertainties <- rbind(uncertainties,
                                 format_uncertainty_as_df(uncertainty_level = 1,
                                                          input = x_backup[i],
                                                          result_mo = lookup(fullname == "Salmonella enterica", "mo", uncertainty = -1)),
                                 stringsAsFactors = FALSE)
          next
        }
      }

      # trivial names known to the field:
      if ("meningococcus" %like_case% x_trimmed[i]) {
        # coerce Neisseria meningitidis
        x[i] <- lookup(fullname == "Neisseria meningitidis", uncertainty = -1)
        next
      }
      if ("gonococcus" %like_case% x_trimmed[i]) {
        # coerce Neisseria gonorrhoeae
        x[i] <- lookup(fullname == "Neisseria gonorrhoeae", uncertainty = -1)
        next
      }
      if ("pneumococcus" %like_case% x_trimmed[i]) {
        # coerce Streptococcus penumoniae
        x[i] <- lookup(fullname == "Streptococcus pneumoniae", uncertainty = -1)
        next
      }
      # }

      # NOW RUN THROUGH DIFFERENT PREVALENCE LEVELS
      check_per_prevalence <- function(data_to_check,
                                       data.old_to_check,
                                       a.x_backup,
                                       b.x_trimmed,
                                       c.x_trimmed_without_group,
                                       d.x_withspaces_start_end,
                                       e.x_withspaces_start_only,
                                       f.x_withspaces_end_only,
                                       g.x_backup_without_spp,
                                       h.x_species,
                                       i.x_trimmed_species) {

        # FIRST TRY FULLNAMES AND CODES ----
        # if only genus is available, return only genus

        if (all(!c(x[i], b.x_trimmed) %like_case% " ")) {
          found <- lookup(fullname_lower %in% c(h.x_species, i.x_trimmed_species),
                          haystack = data_to_check)
          if (!is.na(found)) {
            x[i] <- found[1L]
            return(x[i])
          }
          if (nchar(g.x_backup_without_spp) >= 6) {
            found <- lookup(fullname_lower %like_case% paste0("^", unregex(g.x_backup_without_spp), "[a-z]+"),
                            haystack = data_to_check)
            if (!is.na(found)) {
              x[i] <- found[1L]
              return(x[i])
            }
          }
          # rest of genus only is in allow_uncertain part.
        }

        # allow no codes less than 4 characters long, was already checked for WHONET earlier
        if (nchar(g.x_backup_without_spp) < 4) {
          x[i] <- lookup(mo == "UNKNOWN")
          if (initial_search == TRUE) {
            failures <- c(failures, a.x_backup)
          }
          return(x[i])
        }

        # try probable: trimmed version of fullname ----
        found <- lookup(fullname_lower %in% tolower(g.x_backup_without_spp),
                        haystack = data_to_check)
        if (!is.na(found)) {
          return(found[1L])
        }

        # try any match keeping spaces ----
        if (nchar(g.x_backup_without_spp) >= 6) {
          found <- lookup(fullname_lower %like_case% d.x_withspaces_start_end,
                          haystack = data_to_check)
          if (!is.na(found)) {
            return(found[1L])
          }
        }

        # try any match keeping spaces, not ending with $ ----
        found <- lookup(fullname_lower %like_case% paste0(trimws(e.x_withspaces_start_only), " "),
                        haystack = data_to_check)
        if (!is.na(found)) {
          return(found[1L])
        }
        if (nchar(g.x_backup_without_spp) >= 6) {
          found <- lookup(fullname_lower %like_case% e.x_withspaces_start_only,
                          haystack = data_to_check)
          if (!is.na(found)) {
            return(found[1L])
          }
        }

        # try any match keeping spaces, not start with ^ ----
        found <- lookup(fullname_lower %like_case% paste0(" ", trimws(f.x_withspaces_end_only)),
                        haystack = data_to_check)
        if (!is.na(found)) {
          return(found[1L])
        }

        # try a trimmed version
        if (nchar(g.x_backup_without_spp) >= 6) {
          found <- lookup(fullname_lower %like_case% b.x_trimmed |
                            fullname_lower %like_case% c.x_trimmed_without_group,
                          haystack = data_to_check)
          if (!is.na(found)) {
            return(found[1L])
          }
        }
        
        
        # try splitting of characters in the middle and then find ID ----
        # only when text length is 6 or lower
        # like esco = E. coli, klpn = K. pneumoniae, stau = S. aureus, staaur = S. aureus
        if (nchar(g.x_backup_without_spp) <= 6) {
          x_length <- nchar(g.x_backup_without_spp)
          x_split <- paste0("^",
                            g.x_backup_without_spp %pm>% substr(1, x_length / 2),
                            ".* ",
                            g.x_backup_without_spp %pm>% substr((x_length / 2) + 1, x_length))
          found <- lookup(fullname_lower %like_case% x_split,
                          haystack = data_to_check)
          if (!is.na(found)) {
            return(found[1L])
          }
        }

        # try fullname without start and without nchar limit of >= 6 ----
        # like "K. pneu rhino" >> "Klebsiella pneumoniae (rhinoscleromatis)" = KLEPNERH
        found <- lookup(fullname_lower %like_case% e.x_withspaces_start_only,
                        haystack = data_to_check)
        if (!is.na(found)) {
          return(found[1L])
        }

        # MISCELLANEOUS ----

        # look for old taxonomic names ----
        found <- lookup(fullname_lower %like_case% e.x_withspaces_start_only,
                        column = NULL, # all columns
                        haystack = data.old_to_check)
        if (!all(is.na(found))) {
          # when property is "ref" (which is the case in mo_ref, mo_authors and mo_year), return the old value, so:
          # mo_ref() of "Chlamydia psittaci" will be "Page, 1968" (with warning)
          # mo_ref() of "Chlamydophila psittaci" will be "Everett et al., 1999"
          if (property == "ref") {
            x[i] <- found["ref"]
          } else {
            x[i] <- lookup(fullname == found["fullname_new"], haystack = MO_lookup)
          }
          pkg_env$mo_renamed_last_run <- found["fullname"]
          was_renamed(name_old = found["fullname"],
                      name_new = lookup(fullname == found["fullname_new"], "fullname", haystack = MO_lookup),
                      ref_old = found["ref"],
                      ref_new = lookup(fullname == found["fullname_new"], "ref", haystack = MO_lookup),
                      mo = lookup(fullname == found["fullname_new"], "mo", haystack = MO_lookup))
          return(x[i])
        }

        # check for uncertain results ----
        uncertain_fn <- function(a.x_backup,
                                 b.x_trimmed,
                                 d.x_withspaces_start_end,
                                 e.x_withspaces_start_only,
                                 f.x_withspaces_end_only,
                                 g.x_backup_without_spp,
                                 uncertain.reference_data_to_use) {

          if (uncertainty_level == 0) {
            # do not allow uncertainties
            return(NA_character_)
          }

          # UNCERTAINTY LEVEL 1 ----
          if (uncertainty_level >= 1) {
            now_checks_for_uncertainty_level <- 1

            # (1) look again for old taxonomic names, now for G. species ----
            if (isTRUE(debug)) {
              cat(font_bold("\n[ UNCERTAINTY LEVEL", now_checks_for_uncertainty_level, "] (1) look again for old taxonomic names, now for G. species\n"))
            }
            if (isTRUE(debug)) {
              message("Running '", d.x_withspaces_start_end, "' and '", e.x_withspaces_start_only, "'")
            }
            found <- lookup(fullname_lower %like_case% d.x_withspaces_start_end |
                              fullname_lower %like_case% e.x_withspaces_start_only,
                            column = NULL, # all columns
                            haystack = data.old_to_check)
            if (!all(is.na(found)) & nchar(g.x_backup_without_spp) >= 6) {
              if (property == "ref") {
                # when property is "ref" (which is the case in mo_ref, mo_authors and mo_year), return the old value, so:
                # mo_ref("Chlamydia psittaci") = "Page, 1968" (with warning)
                # mo_ref("Chlamydophila psittaci") = "Everett et al., 1999"
                x <- found["ref"]
              } else {
                x <- lookup(fullname == found["fullname_new"], haystack = MO_lookup)
              }
              was_renamed(name_old = found["fullname"],
                          name_new = lookup(fullname == found["fullname_new"], "fullname", haystack = MO_lookup),
                          ref_old = found["ref"],
                          ref_new = lookup(fullname == found["fullname_new"], "ref", haystack = MO_lookup),
                          mo = lookup(fullname == found["fullname_new"], "mo", haystack = MO_lookup))
              pkg_env$mo_renamed_last_run <- found["fullname"]
              uncertainties <<- rbind(uncertainties,
                                      format_uncertainty_as_df(uncertainty_level = now_checks_for_uncertainty_level,
                                                               input = a.x_backup,
                                                               result_mo = lookup(fullname == found["fullname_new"], "mo", haystack = MO_lookup)),
                                      stringsAsFactors = FALSE)
              return(x)
            }

            # (2) Try with misspelled input ----
            # just rerun with dyslexia_mode = TRUE will used the extensive regex part above
            if (isTRUE(debug)) {
              cat(font_bold("\n[ UNCERTAINTY LEVEL", now_checks_for_uncertainty_level, "] (2) Try with misspelled input\n"))
            }
            if (isTRUE(debug)) {
              message("Running '", a.x_backup, "'")
            }
            # first try without dyslexia mode
            found <- suppressMessages(suppressWarnings(exec_as.mo(a.x_backup, initial_search = FALSE, dyslexia_mode = FALSE, allow_uncertain = FALSE, debug = debug, reference_data_to_use = uncertain.reference_data_to_use, actual_uncertainty = 1, actual_input = a.x_backup)))
            if (empty_result(found)) {
              # then with dyslexia mode
              found <- suppressMessages(suppressWarnings(exec_as.mo(a.x_backup, initial_search = FALSE, dyslexia_mode = TRUE, allow_uncertain = FALSE, debug = debug, reference_data_to_use = uncertain.reference_data_to_use, actual_uncertainty = 1, actual_input = a.x_backup)))
            }
            if (!empty_result(found)) {
              found_result <- found
              uncertainties <<- rbind(uncertainties,
                                      attr(found, which = "uncertainties", exact = TRUE),
                                      stringsAsFactors = FALSE)
              found <- lookup(mo == found)
              return(found)
            }
          }

          # UNCERTAINTY LEVEL 2 ----
          if (uncertainty_level >= 2) {
            now_checks_for_uncertainty_level <- 2

            # (3) look for genus only, part of name ----
            if (isTRUE(debug)) {
              cat(font_bold("\n[ UNCERTAINTY LEVEL", now_checks_for_uncertainty_level, "] (3) look for genus only, part of name\n"))
            }
            if (nchar(g.x_backup_without_spp) > 4 & !b.x_trimmed %like_case% " ") {
              if (!grepl("^[A-Z][a-z]+", b.x_trimmed, ignore.case = FALSE)) {
                if (isTRUE(debug)) {
                  message("Running '", paste(b.x_trimmed, "species"), "'")
                }
                # not when input is like Genustext, because then Neospora would lead to Actinokineospora
                found <- lookup(fullname_lower %like_case% paste(b.x_trimmed, "species"),
                                haystack = uncertain.reference_data_to_use)
                if (!is.na(found)) {
                  found_result <- found
                  found <- lookup(mo == found)
                  uncertainties <<- rbind(uncertainties,
                                          format_uncertainty_as_df(uncertainty_level = now_checks_for_uncertainty_level,
                                                                   input = a.x_backup,
                                                                   result_mo = found_result),
                                          stringsAsFactors = FALSE)
                  return(found)
                }
              }
            }

            # (4) strip values between brackets ----
            if (isTRUE(debug)) {
              cat(font_bold("\n[ UNCERTAINTY LEVEL", now_checks_for_uncertainty_level, "] (4) strip values between brackets\n"))
            }
            a.x_backup_stripped <- gsub("( *[(].*[)] *)", " ", a.x_backup, perl = TRUE)
            a.x_backup_stripped <- trimws(gsub(" +", " ", a.x_backup_stripped, perl = TRUE))
            if (isTRUE(debug)) {
              message("Running '", a.x_backup_stripped, "'")
            }
            # first try without dyslexia mode
            found <- suppressMessages(suppressWarnings(exec_as.mo(a.x_backup_stripped, initial_search = FALSE, dyslexia_mode = FALSE, allow_uncertain = FALSE, debug = debug, reference_data_to_use = uncertain.reference_data_to_use, actual_uncertainty = 2, actual_input = a.x_backup)))
            if (empty_result(found)) {
              # then with dyslexia mode
              found <- suppressMessages(suppressWarnings(exec_as.mo(a.x_backup_stripped, initial_search = FALSE, dyslexia_mode = TRUE, allow_uncertain = FALSE, debug = debug, reference_data_to_use = uncertain.reference_data_to_use, actual_uncertainty = 2, actual_input = a.x_backup)))
            }
            if (!empty_result(found) & nchar(g.x_backup_without_spp) >= 6) {
              found_result <- found
              uncertainties <<- rbind(uncertainties,
                                      attr(found, which = "uncertainties", exact = TRUE),
                                      stringsAsFactors = FALSE)
              found <- lookup(mo == found)
              return(found)
            }

            # (5) inverse input ----
            if (isTRUE(debug)) {
              cat(font_bold("\n[ UNCERTAINTY LEVEL", now_checks_for_uncertainty_level, "] (5) inverse input\n"))
            }
            a.x_backup_inversed <- paste(rev(unlist(strsplit(a.x_backup, split = " "))), collapse = " ")
            if (isTRUE(debug)) {
              message("Running '", a.x_backup_inversed, "'")
            }

            # first try without dyslexia mode
            found <- suppressMessages(suppressWarnings(exec_as.mo(a.x_backup_inversed, initial_search = FALSE, dyslexia_mode = FALSE, allow_uncertain = FALSE, debug = debug, reference_data_to_use = uncertain.reference_data_to_use, actual_uncertainty = 2, actual_input = a.x_backup)))
            if (empty_result(found)) {
              # then with dyslexia mode
              found <- suppressMessages(suppressWarnings(exec_as.mo(a.x_backup_inversed, initial_search = FALSE, dyslexia_mode = TRUE, allow_uncertain = FALSE, debug = debug, reference_data_to_use = uncertain.reference_data_to_use, actual_uncertainty = 2, actual_input = a.x_backup)))
            }
            if (!empty_result(found) & nchar(g.x_backup_without_spp) >= 6) {
              found_result <- found
              uncertainties <<- rbind(uncertainties,
                                      attr(found, which = "uncertainties", exact = TRUE),
                                      stringsAsFactors = FALSE)
              found <- lookup(mo == found)
              return(found)
            }

            # (6) try to strip off half an element from end and check the remains ----
            if (isTRUE(debug)) {
              cat(font_bold("\n[ UNCERTAINTY LEVEL", now_checks_for_uncertainty_level, "] (6) try to strip off half an element from end and check the remains\n"))
            }
            x_strip <- a.x_backup %pm>% strsplit("[ .]") %pm>% unlist()
            if (length(x_strip) > 1) {
              for (i in seq_len(length(x_strip) - 1)) {
                lastword <- x_strip[length(x_strip) - i + 1]
                lastword_half <- substr(lastword, 1, as.integer(nchar(lastword) / 2))
                # remove last half of the second term
                x_strip_collapsed <- paste(c(x_strip[seq_len(length(x_strip) - i)], lastword_half), collapse = " ")
                if (nchar(x_strip_collapsed) >= 4 & nchar(lastword_half) > 2) {
                  if (isTRUE(debug)) {
                    message("Running '", x_strip_collapsed, "'")
                  }
                  # first try without dyslexia mode
                  found <- suppressMessages(suppressWarnings(exec_as.mo(x_strip_collapsed, initial_search = FALSE, dyslexia_mode = FALSE, allow_uncertain = FALSE, debug = debug, reference_data_to_use = uncertain.reference_data_to_use, actual_uncertainty = 2, actual_input = a.x_backup)))
                  if (empty_result(found)) {
                    # then with dyslexia mode
                    found <- suppressMessages(suppressWarnings(exec_as.mo(x_strip_collapsed, initial_search = FALSE, dyslexia_mode = TRUE, allow_uncertain = FALSE, debug = debug, reference_data_to_use = uncertain.reference_data_to_use, actual_uncertainty = 2, actual_input = a.x_backup)))
                  }
                  if (!empty_result(found)) {
                    found_result <- found
                    uncertainties <<- rbind(uncertainties,
                                            attr(found, which = "uncertainties", exact = TRUE),
                                            stringsAsFactors = FALSE)
                    found <- lookup(mo == found)
                    return(found)
                  }
                }
              }
            }
            # (7) try to strip off one element from end and check the remains ----
            if (isTRUE(debug)) {
              cat(font_bold("\n[ UNCERTAINTY LEVEL", now_checks_for_uncertainty_level, "] (7) try to strip off one element from end and check the remains\n"))
            }
            if (length(x_strip) > 1) {
              for (i in seq_len(length(x_strip) - 1)) {
                x_strip_collapsed <- paste(x_strip[seq_len(length(x_strip) - i)], collapse = " ")
                if (nchar(x_strip_collapsed) >= 6) {
                  if (isTRUE(debug)) {
                    message("Running '", x_strip_collapsed, "'")
                  }
                  # first try without dyslexia mode
                  found <- suppressMessages(suppressWarnings(exec_as.mo(x_strip_collapsed, initial_search = FALSE, dyslexia_mode = FALSE, allow_uncertain = FALSE, debug = debug, reference_data_to_use = uncertain.reference_data_to_use, actual_uncertainty = 2, actual_input = a.x_backup)))
                  if (empty_result(found)) {
                    # then with dyslexia mode
                    found <- suppressMessages(suppressWarnings(exec_as.mo(x_strip_collapsed, initial_search = FALSE, dyslexia_mode = TRUE, allow_uncertain = FALSE, debug = debug, reference_data_to_use = uncertain.reference_data_to_use, actual_uncertainty = 2, actual_input = a.x_backup)))
                  }

                  if (!empty_result(found)) {
                    found_result <- found
                    uncertainties <<- rbind(uncertainties,
                                            attr(found, which = "uncertainties", exact = TRUE),
                                            stringsAsFactors = FALSE)
                    found <- lookup(mo == found)
                    return(found)
                  }
                }
              }
            }
            # (8) check for unknown yeasts/fungi ----
            if (isTRUE(debug)) {
              cat(font_bold("\n[ UNCERTAINTY LEVEL", now_checks_for_uncertainty_level, "] (8) check for unknown yeasts/fungi\n"))
            }
            if (b.x_trimmed %like_case% "yeast") {
              found <- "F_YEAST"
              found_result <- found
              found <- lookup(mo == found)
              uncertainties <<- rbind(uncertainties,
                                      format_uncertainty_as_df(uncertainty_level = now_checks_for_uncertainty_level,
                                                               input = a.x_backup,
                                                               result_mo = found_result),
                                      stringsAsFactors = FALSE)
              return(found)
            }
            if (b.x_trimmed %like_case% "(fungus|fungi)" & !b.x_trimmed %like_case% "fungiphrya") {
              found <- "F_FUNGUS"
              found_result <- found
              found <- lookup(mo == found)
              uncertainties <<- rbind(uncertainties,
                                      format_uncertainty_as_df(uncertainty_level = now_checks_for_uncertainty_level,
                                                               input = a.x_backup,
                                                               result_mo = found_result),
                                      stringsAsFactors = FALSE)
              return(found)
            }
            # (9) try to strip off one element from start and check the remains (only allow >= 2-part name outcome) ----
            if (isTRUE(debug)) {
              cat(font_bold("\n[ UNCERTAINTY LEVEL", now_checks_for_uncertainty_level, "] (9) try to strip off one element from start and check the remains (only allow >= 2-part name outcome)\n"))
            }
            x_strip <- a.x_backup %pm>% strsplit("[ .]") %pm>% unlist()
            if (length(x_strip) > 1 & nchar(g.x_backup_without_spp) >= 6) {
              for (i in 2:(length(x_strip))) {
                x_strip_collapsed <- paste(x_strip[i:length(x_strip)], collapse = " ")
                if (isTRUE(debug)) {
                  message("Running '", x_strip_collapsed, "'")
                }
                # first try without dyslexia mode
                found <- suppressMessages(suppressWarnings(exec_as.mo(x_strip_collapsed, initial_search = FALSE, dyslexia_mode = FALSE, allow_uncertain = FALSE, debug = debug, reference_data_to_use = uncertain.reference_data_to_use, actual_uncertainty = 2, actual_input = a.x_backup)))
                if (empty_result(found)) {
                  # then with dyslexia mode
                  found <- suppressMessages(suppressWarnings(exec_as.mo(x_strip_collapsed, initial_search = FALSE, dyslexia_mode = TRUE, allow_uncertain = FALSE, debug = debug, reference_data_to_use = uncertain.reference_data_to_use, actual_uncertainty = 2, actual_input = a.x_backup)))
                }
                if (!empty_result(found)) {
                  found_result <- found
                  # uncertainty level 2 only if searched part contains a space (otherwise it will be found with lvl 3)
                  if (x_strip_collapsed %like_case% " ") {
                    uncertainties <<- rbind(uncertainties,
                                            attr(found, which = "uncertainties", exact = TRUE),
                                            stringsAsFactors = FALSE)
                    found <- lookup(mo == found)
                    return(found)
                  }
                }
              }
            }
          }

          # UNCERTAINTY LEVEL 3 ----
          if (uncertainty_level >= 3) {
            now_checks_for_uncertainty_level <- 3

            # (10) try to strip off one element from start and check the remains (any text size) ----
            if (isTRUE(debug)) {
              cat(font_bold("\n[ UNCERTAINTY LEVEL", now_checks_for_uncertainty_level, "] (10) try to strip off one element from start and check the remains (any text size)\n"))
            }
            x_strip <- a.x_backup %pm>% strsplit("[ .]") %pm>% unlist()
            if (length(x_strip) > 1 & nchar(g.x_backup_without_spp) >= 6) {
              for (i in 2:(length(x_strip))) {
                x_strip_collapsed <- paste(x_strip[i:length(x_strip)], collapse = " ")
                if (isTRUE(debug)) {
                  message("Running '", x_strip_collapsed, "'")
                }
                # first try without dyslexia mode
                found <- suppressMessages(suppressWarnings(exec_as.mo(x_strip_collapsed, initial_search = FALSE, dyslexia_mode = FALSE, allow_uncertain = FALSE, debug = debug, reference_data_to_use = uncertain.reference_data_to_use, actual_uncertainty = 3, actual_input = a.x_backup)))
                if (empty_result(found)) {
                  # then with dyslexia mode
                  found <- suppressMessages(suppressWarnings(exec_as.mo(x_strip_collapsed, initial_search = FALSE, dyslexia_mode = TRUE, allow_uncertain = FALSE, debug = debug, reference_data_to_use = uncertain.reference_data_to_use, actual_uncertainty = 3, actual_input = a.x_backup)))
                }
                if (!empty_result(found)) {
                  found_result <- found
                  uncertainties <<- rbind(uncertainties,
                                          attr(found, which = "uncertainties", exact = TRUE),
                                          stringsAsFactors = FALSE)
                  found <- lookup(mo == found)
                  return(found)
                }
              }
            }
            # (11) try to strip off one element from end and check the remains (any text size) ----
            # (this is in fact 7 but without nchar limit of >=6)
            if (isTRUE(debug)) {
              cat(font_bold("\n[ UNCERTAINTY LEVEL", now_checks_for_uncertainty_level, "] (11) try to strip off one element from end and check the remains (any text size)\n"))
            }
            if (length(x_strip) > 1) {
              for (i in seq_len(length(x_strip) - 1)) {
                x_strip_collapsed <- paste(x_strip[seq_len(length(x_strip) - i)], collapse = " ")
                if (isTRUE(debug)) {
                  message("Running '", x_strip_collapsed, "'")
                }
                # first try without dyslexia mode
                found <- suppressMessages(suppressWarnings(exec_as.mo(x_strip_collapsed, initial_search = FALSE, dyslexia_mode = FALSE, allow_uncertain = FALSE, debug = debug, reference_data_to_use = uncertain.reference_data_to_use, actual_uncertainty = 3, actual_input = a.x_backup)))
                if (empty_result(found)) {
                  # then with dyslexia mode
                  found <- suppressMessages(suppressWarnings(exec_as.mo(x_strip_collapsed, initial_search = FALSE, dyslexia_mode = TRUE, allow_uncertain = FALSE, debug = debug, reference_data_to_use = uncertain.reference_data_to_use, actual_uncertainty = 3, actual_input = a.x_backup)))
                }
                if (!empty_result(found)) {
                  found_result <- found
                  uncertainties <<- rbind(uncertainties,
                                          attr(found, which = "uncertainties", exact = TRUE),
                                          stringsAsFactors = FALSE)
                  found <- lookup(mo == found)
                  return(found)
                }
              }
            }

            # (12) part of a name (very unlikely match) ----
            if (isTRUE(debug)) {
              cat(font_bold("\n[ UNCERTAINTY LEVEL", now_checks_for_uncertainty_level, "] (12) part of a name (very unlikely match)\n"))
            }
            if (isTRUE(debug)) {
              message("Running '", f.x_withspaces_end_only, "'")
            }
            if (nchar(g.x_backup_without_spp) >= 6) {
              found <- lookup(fullname_lower %like_case% f.x_withspaces_end_only, column = "mo")
              if (!is.na(found)) {
                found_result <- lookup(mo == found)
                uncertainties <<- rbind(uncertainties,
                                        attr(found, which = "uncertainties", exact = TRUE),
                                        stringsAsFactors = FALSE)
                found <- lookup(mo == found)
                return(found)
              }
            }
          }


          # didn't found in uncertain results too
          return(NA_character_)
        }

        # uncertain results
        x[i] <- uncertain_fn(a.x_backup = a.x_backup,
                             b.x_trimmed = b.x_trimmed,
                             d.x_withspaces_start_end = d.x_withspaces_start_end,
                             e.x_withspaces_start_only = e.x_withspaces_start_only,
                             f.x_withspaces_end_only = f.x_withspaces_end_only,
                             g.x_backup_without_spp = g.x_backup_without_spp,
                             uncertain.reference_data_to_use = MO_lookup) # MO_lookup[which(MO_lookup$prevalence %in% c(1, 2)), ])
        if (!empty_result(x[i])) {
          return(x[i])
        }
        # x[i] <- uncertain_fn(a.x_backup = a.x_backup,
        #                      b.x_trimmed = b.x_trimmed,
        #                      d.x_withspaces_start_end = d.x_withspaces_start_end,
        #                      e.x_withspaces_start_only = e.x_withspaces_start_only,
        #                      f.x_withspaces_end_only = f.x_withspaces_end_only,
        #                      g.x_backup_without_spp = g.x_backup_without_spp,
        #                      uncertain.reference_data_to_use = MO_lookup[which(MO_lookup$prevalence == 3), ])
        # if (!empty_result(x[i])) {
        #   return(x[i])
        # }

        # didn't found any
        return(NA_character_)
      }

      # CHECK ALL IN ONE GO ----
      x[i] <- check_per_prevalence(data_to_check = MO_lookup,
                                   data.old_to_check = MO.old_lookup,
                                   a.x_backup = x_backup[i],
                                   b.x_trimmed = x_trimmed[i],
                                   c.x_trimmed_without_group = x_trimmed_without_group[i],
                                   d.x_withspaces_start_end = x_withspaces_start_end[i],
                                   e.x_withspaces_start_only = x_withspaces_start_only[i],
                                   f.x_withspaces_end_only = x_withspaces_end_only[i],
                                   g.x_backup_without_spp = x_backup_without_spp[i],
                                   h.x_species = x_species[i],
                                   i.x_trimmed_species = x_trimmed_species[i])
      if (!empty_result(x[i])) {
        next
      }


      # no results found: make them UNKNOWN ----
      x[i] <- lookup(mo == "UNKNOWN", uncertainty = -1)
      if (initial_search == TRUE) {
        failures <- c(failures, x_backup[i])
      }
    }

    if (initial_search == TRUE) {
      close(progress)
    }
  }

  # handling failures ----
  failures <- failures[!failures %in% c(NA, NULL, NaN)]
  if (length(failures) > 0 & initial_search == TRUE) {
    pkg_env$mo_failures <- sort(unique(failures))
    plural <- c("value", "it", "was")
    if (pm_n_distinct(failures) > 1) {
      plural <- c("values", "them", "were")
    }
    x_input_clean <- trimws2(x_input)
    total_failures <- length(x_input_clean[as.character(x_input_clean) %in% as.character(failures) & !x_input %in% c(NA, NULL, NaN)])
    total_n <- length(x_input[!x_input %in% c(NA, NULL, NaN)])
    msg <- paste0(nr2char(pm_n_distinct(failures)), " unique ", plural[1],
                  " (covering ", percentage(total_failures / total_n),
                  ") could not be coerced and ", plural[3], " considered 'unknown'")
    if (pm_n_distinct(failures) <= 10) {
      msg <- paste0(msg, ": ", paste('"', unique(failures), '"', sep = "", collapse = ", "))
    }
    msg <- paste0(msg,
                  ".\nUse mo_failures() to review ", plural[2], ". Edit the `allow_uncertain` argument if needed (see ?as.mo).\n",
                  "You can also use your own reference data, e.g.:\n",
                  '  as.mo("mycode", reference_df = data.frame(own = "mycode", mo = "', MO_lookup$mo[match("Escherichia coli", MO_lookup$fullname)], '"))\n',
                  '  mo_name("mycode", reference_df = data.frame(own = "mycode", mo = "', MO_lookup$mo[match("Escherichia coli", MO_lookup$fullname)], '"))\n')
    warning_(paste0("\n", msg),
             add_fn = font_red,
             call = FALSE,
             immediate = TRUE) # thus will always be shown, even if >= warnings
  }
  # handling uncertainties ----
  if (NROW(uncertainties) > 0 & initial_search == TRUE) {
    uncertainties <- as.list(pm_distinct(uncertainties, input, .keep_all = TRUE))
    pkg_env$mo_uncertainties <- uncertainties

    plural <- c("", "it", "was")
    if (length(uncertainties$input) > 1) {
      plural <- c("s", "them", "were")
    }
    msg <- paste0("Translation to ", nr2char(length(uncertainties$input)), " microorganism", plural[1],
                  " ", plural[3], " guessed with uncertainty. Use mo_uncertainties() to review ", plural[2], ".")
    message_(msg)
  }

  # Becker ----
  if (Becker == TRUE | Becker == "all") {
    # warn when species found that are not in:
    # - Becker et al. 2014, PMID 25278577
    # - Becker et al. 2019, PMID 30872103
    # - Becker et al. 2020, PMID 32056452
    post_Becker <- character(0) # 2020-10-20 currently all are mentioned in above papers (otherwise uncomment below)
    
    # nolint start
    # if (any(x %in% MO_lookup[which(MO_lookup$species %in% post_Becker), property])) {
    #   warning_("Becker ", font_italic("et al."), " (2014, 2019) does not contain these species named after their publication: ",
    #            font_italic(paste("S.",
    #                              sort(mo_species(unique(x[x %in% MO_lookup[which(MO_lookup$species %in% post_Becker), property]]))),
    #                              collapse = ", ")),
    #            ".",
    #            call = FALSE,
    #            immediate = TRUE)
    # }
    # nolint end

    # 'MO_CONS' and 'MO_COPS' are <mo> vectors created in R/zzz.R
    CoNS <- MO_lookup[which(MO_lookup$mo %in% MO_CONS), property, drop = TRUE]
    x[x %in% CoNS] <- lookup(mo == "B_STPHY_CONS", uncertainty = -1)

    CoPS <- MO_lookup[which(MO_lookup$mo %in% MO_COPS), property, drop = TRUE]
    x[x %in% CoPS] <- lookup(mo == "B_STPHY_COPS", uncertainty = -1)

    if (Becker == "all") {
      x[x %in% lookup(fullname %like_case% "^Staphylococcus aureus", n = Inf)] <- lookup(mo == "B_STPHY_COPS", uncertainty = -1)
    }
  }

  # Lancefield ----
  if (Lancefield == TRUE | Lancefield == "all") {
    # group A - S. pyogenes
    x[x %in% lookup(genus == "Streptococcus" & species == "pyogenes", n = Inf)] <- lookup(fullname == "Streptococcus group A", uncertainty = -1)
    # group B - S. agalactiae
    x[x %in% lookup(genus == "Streptococcus" & species == "agalactiae", n = Inf)] <- lookup(fullname == "Streptococcus group B", uncertainty = -1)
    # group C
    x[x %in% lookup(genus == "Streptococcus" &
                      species %in% c("equisimilis", "equi", "zooepidemicus", "dysgalactiae"),
                    n = Inf)] <- lookup(fullname == "Streptococcus group C", uncertainty = -1)
    if (Lancefield == "all") {
      # all Enterococci
      x[x %in% lookup(genus == "Enterococcus", n = Inf)] <- lookup(fullname == "Streptococcus group D", uncertainty = -1)
    }
    # group F - S. anginosus
    x[x %in% lookup(genus == "Streptococcus" & species == "anginosus", n = Inf)] <- lookup(fullname == "Streptococcus group F", uncertainty = -1)
    # group H - S. sanguinis
    x[x %in% lookup(genus == "Streptococcus" & species == "sanguinis", n = Inf)] <- lookup(fullname == "Streptococcus group H", uncertainty = -1)
    # group K - S. salivarius
    x[x %in% lookup(genus == "Streptococcus" & species == "salivarius", n = Inf)] <- lookup(fullname == "Streptococcus group K", uncertainty = -1)
  }

  # Wrap up ----------------------------------------------------------------

  # comply to x, which is also unique and without empty values
  x_input_unique_nonempty <- unique(x_input[!is.na(x_input)
                                            & !is.null(x_input)
                                            & !identical(x_input, "")
                                            & !identical(x_input, "xxx")])

  # left join the found results to the original input values (x_input)
  df_found <- data.frame(input = as.character(x_input_unique_nonempty),
                         found = as.character(x),
                         stringsAsFactors = FALSE)
  df_input <- data.frame(input = as.character(x_input),
                         stringsAsFactors = FALSE)

  # super fast using match() which is a lot faster than merge()
  x <- df_found$found[match(df_input$input, df_found$input)]

  if (property == "mo") {
    x <- set_clean_class(x, new_class = c("mo", "character"))
  }

  if (length(mo_renamed()) > 0) {
    print(mo_renamed())
  }

  if (initial_search == FALSE) {
    # we got here from uncertain_fn().
    if (NROW(uncertainties) == 0) {
      # the stripped/transformed version of x_backup is apparently a full hit, like with: as.mo("Escherichia (hello there) coli")
      uncertainties <- rbind(uncertainties,
                             format_uncertainty_as_df(uncertainty_level = actual_uncertainty,
                                                      input = actual_input,
                                                      result_mo = x,
                                                      candidates = ""),
                             stringsAsFactors = FALSE)
    }
    # this will save the uncertain items as attribute, so they can be bound to `uncertainties` in the uncertain_fn() function
    x <- structure(x, uncertainties = uncertainties)
  } else {
    # keep track of time - give some hints to improve speed if it takes a long time
    end_time <- Sys.time()
    delta_time <- difftime(end_time, start_time, units = "secs")
    if (delta_time >= 30) {
      message_("Using `as.mo()` took ", delta_time, " seconds, which is a long time. Some suggestions to improve speed include:")
      message_(word_wrap("- Try to use as many valid taxonomic names as possible for your input.",
                         extra_indent = 2),
               as_note = FALSE)
      message_(word_wrap("- Save the output and use it as input for future calculations, e.g. create a new variable to your data using `as.mo()`. All functions in this package that rely on microorganism codes will automatically use that new column where possible. All `mo_*()` functions also do not require you to set their `x` argument as long as you have the dplyr package installed and you have a column of class <mo>.",
                         extra_indent = 2),
               as_note = FALSE)
      message_(word_wrap("- Use `set_mo_source()` to continually transform your organisation codes to microorganisms codes used by this package, please see `?mo_source`.",
                         extra_indent = 2),
               as_note = FALSE)
    }
  }
  
  x
}

empty_result <- function(x) {
  all(x %in% c(NA, "UNKNOWN"))
}

was_renamed <- function(name_old, name_new, ref_old = "", ref_new = "", mo = "") {
  newly_set <- data.frame(old_name = name_old,
                          old_ref = ref_old,
                          new_name = name_new,
                          new_ref = ref_new,
                          mo = mo,
                          stringsAsFactors = FALSE)
  already_set <- pkg_env$mo_renamed
  if (!is.null(already_set)) {
    pkg_env$mo_renamed = rbind(already_set,
                               newly_set,
                               stringsAsFactors = FALSE)
  } else {
    pkg_env$mo_renamed <- newly_set
  }
}

format_uncertainty_as_df <- function(uncertainty_level,
                                     input,
                                     result_mo,
                                     candidates = NULL) {
  if (!is.null(pkg_env$mo_renamed_last_run)) {
    fullname <- pkg_env$mo_renamed_last_run
    pkg_env$mo_renamed_last_run <- NULL
    renamed_to <- MO_lookup[match(result_mo, MO_lookup$mo), "fullname", drop = TRUE][1]
  } else {
    fullname <- MO_lookup[match(result_mo, MO_lookup$mo), "fullname", drop = TRUE][1]
    renamed_to <- NA_character_
  }
  data.frame(uncertainty = uncertainty_level,
             input = input,
             fullname = fullname,
             renamed_to = renamed_to,
             mo = result_mo,
             # save max 26 entries: the one to be chosen and 25 more
             candidates = if (length(candidates) > 1) paste(candidates[c(2:min(26, length(candidates)))], collapse = ", ") else "",
             stringsAsFactors = FALSE)
}

# will be exported using s3_register() in R/zzz.R
pillar_shaft.mo <- function(x, ...) {
  out <- format(x)
  # grey out the kingdom (part until first "_")
  out[!is.na(x)] <- gsub("^([A-Z]+_)(.*)", paste0(font_subtle("\\1"), "\\2"), out[!is.na(x)], perl = TRUE)
  # and grey out every _
  out[!is.na(x)] <- gsub("_", font_subtle("_"), out[!is.na(x)])

  # markup NA and UNKNOWN
  out[is.na(x)] <- font_na("  NA")
  out[x == "UNKNOWN"] <- font_na("  UNKNOWN")

  # make it always fit exactly
  max_char <- max(nchar(x))
  if (is.na(max_char)) {
    max_char <- 7
  }
  create_pillar_column(out,
                       align = "left",
                       width = max_char + ifelse(any(x %in% c(NA, "UNKNOWN")), 2, 0))
}

# will be exported using s3_register() in R/zzz.R
type_sum.mo <- function(x, ...) {
  "mo"
}

# will be exported using s3_register() in R/zzz.R
freq.mo <- function(x, ...) {
  x_noNA <- as.mo(x[!is.na(x)]) # as.mo() to get the newest mo codes
  grams <- mo_gramstain(x_noNA, language = NULL)
  digits <- list(...)$digits
  if (is.null(digits)) {
    digits <- 2
  }
  cleaner::freq.default(
    x = x,
    ...,
    .add_header = list(
      `Gram-negative` = paste0(
        format(sum(grams == "Gram-negative", na.rm = TRUE),
               big.mark = ",",
               decimal.mark = "."),
        " (", percentage(sum(grams == "Gram-negative", na.rm = TRUE) / length(grams),
                         digits = digits),
        ")"),
      `Gram-positive` = paste0(
        format(sum(grams == "Gram-positive", na.rm = TRUE),
               big.mark = ",",
               decimal.mark = "."),
        " (", percentage(sum(grams == "Gram-positive", na.rm = TRUE) / length(grams),
                         digits = digits),
        ")"),
      `Nr. of genera` = pm_n_distinct(mo_genus(x_noNA, language = NULL)),
      `Nr. of species` = pm_n_distinct(paste(mo_genus(x_noNA, language = NULL),
                                             mo_species(x_noNA, language = NULL)))))
}

# will be exported using s3_register() in R/zzz.R
get_skimmers.mo <- function(column) {
  skimr::sfl(
    skim_type = "mo",
    unique_total = ~pm_n_distinct(., na.rm = TRUE),
    gram_negative = ~sum(mo_is_gram_negative(stats::na.omit(.))),
    gram_positive = ~sum(mo_is_gram_positive(stats::na.omit(.))),
    top_genus = ~names(sort(-table(mo_genus(stats::na.omit(.), language = NULL))))[1L],
    top_species = ~names(sort(-table(mo_name(stats::na.omit(.), language = NULL))))[1L]
  )
}

#' @method print mo
#' @export
#' @noRd
print.mo <- function(x, print.shortnames = FALSE, ...) {
  cat("Class <mo>\n")
  x_names <- names(x)
  if (is.null(x_names) & print.shortnames == TRUE) {
    x_names <- tryCatch(mo_shortname(x, ...), error = function(e) NULL)
  }
  x <- as.character(x)
  names(x) <- x_names
  print.default(x, quote = FALSE)
}

#' @method summary mo
#' @export
#' @noRd
summary.mo <- function(object, ...) {
  # unique and top 1-3
  x <- as.mo(object) # force again, could be mo from older pkg version
  top <- as.data.frame(table(x), responseName = "n", stringsAsFactors = FALSE)
  top_3 <- top[order(-top$n), 1][1:3]
  value <- c("Class" = "mo",
             "<NA>" = length(x[is.na(x)]),
             "Unique" = pm_n_distinct(x[!is.na(x)]),
             "#1" = top_3[1],
             "#2" = top_3[2],
             "#3" = top_3[3])
  class(value) <- c("summaryDefault", "table")
  value
}

#' @method as.data.frame mo
#' @export
#' @noRd
as.data.frame.mo <- function(x, ...) {
  nm <- deparse1(substitute(x))
  if (!"nm" %in% names(list(...))) {
    as.data.frame.vector(as.mo(x), ..., nm = nm)
  } else {
    as.data.frame.vector(as.mo(x), ...)
  }
}

#' @method [ mo
#' @export
#' @noRd
"[.mo" <- function(x, ...) {
  y <- NextMethod()
  attributes(y) <- attributes(x)
  y
}
#' @method [[ mo
#' @export
#' @noRd
"[[.mo" <- function(x, ...) {
  y <- NextMethod()
  attributes(y) <- attributes(x)
  y
}
#' @method [<- mo
#' @export
#' @noRd
"[<-.mo" <- function(i, j, ..., value) {
  y <- NextMethod()
  attributes(y) <- attributes(i)
  # must only contain valid MOs
  class_integrity_check(y, "microorganism code", c(as.character(microorganisms$mo),
                                                   as.character(microorganisms.translation$mo_old)))
}
#' @method [[<- mo
#' @export
#' @noRd
"[[<-.mo" <- function(i, j, ..., value) {
  y <- NextMethod()
  attributes(y) <- attributes(i)
  # must only contain valid MOs
  class_integrity_check(y, "microorganism code", c(as.character(microorganisms$mo),
                                                   as.character(microorganisms.translation$mo_old)))
}
#' @method c mo
#' @export
#' @noRd
c.mo <- function(x, ...) {
  y <- NextMethod()
  attributes(y) <- attributes(x)
  # must only contain valid MOs
  class_integrity_check(y, "microorganism code", c(as.character(microorganisms$mo),
                                                   as.character(microorganisms.translation$mo_old)))
}

#' @method unique mo
#' @export
#' @noRd
unique.mo <- function(x, incomparables = FALSE, ...) {
  y <- NextMethod()
  attributes(y) <- attributes(x)
  y
}

#' @rdname as.mo
#' @export
mo_failures <- function() {
  pkg_env$mo_failures
}

#' @rdname as.mo
#' @export
mo_uncertainties <- function() {
  if (is.null(pkg_env$mo_uncertainties)) {
    return(NULL)
  }
  set_clean_class(as.data.frame(pkg_env$mo_uncertainties, 
                                stringsAsFactors = FALSE),
                  new_class = c("mo_uncertainties", "data.frame"))
}

#' @method print mo_uncertainties
#' @export
#' @noRd
print.mo_uncertainties <- function(x, ...) {
  if (NROW(x) == 0) {
    return(NULL)
  }
  message_("Matching scores are based on human pathogenic prevalence and the resemblance between the input and the full taxonomic name. Please see ?mo_matching_score.", as_note = FALSE)
  cat("\n")

  msg <- ""
  for (i in seq_len(nrow(x))) {
    if (x[i, ]$candidates != "") {
      candidates <- unlist(strsplit(x[i, ]$candidates, ", ", fixed = TRUE))
      scores <- mo_matching_score(x = x[i, ]$input, n = candidates)
      # sort on descending scores
      candidates <- candidates[order(1 - scores)]
      scores_formatted <- trimws(formatC(round(scores, 3), format = "f", digits = 3))
      n_candidates <- length(candidates)
      candidates <- paste0(candidates, " (", scores_formatted[order(1 - scores)], ")", collapse = ", ")
      # align with input after arrow
      candidates <- paste0("\n",
                           strwrap(paste0("Also matched",
                                          ifelse(n_candidates >= 25, " (max 25)", ""), ": ",
                                          candidates), # this is already max 25 due to format_uncertainty_as_df()
                                   indent = nchar(x[i, ]$input) + 6,
                                   exdent = nchar(x[i, ]$input) + 6,
                                   width = 0.98 * getOption("width")),
                           collapse = "")
      # after strwrap, make taxonomic names italic
      candidates <- gsub("([A-Za-z]+)", font_italic("\\1"), candidates)
      candidates <- gsub(paste(font_italic(c("Also", "matched"), collapse = NULL), collapse = " "),
                         "Also matched",
                         candidates, fixed = TRUE)
      candidates <- gsub(font_italic("max"), "max", candidates, fixed = TRUE)
    } else {
      candidates <- ""
    }
    score <- trimws(formatC(round(mo_matching_score(x = x[i, ]$input,
                                                    n = x[i, ]$fullname),
                                  3),
                            format = "f", digits = 3))
    msg <- paste(msg,
                 paste0(
                   strwrap(
                     paste0('"', x[i, ]$input, '" -> ',
                            paste0(font_bold(font_italic(x[i, ]$fullname)),
                                   ifelse(!is.na(x[i, ]$renamed_to), paste(", renamed to", font_italic(x[i, ]$renamed_to)), ""),
                                   " (", x[i, ]$mo,
                                   ", matching score = ", score,
                                   ") ")),
                     width = 0.98 * getOption("width"),
                     exdent = nchar(x[i, ]$input) + 6),
                   collapse = "\n"),
                 candidates,
                 sep = "\n")
    msg <- paste0(gsub("\n\n", "\n", msg), "\n\n")
  }
  cat(msg)
}

#' @rdname as.mo
#' @export
mo_renamed <- function() {
  items <- pkg_env$mo_renamed
  if (is.null(items)) {
    items <- data.frame(stringsAsFactors = FALSE)
  } else {
    items <- pm_distinct(items, old_name, .keep_all = TRUE)
  }
  set_clean_class(as.data.frame(items,
                                stringsAsFactors = FALSE),
                  new_class = c("mo_renamed", "data.frame"))
}

#' @method print mo_renamed
#' @export
#' @noRd
print.mo_renamed <- function(x, ...) {
  if (NROW(x) == 0) {
    return(invisible())
  }
  for (i in seq_len(nrow(x))) {
    message_(font_italic(x$old_name[i]),
             ifelse(x$old_ref[i] %in% c("", NA),
                    "",
                    paste0(" (",  gsub("et al.", font_italic("et al."), x$old_ref[i]), ")")),
             " was renamed ",
             ifelse(as.integer(gsub("[^0-9]", "", x$new_ref[i])) < as.integer(gsub("[^0-9]", "", x$old_ref[i])),
                    font_bold("back to "),
                    ""),
             font_italic(x$new_name[i]),
             ifelse(x$new_ref[i] %in% c("", NA), 
                    "",
                    paste0(" (",  gsub("et al.", font_italic("et al."), x$new_ref[i]), ")")),
             " [", x$mo[i], "]")
  }
}

nr2char <- function(x) {
  if (x %in% c(1:10)) {
    v <- c("one" = 1, "two" = 2, "three" = 3, "four" = 4, "five" = 5,
           "six" = 6, "seven" = 7, "eight" = 8, "nine" = 9, "ten" = 10)
    names(v[x])
  } else {
    x
  }
}

unregex <- function(x) {
  gsub("[^a-zA-Z0-9 -]", "", x)
}

translate_allow_uncertain <- function(allow_uncertain) {
  if (isTRUE(allow_uncertain)) {
    # default to uncertainty level 2
    allow_uncertain <- 2
  } else {
    allow_uncertain[tolower(allow_uncertain) == "none"] <- 0
    allow_uncertain[tolower(allow_uncertain) == "all"] <- 3
    allow_uncertain <- as.integer(allow_uncertain)
    stop_ifnot(allow_uncertain %in% c(0:3),
               '`allow_uncertain` must be a number between 0 (or "none") and 3 (or "all"), or TRUE (= 2) or FALSE (= 0)', call = FALSE)
  }
  allow_uncertain
}

get_mo_failures_uncertainties_renamed <- function() {
  remember <- list(failures = pkg_env$mo_failures,
                   uncertainties = pkg_env$mo_uncertainties,
                   renamed = pkg_env$mo_renamed)
  # empty them, otherwise mo_shortname("Chlamydophila psittaci") will give 3 notes
  pkg_env$mo_failures <- NULL
  pkg_env$mo_uncertainties <- NULL
  pkg_env$mo_renamed <- NULL
  remember
}

load_mo_failures_uncertainties_renamed <- function(metadata) {
  pkg_env$mo_failures <- metadata$failures
  pkg_env$mo_uncertainties <- metadata$uncertainties
  pkg_env$mo_renamed <- metadata$renamed
}

trimws2 <- function(x) {
  trimws(gsub("[\\s]+", " ", x, perl = TRUE))
}

parse_and_convert <- function(x) {
  tryCatch({
    if (!is.null(dim(x))) {
      if (NCOL(x) > 2) {
        stop("a maximum of two columns is allowed", call. = FALSE)
      } else if (NCOL(x) == 2) {
        # support Tidyverse selection like: df %pm>% select(colA, colB)
        # paste these columns together
        x <- as.data.frame(x, stringsAsFactors = FALSE)
        colnames(x) <- c("A", "B")
        x <- paste(x$A, x$B)
      } else {
        # support Tidyverse selection like: df %pm>% select(colA)
        x <- as.data.frame(x, stringsAsFactors = FALSE)[[1]]
      }
    }
    x[is.null(x)] <- NA
    parsed <- iconv(x, to = "UTF-8")
    parsed[is.na(parsed) & !is.na(x)] <- iconv(x[is.na(parsed) & !is.na(x)], from = "Latin1", to = "ASCII//TRANSLIT")
    parsed <- gsub('"', "", parsed, fixed = TRUE)
  }, error = function(e) stop(e$message, call. = FALSE)) # this will also be thrown when running `as.mo(no_existing_object)`
  parsed
}

replace_old_mo_codes <- function(x, property) {
  if (any(toupper(x) %in% microorganisms.translation$mo_old, na.rm = TRUE)) {
    # get the ones that match
    matched <- match(toupper(x), microorganisms.translation$mo_old)
    # and their new codes
    mo_new <- microorganisms.translation$mo_new[matched]
    # assign on places where a match was found
    x[which(!is.na(matched))] <- mo_new[which(!is.na(matched))]
    n_matched <- length(matched[!is.na(matched)])
    if (property != "mo") {
      message_(font_blue("NOTE: The input contained old microbial codes (from previous package versions). Please update your MO codes with as.mo()."))
    } else {
      if (n_matched == 1) {
        message_(font_blue("NOTE: 1 old microbial code (from previous package versions) was updated to a current used MO code."))
      } else {
        message_(font_blue("NOTE:", n_matched, "old microbial codes (from previous package versions) were updated to current used MO codes.")) 
      }
    }
  }
  x
}

replace_ignore_pattern <- function(x, ignore_pattern) {
  if (!is.null(ignore_pattern) && !identical(trimws2(ignore_pattern), "")) {
    ignore_cases <- x %like% ignore_pattern
    if (sum(ignore_cases) > 0) {
      message_("The following input was ignored by `ignore_pattern = \"", ignore_pattern, "\"`: ",
               paste0("'", sort(unique(x[x %like% ignore_pattern])), "'", collapse = ", "),
               collapse = ", ")
      x[x %like% ignore_pattern] <- NA_character_
    }
  }
  x
}

repair_reference_df <- function(reference_df) {
  # has valid own reference_df
  reference_df <- reference_df %pm>%
    pm_filter(!is.na(mo))
  
  # keep only first two columns, second must be mo
  if (colnames(reference_df)[1] == "mo") {
    reference_df <- reference_df %pm>% pm_select(2, "mo")
  } else {
    reference_df <- reference_df %pm>% pm_select(1, "mo")
  }
  
  # remove factors, just keep characters
  colnames(reference_df)[1] <- "x"
  reference_df[, "x"] <- as.character(reference_df[, "x", drop = TRUE])
  reference_df[, "mo"] <- as.character(reference_df[, "mo", drop = TRUE])
  
  # some microbial codes might be old
  reference_df[, "mo"] <- as.mo(reference_df[, "mo", drop = TRUE])
  reference_df
}
````
