## Supplementary material for "AMR - An R Package for Working with Antimicrobial Resistance Data": Source package: mo_matching_score.R

```
# ==================================================================== #
# TITLE                                                                #
# Antimicrobial Resistance (AMR) Analysis for R                        #
#                                                                      #
# SOURCE                                                               #
# https://github.com/msberends/AMR                                     #
#                                                                      #
# LICENCE                                                              #
# (c) 2018-2021 Berends MS, Luz CF et al.                              #
# Developed at the University of Groningen, the Netherlands, in        #
# collaboration with non-profit organisations Certe Medical            #
# Diagnostics & Advice, and University Medical Center Groningen.       # 
#                                                                      #
# This R package is free software; you can freely use and distribute   #
# it for both personal and commercial purposes under the terms of the  #
# GNU General Public License version 2.0 (GNU GPL-2), as published by  #
# the Free Software Foundation.                                        #
# We created this package for both routine data analysis and academic  #
# research and it was publicly released in the hope that it will be    #
# useful, but it comes WITHOUT ANY WARRANTY OR LIABILITY.              #
#                                                                      #
# Visit our website for the full manual and a complete tutorial about  #
# how to conduct AMR analysis: https://msberends.github.io/AMR/        #
# ==================================================================== #

#' Calculate the matching score for microorganisms
#' 
#' This algorithm is used by [as.mo()] and all the [`mo_*`][mo_property()] functions to determine the most probable match of taxonomic records based on user input. 
#' @inheritSection lifecycle Stable lifecycle
#' @author Matthijs S. Berends
#' @param x Any user input value(s)
#' @param n A full taxonomic name, that exists in [`microorganisms$fullname`][microorganisms]
#' @section Matching score for microorganisms:
#' With ambiguous user input in [as.mo()] and all the [`mo_*`][mo_property()] functions, the returned results are chosen based on their matching score using [mo_matching_score()]. This matching score \eqn{m}, is calculated as:
#' 
#' \deqn{m_{(x, n)} = \frac{l_{n} - 0.5 \cdot \min \begin{cases}l_{n} \\ \textrm{lev}(x, n)\end{cases}}{l_{n} \cdot p_{n} \cdot k_{n}}}{m(x, n) = ( l_n * min(l_n, lev(x, n) ) ) / ( l_n * p_n * k_n )}
#' 
#' where:
#' 
#' * \eqn{x} is the user input;
#' * \eqn{n} is a taxonomic name (genus, species, and subspecies);
#' * \eqn{l_n}{l_n} is the length of \eqn{n};
#' * lev is the [Levenshtein distance function](https://en.wikipedia.org/wiki/Levenshtein_distance), which counts any insertion, deletion and substitution as 1 that is needed to change \eqn{x} into \eqn{n};
#' * \eqn{p_n}{p_n} is the human pathogenic prevalence group of \eqn{n}, as described below;
#' * \eqn{k_n}{p_n} is the taxonomic kingdom of \eqn{n}, set as Bacteria = 1, Fungi = 2, Protozoa = 3, Archaea = 4, others = 5.
#' 
#' The grouping into human pathogenic prevalence (\eqn{p}) is based on experience from several microbiological laboratories in the Netherlands in conjunction with international reports on pathogen prevalence. **Group 1** (most prevalent microorganisms) consists of all microorganisms where the taxonomic class is Gammaproteobacteria or where the taxonomic genus is *Enterococcus*, *Staphylococcus* or *Streptococcus*. This group consequently contains all common Gram-negative bacteria, such as *Pseudomonas* and *Legionella* and all species within the order Enterobacterales. **Group 2** consists of all microorganisms where the taxonomic phylum is Proteobacteria, Firmicutes, Actinobacteria or Sarcomastigophora, or where the taxonomic genus is *Absidia*, *Acremonium*, *Actinotignum*, *Alternaria*, *Anaerosalibacter*, *Apophysomyces*, *Arachnia*, *Aspergillus*, *Aureobacterium*, *Aureobasidium*, *Bacteroides*, *Basidiobolus*, *Beauveria*, *Blastocystis*, *Branhamella*, *Calymmatobacterium*, *Candida*, *Capnocytophaga*, *Catabacter*, *Chaetomium*, *Chryseobacterium*, *Chryseomonas*, *Chrysonilia*, *Cladophialophora*, *Cladosporium*, *Conidiobolus*, *Cryptococcus*, *Curvularia*, *Exophiala*, *Exserohilum*, *Flavobacterium*, *Fonsecaea*, *Fusarium*, *Fusobacterium*, *Hendersonula*, *Hypomyces*, *Koserella*, *Lelliottia*, *Leptosphaeria*, *Leptotrichia*, *Malassezia*, *Malbranchea*, *Mortierella*, *Mucor*, *Mycocentrospora*, *Mycoplasma*, *Nectria*, *Ochroconis*, *Oidiodendron*, *Phoma*, *Piedraia*, *Pithomyces*, *Pityrosporum*, *Prevotella*,\\*Pseudallescheria*, *Rhizomucor*, *Rhizopus*, *Rhodotorula*, *Scolecobasidium*, *Scopulariopsis*, *Scytalidium*,*Sporobolomyces*, *Stachybotrys*, *Stomatococcus*, *Treponema*, *Trichoderma*, *Trichophyton*, *Trichosporon*, *Tritirachium* or *Ureaplasma*. **Group 3** consists of all other microorganisms.
#' 
#' All matches are sorted descending on their matching score and for all user input values, the top match will be returned. This will lead to the effect that e.g., `"E. coli"` will return the microbial ID of *Escherichia coli* (\eqn{m = `r round(mo_matching_score("E. coli", "Escherichia coli"), 3)`}, a highly prevalent microorganism found in humans) and not *Entamoeba coli* (\eqn{m = `r round(mo_matching_score("E. coli", "Entamoeba coli"), 3)`}, a less prevalent microorganism in humans), although the latter would alphabetically come first. 
#' @export
#' @examples 
#' as.mo("E. coli")
#' mo_uncertainties()
#' 
#' mo_matching_score(x = "E. coli",
#'                   n = c("Escherichia coli", "Entamoeba coli"))
mo_matching_score <- function(x, n) {
  meet_criteria(x, allow_class = c("character", "data.frame", "list"))
  meet_criteria(n, allow_class = "character")
  
  x <- parse_and_convert(x)
  # no dots and other non-whitespace characters
  x <- gsub("[^a-zA-Z0-9 \\(\\)]+", "", x)
  # only keep one space
  x <- gsub(" +", " ", x)

  # n is always a taxonomically valid full name
  if (length(n) == 1) {
    n <- rep(n, length(x))
  }
  if (length(x) == 1) {
    x <- rep(x, length(n))
  }
  
  # length of fullname
  l_n <- nchar(n)
  lev <- double(length = length(x))
  l_n.lev <- double(length = length(x))
  for (i in seq_len(length(x))) {
    # determine Levenshtein distance, but maximise to nchar of n
    lev[i] <- utils::adist(x[i], n[i], ignore.case = FALSE, fixed = TRUE)
    # minimum of (l_n, Levenshtein distance)
    l_n.lev[i] <- min(l_n[i], as.double(lev[i]))
  }
  # human pathogenic prevalence (1 to 3), see ?as.mo
  p_n <- MO_lookup[match(n, MO_lookup$fullname), "prevalence", drop = TRUE]
  # kingdom index (Bacteria = 1, Fungi = 2, Protozoa = 3, Archaea = 4, others = 5)
  k_n <- MO_lookup[match(n, MO_lookup$fullname), "kingdom_index", drop = TRUE]
  
  # matching score:
  (l_n - 0.5 * l_n.lev) / (l_n * p_n * k_n)
}
```
