## Supplementary material for "AMR - An R Package for Working with Antimicrobial Resistance Data": Source package: mo_property.R

```
# ==================================================================== #
# TITLE                                                                #
# Antimicrobial Resistance (AMR) Analysis for R                        #
#                                                                      #
# SOURCE                                                               #
# https://github.com/msberends/AMR                                     #
#                                                                      #
# LICENCE                                                              #
# (c) 2018-2021 Berends MS, Luz CF et al.                              #
# Developed at the University of Groningen, the Netherlands, in        #
# collaboration with non-profit organisations Certe Medical            #
# Diagnostics & Advice, and University Medical Center Groningen.       #
#                                                                      #
# This R package is free software; you can freely use and distribute   #
# it for both personal and commercial purposes under the terms of the  #
# GNU General Public License version 2.0 (GNU GPL-2), as published by  #
# the Free Software Foundation.                                        #
# We created this package for both routine data analysis and academic  #
# research and it was publicly released in the hope that it will be    #
# useful, but it comes WITHOUT ANY WARRANTY OR LIABILITY.              #
#                                                                      #
# Visit our website for the full manual and a complete tutorial about  #
# how to conduct AMR analysis: https://msberends.github.io/AMR/        #
# ==================================================================== #

#' Get properties of a microorganism
#'
#' Use these functions to return a specific property of a microorganism based on the latest accepted taxonomy. All input values will be evaluated internally with [as.mo()], which makes it possible to use microbial abbreviations, codes and names as input. Please see *Examples*.
#' @inheritSection lifecycle Stable lifecycle
#' @param x any character (vector) that can be coerced to a valid microorganism code with [as.mo()]. Can be left blank for auto-guessing the column containing microorganism codes when used inside `dplyr` verbs, such as [`filter()`][dplyr::filter()], [`mutate()`][dplyr::mutate()] and [`summarise()`][dplyr::summarise()], please see *Examples*.
#' @param property one of the column names of the [microorganisms] data set: `r paste0('"``', colnames(microorganisms), '\``"', collapse = ", ")`, or must be `"shortname"`
#' @param language language of the returned text, defaults to system language (see [get_locale()]) and can be overwritten by setting the option `AMR_locale`, e.g. `options(AMR_locale = "de")`, see [translate]. Also used to translate text like "no growth". Use `language = NULL` or `language = ""` to prevent translation.
#' @param ... other arguments passed on to [as.mo()], such as 'allow_uncertain' and 'ignore_pattern'
#' @param ab any (vector of) text that can be coerced to a valid antibiotic code with [as.ab()]
#' @param open browse the URL using [`browseURL()`][utils::browseURL()]
#' @details All functions will return the most recently known taxonomic property according to the Catalogue of Life, except for [mo_ref()], [mo_authors()] and [mo_year()]. Please refer to this example, knowing that *Escherichia blattae* was renamed to *Shimwellia blattae* in 2010:
#' - `mo_name("Escherichia blattae")` will return `"Shimwellia blattae"` (with a message about the renaming)
#' - `mo_ref("Escherichia blattae")` will return `"Burgess et al., 1973"` (with a message about the renaming)
#' - `mo_ref("Shimwellia blattae")` will return `"Priest et al., 2010"` (without a message)
#'
#' The short name - [mo_shortname()] - almost always returns the first character of the genus and the full species, like `"E. coli"`. Exceptions are abbreviations of staphylococci (such as *"CoNS"*, Coagulase-Negative Staphylococci) and beta-haemolytic streptococci (such as *"GBS"*, Group B Streptococci). Please bear in mind that e.g. *E. coli* could mean *Escherichia coli* (kingdom of Bacteria) as well as *Entamoeba coli* (kingdom of Protozoa). Returning to the full name will be done using [as.mo()] internally, giving priority to bacteria and human pathogens, i.e. `"E. coli"` will be considered *Escherichia coli*. In other words, `mo_fullname(mo_shortname("Entamoeba coli"))` returns `"Escherichia coli"`.
#'
#' Since the top-level of the taxonomy is sometimes referred to as 'kingdom' and sometimes as 'domain', the functions [mo_kingdom()] and [mo_domain()] return the exact same results.
#'
#' The Gram stain - [mo_gramstain()] - will be determined based on the taxonomic kingdom and phylum. According to Cavalier-Smith (2002, [PMID 11837318](https://pubmed.ncbi.nlm.nih.gov/11837318)), who defined subkingdoms Negibacteria and Posibacteria, only these phyla are Posibacteria: Actinobacteria, Chloroflexi, Firmicutes and Tenericutes. These bacteria are considered Gram-positive - all other bacteria are considered Gram-negative. Species outside the kingdom of Bacteria will return a value `NA`. Functions [mo_is_gram_negative()] and [mo_is_gram_positive()] always return `TRUE` or `FALSE` (except when the input is `NA` or the MO code is `UNKNOWN`), thus always return `FALSE` for species outside the taxonomic kingdom of Bacteria.
#' 
#' Intrinsic resistance - [mo_is_intrinsic_resistant()] - will be determined based on the [intrinsic_resistant] data set, which is based on `r format_eucast_version_nr(3.2)`. The [mo_is_intrinsic_resistant()] can be vectorised over arguments `x` (input for microorganisms) and over `ab` (input for antibiotics).
#'
#' All output will be [translate]d where possible.
#'
#' The function [mo_url()] will return the direct URL to the online database entry, which also shows the scientific reference of the concerned species.
#' @inheritSection mo_matching_score Matching score for microorganisms
#' @inheritSection catalogue_of_life Catalogue of Life
#' @inheritSection as.mo Source
#' @rdname mo_property
#' @name mo_property
#' @return
#' - An [integer] in case of [mo_year()]
#' - A [list] in case of [mo_taxonomy()] and [mo_info()]
#' - A named [character] in case of [mo_url()]
#' - A [double] in case of [mo_snomed()]
#' - A [character] in all other cases
#' @export
#' @seealso [microorganisms]
#' @inheritSection AMR Reference data publicly available
#' @inheritSection AMR Read more on our website!
#' @examples
#' # taxonomic tree -----------------------------------------------------------
#' mo_kingdom("E. coli")         # "Bacteria"
#' mo_phylum("E. coli")          # "Proteobacteria"
#' mo_class("E. coli")           # "Gammaproteobacteria"
#' mo_order("E. coli")           # "Enterobacterales"
#' mo_family("E. coli")          # "Enterobacteriaceae"
#' mo_genus("E. coli")           # "Escherichia"
#' mo_species("E. coli")         # "coli"
#' mo_subspecies("E. coli")      # ""
#'
#' # colloquial properties ----------------------------------------------------
#' mo_name("E. coli")            # "Escherichia coli"
#' mo_fullname("E. coli")        # "Escherichia coli" - same as mo_name()
#' mo_shortname("E. coli")       # "E. coli"
#'
#' # other properties ---------------------------------------------------------
#' mo_gramstain("E. coli")       # "Gram-negative"
#' mo_snomed("E. coli")          # 112283007, 116395006, ... (SNOMED codes)
#' mo_type("E. coli")            # "Bacteria" (equal to kingdom, but may be translated)
#' mo_rank("E. coli")            # "species"
#' mo_url("E. coli")             # get the direct url to the online database entry
#' mo_synonyms("E. coli")        # get previously accepted taxonomic names
#'
#' # scientific reference -----------------------------------------------------
#' mo_ref("E. coli")             # "Castellani et al., 1919"
#' mo_authors("E. coli")         # "Castellani et al."
#' mo_year("E. coli")            # 1919
#'
#' # abbreviations known in the field -----------------------------------------
#' mo_genus("MRSA")              # "Staphylococcus"
#' mo_species("MRSA")            # "aureus"
#' mo_shortname("VISA")          # "S. aureus"
#' mo_gramstain("VISA")          # "Gram-positive"
#'
#' mo_genus("EHEC")              # "Escherichia"
#' mo_species("EHEC")            # "coli"
#'
#' # known subspecies ---------------------------------------------------------
#' mo_name("doylei")             # "Campylobacter jejuni doylei"
#' mo_genus("doylei")            # "Campylobacter"
#' mo_species("doylei")          # "jejuni"
#' mo_subspecies("doylei")       # "doylei"
#'
#' mo_fullname("K. pneu rh")     # "Klebsiella pneumoniae rhinoscleromatis"
#' mo_shortname("K. pneu rh")    # "K. pneumoniae"
#'
#' \donttest{
#' # Becker classification, see ?as.mo ----------------------------------------
#' mo_fullname("S. epi")                     # "Staphylococcus epidermidis"
#' mo_fullname("S. epi", Becker = TRUE)      # "Coagulase-negative Staphylococcus (CoNS)"
#' mo_shortname("S. epi")                    # "S. epidermidis"
#' mo_shortname("S. epi", Becker = TRUE)     # "CoNS"
#'
#' # Lancefield classification, see ?as.mo ------------------------------------
#' mo_fullname("S. pyo")                     # "Streptococcus pyogenes"
#' mo_fullname("S. pyo", Lancefield = TRUE)  # "Streptococcus group A"
#' mo_shortname("S. pyo")                    # "S. pyogenes"
#' mo_shortname("S. pyo", Lancefield = TRUE) # "GAS" (='Group A Streptococci')
#'
#'
#' # language support  --------------------------------------------------------
#' mo_gramstain("E. coli", language = "de")  # "Gramnegativ"
#' mo_gramstain("E. coli", language = "nl")  # "Gram-negatief"
#' mo_gramstain("E. coli", language = "es")  # "Gram negativo"
#'
#' # mo_type is equal to mo_kingdom, but mo_kingdom will remain official
#' mo_kingdom("E. coli")                     # "Bacteria" on a German system
#' mo_type("E. coli")                        # "Bakterien" on a German system
#' mo_type("E. coli")                        # "Bacteria" on an English system
#'
#' mo_fullname("S. pyogenes",
#'             Lancefield = TRUE,
#'             language = "de")              # "Streptococcus Gruppe A"
#' mo_fullname("S. pyogenes",
#'             Lancefield = TRUE,
#'             language = "nl")              # "Streptococcus groep A"
#'
#'
#' # other --------------------------------------------------------------------
#' 
#' # gram stains and intrinsic resistance can also be used as a filter in dplyr verbs
#' if (require("dplyr")) {
#'   example_isolates %>%
#'     filter(mo_is_gram_positive())
#'     
#'   example_isolates %>%
#'     filter(mo_is_intrinsic_resistant(ab = "vanco"))
#' }
#' 
#' 
#' # get a list with the complete taxonomy (from kingdom to subspecies)
#' mo_taxonomy("E. coli")
#' # get a list with the taxonomy, the authors, Gram-stain and URL to the online database
#' mo_info("E. coli")
#' }
mo_name <- function(x, language = get_locale(), ...) {
  if (missing(x)) {
    # this tries to find the data and an <mo> column
    x <- find_mo_col(fn = "mo_name")
  }
  meet_criteria(x, allow_NA = TRUE)
  meet_criteria(language, has_length = 1, is_in = c(LANGUAGES_SUPPORTED, ""), allow_NULL = TRUE, allow_NA = TRUE)
  
  translate_AMR(mo_validate(x = x, property = "fullname", language = language, ...), language = language, only_unknown = FALSE)
}

#' @rdname mo_property
#' @export
mo_fullname <- mo_name

#' @rdname mo_property
#' @export
mo_shortname <- function(x, language = get_locale(), ...) {
  if (missing(x)) {
    # this tries to find the data and an <mo> column
    x <- find_mo_col(fn = "mo_shortname")
  }
  meet_criteria(x, allow_NA = TRUE)
  meet_criteria(language, has_length = 1, is_in = c(LANGUAGES_SUPPORTED, ""), allow_NULL = TRUE, allow_NA = TRUE)
  
  x.mo <- as.mo(x, language = language, ...)
  
  metadata <- get_mo_failures_uncertainties_renamed()
  
  replace_empty <- function(x) {
    x[x == ""] <- "spp."
    x
  }
  
  # get first char of genus and complete species in English
  genera <- mo_genus(x.mo, language = NULL)
  shortnames <- paste0(substr(genera, 1, 1), ". ", replace_empty(mo_species(x.mo, language = NULL)))
  
  # exceptions for where no species is known
  shortnames[shortnames %like% ".[.] spp[.]"] <- genera[shortnames %like% ".[.] spp[.]"]
  # exceptions for staphylococci
  shortnames[shortnames == "S. coagulase-negative"] <- "CoNS"
  shortnames[shortnames == "S. coagulase-positive"] <- "CoPS"
  # exceptions for streptococci: Group A Streptococcus -> GAS
  shortnames[shortnames %like% "S. group [ABCDFGHK]"] <- paste0("G", gsub("S. group ([ABCDFGHK])", "\\1", shortnames[shortnames %like% "S. group [ABCDFGHK]"]), "S")
  # unknown species etc.
  shortnames[shortnames %like% "unknown"] <- paste0("(", trimws(gsub("[^a-zA-Z -]", "", shortnames[shortnames %like% "unknown"])), ")")
  
  shortnames[is.na(x.mo)] <- NA_character_
  load_mo_failures_uncertainties_renamed(metadata)
  translate_AMR(shortnames, language = language, only_unknown = FALSE)
}

#' @rdname mo_property
#' @export
mo_subspecies <- function(x, language = get_locale(), ...) {
  if (missing(x)) {
    # this tries to find the data and an <mo> column
    x <- find_mo_col(fn = "mo_subspecies")
  }
  meet_criteria(x, allow_NA = TRUE)
  meet_criteria(language, has_length = 1, is_in = c(LANGUAGES_SUPPORTED, ""), allow_NULL = TRUE, allow_NA = TRUE)
  
  translate_AMR(mo_validate(x = x, property = "subspecies", language = language, ...), language = language, only_unknown = TRUE)
}

#' @rdname mo_property
#' @export
mo_species <- function(x, language = get_locale(), ...) {
  if (missing(x)) {
    # this tries to find the data and an <mo> column
    x <- find_mo_col(fn = "mo_species")
  }
  meet_criteria(x, allow_NA = TRUE)
  meet_criteria(language, has_length = 1, is_in = c(LANGUAGES_SUPPORTED, ""), allow_NULL = TRUE, allow_NA = TRUE)
  
  translate_AMR(mo_validate(x = x, property = "species", language = language, ...), language = language, only_unknown = TRUE)
}

#' @rdname mo_property
#' @export
mo_genus <- function(x, language = get_locale(), ...) {
  if (missing(x)) {
    # this tries to find the data and an <mo> column
    x <- find_mo_col(fn = "mo_genus")
  }
  meet_criteria(x, allow_NA = TRUE)
  meet_criteria(language, has_length = 1, is_in = c(LANGUAGES_SUPPORTED, ""), allow_NULL = TRUE, allow_NA = TRUE)
  
  translate_AMR(mo_validate(x = x, property = "genus", language = language, ...), language = language, only_unknown = TRUE)
}

#' @rdname mo_property
#' @export
mo_family <- function(x, language = get_locale(), ...) {
  if (missing(x)) {
    # this tries to find the data and an <mo> column
    x <- find_mo_col(fn = "mo_family")
  }
  meet_criteria(x, allow_NA = TRUE)
  meet_criteria(language, has_length = 1, is_in = c(LANGUAGES_SUPPORTED, ""), allow_NULL = TRUE, allow_NA = TRUE)
  
  translate_AMR(mo_validate(x = x, property = "family", language = language, ...), language = language, only_unknown = TRUE)
}

#' @rdname mo_property
#' @export
mo_order <- function(x, language = get_locale(), ...) {
  if (missing(x)) {
    # this tries to find the data and an <mo> column
    x <- find_mo_col(fn = "mo_order")
  }
  meet_criteria(x, allow_NA = TRUE)
  meet_criteria(language, has_length = 1, is_in = c(LANGUAGES_SUPPORTED, ""), allow_NULL = TRUE, allow_NA = TRUE)
  
  translate_AMR(mo_validate(x = x, property = "order", language = language, ...), language = language, only_unknown = TRUE)
}

#' @rdname mo_property
#' @export
mo_class <- function(x, language = get_locale(), ...) {
  if (missing(x)) {
    # this tries to find the data and an <mo> column
    x <- find_mo_col(fn = "mo_class")
  }
  meet_criteria(x, allow_NA = TRUE)
  meet_criteria(language, has_length = 1, is_in = c(LANGUAGES_SUPPORTED, ""), allow_NULL = TRUE, allow_NA = TRUE)
  
  translate_AMR(mo_validate(x = x, property = "class", language = language, ...), language = language, only_unknown = TRUE)
}

#' @rdname mo_property
#' @export
mo_phylum <- function(x, language = get_locale(), ...) {
  if (missing(x)) {
    # this tries to find the data and an <mo> column
    x <- find_mo_col(fn = "mo_phylum")
  }
  meet_criteria(x, allow_NA = TRUE)
  meet_criteria(language, has_length = 1, is_in = c(LANGUAGES_SUPPORTED, ""), allow_NULL = TRUE, allow_NA = TRUE)
  
  translate_AMR(mo_validate(x = x, property = "phylum", language = language, ...), language = language, only_unknown = TRUE)
}

#' @rdname mo_property
#' @export
mo_kingdom <- function(x, language = get_locale(), ...) {
  if (missing(x)) {
    # this tries to find the data and an <mo> column
    x <- find_mo_col(fn = "mo_kingdom")
  }
  meet_criteria(x, allow_NA = TRUE)
  meet_criteria(language, has_length = 1, is_in = c(LANGUAGES_SUPPORTED, ""), allow_NULL = TRUE, allow_NA = TRUE)
  
  translate_AMR(mo_validate(x = x, property = "kingdom", language = language, ...), language = language, only_unknown = TRUE)
}

#' @rdname mo_property
#' @export
mo_domain <- mo_kingdom

#' @rdname mo_property
#' @export
mo_type <- function(x, language = get_locale(), ...) {
  if (missing(x)) {
    # this tries to find the data and an <mo> column
    x <- find_mo_col(fn = "mo_type")
  }
  meet_criteria(x, allow_NA = TRUE)
  meet_criteria(language, has_length = 1, is_in = c(LANGUAGES_SUPPORTED, ""), allow_NULL = TRUE, allow_NA = TRUE)
  
  translate_AMR(mo_validate(x = x, property = "kingdom", language = language, ...), language = language, only_unknown = FALSE)
}

#' @rdname mo_property
#' @export
mo_gramstain <- function(x, language = get_locale(), ...) {
  if (missing(x)) {
    # this tries to find the data and an <mo> column
    x <- find_mo_col(fn = "mo_gramstain")
  }
  meet_criteria(x, allow_NA = TRUE)
  meet_criteria(language, has_length = 1, is_in = c(LANGUAGES_SUPPORTED, ""), allow_NULL = TRUE, allow_NA = TRUE)
  
  x.mo <- as.mo(x, language = language, ...)
  metadata <- get_mo_failures_uncertainties_renamed()
  
  x.phylum <- mo_phylum(x.mo)
  # DETERMINE GRAM STAIN FOR BACTERIA
  # Source: https://itis.gov/servlet/SingleRpt/SingleRpt?search_topic=TSN&search_value=956097
  # It says this:
  # Kingdom Bacteria (Cavalier-Smith, 2002)
  #    Subkingdom Posibacteria (Cavalier-Smith, 2002)
  #   Direct Children:
  #       Phylum  Actinobacteria (Cavalier-Smith, 2002)
  #       Phylum  Chloroflexi (Garrity and Holt, 2002)
  #       Phylum  Firmicutes (corrig. Gibbons and Murray, 1978)
  #       Phylum  Tenericutes (Murray, 1984)
  x <- NA_character_
  # make all bacteria Gram negative
  x[mo_kingdom(x.mo) == "Bacteria"] <- "Gram-negative"
  # overwrite these phyla with Gram positive
  x[x.phylum %in% c("Actinobacteria",
                    "Chloroflexi",
                    "Firmicutes",
                    "Tenericutes")
    | x.mo == "B_GRAMP"] <- "Gram-positive"
  
  load_mo_failures_uncertainties_renamed(metadata)
  translate_AMR(x, language = language, only_unknown = FALSE)
}

#' @rdname mo_property
#' @export
mo_is_gram_negative <- function(x, language = get_locale(), ...) {
  if (missing(x)) {
    # this tries to find the data and an <mo> column
    x <- find_mo_col(fn = "mo_is_gram_negative")
  }
  meet_criteria(x, allow_NA = TRUE)
  meet_criteria(language, has_length = 1, is_in = c(LANGUAGES_SUPPORTED, ""), allow_NULL = TRUE, allow_NA = TRUE)
  
  x.mo <- as.mo(x, language = language, ...)
  metadata <- get_mo_failures_uncertainties_renamed()
  grams <- mo_gramstain(x.mo, language = NULL)
  load_mo_failures_uncertainties_renamed(metadata)
  out <- grams == "Gram-negative" & !is.na(grams)
  out[x.mo %in% c(NA_character_, "UNKNOWN")] <- NA
  out
}

#' @rdname mo_property
#' @export
mo_is_gram_positive <- function(x, language = get_locale(), ...) {
  if (missing(x)) {
    # this tries to find the data and an <mo> column
    x <- find_mo_col(fn = "mo_is_gram_positive")
  }
  meet_criteria(x, allow_NA = TRUE)
  meet_criteria(language, has_length = 1, is_in = c(LANGUAGES_SUPPORTED, ""), allow_NULL = TRUE, allow_NA = TRUE)
  
  x.mo <- as.mo(x, language = language, ...)
  metadata <- get_mo_failures_uncertainties_renamed()
  grams <- mo_gramstain(x.mo, language = NULL)
  load_mo_failures_uncertainties_renamed(metadata)
  out <- grams == "Gram-positive" & !is.na(grams)
  out[x.mo %in% c(NA_character_, "UNKNOWN")] <- NA
  out
}

#' @rdname mo_property
#' @export
mo_is_intrinsic_resistant <- function(x, ab, language = get_locale(), ...) {
  if (missing(x)) {
    # this tries to find the data and an <mo> column
    x <- find_mo_col(fn = "mo_is_intrinsic_resistant")
  }
  meet_criteria(x, allow_NA = TRUE)
  meet_criteria(ab, allow_NA = FALSE)
  meet_criteria(language, has_length = 1, is_in = c(LANGUAGES_SUPPORTED, ""), allow_NULL = TRUE, allow_NA = TRUE)
  
  x <- as.mo(x, language = language, ...)
  ab <- as.ab(ab, language = NULL, flag_multiple_results = FALSE, info = FALSE)
  
  if (length(x) == 1 & length(ab) > 1) {
    x <- rep(x, length(ab))
  } else if (length(ab) == 1 & length(x) > 1) {
    ab <- rep(ab, length(x))
  }
  if (length(x) != length(ab)) {
    stop_("length of `x` and `ab` must be equal, or one of them must be of length 1.")
  }
  
  # show used version number once per session (pkg_env will reload every session)
  if (message_not_thrown_before("intrinsic_resistant_version", entire_session = TRUE)) {
    message_("Determining intrinsic resistance based on ",
             format_eucast_version_nr(3.2, markdown = FALSE), ". ",
             font_red("This note will be shown once per session."))
    remember_thrown_message("intrinsic_resistant_version", entire_session = TRUE)
  }
  
  # runs against internal vector: INTRINSIC_R (see zzz.R)
  paste(x, ab) %in% INTRINSIC_R
}

#' @rdname mo_property
#' @export
mo_snomed <- function(x, language = get_locale(), ...) {
  if (missing(x)) {
    # this tries to find the data and an <mo> column
    x <- find_mo_col(fn = "mo_snomed")
  }
  meet_criteria(x, allow_NA = TRUE)
  meet_criteria(language, has_length = 1, is_in = c(LANGUAGES_SUPPORTED, ""), allow_NULL = TRUE, allow_NA = TRUE)
  
  mo_validate(x = x, property = "snomed", language = language, ...)
}

#' @rdname mo_property
#' @export
mo_ref <- function(x, language = get_locale(), ...) {
  if (missing(x)) {
    # this tries to find the data and an <mo> column
    x <- find_mo_col(fn = "mo_ref")
  }
  meet_criteria(x, allow_NA = TRUE)
  meet_criteria(language, has_length = 1, is_in = c(LANGUAGES_SUPPORTED, ""), allow_NULL = TRUE, allow_NA = TRUE)
  
  mo_validate(x = x, property = "ref", language = language, ...)
}

#' @rdname mo_property
#' @export
mo_authors <- function(x, language = get_locale(), ...) {
  if (missing(x)) {
    # this tries to find the data and an <mo> column
    x <- find_mo_col(fn = "mo_authors")
  }
  meet_criteria(x, allow_NA = TRUE)
  meet_criteria(language, has_length = 1, is_in = c(LANGUAGES_SUPPORTED, ""), allow_NULL = TRUE, allow_NA = TRUE)
  
  x <- mo_validate(x = x, property = "ref", language = language, ...)
  # remove last 4 digits and presumably the comma and space that preceed them
  x[!is.na(x)] <- gsub(",? ?[0-9]{4}", "", x[!is.na(x)])
  suppressWarnings(x)
}

#' @rdname mo_property
#' @export
mo_year <- function(x, language = get_locale(), ...) {
  if (missing(x)) {
    # this tries to find the data and an <mo> column
    x <- find_mo_col(fn = "mo_year")
  }
  meet_criteria(x, allow_NA = TRUE)
  meet_criteria(language, has_length = 1, is_in = c(LANGUAGES_SUPPORTED, ""), allow_NULL = TRUE, allow_NA = TRUE)
  
  x <- mo_validate(x = x, property = "ref", language = language, ...)
  # get last 4 digits
  x[!is.na(x)] <- gsub(".*([0-9]{4})$", "\\1", x[!is.na(x)])
  suppressWarnings(as.integer(x))
}

#' @rdname mo_property
#' @export
mo_rank <- function(x, language = get_locale(), ...) {
  if (missing(x)) {
    # this tries to find the data and an <mo> column
    x <- find_mo_col(fn = "mo_rank")
  }
  meet_criteria(x, allow_NA = TRUE)
  meet_criteria(language, has_length = 1, is_in = c(LANGUAGES_SUPPORTED, ""), allow_NULL = TRUE, allow_NA = TRUE)
  
  mo_validate(x = x, property = "rank", language = language, ...)
}

#' @rdname mo_property
#' @export
mo_taxonomy <- function(x, language = get_locale(),  ...) {
  if (missing(x)) {
    # this tries to find the data and an <mo> column
    x <- find_mo_col(fn = "mo_taxonomy")
  }
  meet_criteria(x, allow_NA = TRUE)
  meet_criteria(language, has_length = 1, is_in = c(LANGUAGES_SUPPORTED, ""), allow_NULL = TRUE, allow_NA = TRUE)
  
  x <- as.mo(x, language = language, ...)
  metadata <- get_mo_failures_uncertainties_renamed()
  
  result <- list(kingdom = mo_kingdom(x, language = language),
                 phylum = mo_phylum(x, language = language),
                 class = mo_class(x, language = language),
                 order = mo_order(x, language = language),
                 family = mo_family(x, language = language),
                 genus = mo_genus(x, language = language),
                 species = mo_species(x, language = language),
                 subspecies = mo_subspecies(x, language = language))
  
  load_mo_failures_uncertainties_renamed(metadata)
  result
}

#' @rdname mo_property
#' @export
mo_synonyms <- function(x, language = get_locale(), ...) {
  if (missing(x)) {
    # this tries to find the data and an <mo> column
    x <- find_mo_col(fn = "mo_synonyms")
  }
  meet_criteria(x, allow_NA = TRUE)
  meet_criteria(language, has_length = 1, is_in = c(LANGUAGES_SUPPORTED, ""), allow_NULL = TRUE, allow_NA = TRUE)
  
  x <- as.mo(x, language = language, ...)
  metadata <- get_mo_failures_uncertainties_renamed()
  
  IDs <- mo_name(x = x, language = NULL)
  syns <- lapply(IDs, function(newname) {
    res <- sort(microorganisms.old[which(microorganisms.old$fullname_new == newname), "fullname"])
    if (length(res) == 0) {
      NULL
    } else {
      res
    }
  })
  if (length(syns) > 1) {
    names(syns) <- mo_name(x)
    result <- syns
  } else {
    result <- unlist(syns)
  }
  
  load_mo_failures_uncertainties_renamed(metadata)
  result
}

#' @rdname mo_property
#' @export
mo_info <- function(x, language = get_locale(),  ...) {
  if (missing(x)) {
    # this tries to find the data and an <mo> column
    x <- find_mo_col(fn = "mo_info")
  }
  meet_criteria(x, allow_NA = TRUE)
  meet_criteria(language, has_length = 1, is_in = c(LANGUAGES_SUPPORTED, ""), allow_NULL = TRUE, allow_NA = TRUE)
  
  x <- as.mo(x, language = language, ...)
  metadata <- get_mo_failures_uncertainties_renamed()
  
  info <- lapply(x, function(y)
    c(mo_taxonomy(y, language = language),
      list(synonyms = mo_synonyms(y),
           gramstain = mo_gramstain(y, language = language),
           url = unname(mo_url(y, open = FALSE)),
           ref = mo_ref(y))))
  if (length(info) > 1) {
    names(info) <- mo_name(x)
    result <- info
  } else {
    result <- info[[1L]]
  }
  
  load_mo_failures_uncertainties_renamed(metadata)
  result
}

#' @rdname mo_property
#' @export
mo_url <- function(x, open = FALSE, language = get_locale(), ...) {
  if (missing(x)) {
    # this tries to find the data and an <mo> column
    x <- find_mo_col(fn = "mo_url")
  }
  meet_criteria(x, allow_NA = TRUE)
  meet_criteria(open, allow_class = "logical", has_length = 1)
  meet_criteria(language, has_length = 1, is_in = c(LANGUAGES_SUPPORTED, ""), allow_NULL = TRUE, allow_NA = TRUE)
  
  mo <- as.mo(x = x, language = language, ... = ...)
  mo_names <- mo_name(mo)
  metadata <- get_mo_failures_uncertainties_renamed()
  
  df <- data.frame(mo, stringsAsFactors = FALSE) %pm>%
    pm_left_join(pm_select(microorganisms, mo, source, species_id), by = "mo")
  df$url <- ifelse(df$source == "CoL",
                   paste0(catalogue_of_life$url_CoL, "details/species/id/", df$species_id, "/"),
                   ifelse(df$source == "DSMZ",
                          paste0(catalogue_of_life$url_DSMZ, "/advanced_search?adv[taxon-name]=", gsub(" ", "+", mo_names), "/"),
                          NA_character_))
  u <- df$url
  names(u) <- mo_names
  
  if (open == TRUE) {
    if (length(u) > 1) {
      warning_("Only the first URL will be opened, as `browseURL()` only suports one string.")
    }
    utils::browseURL(u[1L])
  }
  
  load_mo_failures_uncertainties_renamed(metadata)
  u
}


#' @rdname mo_property
#' @export
mo_property <- function(x, property = "fullname", language = get_locale(), ...) {
  if (missing(x)) {
    # this tries to find the data and an <mo> column
    x <- find_mo_col(fn = "mo_property")
  }
  meet_criteria(x, allow_NA = TRUE)
  meet_criteria(property, allow_class = "character", has_length = 1, is_in = colnames(microorganisms))
  meet_criteria(language, has_length = 1, is_in = c(LANGUAGES_SUPPORTED, ""), allow_NULL = TRUE, allow_NA = TRUE)
  
  translate_AMR(mo_validate(x = x, property = property, language = language, ...), language = language, only_unknown = TRUE)
}

mo_validate <- function(x, property, language, ...) {
  check_dataset_integrity()
  
  if (tryCatch(all(x[!is.na(x)] %in% MO_lookup$mo) & length(list(...)) == 0, error = function(e) FALSE)) {
    # special case for mo_* functions where class is already <mo>
    return(MO_lookup[match(x, MO_lookup$mo), property, drop = TRUE])
  }
  
  dots <- list(...)
  Becker <- dots$Becker
  if (is.null(Becker)) {
    Becker <- FALSE
  }
  Lancefield <- dots$Lancefield
  if (is.null(Lancefield)) {
    Lancefield <- FALSE
  }
  
  # try to catch an error when inputting an invalid argument
  # so the 'call.' can be set to FALSE
  tryCatch(x[1L] %in% MO_lookup[1, property, drop = TRUE],
           error = function(e) stop(e$message, call. = FALSE))
  
  if (is.mo(x)
      & !Becker %in% c(TRUE, "all")
      & !Lancefield %in% c(TRUE, "all")) {
    # this will not reset mo_uncertainties and mo_failures
    # because it's already a valid MO
    x <- exec_as.mo(x, property = property, initial_search = FALSE, language = language, ...)
  } else if (!all(x %in% MO_lookup[, property, drop = TRUE])
             | Becker %in% c(TRUE, "all")
             | Lancefield %in% c(TRUE, "all")) {
    x <- exec_as.mo(x, property = property, language = language, ...)
  }
  
  if (property == "mo") {
    return(set_clean_class(x, new_class = c("mo", "character")))
  } else if (property == "snomed") {
    return(as.double(eval(parse(text = x))))
  } else {
    return(x)
  }
}

find_mo_col <- function(fn) {
  # this function tries to find an mo column in the data the function was called in,
  # which is useful when functions are used within dplyr verbs
  df <- get_current_data(arg_name = "x", call = -3) # will return an error if not found
  mo <- NULL
  try({
    mo <- suppressMessages(search_type_in_df(df, "mo"))
  }, silent = TRUE)
  if (!is.null(df) && !is.null(mo) && is.data.frame(df)) {
    if (message_not_thrown_before(fn = fn)) {
      message_("Using column '", font_bold(mo), "' as input for ", fn, "()")
      remember_thrown_message(fn = fn)
    }
    return(df[, mo, drop = TRUE])
  } else {
    stop_("argument `x` is missing and no column with info about microorganisms could be found.", call = -2)
  }
}
```
