## Supplementary material for "AMR - An R Package for Working with Antimicrobial Resistance Data": Source package: mo_source.R

````
# ==================================================================== #
# TITLE                                                                #
# Antimicrobial Resistance (AMR) Analysis for R                        #
#                                                                      #
# SOURCE                                                               #
# https://github.com/msberends/AMR                                     #
#                                                                      #
# LICENCE                                                              #
# (c) 2018-2021 Berends MS, Luz CF et al.                              #
# Developed at the University of Groningen, the Netherlands, in        #
# collaboration with non-profit organisations Certe Medical            #
# Diagnostics & Advice, and University Medical Center Groningen.       # 
#                                                                      #
# This R package is free software; you can freely use and distribute   #
# it for both personal and commercial purposes under the terms of the  #
# GNU General Public License version 2.0 (GNU GPL-2), as published by  #
# the Free Software Foundation.                                        #
# We created this package for both routine data analysis and academic  #
# research and it was publicly released in the hope that it will be    #
# useful, but it comes WITHOUT ANY WARRANTY OR LIABILITY.              #
#                                                                      #
# Visit our website for the full manual and a complete tutorial about  #
# how to conduct AMR analysis: https://msberends.github.io/AMR/        #
# ==================================================================== #

#' User-defined reference data set for microorganisms
#'
#' @description These functions can be used to predefine your own reference to be used in [as.mo()] and consequently all [`mo_*`][mo_property()] functions (such as [mo_genus()] and [mo_gramstain()]).
#'
#' This is **the fastest way** to have your organisation (or analysis) specific codes picked up and translated by this package, since you don't have to bother about it again after setting it up once.
#' @inheritSection lifecycle Stable lifecycle
#' @param path location of your reference file, see Details. Can be `""`, `NULL` or `FALSE` to delete the reference file.
#' @param destination destination of the compressed data file, default to the user's home directory.
#' @rdname mo_source
#' @name mo_source
#' @aliases set_mo_source get_mo_source
#' @details The reference file can be a text file separated with commas (CSV) or tabs or pipes, an Excel file (either 'xls' or 'xlsx' format) or an \R object file (extension '.rds'). To use an Excel file, you will need to have the `readxl` package installed.
#'
#' [set_mo_source()] will check the file for validity: it must be a [data.frame], must have a column named `"mo"` which contains values from [`microorganisms$mo`][microorganisms] and must have a reference column with your own defined values. If all tests pass, [set_mo_source()] will read the file into \R and will ask to export it to `"~/mo_source.rds"`. The CRAN policy disallows packages to write to the file system, although '*exceptions may be allowed in interactive sessions if the package obtains confirmation from the user*'. For this reason, this function only works in interactive sessions so that the user can **specifically confirm and allow** that this file will be created. The destination of this file can be set with the `destination` argument and defaults to the user's home directory. It can also be set as an \R option, using `options(AMR_mo_source = "my/location/file.rds")`.
#' 
#' The created compressed data file `"mo_source.rds"` will be used at default for MO determination (function [as.mo()] and consequently all `mo_*` functions like [mo_genus()] and [mo_gramstain()]). The location and timestamp of the original file will be saved as an attribute to the compressed data file. 
#' 
#' The function [get_mo_source()] will return the data set by reading `"mo_source.rds"` with [readRDS()]. If the original file has changed (by checking the location and timestamp of the original file), it will call [set_mo_source()] to update the data file automatically if used in an interactive session.
#'
#' Reading an Excel file (`.xlsx`) with only one row has a size of 8-9 kB. The compressed file created with [set_mo_source()] will then have a size of 0.1 kB and can be read by [get_mo_source()] in only a couple of microseconds (millionths of a second).
#' 
#' @section How to setup:
#' 
#' Imagine this data on a sheet of an Excel file (mo codes were looked up in the [microorganisms] data set). The first column contains the organisation specific codes, the second column contains an MO code from this package:
#' 
#' ```
#'   |         A          |       B      |
#' --|--------------------|--------------|
#' 1 | Organisation XYZ   | mo           |
#' 2 | lab_mo_ecoli       | B_ESCHR_COLI |
#' 3 | lab_mo_kpneumoniae | B_KLBSL_PNMN |
#' 4 |                    |              |
#' ```
#'
#' We save it as `"home/me/ourcodes.xlsx"`. Now we have to set it as a source:
#' 
#' ```
#' set_mo_source("home/me/ourcodes.xlsx")
#' #> NOTE: Created mo_source file '/Users/me/mo_source.rds' (0.3 kB) from
#' #>       '/Users/me/Documents/ourcodes.xlsx' (9 kB), columns 
#' #>       "Organisation XYZ" and "mo"
#' ```
#'
#' It has now created a file `"~/mo_source.rds"` with the contents of our Excel file. Only the first column with foreign values and the 'mo' column will be kept when creating the RDS file.
#'
#' And now we can use it in our functions:
#' 
#' ```
#' as.mo("lab_mo_ecoli")
#' #> Class <mo>
#' #> [1] B_ESCHR_COLI
#'
#' mo_genus("lab_mo_kpneumoniae")
#' #> [1] "Klebsiella"
#'
#' # other input values still work too
#' as.mo(c("Escherichia coli", "E. coli", "lab_mo_ecoli"))
#' #> NOTE: Translation to one microorganism was guessed with uncertainty.
#' #>       Use mo_uncertainties() to review it.
#' #> Class <mo>
#' #> [1] B_ESCHR_COLI B_ESCHR_COLI B_ESCHR_COLI
#' ```
#'
#' If we edit the Excel file by, let's say, adding row 4 like this:
#' 
#' ```
#'   |         A          |       B      |
#' --|--------------------|--------------|
#' 1 | Organisation XYZ   | mo           |
#' 2 | lab_mo_ecoli       | B_ESCHR_COLI |
#' 3 | lab_mo_kpneumoniae | B_KLBSL_PNMN |
#' 4 | lab_Staph_aureus   | B_STPHY_AURS |
#' 5 |                    |              |
#' ```
#'
#' ...any new usage of an MO function in this package will update your data file:
#' 
#' ```
#' as.mo("lab_mo_ecoli")
#' #> NOTE: Updated mo_source file '/Users/me/mo_source.rds' (0.3 kB) from 
#' #>       '/Users/me/Documents/ourcodes.xlsx' (9 kB), columns
#' #>        "Organisation XYZ" and "mo"
#' #> Class <mo>
#' #> [1] B_ESCHR_COLI
#'
#' mo_genus("lab_Staph_aureus")
#' #> [1] "Staphylococcus"
#' ```
#'
#' To delete the reference data file, just use `""`, `NULL` or `FALSE` as input for [set_mo_source()]:
#' 
#' ```
#' set_mo_source(NULL)
#' #> Removed mo_source file '/Users/me/mo_source.rds'
#' ```
#' 
#' If the original file (in the previous case an Excel file) is moved or deleted, the `mo_source.rds` file will be removed upon the next use of [as.mo()] or any [`mo_*`][mo_property()] function.
#' @export
#' @inheritSection AMR Read more on our website!
set_mo_source <- function(path, destination = getOption("AMR_mo_source", "~/mo_source.rds")) {
  meet_criteria(path, allow_class = "character", has_length = 1, allow_NULL = TRUE)
  meet_criteria(destination, allow_class = "character", has_length = 1)
  stop_ifnot(destination %like% "[.]rds$", "the `destination` must be a file location with file extension .rds.")
  
  mo_source_destination <- path.expand(destination)
  
  stop_ifnot(interactive(), "this function can only be used in interactive mode, since it must ask for the user's permission to write a file to their home folder.")

  if (is.null(path) || path %in% c(FALSE, "")) {
    pkg_env$mo_source <- NULL
    if (file.exists(mo_source_destination)) {
      unlink(mo_source_destination)
      message_("Removed mo_source file '", font_bold(mo_source_destination), "'",
               add_fn = font_red,
               as_note = FALSE)
    }
    return(invisible())
  }
  
  stop_ifnot(file.exists(path), "file not found: ", path)
  
  if (path %like% "[.]rds$") {
    df <- readRDS(path)
    
  } else if (path %like% "[.]xlsx?$") {
    # is Excel file (old or new)
    stop_ifnot_installed("readxl")
    df <- readxl::read_excel(path)
    
  } else if (path %like% "[.]tsv$") {
    df <- utils::read.table(header = TRUE, sep = "\t", stringsAsFactors = FALSE)
    
  } else {
    # try comma first
    try(
      df <- utils::read.table(header = TRUE, sep = ",", stringsAsFactors = FALSE),
      silent = TRUE)
    if (!check_validity_mo_source(df, stop_on_error = FALSE)) {
      # try tab
      try(
        df <- utils::read.table(header = TRUE, sep = "\t", stringsAsFactors = FALSE),
        silent = TRUE)
    }
    if (!check_validity_mo_source(df, stop_on_error = FALSE)) {
      # try pipe
      try(
        df <- utils::read.table(header = TRUE, sep = "|", stringsAsFactors = FALSE),
        silent = TRUE)
    }
  }
  
  # check integrity
  check_validity_mo_source(df)
  
  df <- subset(df, !is.na(mo))
  
  # keep only first two columns, second must be mo
  if (colnames(df)[1] == "mo") {
    df <- df[, c(colnames(df)[2], "mo")]
  } else {
    df <- df[, c(colnames(df)[1], "mo")]
  }
  
  df <- as.data.frame(df, stringAsFactors = FALSE)
  df[, "mo"] <- set_clean_class(df[, "mo", drop = TRUE], c("mo", "character"))
  
  # success
  if (file.exists(mo_source_destination)) {
    action <- "Updated"
  } else {
    action <- "Created"
    # only ask when file is created, not when it is updated
    txt <- paste0(word_wrap(paste0("This will write create the new file '", 
                                   mo_source_destination, 
                                   "', for which your permission is needed.")),
                  "\n\n",
                  word_wrap("Do you agree that this file will be created?"))
    if ("rsasdtudioapi" %in% rownames(utils::installed.packages())) {
      showQuestion <- import_fn("showQuestion", "rstudioapi")
      q_continue <- showQuestion("Create new file in home directory", txt)
    } else {
      q_continue <- utils::menu(choices = c("OK", "Cancel"), graphics = FALSE, title = txt)
    }
    if (q_continue %in% c(FALSE, 2)) {
      return(invisible())
    }
  }
  attr(df, "mo_source_location") <- path
  attr(df, "mo_source_destination") <- mo_source_destination
  attr(df, "mo_source_timestamp") <- file.mtime(path)
  saveRDS(df, mo_source_destination)
  pkg_env$mo_source <- df
  message_(action, " mo_source file '", font_bold(mo_source_destination),
           "' (", formatted_filesize(mo_source_destination),
           ") from '", font_bold(path),
           "' (", formatted_filesize(path),
           '), columns "', colnames(df)[1], '" and "', colnames(df)[2], '"')
}

#' @rdname mo_source
#' @export
get_mo_source <- function(destination = getOption("AMR_mo_source", "~/mo_source.rds")) {
  if (!file.exists(path.expand(destination))) {
    if (interactive()) {
      # source file might have been deleted, so update reference
      set_mo_source("")
    }
    return(NULL)
  }
  if (is.null(pkg_env$mo_source)) {
    pkg_env$mo_source <- readRDS(path.expand(destination))
  }
  
  old_time <- attributes(pkg_env$mo_source)$mo_source_timestamp
  new_time <- file.mtime(attributes(pkg_env$mo_source)$mo_source_location)
  if (interactive() && !identical(old_time, new_time)) {
    # source file was updated, also update reference
    set_mo_source(attributes(pkg_env$mo_source)$mo_source_location)
  }
  pkg_env$mo_source
}

check_validity_mo_source <- function(x, refer_to_name = "`reference_df`", stop_on_error = TRUE) {
  check_dataset_integrity()
  
  if (paste(deparse(substitute(x)), collapse = "") == "get_mo_source()") {
    return(TRUE)
  }
  if (is.null(pkg_env$mo_source) && (identical(x, get_mo_source()))) {
    return(TRUE)
  }
  if (is.null(x)) {
    if (stop_on_error == TRUE) {
      stop_(refer_to_name, " cannot be NULL", call = FALSE)
    } else {
      return(FALSE)
    }
  }
  if (!is.data.frame(x)) {
    if (stop_on_error == TRUE) {
      stop_(refer_to_name, " must be a data.frame", call = FALSE)
    } else {
      return(FALSE)
    }
  }
  if (!"mo" %in% colnames(x)) {
    if (stop_on_error == TRUE) {
      stop_(refer_to_name, " must contain a column 'mo'", call = FALSE)
    } else {
      return(FALSE)
    }
  }
  if (!all(x$mo %in% c("", microorganisms$mo, microorganisms.translation$mo_old), na.rm = TRUE)) {
    if (stop_on_error == TRUE) {
      invalid <- x[which(!x$mo %in% c("", microorganisms$mo, microorganisms.translation$mo_old)), , drop = FALSE]
      if (nrow(invalid) > 1) {
        plural <- "s"
      } else {
        plural <- ""
      }
      stop_("Value", plural, " ", paste0("'", invalid[, 1, drop = TRUE], "'", collapse = ", "), 
           " found in ", tolower(refer_to_name), 
           ", but with invalid microorganism code", plural, " ", paste0("'", invalid$mo, "'", collapse = ", "),
           call = FALSE)
    } else {
      return(FALSE)
    }
  }
  if (colnames(x)[1] != "mo" & nrow(x) > length(unique(x[, 1, drop = TRUE]))) {
    if (stop_on_error == TRUE) {
      stop_(refer_to_name, " contains duplicate values in column '", colnames(x)[1], "'", call = FALSE)
    } else {
      return(FALSE)
    }
  }
  if (colnames(x)[2] != "mo" & nrow(x) > length(unique(x[, 2, drop = TRUE]))) {
    if (stop_on_error == TRUE) {
      stop_(refer_to_name, " contains duplicate values in column '", colnames(x)[2], "'", call = FALSE)
    } else {
      return(FALSE)
    }
  }
  return(TRUE)
}
````
