## Supplementary material for "AMR - An R Package for Working with Antimicrobial Resistance Data": Source package: pca.R

```
# ==================================================================== #
# TITLE                                                                #
# Antimicrobial Resistance (AMR) Analysis for R                        #
#                                                                      #
# SOURCE                                                               #
# https://github.com/msberends/AMR                                     #
#                                                                      #
# LICENCE                                                              #
# (c) 2018-2021 Berends MS, Luz CF et al.                              #
# Developed at the University of Groningen, the Netherlands, in        #
# collaboration with non-profit organisations Certe Medical            #
# Diagnostics & Advice, and University Medical Center Groningen.       # 
#                                                                      #
# This R package is free software; you can freely use and distribute   #
# it for both personal and commercial purposes under the terms of the  #
# GNU General Public License version 2.0 (GNU GPL-2), as published by  #
# the Free Software Foundation.                                        #
# We created this package for both routine data analysis and academic  #
# research and it was publicly released in the hope that it will be    #
# useful, but it comes WITHOUT ANY WARRANTY OR LIABILITY.              #
#                                                                      #
# Visit our website for the full manual and a complete tutorial about  #
# how to conduct AMR analysis: https://msberends.github.io/AMR/        #
# ==================================================================== #

#' Principal Component Analysis (for AMR)
#' 
#' Performs a principal component analysis (PCA) based on a data set with automatic determination for afterwards plotting the groups and labels, and automatic filtering on only suitable (i.e. non-empty and numeric) variables.
#' @inheritSection lifecycle Maturing lifecycle
#' @param x a [data.frame] containing numeric columns
#' @param ... columns of `x` to be selected for PCA, can be unquoted since it supports quasiquotation.
#' @inheritParams stats::prcomp
#' @details The [pca()] function takes a [data.frame] as input and performs the actual PCA with the \R function [prcomp()].
#' 
#' The result of the [pca()] function is a [prcomp] object, with an additional attribute `non_numeric_cols` which is a vector with the column names of all columns that do not contain numeric values. These are probably the groups and labels, and will be used by [ggplot_pca()].
#' @return An object of classes [pca] and [prcomp]
#' @importFrom stats prcomp
#' @export
#' @examples 
#' # `example_isolates` is a dataset available in the AMR package.
#' # See ?example_isolates.
#'
#' \donttest{
#' 
#' if (require("dplyr")) {
#'   # calculate the resistance per group first 
#'   resistance_data <- example_isolates %>% 
#'     group_by(order = mo_order(mo),       # group on anything, like order
#'              genus = mo_genus(mo)) %>%   #  and genus as we do here
#'     summarise_if(is.rsi, resistance)     # then get resistance of all drugs
#'     
#'   # now conduct PCA for certain antimicrobial agents
#'   pca_result <- resistance_data %>%         
#'     pca(AMC, CXM, CTX, CAZ, GEN, TOB, TMP, SXT) 
#'     
#'   pca_result
#'   summary(pca_result)
#'   biplot(pca_result)
#'   ggplot_pca(pca_result) # a new and convenient plot function
#' }
#' }
pca <- function(x,
                ...,
                retx = TRUE,
                center = TRUE, 
                scale. = TRUE,
                tol = NULL,
                rank. = NULL) {
  meet_criteria(x, allow_class = "data.frame")
  meet_criteria(retx, allow_class = "logical", has_length = 1)
  meet_criteria(center, allow_class = "logical", has_length = 1)
  meet_criteria(scale., allow_class = "logical", has_length = 1)
  meet_criteria(tol, allow_class = "numeric", has_length = 1, allow_NULL = TRUE)
  meet_criteria(rank., allow_class = "numeric", has_length = 1, allow_NULL = TRUE)
  
  # unset data.table, tibble, etc.
  # also removes groups made by dplyr::group_by
  x <- as.data.frame(x, stringsAsFactors = FALSE)
  x.bak <- x
  
  # defuse R expressions, this replaces rlang::enquos()
  dots <- substitute(list(...))
  if (length(dots) > 1) {
    new_list <- list(0)
    for (i in seq_len(length(dots) - 1)) {
      new_list[[i]] <- tryCatch(eval(dots[[i + 1]], envir = x),
                                error = function(e) stop(e$message, call. = FALSE))
      if (length(new_list[[i]]) == 1) {
        if (is.character(new_list[[i]]) & new_list[[i]] %in% colnames(x)) {
          # this is to support quoted variables: df %pm>% pca("mycol1", "mycol2")
          new_list[[i]] <- x[, new_list[[i]]]
        } else {
          # remove item - it's a argument like `center`
          new_list[[i]] <- NULL
        }
      }
    }
    
    x <- as.data.frame(new_list, stringsAsFactors = FALSE)
    if (any(vapply(FUN.VALUE = logical(1), x, function(y) !is.numeric(y)))) {
      warning_("Be sure to first calculate the resistance (or susceptibility) of variables with antimicrobial test results, since PCA works with numeric variables only. Please see Examples in ?pca.")
    }
    
    # set column names
    tryCatch(colnames(x) <- as.character(dots)[2:length(dots)],
             error = function(e) warning("column names could not be set"))
    
    # keep only numeric columns
    x <- x[, vapply(FUN.VALUE = logical(1), x, function(y) is.numeric(y))]
    # bind the data set with the non-numeric columns
    x <- cbind(x.bak[, vapply(FUN.VALUE = logical(1), x.bak, function(y) !is.numeric(y) & !all(is.na(y))), drop = FALSE], x)
  }
  
  x <- pm_ungroup(x)  # would otherwise select the grouping vars
  x <- x[rowSums(is.na(x)) == 0, ] # remove columns containing NAs
  
  pca_data <- x[, which(vapply(FUN.VALUE = logical(1), x, function(x) is.numeric(x)))]
  
  message_("Columns selected for PCA: ", paste0(font_bold(colnames(pca_data)), collapse = "/"),
           ". Total observations available: ", nrow(pca_data), ".")
  
  pca_model <- prcomp(pca_data, retx = retx, center = center, scale. = scale., tol = tol, rank. = rank.)
  attr(pca_model, "non_numeric_cols") <- x[, vapply(FUN.VALUE = logical(1), x, function(y) !is.numeric(y) & !all(is.na(y))), drop = FALSE]
  class(pca_model) <- c("pca", class(pca_model))
  pca_model
}
```
