## Supplementary material for "AMR - An R Package for Working with Antimicrobial Resistance Data": Source package: proportion.R

````
# ==================================================================== #
# TITLE                                                                #
# Antimicrobial Resistance (AMR) Analysis for R                        #
#                                                                      #
# SOURCE                                                               #
# https://github.com/msberends/AMR                                     #
#                                                                      #
# LICENCE                                                              #
# (c) 2018-2021 Berends MS, Luz CF et al.                              #
# Developed at the University of Groningen, the Netherlands, in        #
# collaboration with non-profit organisations Certe Medical            #
# Diagnostics & Advice, and University Medical Center Groningen.       # 
#                                                                      #
# This R package is free software; you can freely use and distribute   #
# it for both personal and commercial purposes under the terms of the  #
# GNU General Public License version 2.0 (GNU GPL-2), as published by  #
# the Free Software Foundation.                                        #
# We created this package for both routine data analysis and academic  #
# research and it was publicly released in the hope that it will be    #
# useful, but it comes WITHOUT ANY WARRANTY OR LIABILITY.              #
#                                                                      #
# Visit our website for the full manual and a complete tutorial about  #
# how to conduct AMR analysis: https://msberends.github.io/AMR/        #
# ==================================================================== #

#' Calculate microbial resistance
#'
#' @description These functions can be used to calculate the (co-)resistance or susceptibility of microbial isolates (i.e. percentage of S, SI, I, IR or R). All functions support quasiquotation with pipes, can be used in `summarise()` from the `dplyr` package and also support grouped variables, please see *Examples*.
#'
#' [resistance()] should be used to calculate resistance, [susceptibility()] should be used to calculate susceptibility.\cr
#' @inheritSection lifecycle Stable lifecycle
#' @param ... one or more vectors (or columns) with antibiotic interpretations. They will be transformed internally with [as.rsi()] if needed. Use multiple columns to calculate (the lack of) co-resistance: the probability where one of two drugs have a resistant or susceptible result. See Examples.
#' @param minimum the minimum allowed number of available (tested) isolates. Any isolate count lower than `minimum` will return `NA` with a warning. The default number of `30` isolates is advised by the Clinical and Laboratory Standards Institute (CLSI) as best practice, see Source.
#' @param as_percent a logical to indicate whether the output must be returned as a hundred fold with % sign (a character). A value of `0.123456` will then be returned as `"12.3%"`.
#' @param only_all_tested (for combination therapies, i.e. using more than one variable for `...`): a logical to indicate that isolates must be tested for all antibiotics, see section *Combination therapy* below
#' @param data a [data.frame] containing columns with class [`rsi`] (see [as.rsi()])
#' @param translate_ab a column name of the [antibiotics] data set to translate the antibiotic abbreviations to, using [ab_property()]
#' @inheritParams ab_property
#' @param combine_SI a logical to indicate whether all values of S and I must be merged into one, so the output only consists of S+I vs. R (susceptible vs. resistant). This used to be the argument `combine_IR`, but this now follows the redefinition by EUCAST about the interpretation of I (increased exposure) in 2019, see section 'Interpretation of S, I and R' below. Default is `TRUE`.
#' @param combine_IR a logical to indicate whether all values of I and R must be merged into one, so the output only consists of S vs. I+R (susceptible vs. non-susceptible). This is outdated, see argument `combine_SI`.
#' @inheritSection as.rsi Interpretation of R and S/I
#' @details
#' The function [resistance()] is equal to the function [proportion_R()]. The function [susceptibility()] is equal to the function [proportion_SI()].
#'  
#' **Remember that you should filter your table to let it contain only first isolates!** This is needed to exclude duplicates and to reduce selection bias. Use [first_isolate()] to determine them in your data set.
#'
#' These functions are not meant to count isolates, but to calculate the proportion of resistance/susceptibility. Use the [`count()`][AMR::count()] functions to count isolates. The function [susceptibility()] is essentially equal to `count_susceptible() / count_all()`. *Low counts can influence the outcome - the `proportion` functions may camouflage this, since they only return the proportion (albeit being dependent on the `minimum` argument).*
#'
#' The function [proportion_df()] takes any variable from `data` that has an [`rsi`] class (created with [as.rsi()]) and calculates the proportions R, I and S. It also supports grouped variables. The function [rsi_df()] works exactly like [proportion_df()], but adds the number of isolates.
#' @section Combination therapy:
#' When using more than one variable for `...` (= combination therapy), use `only_all_tested` to only count isolates that are tested for all antibiotics/variables that you test them for. See this example for two antibiotics, Drug A and Drug B, about how [susceptibility()] works to calculate the %SI:
#'
#' ```
#' --------------------------------------------------------------------
#'                     only_all_tested = FALSE  only_all_tested = TRUE
#'                     -----------------------  -----------------------
#'  Drug A    Drug B   include as  include as   include as  include as
#'                     numerator   denominator  numerator   denominator
#' --------  --------  ----------  -----------  ----------  -----------
#'  S or I    S or I       X            X            X            X
#'    R       S or I       X            X            X            X
#'   <NA>     S or I       X            X            -            -
#'  S or I      R          X            X            X            X
#'    R         R          -            X            -            X
#'   <NA>       R          -            -            -            -
#'  S or I     <NA>        X            X            -            -
#'    R        <NA>        -            -            -            -
#'   <NA>      <NA>        -            -            -            -
#' --------------------------------------------------------------------
#' ```
#'
#' Please note that, in combination therapies, for `only_all_tested = TRUE` applies that:
#' ```
#'     count_S()    +   count_I()    +   count_R()    = count_all()
#'   proportion_S() + proportion_I() + proportion_R() = 1
#' ```
#' and that, in combination therapies, for `only_all_tested = FALSE` applies that:
#' ```
#'     count_S()    +   count_I()    +   count_R()    >= count_all()
#'   proportion_S() + proportion_I() + proportion_R() >= 1
#' ```
#'
#' Using `only_all_tested` has no impact when only using one antibiotic as input.
#' @source **M39 Analysis and Presentation of Cumulative Antimicrobial Susceptibility Test Data, 4th Edition**, 2014, *Clinical and Laboratory Standards Institute (CLSI)*. <https://clsi.org/standards/products/microbiology/documents/m39/>.
#' @seealso [AMR::count()] to count resistant and susceptible isolates.
#' @return A [double] or, when `as_percent = TRUE`, a [character].
#' @rdname proportion
#' @aliases portion
#' @name proportion
#' @export
#' @inheritSection AMR Read more on our website!
#' @examples
#' # example_isolates is a data set available in the AMR package.
#' ?example_isolates
#' 
#' resistance(example_isolates$AMX)     # determines %R
#' susceptibility(example_isolates$AMX) # determines %S+I
#'
#' # be more specific
#' proportion_S(example_isolates$AMX)
#' proportion_SI(example_isolates$AMX)
#' proportion_I(example_isolates$AMX)
#' proportion_IR(example_isolates$AMX)
#' proportion_R(example_isolates$AMX)
#'
#' if (require("dplyr")) {
#'   example_isolates %>%
#'     group_by(hospital_id) %>%
#'     summarise(r = resistance(CIP),
#'               n = n_rsi(CIP)) # n_rsi works like n_distinct in dplyr, see ?n_rsi
#'  
#'   example_isolates %>%
#'     group_by(hospital_id) %>%
#'     summarise(R  = resistance(CIP, as_percent = TRUE),
#'               SI = susceptibility(CIP, as_percent = TRUE),
#'               n1 = count_all(CIP),  # the actual total; sum of all three
#'               n2 = n_rsi(CIP),      # same - analogous to n_distinct
#'               total = n())          # NOT the number of tested isolates!
#'  
#'   # Calculate co-resistance between amoxicillin/clav acid and gentamicin,
#'   # so we can see that combination therapy does a lot more than mono therapy:
#'   example_isolates %>% susceptibility(AMC)  # %SI = 76.3%
#'   example_isolates %>% count_all(AMC)       #   n = 1879
#'  
#'   example_isolates %>% susceptibility(GEN)  # %SI = 75.4%
#'   example_isolates %>% count_all(GEN)       #   n = 1855
#'  
#'   example_isolates %>% susceptibility(AMC, GEN) # %SI = 94.1%
#'   example_isolates %>% count_all(AMC, GEN)      #   n = 1939
#'  
#'  
#'   # See Details on how `only_all_tested` works. Example:
#'   example_isolates %>%
#'     summarise(numerator = count_susceptible(AMC, GEN),
#'               denominator = count_all(AMC, GEN),
#'               proportion = susceptibility(AMC, GEN))
#'  
#'   example_isolates %>%
#'     summarise(numerator = count_susceptible(AMC, GEN, only_all_tested = TRUE),
#'               denominator = count_all(AMC, GEN, only_all_tested = TRUE),
#'               proportion = susceptibility(AMC, GEN, only_all_tested = TRUE))
#'  
#'  
#'   example_isolates %>%
#'     group_by(hospital_id) %>%
#'     summarise(cipro_p = susceptibility(CIP, as_percent = TRUE),
#'               cipro_n = count_all(CIP),
#'               genta_p = susceptibility(GEN, as_percent = TRUE),
#'               genta_n = count_all(GEN),
#'               combination_p = susceptibility(CIP, GEN, as_percent = TRUE),
#'               combination_n = count_all(CIP, GEN))
#'  
#'   # Get proportions S/I/R immediately of all rsi columns
#'   example_isolates %>%
#'     select(AMX, CIP) %>%
#'     proportion_df(translate = FALSE)
#'  
#'   # It also supports grouping variables
#'   example_isolates %>%
#'     select(hospital_id, AMX, CIP) %>%
#'     group_by(hospital_id) %>%
#'     proportion_df(translate = FALSE)
#' }
resistance <- function(...,
                       minimum = 30,
                       as_percent = FALSE,
                       only_all_tested = FALSE) {
  rsi_calc(...,
           ab_result = "R",
           minimum = minimum,
           as_percent = as_percent,
           only_all_tested = only_all_tested,
           only_count = FALSE)
}

#' @rdname proportion
#' @export
susceptibility <- function(...,
                           minimum = 30,
                           as_percent = FALSE,
                           only_all_tested = FALSE) {
  rsi_calc(...,
           ab_result = c("S", "I"),
           minimum = minimum,
           as_percent = as_percent,
           only_all_tested = only_all_tested,
           only_count = FALSE)
}

#' @rdname proportion
#' @export
proportion_R <- function(...,
                         minimum = 30,
                         as_percent = FALSE,
                         only_all_tested = FALSE) {
  rsi_calc(...,
           ab_result = "R",
           minimum = minimum,
           as_percent = as_percent,
           only_all_tested = only_all_tested,
           only_count = FALSE)
}

#' @rdname proportion
#' @export
proportion_IR <- function(...,
                          minimum = 30,
                          as_percent = FALSE,
                          only_all_tested = FALSE) {
  rsi_calc(...,
           ab_result = c("I", "R"),
           minimum = minimum,
           as_percent = as_percent,
           only_all_tested = only_all_tested,
           only_count = FALSE)
}

#' @rdname proportion
#' @export
proportion_I <- function(...,
                         minimum = 30,
                         as_percent = FALSE,
                         only_all_tested = FALSE) {
  rsi_calc(...,
           ab_result = "I",
           minimum = minimum,
           as_percent = as_percent,
           only_all_tested = only_all_tested,
           only_count = FALSE)
}

#' @rdname proportion
#' @export
proportion_SI <- function(...,
                          minimum = 30,
                          as_percent = FALSE,
                          only_all_tested = FALSE) {
  rsi_calc(...,
           ab_result = c("S", "I"),
           minimum = minimum,
           as_percent = as_percent,
           only_all_tested = only_all_tested,
           only_count = FALSE)
}

#' @rdname proportion
#' @export
proportion_S <- function(...,
                         minimum = 30,
                         as_percent = FALSE,
                         only_all_tested = FALSE) {
  rsi_calc(...,
           ab_result = "S",
           minimum = minimum,
           as_percent = as_percent,
           only_all_tested = only_all_tested,
           only_count = FALSE)
}

#' @rdname proportion
#' @export
proportion_df <- function(data,
                          translate_ab = "name",
                          language = get_locale(),
                          minimum = 30,
                          as_percent = FALSE,
                          combine_SI = TRUE,
                          combine_IR = FALSE) {
  rsi_calc_df(type = "proportion",
              data = data,
              translate_ab = translate_ab,
              language = language,
              minimum = minimum,
              as_percent = as_percent,
              combine_SI = combine_SI,
              combine_IR = combine_IR,
              combine_SI_missing = missing(combine_SI))
}
````
