## Supplementary material for "AMR - An R Package for Working with Antimicrobial Resistance Data": Source package: random.R

```
# ==================================================================== #
# TITLE                                                                #
# Antimicrobial Resistance (AMR) Analysis for R                        #
#                                                                      #
# SOURCE                                                               #
# https://github.com/msberends/AMR                                     #
#                                                                      #
# LICENCE                                                              #
# (c) 2018-2021 Berends MS, Luz CF et al.                              #
# Developed at the University of Groningen, the Netherlands, in        #
# collaboration with non-profit organisations Certe Medical            #
# Diagnostics & Advice, and University Medical Center Groningen.       # 
#                                                                      #
# This R package is free software; you can freely use and distribute   #
# it for both personal and commercial purposes under the terms of the  #
# GNU General Public License version 2.0 (GNU GPL-2), as published by  #
# the Free Software Foundation.                                        #
# We created this package for both routine data analysis and academic  #
# research and it was publicly released in the hope that it will be    #
# useful, but it comes WITHOUT ANY WARRANTY OR LIABILITY.              #
#                                                                      #
# Visit our website for the full manual and a complete tutorial about  #
# how to conduct AMR analysis: https://msberends.github.io/AMR/        #
# ==================================================================== #

#' Random MIC values/disk zones/RSI generation
#'
#' These functions can be used for generating random MIC values and disk diffusion diameters, for AMR analysis practice.
#' @inheritSection lifecycle Maturing lifecycle 
#' @param size desired size of the returned vector
#' @param mo any character that can be coerced to a valid microorganism code with [as.mo()]
#' @param ab any character that can be coerced to a valid antimicrobial agent code with [as.ab()]
#' @param prob_RSI a vector of length 3: the probabilities for R (1st value), S (2nd value) and I (3rd value)
#' @param ... extension for future versions, not used at the moment
#' @details The base R function [sample()] is used for generating values.
#' 
#' Generated values are based on the latest EUCAST guideline implemented in the [rsi_translation] data set. To create specific generated values per bug or drug, set the `mo` and/or `ab` argument.
#' @return class `<mic>` for [random_mic()] (see [as.mic()]) and class `<disk>` for [random_disk()] (see [as.disk()])
#' @name random
#' @rdname random
#' @export
#' @inheritSection AMR Read more on our website!
#' @examples
#' random_mic(100)
#' random_disk(100)
#' random_rsi(100)
#' 
#' \donttest{
#' # make the random generation more realistic by setting a bug and/or drug:
#' random_mic(100, "Klebsiella pneumoniae")                 # range 0.0625-64
#' random_mic(100, "Klebsiella pneumoniae", "meropenem")    # range 0.0625-16
#' random_mic(100, "Streptococcus pneumoniae", "meropenem") # range 0.0625-4
#' 
#' random_disk(100, "Klebsiella pneumoniae")                  # range 11-50
#' random_disk(100, "Klebsiella pneumoniae", "ampicillin")    # range 6-14
#' random_disk(100, "Streptococcus pneumoniae", "ampicillin") # range 16-22
#' }
random_mic <- function(size, mo = NULL, ab = NULL, ...) {
  random_exec("MIC", size = size, mo = mo, ab = ab)
}

#' @rdname random
#' @export
random_disk <- function(size, mo = NULL, ab = NULL, ...) {
  random_exec("DISK", size = size, mo = mo, ab = ab)
}

#' @rdname random
#' @export
random_rsi <- function(size, prob_RSI = c(0.33, 0.33, 0.33), ...) {
  sample(as.rsi(c("R", "S", "I")), size = size, replace = TRUE, prob = prob_RSI)
}

random_exec <- function(type, size, mo = NULL, ab = NULL) {
  df <- rsi_translation %pm>%
    pm_filter(guideline %like% "EUCAST") %pm>%
    pm_arrange(pm_desc(guideline)) %pm>%
    subset(guideline == max(guideline) &
              method == type)
  
  if (!is.null(mo)) {
    mo_coerced <- as.mo(mo)
    mo_include <- c(mo_coerced,
                    as.mo(mo_genus(mo_coerced)),
                    as.mo(mo_family(mo_coerced)),
                    as.mo(mo_order(mo_coerced)))
    df_new <- df %pm>%
      subset(mo %in% mo_include)
    if (nrow(df_new) > 0) {
      df <- df_new
    } else {
      warning_("No rows found that match mo '", mo, "', ignoring argument `mo`", call = FALSE)
    }
  }
  
  if (!is.null(ab)) {
    ab_coerced <- as.ab(ab)
    df_new <- df %pm>%
      subset(ab %in% ab_coerced)
    if (nrow(df_new) > 0) {
      df <- df_new
    } else {
      warning_("No rows found that match ab '", ab, "', ignoring argument `ab`", call = FALSE)
    }
  }
  
  if (type == "MIC") {
    # all valid MIC levels
    valid_range <- as.mic(levels(as.mic(1)))
    set_range_max <- max(df$breakpoint_R)
    if (log(set_range_max, 2) %% 1 == 0) {
      # return powers of 2
      valid_range <- unique(as.double(valid_range))
      # add one higher MIC level to set_range_max
      set_range_max <- 2 ^ (log(set_range_max, 2) + 1)
      set_range <- as.mic(valid_range[log(valid_range, 2) %% 1 == 0 & valid_range <= set_range_max])
    } else {
      # no power of 2, return factors of 2 to left and right side
      valid_mics <- suppressWarnings(as.mic(set_range_max / (2 ^ c(-3:3))))
      set_range <- valid_mics[!is.na(valid_mics)]
    }
    return(as.mic(sample(set_range, size = size, replace = TRUE)))
  } else if (type == "DISK") {
    set_range <- seq(from = as.integer(min(df$breakpoint_R)),
                     to = as.integer(max(df$breakpoint_S)),
                     by = 1)
    out <- sample(set_range, size = size, replace = TRUE)
    out[out < 6] <- sample(c(6:10), length(out[out < 6]), replace = TRUE)
    out[out > 50] <- sample(c(40:50), length(out[out > 50]), replace = TRUE)
    return(as.disk(out))
  }
}
```
