## Supplementary material for "AMR - An R Package for Working with Antimicrobial Resistance Data": Source package: resistance_predict.R

```
# ==================================================================== #
# TITLE                                                                #
# Antimicrobial Resistance (AMR) Analysis for R                        #
#                                                                      #
# SOURCE                                                               #
# https://github.com/msberends/AMR                                     #
#                                                                      #
# LICENCE                                                              #
# (c) 2018-2021 Berends MS, Luz CF et al.                              #
# Developed at the University of Groningen, the Netherlands, in        #
# collaboration with non-profit organisations Certe Medical            #
# Diagnostics & Advice, and University Medical Center Groningen.       # 
#                                                                      #
# This R package is free software; you can freely use and distribute   #
# it for both personal and commercial purposes under the terms of the  #
# GNU General Public License version 2.0 (GNU GPL-2), as published by  #
# the Free Software Foundation.                                        #
# We created this package for both routine data analysis and academic  #
# research and it was publicly released in the hope that it will be    #
# useful, but it comes WITHOUT ANY WARRANTY OR LIABILITY.              #
#                                                                      #
# Visit our website for the full manual and a complete tutorial about  #
# how to conduct AMR analysis: https://msberends.github.io/AMR/        #
# ==================================================================== #

#' Predict antimicrobial resistance
#'
#' Create a prediction model to predict antimicrobial resistance for the next years on statistical solid ground. Standard errors (SE) will be returned as columns `se_min` and `se_max`. See *Examples* for a real live example.
#' @inheritSection lifecycle Maturing lifecycle
#' @param col_ab column name of `x` containing antimicrobial interpretations (`"R"`, `"I"` and `"S"`)
#' @param col_date column name of the date, will be used to calculate years if this column doesn't consist of years already, defaults to the first column of with a date class
#' @param year_min lowest year to use in the prediction model, dafaults to the lowest year in `col_date`
#' @param year_max highest year to use in the prediction model, defaults to 10 years after today
#' @param year_every unit of sequence between lowest year found in the data and `year_max`
#' @param minimum minimal amount of available isolates per year to include. Years containing less observations will be estimated by the model.
#' @param model the statistical model of choice. This could be a generalised linear regression model with binomial distribution (i.e. using `glm(..., family = binomial)``, assuming that a period of zero resistance was followed by a period of increasing resistance leading slowly to more and more resistance. See Details for all valid options.
#' @param I_as_S a logical to indicate whether values `"I"` should be treated as `"S"` (will otherwise be treated as `"R"`). The default, `TRUE`, follows the redefinition by EUCAST about the interpretation of I (increased exposure) in 2019, see section *Interpretation of S, I and R* below. 
#' @param preserve_measurements a logical to indicate whether predictions of years that are actually available in the data should be overwritten by the original data. The standard errors of those years will be `NA`.
#' @param info a logical to indicate whether textual analysis should be printed with the name and [summary()] of the statistical model.
#' @param main title of the plot
#' @param ribbon a logical to indicate whether a ribbon should be shown (default) or error bars
#' @param ... arguments passed on to functions
#' @inheritSection as.rsi Interpretation of R and S/I
#' @inheritParams first_isolate
#' @inheritParams graphics::plot
#' @details Valid options for the statistical model (argument `model`) are:
#' - `"binomial"` or `"binom"` or `"logit"`: a generalised linear regression model with binomial distribution
#' - `"loglin"` or `"poisson"`: a generalised log-linear regression model with poisson distribution
#' - `"lin"` or `"linear"`: a linear regression model
#' @return A [data.frame] with extra class [`resistance_predict`] with columns:
#' - `year`
#' - `value`, the same as `estimated` when `preserve_measurements = FALSE`, and a combination of `observed` and `estimated` otherwise
#' - `se_min`, the lower bound of the standard error with a minimum of `0` (so the standard error will never go below 0%)
#' - `se_max` the upper bound of the standard error with a maximum of `1` (so the standard error will never go above 100%)
#' - `observations`, the total number of available observations in that year, i.e. \eqn{S + I + R}
#' - `observed`, the original observed resistant percentages
#' - `estimated`, the estimated resistant percentages, calculated by the model
#'   
#' Furthermore, the model itself is available as an attribute: `attributes(x)$model`, please see *Examples*.
#' @seealso The [proportion()] functions to calculate resistance
#' 
#' Models: [lm()] [glm()]
#' @rdname resistance_predict
#' @export
#' @importFrom stats predict glm lm
#' @inheritSection AMR Read more on our website!
#' @examples
#' x <- resistance_predict(example_isolates, 
#'                         col_ab = "AMX",
#'                         year_min = 2010,
#'                         model = "binomial")
#' plot(x)
#' if (require("ggplot2")) {
#'   ggplot_rsi_predict(x)
#' }
#'
#' # using dplyr:
#' if (require("dplyr")) {
#'   x <- example_isolates %>%
#'     filter_first_isolate() %>%
#'     filter(mo_genus(mo) == "Staphylococcus") %>%
#'     resistance_predict("PEN", model = "binomial")
#'   plot(x)
#'
#'   # get the model from the object
#'   mymodel <- attributes(x)$model
#'   summary(mymodel)
#' }
#'
#' # create nice plots with ggplot2 yourself
#' if (require("dplyr") & require("ggplot2")) {
#' 
#'   data <- example_isolates %>%
#'     filter(mo == as.mo("E. coli")) %>%
#'     resistance_predict(col_ab = "AMX",
#'                        col_date = "date",
#'                        model = "binomial",
#'                        info = FALSE,
#'                        minimum = 15)
#'
#'   ggplot(data,
#'          aes(x = year)) +
#'     geom_col(aes(y = value),
#'              fill = "grey75") +
#'     geom_errorbar(aes(ymin = se_min,
#'                       ymax = se_max),
#'                   colour = "grey50") +
#'     scale_y_continuous(limits = c(0, 1),
#'                        breaks = seq(0, 1, 0.1),
#'                        labels = paste0(seq(0, 100, 10), "%")) +
#'     labs(title = expression(paste("Forecast of Amoxicillin Resistance in ",
#'                                   italic("E. coli"))),
#'          y = "%R",
#'          x = "Year") +
#'     theme_minimal(base_size = 13)
#' }
resistance_predict <- function(x,
                               col_ab,
                               col_date = NULL,
                               year_min = NULL,
                               year_max = NULL,
                               year_every = 1,
                               minimum = 30,
                               model = NULL,
                               I_as_S = TRUE,
                               preserve_measurements = TRUE,
                               info = interactive(),
                               ...) {
  meet_criteria(x, allow_class = "data.frame")
  meet_criteria(col_ab, allow_class = "character", has_length = 1, is_in = colnames(x))
  meet_criteria(col_date, allow_class = "character", has_length = 1, is_in = colnames(x), allow_NULL = TRUE)
  meet_criteria(year_min, allow_class = c("numeric", "integer"), has_length = 1, allow_NULL = TRUE)
  meet_criteria(year_max, allow_class = c("numeric", "integer"), has_length = 1, allow_NULL = TRUE)
  meet_criteria(year_every, allow_class = c("numeric", "integer"), has_length = 1)
  meet_criteria(minimum, allow_class = c("numeric", "integer"), has_length = 1)
  meet_criteria(model, allow_class = c("character", "function"), has_length = 1, allow_NULL = TRUE)
  meet_criteria(I_as_S, allow_class = "logical", has_length = 1)
  meet_criteria(preserve_measurements, allow_class = "logical", has_length = 1)
  meet_criteria(info, allow_class = "logical", has_length = 1)
  
  stop_if(is.null(model), 'choose a regression model with the `model` argument, e.g. resistance_predict(..., model = "binomial")')

  dots <- unlist(list(...))
  if (length(dots) != 0) {
    # backwards compatibility with old arguments
    dots.names <- dots %pm>% names()
    if ("tbl" %in% dots.names) {
      x <- dots[which(dots.names == "tbl")]
    }
    if ("I_as_R" %in% dots.names) {
      warning_("`I_as_R is deprecated - use I_as_S instead.", call = FALSE)
    }
  }
  
  # -- date
  if (is.null(col_date)) {
    col_date <- search_type_in_df(x = x, type = "date")
    stop_if(is.null(col_date), "`col_date` must be set")
  }
  stop_ifnot(col_date %in% colnames(x),
             "column '", col_date, "' not found")
  
  # no grouped tibbles
  x <- as.data.frame(x, stringsAsFactors = FALSE)
  
  year <- function(x) {
    # don't depend on lubridate or so, would be overkill for only this function
    if (all(grepl("^[0-9]{4}$", x))) {
      as.integer(x)
    } else {
      as.integer(format(as.Date(x), "%Y"))
    }
  }
  
  df <- x
  df[, col_ab] <- droplevels(as.rsi(df[, col_ab, drop = TRUE]))
  if (I_as_S == TRUE) {
    # then I as S
    df[, col_ab] <- gsub("I", "S", df[, col_ab, drop = TRUE])
  } else {
    # then I as R
    df[, col_ab] <- gsub("I", "R", df[, col_ab, drop = TRUE])
  }
  df[, col_ab] <- ifelse(is.na(df[, col_ab, drop = TRUE]), 0, df[, col_ab, drop = TRUE])
  
  # remove rows with NAs
  df <- subset(df, !is.na(df[, col_ab, drop = TRUE]))
  df$year <- year(df[, col_date, drop = TRUE])
  df <- as.data.frame(rbind(table(df[, c("year", col_ab)])),
                      stringsAsFactors = FALSE)
  df$year <- as.integer(rownames(df))
  rownames(df) <- NULL
  
  df <- subset(df, sum(df$R + df$S, na.rm = TRUE) >= minimum)
  # nolint start
  df_matrix <- as.matrix(df[, c("R", "S"), drop = FALSE])
  # nolint end
  
  stop_if(NROW(df) == 0, "there are no observations")
  
  year_lowest <- min(df$year)
  if (is.null(year_min)) {
    year_min <- year_lowest
  } else {
    year_min <- max(year_min, year_lowest, na.rm = TRUE)
  }
  if (is.null(year_max)) {
    year_max <- year(Sys.Date()) + 10
  }
  
  years <- list(year = seq(from = year_min, to = year_max, by = year_every))
  
  if (model %in% c("binomial", "binom", "logit")) {
    model <- "binomial"
    model_lm <- with(df, glm(df_matrix ~ year, family = binomial))
    if (info == TRUE) {
      cat("\nLogistic regression model (logit) with binomial distribution")
      cat("\n------------------------------------------------------------\n")
      print(summary(model_lm))
    }
    
    predictmodel <- predict(model_lm, newdata = years, type = "response", se.fit = TRUE)
    prediction <- predictmodel$fit
    se <- predictmodel$se.fit
    
  } else if (model %in% c("loglin", "poisson")) {
    model <- "poisson"
    model_lm <- with(df, glm(R ~ year, family = poisson))
    if (info == TRUE) {
      cat("\nLog-linear regression model (loglin) with poisson distribution")
      cat("\n--------------------------------------------------------------\n")
      print(summary(model_lm))
    }
    
    predictmodel <- predict(model_lm, newdata = years, type = "response", se.fit = TRUE)
    prediction <- predictmodel$fit
    se <- predictmodel$se.fit
    
  } else if (model %in% c("lin", "linear")) {
    model <- "linear"
    model_lm <- with(df, lm((R / (R + S)) ~ year))
    if (info == TRUE) {
      cat("\nLinear regression model")
      cat("\n-----------------------\n")
      print(summary(model_lm))
    }
    
    predictmodel <- predict(model_lm, newdata = years, se.fit = TRUE)
    prediction <- predictmodel$fit
    se <- predictmodel$se.fit
    
  } else {
    stop("no valid model selected. See ?resistance_predict.")
  }
  
  # prepare the output dataframe
  df_prediction <- data.frame(year = unlist(years),
                              value = prediction,
                              se_min = prediction - se,
                              se_max = prediction + se,
                              stringsAsFactors = FALSE)
  
  if (model == "poisson") {
    df_prediction$value <- as.integer(format(df_prediction$value, scientific = FALSE))
    df_prediction$se_min <- as.integer(df_prediction$se_min)
    df_prediction$se_max <- as.integer(df_prediction$se_max)
    
  } else {
    # se_max not above 1
    df_prediction$se_max <- ifelse(df_prediction$se_max > 1, 1, df_prediction$se_max)
  }
  # se_min not below 0
  df_prediction$se_min <- ifelse(df_prediction$se_min < 0, 0, df_prediction$se_min)
  
  df_observations <- data.frame(year = df$year,
                                observations = df$R + df$S,
                                observed = df$R / (df$R + df$S),
                                stringsAsFactors = FALSE)
  df_prediction <- df_prediction %pm>%
    pm_left_join(df_observations, by = "year")
  df_prediction$estimated <- df_prediction$value
  
  if (preserve_measurements == TRUE) {
    # replace estimated data by observed data
    df_prediction$value <- ifelse(!is.na(df_prediction$observed), df_prediction$observed, df_prediction$value)
    df_prediction$se_min <- ifelse(!is.na(df_prediction$observed), NA, df_prediction$se_min)
    df_prediction$se_max <- ifelse(!is.na(df_prediction$observed), NA, df_prediction$se_max)
  }
  
  df_prediction$value <- ifelse(df_prediction$value > 1, 1, ifelse(df_prediction$value < 0, 0, df_prediction$value))
  df_prediction <- df_prediction[order(df_prediction$year), ]
  
  structure(
    .Data = df_prediction,
    class = c("resistance_predict", "data.frame"),
    I_as_S = I_as_S,
    model_title = model,
    model = model_lm,
    ab = col_ab
  )
}

#' @rdname resistance_predict
#' @export
rsi_predict <- resistance_predict

#' @method plot resistance_predict
#' @export
#' @importFrom graphics plot axis arrows points
#' @rdname resistance_predict
plot.resistance_predict <- function(x, main = paste("Resistance Prediction of", x_name), ...) {
  x_name <- paste0(ab_name(attributes(x)$ab), " (", attributes(x)$ab, ")")
  meet_criteria(main, allow_class = "character", has_length = 1)
  
  if (attributes(x)$I_as_S == TRUE) {
    ylab <- "%R"
  } else {
    ylab <- "%IR"
  }

  plot(x = x$year,
       y = x$value,
       ylim = c(0, 1),
       yaxt = "n", # no y labels
       pch = 19, # closed dots
       ylab = paste0("Percentage (", ylab, ")"),
       xlab = "Year",
       main = main,
       sub = paste0("(n = ", sum(x$observations, na.rm = TRUE),
                    ", model: ", attributes(x)$model_title, ")"),
       cex.sub = 0.75)
  
  
  axis(side = 2, at = seq(0, 1, 0.1), labels = paste0(0:10 * 10, "%"))
  
  # hack for error bars: https://stackoverflow.com/a/22037078/4575331
  arrows(x0 = x$year,
         y0 = x$se_min,
         x1 = x$year,
         y1 = x$se_max,
         length = 0.05, angle = 90, code = 3, lwd = 1.5)
  
  # overlay grey points for prediction
  points(x = subset(x, is.na(observations))$year,
         y = subset(x, is.na(observations))$value,
         pch = 19,
         col = "grey40")
}

#' @rdname resistance_predict
#' @export
ggplot_rsi_predict <- function(x,
                               main = paste("Resistance Prediction of", x_name),
                               ribbon = TRUE,
                               ...) {
  x_name <- paste0(ab_name(attributes(x)$ab), " (", attributes(x)$ab, ")")
  meet_criteria(main, allow_class = "character", has_length = 1)
  meet_criteria(ribbon, allow_class = "logical", has_length = 1)
  
  stop_ifnot_installed("ggplot2")
  stop_ifnot(inherits(x, "resistance_predict"), "`x` must be a resistance prediction model created with resistance_predict()")
  
  
  if (attributes(x)$I_as_S == TRUE) {
    ylab <- "%R"
  } else {
    ylab <- "%IR"
  }
  
  p <- ggplot2::ggplot(x, ggplot2::aes(x = year, y = value)) +
    ggplot2::geom_point(data = subset(x, !is.na(observations)),
                        size = 2) +
    scale_y_percent(limits = c(0, 1)) +
    ggplot2::labs(title = main,
                  y = paste0("Percentage (", ylab, ")"),
                  x = "Year",
                  caption = paste0("(n = ", sum(x$observations, na.rm = TRUE),
                                   ", model: ", attributes(x)$model_title, ")"))
  
  if (ribbon == TRUE) {
    p <- p + ggplot2::geom_ribbon(ggplot2::aes(ymin = se_min, ymax = se_max), alpha = 0.25)
  } else {
    p <- p + ggplot2::geom_errorbar(ggplot2::aes(ymin = se_min, ymax = se_max), na.rm = TRUE, width = 0.5)
  }
  p <- p +
    # overlay grey points for prediction
    ggplot2::geom_point(data = subset(x, is.na(observations)),
                        size = 2,
                        colour = "grey40")
  p
}
```
