## Supplementary material for "AMR - An R Package for Working with Antimicrobial Resistance Data": Source package: rsi.R

````
# ==================================================================== #
# TITLE                                                                #
# Antimicrobial Resistance (AMR) Analysis for R                        #
#                                                                      #
# SOURCE                                                               #
# https://github.com/msberends/AMR                                     #
#                                                                      #
# LICENCE                                                              #
# (c) 2018-2021 Berends MS, Luz CF et al.                              #
# Developed at the University of Groningen, the Netherlands, in        #
# collaboration with non-profit organisations Certe Medical            #
# Diagnostics & Advice, and University Medical Center Groningen.       # 
#                                                                      #
# This R package is free software; you can freely use and distribute   #
# it for both personal and commercial purposes under the terms of the  #
# GNU General Public License version 2.0 (GNU GPL-2), as published by  #
# the Free Software Foundation.                                        #
# We created this package for both routine data analysis and academic  #
# research and it was publicly released in the hope that it will be    #
# useful, but it comes WITHOUT ANY WARRANTY OR LIABILITY.              #
#                                                                      #
# Visit our website for the full manual and a complete tutorial about  #
# how to conduct AMR analysis: https://msberends.github.io/AMR/        #
# ==================================================================== #

#' Interpret MIC and disk values, or clean raw R/SI data
#'
#' Interpret minimum inhibitory concentration (MIC) values and disk diffusion diameters according to EUCAST or CLSI, or clean up existing R/SI values. This transforms the input to a new class [`rsi`], which is an ordered factor with levels `S < I < R`. Values that cannot be interpreted will be returned as `NA` with a warning.
#' @inheritSection lifecycle Stable lifecycle
#' @rdname as.rsi
#' @param x vector of values (for class [`mic`]: an MIC value in mg/L, for class [`disk`]: a disk diffusion radius in millimetres)
#' @param mo any (vector of) text that can be coerced to a valid microorganism code with [as.mo()], will be determined automatically if the `dplyr` package is installed
#' @param ab any (vector of) text that can be coerced to a valid antimicrobial code with [as.ab()]
#' @param uti (Urinary Tract Infection) A vector with [logical]s (`TRUE` or `FALSE`) to specify whether a UTI specific interpretation from the guideline should be chosen. For using [as.rsi()] on a [data.frame], this can also be a column containing [logical]s or when left blank, the data set will be searched for a 'specimen' and rows containing 'urin' (such as 'urine', 'urina') in that column will be regarded isolates from a UTI. See *Examples*.
#' @inheritParams first_isolate
#' @param guideline defaults to the latest included EUCAST guideline, see Details for all options
#' @param conserve_capped_values a logical to indicate that MIC values starting with `">"` (but not `">="`) must always return "R" , and that MIC values starting with `"<"` (but not `"<="`) must always return "S"
#' @param add_intrinsic_resistance *(only useful when using a EUCAST guideline)* a logical to indicate whether intrinsic antibiotic resistance must also be considered for applicable bug-drug combinations, meaning that e.g. ampicillin will always return "R" in *Klebsiella* species. Determination is based on the [intrinsic_resistant] data set, that itself is based on `r format_eucast_version_nr(3.2)`.
#' @param reference_data a [data.frame] to be used for interpretation, which defaults to the [rsi_translation] data set. Changing this argument allows for using own interpretation guidelines. This argument must contain a data set that is equal in structure to the [rsi_translation] data set (same column names and column types). Please note that the `guideline` argument will be ignored when `reference_data` is manually set.
#' @param threshold maximum fraction of invalid antimicrobial interpretations of `x`, please see *Examples*
#' @param ... for using on a [data.frame]: names of columns to apply [as.rsi()] on (supports tidy selection like `AMX:VAN`). Otherwise: arguments passed on to methods.
#' @details 
#' ## How it works
#' 
#' The [as.rsi()] function works in four ways:
#' 
#' 1. For **cleaning raw / untransformed data**. The data will be cleaned to only contain values S, I and R and will try its best to determine this with some intelligence. For example, mixed values with R/SI interpretations and MIC values such as `"<0.25; S"` will be coerced to `"S"`. Combined interpretations for multiple test methods (as seen in laboratory records) such as `"S; S"` will be coerced to `"S"`, but a value like `"S; I"` will return `NA` with a warning that the input is unclear.
#' 
#' 2. For **interpreting minimum inhibitory concentration (MIC) values** according to EUCAST or CLSI. You must clean your MIC values first using [as.mic()], that also gives your columns the new data class [`mic`]. Also, be sure to have a column with microorganism names or codes. It will be found automatically, but can be set manually using the `mo` argument.
#'    * Using `dplyr`, R/SI interpretation can be done very easily with either: 
#'      ```
#'      your_data %>% mutate_if(is.mic, as.rsi)             # until dplyr 1.0.0
#'      your_data %>% mutate(across(where(is.mic), as.rsi)) # since dplyr 1.0.0
#'      ```
#'    * Operators like "<=" will be stripped before interpretation. When using `conserve_capped_values = TRUE`, an MIC value of e.g. ">2" will always return "R", even if the breakpoint according to the chosen guideline is ">=4". This is to prevent that capped values from raw laboratory data would not be treated conservatively. The default behaviour (`conserve_capped_values = FALSE`) considers ">2" to be lower than ">=4" and might in this case return "S" or "I".
#'      
#' 3. For **interpreting disk diffusion diameters** according to EUCAST or CLSI. You must clean your disk zones first using [as.disk()], that also gives your columns the new data class [`disk`]. Also, be sure to have a column with microorganism names or codes. It will be found automatically, but can be set manually using the `mo` argument.
#'    * Using `dplyr`, R/SI interpretation can be done very easily with either: 
#'      ```
#'      your_data %>% mutate_if(is.disk, as.rsi)             # until dplyr 1.0.0
#'      your_data %>% mutate(across(where(is.disk), as.rsi)) # since dplyr 1.0.0
#'      ```
#' 
#' 4. For **interpreting a complete data set**, with automatic determination of MIC values, disk diffusion diameters, microorganism names or codes, and antimicrobial test results. This is done very simply by running `as.rsi(data)`.
#' 
#' ## Supported guidelines
#' 
#' For interpreting MIC values as well as disk diffusion diameters, supported guidelines to be used as input for the `guideline` argument are: `r paste0('"', sort(unique(AMR::rsi_translation$guideline)), '"', collapse = ", ")`.
#' 
#' Simply using `"CLSI"` or `"EUCAST"` as input will automatically select the latest version of that guideline. You can set your own data set using the `reference_data` argument. The `guideline` argument will then be ignored.
#' 
#' ## After interpretation
#' 
#' After using [as.rsi()], you can use the [eucast_rules()] defined by EUCAST to (1) apply inferred susceptibility and resistance based on results of other antimicrobials and (2) apply intrinsic resistance based on taxonomic properties of a microorganism.
#' 
#' ## Machine readable interpretation guidelines
#' 
#' The repository of this package [contains a machine readable version](https://github.com/msberends/AMR/blob/master/data-raw/rsi_translation.txt) of all guidelines. This is a CSV file consisting of `r format(nrow(AMR::rsi_translation), big.mark = ",")` rows and `r ncol(AMR::rsi_translation)` columns. This file is machine readable, since it contains one row for every unique combination of the test method (MIC or disk diffusion), the antimicrobial agent and the microorganism. **This allows for easy implementation of these rules in laboratory information systems (LIS)**. Note that it only contains interpretation guidelines for humans - interpretation guidelines from CLSI for animals were removed.
#'
#' ## Other
#'
#' The function [is.rsi.eligible()] returns `TRUE` when a columns contains at most 5% invalid antimicrobial interpretations (not S and/or I and/or R), and `FALSE` otherwise. The threshold of 5% can be set with the `threshold` argument.
#' @section Interpretation of R and S/I:
#' In 2019, the European Committee on Antimicrobial Susceptibility Testing (EUCAST) has decided to change the definitions of susceptibility testing categories R and S/I as shown below (<https://www.eucast.org/newsiandr/>).
#'
#' - **R = Resistant**\cr
#'   A microorganism is categorised as *Resistant* when there is a high likelihood of therapeutic failure even when there is increased exposure. Exposure is a function of how the mode of administration, dose, dosing interval, infusion time, as well as distribution and excretion of the antimicrobial agent will influence the infecting organism at the site of infection.
#' - **S = Susceptible**\cr
#'   A microorganism is categorised as *Susceptible, standard dosing regimen*, when there is a high likelihood of therapeutic success using a standard dosing regimen of the agent.
#' - **I = Increased exposure, but still susceptible**\cr
#'   A microorganism is categorised as *Susceptible, Increased exposure* when there is a high likelihood of therapeutic success because exposure to the agent is increased by adjusting the dosing regimen or by its concentration at the site of infection.
#'
#' This AMR package honours this new insight. Use [susceptibility()] (equal to [proportion_SI()]) to determine antimicrobial susceptibility and [count_susceptible()] (equal to [count_SI()]) to count susceptible isolates.
#' @return Ordered [factor] with new class [`rsi`]
#' @aliases rsi
#' @export
#' @seealso [as.mic()], [as.disk()], [as.mo()]
#' @inheritSection AMR Reference data publicly available
#' @inheritSection AMR Read more on our website!
#' @examples
#' summary(example_isolates) # see all R/SI results at a glance
#' 
#' if (require("skimr")) {
#'   # class <rsi> supported in skim() too:
#'   skim(example_isolates)
#' }
#' 
#' # For INTERPRETING disk diffusion and MIC values -----------------------
#'        
#' # a whole data set, even with combined MIC values and disk zones
#' df <- data.frame(microorganism = "Escherichia coli",
#'                  AMP = as.mic(8),
#'                  CIP = as.mic(0.256),
#'                  GEN = as.disk(18),
#'                  TOB = as.disk(16),
#'                  NIT = as.mic(32),
#'                  ERY = "R")
#' as.rsi(df)
#' 
#' # for single values
#' as.rsi(x = as.mic(2),
#'        mo = as.mo("S. pneumoniae"),
#'        ab = "AMP",
#'        guideline = "EUCAST")
#'
#' as.rsi(x = as.disk(18),
#'        mo = "Strep pneu",  # `mo` will be coerced with as.mo()
#'        ab = "ampicillin",  # and `ab` with as.ab()
#'        guideline = "EUCAST")
#'
#' \donttest{
#' # the dplyr way
#' if (require("dplyr")) {
#'   df %>% mutate_if(is.mic, as.rsi)
#'   df %>% mutate_if(function(x) is.mic(x) | is.disk(x), as.rsi)
#'   df %>% mutate(across(where(is.mic), as.rsi))
#'   df %>% mutate_at(vars(AMP:TOB), as.rsi)
#'   df %>% mutate(across(AMP:TOB, as.rsi))
#'  
#'   df %>%
#'     mutate_at(vars(AMP:TOB), as.rsi, mo = .$microorganism)
#'     
#'   # to include information about urinary tract infections (UTI)
#'   data.frame(mo = "E. coli",
#'              NIT = c("<= 2", 32),
#'              from_the_bladder = c(TRUE, FALSE)) %>%
#'     as.rsi(uti = "from_the_bladder")
#'     
#'   data.frame(mo = "E. coli",
#'              NIT = c("<= 2", 32),
#'              specimen = c("urine", "blood")) %>%
#'     as.rsi() # automatically determines urine isolates
#'   
#'   df %>%
#'     mutate_at(vars(AMP:NIT), as.rsi, mo = "E. coli", uti = TRUE)  
#' }
#'
#' # For CLEANING existing R/SI values ------------------------------------
#' 
#' as.rsi(c("S", "I", "R", "A", "B", "C"))
#' as.rsi("<= 0.002; S") # will return "S"

#' rsi_data <- as.rsi(c(rep("S", 474), rep("I", 36), rep("R", 370)))
#' is.rsi(rsi_data)
#' plot(rsi_data)    # for percentages
#' barplot(rsi_data) # for frequencies
#'
#' # the dplyr way
#' if (require("dplyr")) {
#'   example_isolates %>%
#'     mutate_at(vars(PEN:RIF), as.rsi)
#'   # same:   
#'   example_isolates %>%
#'     as.rsi(PEN:RIF)
#'  
#'   # fastest way to transform all columns with already valid AMR results to class `rsi`:
#'   example_isolates %>%
#'     mutate_if(is.rsi.eligible, as.rsi)
#'     
#'   # note: from dplyr 1.0.0 on, this will be: 
#'   # example_isolates %>%
#'   #   mutate(across(where(is.rsi.eligible), as.rsi))
#' }
#' }
as.rsi <- function(x, ...) {
  UseMethod("as.rsi")
}

#' @rdname as.rsi
#' @export
is.rsi <- function(x) {
  inherits(x, "rsi")
}

#' @rdname as.rsi
#' @export
is.rsi.eligible <- function(x, threshold = 0.05) {
  meet_criteria(threshold, allow_class = "numeric", has_length = 1)
  
  stop_if(NCOL(x) > 1, "`x` must be a one-dimensional vector.")
  if (any(c("logical",
            "numeric",
            "integer",
            "mo",
            "Date",
            "POSIXct",
            "rsi",
            "raw",
            "hms")
          %in% class(x))) {
    # no transformation needed
    FALSE
  } else {
    x <- x[!is.na(x) & !is.null(x) & !identical(x, "")]
    if (length(x) == 0) {
      return(FALSE)
    }
    checked <- suppressWarnings(as.rsi(x))
    outcome <- sum(is.na(checked)) / length(x)
    outcome <= threshold
  }
}

#' @export
as.rsi.default <- function(x, ...) {
  if (is.rsi(x)) {
    return(x)
  }
  
  if (inherits(x, "integer") & all(x %in% c(1:3, NA))) {
    x[x == 1] <- "S"
    x[x == 2] <- "I"
    x[x == 3] <- "R"
    
  } else if (!all(is.na(x)) && !identical(levels(x), c("S", "I", "R"))) {
    
    if (!any(x %like% "(R|S|I)", na.rm = TRUE)) {
      # check if they are actually MICs or disks now that the antibiotic name is valid
      if (all_valid_mics(x)) {
        warning_("The input seems to be MIC values. Transform them with as.mic() before running as.rsi() to interpret them.")
      } else if (all_valid_disks(x)) {
        warning_("The input seems to be disk diffusion values. Transform them with as.disk() before running as.rsi() to interpret them.")
      }
    }
    
    x <- as.character(unlist(x))
    x.bak <- x
    
    na_before <- length(x[is.na(x) | x == ""])
    # remove all spaces
    x <- gsub(" +", "", x)
    # remove all MIC-like values: numbers, operators and periods
    x <- gsub("[0-9.,;:<=>]+", "", x)
    # remove everything between brackets, and 'high' and 'low'
    x <- gsub("([(].*[)])", "", x)
    x <- gsub("(high|low)", "", x, ignore.case = TRUE)
    # disallow more than 3 characters
    x[nchar(x) > 3] <- NA
    # set to capitals
    x <- toupper(x)
    # remove all invalid characters
    x <- gsub("[^RSI]+", "", x)
    # in cases of "S;S" keep S, but in case of "S;I" make it NA
    x <- gsub("^S+$", "S", x)
    x <- gsub("^I+$", "I", x)
    x <- gsub("^R+$", "R", x)
    x[!x %in% c("S", "I", "R")] <- NA
    na_after <- length(x[is.na(x) | x == ""])
    
    if (!isFALSE(list(...)$warn)) { # so as.rsi(..., warn = FALSE) will never throw a warning
      if (na_before != na_after) {
        list_missing <- x.bak[is.na(x) & !is.na(x.bak) & x.bak != ""] %pm>%
          unique() %pm>%
          sort()
        list_missing <- paste0('"', list_missing, '"', collapse = ", ")
        warning_(na_after - na_before, " results truncated (",
                 round(((na_after - na_before) / length(x)) * 100),
                 "%) that were invalid antimicrobial interpretations: ",
                 list_missing, call = FALSE)
      }
    }
  }
  
  set_clean_class(factor(x, levels = c("S", "I", "R"), ordered = TRUE),
                  new_class =  c("rsi", "ordered", "factor"))
}

#' @rdname as.rsi
#' @export
as.rsi.mic <- function(x,
                       mo = NULL, 
                       ab = deparse(substitute(x)), 
                       guideline = "EUCAST", 
                       uti = FALSE,
                       conserve_capped_values = FALSE,
                       add_intrinsic_resistance = FALSE,
                       reference_data = AMR::rsi_translation,
                       ...) {
  meet_criteria(x)
  meet_criteria(mo, allow_class = c("mo", "character"), allow_NULL = TRUE)
  meet_criteria(ab, allow_class = c("ab", "character"))
  meet_criteria(guideline, allow_class = "character", has_length = 1)
  meet_criteria(uti, allow_class = "logical", has_length = c(1, length(x)))
  meet_criteria(conserve_capped_values, allow_class = "logical", has_length = 1)
  meet_criteria(add_intrinsic_resistance, allow_class = "logical", has_length = 1)
  meet_criteria(reference_data, allow_class = "data.frame")
  check_reference_data(reference_data)
  
  # for dplyr's across()
  cur_column_dplyr <- import_fn("cur_column", "dplyr", error_on_fail = FALSE)
  if (!is.null(cur_column_dplyr)) {
    # try to get current column, which will only be available when in across()
    ab <- tryCatch(cur_column_dplyr(),
                   error = function(e) ab)
  }
  
  # for auto-determining mo
  mo_var_found <- ""
  if (is.null(mo)) {
    tryCatch({
      df <- get_current_data(arg_name = "mo", call = -3) # will return an error if not found
      mo <- NULL
      try({
        mo <- suppressMessages(search_type_in_df(df, "mo"))
      }, silent = TRUE)
      if (!is.null(df) && !is.null(mo) && is.data.frame(df)) {
        mo_var_found <- paste0(" based on column '", font_bold(mo), "'")
        mo <- df[, mo, drop = TRUE]
      }
    }, error = function(e) 
      stop_('No information was supplied about the microorganisms (missing argument `mo`). See ?as.rsi.\n\n',
            "To transform certain columns with e.g. mutate_at(), use `data %>% mutate_at(vars(...), as.rsi, mo = .$x)`, where x is your column with microorganisms.\n",
            "To tranform all disk diffusion zones in a data set, use `data %>% as.rsi()` or data %>% mutate_if(is.disk, as.rsi).", call = FALSE)
    )
  }
  if (length(ab) == 1 && ab %like% "as.mic") {
    stop_('No unambiguous name was supplied about the antibiotic (argument `ab`). See ?as.rsi.', call = FALSE)
  }
  
  ab_coerced <- suppressWarnings(as.ab(ab))
  mo_coerced <- suppressWarnings(as.mo(mo))
  guideline_coerced <- get_guideline(guideline, reference_data)
  if (is.na(ab_coerced)) {
    message_("Returning NAs for unknown drug: '", font_bold(ab),
             "'. Rename this column to a drug name or code, and check the output with `as.ab()`.", 
             add_fn = font_red, 
             as_note = FALSE)
    return(as.rsi(rep(NA, length(x))))
  }
  if (length(mo_coerced) == 1) {
    mo_coerced <- rep(mo_coerced, length(x))
  }
  if (length(uti) == 1) {
    uti <- rep(uti, length(x))
  }
  
  message_("=> Interpreting MIC values of ", ifelse(isTRUE(list(...)$is_data.frame), "column ", ""), "'", font_bold(ab), "' (",
           ifelse(ab_coerced != ab, paste0(ab_coerced, ", "), ""),
           ab_name(ab_coerced, tolower = TRUE), ")", mo_var_found, 
           " according to ", ifelse(identical(reference_data, AMR::rsi_translation),
                                    font_bold(guideline_coerced),
                                    "manually defined 'reference_data'"),
           " ... ",
           appendLF = FALSE,
           as_note = FALSE)
  
  result <- exec_as.rsi(method = "mic",
                        x = x,
                        mo = mo_coerced,
                        ab = ab_coerced,
                        guideline = guideline_coerced,
                        uti = uti,
                        conserve_capped_values = conserve_capped_values,
                        add_intrinsic_resistance = add_intrinsic_resistance,
                        reference_data = reference_data) # exec_as.rsi will return message 'OK'
  result
}

#' @rdname as.rsi
#' @export
as.rsi.disk <- function(x,
                        mo = NULL, 
                        ab = deparse(substitute(x)), 
                        guideline = "EUCAST", 
                        uti = FALSE,
                        add_intrinsic_resistance = FALSE,
                        reference_data = AMR::rsi_translation,
                        ...) {
  meet_criteria(x)
  meet_criteria(mo, allow_class = c("mo", "character"), allow_NULL = TRUE)
  meet_criteria(ab, allow_class = c("ab", "character"))
  meet_criteria(guideline, allow_class = "character", has_length = 1)
  meet_criteria(uti, allow_class = "logical", has_length = c(1, length(x)))
  meet_criteria(add_intrinsic_resistance, allow_class = "logical", has_length = 1)
  meet_criteria(reference_data, allow_class = "data.frame")
  check_reference_data(reference_data)
  
  # for dplyr's across()
  cur_column_dplyr <- import_fn("cur_column", "dplyr", error_on_fail = FALSE)
  if (!is.null(cur_column_dplyr)) {
    # try to get current column, which will only be available when in across()
    ab <- tryCatch(cur_column_dplyr(),
                   error = function(e) ab)
  }
  
  # for auto-determining mo
  mo_var_found <- ""
  if (is.null(mo)) {
    tryCatch({
      df <- get_current_data(arg_name = "mo", call = -3) # will return an error if not found
      mo <- NULL
      try({
        mo <- suppressMessages(search_type_in_df(df, "mo"))
      }, silent = TRUE)
      if (!is.null(df) && !is.null(mo) && is.data.frame(df)) {
        mo_var_found <- paste0(" based on column '", font_bold(mo), "'")
        mo <- df[, mo, drop = TRUE]
      }
    }, error = function(e) 
      stop_('No information was supplied about the microorganisms (missing argument `mo`). See ?as.rsi.\n\n',
            "To transform certain columns with e.g. mutate_at(), use `data %>% mutate_at(vars(...), as.rsi, mo = .$x)`, where x is your column with microorganisms.\n",
            "To tranform all disk diffusion zones in a data set, use `data %>% as.rsi()` or data %>% mutate_if(is.disk, as.rsi).", call = FALSE)
    )
  }
  if (length(ab) == 1 && ab %like% "as.disk") {
    stop_('No unambiguous name was supplied about the antibiotic (argument `ab`). See ?as.rsi.', call = FALSE)
  }
  
  ab_coerced <- suppressWarnings(as.ab(ab))
  mo_coerced <- suppressWarnings(as.mo(mo))
  guideline_coerced <- get_guideline(guideline, reference_data)
  if (is.na(ab_coerced)) {
    message_("Returning NAs for unknown drug: '", font_bold(ab),
             "'. Rename this column to a drug name or code, and check the output with `as.ab()`.", 
             add_fn = font_red, 
             as_note = FALSE)
    return(as.rsi(rep(NA, length(x))))
  }
  if (length(mo_coerced) == 1) {
    mo_coerced <- rep(mo_coerced, length(x))
  }
  if (length(uti) == 1) {
    uti <- rep(uti, length(x))
  }
  
  message_("=> Interpreting disk zones of ", ifelse(isTRUE(list(...)$is_data.frame), "column ", ""), "'", font_bold(ab), "' (",
           ifelse(ab_coerced != ab, paste0(ab_coerced, ", "), ""),
           ab_name(ab_coerced, tolower = TRUE), ")", mo_var_found, 
           " according to ", ifelse(identical(reference_data, AMR::rsi_translation),
                                    font_bold(guideline_coerced),
                                    "manually defined 'reference_data'"),
           " ... ",
           appendLF = FALSE,
           as_note = FALSE)
  
  result <- exec_as.rsi(method = "disk",
                        x = x,
                        mo = mo_coerced,
                        ab = ab_coerced,
                        guideline = guideline_coerced,
                        uti = uti,
                        conserve_capped_values = FALSE,
                        add_intrinsic_resistance = add_intrinsic_resistance,
                        reference_data = reference_data) # exec_as.rsi will return message 'OK'
  result
}

#' @rdname as.rsi
#' @export
as.rsi.data.frame <- function(x,
                              ..., 
                              col_mo = NULL, 
                              guideline = "EUCAST",
                              uti = NULL,
                              conserve_capped_values = FALSE,
                              add_intrinsic_resistance = FALSE,
                              reference_data = rsi_translation) {
  meet_criteria(x, allow_class = "data.frame") # will also check for dimensions > 0
  meet_criteria(col_mo, allow_class = "character", is_in = colnames(x), allow_NULL = TRUE)
  meet_criteria(guideline, allow_class = "character", has_length = 1)
  meet_criteria(uti, allow_class = c("logical", "character"), allow_NULL = TRUE)
  meet_criteria(conserve_capped_values, allow_class = "logical", has_length = 1)
  meet_criteria(add_intrinsic_resistance, allow_class = "logical", has_length = 1)
  meet_criteria(reference_data, allow_class = "data.frame")

  x.bak <- x
  for (i in seq_len(ncol(x))) {
    # don't keep factors
    if (is.factor(x[, i, drop = TRUE])) {
      x[, i] <- as.character(x[, i, drop = TRUE])
    }
  }
  
  # -- MO
  col_mo.bak <- col_mo
  if (is.null(col_mo)) {
    col_mo <- search_type_in_df(x = x, type = "mo", info = FALSE)
  }

  # -- UTIs
  col_uti <- uti
  if (is.null(col_uti)) {
    col_uti <- search_type_in_df(x = x, type = "uti")
  }
  if (!is.null(col_uti)) {
    if (is.logical(col_uti)) {
      # already a logical vector as input
      if (length(col_uti) == 1) {
        uti <- rep(col_uti, NROW(x))
      } else {
        uti <- col_uti
      }
    } else {
      # column found, transform to logical
      stop_if(length(col_uti) != 1 | !col_uti %in% colnames(x),
              "argument `uti` must be a logical vector, of must be a single column name of `x`")
      uti <- as.logical(x[, col_uti, drop = TRUE])
    }
  } else {
    # look for specimen column and make logicals of the urines
    col_specimen <- suppressMessages(search_type_in_df(x = x, type = "specimen"))
    if (!is.null(col_specimen)) {
      uti <- x[, col_specimen, drop = TRUE] %like% "urin"
      values <- sort(unique(x[uti, col_specimen, drop = TRUE]))
      if (length(values) > 1) {
        plural <- c("s", "", "")
      } else {
        plural <- c("", "s", "a ")
      }
      message_("Assuming value", plural[1], " ", 
               paste(paste0('"', values, '"'), collapse = ", "),
               " in column '", font_bold(col_specimen),
               "' reflect", plural[2], " ", plural[3], "urinary tract infection", plural[1],
               ".\n  Use `as.rsi(uti = FALSE)` to prevent this.")
    } else {
      # no data about UTI's found
      uti <- FALSE
    }
  }

  i <- 0
  sel <- colnames(pm_select(x, ...))
  if (!is.null(col_mo)) {
    sel <- sel[sel != col_mo]
  }

  ab_cols <- colnames(x)[vapply(FUN.VALUE = logical(1), x, function(y) {
    i <<- i + 1
    check <- is.mic(y) | is.disk(y)
    ab <- colnames(x)[i]
    if (!is.null(col_mo) && ab == col_mo) {
      return(FALSE)
    }
    if (!is.null(col_uti) && ab == col_uti) {
      return(FALSE)
    }
    if (length(sel) == 0 || (length(sel) > 0 && ab %in% sel)) {
      ab_coerced <- suppressWarnings(as.ab(ab))
      if (is.na(ab_coerced) || (length(sel) > 0 & !ab %in% sel)) {
        # not even a valid AB code
        return(FALSE)
      } else {
        return(TRUE)
      }
    } else {
      return(FALSE)
    }
  })]

  stop_if(length(ab_cols) == 0,
          "no columns with MIC values, disk zones or antibiotic column names found in this data set. Use as.mic() or as.disk() to transform antimicrobial columns.")
  # set type per column
  types <- character(length(ab_cols))
  types[vapply(FUN.VALUE = logical(1), x.bak[, ab_cols, drop = FALSE], is.disk)] <- "disk"
  types[vapply(FUN.VALUE = logical(1), x.bak[, ab_cols, drop = FALSE], is.mic)] <- "mic"
  types[types == "" & vapply(FUN.VALUE = logical(1), x[, ab_cols, drop = FALSE], all_valid_disks)] <- "disk"
  types[types == "" & vapply(FUN.VALUE = logical(1), x[, ab_cols, drop = FALSE], all_valid_mics)] <- "mic"
  types[types == "" & !vapply(FUN.VALUE = logical(1), x.bak[, ab_cols, drop = FALSE], is.rsi)] <- "rsi"
  if (any(types %in% c("mic", "disk"), na.rm = TRUE)) {
    # now we need an mo column
    stop_if(is.null(col_mo), "`col_mo` must be set")
    # if not null, we already found it, now find again so a message will show
    if (is.null(col_mo.bak)) {
      col_mo <- search_type_in_df(x = x, type = "mo")
    }
  }

  x_mo <- as.mo(x %pm>% pm_pull(col_mo))

  for (i in seq_len(length(ab_cols))) {
    if (types[i] == "mic") {
      x[, ab_cols[i]] <- as.rsi(x = x %pm>% 
                                  pm_pull(ab_cols[i]) %pm>% 
                                  as.character() %pm>%
                                  as.mic(),
                                mo = x_mo,
                                ab = ab_cols[i],
                                guideline = guideline,
                                uti = uti,
                                conserve_capped_values = conserve_capped_values,
                                add_intrinsic_resistance = add_intrinsic_resistance,
                                reference_data = reference_data,
                                is_data.frame = TRUE)
    } else if (types[i] == "disk") {
      x[, ab_cols[i]] <- as.rsi(x = x %pm>% 
                                  pm_pull(ab_cols[i]) %pm>% 
                                  as.character() %pm>%
                                  as.disk(),
                                mo = x_mo,
                                ab = ab_cols[i],
                                guideline = guideline,
                                uti = uti,
                                add_intrinsic_resistance = add_intrinsic_resistance,
                                reference_data = reference_data,
                                is_data.frame = TRUE)
    } else if (types[i] == "rsi") {
      show_message <- FALSE
      ab <- ab_cols[i]
      ab_coerced <- suppressWarnings(as.ab(ab))
      if (!all(x[, ab_cols[i], drop = TRUE] %in% c("R", "S", "I"), na.rm = TRUE)) {
        show_message <- TRUE
        # only print message if values are not already clean
        message_("=> Cleaning values in column '", font_bold(ab), "' (",
                 ifelse(ab_coerced != ab, paste0(ab_coerced, ", "), ""),
                 ab_name(ab_coerced, tolower = TRUE), ")... ",
                 appendLF = FALSE,
                 as_note = FALSE)
      } else if (!is.rsi(x.bak[, ab_cols[i], drop = TRUE])) {
        show_message <- TRUE
        # only print message if class not already set
        message_("=> Assigning class <rsi> to already clean column '", font_bold(ab), "' (",
                 ifelse(ab_coerced != ab, paste0(ab_coerced, ", "), ""),
                 ab_name(ab_coerced, tolower = TRUE), ")... ",
                 appendLF = FALSE,
                 as_note = FALSE)
      }
      x[, ab_cols[i]] <- as.rsi.default(x = as.character(x[, ab_cols[i], drop = TRUE]))
      if (show_message == TRUE) {
        message_(" OK.", add_fn = list(font_green, font_bold), as_note = FALSE)
      }
    }
  }
  
  x
}

get_guideline <- function(guideline, reference_data) {
  if (!identical(reference_data, AMR::rsi_translation)) {
    return(guideline)
  }
  guideline_param <- toupper(guideline)
  if (guideline_param %in% c("CLSI", "EUCAST")) {
    guideline_param <- rev(sort(subset(reference_data, guideline %like% guideline_param)$guideline))[1L]
  }
  if (!guideline_param %like% " ") {
    # like 'EUCAST2020', should be 'EUCAST 2020'
    guideline_param <- gsub("([a-z]+)([0-9]+)", "\\1 \\2", guideline_param, ignore.case = TRUE)
  }
  
  stop_ifnot(guideline_param %in% reference_data$guideline,
             "invalid guideline: '", guideline,
             "'.\nValid guidelines are: ", paste0("'", unique(reference_data$guideline), "'", collapse = ", "), call = FALSE)
  
  guideline_param
  
}

exec_as.rsi <- function(method,
                        x,
                        mo,
                        ab,
                        guideline,
                        uti,
                        conserve_capped_values, 
                        add_intrinsic_resistance,
                        reference_data) {
  
  metadata_mo <- get_mo_failures_uncertainties_renamed()
  
  x_bak <- data.frame(x_mo = paste0(x, mo), stringsAsFactors = FALSE)
  df <- unique(data.frame(x, mo), stringsAsFactors = FALSE)
  x <- df$x
  mo <- df$mo
  
  if (method == "mic") {
    x <- as.mic(x) # when as.rsi.mic is called directly
  } else if (method == "disk") {
    x <- as.disk(x) # when as.rsi.disk is called directly
  }
  
  warned <- FALSE
  method_param <- toupper(method)
  
  genera <- mo_genus(mo)
  mo_genus <- as.mo(genera)
  mo_family <- as.mo(mo_family(mo))
  mo_order <- as.mo(mo_order(mo))
  if (any(genera == "Staphylococcus", na.rm = TRUE)) {
    mo_becker <- as.mo(mo, Becker = TRUE)
  } else {
    mo_becker <- mo
  }
  if (any(genera == "Streptococcus", na.rm = TRUE)) {
    mo_lancefield <- as.mo(mo, Lancefield = TRUE)
  } else {
    mo_lancefield <- mo
  }
  mo_other <- as.mo(rep("UNKNOWN", length(mo)))
  
  guideline_coerced <- get_guideline(guideline, reference_data)
  if (guideline_coerced != guideline) {
    if (message_not_thrown_before("as.rsi")) {
      message_("Using guideline ", font_bold(guideline_coerced), " as input for `guideline`.")
      remember_thrown_message("as.rsi")
    }
  }
  
  new_rsi <- rep(NA_character_, length(x))
  ab_param <- ab
  if (identical(reference_data, AMR::rsi_translation)) {
    trans <- reference_data %pm>%
      subset(guideline == guideline_coerced & method == method_param & ab == ab_param)
  } else {
    trans <- reference_data %pm>%
      subset(method == method_param & ab == ab_param)
  }
  trans$lookup <- paste(trans$mo, trans$ab)
  
  lookup_mo <- paste(mo, ab)
  lookup_genus <- paste(mo_genus, ab)
  lookup_family <- paste(mo_family, ab)
  lookup_order <- paste(mo_order, ab)
  lookup_becker <- paste(mo_becker, ab)
  lookup_lancefield <- paste(mo_lancefield, ab)
  lookup_other <- paste(mo_other, ab)
  
  if (all(trans$uti == TRUE, na.rm = TRUE) & all(uti == FALSE)) {
    message_("WARNING.", add_fn = list(font_yellow, font_bold), as_note = FALSE)
    warning_("Introducing NA: interpretation of ", font_bold(ab_name(ab, tolower = TRUE)), " for some microorganisms is only available for (uncomplicated) urinary tract infections (UTI). Use argument `uti` to set which isolates are from urine. See ?as.rsi.", call = FALSE)
    warned <- TRUE
  }
  
  any_is_intrinsic_resistant <- FALSE
  
  for (i in seq_len(length(x))) {
    is_intrinsic_r <- paste(mo[i], ab) %in% INTRINSIC_R
    any_is_intrinsic_resistant <- any_is_intrinsic_resistant | is_intrinsic_r
    
    if (isTRUE(add_intrinsic_resistance) & is_intrinsic_r) {
      if (!guideline_coerced %like% "EUCAST") {
        if (message_not_thrown_before("as.rsi2")) {
          warning_("Using 'add_intrinsic_resistance' is only useful when using EUCAST guidelines, since the rules for intrinsic resistance are based on EUCAST.", call = FALSE)
          remember_thrown_message("as.rsi2")
        }
      } else {
        new_rsi[i] <- "R"
        next
      }
    }
    
    get_record <- trans %pm>%
      # no subsetting to UTI for now
      subset(lookup %in% c(lookup_mo[i],
                           lookup_genus[i],
                           lookup_family[i],
                           lookup_order[i],
                           lookup_becker[i],
                           lookup_lancefield[i],
                           lookup_other[i]))
    
    if (isTRUE(uti[i])) {
      get_record <- get_record %pm>% 
        # be as specific as possible (i.e. prefer species over genus):
        # pm_desc(uti) = TRUE on top and FALSE on bottom
        pm_arrange(pm_desc(uti), pm_desc(nchar(mo))) # 'uti' is a column in data set 'rsi_translation'
    } else {
      get_record <- get_record %pm>% 
        pm_filter(uti == FALSE) %pm>% # 'uti' is a column in rsi_translation
        pm_arrange(pm_desc(nchar(mo)))
    }
    get_record <- get_record[1L, , drop = FALSE]
    
    if (NROW(get_record) > 0) {
      if (is.na(x[i])) {
        new_rsi[i] <- NA_character_
      } else if (method == "mic") {
        mic_input <- x[i]
        mic_S <- as.mic(get_record$breakpoint_S)
        mic_R <- as.mic(get_record$breakpoint_R)
        new_rsi[i] <- quick_case_when(isTRUE(conserve_capped_values) & mic_input %like% "^<[0-9]" ~ "S",
                                      isTRUE(conserve_capped_values) & mic_input %like% "^>[0-9]" ~ "R",
                                      # start interpreting: EUCAST uses <= S and > R, CLSI uses <=S and >= R
                                      isTRUE(which(levels(mic_input) == mic_input) <= which(levels(mic_S) == mic_S)) ~ "S",
                                      guideline_coerced %like% "EUCAST" &
                                        isTRUE(which(levels(mic_input) == mic_input) > which(levels(mic_R) == mic_R)) ~ "R",
                                      guideline_coerced %like% "CLSI" &
                                        isTRUE(which(levels(mic_input) == mic_input) >= which(levels(mic_R) == mic_R)) ~ "R",
                                      # return "I" when not match the bottom or top
                                      !is.na(get_record$breakpoint_S) & !is.na(get_record$breakpoint_R) ~ "I",
                                      # and NA otherwise
                                      TRUE ~ NA_character_)
      } else if (method == "disk") {
        new_rsi[i] <- quick_case_when(isTRUE(as.double(x[i]) >= as.double(get_record$breakpoint_S)) ~ "S",
                                      # start interpreting: EUCAST uses >= S and < R, CLSI uses >=S and <= R
                                      guideline_coerced %like% "EUCAST" &
                                        isTRUE(as.double(x[i]) < as.double(get_record$breakpoint_R)) ~ "R",
                                      guideline_coerced %like% "CLSI" &
                                        isTRUE(as.double(x[i]) <= as.double(get_record$breakpoint_R)) ~ "R",
                                      # return "I" when not match the bottom or top
                                      !is.na(get_record$breakpoint_S) & !is.na(get_record$breakpoint_R) ~ "I",
                                      # and NA otherwise
                                      TRUE ~ NA_character_)
      }
    }
  }
  
  if (any_is_intrinsic_resistant & guideline_coerced %like% "EUCAST" & !isTRUE(add_intrinsic_resistance)) {
    # found some intrinsic resistance, but was not applied
    message_("WARNING.", add_fn = list(font_yellow, font_bold), as_note = FALSE)
    if (message_not_thrown_before("as.rsi3")) {
      warning_("Found intrinsic resistance in some bug/drug combinations, although it was not applied.\nUse `as.rsi(..., add_intrinsic_resistance = TRUE)` to apply it.", call = FALSE)
      remember_thrown_message("as.rsi3")
    }
    warned <- TRUE
  }
  
  new_rsi <- x_bak %pm>%
    pm_left_join(data.frame(x_mo = paste0(df$x, df$mo), new_rsi,
                            stringsAsFactors = FALSE),
                 by = "x_mo") %pm>%
    pm_pull(new_rsi)
  
  if (warned == FALSE) {
    message_(" OK.", add_fn = list(font_green, font_bold), as_note = FALSE)
  }
  
  load_mo_failures_uncertainties_renamed(metadata_mo)
  
  set_clean_class(factor(new_rsi, levels = c("S", "I", "R"), ordered = TRUE),
                  new_class =  c("rsi", "ordered", "factor"))
}

# will be exported using s3_register() in R/zzz.R
pillar_shaft.rsi <- function(x, ...) {
  out <- trimws(format(x))
  out[is.na(x)] <- font_grey(" NA")
  out[x == "R"] <- font_rsi_R_bg(font_black("  R  "))
  out[x == "S"] <- font_rsi_S_bg(font_black("  S  "))
  out[x == "I"] <- font_rsi_I_bg(font_black("  I  "))
  create_pillar_column(out, align = "left", width = 5)
}

# will be exported using s3_register() in R/zzz.R
type_sum.rsi <- function(x, ...) {
  "rsi"
}

# will be exported using s3_register() in R/zzz.R
freq.rsi <- function(x, ...) {
  x_name <- deparse(substitute(x))
  x_name <- gsub(".*[$]", "", x_name)
  if (x_name %in% c("x", ".")) {
    # try again going through system calls
    x_name <- stats::na.omit(vapply(FUN.VALUE = character(1),
                                    sys.calls(), 
                                    function(call) {
                                      call_txt <- as.character(call)
                                      ifelse(call_txt[1] %like% "freq$", call_txt[length(call_txt)], character(0))
                                    }))[1L]
  }
  ab <- suppressMessages(suppressWarnings(as.ab(x_name)))
  digits <- list(...)$digits
  if (is.null(digits)) {
    digits <- 2
  }
  if (!is.na(ab)) {
    cleaner::freq.default(x = x, ...,
                          .add_header = list(
                            Drug = paste0(ab_name(ab, language = NULL), " (", ab, ", ", ab_atc(ab), ")"),
                            `Drug group` = ab_group(ab, language = NULL),
                            `%SI` = percentage(susceptibility(x, minimum = 0, as_percent = FALSE),
                                               digits = digits)))
  } else {
    cleaner::freq.default(x = x, ...,
                          .add_header = list(
                            `%SI` = percentage(susceptibility(x, minimum = 0, as_percent = FALSE),
                                               digits = digits)))
  }
}


# will be exported using s3_register() in R/zzz.R
get_skimmers.rsi <- function(column) {
  # get the variable name 'skim_variable'
  name_call <- function(.data) {
    calls <- sys.calls()
    calls_txt <- vapply(calls, function(x) paste(deparse(x), collapse = ""), FUN.VALUE = character(1))
    if (any(calls_txt %like% "skim_variable", na.rm = TRUE)) {
      ind <- which(calls_txt %like% "skim_variable")[1L]
      vars <- tryCatch(eval(parse(text = ".data$skim_variable"), envir = sys.frame(ind)), 
                       error = function(e) NULL)
    } else {
      vars <- NULL
    }
    i <- tryCatch(attributes(calls[[length(calls)]])$position, 
                  error = function(e) NULL)
    if (is.null(vars) | is.null(i)) {
      NA_character_
    } else {
      lengths <- vapply(FUN.VALUE = double(1), vars, length)
      when_starts_rsi <- which(names(vapply(FUN.VALUE = double(1), vars, length)) == "rsi")
      offset <- sum(lengths[c(1:when_starts_rsi - 1)])
      var <- vars$rsi[i - offset]
      if (!isFALSE(var == "data")) {
        NA_character_
      } else{
        ab_name(var)
      }
    }
  }
  
  skimr::sfl(
    skim_type = "rsi",
    ab_name = name_call,
    count_R = count_R,
    count_S = count_susceptible,
    count_I = count_I,
    prop_R = ~proportion_R(., minimum = 0),
    prop_S = ~susceptibility(., minimum = 0),
    prop_I = ~proportion_I(., minimum = 0)
  )
}

#' @method print rsi
#' @export
#' @noRd
print.rsi <- function(x, ...) {
  cat("Class <rsi>\n")
  print(as.character(x), quote = FALSE)
}

#' @method droplevels rsi
#' @export
#' @noRd
droplevels.rsi <- function(x, exclude = if (any(is.na(levels(x)))) NULL else NA, ...) {
  x <- droplevels.factor(x, exclude = exclude, ...)
  class(x) <- c("rsi", "ordered", "factor")
  x
}

#' @method summary rsi
#' @export
#' @noRd
summary.rsi <- function(object, ...) {
  x <- object
  n <- sum(!is.na(x))
  S <- sum(x == "S", na.rm = TRUE)
  I <- sum(x == "I", na.rm = TRUE)
  R <- sum(x == "R", na.rm = TRUE)
  pad <- function(x) {
    if (x == "0%") {
      x <- " 0.0%"
    }
    if (nchar(x) < 5) {
      x <- paste0(rep(" ", 5 - nchar(x)), x)
    }
    x
  }
  value <- c(
    "Class" = "rsi",
    "%R" = paste0(pad(percentage(R / n, digits = 1)), " (n=", R, ")"),
    "%SI" = paste0(pad(percentage((S + I) / n, digits = 1)), " (n=", S + I, ")"),
    "- %S" = paste0(pad(percentage(S / n, digits = 1)), " (n=", S, ")"),
    "- %I" = paste0(pad(percentage(I / n, digits = 1)), " (n=", I, ")")
  )
  class(value) <- c("summaryDefault", "table")
  value
}

#' @method plot rsi
#' @export
#' @importFrom graphics plot text axis
#' @rdname plot
plot.rsi <- function(x,
                     lwd = 2,
                     ylim = NULL,
                     ylab = "Percentage",
                     xlab = "Antimicrobial Interpretation",
                     main = paste("Resistance Overview of", deparse(substitute(x))),
                     axes = FALSE,
                     ...) {
  meet_criteria(lwd, allow_class = c("numeric", "integer"), has_length = 1)
  meet_criteria(ylim, allow_class = c("numeric", "integer"), allow_NULL = TRUE)
  meet_criteria(ylab, allow_class = "character", has_length = 1)
  meet_criteria(xlab, allow_class = "character", has_length = 1)
  meet_criteria(main, allow_class = "character", has_length = 1)
  meet_criteria(axes, allow_class = "logical", has_length = 1)
  
  data <- as.data.frame(table(x), stringsAsFactors = FALSE)
  colnames(data) <- c("x", "n")
  data$s <- round((data$n / sum(data$n)) * 100, 1)
  
  if (!"S" %in% data$x) {
    data <- rbind(data, data.frame(x = "S", n = 0, s = 0, stringsAsFactors = FALSE),
                  stringsAsFactors = FALSE)
  }
  if (!"I" %in% data$x) {
    data <- rbind(data, data.frame(x = "I", n = 0, s = 0, stringsAsFactors = FALSE),
                  stringsAsFactors = FALSE)
  }
  if (!"R" %in% data$x) {
    data <- rbind(data, data.frame(x = "R", n = 0, s = 0, stringsAsFactors = FALSE),
                  stringsAsFactors = FALSE)
  }
  
  # don't use as.rsi() here, it will confuse plot()
  data$x <- factor(data$x, levels = c("S", "I", "R"), ordered = TRUE)
  
  ymax <- pm_if_else(max(data$s) > 95, 105, 100)
  
  plot(x = data$x,
       y = data$s,
       lwd = lwd,
       ylim = c(0, ymax),
       ylab = ylab,
       xlab = xlab,
       main = main,
       axes = axes,
       ...)
  # x axis
  axis(side = 1, at = 1:pm_n_distinct(data$x), labels = levels(data$x), lwd = 0)
  # y axis, 0-100%
  axis(side = 2, at = seq(0, 100, 5))
  
  text(x = data$x,
       y = data$s + 4,
       labels = paste0(data$s, "% (n = ", data$n, ")"))
}


#' @method barplot rsi
#' @export
#' @importFrom graphics barplot axis par
#' @rdname plot
barplot.rsi <- function(height,
                        col = c("chartreuse4", "chartreuse3", "brown3"),
                        xlab = ifelse(beside, "Antimicrobial Interpretation", ""),
                        main = paste("Resistance Overview of", deparse(substitute(height))),
                        ylab = "Frequency",
                        beside = TRUE,
                        axes = beside,
                        ...) {
  meet_criteria(col, allow_class = "character", has_length = 3)
  meet_criteria(xlab, allow_class = "character", has_length = 1)
  meet_criteria(main, allow_class = "character", has_length = 1)
  meet_criteria(ylab, allow_class = "character", has_length = 1)
  meet_criteria(beside, allow_class = "logical", has_length = 1)
  meet_criteria(axes, allow_class = "logical", has_length = 1)
  
  if (axes == TRUE) {
    par(mar =  c(5, 4, 4, 2) + 0.1)
  } else {
    par(mar =  c(2, 4, 4, 2) + 0.1)
  }
  
  barplot(as.matrix(table(height)),
          col = col,
          xlab = xlab,
          main = main,
          ylab = ylab,
          beside = beside,
          axes = FALSE,
          ...)
  # y axis, 0-100%
  axis(side = 2, at = seq(0, max(table(height)) + max(table(height)) * 1.1, by = 25))
  if (axes == TRUE && beside == TRUE) {
    axis(side = 1, labels = levels(height), at = c(1, 2, 3) + 0.5, lwd = 0)
  }
}

#' @method [<- rsi
#' @export
#' @noRd
"[<-.rsi" <- function(i, j, ..., value) {
  value <- as.rsi(value)
  y <- NextMethod()
  attributes(y) <- attributes(i)
  y
}
#' @method [[<- rsi
#' @export
#' @noRd
"[[<-.rsi" <- function(i, j, ..., value) {
  value <- as.rsi(value)
  y <- NextMethod()
  attributes(y) <- attributes(i)
  y
}
#' @method c rsi
#' @export
#' @noRd
c.rsi <- function(x, ...) {
  y <- unlist(lapply(list(...), as.character))
  x <- as.character(x)
  as.rsi(c(x, y))
}

#' @method unique rsi
#' @export
#' @noRd
unique.rsi <- function(x, incomparables = FALSE, ...) {
  y <- NextMethod()
  attributes(y) <- attributes(x)
  y
}

check_reference_data <- function(reference_data) {
  if (!identical(reference_data, AMR::rsi_translation)) {
    class_rsi <- vapply(FUN.VALUE = character(1), rsi_translation, function(x) paste0("<", class(x), ">", collapse = " and "))
    class_ref <- vapply(FUN.VALUE = character(1), reference_data, function(x) paste0("<", class(x), ">", collapse = " and "))
    if (!all(names(class_rsi) == names(class_ref))) {
      stop_("`reference_data` must have the same column names as the 'rsi_translation' data set.", call = -2)
    }
    if (!all(class_rsi == class_ref)) {
      class_rsi[class_rsi != class_ref][1]
      stop_("`reference_data` must be the same structure as the 'rsi_translation' data set. Column '", names(class_ref[class_rsi != class_ref][1]), "' is of class ", class_ref[class_rsi != class_ref][1], ", but should be of class ", class_rsi[class_rsi != class_ref][1], ".", call = -2)
    }
  }
}
````
