## Supplementary material for "AMR - An R Package for Working with Antimicrobial Resistance Data": Source package: rsi_calc.R

```
# ==================================================================== #
# TITLE                                                                #
# Antimicrobial Resistance (AMR) Analysis for R                        #
#                                                                      #
# SOURCE                                                               #
# https://github.com/msberends/AMR                                     #
#                                                                      #
# LICENCE                                                              #
# (c) 2018-2021 Berends MS, Luz CF et al.                              #
# Developed at the University of Groningen, the Netherlands, in        #
# collaboration with non-profit organisations Certe Medical            #
# Diagnostics & Advice, and University Medical Center Groningen.       # 
#                                                                      #
# This R package is free software; you can freely use and distribute   #
# it for both personal and commercial purposes under the terms of the  #
# GNU General Public License version 2.0 (GNU GPL-2), as published by  #
# the Free Software Foundation.                                        #
# We created this package for both routine data analysis and academic  #
# research and it was publicly released in the hope that it will be    #
# useful, but it comes WITHOUT ANY WARRANTY OR LIABILITY.              #
#                                                                      #
# Visit our website for the full manual and a complete tutorial about  #
# how to conduct AMR analysis: https://msberends.github.io/AMR/        #
# ==================================================================== #

dots2vars <- function(...) {
  # this function is to give more informative output about 
  # variable names in count_* and proportion_* functions
  dots <- substitute(list(...))
  paste(as.character(dots)[2:length(dots)], collapse = ", ")
}

rsi_calc <- function(...,
                     ab_result,
                     minimum = 0,
                     as_percent = FALSE,
                     only_all_tested = FALSE,
                     only_count = FALSE) {
  meet_criteria(ab_result, allow_class = c("character", "numeric", "integer"), has_length = c(1, 2, 3), .call_depth = 1)
  meet_criteria(minimum, allow_class = c("numeric", "integer"), has_length = 1, .call_depth = 1)
  meet_criteria(as_percent, allow_class = "logical", has_length = 1, .call_depth = 1)
  meet_criteria(only_all_tested, allow_class = "logical", has_length = 1, .call_depth = 1)
  meet_criteria(only_count, allow_class = "logical", has_length = 1, .call_depth = 1)
  
  data_vars <- dots2vars(...)
  
  dots_df <- switch(1, ...)
  if (is.data.frame(dots_df)) {
    # make sure to remove all other classes like tibbles, data.tables, etc
    dots_df <- as.data.frame(dots_df, stringsAsFactors = FALSE)
  }
  
  dots <- eval(substitute(alist(...)))
  stop_if(length(dots) == 0, "no variables selected", call = -2)
  
  stop_if("also_single_tested" %in% names(dots),
          "`also_single_tested` was replaced by `only_all_tested`.\n",
          "Please read Details in the help page (`?proportion`) as this may have a considerable impact on your analysis.", call = -2)
  ndots <- length(dots)
  
  if (is.data.frame(dots_df)) {
    # data.frame passed with other columns, like: example_isolates %pm>% proportion_S(AMC, GEN)
    
    dots <- as.character(dots)
    # remove first element, it's the data.frame
    if (length(dots) == 1) {
      dots <- character(0)
    } else {
      dots <- dots[2:length(dots)]
    }
    if (length(dots) == 0 | all(dots == "df")) {
     # for complete data.frames, like example_isolates %pm>% select(AMC, GEN) %pm>% proportion_S()
      # and the old rsi function, which has "df" as name of the first argument
      x <- dots_df
    } else {
      # get dots that are in column names already, and the ones that will be once evaluated using dots_df or global env
      # this is to support susceptibility(example_isolates, AMC, any_of(some_vector_with_AB_names))
      dots <- c(dots[dots %in% colnames(dots_df)],
                eval(parse(text = dots[!dots %in% colnames(dots_df)]), envir = dots_df, enclos = globalenv()))
      dots_not_exist <- dots[!dots %in% colnames(dots_df)]
      stop_if(length(dots_not_exist) > 0, "column(s) not found: ", paste0("'", dots_not_exist, "'", collapse = ", "), call = -2)
      x <- dots_df[, dots, drop = FALSE]
    }
  } else if (ndots == 1) {
    # only 1 variable passed (can also be data.frame), like: proportion_S(example_isolates$AMC) and example_isolates$AMC %pm>% proportion_S()
    x <- dots_df
  } else {
    # multiple variables passed without pipe, like: proportion_S(example_isolates$AMC, example_isolates$GEN)
    x <- NULL
    try(x <- as.data.frame(dots, stringsAsFactors = FALSE), silent = TRUE)
    if (is.null(x)) {
      # support for example_isolates %pm>% group_by(hospital_id) %pm>% summarise(amox = susceptibility(GEN, AMX))
      x <- as.data.frame(list(...), stringsAsFactors = FALSE)
    }
  }
  
  if (is.null(x)) {
    warning_("argument is NULL (check if columns exist): returning NA", call = FALSE)
    if (as_percent == TRUE) {
      return(NA_character_)
    } else {
      return(NA_real_)
    }
  }
  
  print_warning <- FALSE
  
  ab_result <- as.rsi(ab_result)
  
  if (is.data.frame(x)) {
    rsi_integrity_check <- character(0)
    for (i in seq_len(ncol(x))) {
      # check integrity of columns: force <rsi> class
      if (!is.rsi(x[, i, drop = TRUE])) {
        rsi_integrity_check <- c(rsi_integrity_check, as.character(x[, i, drop = TRUE]))
        x[, i] <- suppressWarnings(as.rsi(x[, i, drop = TRUE])) # warning will be given later
        print_warning <- TRUE
      }
    }
    if (length(rsi_integrity_check) > 0) {
      # this will give a warning for invalid results, of all input columns (so only 1 warning)
      rsi_integrity_check <- as.rsi(rsi_integrity_check)
    }
    
    x_transposed <- as.list(as.data.frame(t(x), stringsAsFactors = FALSE))
    if (only_all_tested == TRUE) {
      # no NAs in any column
      y <- apply(X = as.data.frame(lapply(x, as.integer), stringsAsFactors = FALSE),
                 MARGIN = 1,
                 FUN = min)
      numerator <- sum(as.integer(y) %in% as.integer(ab_result), na.rm = TRUE)
      denominator <- sum(vapply(FUN.VALUE = logical(1), x_transposed, function(y) !(any(is.na(y)))))
    } else {
      # may contain NAs in any column
      other_values <- setdiff(c(NA, levels(ab_result)), ab_result)
      numerator <- sum(vapply(FUN.VALUE = logical(1), x_transposed, function(y) any(y %in% ab_result, na.rm = TRUE)))
      denominator <- sum(vapply(FUN.VALUE = logical(1), x_transposed, function(y) !(all(y %in% other_values) & any(is.na(y)))))
    }
  } else {
    # x is not a data.frame
    if (!is.rsi(x)) {
      x <- as.rsi(x)
      print_warning <- TRUE
    }
    numerator <- sum(x %in% ab_result, na.rm = TRUE)
    denominator <- sum(x %in% levels(ab_result), na.rm = TRUE)
  }
  
  if (print_warning == TRUE) {
    if (message_not_thrown_before("rsi_calc")) {
      warning_("Increase speed by transforming to class <rsi> on beforehand: your_data %>% mutate_if(is.rsi.eligible, as.rsi)",
               call = FALSE)
      remember_thrown_message("rsi_calc")
    }
  }
  
  if (only_count == TRUE) {
    return(numerator)
  }
  
  if (denominator < minimum) {
    if (data_vars != "") {
      data_vars <- paste(" for", data_vars)
    }
    warning_("Introducing NA: only ", denominator, " results available", data_vars, " (`minimum` = ", minimum, ").", call = FALSE)
    fraction <- NA_real_
  } else {
    fraction <- numerator / denominator
    fraction[is.nan(fraction)] <- NA_real_
  }
  
  if (as_percent == TRUE) {
    percentage(fraction, digits = 1)
  } else {
    fraction
  }
}

rsi_calc_df <- function(type, # "proportion", "count" or "both"
                        data,
                        translate_ab = "name",
                        language = get_locale(),
                        minimum = 30,
                        as_percent = FALSE,
                        combine_SI = TRUE,
                        combine_IR = FALSE,
                        combine_SI_missing = FALSE) {
  meet_criteria(type, is_in = c("proportion", "count", "both"), has_length = 1, .call_depth = 1)
  meet_criteria(data, allow_class = "data.frame", contains_column_class = "rsi", .call_depth = 1)
  meet_criteria(translate_ab, allow_class = c("character", "logical"), has_length = 1, allow_NA = TRUE, .call_depth = 1)
  meet_criteria(language, has_length = 1, is_in = c(LANGUAGES_SUPPORTED, ""), allow_NULL = TRUE, allow_NA = TRUE, .call_depth = 1)
  meet_criteria(minimum, allow_class = c("numeric", "integer"), has_length = 1, .call_depth = 1)
  meet_criteria(as_percent, allow_class = "logical", has_length = 1, .call_depth = 1)
  meet_criteria(combine_SI, allow_class = "logical", has_length = 1, .call_depth = 1)
  meet_criteria(combine_SI_missing, allow_class = "logical", has_length = 1, .call_depth = 1)
  
  check_dataset_integrity()
  
  if (isTRUE(combine_IR) & isTRUE(combine_SI_missing)) {
    combine_SI <- FALSE
  }
  stop_if(isTRUE(combine_SI) & isTRUE(combine_IR), "either `combine_SI` or `combine_IR` can be TRUE, not both", call = -2)
  
  translate_ab <- get_translate_ab(translate_ab)
  
  # select only groups and antibiotics
  if (inherits(data, "grouped_df")) {
    data_has_groups <- TRUE
    groups <- setdiff(names(attributes(data)$groups), ".rows")
    data <- data[, c(groups, colnames(data)[vapply(FUN.VALUE = logical(1), data, is.rsi)]), drop = FALSE]
  } else {
    data_has_groups <- FALSE
    data <- data[, colnames(data)[vapply(FUN.VALUE = logical(1), data, is.rsi)], drop = FALSE]
  }
  
  data <- as.data.frame(data, stringsAsFactors = FALSE)
  if (isTRUE(combine_SI) | isTRUE(combine_IR)) {
    for (i in seq_len(ncol(data))) {
      if (is.rsi(data[, i, drop = TRUE])) {
        data[, i] <- as.character(data[, i, drop = TRUE])
        if (isTRUE(combine_SI)) {
          data[, i] <- gsub("(I|S)", "SI", data[, i, drop = TRUE])
        } else if (isTRUE(combine_IR)) {
          data[, i] <- gsub("(I|R)", "IR", data[, i, drop = TRUE])
        }
      }
    }
  }
  
  sum_it <- function(.data) {
    out <- data.frame(antibiotic = character(0),
                      interpretation = character(0),
                      value = double(0),
                      isolates = integer(0),
                      stringsAsFactors = FALSE)
    if (data_has_groups) {
      group_values <- unique(.data[, which(colnames(.data) %in% groups), drop = FALSE])
      rownames(group_values) <- NULL
      .data <- .data[, which(!colnames(.data) %in% groups), drop = FALSE]
    }
    for (i in seq_len(ncol(.data))) {
      values <- .data[, i, drop = TRUE]
      if (isTRUE(combine_SI)) {
        values <- factor(values, levels = c("SI", "R"), ordered = TRUE)
      } else if (isTRUE(combine_IR)) {
        values <- factor(values, levels = c("S", "IR"), ordered = TRUE)
      } else {
        values <- factor(values, levels = c("S", "I", "R"), ordered = TRUE)
      }
      col_results <- as.data.frame(as.matrix(table(values)), stringsAsFactors = FALSE)
      col_results$interpretation <- rownames(col_results)
      col_results$isolates <- col_results[, 1, drop = TRUE]
      if (NROW(col_results) > 0 && sum(col_results$isolates, na.rm = TRUE) > 0) {
        if (sum(col_results$isolates, na.rm = TRUE) >= minimum) {
          col_results$value <- col_results$isolates / sum(col_results$isolates, na.rm = TRUE)
        } else {
          col_results$value <- rep(NA_real_, NROW(col_results))
        }
        out_new <- data.frame(antibiotic = ifelse(isFALSE(translate_ab),
                                                  colnames(.data)[i],
                                                  ab_property(colnames(.data)[i], property = translate_ab, language = language)),
                              interpretation = col_results$interpretation,
                              value = col_results$value,
                              isolates = col_results$isolates,
                              stringsAsFactors = FALSE)
        if (data_has_groups) {
          if (nrow(group_values) < nrow(out_new)) {
            # repeat group_values for the number of rows in out_new
            repeated <- rep(seq_len(nrow(group_values)),
                            each = nrow(out_new) / nrow(group_values))
            group_values <- group_values[repeated, , drop = FALSE]
          }
          out_new <- cbind(group_values, out_new)
        }
        out <- rbind(out, out_new, stringsAsFactors = FALSE)
      }
    }
    out
  }
  
  # based on pm_apply_grouped_function
  apply_group <- function(.data, fn, groups, drop = FALSE, ...) {
    grouped <- pm_split_into_groups(.data, groups, drop)
    res <- do.call(rbind, unname(lapply(grouped, fn, ...)))
    if (any(groups %in% colnames(res))) {
      class(res) <- c("grouped_data", class(res))
      res <- pm_set_groups(res, groups[groups %in% colnames(res)])
    }
    res
  }
  
  if (data_has_groups) {
    out <- apply_group(data, "sum_it", groups)
  } else {
    out <- sum_it(data)
  }
  
  # apply factors for right sorting in interpretation
  if (isTRUE(combine_SI)) {
    out$interpretation <- factor(out$interpretation, levels = c("SI", "R"), ordered = TRUE)
  } else if (isTRUE(combine_IR)) {
    out$interpretation <- factor(out$interpretation, levels = c("S", "IR"), ordered = TRUE)
  } else {
    # don't use as.rsi() here, as it would add the class <rsi> and we would like
    # the same data structure as output, regardless of input
    out$interpretation <- factor(out$interpretation, levels = c("S", "I", "R"), ordered = TRUE)
  }
  
  if (data_has_groups) {
    # ordering by the groups and two more: "antibiotic" and "interpretation"
   out <-  pm_ungroup(out[do.call("order", out[, seq_len(length(groups) + 2)]), ])
  } else {
    out <- out[order(out$antibiotic, out$interpretation), ]
  }
  
  if (type == "proportion") {
    out <- subset(out, select = -c(isolates))
  } else if (type == "count") {
    out$value <- out$isolates
    out <- subset(out, select = -c(isolates))
  } 
  
  rownames(out) <- NULL
  out
}

get_translate_ab <- function(translate_ab) {
  translate_ab <- as.character(translate_ab)[1L]
  if (translate_ab %in% c("TRUE", "official")) {
    return("name")
  } else if (translate_ab %in% c(NA_character_, "FALSE")) {
    return(FALSE)
  } else {
    translate_ab <- tolower(translate_ab)
    stop_ifnot(translate_ab %in% colnames(AMR::antibiotics),
               "invalid value for 'translate_ab', this must be a column name of the antibiotics data set\n",
               "or TRUE (equals 'name') or FALSE to not translate at all.",
               call = FALSE)
    translate_ab
  }
}
```
