## Supplementary material for "AMR - An R Package for Working with Antimicrobial Resistance Data": Source package: translate.R

```
# ==================================================================== #
# TITLE                                                                #
# Antimicrobial Resistance (AMR) Analysis for R                        #
#                                                                      #
# SOURCE                                                               #
# https://github.com/msberends/AMR                                     #
#                                                                      #
# LICENCE                                                              #
# (c) 2018-2021 Berends MS, Luz CF et al.                              #
# Developed at the University of Groningen, the Netherlands, in        #
# collaboration with non-profit organisations Certe Medical            #
# Diagnostics & Advice, and University Medical Center Groningen.       # 
#                                                                      #
# This R package is free software; you can freely use and distribute   #
# it for both personal and commercial purposes under the terms of the  #
# GNU General Public License version 2.0 (GNU GPL-2), as published by  #
# the Free Software Foundation.                                        #
# We created this package for both routine data analysis and academic  #
# research and it was publicly released in the hope that it will be    #
# useful, but it comes WITHOUT ANY WARRANTY OR LIABILITY.              #
#                                                                      #
# Visit our website for the full manual and a complete tutorial about  #
# how to conduct AMR analysis: https://msberends.github.io/AMR/        #
# ==================================================================== #

#' Translate strings from AMR package
#'
#' For language-dependent output of AMR functions, like [mo_name()], [mo_gramstain()], [mo_type()] and [ab_name()].
#' @inheritSection lifecycle Stable lifecycle
#' @details Strings will be translated to foreign languages if they are defined in a local translation file. Additions to this file can be suggested at our repository. The file can be found here: <https://github.com/msberends/AMR/blob/master/data-raw/translations.tsv>. This file will be read by all functions where a translated output can be desired, like all [`mo_*`][mo_property()] functions (such as [mo_name()], [mo_gramstain()], [mo_type()], etc.) and [`ab_*`][ab_property()] functions (such as [ab_name()], [ab_group()], etc.). 
#'
#' Currently supported languages are: `r paste(sort(gsub(";.*", "", ISOcodes::ISO_639_2[which(ISOcodes::ISO_639_2$Alpha_2 %in% LANGUAGES_SUPPORTED), "Name"])), collapse = ", ")`. Please note that currently not all these languages have translations available for all antimicrobial agents and colloquial microorganism names. 
#'
#' Please suggest your own translations [by creating a new issue on our repository](https://github.com/msberends/AMR/issues/new?title=Translations).
#'
#' ## Changing the default language
#' The system language will be used at default (as returned by `Sys.getenv("LANG")` or, if `LANG` is not set, [Sys.getlocale()]), if that language is supported. But the language to be used can be overwritten in two ways and will be checked in this order:
#' 
#'   1. Setting the R option `AMR_locale`, e.g. by running `options(AMR_locale = "de")`
#'   2. Setting the system variable `LANGUAGE` or `LANG`, e.g. by adding `LANGUAGE="de_DE.utf8"` to your `.Renviron` file in your home directory
#' 
#' So if the R option `AMR_locale` is set, the system variables `LANGUAGE` and `LANG` will be ignored.
#' @inheritSection AMR Read more on our website!
#' @rdname translate
#' @name translate
#' @export
#' @examples
#' # The 'language' argument of below functions
#' # will be set automatically to your system language
#' # with get_locale()
#'
#' # English
#' mo_name("CoNS", language = "en")
#' #> "Coagulase-negative Staphylococcus (CoNS)"
#'
#' # German
#' mo_name("CoNS", language = "de")
#' #> "Koagulase-negative Staphylococcus (KNS)"
#'
#' # Dutch
#' mo_name("CoNS", language = "nl")
#' #> "Coagulase-negatieve Staphylococcus (CNS)"
#'
#' # Spanish
#' mo_name("CoNS", language = "es")
#' #> "Staphylococcus coagulasa negativo (SCN)"
#'
#' # Italian
#' mo_name("CoNS", language = "it")
#' #> "Staphylococcus negativo coagulasi (CoNS)"
#'
#' # Portuguese
#' mo_name("CoNS", language = "pt")
#' #> "Staphylococcus coagulase negativo (CoNS)"
get_locale <- function() {
  # AMR versions 1.3.0 and prior used the environmental variable:
  if (!identical("", Sys.getenv("AMR_locale"))) {
    options(AMR_locale = Sys.getenv("AMR_locale"))
  }
  
  if (!is.null(getOption("AMR_locale", default = NULL))) {
    lang <- getOption("AMR_locale")
    if (lang %in% LANGUAGES_SUPPORTED) {
      return(lang)
    } else {
      stop_("unsupported language set as option 'AMR_locale': '", lang, "' - use one of: ",
            paste0("'", LANGUAGES_SUPPORTED, "'", collapse = ", "))
    }
  } else {
    # we now support the LANGUAGE system variable - return it if set
    if (!identical("", Sys.getenv("LANGUAGE"))) {
      return(coerce_language_setting(Sys.getenv("LANGUAGE")))
    }
    if (!identical("", Sys.getenv("LANG"))) {
      return(coerce_language_setting(Sys.getenv("LANG")))
    }
  }
  
  coerce_language_setting(Sys.getlocale("LC_COLLATE"))
}

coerce_language_setting <- function(lang) {
  # grepl() with ignore.case = FALSE is faster than %like%
  if (grepl("^(English|en_|EN_)", lang, ignore.case = FALSE, perl = TRUE)) {
    # as first option to optimise speed
    "en"
  } else if (grepl("^(German|Deutsch|de_|DE_)", lang, ignore.case = FALSE, perl = TRUE)) {
    "de"
  } else if (grepl("^(Dutch|Nederlands|nl_|NL_)", lang, ignore.case = FALSE, perl = TRUE)) {
    "nl"
  } else if (grepl("^(Spanish|Espa.+ol|es_|ES_)", lang, ignore.case = FALSE, perl = TRUE)) {
    "es"
  } else if (grepl("^(Italian|Italiano|it_|IT_)", lang, ignore.case = FALSE, perl = TRUE)) {
    "it"
  } else if (grepl("^(French|Fran.+ais|fr_|FR_)", lang, ignore.case = FALSE, perl = TRUE)) {
    "fr"
  } else if (grepl("^(Portuguese|Portugu.+s|pt_|PT_)", lang, ignore.case = FALSE, perl = TRUE)) {
    "pt"
  } else {
    # other language -> set to English
    "en"
  }
}

# translate strings based on inst/translations.tsv
translate_AMR <- function(from, language = get_locale(), only_unknown = FALSE) {
  
  if (is.null(language)) {
    return(from)
  }
  if (language %in% c("en", "", NA)) {
    return(from)
  }
  
  df_trans <- translations_file # internal data file
  from.bak <- from
  from_unique <- unique(from)
  from_unique_translated <- from_unique
  
  stop_ifnot(language %in% LANGUAGES_SUPPORTED,
             "unsupported language: '", language, "' - use one of: ",
             paste0("'", LANGUAGES_SUPPORTED, "'", collapse = ", "),
             call = FALSE)
  
  df_trans <- subset(df_trans, lang == language)
  if (only_unknown == TRUE) {
    df_trans <- subset(df_trans, pattern %like% "unknown")
  }
  
  # default case sensitive if value if 'ignore.case' is missing:
  df_trans$ignore.case[is.na(df_trans$ignore.case)] <- FALSE
  # default not using regular expressions (fixed = TRUE) if 'fixed' is missing:
  df_trans$fixed[is.na(df_trans$fixed)] <- TRUE
  
  # check if text to look for is in one of the patterns
  any_form_in_patterns <- tryCatch(any(from_unique %like% paste0("(", paste(df_trans$pattern, collapse = "|"), ")")),
                                   error = function(e) {
                                     warning_("Translation not possible. Please open an issue on GitHub (https://github.com/msberends/AMR/issues).", call = FALSE)
                                     return(FALSE)
                                   })
  if (NROW(df_trans) == 0 | !any_form_in_patterns) {
    return(from)
  }
  
  lapply(seq_len(nrow(df_trans)), 
         function(i) from_unique_translated <<- gsub(pattern = df_trans$pattern[i],
                                                     replacement = df_trans$replacement[i],
                                                     x = from_unique_translated,
                                                     ignore.case = df_trans$ignore.case[i], 
                                                     fixed = df_trans$fixed[i]))
  
  # force UTF-8 for diacritics
  from_unique_translated <- enc2utf8(from_unique_translated)

  # a kind of left join to get all results back
  from_unique_translated[match(from.bak, from_unique)]
}
```
