## Supplementary material for "AMR - An R Package for Working with Antimicrobial Resistance Data": Source package: zzz.R

```
# ==================================================================== #
# TITLE                                                                #
# Antimicrobial Resistance (AMR) Analysis for R                        #
#                                                                      #
# SOURCE                                                               #
# https://github.com/msberends/AMR                                     #
#                                                                      #
# LICENCE                                                              #
# (c) 2018-2021 Berends MS, Luz CF et al.                              #
# Developed at the University of Groningen, the Netherlands, in        #
# collaboration with non-profit organisations Certe Medical            #
# Diagnostics & Advice, and University Medical Center Groningen.       # 
#                                                                      #
# This R package is free software; you can freely use and distribute   #
# it for both personal and commercial purposes under the terms of the  #
# GNU General Public License version 2.0 (GNU GPL-2), as published by  #
# the Free Software Foundation.                                        #
# We created this package for both routine data analysis and academic  #
# research and it was publicly released in the hope that it will be    #
# useful, but it comes WITHOUT ANY WARRANTY OR LIABILITY.              #
#                                                                      #
# Visit our website for the full manual and a complete tutorial about  #
# how to conduct AMR analysis: https://msberends.github.io/AMR/        #
# ==================================================================== #

# set up package environment, used by numerous AMR functions
pkg_env <- new.env(hash = FALSE)

.onLoad <- function(libname, pkgname) {
  
  assign(x = "AB_lookup",
         value = create_AB_lookup(),
         envir = asNamespace("AMR"))
  
  assign(x = "MO_lookup",
         value = create_MO_lookup(),
         envir = asNamespace("AMR"))
  
  assign(x = "MO.old_lookup",
         value = create_MO.old_lookup(),
         envir = asNamespace("AMR"))
  
  assign(x = "INTRINSIC_R",
         value = create_intr_resistance(),
         envir = asNamespace("AMR"))
  
  assign(x = "LANGUAGES_SUPPORTED",
         value = sort(c("en", unique(translations_file$lang))),
         envir = asNamespace("AMR"))
  
  assign(x = "MO_CONS",
         value = create_species_cons_cops("CoNS"),
         envir = asNamespace("AMR"))
  
  assign(x = "MO_COPS",
         value = create_species_cons_cops("CoPS"),
         envir = asNamespace("AMR"))

  # Support for tibble headers (type_sum) and tibble columns content (pillar_shaft)
  # without the need to depend on other packages. This was suggested by the 
  # developers of the vctrs package: 
  # https://github.com/r-lib/vctrs/blob/05968ce8e669f73213e3e894b5f4424af4f46316/R/register-s3.R
  s3_register("pillar::pillar_shaft", "ab")
  s3_register("pillar::pillar_shaft", "mo")
  s3_register("pillar::pillar_shaft", "rsi")
  s3_register("pillar::pillar_shaft", "mic")
  s3_register("pillar::pillar_shaft", "disk")
  s3_register("tibble::type_sum", "ab")
  s3_register("tibble::type_sum", "mo")
  s3_register("tibble::type_sum", "rsi")
  s3_register("tibble::type_sum", "mic")
  s3_register("tibble::type_sum", "disk")
  # Support for frequency tables from the cleaner package
  s3_register("cleaner::freq", "mo")
  s3_register("cleaner::freq", "rsi")
  # Support from skim() from the skimr package
  s3_register("skimr::get_skimmers", "mo")
  s3_register("skimr::get_skimmers", "rsi")
  s3_register("skimr::get_skimmers", "mic")
  s3_register("skimr::get_skimmers", "disk")
  
  # if mo source exists, fire it up (see mo_source())
  try({
    if (file.exists(getOption("AMR_mo_source", "~/mo_source.rds"))) {
      invisible(get_mo_source())
    }
  }, silent = TRUE)
}

.onAttach <- function(...) {
  # show notice in 10% of cases in interactive session
  if (!interactive() || stats::runif(1) > 0.1 || isTRUE(as.logical(getOption("AMR_silentstart", FALSE)))) {
    return()
  }
  packageStartupMessage(word_wrap("Thank you for using the AMR package! ",
                                  "If you have a minute, please anonymously fill in this short questionnaire to improve the package and its functionalities: ",
                                  font_blue("https://msberends.github.io/AMR/survey.html\n"),
                                  "[prevent his notice with ",
                                  font_bold("suppressPackageStartupMessages(library(AMR))"),
                                  " or use ",
                                  font_bold("options(AMR_silentstart = TRUE)"), "]"))
}

create_intr_resistance <- function() {
  # for mo_is_intrinsic_resistant() - saves a lot of time when executed on this vector
  paste(AMR::microorganisms[match(AMR::intrinsic_resistant$microorganism, AMR::microorganisms$fullname), "mo", drop = TRUE],
        AMR::antibiotics[match(AMR::intrinsic_resistant$antibiotic, AMR::antibiotics$name), "ab", drop = TRUE])
}

create_species_cons_cops <- function(type = c("CoNS", "CoPS")) {
  # Determination of which staphylococcal species are CoNS/CoPS according to:
  # - Becker et al. 2014, PMID 25278577
  # - Becker et al. 2019, PMID 30872103
  # - Becker et al. 2020, PMID 32056452
  # this function returns class <mo>
  MO_staph <- AMR::microorganisms
  MO_staph <- MO_staph[which(MO_staph$genus == "Staphylococcus"), , drop = FALSE]
  if (type == "CoNS") {
    MO_staph[which(MO_staph$species %in% c("coagulase-negative", "argensis", "arlettae",
                                           "auricularis", "caeli", "capitis", "caprae", 
                                           "carnosus", "chromogenes", "cohnii", "condimenti",
                                           "debuckii", "devriesei", "edaphicus", "epidermidis",
                                           "equorum", "felis", "fleurettii", "gallinarum",
                                           "haemolyticus", "hominis", "jettensis", "kloosii",
                                           "lentus", "lugdunensis", "massiliensis", "microti",
                                           "muscae", "nepalensis", "pasteuri", "petrasii",
                                           "pettenkoferi", "piscifermentans", "pseudoxylosus",
                                           "rostri", "saccharolyticus", "saprophyticus",
                                           "sciuri", "simulans", "stepanovicii", "succinus",
                                           "vitulinus", "warneri", "xylosus")
                   | (MO_staph$species == "schleiferi" & MO_staph$subspecies %in% c("schleiferi", ""))),
             "mo", drop = TRUE]
  } else if (type == "CoPS") {
    MO_staph[which(MO_staph$species %in% c("coagulase-positive",
                                           "simiae", "agnetis",
                                           "delphini", "lutrae",
                                           "hyicus", "intermedius",
                                           "pseudintermedius", "pseudointermedius",
                                           "schweitzeri", "argenteus")
                   | (MO_staph$species == "schleiferi" & MO_staph$subspecies == "coagulans")),
             "mo", drop = TRUE]
  }
}

create_AB_lookup <- function() {
  AB_lookup <- AMR::antibiotics
  AB_lookup$generalised_name <- generalise_antibiotic_name(AB_lookup$name)
  AB_lookup$generalised_synonyms <- lapply(AB_lookup$synonyms, generalise_antibiotic_name)
  AB_lookup$generalised_abbreviations <- lapply(AB_lookup$abbreviations, generalise_antibiotic_name)
  AB_lookup$generalised_loinc <- lapply(AB_lookup$loinc, generalise_antibiotic_name)
  AB_lookup
}

create_MO_lookup <- function() {
  MO_lookup <- AMR::microorganisms
  
  MO_lookup$kingdom_index <- NA_real_
  MO_lookup[which(MO_lookup$kingdom == "Bacteria" | MO_lookup$mo == "UNKNOWN"), "kingdom_index"] <- 1
  MO_lookup[which(MO_lookup$kingdom == "Fungi"), "kingdom_index"] <- 2
  MO_lookup[which(MO_lookup$kingdom == "Protozoa"), "kingdom_index"] <- 3
  MO_lookup[which(MO_lookup$kingdom == "Archaea"), "kingdom_index"] <- 4
  # all the rest
  MO_lookup[which(is.na(MO_lookup$kingdom_index)), "kingdom_index"] <- 5
  
  # use this paste instead of `fullname` to work with Viridans Group Streptococci, etc.
  MO_lookup$fullname_lower <- tolower(trimws(paste(MO_lookup$genus, 
                                                   MO_lookup$species,
                                                   MO_lookup$subspecies)))
  ind <- MO_lookup$genus == "" | grepl("^[(]unknown ", MO_lookup$fullname)
  MO_lookup[ind, "fullname_lower"] <- tolower(MO_lookup[ind, "fullname"])
  MO_lookup$fullname_lower <- trimws(gsub("[^.a-z0-9/ \\-]+", "", MO_lookup$fullname_lower, perl = TRUE))
  
  # add a column with only "e coli" like combinations
  MO_lookup$g_species <- gsub("^([a-z])[a-z]+ ([a-z]+) ?.*", "\\1 \\2", MO_lookup$fullname_lower, perl = TRUE)
  
  # so arrange data on prevalence first, then kingdom, then full name
  MO_lookup[order(MO_lookup$prevalence, MO_lookup$kingdom_index, MO_lookup$fullname_lower), ]
}

create_MO.old_lookup <- function() {
  MO.old_lookup <- AMR::microorganisms.old
  MO.old_lookup$fullname_lower <- trimws(gsub("[^.a-z0-9/ \\-]+", "", tolower(trimws(MO.old_lookup$fullname))))
  
  # add a column with only "e coli"-like combinations
  MO.old_lookup$g_species <- trimws(gsub("^([a-z])[a-z]+ ([a-z]+) ?.*", "\\1 \\2", MO.old_lookup$fullname_lower))
  
  # so arrange data on prevalence first, then full name
  MO.old_lookup[order(MO.old_lookup$prevalence, MO.old_lookup$fullname_lower), ]
}
```
