## Supplementary material for "AMR - An R Package for Working with Antimicrobial Resistance Data": Source package: README.md

% AMR (for R)

# `AMR` (for R)

[![CRAN](https://www.r-pkg.org/badges/version-ago/AMR)](https://cran.r-project.org/package=AMR)
[![CRANlogs](https://cranlogs.r-pkg.org/badges/grand-total/AMR)](https://cran.r-project.org/package=AMR)
![R-code-check](https://github.com/msberends/AMR/workflows/R-code-check/badge.svg?branch=master)
[![CodeFactor](https://www.codefactor.io/repository/github/msberends/amr/badge)](https://www.codefactor.io/repository/github/msberends/amr)
[![Codecov](https://codecov.io/gh/msberends/AMR/branch/master/graph/badge.svg)](https://codecov.io/gh/msberends/AMR?branch=master)

<img src="https://msberends.github.io/AMR/works_great_on.png" align="center" height="150px" />

`AMR` is a free, open-source and independent R package to simplify the analysis and prediction of Antimicrobial Resistance (AMR) and to work with microbial and antimicrobial data and properties, by using evidence-based methods. Our aim is to provide a standard for clean and reproducible antimicrobial resistance data analysis, that can therefore empower epidemiological analyses to continuously enable surveillance and treatment evaluation in any setting.
 
After installing this package, R knows ~70,000 distinct microbial species and all ~550 antibiotic, antimycotic and antiviral drugs by name and code (including ATC, EARS-NET, LOINC and SNOMED CT), and knows all about valid R/SI and MIC values. It supports any data format, including WHONET/EARS-Net data.

This package is fully independent of any other R package and works on Windows, macOS and Linux with all versions of R since R-3.0.0 (April 2013). It was designed to work in any setting, including those with very limited resources. It was created for both routine data analysis and academic research at the Faculty of Medical Sciences of the University of Groningen, in collaboration with non-profit organisations Certe Medical Diagnostics and Advice and University Medical Center Groningen. This R package is actively maintained and free software; you can freely use and distribute it for both personal and commercial (but not patent) purposes under the terms of the GNU General Public License version 2.0 (GPL-2), as published by the Free Software Foundation.

This is the development source of the `AMR` package for R. Not a developer? Then please visit our website [https://msberends.github.io/AMR/](https://msberends.github.io/AMR/) to read more about this package.

*NOTE: this source code is on GitHub (https://github.com/msberends/AMR), but also automatically mirrored to GitLab (https://gitlab.com/msberends/AMR).*

### How to get this package
Please see [our website](https://msberends.github.io/AMR/#get-this-package).

Bottom line: `install.packages("AMR")`

### Copyright

This R package is licensed under the [GNU General Public License (GPL) v2.0](https://github.com/msberends/AMR/blob/master/LICENSE). In a nutshell, this means that this package:

- May be used for commercial purposes

- May be used for private purposes

- May **not** be used for patent purposes

- May be modified, although:

  - Modifications **must** be released under the same license when distributing the package
  - Changes made to the code **must** be documented

- May be distributed, although:

  - Source code **must** be made available when the package is distributed
  - A copy of the license and copyright notice **must** be included with the package.

- Comes with a LIMITATION of liability

- Comes with NO warranty
