## Supplementary material for "AMR - An R Package for Working with Antimicrobial Resistance Data": Source package: test-_misc.R

```
# ==================================================================== #
# TITLE                                                                #
# Antimicrobial Resistance (AMR) Analysis for R                        #
#                                                                      #
# SOURCE                                                               #
# https://github.com/msberends/AMR                                     #
#                                                                      #
# LICENCE                                                              #
# (c) 2018-2021 Berends MS, Luz CF et al.                              #
# Developed at the University of Groningen, the Netherlands, in        #
# collaboration with non-profit organisations Certe Medical            #
# Diagnostics & Advice, and University Medical Center Groningen.       # 
#                                                                      #
# This R package is free software; you can freely use and distribute   #
# it for both personal and commercial purposes under the terms of the  #
# GNU General Public License version 2.0 (GNU GPL-2), as published by  #
# the Free Software Foundation.                                        #
# We created this package for both routine data analysis and academic  #
# research and it was publicly released in the hope that it will be    #
# useful, but it comes WITHOUT ANY WARRANTY OR LIABILITY.              #
#                                                                      #
# Visit our website for the full manual and a complete tutorial about  #
# how to conduct AMR analysis: https://msberends.github.io/AMR/        #
# ==================================================================== #

context("aa_helper_functions.R")

test_that("percentages works", {
  skip_on_cran()
  expect_equal(percentage(0.25), "25%")
  expect_equal(percentage(0.5), "50%")
  expect_equal(percentage(0.500, digits = 1), "50.0%")
  expect_equal(percentage(0.1234), "12.3%")
  # round up 0.5
  expect_equal(percentage(0.0054), "0.5%")
  expect_equal(percentage(0.0055), "0.6%")
})

test_that("functions missing in older R versions work", {
  skip_on_cran()
  expect_equal(strrep("A", 5), "AAAAA")
  expect_equal(strrep(c("A", "B"), c(5, 2)), c("AAAAA", "BB"))
  expect_equal(trimws(" test "), "test")
  expect_equal(trimws(" test ", "l"), "test ")
  expect_equal(trimws(" test ", "r"), " test")
})

test_that("looking up ab columns works", {
  skip_on_cran()
  expect_warning(generate_warning_abs_missing(c("AMP", "AMX")))
  expect_warning(generate_warning_abs_missing(c("AMP", "AMX"), any = TRUE))
  expect_warning(get_column_abx(example_isolates, hard_dependencies = "FUS"))
  expect_message(get_column_abx(example_isolates, soft_dependencies = "FUS"))
  expect_warning(get_column_abx(dplyr::rename(example_isolates, thisone = AMX), amox = "thisone", tmp = "thisone", verbose = TRUE))
  expect_warning(get_column_abx(dplyr::rename(example_isolates, thisone = AMX), amox = "thisone", tmp = "thisone", verbose = FALSE))
})
```
