## Supplementary material for "AMR - An R Package for Working with Antimicrobial Resistance Data": Source package: test-ab.R

```
# ==================================================================== #
# TITLE                                                                #
# Antimicrobial Resistance (AMR) Analysis for R                        #
#                                                                      #
# SOURCE                                                               #
# https://github.com/msberends/AMR                                     #
#                                                                      #
# LICENCE                                                              #
# (c) 2018-2021 Berends MS, Luz CF et al.                              #
# Developed at the University of Groningen, the Netherlands, in        #
# collaboration with non-profit organisations Certe Medical            #
# Diagnostics & Advice, and University Medical Center Groningen.       # 
#                                                                      #
# This R package is free software; you can freely use and distribute   #
# it for both personal and commercial purposes under the terms of the  #
# GNU General Public License version 2.0 (GNU GPL-2), as published by  #
# the Free Software Foundation.                                        #
# We created this package for both routine data analysis and academic  #
# research and it was publicly released in the hope that it will be    #
# useful, but it comes WITHOUT ANY WARRANTY OR LIABILITY.              #
#                                                                      #
# Visit our website for the full manual and a complete tutorial about  #
# how to conduct AMR analysis: https://msberends.github.io/AMR/        #
# ==================================================================== #

context("ab.R")

test_that("as.ab works", {
  skip_on_cran()
  
  expect_equal(as.character(as.ab(c("J01FA01",
                                    "J 01 FA 01",
                                    "Erythromycin",
                                    "eryt",
                                    "   eryt 123",
                                    "ERYT",
                                    "ERY",
                                    "erytromicine",
                                    "Erythrocin",
                                    "Romycin"))),
               rep("ERY", 10))

  expect_identical(class(as.ab("amox")), c("ab", "character"))
  expect_identical(class(antibiotics$ab), c("ab", "character"))
  expect_true(is.ab(as.ab("amox")))
  expect_output(print(as.ab("amox")))
  expect_output(print(data.frame(a = as.ab("amox"))))

  expect_warning(as.ab("J00AA00")) # ATC not yet available in data set
  expect_warning(as.ab("UNKNOWN"))
  expect_warning(as.ab(""))

  expect_output(print(as.ab("amox")))

  expect_equal(as.character(as.ab("Phloxapen")),
               "FLC")

  expect_equal(suppressWarnings(as.character(as.ab(c("Bacteria", "Bacterial")))),
               c(NA, "TMP"))
  
  expect_equal(as.character(as.ab("Amoxy + clavulaanzuur")),
               "AMC")
  
  expect_equal(as.character(as.ab(c("mreopenem", "co-maoxiclav"))),
               c("MEM", "AMC"))
  
  expect_message(as.ab("cipro mero"))
  
  # assigning and subsetting
  x <- antibiotics$ab
  expect_s3_class(x[1], "ab")
  expect_s3_class(x[[1]], "ab")
  expect_s3_class(c(x[1], x[9]), "ab")
  expect_s3_class(unique(x[1], x[9]), "ab")
  expect_warning(x[1] <- "invalid code")
  expect_warning(x[[1]] <- "invalid code")
  expect_warning(c(x[1], "test"))
})
```
