## Supplementary material for "AMR - An R Package for Working with Antimicrobial Resistance Data": Source package: test-ab_property.R

```
# ==================================================================== #
# TITLE                                                                #
# Antimicrobial Resistance (AMR) Analysis for R                        #
#                                                                      #
# SOURCE                                                               #
# https://github.com/msberends/AMR                                     #
#                                                                      #
# LICENCE                                                              #
# (c) 2018-2021 Berends MS, Luz CF et al.                              #
# Developed at the University of Groningen, the Netherlands, in        #
# collaboration with non-profit organisations Certe Medical            #
# Diagnostics & Advice, and University Medical Center Groningen.       # 
#                                                                      #
# This R package is free software; you can freely use and distribute   #
# it for both personal and commercial purposes under the terms of the  #
# GNU General Public License version 2.0 (GNU GPL-2), as published by  #
# the Free Software Foundation.                                        #
# We created this package for both routine data analysis and academic  #
# research and it was publicly released in the hope that it will be    #
# useful, but it comes WITHOUT ANY WARRANTY OR LIABILITY.              #
#                                                                      #
# Visit our website for the full manual and a complete tutorial about  #
# how to conduct AMR analysis: https://msberends.github.io/AMR/        #
# ==================================================================== #

context("ab_property.R")

test_that("ab_property works", {
  skip_on_cran()
  
  expect_identical(ab_name("AMX", language = NULL), "Amoxicillin")
  expect_identical(as.character(ab_atc("AMX")), "J01CA04")
  expect_identical(ab_cid("AMX"), as.integer(33613))

  expect_equal(class(ab_tradenames("AMX")), "character")
  expect_equal(class(ab_tradenames(c("AMX", "AMX"))), "list")

  expect_identical(ab_group("AMX", language = NULL), "Beta-lactams/penicillins")
  expect_identical(ab_atc_group1("AMX", language = NULL), "Beta-lactam antibacterials, penicillins")
  expect_identical(ab_atc_group2("AMX", language = NULL), "Penicillins with extended spectrum")

  expect_identical(ab_name("Fluclox", language = NULL), "Flucloxacillin")
  expect_identical(ab_name("fluklox", language = NULL), "Flucloxacillin")
  expect_identical(ab_name("floxapen", language = NULL), "Flucloxacillin")
  expect_identical(ab_name(21319, language = NULL), "Flucloxacillin")
  expect_identical(ab_name("J01CF05", language = NULL), "Flucloxacillin")

  expect_identical(ab_ddd("AMX", "oral"), 1.5)
  expect_identical(ab_ddd("AMX", "oral", units = TRUE), "g")
  expect_identical(ab_ddd("AMX", "iv"), 3)
  expect_identical(ab_ddd("AMX", "iv", units = TRUE), "g")

  expect_identical(ab_name(x = c("AMC", "PLB"), language = NULL), c("Amoxicillin/clavulanic acid", "Polymyxin B"))
  expect_identical(ab_name(x = c("AMC", "PLB"), tolower = TRUE, language = NULL),
                   c("amoxicillin/clavulanic acid", "polymyxin B"))

  expect_equal(class(ab_info("AMX")), "list")

  expect_error(ab_property("amox", "invalid property"))
  expect_error(ab_name("amox", language = "INVALID"))
  expect_output(print(ab_name("amox", language = NULL)))
  
  expect_equal(ab_name("21066-6", language = NULL), "Ampicillin")
  expect_equal(ab_loinc("ampicillin"),
               c("21066-6", "3355-5", "33562-0", "33919-2", "43883-8", "43884-6", "87604-5"))
  
  expect_true(ab_url("AMX") %like% "whocc.no")
  expect_warning(ab_url("ASP"))
})
```
