## Supplementary material for "AMR - An R Package for Working with Antimicrobial Resistance Data": Source package: test-age.R

```
# ==================================================================== #
# TITLE                                                                #
# Antimicrobial Resistance (AMR) Analysis for R                        #
#                                                                      #
# SOURCE                                                               #
# https://github.com/msberends/AMR                                     #
#                                                                      #
# LICENCE                                                              #
# (c) 2018-2021 Berends MS, Luz CF et al.                              #
# Developed at the University of Groningen, the Netherlands, in        #
# collaboration with non-profit organisations Certe Medical            #
# Diagnostics & Advice, and University Medical Center Groningen.       # 
#                                                                      #
# This R package is free software; you can freely use and distribute   #
# it for both personal and commercial purposes under the terms of the  #
# GNU General Public License version 2.0 (GNU GPL-2), as published by  #
# the Free Software Foundation.                                        #
# We created this package for both routine data analysis and academic  #
# research and it was publicly released in the hope that it will be    #
# useful, but it comes WITHOUT ANY WARRANTY OR LIABILITY.              #
#                                                                      #
# Visit our website for the full manual and a complete tutorial about  #
# how to conduct AMR analysis: https://msberends.github.io/AMR/        #
# ==================================================================== #

context("age.R")

test_that("age works", {
  skip_on_cran()
  expect_equal(age(x = c("1980-01-01", "1985-01-01", "1990-01-01"),
                   reference = "2019-01-01"),
               c(39, 34, 29))

  expect_equal(age(x = c("2019-01-01", "2019-04-01", "2019-07-01"),
                   reference = "2019-09-01",
                   exact = TRUE),
               c(0.6656393, 0.4191781, 0.1698630),
               tolerance = 0.001)

  expect_error(age(x = c("1980-01-01", "1985-01-01", "1990-01-01"),
                   reference = c("2019-01-01", "2019-01-01")))

  expect_warning(age(x = c("1980-01-01", "1985-01-01", "1990-01-01"),
                   reference = "1975-01-01"))

  expect_warning(age(x = c("1800-01-01", "1805-01-01", "1810-01-01"),
                     reference = "2019-01-01"))
  
  expect_equal(length(age(x = c("2019-01-01", NA), na.rm = TRUE)),
               1)
  
})

test_that("age_groups works", {
  skip_on_cran()
  ages <- c(3, 8, 16, 54, 31, 76, 101, 43, 21)

  expect_equal(length(unique(age_groups(ages, 50))),
               2)
  expect_equal(length(unique(age_groups(ages, c(50, 60)))),
               3)
  expect_identical(class(age_groups(ages, "child")),
                   c("ordered", "factor"))

  expect_identical(class(age_groups(ages, "elderly")),
                   c("ordered", "factor"))

  expect_identical(class(age_groups(ages, "tens")),
                   c("ordered", "factor"))

  expect_identical(class(age_groups(ages, "fives")),
                   c("ordered", "factor"))
  
  expect_equal(length(age_groups(c(10, 20, 30, NA), na.rm = TRUE)),
               3)

})
```
