## Supplementary material for "AMR - An R Package for Working with Antimicrobial Resistance Data": Source package: test-count.R

```
# ==================================================================== #
# TITLE                                                                #
# Antimicrobial Resistance (AMR) Analysis for R                        #
#                                                                      #
# SOURCE                                                               #
# https://github.com/msberends/AMR                                     #
#                                                                      #
# LICENCE                                                              #
# (c) 2018-2021 Berends MS, Luz CF et al.                              #
# Developed at the University of Groningen, the Netherlands, in        #
# collaboration with non-profit organisations Certe Medical            #
# Diagnostics & Advice, and University Medical Center Groningen.       # 
#                                                                      #
# This R package is free software; you can freely use and distribute   #
# it for both personal and commercial purposes under the terms of the  #
# GNU General Public License version 2.0 (GNU GPL-2), as published by  #
# the Free Software Foundation.                                        #
# We created this package for both routine data analysis and academic  #
# research and it was publicly released in the hope that it will be    #
# useful, but it comes WITHOUT ANY WARRANTY OR LIABILITY.              #
#                                                                      #
# Visit our website for the full manual and a complete tutorial about  #
# how to conduct AMR analysis: https://msberends.github.io/AMR/        #
# ==================================================================== #

context("count.R")

test_that("counts work", {
  skip_on_cran()
  
  expect_equal(count_resistant(example_isolates$AMX), count_R(example_isolates$AMX))
  expect_equal(count_susceptible(example_isolates$AMX), count_SI(example_isolates$AMX))
  expect_equal(count_all(example_isolates$AMX), n_rsi(example_isolates$AMX))
  
  # AMX resistance in `example_isolates`
  expect_equal(count_R(example_isolates$AMX), 804)
  expect_equal(count_I(example_isolates$AMX), 3)
  expect_equal(suppressWarnings(count_S(example_isolates$AMX)), 543)
  expect_equal(count_R(example_isolates$AMX) + count_I(example_isolates$AMX),
               suppressWarnings(count_IR(example_isolates$AMX)))
  expect_equal(suppressWarnings(count_S(example_isolates$AMX)) + count_I(example_isolates$AMX),
               count_SI(example_isolates$AMX))
  
  library(dplyr, warn.conflicts = FALSE)
  expect_equal(example_isolates %>% count_susceptible(AMC), 1433)
  expect_equal(example_isolates %>% count_susceptible(AMC, GEN, only_all_tested = TRUE), 1687)
  expect_equal(example_isolates %>% count_susceptible(AMC, GEN, only_all_tested = FALSE), 1764)
  expect_equal(example_isolates %>% count_all(AMC, GEN, only_all_tested = TRUE), 1798)
  expect_equal(example_isolates %>% count_all(AMC, GEN, only_all_tested = FALSE), 1936)
  expect_identical(example_isolates %>% count_all(AMC, GEN, only_all_tested = TRUE),
                   example_isolates %>% count_susceptible(AMC, GEN, only_all_tested = TRUE) +
                     example_isolates %>% count_resistant(AMC, GEN, only_all_tested = TRUE))
  
  # count of cases
  expect_equal(example_isolates %>%
                 group_by(hospital_id) %>%
                 summarise(cipro = count_susceptible(CIP),
                           genta = count_susceptible(GEN),
                           combination = count_susceptible(CIP, GEN)) %>%
                 pull(combination),
               c(253, 465, 192, 558))
  
  # count_df
  expect_equal(
    example_isolates %>% select(AMX) %>% count_df() %>% pull(value),
    c(example_isolates$AMX %>% count_susceptible(),
      example_isolates$AMX %>% count_resistant())
  )
  expect_equal(
    example_isolates %>% select(AMX) %>% count_df(combine_IR = TRUE) %>% pull(value),
    c(suppressWarnings(example_isolates$AMX %>% count_S()),
      suppressWarnings(example_isolates$AMX %>% count_IR()))
  )
  expect_equal(
    example_isolates %>% select(AMX) %>% count_df(combine_SI = FALSE) %>% pull(value),
    c(suppressWarnings(example_isolates$AMX %>% count_S()),
      example_isolates$AMX %>% count_I(),
      example_isolates$AMX %>% count_R())
  )
  
  # warning for speed loss
  reset_all_thrown_messages()
  expect_warning(count_resistant(as.character(example_isolates$AMC)))
  reset_all_thrown_messages()
  expect_warning(count_resistant(example_isolates$AMC,
                                 as.character(example_isolates$GEN)))
  
  # check for errors
  expect_error(count_resistant("test", minimum = "test"))
  expect_error(count_resistant("test", as_percent = "test"))
  expect_error(count_susceptible("test", minimum = "test"))
  expect_error(count_susceptible("test", as_percent = "test"))
  
  expect_error(count_df(c("A", "B", "C")))
  expect_error(count_df(example_isolates[, "date"]))
  
  # grouping in rsi_calc_df() (= backbone of rsi_df())
  expect_true("hospital_id" %in% (example_isolates %>% 
                group_by(hospital_id) %>% 
                select(hospital_id, AMX, CIP, gender) %>%
                rsi_df() %>% 
                colnames()))
  
})
```
