## Supplementary material for "AMR - An R Package for Working with Antimicrobial Resistance Data": Source package: test-data.R

```
# ==================================================================== #
# TITLE                                                                #
# Antimicrobial Resistance (AMR) Analysis for R                        #
#                                                                      #
# SOURCE                                                               #
# https://github.com/msberends/AMR                                     #
#                                                                      #
# LICENCE                                                              #
# (c) 2018-2021 Berends MS, Luz CF et al.                              #
# Developed at the University of Groningen, the Netherlands, in        #
# collaboration with non-profit organisations Certe Medical            #
# Diagnostics & Advice, and University Medical Center Groningen.       # 
#                                                                      #
# This R package is free software; you can freely use and distribute   #
# it for both personal and commercial purposes under the terms of the  #
# GNU General Public License version 2.0 (GNU GPL-2), as published by  #
# the Free Software Foundation.                                        #
# We created this package for both routine data analysis and academic  #
# research and it was publicly released in the hope that it will be    #
# useful, but it comes WITHOUT ANY WARRANTY OR LIABILITY.              #
#                                                                      #
# Visit our website for the full manual and a complete tutorial about  #
# how to conduct AMR analysis: https://msberends.github.io/AMR/        #
# ==================================================================== #

context("data.R")

test_that("data sets are valid", {
  skip_on_cran()
  expect_true(check_dataset_integrity()) # in misc.R
  
  # IDs should always be unique
  expect_identical(nrow(microorganisms), length(unique(microorganisms$mo)))
  expect_identical(class(microorganisms$mo), c("mo", "character"))
  expect_identical(nrow(antibiotics), length(unique(antibiotics$ab)))
  expect_identical(class(antibiotics$ab), c("ab", "character"))
  
  # check cross table reference
  expect_true(all(microorganisms.codes$mo %in% microorganisms$mo))
  expect_true(all(example_isolates$mo %in% microorganisms$mo))
  expect_true(all(microorganisms.translation$mo_new %in% microorganisms$mo))
  expect_true(all(rsi_translation$mo %in% microorganisms$mo))
  expect_true(all(intrinsic_resistant$microorganism %in% microorganisms$fullname)) # also important for mo_is_intrinsic_resistant()
  expect_true(all(intrinsic_resistant$antibiotic %in% antibiotics$name))
  expect_false(any(is.na(microorganisms.codes$code)))
  expect_false(any(is.na(microorganisms.codes$mo)))
  expect_false(any(microorganisms.translation$mo_old %in% microorganisms$mo))
  
  # antibiotic names must always be coercible to their original AB code
  expect_identical(antibiotics$ab, as.ab(antibiotics$name))
  
  # there should be no diacritics (i.e. non ASCII) characters in the datasets (CRAN policy)
  datasets <- data(package = "AMR", envir = asNamespace("AMR"))$results[, "Item"]
  for (i in seq_len(length(datasets))) {
    dataset <- get(datasets[i], envir = asNamespace("AMR"))
    expect_identical(dataset_UTF8_to_ASCII(dataset), dataset, label = datasets[i])
  }
})

test_that("creation of data sets is valid", {
  skip_on_cran()
  
  df <- create_MO_lookup()
  expect_lt(nrow(df[which(df$prevalence == 1), ]), nrow(df[which(df$prevalence == 2), ]))
  expect_lt(nrow(df[which(df$prevalence == 2), ]), nrow(df[which(df$prevalence == 3), ]))
  expect_true(all(c("mo", "fullname",
                    "kingdom", "phylum", "class", "order", "family", "genus", "species", "subspecies",
                    "rank", "ref", "species_id", "source", "prevalence", "snomed",
                    "kingdom_index", "fullname_lower", "g_species") %in% colnames(df)))
  
  olddf <- create_MO.old_lookup()
  expect_true(all(c("fullname", "fullname_new", "ref", "prevalence",
                    "fullname_lower", "g_species") %in% colnames(olddf)))
  
  expect_s3_class(create_species_cons_cops("CoNS"), "mo")
  
})

test_that("CoL version info works", {
  skip_on_cran()
  
  expect_identical(class(catalogue_of_life_version()),
                   c("catalogue_of_life_version", "list"))
  
  expect_output(print(catalogue_of_life_version()))
})
```
