## Supplementary material for "AMR - An R Package for Working with Antimicrobial Resistance Data": Source package: test-episode.R

```
# ==================================================================== #
# TITLE                                                                #
# Antimicrobial Resistance (AMR) Analysis for R                        #
#                                                                      #
# SOURCE                                                               #
# https://github.com/msberends/AMR                                     #
#                                                                      #
# LICENCE                                                              #
# (c) 2018-2021 Berends MS, Luz CF et al.                              #
# Developed at the University of Groningen, the Netherlands, in        #
# collaboration with non-profit organisations Certe Medical            #
# Diagnostics & Advice, and University Medical Center Groningen.       # 
#                                                                      #
# This R package is free software; you can freely use and distribute   #
# it for both personal and commercial purposes under the terms of the  #
# GNU General Public License version 2.0 (GNU GPL-2), as published by  #
# the Free Software Foundation.                                        #
# We created this package for both routine data analysis and academic  #
# research and it was publicly released in the hope that it will be    #
# useful, but it comes WITHOUT ANY WARRANTY OR LIABILITY.              #
#                                                                      #
# Visit our website for the full manual and a complete tutorial about  #
# how to conduct AMR analysis: https://msberends.github.io/AMR/        #
# ==================================================================== #

context("episode.R")

test_that("episodes work", {
  skip_on_cran()
  
  test_df <- rbind(
    data.frame(
      date = as.Date(c("2015-01-01", "2015-10-01", "2016-02-04", "2016-12-31", "2017-01-01", "2017-02-01", "2017-02-05", "2020-01-01")),
      patient_id = "A"
    ),
    data.frame(
      date = as.Date(c("2015-01-01", "2016-02-01", "2016-12-31", "2017-01-01", "2017-02-03")),
      patient_id = "B"
    ))
  
  expect_equal(get_episode(test_df$date, 365),
               c(1, 1, 2, 2, 2, 3, 3, 4, 1, 2, 2, 2, 3))
  
  library(dplyr)
  expect_identical(test_df %>% group_by(patient_id) %>% mutate(f = is_new_episode(date, 365)) %>% pull(f),
                   c(TRUE, FALSE, TRUE, FALSE, FALSE, FALSE, TRUE, TRUE, TRUE, TRUE, FALSE, FALSE, TRUE))
  
  suppressMessages(
    x <- example_isolates %>%
      mutate(out = first_isolate(., include_unknown = TRUE, info = FALSE))
  )
  
  y <- example_isolates %>%
    group_by(patient_id, mo) %>%
    mutate(out = is_new_episode(date, 365))
  
  expect_identical(which(x$out), which(y$out))
})
```
