## Supplementary material for "AMR - An R Package for Working with Antimicrobial Resistance Data": Source package: test-eucast_rules.R

```
# ==================================================================== #
# TITLE                                                                #
# Antimicrobial Resistance (AMR) Analysis for R                        #
#                                                                      #
# SOURCE                                                               #
# https://github.com/msberends/AMR                                     #
#                                                                      #
# LICENCE                                                              #
# (c) 2018-2021 Berends MS, Luz CF et al.                              #
# Developed at the University of Groningen, the Netherlands, in        #
# collaboration with non-profit organisations Certe Medical            #
# Diagnostics & Advice, and University Medical Center Groningen.       # 
#                                                                      #
# This R package is free software; you can freely use and distribute   #
# it for both personal and commercial purposes under the terms of the  #
# GNU General Public License version 2.0 (GNU GPL-2), as published by  #
# the Free Software Foundation.                                        #
# We created this package for both routine data analysis and academic  #
# research and it was publicly released in the hope that it will be    #
# useful, but it comes WITHOUT ANY WARRANTY OR LIABILITY.              #
#                                                                      #
# Visit our website for the full manual and a complete tutorial about  #
# how to conduct AMR analysis: https://msberends.github.io/AMR/        #
# ==================================================================== #

context("eucast_rules.R")

test_that("EUCAST rules work", {
  
  skip_on_cran()
  
  # thoroughly check input table
  expect_equal(colnames(eucast_rules_file),
               c("if_mo_property", "like.is.one_of", "this_value",
                 "and_these_antibiotics", "have_these_values",
                 "then_change_these_antibiotics", "to_value",
                 "reference.rule", "reference.rule_group",
                 "reference.version",
                 "note"))
  MOs_mentioned <- unique(eucast_rules_file$this_value)
  MOs_mentioned <- sort(trimws(unlist(strsplit(MOs_mentioned[!is_possibly_regex(MOs_mentioned)], ",", fixed = TRUE))))
  MOs_test <- suppressWarnings(suppressMessages(mo_name(MOs_mentioned)))
  expect_length(MOs_mentioned[MOs_test != MOs_mentioned], 0)
  
  expect_error(suppressWarnings(eucast_rules(example_isolates, col_mo = "Non-existing")))
  expect_error(eucast_rules(x = "text"))
  expect_error(eucast_rules(data.frame(a = "test")))
  expect_error(eucast_rules(data.frame(mo = "test"), rules = "invalid rules set"))
  
  expect_warning(eucast_rules(data.frame(mo = "Escherichia coli", vancomycin = "S", stringsAsFactors = TRUE)))
  
  expect_identical(colnames(example_isolates),
                   colnames(suppressWarnings(eucast_rules(example_isolates))))
  
  a <- data.frame(mo = c("Klebsiella pneumoniae",
                         "Pseudomonas aeruginosa",
                         "Enterobacter aerogenes"),
                  amox = "-",           # Amoxicillin
                  stringsAsFactors = FALSE)
  b <- data.frame(mo = c("Klebsiella pneumoniae",
                         "Pseudomonas aeruginosa",
                         "Enterobacter aerogenes"),
                  amox = "R",       # Amoxicillin
                  stringsAsFactors = FALSE)
  expect_identical(suppressWarnings(eucast_rules(a, "mo", info = FALSE)), b)
  expect_identical(suppressWarnings(eucast_rules(a, "mo", info = TRUE)), b)
  
  a <- data.frame(mo = c("Staphylococcus aureus",
                         "Streptococcus group A"),
                  COL = "-",       # Colistin
                  stringsAsFactors = FALSE)
  b <- data.frame(mo = c("Staphylococcus aureus",
                         "Streptococcus group A"),
                  COL = "R",       # Colistin
                  stringsAsFactors = FALSE)
  expect_equal(suppressWarnings(eucast_rules(a, "mo", info = FALSE)), b)
  
  # piperacillin must be R in Enterobacteriaceae when tica is R
  library(dplyr, warn.conflicts = FALSE)
  expect_equal(suppressWarnings(
    example_isolates %>%
      mutate(TIC = as.rsi("R"),
             PIP = as.rsi("S")) %>%
      eucast_rules(col_mo = "mo", version_expertrules = 3.1) %>%
      left_join_microorganisms() %>%
      filter(family == "Enterobacteriaceae") %>%
      pull(PIP) %>%
      unique() %>%
      as.character()),
    "R")
  
  # Azithromycin and Clarythromycin must be equal to Erythromycin
  a <- suppressWarnings(as.rsi(eucast_rules(data.frame(mo = example_isolates$mo,
                                                       ERY = example_isolates$ERY,
                                                       AZM = as.rsi("R"),
                                                       CLR = factor("R"),
                                                       stringsAsFactors = FALSE),
                                            version_expertrules = 3.1)$CLR))
  b <- example_isolates$ERY
  expect_identical(a[!is.na(b)],
                   b[!is.na(b)])
  
  # amox is inferred by benzylpenicillin in Kingella kingae
  expect_equal(
    suppressWarnings(
      as.list(eucast_rules(
        data.frame(mo = as.mo("Kingella kingae"),
                   PEN = "S",
                   AMX = "-",
                   stringsAsFactors = FALSE)
        , info = FALSE))$AMX
    ),
    "S")
  
  # also test norf
  expect_output(suppressWarnings(eucast_rules(example_isolates %>% mutate(NOR = "S", NAL = "S"), info = TRUE)))
  
  # check verbose output
  expect_output(suppressWarnings(eucast_rules(example_isolates, verbose = TRUE, rules = "all", info = TRUE)))
  
  # AmpC de-repressed cephalo mutants
  expect_identical(
    eucast_rules(data.frame(mo = c("Escherichia coli", "Enterobacter cloacae"),
                            cefotax = as.rsi(c("S", "S"))),
                 ampc_cephalosporin_resistance = "R",
                 info = FALSE)$cefotax,
    as.rsi(c("S", "R")))
  expect_identical(
    eucast_rules(data.frame(mo = c("Escherichia coli", "Enterobacter cloacae"),
                            cefotax = as.rsi(c("S", "S"))),
                 ampc_cephalosporin_resistance = NA,
                 info = FALSE)$cefotax,
    as.rsi(c("S", NA)))
  expect_identical(
    eucast_rules(data.frame(mo = c("Escherichia coli", "Enterobacter cloacae"),
                            cefotax = as.rsi(c("S", "S"))),
                 ampc_cephalosporin_resistance = NULL,
                 info = FALSE)$cefotax,
    as.rsi(c("S", "S")))
  
})
```
