## Supplementary material for "AMR - An R Package for Working with Antimicrobial Resistance Data": Source package: test-filter_ab_class.R

```
# ==================================================================== #
# TITLE                                                                #
# Antimicrobial Resistance (AMR) Analysis for R                        #
#                                                                      #
# SOURCE                                                               #
# https://github.com/msberends/AMR                                     #
#                                                                      #
# LICENCE                                                              #
# (c) 2018-2021 Berends MS, Luz CF et al.                              #
# Developed at the University of Groningen, the Netherlands, in        #
# collaboration with non-profit organisations Certe Medical            #
# Diagnostics & Advice, and University Medical Center Groningen.       # 
#                                                                      #
# This R package is free software; you can freely use and distribute   #
# it for both personal and commercial purposes under the terms of the  #
# GNU General Public License version 2.0 (GNU GPL-2), as published by  #
# the Free Software Foundation.                                        #
# We created this package for both routine data analysis and academic  #
# research and it was publicly released in the hope that it will be    #
# useful, but it comes WITHOUT ANY WARRANTY OR LIABILITY.              #
#                                                                      #
# Visit our website for the full manual and a complete tutorial about  #
# how to conduct AMR analysis: https://msberends.github.io/AMR/        #
# ==================================================================== #

context("filter_ab_class.R")

test_that("ATC-group filtering works", {
  skip_on_cran()
  
  library(dplyr)
  expect_gt(example_isolates %>% filter_ab_class("carbapenem") %>% nrow(), 0)
  expect_gt(example_isolates %>% filter_aminoglycosides() %>% nrow(), 0)
  expect_gt(example_isolates %>% filter_carbapenems() %>% nrow(), 0)
  expect_gt(example_isolates %>% filter_cephalosporins() %>% nrow(), 0)
  expect_gt(example_isolates %>% filter_1st_cephalosporins() %>% nrow(), 0)
  expect_gt(example_isolates %>% filter_2nd_cephalosporins() %>% nrow(), 0)
  expect_gt(example_isolates %>% filter_3rd_cephalosporins() %>% nrow(), 0)
  expect_gt(example_isolates %>% filter_4th_cephalosporins() %>% nrow(), 0)
  expect_gt(example_isolates %>% filter_fluoroquinolones() %>% nrow(), 0)
  expect_gt(example_isolates %>% filter_glycopeptides() %>% nrow(), 0)
  expect_gt(example_isolates %>% filter_macrolides() %>% nrow(), 0)
  expect_gt(example_isolates %>% filter_tetracyclines() %>% nrow(), 0)

  expect_gt(example_isolates %>% filter_carbapenems("R", "all") %>% nrow(), 0)

  expect_error(example_isolates %>% filter_carbapenems(result = "test"))
  expect_error(example_isolates %>% filter_carbapenems(scope = "test"))
  expect_message(example_isolates %>% select(1:3) %>% filter_carbapenems())
})
```
