## Supplementary material for "AMR - An R Package for Working with Antimicrobial Resistance Data": Source package: test-first_isolate.R

```
# ==================================================================== #
# TITLE                                                                #
# Antimicrobial Resistance (AMR) Analysis for R                        #
#                                                                      #
# SOURCE                                                               #
# https://github.com/msberends/AMR                                     #
#                                                                      #
# LICENCE                                                              #
# (c) 2018-2021 Berends MS, Luz CF et al.                              #
# Developed at the University of Groningen, the Netherlands, in        #
# collaboration with non-profit organisations Certe Medical            #
# Diagnostics & Advice, and University Medical Center Groningen.       # 
#                                                                      #
# This R package is free software; you can freely use and distribute   #
# it for both personal and commercial purposes under the terms of the  #
# GNU General Public License version 2.0 (GNU GPL-2), as published by  #
# the Free Software Foundation.                                        #
# We created this package for both routine data analysis and academic  #
# research and it was publicly released in the hope that it will be    #
# useful, but it comes WITHOUT ANY WARRANTY OR LIABILITY.              #
#                                                                      #
# Visit our website for the full manual and a complete tutorial about  #
# how to conduct AMR analysis: https://msberends.github.io/AMR/        #
# ==================================================================== #

context("first_isolate.R")

test_that("first isolates work", {
  skip_on_cran()
  
  # first isolates
  expect_equal(
    sum(
      first_isolate(x = example_isolates,
                    col_date = "date",
                    col_patient_id = "patient_id",
                    col_mo = "mo",
                    info = TRUE),
      na.rm = TRUE),
    1300)

  # first weighted isolates
  ex_iso_with_keyab <- example_isolates
  ex_iso_with_keyab$keyab <- key_antibiotics(example_isolates, warnings = FALSE)
  expect_equal(
    suppressWarnings(
      sum(
        first_isolate(x = ex_iso_with_keyab,
                      # let syntax determine arguments automatically
                      type = "keyantibiotics",
                      info = TRUE),
        na.rm = TRUE)),
    1396)

  # when not ignoring I
  expect_equal(
    suppressWarnings(
      sum(
        first_isolate(x = ex_iso_with_keyab,
                      col_date = "date",
                      col_patient_id = "patient_id",
                      col_mo = "mo",
                      col_keyantibiotics = "keyab",
                      ignore_I = FALSE,
                      type = "keyantibiotics",
                      info = TRUE),
        na.rm = TRUE)),
    1419)
  # when using points
  expect_equal(
    suppressWarnings(
      sum(
        first_isolate(x = ex_iso_with_keyab,
                      col_date = "date",
                      col_patient_id = "patient_id",
                      col_mo = "mo",
                      col_keyantibiotics = "keyab",
                      type = "points",
                      info = TRUE),
        na.rm = TRUE)),
    1399)

  # first non-ICU isolates
  expect_equal(
    sum(
      first_isolate(example_isolates,
                    col_mo = "mo",
                    col_date = "date",
                    col_patient_id = "patient_id",
                    col_icu = "ward_icu",
                    info = TRUE,
                    icu_exclude = TRUE),
      na.rm = TRUE),
    881)

  # set 1500 random observations to be of specimen type 'Urine'
  random_rows <- sample(x = 1:2000, size = 1500, replace = FALSE)
  x <- example_isolates
  x$specimen <- "Other"
  x[random_rows, "specimen"] <- "Urine"
  expect_lt(
    sum(
      first_isolate(x = x,
                    col_date = "date",
                    col_patient_id = "patient_id",
                    col_mo = "mo",
                    col_specimen = "specimen",
                    filter_specimen = "Urine",
                    info = TRUE),
      na.rm = TRUE),
    1501)
  # same, but now exclude ICU
  expect_lt(
    sum(
      first_isolate(x = x,
                    col_date = "date",
                    col_patient_id = "patient_id",
                    col_mo = "mo",
                    col_specimen = "specimen",
                    filter_specimen = "Urine",
                    col_icu = "ward_icu",
                    icu_exclude = TRUE,
                    info = TRUE),
      na.rm = TRUE),
    1501)

  # "No isolates found"
  test_iso <- example_isolates
  test_iso$specimen <- "test"
  expect_message(first_isolate(test_iso, 
                               "date", 
                               "patient_id",
                               col_mo = "mo",
                               col_specimen = "specimen",
                               filter_specimen = "something_unexisting",
                               info = TRUE))

  # printing of exclusion message
  expect_message(first_isolate(example_isolates,
                                col_date = "date",
                                col_mo = "mo",
                                col_patient_id = "patient_id",
                                col_testcode = "gender",
                                testcodes_exclude = "M",
                                info = TRUE))

  # errors
  expect_error(first_isolate("date", "patient_id", col_mo = "mo"))
  expect_error(first_isolate(example_isolates,
                             col_date = "non-existing col",
                             col_mo = "mo"))

  require("dplyr")
  
  # if mo is not an mo class, result should be the same
  expect_identical(example_isolates %>%
                     mutate(mo = as.character(mo)) %>%
                     first_isolate(col_date = "date",
                                   col_mo = "mo",
                                   col_patient_id = "patient_id"),
                   example_isolates %>%
                     first_isolate(col_date = "date",
                                   col_mo = "mo",
                                   col_patient_id = "patient_id"))
  
  # support for WHONET
  expect_message(example_isolates %>%
                   select(-patient_id) %>%
                   mutate(`First name` = "test",
                          `Last name` = "test", 
                          Sex = "Female") %>% 
                   first_isolate(info = TRUE))

  # missing dates should be no problem
  df <- example_isolates
  df[1:100, "date"] <- NA
  expect_equal(
    sum(
      first_isolate(x = df,
                    col_date = "date",
                    col_patient_id = "patient_id",
                    col_mo = "mo",
                    info = TRUE),
      na.rm = TRUE),
    1305)
  
  # unknown MOs
  test_unknown <- example_isolates
  test_unknown$mo <- ifelse(test_unknown$mo == "B_ESCHR_COLI", "UNKNOWN", test_unknown$mo)
  expect_equal(sum(first_isolate(test_unknown, include_unknown = FALSE)), 
               1045)
  expect_equal(sum(first_isolate(test_unknown, include_unknown = TRUE)),
               1528)
  
  test_unknown$mo <- ifelse(test_unknown$mo == "UNKNOWN", NA, test_unknown$mo)
  expect_equal(sum(first_isolate(test_unknown)),
               1045)
  
  # shortcuts
  expect_identical(filter_first_isolate(example_isolates),
                   subset(example_isolates, first_isolate(example_isolates)))
  ex <- example_isolates
  ex$keyab <- key_antibiotics(ex)
  expect_identical(filter_first_weighted_isolate(example_isolates),
                   subset(example_isolates, first_isolate(ex)))
  
  # notice that all mo's are distinct, so all are TRUE
  expect_true(all(example_isolates %pm>%
                    pm_distinct(mo, .keep_all = TRUE) %pm>%
                    first_isolate(info = TRUE) == TRUE))
  
  # only one isolate, so return fast
  expect_true(first_isolate(data.frame(mo = "Escherichia coli", date = Sys.Date(), patient = "patient"), info = TRUE))

})
```
