## Supplementary material for "AMR - An R Package for Working with Antimicrobial Resistance Data": Source package: test-g.test.R

```
# ==================================================================== #
# TITLE                                                                #
# Antimicrobial Resistance (AMR) Analysis for R                        #
#                                                                      #
# SOURCE                                                               #
# https://github.com/msberends/AMR                                     #
#                                                                      #
# LICENCE                                                              #
# (c) 2018-2021 Berends MS, Luz CF et al.                              #
# Developed at the University of Groningen, the Netherlands, in        #
# collaboration with non-profit organisations Certe Medical            #
# Diagnostics & Advice, and University Medical Center Groningen.       # 
#                                                                      #
# This R package is free software; you can freely use and distribute   #
# it for both personal and commercial purposes under the terms of the  #
# GNU General Public License version 2.0 (GNU GPL-2), as published by  #
# the Free Software Foundation.                                        #
# We created this package for both routine data analysis and academic  #
# research and it was publicly released in the hope that it will be    #
# useful, but it comes WITHOUT ANY WARRANTY OR LIABILITY.              #
#                                                                      #
# Visit our website for the full manual and a complete tutorial about  #
# how to conduct AMR analysis: https://msberends.github.io/AMR/        #
# ==================================================================== #

context("g.test.R")

test_that("G-test works", {
  skip_on_cran()

  # GOODNESS-OF-FIT

  # example 1: clearfield rice vs. red rice
  x <- c(772, 1611, 737)
  expect_equal(g.test(x, p = c(0.25, 0.50, 0.25))$p.value,
               expected = 0.12574,
               tolerance = 0.00001)

  # example 2: red crossbills
  x <- c(1752, 1895)
  expect_equal(g.test(x)$p.value,
               expected = 0.01787343,
               tolerance = 0.00000001)

  expect_error(g.test(0))
  expect_error(g.test(c(0, 1), 0))
  expect_error(g.test(c(1, 2, 3, 4), p = c(0.25, 0.25)))
  expect_error(g.test(c(1, 2, 3, 4), p = c(0.25, 0.25, 0.25, 0.24)))
  expect_warning(g.test(c(1, 2, 3, 4), p = c(0.25, 0.25, 0.25, 0.24), rescale.p = TRUE))

  # INDEPENDENCE

  x <- as.data.frame(
    matrix(data = round(runif(4) * 100000, 0),
           ncol = 2,
           byrow = TRUE)
  )
 
  # fisher.test() is always better for 2x2 tables:
  expect_warning(g.test(x))
  expect_lt(suppressWarnings(g.test(x)$p.value),
            1)

  expect_warning(g.test(x = c(772, 1611, 737),
                        y = c(780, 1560, 780),
                        rescale.p = TRUE))

  expect_error(g.test(matrix(data = c(-1, -2, -3, -4), ncol = 2, byrow = TRUE)))
  expect_error(g.test(matrix(data = c(0, 0, 0, 0), ncol = 2, byrow = TRUE)))

})
```
