## Supplementary material for "AMR - An R Package for Working with Antimicrobial Resistance Data": Source package: test-ggplot_rsi.R

```
# ==================================================================== #
# TITLE                                                                #
# Antimicrobial Resistance (AMR) Analysis for R                        #
#                                                                      #
# SOURCE                                                               #
# https://github.com/msberends/AMR                                     #
#                                                                      #
# LICENCE                                                              #
# (c) 2018-2021 Berends MS, Luz CF et al.                              #
# Developed at the University of Groningen, the Netherlands, in        #
# collaboration with non-profit organisations Certe Medical            #
# Diagnostics & Advice, and University Medical Center Groningen.       # 
#                                                                      #
# This R package is free software; you can freely use and distribute   #
# it for both personal and commercial purposes under the terms of the  #
# GNU General Public License version 2.0 (GNU GPL-2), as published by  #
# the Free Software Foundation.                                        #
# We created this package for both routine data analysis and academic  #
# research and it was publicly released in the hope that it will be    #
# useful, but it comes WITHOUT ANY WARRANTY OR LIABILITY.              #
#                                                                      #
# Visit our website for the full manual and a complete tutorial about  #
# how to conduct AMR analysis: https://msberends.github.io/AMR/        #
# ==================================================================== #

context("ggplot_rsi.R")

test_that("ggplot_rsi works", {

  skip_on_cran()
  
  skip_if_not_installed("ggplot2")

  library(dplyr, warn.conflicts = FALSE)
  library(ggplot2)
  
  pdf(NULL) # prevent Rplots.pdf being created

  # data should be equal
  expect_equal(
    (example_isolates %>% select(AMC, CIP) %>% ggplot_rsi())$data %>% summarise_all(resistance) %>% as.double(),
    example_isolates %>% select(AMC, CIP) %>% summarise_all(resistance) %>% as.double()
  )

  print(example_isolates %>% select(AMC, CIP) %>% ggplot_rsi(x = "interpretation", facet = "antibiotic"))
  print(example_isolates %>% select(AMC, CIP) %>% ggplot_rsi(x = "antibiotic", facet = "interpretation"))

  expect_equal(
    (example_isolates %>% select(AMC, CIP) %>% ggplot_rsi(x = "interpretation", facet = "antibiotic"))$data %>% summarise_all(resistance) %>% as.double(),
    example_isolates %>% select(AMC, CIP) %>% summarise_all(resistance) %>% as.double()
  )

  expect_equal(
    (example_isolates %>% select(AMC, CIP) %>% ggplot_rsi(x = "antibiotic", facet = "interpretation"))$data %>% summarise_all(resistance) %>% as.double(),
    example_isolates %>% select(AMC, CIP) %>% summarise_all(resistance) %>% as.double()
  )

  expect_equal(
    (example_isolates %>% select(AMC, CIP) %>% ggplot_rsi(x = "antibiotic", facet = "interpretation"))$data %>% summarise_all(count_resistant) %>% as.double(),
    example_isolates %>% select(AMC, CIP) %>% summarise_all(count_resistant) %>% as.double()
  )

  # support for scale_type ab and mo
  expect_equal(class((data.frame(mo = as.mo(c("e. coli", "s aureus")),
                                 n = c(40, 100)) %>%
                        ggplot(aes(x = mo, y = n)) +
                        geom_col())$data),
               "data.frame")
  expect_equal(class((data.frame(ab = as.ab(c("amx", "amc")),
                                 n = c(40, 100)) %>%
                        ggplot(aes(x = ab, y = n)) +
                        geom_col())$data),
               "data.frame")

})
```
