## Supplementary material for "AMR - An R Package for Working with Antimicrobial Resistance Data": Source package: test-join_microorganisms.R

```
# ==================================================================== #
# TITLE                                                                #
# Antimicrobial Resistance (AMR) Analysis for R                        #
#                                                                      #
# SOURCE                                                               #
# https://github.com/msberends/AMR                                     #
#                                                                      #
# LICENCE                                                              #
# (c) 2018-2021 Berends MS, Luz CF et al.                              #
# Developed at the University of Groningen, the Netherlands, in        #
# collaboration with non-profit organisations Certe Medical            #
# Diagnostics & Advice, and University Medical Center Groningen.       # 
#                                                                      #
# This R package is free software; you can freely use and distribute   #
# it for both personal and commercial purposes under the terms of the  #
# GNU General Public License version 2.0 (GNU GPL-2), as published by  #
# the Free Software Foundation.                                        #
# We created this package for both routine data analysis and academic  #
# research and it was publicly released in the hope that it will be    #
# useful, but it comes WITHOUT ANY WARRANTY OR LIABILITY.              #
#                                                                      #
# Visit our website for the full manual and a complete tutorial about  #
# how to conduct AMR analysis: https://msberends.github.io/AMR/        #
# ==================================================================== #

context("join_microorganisms.R")

test_that("joins work", {
  skip_on_cran()
  unjoined <- example_isolates
  inner <- example_isolates %>% inner_join_microorganisms()
  left <- example_isolates %>% left_join_microorganisms()
  semi <- example_isolates %>% semi_join_microorganisms()
  anti <- example_isolates %>% anti_join_microorganisms()
  suppressWarnings(right <- example_isolates %>% right_join_microorganisms())
  suppressWarnings(full <- example_isolates %>% full_join_microorganisms())

  expect_true(ncol(unjoined) < ncol(inner))
  expect_true(nrow(unjoined) == nrow(inner))

  expect_true(ncol(unjoined) < ncol(left))
  expect_true(nrow(unjoined) == nrow(left))

  expect_true(ncol(semi) == ncol(semi))
  expect_true(nrow(semi) == nrow(semi))

  expect_true(nrow(anti) == 0)

  expect_true(nrow(unjoined) < nrow(right))
  expect_true(nrow(unjoined) < nrow(full))

  expect_equal(nrow(inner_join_microorganisms("B_ESCHR_COLI")), 1)
  expect_equal(nrow(inner_join_microorganisms("B_ESCHR_COLI", by = c("mo" = "mo"))), 1)

  expect_equal(nrow(left_join_microorganisms("B_ESCHR_COLI")), 1)

  expect_equal(nrow(semi_join_microorganisms("B_ESCHR_COLI")), 1)
  expect_equal(nrow(anti_join_microorganisms("B_ESCHR_COLI")), 0)

  expect_warning(right_join_microorganisms("B_ESCHR_COLI"))
  expect_warning(full_join_microorganisms("B_ESCHR_COLI"))
  
  library(dplyr, warn.conflicts = FALSE)
  x <- tibble(bact = as.mo("E.coli"))
  expect_warning(left_join_microorganisms(x %>% group_by(bact), "bact"))

})
```
