## Supplementary material for "AMR - An R Package for Working with Antimicrobial Resistance Data": Source package: test-mdro.R

```
# ==================================================================== #
# TITLE                                                                #
# Antimicrobial Resistance (AMR) Analysis for R                        #
#                                                                      #
# SOURCE                                                               #
# https://github.com/msberends/AMR                                     #
#                                                                      #
# LICENCE                                                              #
# (c) 2018-2021 Berends MS, Luz CF et al.                              #
# Developed at the University of Groningen, the Netherlands, in        #
# collaboration with non-profit organisations Certe Medical            #
# Diagnostics & Advice, and University Medical Center Groningen.       # 
#                                                                      #
# This R package is free software; you can freely use and distribute   #
# it for both personal and commercial purposes under the terms of the  #
# GNU General Public License version 2.0 (GNU GPL-2), as published by  #
# the Free Software Foundation.                                        #
# We created this package for both routine data analysis and academic  #
# research and it was publicly released in the hope that it will be    #
# useful, but it comes WITHOUT ANY WARRANTY OR LIABILITY.              #
#                                                                      #
# Visit our website for the full manual and a complete tutorial about  #
# how to conduct AMR analysis: https://msberends.github.io/AMR/        #
# ==================================================================== #

context("mdro.R")

test_that("mdro works", {
  
  skip_on_cran()
  
  expect_error(suppressWarnings(mdro(example_isolates, country = "invalid", col_mo = "mo", info = TRUE)))
  expect_error(suppressWarnings(mdro(example_isolates, country = "fr", info = TRUE)))
  expect_error(mdro(example_isolates, guideline = c("BRMO", "MRGN"), info = TRUE))
  expect_error(mdro(example_isolates, col_mo = "invalid", info = TRUE))

  outcome <- suppressWarnings(mdro(example_isolates))
  outcome <- mdro(example_isolates, "eucast3.1", info = TRUE)
  outcome <- eucast_exceptional_phenotypes(example_isolates, info = TRUE)
  # check class
  expect_equal(class(outcome), c("ordered", "factor"))

  outcome <- mdro(example_isolates, "nl", info = TRUE)
  # check class
  expect_equal(class(outcome), c("ordered", "factor"))

  library(dplyr)
  # example_isolates should have these finding using Dutch guidelines
  expect_equal(as.double(table(outcome)),
               c(1969, 25, 6)) # 1969 neg, 25 unconfirmed, 6 pos

  expect_equal(brmo(example_isolates, info = FALSE),
               mdro(example_isolates, guideline = "BRMO", info = FALSE))

  # test Dutch P. aeruginosa MDRO
  expect_equal(
    as.character(mdro(data.frame(mo = as.mo("P. aeruginosa"),
                                 cfta = "S",
                                 cipr = "S",
                                 mero = "S",
                                 imip = "S",
                                 gent = "S",
                                 tobr = "S",
                                 pita = "S"),
                      guideline = "BRMO",
                      col_mo = "mo",
                      info = FALSE)),
    "Negative")
  expect_equal(
    as.character(mdro(data.frame(mo = as.mo("P. aeruginosa"),
                                 cefta = "R",
                                 cipr = "R",
                                 mero = "R",
                                 imip = "R",
                                 gent = "R",
                                 tobr = "R",
                                 pita = "R"),
                      guideline = "BRMO",
                      col_mo = "mo",
                      info = FALSE)),
    "Positive")
  
  # German 3MRGN and 4MRGN
  expect_equal(as.character(mrgn(
    data.frame(mo = c("E. coli", "E. coli", "K. pneumoniae", "E. coli",
                      "A. baumannii", "A. baumannii", "A. baumannii",
                      "P. aeruginosa", "P. aeruginosa", "P. aeruginosa"), 
               PIP = c("S", "R", "R", "S",
                       "S", "R", "R",
                       "S", "R", "R"),
               CTX = c("S", "R", "R", "S",
                       "R", "R", "R",
                       "R", "R", "R"),
               IPM = c("S", "R", "S", "R",
                       "R", "R", "S",
                       "S", "R", "R"),
               CIP = c("S", "R", "R", "S",
                       "R", "R", "R",
                       "R", "S", "R"),
               stringsAsFactors = FALSE))),
    c("Negative", "4MRGN", "3MRGN", "4MRGN",  "4MRGN",  "4MRGN", "3MRGN", "Negative", "3MRGN", "4MRGN"))
  
  # MDR TB
  expect_equal(
    # select only rifampicine, mo will be determined automatically (as M. tuberculosis),
    # number of mono-resistant strains should be equal to number of rifampicine-resistant strains
    as.double(table(mdr_tb(example_isolates[, "RIF", drop = FALSE])))[2],
    count_R(example_isolates$RIF))

  sample_rsi <- function() {
    sample(c("S", "I", "R"),
           size = 5000,
           prob = c(0.5, 0.1, 0.4),
           replace = TRUE)
  }
  x <- data.frame(rifampicin = sample_rsi(),
                     inh = sample_rsi(),
                     gatifloxacin = sample_rsi(),
                     eth = sample_rsi(),
                     pza = sample_rsi(),
                     MFX = sample_rsi(),
                     KAN = sample_rsi())
  expect_gt(n_distinct(mdr_tb(x)), 2)
  
  # check the guideline by Magiorakos  et al. (2012), the default guideline
  stau <- data.frame(mo = c("S. aureus", "S. aureus", "S. aureus", "S. aureus"), 
                     GEN = c("R", "R", "S", "R"), 
                     RIF = c("S", "R", "S", "R"), 
                     CPT = c("S", "R", "R", "R"), 
                     OXA = c("S", "R", "R", "R"), 
                     CIP = c("S", "S", "R", "R"), 
                     MFX = c("S", "S", "R", "R"),
                     SXT = c("S", "S", "R", "R"), 
                     FUS = c("S", "S", "R", "R"),     
                     VAN = c("S", "S", "R", "R"), 
                     TEC = c("S", "S", "R", "R"),     
                     TLV = c("S", "S", "R", "R"), 
                     TGC = c("S", "S", "R", "R"),     
                     CLI = c("S", "S", "R", "R"), 
                     DAP = c("S", "S", "R", "R"),     
                     ERY = c("S", "S", "R", "R"), 
                     LNZ = c("S", "S", "R", "R"),     
                     CHL = c("S", "S", "R", "R"), 
                     FOS = c("S", "S", "R", "R"),     
                     QDA = c("S", "S", "R", "R"), 
                     TCY = c("S", "S", "R", "R"),     
                     DOX = c("S", "S", "R", "R"), 
                     MNO = c("S", "S", "R", "R"),
                     stringsAsFactors = FALSE)
  expect_equal(as.integer(mdro(stau)), c(1:4))
  expect_s3_class(mdro(stau, verbose = TRUE), "data.frame")
  
  ente <- data.frame(mo = c("Enterococcus", "Enterococcus", "Enterococcus", "Enterococcus"), 
                     GEH = c("R", "R", "S", "R"), 
                     STH = c("S", "R", "S", "R"), 
                     IPM = c("S", "R", "R", "R"), 
                     MEM = c("S", "R", "R", "R"), 
                     DOR = c("S", "S", "R", "R"), 
                     CIP = c("S", "S", "R", "R"), 
                     LVX = c("S", "S", "R", "R"), 
                     MFX = c("S", "S", "R", "R"),     
                     VAN = c("S", "S", "R", "R"), 
                     TEC = c("S", "S", "R", "R"),     
                     TGC = c("S", "S", "R", "R"), 
                     DAP = c("S", "S", "R", "R"),     
                     LNZ = c("S", "S", "R", "R"), 
                     AMP = c("S", "S", "R", "R"),     
                     QDA = c("S", "S", "R", "R"), 
                     DOX = c("S", "S", "R", "R"),     
                     MNO = c("S", "S", "R", "R"),
                     stringsAsFactors = FALSE)
  expect_equal(as.integer(mdro(ente)), c(1:4))
  expect_s3_class(mdro(ente, verbose = TRUE), "data.frame")
  
  entero <- data.frame(mo = c("E. coli", "E. coli", "E. coli", "E. coli"),
                       GEN = c("R", "R", "S", "R"), TOB = c("S", "R", "S", "R"), 
                       AMK = c("S", "R", "R", "R"), NET = c("S", "R", "R", "R"), 
                       CPT = c("S", "R", "R", "R"), TCC = c("S", "R", "R", "R"), 
                       TZP = c("S", "S", "R", "R"), ETP = c("S", "S", "R", "R"), 
                       IPM = c("S", "S", "R", "R"), MEM = c("S", "S", "R", "R"), 
                       DOR = c("S", "S", "R", "R"), CZO = c("S", "S", "R", "R"), 
                       CXM = c("S", "S", "R", "R"), CTX = c("S", "S", "R", "R"), 
                       CAZ = c("S", "S", "R", "R"), FEP = c("S", "S", "R", "R"), 
                       FOX = c("S", "S", "R", "R"), CTT = c("S", "S", "R", "R"), 
                       CIP = c("S", "S", "R", "R"), SXT = c("S", "S", "R", "R"), 
                       TGC = c("S", "S", "R", "R"), ATM = c("S", "S", "R", "R"), 
                       AMP = c("S", "S", "R", "R"), AMC = c("S", "S", "R", "R"), 
                       SAM = c("S", "S", "R", "R"), CHL = c("S", "S", "R", "R"), 
                       FOS = c("S", "S", "R", "R"), COL = c("S", "S", "R", "R"), 
                       TCY = c("S", "S", "R", "R"), DOX = c("S", "S", "R", "R"), 
                       MNO = c("S", "S", "R", "R"),
                       stringsAsFactors = FALSE)
  expect_equal(as.integer(mdro(entero)), c(1:4))
  expect_s3_class(mdro(entero, verbose = TRUE), "data.frame")
  
  pseud <- data.frame(mo = c("P. aeruginosa", "P. aeruginosa", "P. aeruginosa", "P. aeruginosa"),
                      GEN = c("R", "R", "S", "R"), TOB = c("S", "S", "S", "R"), 
                      AMK = c("S", "S", "R", "R"), NET = c("S", "S", "R", "R"),
                      IPM = c("S", "R", "R", "R"), MEM = c("S", "S", "R", "R"), 
                      DOR = c("S", "S", "R", "R"), CAZ = c("S", "S", "R", "R"), 
                      FEP = c("S", "R", "R", "R"), CIP = c("S", "S", "R", "R"), 
                      LVX = c("S", "S", "R", "R"), TCC = c("S", "S", "R", "R"), 
                      TZP = c("S", "S", "R", "R"), ATM = c("S", "S", "R", "R"), 
                      FOS = c("S", "S", "R", "R"), COL = c("S", "S", "R", "R"), 
                      PLB = c("S", "S", "R", "R"),
                      stringsAsFactors = FALSE)
  expect_equal(as.integer(mdro(pseud)), c(1:4))
  expect_s3_class(mdro(pseud, verbose = TRUE), "data.frame")
  
  acin <- data.frame(mo = c("A. baumannii", "A. baumannii", "A. baumannii", "A. baumannii"), 
                     GEN = c("R", "R", "S", "R"), TOB = c("S", "R", "S", "R"), 
                     AMK = c("S", "R", "R", "R"), NET = c("S", "R", "R", "R"), 
                     IPM = c("S", "S", "R", "R"), MEM = c("S", "R", "R", "R"),
                     DOR = c("S", "S", "R", "R"), CIP = c("S", "S", "R", "R"), 
                     LVX = c("S", "S", "R", "R"), TZP = c("S", "S", "R", "R"), 
                     TCC = c("S", "S", "R", "R"), CTX = c("S", "S", "R", "R"), 
                     CRO = c("S", "S", "R", "R"), CAZ = c("S", "S", "R", "R"), 
                     FEP = c("S", "R", "R", "R"), SXT = c("S", "S", "R", "R"), 
                     SAM = c("S", "S", "R", "R"), COL = c("S", "S", "R", "R"), 
                     PLB = c("S", "S", "R", "R"), TCY = c("S", "S", "R", "R"), 
                     DOX = c("S", "S", "R", "R"), MNO = c("S", "S", "R", "R"),
                     stringsAsFactors = FALSE)
  expect_equal(as.integer(mdro(acin)), c(1:4))
  expect_s3_class(mdro(acin, verbose = TRUE), "data.frame")
  
})
```
