## Supplementary material for "AMR - An R Package for Working with Antimicrobial Resistance Data": Source package: test-mic.R

```
# ==================================================================== #
# TITLE                                                                #
# Antimicrobial Resistance (AMR) Analysis for R                        #
#                                                                      #
# SOURCE                                                               #
# https://github.com/msberends/AMR                                     #
#                                                                      #
# LICENCE                                                              #
# (c) 2018-2021 Berends MS, Luz CF et al.                              #
# Developed at the University of Groningen, the Netherlands, in        #
# collaboration with non-profit organisations Certe Medical            #
# Diagnostics & Advice, and University Medical Center Groningen.       # 
#                                                                      #
# This R package is free software; you can freely use and distribute   #
# it for both personal and commercial purposes under the terms of the  #
# GNU General Public License version 2.0 (GNU GPL-2), as published by  #
# the Free Software Foundation.                                        #
# We created this package for both routine data analysis and academic  #
# research and it was publicly released in the hope that it will be    #
# useful, but it comes WITHOUT ANY WARRANTY OR LIABILITY.              #
#                                                                      #
# Visit our website for the full manual and a complete tutorial about  #
# how to conduct AMR analysis: https://msberends.github.io/AMR/        #
# ==================================================================== #

context("mic.R")

test_that("mic works", {
  skip_on_cran()
  expect_true(as.mic(8) == as.mic("8"))
  expect_true(as.mic("1") > as.mic("<=0.0625"))
  expect_true(as.mic("1") < as.mic(">=32"))
  expect_true(is.mic(as.mic(8)))

  expect_equal(as.double(as.mic(">=32")), 32)
  expect_equal(as.numeric(as.mic(">=32")), 32)
  expect_equal(as.integer(as.mic(">=32")), 32)
  expect_equal(suppressWarnings(as.logical(as.mic("INVALID VALUE"))), NA)

  # all levels should be valid MICs
  x <- as.mic(c(2, 4))
  expect_s3_class(x[1], "mic")
  expect_s3_class(x[[1]], "mic")
  expect_s3_class(c(x[1], x[9]), "mic")
  expect_s3_class(unique(x[1], x[9]), "mic")
  expect_s3_class(droplevels(c(x[1], x[9])), "mic")
  x[2] <- 32
  expect_s3_class(x, "mic")
  expect_warning(as.mic("INVALID VALUE"))
  
  pdf(NULL) # prevent Rplots.pdf being created
  expect_silent(barplot(as.mic(c(1, 2, 4, 8))))
  expect_silent(plot(as.mic(c(1, 2, 4, 8))))
  expect_output(print(as.mic(c(1, 2, 4, 8))))
  
  expect_equal(summary(as.mic(c(2, 8))), 
               structure(c("Class" = "mic",
                           "<NA>" = "0",
                           "Min." = "2",
                           "Max." = "8"), class = c("summaryDefault", "table")))
  
  library(dplyr, warn.conflicts = FALSE)
  expect_output(print(tibble(m = as.mic(2:4))))
})
```
