## Supplementary material for "AMR - An R Package for Working with Antimicrobial Resistance Data": Source package: test-mo.R

```
# ==================================================================== #
# TITLE                                                                #
# Antimicrobial Resistance (AMR) Analysis for R                        #
#                                                                      #
# SOURCE                                                               #
# https://github.com/msberends/AMR                                     #
#                                                                      #
# LICENCE                                                              #
# (c) 2018-2021 Berends MS, Luz CF et al.                              #
# Developed at the University of Groningen, the Netherlands, in        #
# collaboration with non-profit organisations Certe Medical            #
# Diagnostics & Advice, and University Medical Center Groningen.       # 
#                                                                      #
# This R package is free software; you can freely use and distribute   #
# it for both personal and commercial purposes under the terms of the  #
# GNU General Public License version 2.0 (GNU GPL-2), as published by  #
# the Free Software Foundation.                                        #
# We created this package for both routine data analysis and academic  #
# research and it was publicly released in the hope that it will be    #
# useful, but it comes WITHOUT ANY WARRANTY OR LIABILITY.              #
#                                                                      #
# Visit our website for the full manual and a complete tutorial about  #
# how to conduct AMR analysis: https://msberends.github.io/AMR/        #
# ==================================================================== #

context("mo.R")

test_that("as.mo works", {
  
  skip_on_cran()
  
  library(dplyr, warn.conflicts = FALSE)
 
  MOs <- microorganisms %>% filter(!is.na(mo), nchar(mo) > 3)
  expect_identical(as.character(MOs$mo), as.character(as.mo(MOs$mo)))
  
  expect_identical(
    as.character(as.mo(c("E. coli", "H. influenzae"))),
    c("B_ESCHR_COLI", "B_HMPHL_INFL"))
  
  expect_equal(as.character(as.mo("Escherichia coli")), "B_ESCHR_COLI")
  expect_equal(as.character(as.mo("Escherichia  coli")), "B_ESCHR_COLI")
  expect_equal(as.character(as.mo(112283007)), "B_ESCHR_COLI")
  expect_equal(as.character(as.mo("Escherichia  species")), "B_ESCHR")
  expect_equal(as.character(as.mo("Escherichia")), "B_ESCHR")
  expect_equal(as.character(as.mo("Esch spp.")), "B_ESCHR")
  expect_equal(as.character(as.mo(" B_ESCHR_COLI ")), "B_ESCHR_COLI")
  expect_equal(as.character(as.mo("e coli")), "B_ESCHR_COLI") # not Campylobacter
  expect_equal(as.character(as.mo("klpn")), "B_KLBSL_PNMN")
  expect_equal(as.character(as.mo("Klebsiella")), "B_KLBSL")
  expect_equal(as.character(as.mo("K. pneu rhino")), "B_KLBSL_PNMN_RHNS") # K. pneumoniae subspp. rhinoscleromatis
  expect_equal(as.character(as.mo("Bartonella")), "B_BRTNL")
  expect_equal(as.character(as.mo("C. difficile")), "B_CRDDS_DFFC")
  expect_equal(as.character(as.mo("L. pneumophila")), "B_LGNLL_PNMP")
  expect_equal(as.character(as.mo("Strepto")), "B_STRPT")
  expect_equal(as.character(as.mo("Streptococcus")), "B_STRPT") # not Peptostreptoccus
  expect_equal(as.character(as.mo("Estreptococos grupo B")), "B_STRPT_GRPB")
  expect_equal(as.character(as.mo("Group B Streptococci")), "B_STRPT_GRPB")
  expect_equal(as.character(suppressWarnings(as.mo("B_STRPT_PNE"))), "B_STRPT_PNMN") # old MO code (<=v0.8.0)
  expect_equal(as.character(as.mo(c("mycobacterie", "mycobakterium"))), c("B_MYCBC", "B_MYCBC"))
  
  expect_equal(as.character(as.mo(c("GAS", "GBS", "a MGS", "haemoly strep"))), c("B_STRPT_GRPA", "B_STRPT_GRPB", "B_STRPT_MILL", "B_STRPT_HAEM"))

  
  expect_equal(as.character(as.mo("S. pyo")), "B_STRPT_PYGN") # not Actinomyces pyogenes
  
  # GLIMS
  expect_equal(as.character(as.mo("bctfgr")), "B_BCTRD_FRGL")
  
  expect_equal(as.character(as.mo("MRSE")), "B_STPHY_EPDR")
  expect_equal(as.character(as.mo("VRE")), "B_ENTRC")
  expect_equal(as.character(as.mo("MRPA")), "B_PSDMN_AERG")
  expect_equal(as.character(as.mo("PISP")), "B_STRPT_PNMN")
  expect_equal(as.character(as.mo("PRSP")), "B_STRPT_PNMN")
  expect_equal(as.character(as.mo("VISP")), "B_STRPT_PNMN")
  expect_equal(as.character(as.mo("VRSP")), "B_STRPT_PNMN")
  
  expect_equal(as.character(as.mo("CNS")), "B_STPHY_CONS")
  expect_equal(as.character(as.mo("CoNS")), "B_STPHY_CONS")
  expect_equal(as.character(as.mo("CPS")), "B_STPHY_COPS")
  expect_equal(as.character(as.mo("CoPS")), "B_STPHY_COPS")
  expect_equal(as.character(as.mo("VGS")), "B_STRPT_VIRI")
  expect_equal(as.character(as.mo("streptococcus milleri")), "B_STRPT_MILL")
  
  
  expect_equal(as.character(as.mo(c("Gram negative", "Gram positive"))), c("B_GRAMN", "B_GRAMP"))
  
  # prevalent MO
  expect_identical(
    suppressWarnings(as.character(
      as.mo(c("stau",
              "STAU",
              "staaur",
              "S. aureus",
              "S aureus",
              "Sthafilokkockus aureeuzz",
              "Staphylococcus aureus",
              "MRSA",
              "VISA")))),
    rep("B_STPHY_AURS", 9))
  expect_identical(
    as.character(
      as.mo(c("EHEC", "EPEC", "EIEC", "STEC", "ATEC", "UPEC"))),
    rep("B_ESCHR_COLI", 6))
  # unprevalent MO
  expect_identical(
    as.character(
      as.mo(c("parnod",
              "P. nodosa",
              "P nodosa",
              "Paraburkholderia nodosa"))),
    rep("B_PRBRK_NODS", 4))
  
  # empty values
  expect_identical(as.character(as.mo(c("", "  ", NA, NaN))), rep(NA_character_, 4))
  expect_identical(as.character(as.mo("  ")), NA_character_)
  # too few characters
  expect_warning(as.mo("ab"))
  
  expect_equal(suppressWarnings(as.character(as.mo(c("Qq species", "", "CRSM", "K. pneu rhino", "esco")))),
               c("UNKNOWN", NA_character_, "B_STNTR_MLTP", "B_KLBSL_PNMN_RHNS", "B_ESCHR_COLI"))
  
  # check for Becker classification
  expect_identical(as.character(as.mo("S. epidermidis",  Becker = FALSE)), "B_STPHY_EPDR")
  expect_identical(as.character(as.mo("S. epidermidis",  Becker = TRUE)),  "B_STPHY_CONS")
  expect_identical(as.character(as.mo("STAEPI",          Becker = TRUE)),  "B_STPHY_CONS")
  expect_identical(as.character(as.mo("S. intermedius",  Becker = FALSE)), "B_STPHY_INTR")
  expect_identical(as.character(as.mo("Sta intermedius", Becker = FALSE)), "B_STPHY_INTR")
  expect_identical(as.character(as.mo("Sta intermedius", Becker = TRUE)),  "B_STPHY_COPS")
  expect_identical(as.character(as.mo("STAINT",          Becker = TRUE)),  "B_STPHY_COPS")
  # aureus must only be influenced if Becker = "all"
  expect_identical(as.character(as.mo("STAAUR", Becker = FALSE)), "B_STPHY_AURS")
  expect_identical(as.character(as.mo("STAAUR", Becker = TRUE)),  "B_STPHY_AURS")
  expect_identical(as.character(as.mo("STAAUR", Becker = "all")), "B_STPHY_COPS")
  
  # check for Lancefield classification
  expect_identical(as.character(as.mo("S. pyogenes", Lancefield = FALSE)),    "B_STRPT_PYGN")
  expect_identical(as.character(as.mo("S. pyogenes", Lancefield = TRUE)),     "B_STRPT_GRPA")
  expect_identical(as.character(as.mo("STCPYO",      Lancefield = TRUE)),     "B_STRPT_GRPA") # group A
  expect_identical(as.character(as.mo("S. agalactiae",  Lancefield = FALSE)), "B_STRPT_AGLC")
  expect_identical(as.character(as.mo("S. agalactiae",  Lancefield = TRUE)),  "B_STRPT_GRPB") # group B
  expect_identical(as.character(suppressWarnings(as.mo("estreptococos grupo B"))), "B_STRPT_GRPB")
  expect_identical(as.character(as.mo("S. equisimilis", Lancefield = FALSE)), "B_STRPT_DYSG_EQSM")
  expect_identical(as.character(as.mo("S. equisimilis", Lancefield = TRUE)),  "B_STRPT_GRPC") # group C
  # Enterococci must only be influenced if Lancefield = "all"
  expect_identical(as.character(as.mo("E. faecium", Lancefield = FALSE)),     "B_ENTRC_FACM")
  expect_identical(as.character(as.mo("E. faecium", Lancefield = TRUE)),      "B_ENTRC_FACM")
  expect_identical(as.character(as.mo("E. faecium", Lancefield = "all")),     "B_STRPT_GRPD") # group D
  expect_identical(as.character(as.mo("S. anginosus",   Lancefield = FALSE)), "B_STRPT_ANGN")
  expect_identical(as.character(as.mo("S. anginosus",   Lancefield = TRUE)),  "B_STRPT_GRPF") # group F
  expect_identical(as.character(as.mo("S. sanguinis",   Lancefield = FALSE)), "B_STRPT_SNGN")
  expect_identical(as.character(as.mo("S. sanguinis",   Lancefield = TRUE)),  "B_STRPT_GRPH") # group H
  expect_identical(as.character(as.mo("S. salivarius",  Lancefield = FALSE)), "B_STRPT_SLVR")
  expect_identical(as.character(as.mo("S. salivarius",  Lancefield = TRUE)),  "B_STRPT_GRPK") # group K

  # select with one column
  expect_identical(
    example_isolates[1:10, ] %>%
      left_join_microorganisms() %>%
      select(genus) %>%
      as.mo() %>%
      as.character(),
    c("B_ESCHR", "B_ESCHR", "B_STPHY", "B_STPHY", "B_STPHY",
      "B_STPHY", "B_STPHY", "B_STPHY", "B_STPHY", "B_STPHY"))
  
  # select with two columns
  expect_identical(
    example_isolates[1:10, ] %>%
      pull(mo),
    example_isolates[1:10, ] %>%
      left_join_microorganisms() %>%
      select(genus, species) %>%
      as.mo())

  # unknown results
  expect_warning(as.mo(c("INVALID", "Yeah, unknown")))
  
  # too many columns
  expect_error(example_isolates %>% select(1:3) %>% as.mo())
  
  # print
  expect_output(print(as.mo(c("B_ESCHR_COLI", NA))))
  
  # test pull
  expect_equal(nrow(example_isolates %>% mutate(mo = as.mo(mo))),
               2000)
  
  # test data.frame
  expect_equal(nrow(data.frame(test = as.mo("B_ESCHR_COLI"))),
               1)
  
  # check empty values
  expect_equal(as.character(suppressWarnings(as.mo(""))),
               NA_character_)
  
  # check less prevalent MOs
  expect_equal(as.character(as.mo("Gomphosphaeria aponina delicatula")), "B_GMPHS_APNN_DLCT")
  expect_equal(as.character(as.mo("Gomphosphaeria apo del")), "B_GMPHS_APNN_DLCT")
  expect_equal(as.character(as.mo("G apo deli")), "B_GMPHS_APNN_DLCT")
  expect_equal(as.character(as.mo("Gomphosphaeria  aponina")), "B_GMPHS_APNN")
  expect_equal(as.character(as.mo("Gomphosphaeria  species")), "B_GMPHS")
  expect_equal(as.character(as.mo("Gomphosphaeria")), "B_GMPHS")
  expect_equal(as.character(as.mo(" B_GMPHS_APNN ")), "B_GMPHS_APNN")
  expect_equal(as.character(as.mo("g aponina")), "B_GMPHS_APNN")
  
  # check old names
  expect_equal(suppressMessages(as.character(as.mo("Escherichia blattae"))), "B_SHMWL_BLTT")
  print(mo_renamed())
  expect_equal(suppressMessages(as.character(as.mo(c("E. coli", "Chlamydo psittaci")))), c("B_ESCHR_COLI", "B_CHLMY_PSTT"))
  
  # check uncertain names
  expect_equal(suppressMessages(as.character(as.mo("staaur extratest", allow_uncertain = TRUE))), "B_STPHY_AURS")
  expect_equal(suppressWarnings(as.character(as.mo("staaur extratest", allow_uncertain = FALSE))), "UNKNOWN")
  expect_message(as.mo("e coli extra_text", allow_uncertain = TRUE))
  expect_equal(suppressMessages(as.character(as.mo("unexisting aureus", allow_uncertain = 3))), "B_STPHY_AURS")
  expect_equal(suppressMessages(as.character(as.mo("unexisting staphy", allow_uncertain = 3))), "B_STPHY_COPS")
  expect_equal(suppressMessages(as.character(as.mo(c("s aur THISISATEST", "Staphylococcus aureus unexisting"), allow_uncertain = 3))), c("B_STPHY_AURS_ANRB", "B_STPHY_AURS_ANRB"))
  
  # predefined reference_df
  expect_equal(as.character(as.mo("TestingOwnID",
                                  reference_df = data.frame(mycol = "TestingOwnID", mo = "B_ESCHR_COLI"))),
               "B_ESCHR_COLI")
  expect_equal(as.character(as.mo(c("TestingOwnID", "E. coli"),
                                  reference_df = data.frame(mycol = "TestingOwnID", mo = "B_ESCHR_COLI"))),
               c("B_ESCHR_COLI", "B_ESCHR_COLI"))
  expect_warning(as.mo("TestingOwnID", reference_df = NULL))
  expect_error(as.mo("E. coli", reference_df = data.frame(mycol = "TestingOwnID")))
  
  # combination of existing mo and other code
  expect_identical(as.character(as.mo(c("B_ESCHR_COL", "ESCCOL"))),
                   c("B_ESCHR_COLI", "B_ESCHR_COLI"))
  
  # from different sources
  expect_equal(as.character(as.mo(
    c("PRTMIR", "bclcer", "B_ESCHR_COLI"))),
    c("B_PROTS_MRBL", "B_BCLLS_CERS", "B_ESCHR_COLI"))
  
  # hard to find
  expect_equal(as.character(suppressMessages(as.mo(
    c("Microbacterium paraoxidans",
      "Streptococcus suis (bovis gr)",
      "Raoultella (here some text) terrigena")))),
    c("B_MCRBC_PRXY", "B_STRPT_SUIS", "B_RLTLL_TRRG"))
  expect_output(print(mo_uncertainties()))
  x <- as.mo("S. aur")
  # many hits
  expect_output(print(mo_uncertainties()))

  # Salmonella (City) are all actually Salmonella enterica spp (City)
  expect_equal(suppressMessages(mo_name(c("Salmonella Goettingen", "Salmonella Typhimurium", "Salmonella Group A"))),
               c("Salmonella enterica", "Salmonella enterica", "Salmonella"))
  
  # no virusses
  expect_equal(as.character(as.mo("Virus")), NA_character_)
  
  # summary
  expect_equal(length(summary(example_isolates$mo)), 6)
  
  # WHONET codes and NA/NaN
  expect_equal(as.character(as.mo(c("xxx", "na", "nan"), debug = TRUE)),
               rep(NA_character_, 3))
  expect_equal(as.character(as.mo("con")), "UNKNOWN")
  expect_equal(as.character(as.mo("xxx")), NA_character_)
  expect_equal(as.character(as.mo(c("xxx", "con", "eco"))), c(NA_character_, "UNKNOWN", "B_ESCHR_COLI"))
  expect_equal(as.character(as.mo(c("other", "none", "unknown"))),
               rep("UNKNOWN", 3))
  
  expect_null(mo_failures())
  expect_true(example_isolates %>% pull(mo) %>% is.mo())
  
  expect_error(translate_allow_uncertain(5))

  # debug mode
  expect_output(print(suppressMessages(suppressWarnings(as.mo("kshgcjkhsdgkshjdfsfvsdfv", debug = TRUE, allow_uncertain = 3)))))
  
  # ..coccus
  expect_equal(as.character(as.mo(c("meningococ", "gonococ", "pneumococ"))), 
               c("B_NESSR_MNNG", "B_NESSR_GNRR", "B_STRPT_PNMN"))
  # yeasts and fungi
  expect_equal(suppressWarnings(as.character(as.mo(c("yeasts", "fungi")))), 
               c("F_YEAST", "F_FUNGUS"))
  
  # print tibble
  expect_output(print(tibble(mo = as.mo("B_ESCHR_COLI"))))
  
  # assigning and subsetting
  x <- example_isolates$mo
  expect_s3_class(x[1], "mo")
  expect_s3_class(x[[1]], "mo")
  expect_s3_class(c(x[1], x[9]), "mo")
  expect_warning(x[1] <- "invalid code")
  expect_warning(x[[1]] <- "invalid code")
  expect_warning(c(x[1], "test"))
  
  # ignoring patterns
  expect_equal(as.character(as.mo(c("E. coli", "E. coli ignorethis"), ignore_pattern = "this")),
               c("B_ESCHR_COLI", NA))
  
  # frequency tables
  if (require("cleaner")) {
    expect_s3_class(cleaner::freq(example_isolates$mo), "freq")
  }
  
})
```
