## Supplementary material for "AMR - An R Package for Working with Antimicrobial Resistance Data": Source package: test-mo_property.R

```
# ==================================================================== #
# TITLE                                                                #
# Antimicrobial Resistance (AMR) Analysis for R                        #
#                                                                      #
# SOURCE                                                               #
# https://github.com/msberends/AMR                                     #
#                                                                      #
# LICENCE                                                              #
# (c) 2018-2021 Berends MS, Luz CF et al.                              #
# Developed at the University of Groningen, the Netherlands, in        #
# collaboration with non-profit organisations Certe Medical            #
# Diagnostics & Advice, and University Medical Center Groningen.       # 
#                                                                      #
# This R package is free software; you can freely use and distribute   #
# it for both personal and commercial purposes under the terms of the  #
# GNU General Public License version 2.0 (GNU GPL-2), as published by  #
# the Free Software Foundation.                                        #
# We created this package for both routine data analysis and academic  #
# research and it was publicly released in the hope that it will be    #
# useful, but it comes WITHOUT ANY WARRANTY OR LIABILITY.              #
#                                                                      #
# Visit our website for the full manual and a complete tutorial about  #
# how to conduct AMR analysis: https://msberends.github.io/AMR/        #
# ==================================================================== #

context("mo_property.R")

test_that("mo_property works", {
  
  skip_on_cran()
  
  expect_equal(mo_kingdom("Escherichia coli"), "Bacteria")
  expect_equal(mo_kingdom("Escherichia coli"), mo_domain("Escherichia coli"))
  expect_equal(mo_phylum("Escherichia coli"), "Proteobacteria")
  expect_equal(mo_class("Escherichia coli"), "Gammaproteobacteria")
  expect_equal(mo_order("Escherichia coli"), "Enterobacterales")
  expect_equal(mo_family("Escherichia coli"), "Enterobacteriaceae")
  expect_equal(mo_genus("Escherichia coli"), "Escherichia")
  expect_equal(mo_species("Escherichia coli"), "coli")
  expect_equal(mo_subspecies("Escherichia coli"), "")
  expect_equal(mo_fullname("Escherichia coli"), "Escherichia coli")
  expect_equal(mo_name("Escherichia coli"), "Escherichia coli")
  expect_equal(mo_type("Escherichia coli", language = "en"), "Bacteria")
  expect_equal(mo_gramstain("Escherichia coli", language = "en"), "Gram-negative")
  expect_equal(class(mo_taxonomy("Escherichia coli")), "list")
  expect_equal(names(mo_taxonomy("Escherichia coli")), c("kingdom", "phylum", "class", "order",
                                                "family", "genus", "species", "subspecies"))
  expect_equal(mo_synonyms("Escherichia coli"), NULL)
  expect_gt(length(mo_synonyms("Candida albicans")), 1)
  expect_equal(class(mo_synonyms(c("Candida albicans", "Escherichia coli"))), "list")
  expect_equal(names(mo_info("Escherichia coli")), c("kingdom", "phylum", "class", "order",
                                            "family", "genus", "species", "subspecies",
                                            "synonyms", "gramstain", "url", "ref"))
  expect_equal(class(mo_info(c("Escherichia coli", "Staphylococcus aureus"))), "list")

  expect_equal(mo_ref("Escherichia coli"), "Castellani et al., 1919")
  expect_equal(mo_authors("Escherichia coli"), "Castellani et al.")
  expect_equal(mo_year("Escherichia coli"), 1919)

  expect_equal(mo_shortname("Escherichia coli"), "E. coli")
  expect_equal(mo_shortname("Escherichia"), "Escherichia")
  expect_equal(mo_shortname("Staphylococcus aureus"), "S. aureus")
  expect_equal(mo_shortname("Staphylococcus aureus", Becker = TRUE), "S. aureus")
  expect_equal(mo_shortname("Staphylococcus aureus", Becker = "all", language = "en"), "CoPS")
  expect_equal(mo_shortname("Streptococcus agalactiae"), "S. agalactiae")
  expect_equal(mo_shortname("Streptococcus agalactiae", Lancefield = TRUE), "GBS")

  expect_true(mo_url("Escherichia coli") %like% "www.catalogueoflife.org")

  # test integrity
  MOs <- microorganisms
  expect_identical(MOs$fullname, mo_fullname(MOs$fullname, language = "en"))

  # check languages
  expect_equal(mo_type("Escherichia coli", language = "de"), "Bakterien")
  expect_equal(mo_gramstain("Escherichia coli", language = "nl"), "Gram-negatief")

  expect_output(print(mo_gramstain("Escherichia coli", language = "en")))
  expect_output(print(mo_gramstain("Escherichia coli", language = "de")))
  expect_output(print(mo_gramstain("Escherichia coli", language = "nl")))
  expect_output(print(mo_gramstain("Escherichia coli", language = "es")))
  expect_output(print(mo_gramstain("Escherichia coli", language = "pt")))
  expect_output(print(mo_gramstain("Escherichia coli", language = "it")))
  expect_output(print(mo_gramstain("Escherichia coli", language = "fr")))

  expect_error(mo_gramstain("Escherichia coli", language = "UNKNOWN"))

  # manual property function
  expect_error(mo_property("Escherichia coli", property = c("tsn", "fullname")))
  expect_error(mo_property("Escherichia coli", property = "UNKNOWN"))
  expect_identical(mo_property("Escherichia coli", property = "fullname"),
                   mo_fullname("Escherichia coli"))
  expect_identical(mo_property("Escherichia coli", property = "genus"),
                   mo_genus("Escherichia coli"))
  expect_identical(mo_property("Escherichia coli", property = "species"),
                   mo_species("Escherichia coli"))

  expect_identical(suppressWarnings(mo_ref("Chlamydia psittaci")), "Page, 1968")
  expect_identical(mo_ref("Chlamydophila psittaci"), "Everett et al., 1999")

  expect_equal(mo_snomed("Escherichia coli"), 112283007)
  
  # old codes must throw a warning in mo_* family
  expect_message(mo_name(c("B_ESCHR_COL", "B_STPHY_AUR")))
  
  # outcome of mo_fullname must always return the fullname from the data set
  x <- data.frame(mo = microorganisms$mo,
                  # fullname from the original data:
                  f1 = microorganisms$fullname,
                  # newly created fullname based on MO code:
                  f2 = mo_fullname(microorganisms$mo, language = "en"),
                  stringsAsFactors = FALSE)
  expect_equal(nrow(subset(x, f1 != f2)), 0)
  
  # is gram pos/neg (also return FALSE for all non-bacteria)
  expect_equal(mo_is_gram_negative(c("Escherichia coli", "Staphylococcus aureus", "Candida albicans")),
               c(TRUE, FALSE, FALSE))
  expect_equal(mo_is_gram_positive(c("Escherichia coli", "Staphylococcus aureus", "Candida albicans")),
               c(FALSE, TRUE, FALSE))
  # is intrinsic resistant
  expect_equal(mo_is_intrinsic_resistant(c("Escherichia coli", "Staphylococcus aureus", "Candida albicans"),
                                         "vanco"),
               c(TRUE, FALSE, FALSE))
  
  # with reference data
  expect_equal(mo_name("test", reference_df = data.frame(col1 = "test", mo = "B_ESCHR_COLI")), 
               "Escherichia coli")
  
  library(dplyr)
  expect_equal(example_isolates %>% filter(mo_is_gram_negative()) %>% nrow(),
               730)
  expect_equal(example_isolates %>% filter(mo_is_gram_positive()) %>% nrow(),
               1238)
  expect_equal(example_isolates %>% filter(mo_is_intrinsic_resistant(ab = "Vancomycin")) %>% nrow(),
               710)
})
```
