## Supplementary material for "AMR - An R Package for Working with Antimicrobial Resistance Data": Source package: test-pca.R

```
# ==================================================================== #
# TITLE                                                                #
# Antimicrobial Resistance (AMR) Analysis for R                        #
#                                                                      #
# SOURCE                                                               #
# https://github.com/msberends/AMR                                     #
#                                                                      #
# LICENCE                                                              #
# (c) 2018-2021 Berends MS, Luz CF et al.                              #
# Developed at the University of Groningen, the Netherlands, in        #
# collaboration with non-profit organisations Certe Medical            #
# Diagnostics & Advice, and University Medical Center Groningen.       # 
#                                                                      #
# This R package is free software; you can freely use and distribute   #
# it for both personal and commercial purposes under the terms of the  #
# GNU General Public License version 2.0 (GNU GPL-2), as published by  #
# the Free Software Foundation.                                        #
# We created this package for both routine data analysis and academic  #
# research and it was publicly released in the hope that it will be    #
# useful, but it comes WITHOUT ANY WARRANTY OR LIABILITY.              #
#                                                                      #
# Visit our website for the full manual and a complete tutorial about  #
# how to conduct AMR analysis: https://msberends.github.io/AMR/        #
# ==================================================================== #

context("pca.R")

test_that("PCA works", {
  
  skip_on_cran()
  
  resistance_data <- structure(list(order = c("Bacillales", "Enterobacterales", "Enterobacterales"),
                                    genus = c("Staphylococcus", "Escherichia", "Klebsiella"), 
                                    AMC = c(0.00425, 0.13062, 0.10344),
                                    CXM = c(0.00425, 0.05376, 0.10344),
                                    CTX = c(0.00000, 0.02396, 0.05172), 
                                    TOB = c(0.02325, 0.02597, 0.10344),
                                    TMP = c(0.08387, 0.39141, 0.18367)),
                               class = c("grouped_df", "tbl_df", "tbl", "data.frame"),
                               row.names = c(NA, -3L), 
                               groups = structure(list(order = c("Bacillales", "Enterobacterales"),
                                                       .rows = list(1L, 2:3)),
                                                  row.names = c(NA, -2L),
                                                  class = c("tbl_df", "tbl", "data.frame"), 
                                                  .drop = TRUE))
  
  pca_model <- pca(resistance_data)
  
  expect_s3_class(pca_model, "pca")
  
  pdf(NULL) # prevent Rplots.pdf being created
  ggplot_pca(pca_model, ellipse = TRUE)
  ggplot_pca(pca_model, arrows_textangled = FALSE)
  
  if (require("dplyr")) {
    resistance_data <- example_isolates %>% 
      group_by(order = mo_order(mo),
               genus = mo_genus(mo)) %>%
      summarise_if(is.rsi, resistance, minimum = 0)
    pca_result <- resistance_data %>%         
      pca(AMC, CXM, CTX, CAZ, GEN, TOB, TMP, "SXT") 
    expect_s3_class(pca_result, "prcomp")
    ggplot_pca(pca_result, ellipse = TRUE)
    ggplot_pca(pca_result, ellipse = FALSE, arrows_textangled = FALSE, scale = FALSE)
  }
})
```
