## Supplementary material for "AMR - An R Package for Working with Antimicrobial Resistance Data": Source package: test-proportion.R

```
# ==================================================================== #
# TITLE                                                                #
# Antimicrobial Resistance (AMR) Analysis for R                        #
#                                                                      #
# SOURCE                                                               #
# https://github.com/msberends/AMR                                     #
#                                                                      #
# LICENCE                                                              #
# (c) 2018-2021 Berends MS, Luz CF et al.                              #
# Developed at the University of Groningen, the Netherlands, in        #
# collaboration with non-profit organisations Certe Medical            #
# Diagnostics & Advice, and University Medical Center Groningen.       # 
#                                                                      #
# This R package is free software; you can freely use and distribute   #
# it for both personal and commercial purposes under the terms of the  #
# GNU General Public License version 2.0 (GNU GPL-2), as published by  #
# the Free Software Foundation.                                        #
# We created this package for both routine data analysis and academic  #
# research and it was publicly released in the hope that it will be    #
# useful, but it comes WITHOUT ANY WARRANTY OR LIABILITY.              #
#                                                                      #
# Visit our website for the full manual and a complete tutorial about  #
# how to conduct AMR analysis: https://msberends.github.io/AMR/        #
# ==================================================================== #

context("proportion.R")

test_that("proportions works", {
  
  skip_on_cran()
  
  expect_equal(proportion_R(example_isolates$AMX), resistance(example_isolates$AMX))
  expect_equal(proportion_SI(example_isolates$AMX), susceptibility(example_isolates$AMX))
  
  # AMX resistance in `example_isolates`
  expect_equal(proportion_R(example_isolates$AMX), 0.5955556, tolerance = 0.0001)
  expect_equal(proportion_I(example_isolates$AMX), 0.002222222, tolerance = 0.0001)
  expect_equal(1 - proportion_R(example_isolates$AMX) - proportion_I(example_isolates$AMX),
               proportion_S(example_isolates$AMX))
  expect_equal(proportion_R(example_isolates$AMX) + proportion_I(example_isolates$AMX),
               proportion_IR(example_isolates$AMX))
  expect_equal(proportion_S(example_isolates$AMX) + proportion_I(example_isolates$AMX),
               proportion_SI(example_isolates$AMX))

  expect_equal(example_isolates %>% proportion_SI(AMC),
               0.7626397,
               tolerance = 0.0001)
  expect_equal(example_isolates %>% proportion_SI(AMC, GEN),
               0.9408,
               tolerance = 0.0001)
  expect_equal(example_isolates %>% proportion_SI(AMC, GEN, only_all_tested = TRUE),
               0.9382647,
               tolerance = 0.0001)

  library(dplyr)
  # percentages
  expect_equal(example_isolates %>%
                 group_by(hospital_id) %>%
                 summarise(R = proportion_R(CIP, as_percent = TRUE),
                           I = proportion_I(CIP, as_percent = TRUE),
                           S = proportion_S(CIP, as_percent = TRUE),
                           n = n_rsi(CIP),
                           total = n()) %>%
                 pull(n) %>%
                 sum(),
               1409)

  # count of cases
  expect_equal(example_isolates %>%
                 group_by(hospital_id) %>%
                 summarise(cipro_p = proportion_SI(CIP, as_percent = TRUE),
                           cipro_n = n_rsi(CIP),
                           genta_p = proportion_SI(GEN, as_percent = TRUE),
                           genta_n = n_rsi(GEN),
                           combination_p = proportion_SI(CIP, GEN, as_percent = TRUE),
                           combination_n = n_rsi(CIP, GEN)) %>%
                 pull(combination_n),
               c(305, 617, 241, 711))
  
  reset_all_thrown_messages()
  expect_warning(proportion_R(as.character(example_isolates$AMC)))
  reset_all_thrown_messages()
  expect_warning(proportion_S(as.character(example_isolates$AMC)))
  reset_all_thrown_messages()
  expect_warning(proportion_S(as.character(example_isolates$AMC,
                                           example_isolates$GEN)))
  reset_all_thrown_messages()
  expect_warning(n_rsi(as.character(example_isolates$AMC,
                                    example_isolates$GEN)))
  expect_equal(suppressWarnings(n_rsi(as.character(example_isolates$AMC,
                                                   example_isolates$GEN))),
               1879)

  # check for errors
  expect_error(proportion_IR("test", minimum = "test"))
  expect_error(proportion_IR("test", as_percent = "test"))
  expect_error(proportion_I("test", minimum = "test"))
  expect_error(proportion_I("test", as_percent = "test"))
  expect_error(proportion_S("test", minimum = "test"))
  expect_error(proportion_S("test", as_percent = "test"))
  expect_error(proportion_S("test", also_single_tested = TRUE))

  # check too low amount of isolates
  expect_identical(suppressWarnings(proportion_R(example_isolates$AMX, minimum = nrow(example_isolates) + 1)),
                   NA_real_)
  expect_identical(suppressWarnings(proportion_I(example_isolates$AMX, minimum = nrow(example_isolates) + 1)),
                   NA_real_)
  expect_identical(suppressWarnings(proportion_S(example_isolates$AMX, minimum = nrow(example_isolates) + 1)),
                   NA_real_)

  # warning for speed loss
  reset_all_thrown_messages()
  expect_warning(proportion_R(as.character(example_isolates$GEN)))
  reset_all_thrown_messages()
  expect_warning(proportion_I(as.character(example_isolates$GEN)))
  reset_all_thrown_messages()
  expect_warning(proportion_S(example_isolates$AMC, as.character(example_isolates$GEN)))

  # proportion_df
  expect_equal(
    example_isolates %>% select(AMX) %>% proportion_df() %>% pull(value),
    c(example_isolates$AMX %>% proportion_SI(),
      example_isolates$AMX %>% proportion_R())
  )
  expect_equal(
    example_isolates %>% select(AMX) %>% proportion_df(combine_IR = TRUE) %>% pull(value),
    c(example_isolates$AMX %>% proportion_S(),
      example_isolates$AMX %>% proportion_IR())
  )
  expect_equal(
    example_isolates %>% select(AMX) %>% proportion_df(combine_SI = FALSE) %>% pull(value),
    c(example_isolates$AMX %>% proportion_S(),
      example_isolates$AMX %>% proportion_I(),
      example_isolates$AMX %>% proportion_R())
  )
  
  expect_error(proportion_df(c("A", "B", "C")))
  expect_error(proportion_df(example_isolates[, "date"]))
})
```
