## Supplementary material for "AMR - An R Package for Working with Antimicrobial Resistance Data": Source package: test-random.R

```
# ==================================================================== #
# TITLE                                                                #
# Antimicrobial Resistance (AMR) Analysis for R                        #
#                                                                      #
# SOURCE                                                               #
# https://github.com/msberends/AMR                                     #
#                                                                      #
# LICENCE                                                              #
# (c) 2018-2021 Berends MS, Luz CF et al.                              #
# Developed at the University of Groningen, the Netherlands, in        #
# collaboration with non-profit organisations Certe Medical            #
# Diagnostics & Advice, and University Medical Center Groningen.       # 
#                                                                      #
# This R package is free software; you can freely use and distribute   #
# it for both personal and commercial purposes under the terms of the  #
# GNU General Public License version 2.0 (GNU GPL-2), as published by  #
# the Free Software Foundation.                                        #
# We created this package for both routine data analysis and academic  #
# research and it was publicly released in the hope that it will be    #
# useful, but it comes WITHOUT ANY WARRANTY OR LIABILITY.              #
#                                                                      #
# Visit our website for the full manual and a complete tutorial about  #
# how to conduct AMR analysis: https://msberends.github.io/AMR/        #
# ==================================================================== #

context("random.R")

test_that("random works", {
  skip_on_cran()
  
  expect_s3_class(random_mic(100), "mic")
  expect_s3_class(random_mic(100, mo = "Klebsiella pneumoniae"), "mic")
  expect_s3_class(random_mic(100, mo = "Klebsiella pneumoniae", ab = "meropenem"), "mic")
  expect_s3_class(random_mic(100, ab = "meropenem"), "mic")
  # no normal factors of 2
  expect_s3_class(random_mic(100, "Haemophilus influenzae", "ceftaroline"), "mic")
  
  expect_s3_class(random_disk(100), "disk")
  expect_s3_class(random_disk(100, mo = "Klebsiella pneumoniae"), "disk")
  expect_s3_class(random_disk(100, mo = "Klebsiella pneumoniae", ab = "meropenem"), "disk")
  expect_s3_class(random_disk(100, ab = "meropenem"), "disk")
  
  expect_s3_class(random_rsi(100), "rsi")
})
```
