## Supplementary material for "AMR - An R Package for Working with Antimicrobial Resistance Data": Source package: test-resistance_predict.R

```
# ==================================================================== #
# TITLE                                                                #
# Antimicrobial Resistance (AMR) Analysis for R                        #
#                                                                      #
# SOURCE                                                               #
# https://github.com/msberends/AMR                                     #
#                                                                      #
# LICENCE                                                              #
# (c) 2018-2021 Berends MS, Luz CF et al.                              #
# Developed at the University of Groningen, the Netherlands, in        #
# collaboration with non-profit organisations Certe Medical            #
# Diagnostics & Advice, and University Medical Center Groningen.       # 
#                                                                      #
# This R package is free software; you can freely use and distribute   #
# it for both personal and commercial purposes under the terms of the  #
# GNU General Public License version 2.0 (GNU GPL-2), as published by  #
# the Free Software Foundation.                                        #
# We created this package for both routine data analysis and academic  #
# research and it was publicly released in the hope that it will be    #
# useful, but it comes WITHOUT ANY WARRANTY OR LIABILITY.              #
#                                                                      #
# Visit our website for the full manual and a complete tutorial about  #
# how to conduct AMR analysis: https://msberends.github.io/AMR/        #
# ==================================================================== #

context("resistance_predict.R")

test_that("prediction of rsi works", {
  skip_on_cran()
  AMX_R <- example_isolates %>%
    filter(mo == "B_ESCHR_COLI") %>%
    rsi_predict(col_ab = "AMX",
                col_date = "date",
                model = "binomial",
                minimum = 10,
                info = TRUE) %>%
    pull("value")
  # AMX resistance will increase according to data set `example_isolates`
  expect_true(AMX_R[3] < AMX_R[20])

  x <- resistance_predict(example_isolates, col_ab = "AMX", year_min = 2010, model = "binomial")
  pdf(NULL) # prevent Rplots.pdf being created
  expect_silent(plot(x))
  expect_silent(ggplot_rsi_predict(x))
  expect_error(ggplot_rsi_predict(example_isolates))

  library(dplyr, warn.conflicts = FALSE)

  expect_output(rsi_predict(x = filter(example_isolates, mo == "B_ESCHR_COLI"),
                            model = "binomial",
                            col_ab = "AMX",
                            col_date = "date",
                            info = TRUE))
  expect_output(rsi_predict(x = filter(example_isolates, mo == "B_ESCHR_COLI"),
                            model = "loglin",
                            col_ab = "AMX",
                            col_date = "date",
                            info = TRUE))
  expect_output(rsi_predict(x = filter(example_isolates, mo == "B_ESCHR_COLI"),
                            model = "lin",
                            col_ab = "AMX",
                            col_date = "date",
                            info = TRUE))

  expect_error(rsi_predict(x = filter(example_isolates, mo == "B_ESCHR_COLI"),
                           model = "INVALID MODEL",
                           col_ab = "AMX",
                           col_date = "date",
                           info = TRUE))
  expect_error(rsi_predict(x = filter(example_isolates, mo == "B_ESCHR_COLI"),
                           model = "binomial",
                           col_ab = "NOT EXISTING COLUMN",
                           col_date = "date",
                           info = TRUE))
  expect_error(rsi_predict(x = filter(example_isolates, mo == "B_ESCHR_COLI"),
                           model = "binomial",
                           col_ab = "AMX",
                           col_date = "NOT EXISTING COLUMN",
                           info = TRUE))
  expect_error(rsi_predict(x = filter(example_isolates, mo == "B_ESCHR_COLI"),
                           col_ab = "AMX",
                           col_date = "NOT EXISTING COLUMN",
                           info = TRUE))
  expect_error(rsi_predict(x = filter(example_isolates, mo == "B_ESCHR_COLI"),
                           col_ab = "AMX",
                           col_date = "date",
                           info = TRUE))
  # almost all E. coli are MEM S in the Netherlands :)
  expect_error(resistance_predict(x = filter(example_isolates, mo == "B_ESCHR_COLI"),
                                  model = "binomial",
                                  col_ab = "MEM",
                                  col_date = "date",
                                  info = TRUE))
})
```
