## Supplementary material for "AMR - An R Package for Working with Antimicrobial Resistance Data": Source package: test-rsi.R

```
# ==================================================================== #
# TITLE                                                                #
# Antimicrobial Resistance (AMR) Analysis for R                        #
#                                                                      #
# SOURCE                                                               #
# https://github.com/msberends/AMR                                     #
#                                                                      #
# LICENCE                                                              #
# (c) 2018-2021 Berends MS, Luz CF et al.                              #
# Developed at the University of Groningen, the Netherlands, in        #
# collaboration with non-profit organisations Certe Medical            #
# Diagnostics & Advice, and University Medical Center Groningen.       # 
#                                                                      #
# This R package is free software; you can freely use and distribute   #
# it for both personal and commercial purposes under the terms of the  #
# GNU General Public License version 2.0 (GNU GPL-2), as published by  #
# the Free Software Foundation.                                        #
# We created this package for both routine data analysis and academic  #
# research and it was publicly released in the hope that it will be    #
# useful, but it comes WITHOUT ANY WARRANTY OR LIABILITY.              #
#                                                                      #
# Visit our website for the full manual and a complete tutorial about  #
# how to conduct AMR analysis: https://msberends.github.io/AMR/        #
# ==================================================================== #

context("rsi.R")

test_that("rsi works", {
  
  skip_on_cran()
  
  expect_true(as.rsi("S") < as.rsi("I"))
  expect_true(as.rsi("I") < as.rsi("R"))
  expect_true(is.rsi(as.rsi("S")))
  
  x <- example_isolates$AMX
  expect_s3_class(x[1], "rsi")
  expect_s3_class(x[[1]], "rsi")
  expect_s3_class(c(x[1], x[9]), "rsi")
  expect_s3_class(unique(x[1], x[9]), "rsi")
  
  pdf(NULL) # prevent Rplots.pdf being created
  expect_silent(barplot(as.rsi(c("S", "I", "R"))))
  expect_silent(plot(as.rsi(c("S", "I", "R"))))
  expect_output(print(as.rsi(c("S", "I", "R"))))
  
  expect_equal(as.character(as.rsi(c(1:3))), c("S", "I", "R"))
  
  expect_equal(suppressWarnings(as.logical(as.rsi("INVALID VALUE"))), NA)
  
  expect_equal(summary(as.rsi(c("S", "R"))),
               structure(c("Class" = "rsi",
                           "%R" = "50.0% (n=1)",
                           "%SI" = "50.0% (n=1)",
                           "- %S" = "50.0% (n=1)",
                           "- %I" = " 0.0% (n=0)"), class = c("summaryDefault", "table")))
  
  expect_identical(as.logical(lapply(example_isolates, is.rsi.eligible)),
                   rep(FALSE, length(example_isolates)))
  
  library(dplyr, warn.conflicts = FALSE)
  # 40 rsi columns
  expect_equal(example_isolates %>%
                 mutate_at(vars(PEN:RIF), as.character) %>%
                 lapply(is.rsi.eligible) %>%
                 as.logical() %>%
                 sum(),
               40)
  
  expect_output(print(tibble(ab = as.rsi("S"))))
  
  expect_error(as.rsi.mic(as.mic(16)))
  expect_error(as.rsi.disk(as.disk(16)))
  
  expect_error(get_guideline("this one does not exist"))
  
  expect_s3_class(example_isolates %>%
                    mutate(m = as.mic(2),
                           d = as.disk(20)) %>% 
                    skimr::skim(),
                  "data.frame")
  expect_s3_class(skimr::skim(example_isolates),
                  "data.frame")

})

test_that("mic2rsi works", {
  
  skip_on_cran()
  
  # S. pneumoniae/ampicillin in EUCAST 2020: 0.5-2 ug/ml (R is only > 2)
  expect_equal(as.character(
    as.rsi(x = as.mic(c(0.125, 0.5, 1, 2, 4)),
           mo = "B_STRPT_PNMN",
           ab = "AMP",
           guideline = "EUCAST 2020")),
    c("S", "S", "I", "I", "R"))
  # S. pneumoniae/amoxicillin in CLSI 2019: 2-8 ug/ml (R is 8 and > 8)
  expect_equal(as.character(
    as.rsi(x = as.mic(c(1, 2, 4, 8, 16)),
           mo = "B_STRPT_PNMN",
           ab = "AMX",
           guideline = "CLSI 2019")),
    c("S", "S", "I", "R", "R"))

  # cutoffs at MIC = 8
  expect_equal(as.rsi(as.mic(2), "E. coli", "ampicillin", guideline = "EUCAST 2020"),
               as.rsi("S"))
  expect_equal(as.rsi(as.mic(32), "E. coli", "ampicillin", guideline = "EUCAST 2020"),
               as.rsi("R"))
  
  expect_true(suppressWarnings(example_isolates %>%
                                 mutate(amox_mic = as.mic(2)) %>%
                                 select(mo, amox_mic) %>%
                                 as.rsi() %>%
                                 pull(amox_mic) %>%
                                 is.rsi()))
})

test_that("disk2rsi works", {
  
  skip_on_cran()
  
  expect_equal(as.character(
    as.rsi(x = as.disk(22),
           mo = "B_STRPT_PNMN",
           ab = "ERY",
           guideline = "CLSI")),
    "S")
  expect_equal(as.character(
    as.rsi(x = as.disk(18),
           mo = "B_STRPT_PNMN",
           ab = "ERY",
           guideline = "CLSI")),
    "I")
  expect_equal(as.character(
    as.rsi(x = as.disk(10),
           mo = "B_STRPT_PNMN",
           ab = "ERY",
           guideline = "CLSI")),
    "R")

  expect_true(example_isolates %>%
                mutate(amox_disk = as.disk(15)) %>%
                select(mo, amox_disk) %>%
                as.rsi(guideline = "CLSI") %>%
                pull(amox_disk) %>%
                is.rsi())
  
  # frequency tables
  if (require("cleaner")) {
    expect_s3_class(cleaner::freq(example_isolates$AMX), "freq")
  }
})

test_that("data.frame2rsi works", {
  
  skip_on_cran()
  
  df <- data.frame(microorganism = "Escherichia coli",
                   AMP = as.mic(8),
                   CIP = as.mic(0.256),
                   GEN = as.disk(18),
                   TOB = as.disk(16),
                   ERY = "R", # note about assigning <rsi> class
                   CLR = "V") # note about cleaning
  expect_s3_class(suppressWarnings(as.rsi(df)), "data.frame")
  
  expect_s3_class(suppressWarnings(as.rsi(data.frame(mo = "Escherichia coli",
                                                     amoxi = c("R", "S", "I", "invalid")))$amoxi), "rsi")
  expect_warning(data.frame(mo = "E. coli",
                            NIT = c("<= 2", 32)) %>%
                   as.rsi())
  expect_message(data.frame(mo = "E. coli",
                            NIT = c("<= 2", 32),
                            uti = TRUE) %>%
                   as.rsi())
  expect_message(
    data.frame(mo = "E. coli",
               NIT = c("<= 2", 32),
               specimen = c("urine", "blood")) %>%
      as.rsi())
})
```
