## Supplementary material for "AMR - An R Package for Working with Antimicrobial Resistance Data": Source package: test-zzz.R

```
# ==================================================================== #
# TITLE                                                                #
# Antimicrobial Resistance (AMR) Analysis for R                        #
#                                                                      #
# SOURCE                                                               #
# https://github.com/msberends/AMR                                     #
#                                                                      #
# LICENCE                                                              #
# (c) 2018-2021 Berends MS, Luz CF et al.                              #
# Developed at the University of Groningen, the Netherlands, in        #
# collaboration with non-profit organisations Certe Medical            #
# Diagnostics & Advice, and University Medical Center Groningen.       # 
#                                                                      #
# This R package is free software; you can freely use and distribute   #
# it for both personal and commercial purposes under the terms of the  #
# GNU General Public License version 2.0 (GNU GPL-2), as published by  #
# the Free Software Foundation.                                        #
# We created this package for both routine data analysis and academic  #
# research and it was publicly released in the hope that it will be    #
# useful, but it comes WITHOUT ANY WARRANTY OR LIABILITY.              #
#                                                                      #
# Visit our website for the full manual and a complete tutorial about  #
# how to conduct AMR analysis: https://msberends.github.io/AMR/        #
# ==================================================================== #

context("zzz.R")

test_that("imports work", {
  skip_on_cran()
  
  # Check if these function still exist in the package (all are in Suggests field)
  # Since GitHub Action runs every night, we will get emailed when a dependency fails based on this unit test
  
  # functions used by import_fn()
  import_functions <- c(
    "anti_join" = "dplyr",
    "cur_column" = "dplyr",
    "full_join" = "dplyr",
    "has_internet" = "curl",
    "html_attr" = "rvest",
    "html_children" = "rvest",
    "html_node" = "rvest",
    "html_nodes" = "rvest",
    "html_table" = "rvest",
    "html_text" = "rvest",
    "inner_join" = "dplyr",
    "insertText" = "rstudioapi",
    "left_join" = "dplyr",
    "new_pillar_shaft_simple" = "pillar",
    "read_html" = "xml2",
    "right_join" = "dplyr",
    "semi_join" = "dplyr",
    "showQuestion" = "rstudioapi")
  
  # functions that are called directly
  call_functions <- c(
    # cleaner
    "freq.default" = "cleaner",
    # skmir
    "inline_hist" = "skimr",
    "sfl" = "skimr",
    # set_mo_source
    "read_excel" = "readxl",
    # ggplot_rsi
    "aes_string" = "ggplot2",
    "element_blank" = "ggplot2",
    "element_line" = "ggplot2",
    "element_text" = "ggplot2",
    "facet_wrap" = "ggplot2",
    "geom_text" = "ggplot2",
    "ggplot" = "ggplot2",
    "labs" = "ggplot2",
    "layer" = "ggplot2",
    "position_fill" = "ggplot2",
    "scale_fill_manual" = "ggplot2",
    "scale_y_continuous" = "ggplot2",
    "theme" = "ggplot2",
    "theme_minimal" = "ggplot2",
    # ggplot_pca
    "aes" = "ggplot2",
    "arrow" = "ggplot2",
    "element_blank" = "ggplot2",
    "element_line" = "ggplot2",
    "element_text" = "ggplot2",
    "expand_limits" = "ggplot2",
    "geom_path" = "ggplot2",
    "geom_point" = "ggplot2",
    "geom_segment" = "ggplot2",
    "geom_text" = "ggplot2",
    "ggplot" = "ggplot2",
    "labs" = "ggplot2",
    "theme" = "ggplot2",
    "theme_minimal" = "ggplot2",
    "unit" = "ggplot2",
    "xlab" = "ggplot2",
    "ylab" = "ggplot2",
    # resistance_predict
    "aes" = "ggplot2",
    "geom_errorbar" = "ggplot2",
    "geom_point" = "ggplot2",
    "geom_ribbon" = "ggplot2",
    "ggplot" = "ggplot2",
    "labs" = "ggplot2"
  )
  
  import_functions <- c(import_functions, call_functions)
  
  for (i in seq_len(length(import_functions))) {
    fn <- names(import_functions)[i]
    pkg <- unname(import_functions[i])
    # function should exist in foreign pkg namespace
    expect(!is.null(import_fn(name = fn, pkg = pkg, error_on_fail = FALSE)),
           failure_message = paste0("Function ", pkg, "::", fn, "() does not exist anymore"))
  }
})
```
