## Supplementary material for "AMR - An R Package for Working with Antimicrobial Resistance Data": Source package: datasets.Rmd

````
---
title: "Data sets for download / own use"
output: 
  rmarkdown::html_vignette:
    toc: true
    toc_depth: 1
vignette: >
  %\VignetteIndexEntry{Data sets for download / own use}
  %\VignetteEncoding{UTF-8}
  %\VignetteEngine{knitr::rmarkdown}
editor_options: 
  chunk_output_type: console
---

```{r setup, include = FALSE, results = 'markup'}
knitr::opts_chunk$set(
  warning = FALSE,
  collapse = TRUE,
  comment = "#",
  fig.width = 7.5,
  fig.height = 5
)

library(AMR)
library(dplyr)

options(knitr.kable.NA = '')

structure_txt <- function(dataset) {
  paste0("A data set with ",
         format(nrow(dataset), big.mark = ","), " rows and ", 
         ncol(dataset), " columns, containing the following column names:  \n*",
         paste0("'", colnames(dataset), "'", collapse = ", "), "*.")
}

download_txt <- function(filename) {
  msg <- paste0("It was last updated on ", 
                trimws(format(file.mtime(paste0("../data/", filename, ".rda")), "%e %B %Y %H:%M:%S %Z")), 
                ". Find more info about the structure of this data set [here](https://msberends.github.io/AMR/reference/", ifelse(filename == "antivirals", "antibiotics", filename), ".html).\n")
  github_base <- "https://github.com/msberends/AMR/raw/master/data-raw/"
  filename <- paste0("../data-raw/", filename)
  txt <- paste0(filename, ".txt")
  rds <- paste0(filename, ".rds")
  spss <- paste0(filename, ".sav")
  stata <- paste0(filename, ".dta")
  sas <- paste0(filename, ".sas")
  excel <- paste0(filename, ".xlsx")
  create_txt <- function(filename, type, software) {
    paste0("* Download as [", software, " file](", github_base, filename, ") (", AMR:::formatted_filesize(filename), ")  \n")
  }

  if (any(file.exists(rds),
          file.exists(excel),
          file.exists(txt),
          file.exists(sas),
          file.exists(spss),
          file.exists(stata))) {
    msg <- c(msg, "\n**Direct download links:**\n\n")
  }
  if (file.exists(rds)) msg <- c(msg, create_txt(rds, "rds", "R"))
  if (file.exists(excel)) msg <- c(msg, create_txt(excel, "xlsx", "Excel"))
  if (file.exists(txt)) msg <- c(msg, create_txt(txt, "txt", "plain text"))
  if (file.exists(sas)) msg <- c(msg, create_txt(sas, "sas", "SAS"))
  if (file.exists(spss)) msg <- c(msg, create_txt(spss, "sav", "SPSS"))
  if (file.exists(stata)) msg <- c(msg, create_txt(stata, "dta", "Stata"))
  paste0(msg, collapse = "")
}

print_df <- function(x, rows = 6) {
  x %>% 
    head(n = rows) %>% 
    mutate_all(function(x) {
      if (is.list(x)) {
        sapply(x, function(y) {
          if (length(y) > 3) {
            paste0(paste(y[1:3], collapse = ", "), ", ...")
          } else if (length(y) == 0 || all(is.na(y))) {
            ""
          } else {
            paste(y, collapse = ", ")
          }
        })
      } else {
        x
      }
    }) %>%
    knitr::kable(align = "c")
}

```

All reference data (about microorganisms, antibiotics, R/SI interpretation, EUCAST rules, etc.) in this `AMR` package are reliable, up-to-date and freely available. We continually export our data sets to formats for use in R, SPSS, SAS, Stata and Excel. We also supply  tab separated files that are machine-readable and suitable for input in any software program, such as laboratory information systems. 

On this page, we explain how to download them and how the structure of the data sets look like. 

<p class="dataset-within-r">If you are reading this page from within R, please <a href="https://msberends.github.io/AMR/articles/datasets.html">visit our website</a>, which is automatically updated with every code change.</p>

## Microorganisms (currently accepted names)

`r structure_txt(microorganisms)`

This data set is in R available as `microorganisms`, after you load the `AMR` package.

`r download_txt("microorganisms")`

### Source

Our full taxonomy of microorganisms is based on the authoritative and comprehensive:

* [Catalogue of Life](http://www.catalogueoflife.org) (included version: `r AMR:::catalogue_of_life$year`)
* [List of Prokaryotic names with Standing in Nomenclature](https://lpsn.dsmz.de) (LPSN, included version: `r AMR:::catalogue_of_life$yearmonth_DSMZ`)

### Example content

Included (sub)species per taxonomic kingdom:

```{r, echo = FALSE}
microorganisms %>% 
  pull(kingdom) %>% 
  table() %>% 
  as.data.frame() %>% 
  mutate(Freq = format(Freq, big.mark = ",")) %>% 
  setNames(c("Kingdom", "Number of (sub)species")) %>% 
  print_df()
```

Example rows when filtering on genus *Escherichia*:

```{r, echo = FALSE}
microorganisms %>%
  filter(genus == "Escherichia") %>% 
  print_df()
```

## Microorganisms (previously accepted names)

`r structure_txt(microorganisms.old)`

**Note:** remember that the 'ref' columns contains the scientific reference to the old taxonomic entries, i.e. of column *'fullname'*. For the scientific reference of the new names, i.e. of column *'fullname_new'*, see the `microorganisms` data set.

This data set is in R available as `microorganisms.old`, after you load the `AMR` package.

`r download_txt("microorganisms.old")`

### Source

This data set contains old, previously accepted taxonomic names. The data sources are the same as the `microorganisms` data set:

* [Catalogue of Life](http://www.catalogueoflife.org) (included version: `r AMR:::catalogue_of_life$year`)
* [List of Prokaryotic names with Standing in Nomenclature](https://lpsn.dsmz.de) (LPSN, included version: `r AMR:::catalogue_of_life$yearmonth_DSMZ`)

### Example content

Example rows when filtering on *Escherichia*:

```{r, echo = FALSE}
microorganisms.old %>%
  filter(fullname %like% "^Escherichia") %>% 
  print_df()
```


## Antibiotic agents

`r structure_txt(antibiotics)`

This data set is in R available as `antibiotics`, after you load the `AMR` package.

`r download_txt("antibiotics")`

### Source

This data set contains all EARS-Net and ATC codes gathered from WHO and WHONET, and all compound IDs from PubChem. It also contains all brand names (synonyms) as found on PubChem and Defined Daily Doses (DDDs) for oral and parenteral administration.

* [ATC/DDD index from WHO Collaborating Centre for Drug Statistics Methodology](https://www.whocc.no/atc_ddd_index/) (note: this may not be used for commercial purposes, but is frelly available from the WHO CC website for personal use)
* [PubChem by the US National Library of Medicine](https://pubchem.ncbi.nlm.nih.gov)
* [WHONET software 2019](https://whonet.org)

### Example content

```{r, echo = FALSE}
antibiotics %>%
  filter(ab %in% colnames(example_isolates)) %>% 
  print_df()
```


## Antiviral agents

`r structure_txt(antivirals)`

This data set is in R available as `antivirals`, after you load the `AMR` package.

`r download_txt("antivirals")`

### Source

This data set contains all ATC codes gathered from WHO and all compound IDs from PubChem. It also contains all brand names (synonyms) as found on PubChem and Defined Daily Doses (DDDs) for oral and parenteral administration.

* [ATC/DDD index from WHO Collaborating Centre for Drug Statistics Methodology](https://www.whocc.no/atc_ddd_index/) (note: this may not be used for commercial purposes, but is frelly available from the WHO CC website for personal use)
* [PubChem by the US National Library of Medicine](https://pubchem.ncbi.nlm.nih.gov)

### Example content

```{r, echo = FALSE}
antivirals %>%
  print_df()
```


## Intrinsic bacterial resistance

`r structure_txt(intrinsic_resistant)`

This data set is in R available as `intrinsic_resistant`, after you load the `AMR` package.

`r download_txt("intrinsic_resistant")`

### Source

This data set contains all defined intrinsic resistance by EUCAST of all bug-drug combinations, and is based on '`r AMR:::EUCAST_VERSION_EXPERT_RULES[["3.2"]]$title`', `r AMR:::EUCAST_VERSION_EXPERT_RULES[["3.2"]]$version_txt` from `r AMR:::EUCAST_VERSION_EXPERT_RULES[["3.2"]]$year`.

### Example content

Example rows when filtering on *Enterobacter cloacae*:

```{r, echo = FALSE}
intrinsic_resistant %>%
  filter(microorganism == "Enterobacter cloacae") %>% 
  print_df(rows = Inf)
```


## Interpretation from MIC values / disk diameters to R/SI

`r structure_txt(rsi_translation)`

This data set is in R available as `rsi_translation`, after you load the `AMR` package.

`r download_txt("rsi_translation")`

### Source

This data set contains interpretation rules for MIC values and disk diffusion diameters. Included guidelines are CLSI (`r min(as.integer(gsub("[^0-9]", "", subset(rsi_translation, guideline %like% "CLSI")$guideline)))`-`r max(as.integer(gsub("[^0-9]", "", subset(rsi_translation, guideline %like% "CLSI")$guideline)))`) and EUCAST (`r min(as.integer(gsub("[^0-9]", "", subset(rsi_translation, guideline %like% "EUCAST")$guideline)))`-`r max(as.integer(gsub("[^0-9]", "", subset(rsi_translation, guideline %like% "EUCAST")$guideline)))`).

### Example content

```{r, echo = FALSE}
rsi_translation %>% 
  mutate(ab = ab_name(ab), mo = mo_name(mo)) %>% 
  print_df()
```
````
