## Supplementary figures and images for "AMR - An R Package for Working with Antimicrobial Resistance Data"

### logo.png

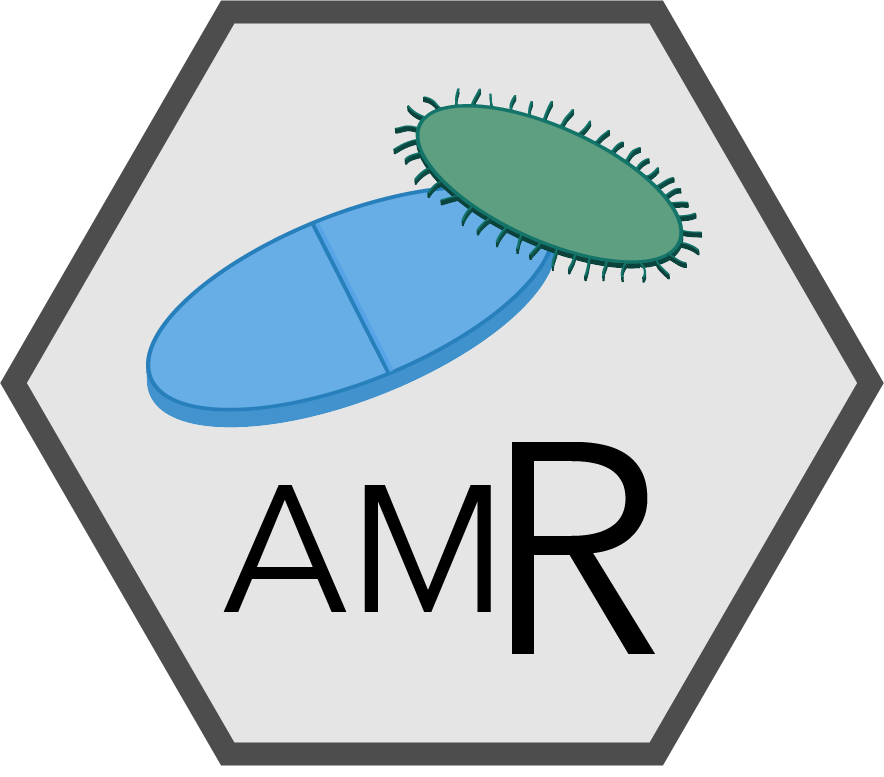

### logo_certe.png

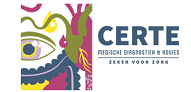

### logo_col.png

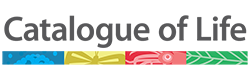

### logo_eh1h.png

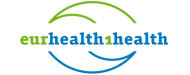

### logo_interreg.png

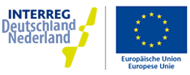

### logo_rug.png

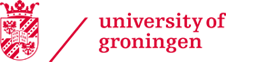

### logo_umcg.png

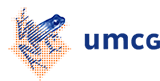

### logo_who.png

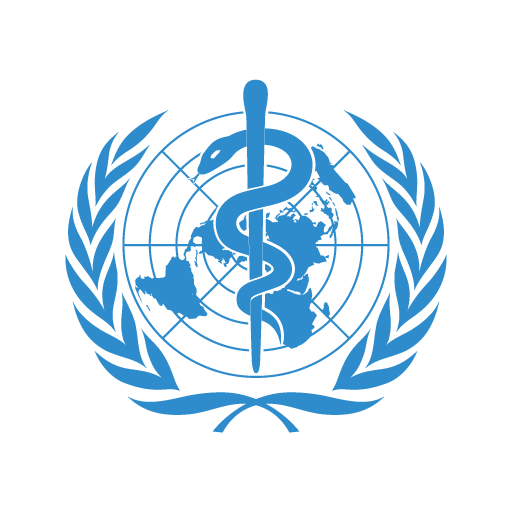

### mo_matching_score.png

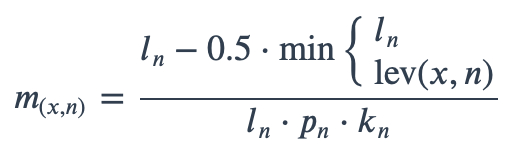
